# Rewiring phospholipid biosynthesis reveals robustness in membrane homeostasis and uncovers lipid regulatory players

**DOI:** 10.1101/2021.07.20.453065

**Authors:** Arun T. John Peter, Sabine N. S. van Schie, Ngaam J. Cheung, Agnès H. Michel, Matthias Peter, Benoît Kornmann

**Author notes:** These authors contributed equally (Arun T. John Peter, Sabine N. S. van Schie).

## Abstract

Intracellular transport of lipids by Lipid Transport Proteins (LTPs) is thought to work alongside vesicular transport to shuttle lipids from their place of synthesis to their destinations. Whereas many LTPs have been identified, it is largely unknown which routes and which LTPs a given lipid utilizes to navigate the multiple membranes of eukaryotic cells. The major and essential phospholipids, phosphatidylethanolamine (PE) and phosphatidylcholine (PC) can be produced by multiple pathways and, in the case of PE, also at multiple locations. Here, we present an approach in which we simplify and rewire yeast phospholipid synthesis by redirecting PE and PC synthesis reactions to distinct subcellular locations using chimeric enzymes fused to specific organelle targeting motifs. In rewired conditions, viability is expected to depend on homeostatic adaptation to the ensuing lipostatic perturbations and on efficient interorganelle lipid transport. We therefore performed genetic screens to identify factors involved in both of these processes. Among the candidates identified, we find genes linked to transcriptional regulation of lipid homeostasis, lipid metabolism and transport. In particular, we identify a requirement for Csf1 –an uncharacterized protein harboring a Chorein-N lipid transport domain- for survival under certain rewired conditions as well as lipidomic adaptation to cold, implicating Csf1 in interorganelle lipid transport and homeostatic adaptation.

## Introduction

Lipid exchange between membrane-bound organelles is vital to establish the characteristic membrane composition underlying organelle function and enable cell growth. Surprisingly, a large fraction of the molecular components and pathways in this fundamental cellular process are still elusive. Vesicular trafficking of lipids occurs in parallel with non-vesicular transport mediated by Lipid Transport Proteins (LTPs), catalyzing the transfer of lipids between two membranes by shielding them from the aqueous environment. LTPs often function at regions of close apposition (10-30 nM) between organelles – termed Membrane Contact Sites (MCSs) – found between various compartments, including membranes connected by vesicular trafficking pathways (Gatta & Levine, 2017; Reinisch & Prinz, 2021). Indeed, phospholipids and sterols, are transported between the ER and the plasma membrane even after inhibition of the secretory pathway, indicating that non-vesicular trafficking pathways contribute to lipid transfer at these sites (Vance *et al*, 1991; Quon *et al*, 2018; Schnabl *et al*, 2005).

Moreover, redundancy in lipid exchange between cellular compartments is observed even within non-vesicular trafficking pathways, as exemplified with mitochondria. In the yeast *Saccharomyces cerevisiae,* two pathways have been implicated in lipid transport at mitochondria; the ER-Mitochondria Encounter Structure (ERMES) pathway and the Mcp1-Vps13 pathway (Kornmann *et al*, 2009; John Peter *et al*, 2017; Lang *et al*, 2015; Park *et al*, 2016). Structural analysis revealed hydrophobic cavities, both in the SMP domains of multiple ERMES components and in the Chorein-N domain of Vps13, which facilitates lipid exchange *in vitro* (AhYoung *et al*, 2017; Jeong *et al*, 2017, 2016; Kumar *et al*, 2018; Li *et al*, 2020). The fact that mitochondria are also in contact with multiple organelles including the PM, peroxisomes, and lipid droplets (Hönscher *et al*, 2014; Schuldiner & Bohnert, 2017; Elbaz-Alon *et al*, 2014; Shai *et al*, 2018) makes it plausible that these organelles could also function as potential donor and/or acceptor organelles for lipid exchange. These redundancies in lipid exchange likely exist for other organelles as well.

Finally, an additional layer of redundancy is observed at the level of lipid synthesis; two pathways, namely the Kennedy and the cytidyl-diphosphate-diacylglycerol (CDP-DAG) pathways (Fig 1A, left panel) produce phosphatidylethanolamine (PE) and phosphatidylcholine (PC). The Kennedy pathway utilizes ethanolamine and choline – primarily obtained from the medium – to convert diacylglycerol (DAG) directly to PE and PC, respectively (Fig 1A, left panel) (Kennedy & Weiss, 1956). By contrast, the CDP-DAG pathway uses CDP-DAG generated from phosphatidic acid (PA) and a series of enzymatic reactions to produce phosphatidylserine (PS), PE and PC in sequential order (Carman & Henry, 1999). In yeast, the dually-localized PS decarboxylase Psd1 produces PE in the mitochondrial inner membrane (MIM) and ER, while Psd2 synthesizes PE at endosomes (Friedman *et al*, 2018; Gulshan *et al*, 2010). The production of PC by trimethylation of PE is performed by the ER-localized enzymes, Cho2 and Opi3. Either pathway alone is sufficient to support growth, indicating that lipids are distributed throughout the cell from different locations. Importantly, cells are likely to adapt to the absence of one of these pathways by activating homeostatic responses. Indeed, mechanisms are known for sensing lipid levels and lipid packing/saturation (Young *et al*, 2010; Covino *et al*, 2016), which are coupled to transcriptional, post-transcriptional or allosteric responses.

**Fig 1.**
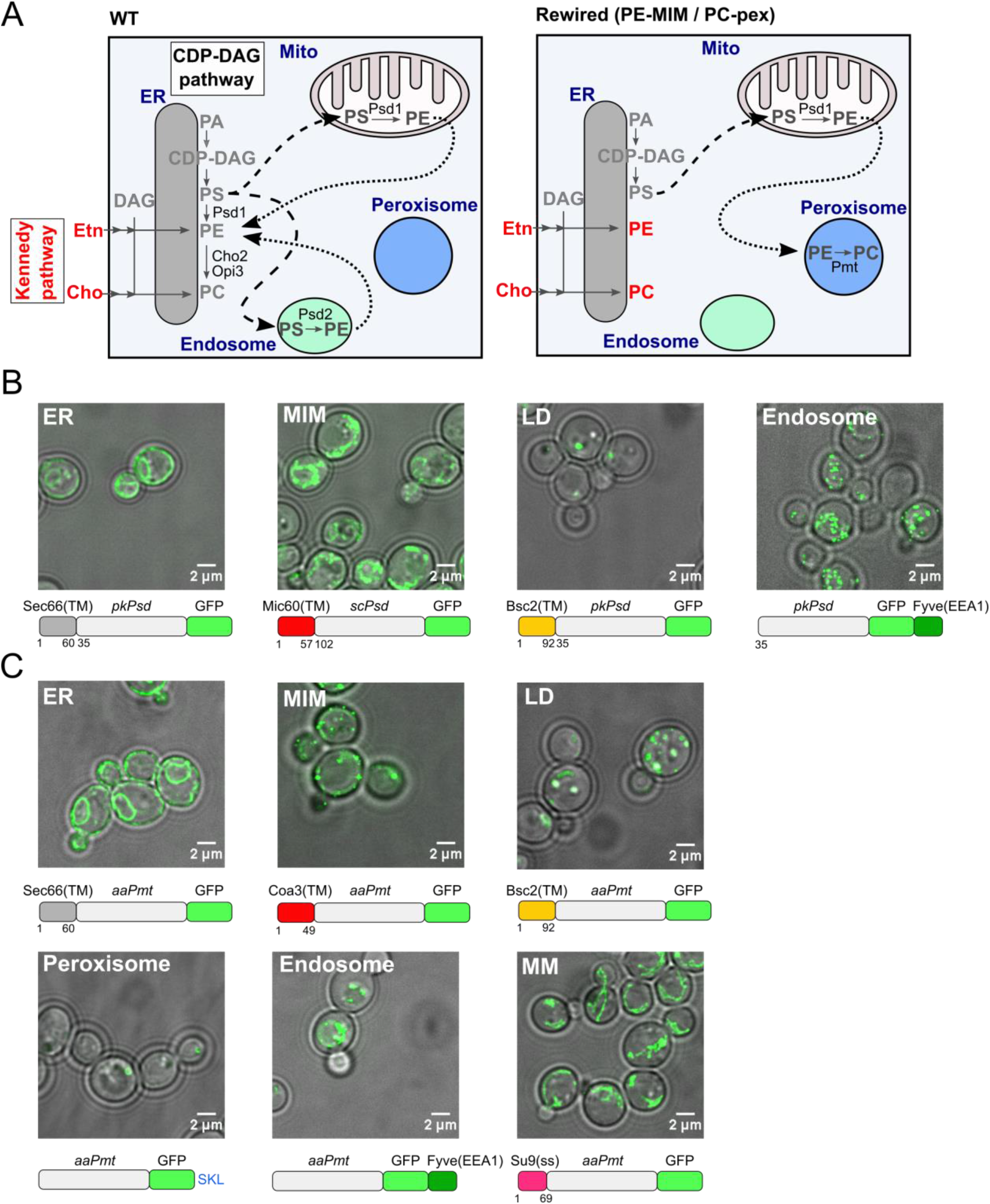
Rewiring PE and PC synthesis to expose lipid trafficking routes. **A)** Schematic depicting the topology of the CDP-DAG-and Kennedy (red) pathways producing PE and PC in wild-type (WT) yeast cells (left panel), and an example of a rewired condition in which PE synthesis is directed to the MIM and PC synthesis to peroxisomes (pex) (right panel). Black arrows indicate PS (dashed) and PE (dotted) lipid transport that must occur to enable the sequential enzymatic reactions. **B)** Localization of the GFP-tagged PE synthesizing enzyme (Psd) targeted to the indicated organelles in *choppΔ* cells, grown in SD medium supplemented with 10 mM choline. Images shown are either a single Z-slice (ER construct) or a maximum intensity projection of several Z-sections (other constructs). A schematic depicting each chimeric Psd construct is shown under the corresponding microscopy image. **C)** Localization of the GFP-tagged PC synthesizing enzyme (Pmt) targeted to the indicated organelles in *choppΔ* cells, grown in SD medium supplemented with 10 mM ethanolamine. Images shown are either a single Z-slice (ER construct) or a maximum intensity projection of several Z-sections (other constructs). A schematic depicting each chimeric Pmt construct is shown under the corresponding microscopy image.

Genetically uncovering lipid transport routes and homeostatic mechanisms requires removal of the multiple layers of redundancy. One way to achieve this goal is to delete endogenous enzymes and redirect synthesis of specific phospholipids to defined organelles through the expression of chimeric enzymes, as was successfully used to demonstrate PE transport from peroxisomes (Raychaudhuri & Prinz, 2008) or PS transport from non-native organelles (Shiino *et al*, 2021). Importantly, in this strategy, data interpretation depends on the targeting accuracy of the chimeric enzymes to the intended organelles.

Here, we extended this rewiring approach (i.e. minimalizing and rerouting phospholipid synthesis) by confining PE and PC synthesis to distinct combinations of organelles. We then performed genetic screens to identify factors that result in a fitness gain or defect in rewired conditions. We focus on *CSF1*, encoding a protein with a Chorein-N lipid transport domain (Levine, 2019; Lees & Reinisch, 2020) and uncover its role in homeostatic adaptation of the lipidome.

## Results

### Rewired yeast phospholipid synthesis as a strategy to remove layers of redundancies

We postulated that we could i) remove layers of redundancy and ii) expose trafficking routes between organelles of interest by constructing yeast strains with simplified and rewired phospholipid synthesis pathways (Fig 1A, right panel). To do this, we first generated a *<u>ch</u>o2Δ <u>op</u>i3Δ <u>p</u>sd1Δ <u>p</u>sd2Δ* strain (hereafter referred to as *choppΔ*) that relies on supplementation with ethanolamine and choline for growth. To synthesize PE and PC in distinct pairs of (or in the same) organelles, we engineered lipid synthesizing enzymes by fusing them to specific targeting sequences. Specifically, we utilized a PS decarboxylase (Psd) lacking the transmembrane domain, either from yeast or *Plasmodium knowlesi* (*pk*Psd) and a soluble PE methyltransferase from *Acetobacter aceti* (*aa*Pmt, hereafter referred to as Pmt) for the production of PE and PC, respectively (Choi *et al*, 2012; Hanada *et al*, 2001; Kobayashi *et al*, 2014). We directed the Psd and Pmt enzymes to the ER, the mitochondrial inner membrane (MIM), endosomes (endo) and lipid droplets (LD), with the Pmt enzyme targeted in addition to the peroxisomal lumen (pex) and the mitochondrial matrix (MM). We engineered a total of 24 combinations in which PE and PC were synthesized at various organelles.

We expressed the chimeric enzymes on a plasmid under the control of the TEF promoter in the *choppΔ* strain, except for yeast *PSD1*, which was expressed from its own promoter (Friedman *et al*, 2018). Correct targeting of the enzymes was verified by microscopy (Fig 1B and 1C, Fig S1-13). Indeed, the ER and mitochondria-targeted enzymes revealed fluorescence patterns characteristic of these organelles (Fig 1B and 1C, Fig S1A, S4-5, S10, S13). Likewise, Pmt targeted to the MIM stained the mitochondrial outline (Fig S1B and S9), but also accumulated as bright puncta, suggesting aggregation. This did not however preclude enzyme activity (see below). Both lipid droplet (LD) and peroxisome (pex)-targeted enzymes co-localized with a LD (Erg6-mCherry) and a peroxisome marker (mCherry-SKL), respectively (Fig S2). A minor fraction of LD- and peroxisome-targeted Pmt was detected in the ER in some cells (Fig S6 and S8). To assess localization to endocytic organelles, we co-stained cells with the amphiphilic dye FM4-64 (Fig S3). Both endosome-targeted Psd and Pmt (Psd-FYVE and Pmt-FYVE) assembled in FM4-64-positive perivacuolar puncta consistent with late endosomes. In addition, the Pmt-FYVE construct localized to vacuoles themselves (Fig 1C, Fig S7). Thus, both engineered enzymes were successfully re-targeted to endocytic compartments.

Since an altered lipid composition in the rewired strains could affect organelle integrity and interfere with protein targeting, we validated the localization of all enzyme constructs in the various combinations of PE and PC synthesis (Fig S4-S13). As both Psd and Pmt enzymes were GFP-tagged, we mutated the GFP of one of the enzymes to a “dark” state to verify proper targeting of the other enzyme. We did not observe gross differences in enzyme targeting in any rewired condition.

### Rewired yeast strains are viable and produce all phospholipids

To examine whether the rewired CDP-DAG pathway supports growth when the Kennedy pathway is inactive, we performed growth assays of all combinations either in the presence or absence of exogenous ethanolamine and choline (conditions hereafter termed Kennedy_ON_ and Kennedy_OFF_, respectively). As expected, the *choppΔ* strain could not grow in the Kennedy_OFF_ conditions (Fig 2A). Remarkably, growth was restored in all rewired strains, albeit to varying extents, indicating that the chimeric enzymes produce phospholipids, bypassing the need for the Kennedy pathway. Growth was most robust when Psd was localized to the mitochondrial inner membrane (MIM), which is normally the major site of PE synthesis (Rosenberger *et al*, 2009). Cell growth was especially limited when Pmt was targeted to the mitochondrial matrix (MM). This was not likely due to the inability of matrix-targeted Pmt to meet PC synthesis demand, since in some rewired conditions, cells grew even slower in the Kennedy_ON_ conditions (Fig 2A). Instead, slow growth was possibly due to lipostatic or proteostatic challenges.

**Fig 2.**
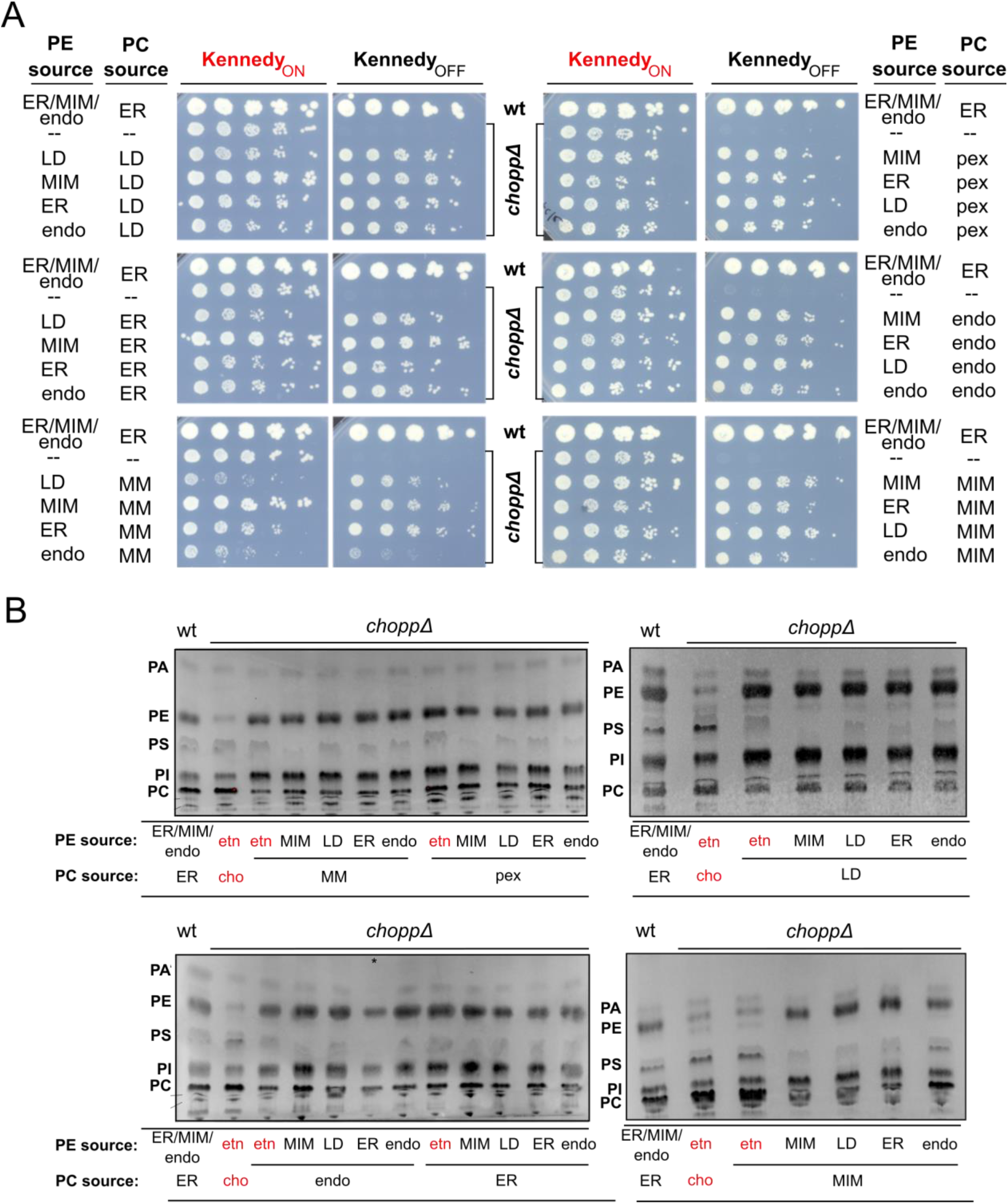
Rewired yeast cells are viable and produce all phospholipids. **A)** Five-fold serial dilutions of strains of the indicated genotypes (wt cells and *chopp*Δ cells harboring empty vectors or *chopp*Δ cells expressing chimeric Psd and Pmt enzymes). Cells were grown on SD-HIS-LEU (-HL) medium (Kennedy_OFF_) or SD-HL containing 10 mM ethanolamine and 10 mM choline (Kennedy_ON_). **B)**Thin layer chromatography (TLC) analysis of steady-state phospholipid profiles of cells of the indicated genotypes. When indicated, cells were grown in the presence of 10 mM ethanolamine (etn) and/or 10 mM choline (cho). Bands corresponding to the phospholipids PA, PS, PE, PI and PC are indicated.

To confirm that the rewired strains produced PE and PC, we used lipid thin layer chromatography (TLC). In agreement with their ability to grow, all rewired strains efficiently produced both lipids (Fig 2B). Differences in overall lipid profiles were nevertheless apparent. For example, phosphatidylinositol (PI) levels appear to be higher in most rewired strains. This increase could be a compensatory mechanism to deal with altered PE/PC ratios. Moreover, in the strain expressing the mito-Psd construct, PS levels were consistently low, suggesting that PS is most efficiently metabolized into PE in these conditions. Altogether, the growth and lipid profile of rewired yeast strains indicate that yeast cells can tolerate topological rewiring of their PE and PC biosynthesis pathways. This suggests that cells must have transport mechanisms to re-distribute lipids from these non-native locations to other membranes around the cell, as well as mechanisms to cope with lipid imbalance.

### A transposon mutagenesis screen to identify genetic components essential for survival of rewired yeast

To identify the genetic components necessary to reroute lipid trafficking and handle lipid imbalances in the rewired strains, we performed genetic screens using SAturated Transposon Analysis in Yeast (SATAY) (Fig 3A) (Michel *et al*, 2019, 2017). Briefly, we transformed rewired yeast cells with a plasmid containing a galactose-inducible transposase (TPase) and a transposon (TN) disrupting the *TRP1* gene. Transposition results in the repair of the *TRP1* gene and allows transposed cells to grow in medium lacking tryptophan. Transposition was induced by growing cells in galactose-containing medium for ~56 hours. Since transposition occurs predominantly in saturated phase (Michel *et al*, 2019), this generates millions of independent transposition events. The transposon libraries for a given genotype were then grown for several cycles in medium lacking tryptophan, in the Kennedy_ON_ or Kennedy_OFF_ conditions to i) select for cells that have successfully transposed and ii) to select against mutants that cannot grow in the rewired conditions. Transposon libraries were prepared for Next Generation Sequencing to map and quantify the transposon insertion sites (TNs) across the genome. ORFs devoid of transposons or enriched for transposons correspond to genes that are essential or harmful in the given conditions, respectively.

**Fig 3.**
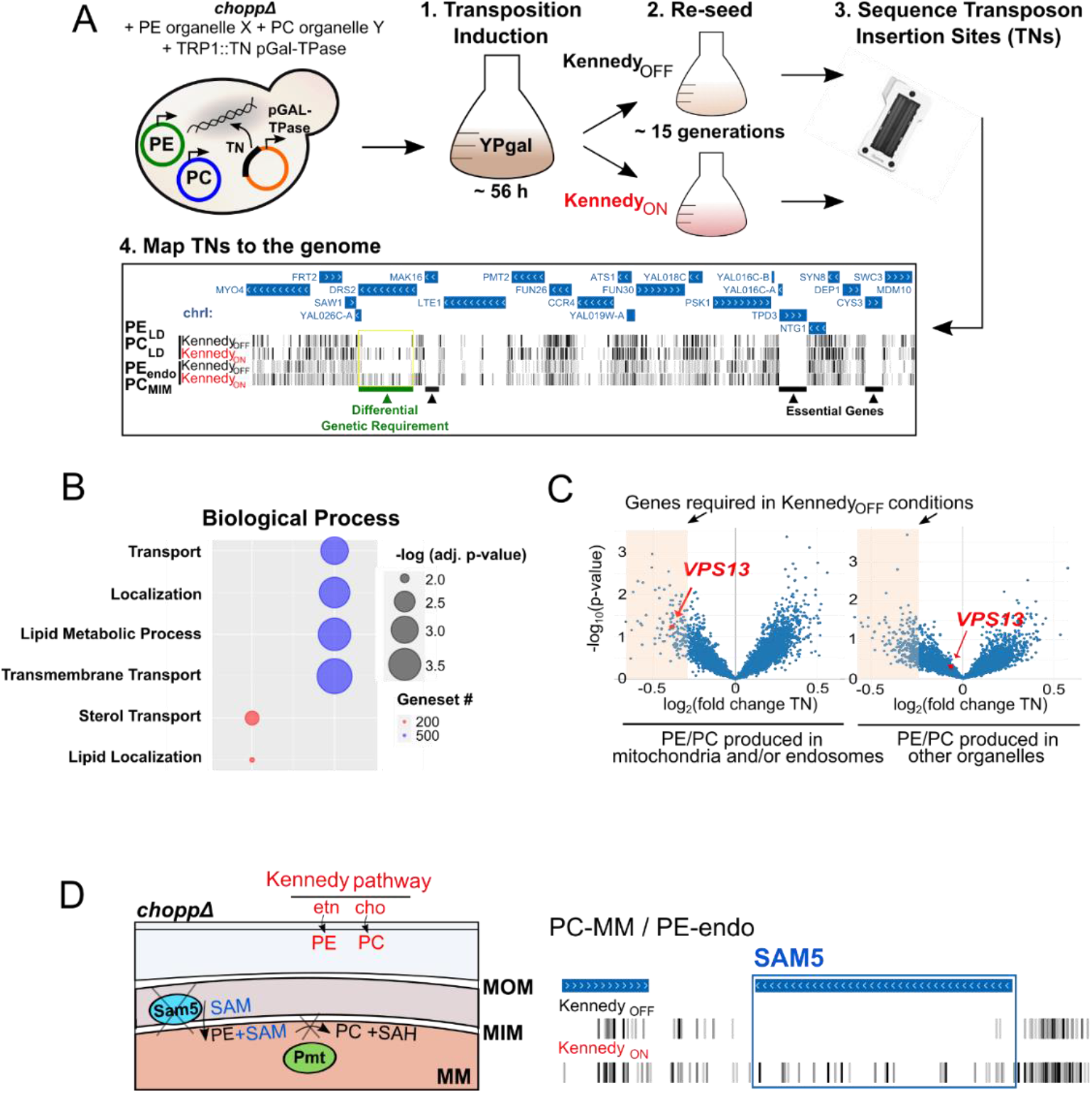
Transposon mutagenesis screens to identify genes required for adaptation and lipid trafficking in the rewired strains. **A)** Outline of the SATAY screening procedure. Cells expressing plasmids containing the PE and PC synthesizing enzymes and a plasmid harboring the galactose inducible transposase (TPase) and a transposon (TN) disrupting the *TRP1* gene were grown for ~56h in galactose-containing medium to induce transposition (step 1). Cells were inoculated in synthetic medium lacking tryptophan in either Kennedy_ON_ or _OFF_ conditions, grown for several generations (step 2) and harvested for DNA extraction and sequencing of transposon insertion sites (TNs) (step 3). TNs are mapped to the genome to identify genes that become required under certain rewired conditions (step 4). An illustrative example of TN insertions of four libraries visualized in the UCSC genome browser highlights a genetic requirement for *DRS2* (green bar) in Kennedy_OFF_ conditions. **B)** GO term enrichments in the top 200 (red) and top 500 (blue) most variable genes (defined as genes with the highest standard deviation of TN insertions across libraries) identified using the yeastmine web server. Significance was determined using Holm-Bonferroni corrected *p*-values with a cut-off of 0.05, the size of the circles is inversely proportional to the *p*-values (i.e. bigger circles indicate higher significance). **C)** *VPS13* is required in Kennedy_OFF_ conditions when chimeric enzymes are targeted to mitochondria or endosomes. Volcano plots show the fold change of number of transposon insertions per gene of libraries grown in Kennedy_OFF_ versus ON conditions. This comparison included all six libraries where the enzymes are targeted to endosomes and/or mitochondria (left panel) or six libraries with enzymes targeted to other organelles (right panel). Data-points for *VPS13* are highlighted in red. **D)** Schematic (left panel) illustrating the requirement for SAM transport across the mitochondrial inner membrane by Sam5 for PC production by Pmt in the mitochondrial matrix (MM). Lipids can be produced by the Kennedy pathway (red) independent of Sam5 activity. Transposon insertion maps generated in the UCSC genome browser for *SAM5* for the library where PC is produced in the MM in either Kennedy_ON_ or _OFF_ conditions (right panel).

We generated a total of 24 libraries comprising 12 rewired strains grown in both Kennedy_ON_ and _OFF_ conditions. Transposition insertion sites mapping across the entire yeast genome (Suppl. Data 1) can be viewed in the genome browser (http://genome-euro.ucsc.edu/s/vsabine/Rewiring_UCSC_genomebrowser, Suppl. Data 1). Additionally, we generated multiple interactive volcano plots available for browsing (Suppl. Data 2 and https://kornmann.bioch.ox.ac.uk/satay/rewiring) in which we plotted the fold change of the average number of transposons per gene of a test set and a reference set against a *p*-value associated with this difference to identify genes that provide a fitness advantage or disadvantage in certain libraries. Using this analysis, we compared different sets of libraries against each other (e.g. individual libraries against all other libraries and libraries with a common condition, such as the expression of the same chimeric enzyme, or Kennedy_ON_ vs. _OFF_, against other libraries).

We performed hierarchical clustering of the genes according to the number of transposons targeting them (Fig S14). Libraries with the same genotypes, in the Kennedy_ON_ and _OFF_ conditions, cluster as nearest neighbors. Since these libraries are replicates grown under different conditions, and the number of transposons in most genes is unaffected by the activity of the Kennedy pathway, this clustering may be dominated by technical variation. Nevertheless, independent libraries generated in strains expressing the same PE or PC producing enzymes also tended to cluster together. In some cases, the location of the PE producing enzyme dominates in the clustering (e.g. PE-LD), and in other cases it is that of the PC-producing enzyme (PC-pex). Yet in general, there is no pattern based on the location of one or the other enzyme. Overall, this clustering implies that the genotype dominates in determining fitness and genetic requirements rather than the availability of choline and ethanolamine.

To identify genes required to adapt to the rewired conditions, we first sought to filter for those with the most variable number of transposons across the whole dataset, using the standard deviation (Suppl. Data 3). This subset may omit genes that become essential in only one or very few conditions yet is an unbiased way of identifying genes that are generally affecting fitness in rewired conditions.

We selected the top 200 and top 500 most variable genes and searched for GO term enrichment using the yeastmine web server (Balakrishnan *et al*, 2012). Remarkably, these genes were enriched for GO terms related to lipid homeostasis (lipid transport, localization and metabolism), and other processes such as intracellular transport (Fig 3B). Particularly striking examples of variable genes are *DRS2*, *DFN2* and *LRO1*, which are completely devoid of transposons in some conditions (Fig S15). *DRS2* and *DFN2* both encode phospholipid translocases required for maintaining phospholipid asymmetry (Sebastian *et al*, 2012). They may be required to maintain proper lipid bilayer distribution in rewired strains. Lro1 is an acyltransferase that converts DAG to triacylglycerol (TAG) using phospholipids as acyl chain donors (Barbosa *et al*, 2019). Thus, Lro1 may be required to degrade excess membrane. Importantly, these genes had variable but non-similar patterns of transposon coverage across libraries, indicating that it was not a common condition (e.g. Kennedy_ON_ or _OFF_) that made them more or less required.

Intriguingly, the lipid transporter Vps13, even though never completely essential, appears to be consistently less covered in Kennedy_OFF_ conditions (Fig S16). Importantly, this is only the case for libraries in which Psd and Pmt are targeted to mitochondria and/or endosomes, organelles to which this lipid transporter localizes (Lang *et al*, 2015; John Peter *et al*, 2017; Park *et al*, 2016) (Fig 3C, Fig S16). Together, these results suggest that Vps13 may have a more prominent role at these organelles in Kennedy_OFF_ conditions, and underlines that lipid transporters could become limiting upon rewiring at their native sites. Notably, we find that the inner mitochondrial S-adenosylmethionine (SAM) transporter *SAM5* (Marobbio *et al*, 2003) is essential in Kennedy_OFF_ conditions when PC synthesis is targeted to the mitochondrial matrix (Fig 3D). SAM is the methyl group donor for the conversion of PE to PC, and thus SAM is necessary in the matrix for PC production, making Sam5 indispensable. This finding excludes the possibility that growth of this strain is attributable to PC production by a small fraction of mistargeted enzymes. Thus, our rewiring approach can identify genes that maintain cell viability in rewired conditions and these genes are, as expected, enriched for lipid and membrane-related processes.

### Revealing genes with common patterns across rewired conditions

Previous genome-wide analyses have shown that genes that are required under the same set of conditions usually contribute to the same biological process, pathway or protein complex (Costanzo *et al*, 2016). Therefore, we asked if we could identify genes that have similar patterns of transposon insertion across different rewired libraries. We selected the most variable genes and computed the Pearson correlation coefficient for pairs of genes across our libraries and performed hierarchical clustering on the correlation matrix (Fig 4A, Suppl. Data 2).

**Fig 4.**
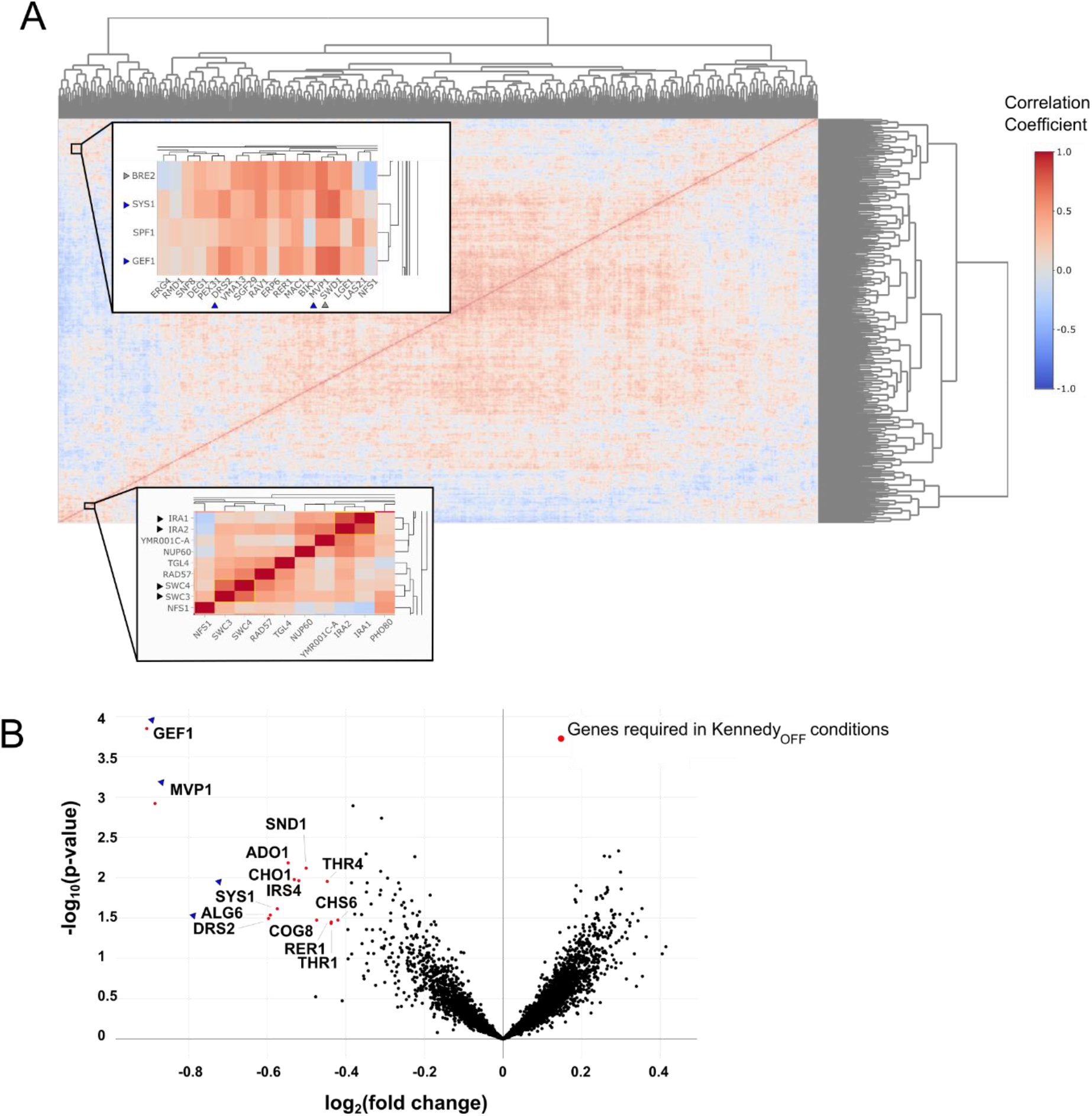
Clustering of gene correlations across libraries reveal gene groups that work together to handle rewired conditions. **A)** Hierarchical clustering of the Pearson correlation coefficients of transposon insertion profiles for gene pairs. Insets show clustering of the *IRA1*/*IRA2* and *SWC3*/*SWC4* gene pairs (bottom inset, black arrows) and high correlation of the *SYS1*/*GEF1*/*SPF1* cluster with *DRS2* and *MVP1* (top inset, blue arrows). **B)** Volcano plot comparing the number of transposon insertions per gene of all 12 libraries in Kennedy_ON_ and _OFF_ conditions. Genes significantly required in Kennedy_OFF_ conditions are highlighted in red (p-value < 0.05, log_2_(fold change) < −0.5). Genes highlighted in the clusterogram in A (top inset) are indicated with blue arrows.

We find clustering of highly correlated gene pairs consistent with their known function and/or profile similarities computed in the whole-genome genetic interaction map (Costanzo *et al*, 2016) (Fig 4A). Examples include *IRA1* and *IRA2* (Tanaka *et al*, 1990), paralogs that negatively regulate the RAS-cAMP pathway and *SWC3* and *SWC4*, both components of the chromatin remodeling SWR Complex (Mizuguchi *et al*, 2004) (Fig 4A, black arrows). We also find correlation between *BRE2* and *SWD1* (Fig 4A, grey arrows), both members of the same histone methylation complex (COMPASS) (Krogan *et al*, 2002).

A striking example, also consistent with the Costanzo dataset (Costanzo *et al*, 2016), is the cluster containing *SYS1*, *SPF1*, *GEF1* and *MVP1* (Fig 4A-B, blue arrows). We found that this cluster became essential in most Kennedy_OFF_ conditions, as shown by a volcano plot in which libraries grown with and without ethanolamine and choline are compared against each other (Fig 4B). This implies that these genes function together and act redundantly with the Kennedy pathway to maintain cell viability in the rewired strains. Given their known roles in vesicular trafficking and protein sorting, it is possible that the Kennedy pathway is required for efficient vesicular trafficking in rewired conditions, or that vesicular trafficking is needed to redistribute lipids in the cell. Alternatively, vesicular trafficking may affect proper localization of other factors required for lipid distribution (e.g. Drs2). These examples demonstrate that the transposon insertion patterns across libraries can reveal genes that function together to maintain cell viability upon lipostatic challenges.

### Uncovering adaptations in the rewired libraries

Besides unique adaptations to specific conditions, rewired strains may have common strategies to deal with general phospholipid and membrane stress. To identify such genes, we compared the rewired libraries generated in this study, with two previously generated wild-type libraries (Fig 5A) (Michel *et al*, 2019). Expected changes in transposon insertion profiles were associated with technical differences in library generation, in the present and previous studies (e.g. genes for histidine, leucine, tryptophan and adenine prototrophy, leucine transporters) (Fig5A, grey dots). Besides these, striking differences were found in several genes involved in transcriptional adaptation (Fig 5A, Suppl. Data 4). One of the best hits is *OPI1*, a transcriptional repressor for a variety of lipid biosynthesis genes under the control of the Ino2-Ino4 transcription activation complex (e.g. *INO1*, *CHO1*, *ITR1*, genes of the Kennedy pathway) (Henry *et al*, 2012). *OPI1* is required in all rewired libraries, indicating an increased need for transcriptional regulation of phospholipid biosynthesis. Another example is the transcription factor *CBF1*, which is implicated in enhancing Ino2-Ino4 transcriptional activation (Shetty & Lopes, 2010), mitochondrial respiration, and repressing ceramide biosynthesis (DeMille *et al*, 2019). Examples of genes required for growth only in specific libraries, include *NEM1*, a phosphatase controlling phospholipid biosynthesis (Siniossoglou *et al*, 1998), and *MGA2*, a transmembrane transcription factor involved in sensing membrane saturation (Covino *et al*, 2016) (Fig S17). These results suggest that distinct rewired conditions cause both common and unique membrane challenges that require a combination of adaptive responses.

**Fig 5.**
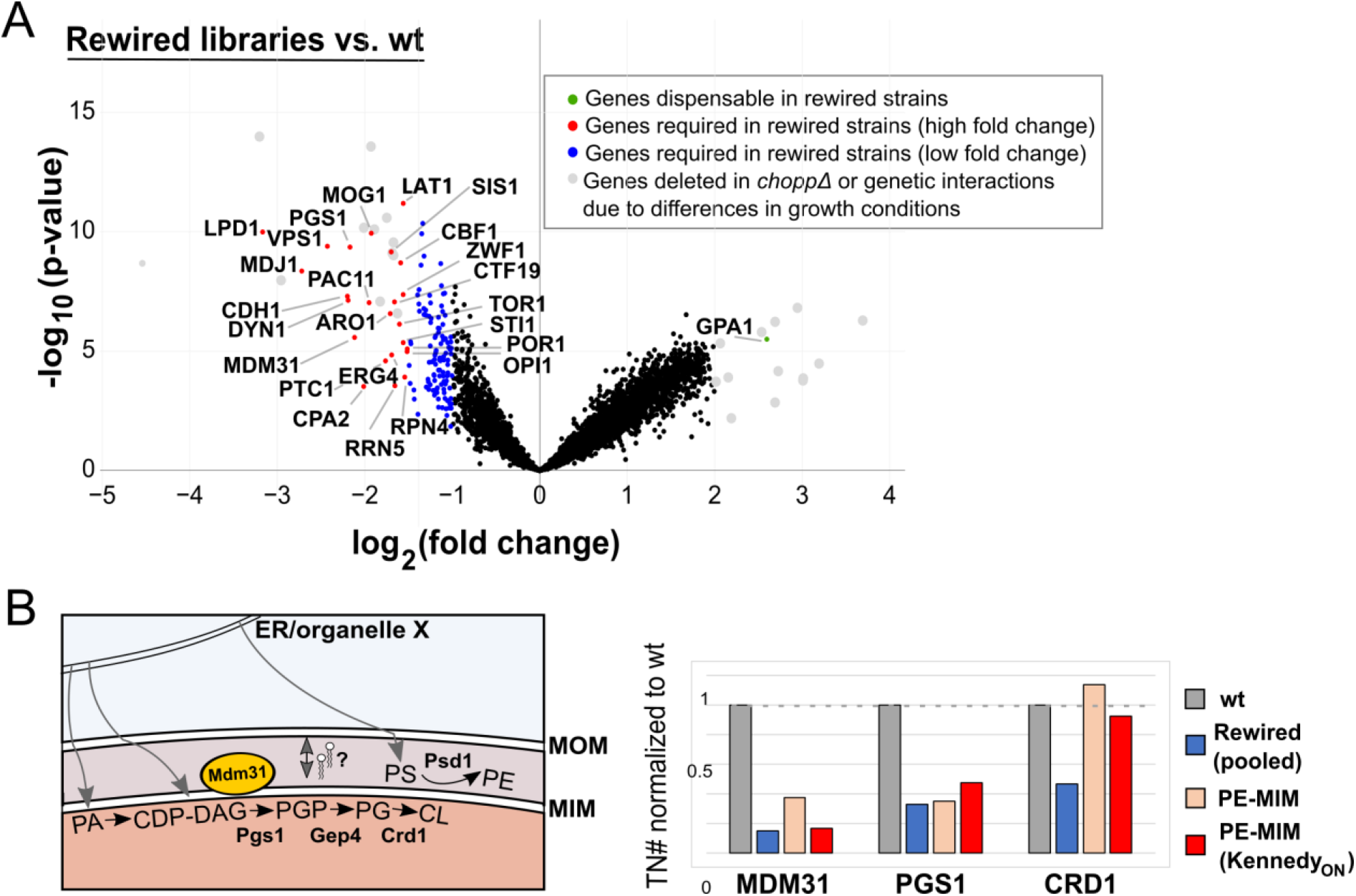
Differential genetic requirements common to rewired libraries. **A)** Volcano plot comparing the number of transposon insertions per gene of all 24 rewired libraries and two wild-type (wt) libraries. Genes that are significantly (*p*-value < 0.05) required (red, log2(fold change) < −1.5; blue, log_2_(fold change) < −1) or dispensable (green, log_2_(fold change > 2)) in the rewired libraries are highlighted. Genes depicted in grey are essential due to differential growth requirements in the preparation of the two wt libraries compared to the rewired strains, or correspond to genes deleted in the rewired strains. **B)** Requirement for mitochondrial lipid transport and biosynthesis genes in rewired libraries. Left panel, schematic depicting PE and cardiolipin (CL) synthesis in mitochondria. The inner mitochondrial putative lipid transporter Mdm31 is depicted in yellow. Grey arrows indicate the need for lipid transport from the ER or other organelles to mitochondria and transport between the MOM and MIM. Right panel, transposon numbers (normalized to wt libraries) for the indicated (set of) library/libraries for *MDM31*, *PGS1*, and *CRD1*.

In addition to transcriptional adaptation to altered lipidome, we observed many additional changes in the genetic requirement of rewired strains (Fig 5A, Suppl. Data 4).

These include an increased reliance on proteostasis regulators (*RPN4*, *MDJ1*), and on otherwise non-essential lipid biosynthesis pathways, such as cardiolipin (CL) synthesis (*PGS1*, *CRD1*) and the final step of ergosterol synthesis (*ERG4*) (Fig 5A). The mitochondrial inner membrane-targeted Psd rescues the requirement for CL synthesis in the transposon screen, consistent with previously published work highlighting redundant functions for PE and CL in mitochondria (Fig 5B) (Joshi *et al*, 2012; Gohil *et al*, 2005). In contrast, synthesis of the cardiolipin precursor phosphatidylglycerol (PG) is required in all libraries (Fig 5B). Moreover, analysis of genes involved in lipid biosynthetic pathways suggest a variable requirement for the enzymes involved in the synthesis of phospholipids such as PA, PI, PE, CL and the neutral lipid TAG (Fig S17).

Interestingly, transposons in the gene encoding the inner mitochondrial membrane protein Mdm31 are tolerated in wild-type libraries, but not in the rewired strains. *MDM31* genetically interacts with *PSD1* and is crucial for mitochondrial lipid homeostasis (Miyata *et al*, 2017). Notably, PE synthesis at the mitochondrial inner membrane by our chimeric construct (resembling mitochondrial localization of Psd1) only marginally rescues the number of transposons in the *chopp*Δ background (Fig 5B), implying that *mdm31* mutants are particularly sensitive to altered PE/PC levels and distribution in the absence of the endogenous CDP-DAG pathway. Mdm31 harbors a Chorein-N domain, a signature domain found in lipid transport proteins including Vps13 (Levine, 2019). Thus, it is tempting to speculate that the requirement for *MDM31* in the rewired conditions is due to a requirement for lipid transport across the mitochondrial intermembrane space.

### Csf1 is required for lipid adaptation in rewired conditions

As we relocalized lipid synthesis to distinct compartments, we expected that LTPs might become particularly necessary to distribute lipids in specific rewired Kennedy_OFF_ conditions. Genes satisfying this condition were scrutinized based on previous literature and structural predictions to identify LTP candidates. Csf1 emerged as a promising hit, as a HHpred analysis (Zimmermann *et al*, 2018) predicts an N-terminal region homologous to the Chorein-N domain found in Vps13, Atg2 and other potential lipid transporters including Mdm31 (Fig 6A) (Li *et al*, 2020; Levine, 2019). Transposons targeting the *CSF1* ORF were strongly depleted in Kennedy_OFF_ conditions (Fig 6A, S18) in the strain where PE production is targeted to the MIM and PC production to peroxisomes (hereafter referred to as PE-MIM/PC-pex). However, contrary to *VPS13*, which becomes required for growth in most libraries involving PE or PC synthesis at endosomes or mitochondria (Fig. 3C, 6A), the requirement for *CSF1* was specific to the PE-MIM/PC-pex conditions.

**Fig 6.**
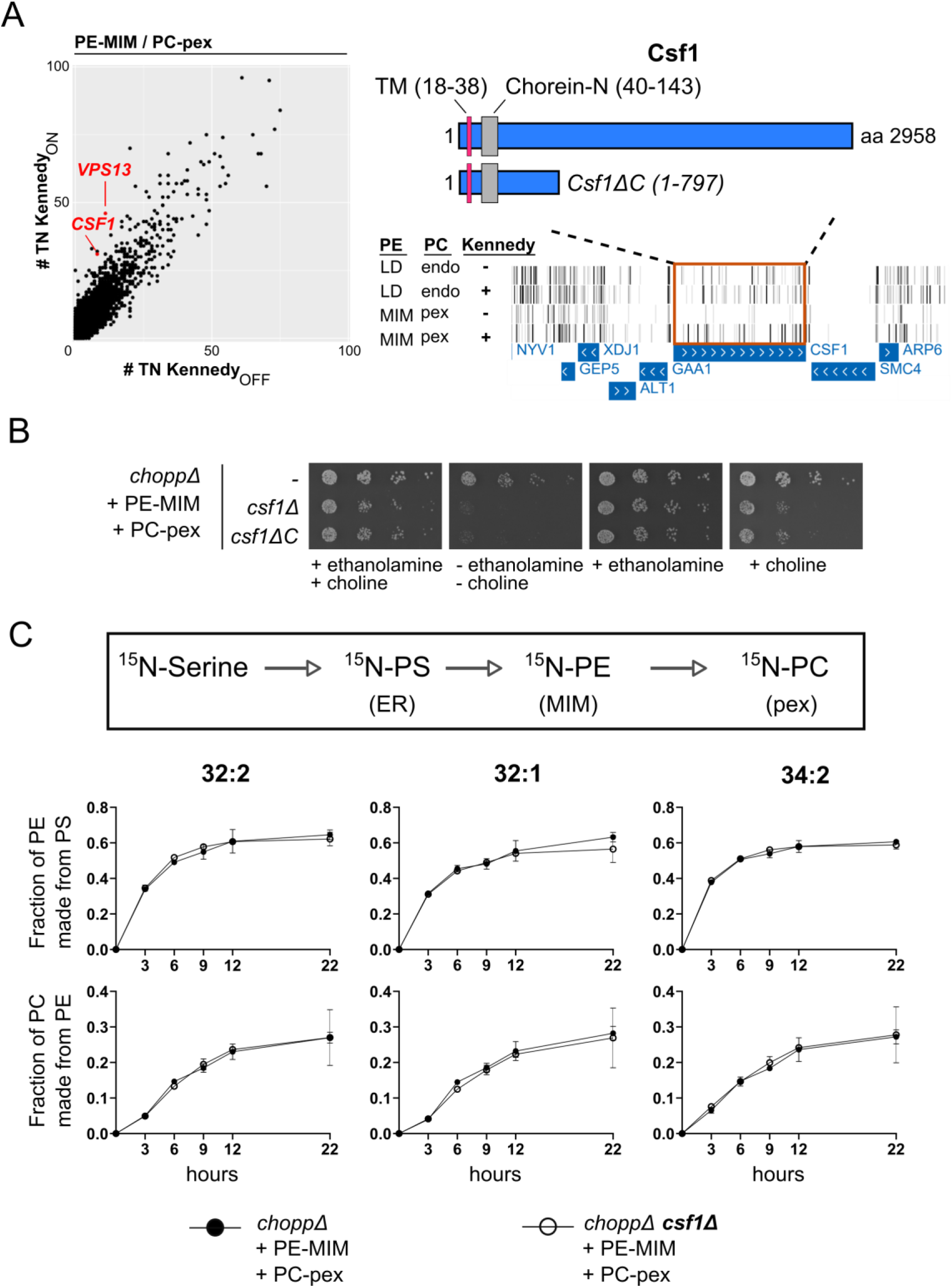
Csf1 is required specifically in the PE-MIM/PC-pex rewired condition. **A)** Left: comparison of the number of transposon insertions per gene (dots) for libraries with PE synthesis targeted to the MIM and PC synthesis targeted to peroxisomes, in Kennedy_ON_ and OFF conditions. *CSF1* and *VPS13* (highlighted in red) are more required in Kennedy_OFF_ conditions. Right: Transposon insertion profile displayed in the UCSC genome browser for the indicated libraries for the genomic region of *CSF1*. Bars correspond to transposons, rows to libraries. The boundaries of the full-length Csf1 protein, the truncated Csf1ΔC protein, the Chorein-N domain and the transmembrane (TM) domain are indicated. **B)** Five-fold serial dilutions of strains of the indicated genotypes grown at 30°C on SD medium supplemented with 10 mM ethanolamine and/or 10 mM choline. Growth assays are representative images of biological replicates. **C)** PS and PE transfer assay. Rewired strains were pulse labeled with ^15^N-serine. Lipids were extracted at the indicated time-points and measured by mass spectrometry. Graphs depict quantification of the appearance of labeled species of the indicated phospholipids normalized by the amount of its precursor lipid.

To validate the screen results, we generated a strain in which *CSF1* was deleted in the *choppΔ* PE-MIM/PC-pex background. As expected, *CSF1* was required for growth in Kennedy_OFF_ conditions (Fig 6B), but dispensable when PE production was redirected to endosomes and PC production to lipid droplets (Fig S19). We observed variable phenotypes when *CSF1* was deleted with different gene replacement cassettes, a phenomenon that might be attributed to the presence of essential genes at both ends of *CSF1*, the expression of which might be affected by the full deletion alleles. To avoid interference with neighboring genes, we truncated (rather than deleting) *CSF1* by introducing a stop codon after amino acid 797. This allele (Csf1ΔC) recapitulated the growth phenotype of the *CSF1* deletion in the rewired Kennedy_OFF_ condition and since it is less likely to interfere with neighboring genes, we used it for further loss-of-function experiments.

Interestingly, we found that ethanolamine supplementation promotes growth of rewired *csf1* mutants better than choline (Fig 6B), indicating that homeostasis of PE, and not PC, likely underlies the growth defect of the *csf1* mutant in the rewired conditions. Yet, no detectable change in phospholipid abundance could be uncovered by thin-layer chromatography of whole cell lipid extracts of *csf1ΔC* cells (Fig S20). To assess if Csf1 was involved in lipid transport between the ER and the MIM-targeted Psd, or between the MIM-targeted Psd and the peroxisome-targeted Pmt, we pulse-labeled the rewired strain with ^15^N-serine and monitored the production of PS, PE and PC over time in the absence of exogenous ethanolamine and choline. The appearance of ^15^N-labelled PE and PC should reflect on both the activities of the enzymes and on lipid transport rates between them. Surprisingly, the synthesis rates of PE and PC were identical in Csf1-proficient and -deficient cells, indicating that the growth defect of the rewired strain in the absence of Csf1 is not likely due to decreased lipid transport between ER, mitochondria and peroxisomes (Fig 6C).

To examine Csf1’s subcellular localization in wild-type cells, we genomically tagged its C-terminus with GFP. To test whether the Csf1-GFP allele is functional, we took advantage of the cold-sensitive phenotype of *csf1* mutants (Tokai *et al*, 2000). While *csf1ΔC* cells showed a severe growth phenotype at 16 ^o^C, cells expressing Csf1-GFP grew indistinguishably from wild-type cells, indicating that the GFP-tagged protein was functional (Fig 7B). Csf1-GFP localized to puncta that co-localized with an ER marker and are predominantly found near the cell periphery (Fig 7A). We found no evidence for obvious co-localization with either mitochondria or peroxisomes, and no indication of localization at contact sites between the ER and these organelles (Fig 7A). Given that Csf1 affects neither steady-state lipid class abundance nor transport to mitochondria and peroxisomes, and that Csf1 does not appear to localize at these organelles, we conclude that Csf1 function is likely affecting the lipid distribution in more indirect fashions, which may particularly affect strains with PE made in mitochondria and PC made in peroxisomes.

**Fig 7:**
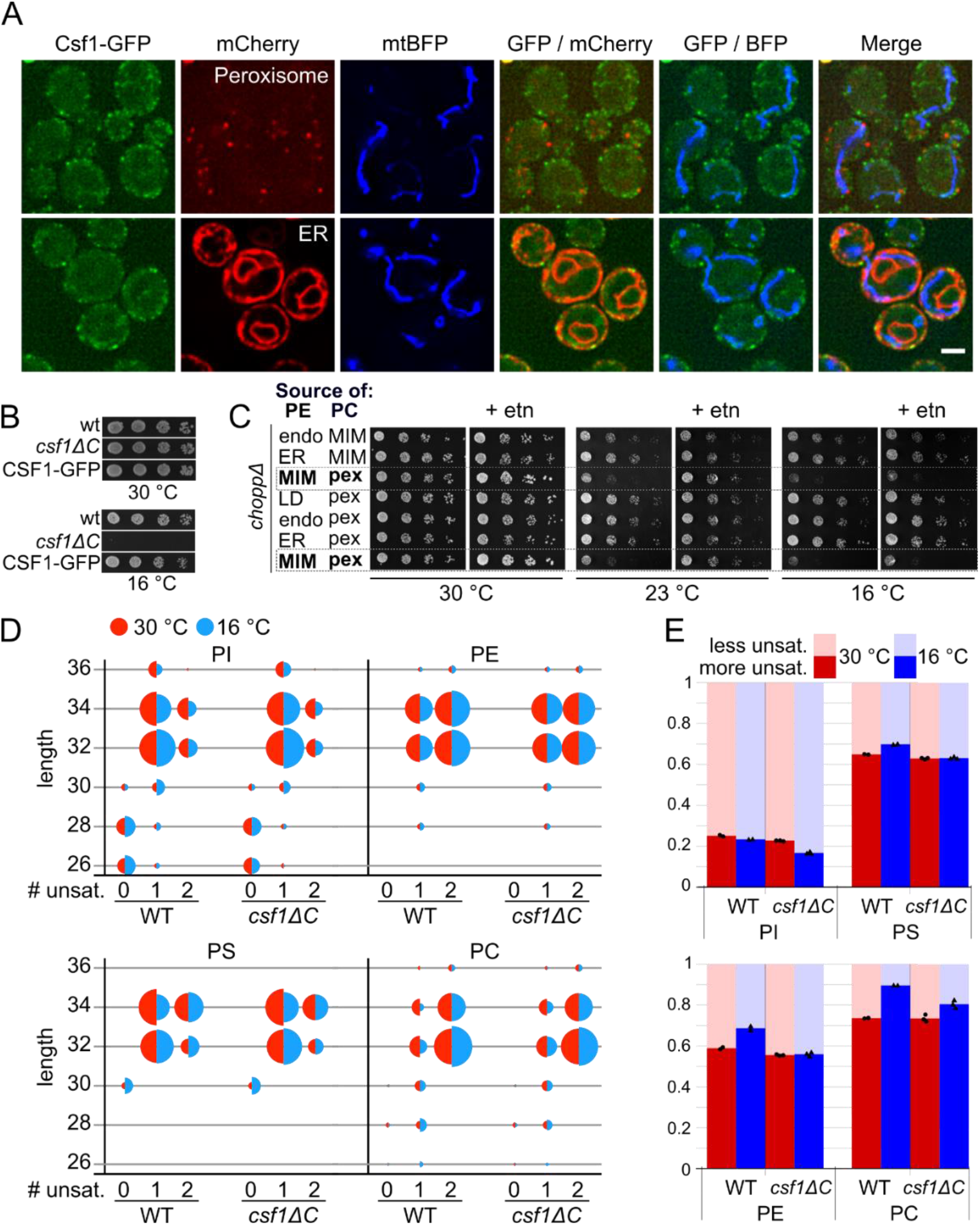
Csf1 is involved in phospholipid remodeling during cold stress. **A)** Top: Co-localization of Csf1-GFP cells in log phase with genomically integrated Pex10-mCherry (peroxisome marker) or mtBFP plasmid (mito marker). Bottom: Co-localization of log growing Csf1-GFP cells with mCherry-Ubc6(tail-anchor) (ER marker) or mtBFP plasmid (mito marker). **B)** Five-fold serial dilutions of strains of the indicated genotypes grown on YPD plates at 30 °C or at 16 °C. Growth assays are representative images of biological replicates. **C)** Five-fold serial dilutions of strains of the indicated genotypes grown at indicated temperatures, in the presence or absence of 10 mM ethanolamine (etn). **D)** Bubble plots depicting the changes in abundance for each phospholipid class, classified based on the fatty acid length and degree of unsaturation, in wild-type (WT) and *csf1ΔC* cells grown at 30°C (red-half) or at 16°C for 14.5 hours (blue-half). **E)** Bar graphs depicting the fraction of more saturated species (defined as fatty acid chains with one double bond for ≤30 carbon chains or with two double bonds for ≥32), and less unsaturated species (defined as fatty acid chains with zero double bond for ≤30 carbon chains or with one double bond for ≥32) for each phospholipid class with different head groups.

Apart from the lethality in rewired conditions reported here, the only other well-documented phenotype of *csf1* mutants is cold sensitivity (Tokai *et al*, 2000). While most cold-sensitive mutations affect lipid-independent processes like RNA metabolism (Noble & Guthrie, 1996), yeast cells also need to adapt their lipidome in cold conditions to maintain membrane fluidity (Klose *et al*, 2012). This raised the possibility that the PE-MIM/PC-pex rewired condition may yield a lipidome that is ill-adapted to cold, and that this inadequacy is synergistic with the effect of *csf1* mutation. Indeed, we find that PE-MIM/PC-pex rewiring causes a cold sensitivity at both 23 and 16 °C and the phenotype at 23 °C is rescued by ethanolamine supplementation (Fig 7C, dashed boxes). This cold-sensitive lipidome of PE-MIM/PC-pex cells may therefore explain the synthetic lethality with *CSF1.*

We thus wondered if loss of *CSF1* interferes with lipidomic adaptation to cold in cells with normal PE and PC biosynthesis. We quantified the levels of different phospholipid species by mass spectrometry in cells grown at 16 °C for 14.5 hours. We find that cells adapt their lipidome upon cold treatment by generally shortening the acyl chains and increasing the level of unsaturation (Fig 7D, Suppl. Data 5), consistent with previous knowledge (Klose *et al*, 2012). Interestingly, this adaptive response was blunted in *csf1ΔC* cells. Quantifying the degree of unsaturation (0 vs 1 double-bond for lipids with ≤30 carbons, or 1 vs. 2 double-bonds for lipids with ≥32 carbons) revealed that while wild-type cells undergo fatty-acid remodeling, *csf1* mutants failed to do so (Fig 7E). These data suggest that a defect in lipid remodeling from saturated to mono- and mono- to di-unsaturated species underlies the cold-sensitive phenotype of *csf1* mutants.

## Discussion

Here we show that synthesis of the phospholipids PE and PC can be re-localized to almost any combination of organelles, supporting yeast growth without the need for the Kennedy pathway. Although this surprising result implies that lipids must constantly shuttle from membrane to membrane using vesicular and non-vesicular routes, knowledge of the extent and direction of interorganelle lipid exchange remains limited. The viability of the rewired strains implies bidirectional transport of lipids to and from the organelles assayed here, which include mitochondria, ER, peroxisomes, lipid droplets, and endosomes. This conclusion is contingent on the proper targeting of the lipid modifying enzymes. While we have observed incomplete targeting of some constructs (peroxisome- and LD-targeted Pmt), in the case of the matrix-targeted Pmt, growth restoration is dependent on the mitochondrial SAM transporter Sam5, indicating that growth restoration is due to the activity of the properly targeted enzyme. Moreover, the uniqueness of transposon insertion profiles indicates that the localization of the enzymes matter, and therefore that the targeted fraction dominates over any mistargeted enzymes.

Bidirectional lipid transport could be direct (between the two organelles), indirect (using one or more intermediary organelle(s)), or a combination thereof. Regardless of the precise route(s) taken, our data suggest that the lipid transport network is more complex than previously thought and organelles considered to be mere lipid importers are also capable of export.

Contact sites have been identified for more than half of all possible pairwise combinations between organelles (Gatta & Levine, 2017). However, non-vesicular phospholipid trafficking has only been implied for a fraction of them. Whether all pairwise combinations of contact sites exist, which of these combinations contribute to phospholipid distribution in cells, and what determines transport directionality, will surely be the focus of many future studies. The approach of rewiring phospholipid biosynthesis pioneered here allows to isolate specific routes and thus facilitates tackling these pressing questions.

What molecular components allow yeast cells to survive rewiring and adapt to changes in organelle membrane composition? We identified genes required for survival by mapping genome-wide differential genetic requirements in the rewired strains using transposon mutagenesis. Interestingly, we found enrichment for genes involved in lipid metabolism, lipid trafficking, proteostasis, and vesicular trafficking, suggesting an important role for these pathways in maintaining functional membranes and organelles upon perturbations of the lipidome. We anticipated that cellular adaptation to the rewiring conditions necessitates sensors of membrane composition and stress. Membrane-associated proteins have been identified that sense and respond to lipid packing (Covino *et al*, 2016) and phospholipid composition (Henry *et al*, 2012). We find the ER lipid packing sensor Mga2 to be required in some of the rewired conditions, implying that the location of phospholipid synthesis can alter the properties of the ER membrane, thus necessitating Mga2 activation. On the other hand, Opi1, which senses the amount and protonation state of PA, is required in all rewired strains. It has been observed that PA protonation does not only depend on intracellular pH, but also on membrane composition, and in particular on the presence of PE head groups (Young *et al*, 2010; Kooijman *et al*, 2005). Possibly, Opi1 can sense altered PA levels or PE/PC ratios via changes in PA protonation.

Could we identify phospholipid transporters between organelles? As a proof of principle, the screen showed that *VPS13* is particularly important in Kennedy_OFF_ conditions in libraries where PE and PC synthesis are redirected to mitochondria and endosomes, two known localizations of this lipid transporter. In addition to Vps13, we identified another Chorein-N domain protein, Csf1, that is required in a specific rewired condition (PE-MIM/PC-pex). Surprisingly, Csf1 was neither directly influencing the global lipidome nor lipid transport rates between the PE and PC-producing enzymes. Instead, our analyses reveal a connection between Csf1 and adaptation to cold. *csf1* mutants are unable to remodel their lipidome in cold stress by increasing the levels of fatty acid unsaturation in phospholipids. Importantly, PE-MIM/PC-pex cells, like *csf1Δ*, were cold-sensitive, and both their cold-sensitivity and synthetic lethality with *csf1* were rescued by ethanolamine supplementation, indicating that defects in lipid homeostasis underlie *csf1* phenotypes.

The mechanism of Csf1 in lipidome remodeling remains unclear. Csf1 might be required for lipid transport from other organelles to the ER to allow their remodeling via ER-localized phospholipases and acyltransferases. Intriguingly, Csf1 homologues in higher organisms are implicated in lipid storage (lpd-3, *C. elegans*) and endosomal trafficking (*tweek* and KIA1109, *D. melanogaster* and *H. Sapiens*, respectively), consistent with a role in lipid transport. Mutations in KIA1109 are associated with the Alkuraya-Kučinskas syndrome, a severe neurological malformation disorder (McKay *et al*, 2003; Kane *et al*, 2019; Verstreken *et al*, 2009). Therefore, it will be important to understand whether and how Csf1 is involved in lipid transport.

Taken together, the extensive genetic interaction network reported here uncovers numerous lipid trafficking routes and genes required to handle lipid imbalances generated by specific lipid rewiring conditions. The data will serve as a rich source for future studies. They are accessible in the genome browser and can be interrogated via interactive plots that reveal significant differential genetic requirements. Correlation of the transposon insertion profiles allows identification of functionally related genes, and the conditions in which these are essential. We have shown that the data provide insight into general genetic requirements for handling membrane stress and can identify gene products that are essential in specific rewired conditions. Specifically, we uncovered a role for the conserved Chorein-N domain-containing protein Csf1 in lipidome remodeling upon cold treatment. This finding highlights a previously unappreciated role for Csf1 in lipid homeostasis, and provides a promising starting point to elucidate its function and regulation in yeast and other organisms.

## Materials and Methods

### Yeast strains and plasmids

Yeast strains, plasmids and primers used in this study are listed in Table S1, S2 and S3, respectively. Gene deletions were performed using genomic integration of PCR fragments (using oligos in Table S3) by homologous recombination following common procedures. We either used gene substitution by a selection marker using the Janke and Longtine toolboxes, as described previously (Janke *et al*, 2004; Longtine *et al*, 1998), or CRISPR-Cas9 mediated marker-less deletion of the ORF (Laughery *et al*, 2015). For CRISPR-Cas9 mediated gene deletions, we supplied the cells simultaneously with three DNA fragments; i) a Swa1-linearized guide RNA (gRNA) expression vector ii) annealed gRNA oligos with homology to the vector iii) a repair DNA template to delete the ORF. The DNA template was generated by amplifying ~ 500bp up- and downstream of the ORF and annealing the two fragments using extension overlap PCR as in (Walter *et al*, 2019). Colonies that had repaired the gRNA expression vector were selected. All gene deletions were confirmed by colony PCR.

Plasmids were generated using yeast gap-repair cloning. For *PSD* and *PMT* plasmids, overlapping DNA fragments were transformed together with linearized pRS415-TEFpr (Mumberg *et al*, 1995) using XbaI and XhoI. The pk*PSD* and aa*PMT* DNA sequences were ordered as gBlocks^®^ from IDT. The GFP sequence was amplified from pYM25 (Janke *et al*, 2004). Targeting signals were PCR-amplified directly from yeast genomic DNA, except for peroxisomes and endosomes, where the signal was incorporated in the primer sequence and PCR-amplified from plasmid pBK196, respectively. The *LEU2* markers of PSD plasmids were exchanged with *HIS3* (from pRS413) and/or *URA3* (from pRS416) to allow simultaneous expression of *PSD* and *PMT* (and other) plasmids. Briefly, pRS415-based *PSD* plasmids were digested with AgeI and transformed into yeast with *HIS3* or *URA3* PCR fragments with overlapping sequences to the vector. To generate the peroxisome marker, pRS416-TEFpr (Mumberg *et al*, 1995) was digested with XbaI and XhoI and transformed with a mCherry-SKL PCR product with overlapping sequences to the linearized vector. To generate the Lipid Droplet marker, the pRS413 vector was digested with SacI and transformed along with two overlapping PCR fragments containing the *ERG6* ORF with its native promoter, and mCherry DNA sequences.

### Growth assays

Cells were grown to mid-log phase in the appropriate synthetic defined (SD) drop-out medium (to select for the plasmids), diluted, and spotted on appropriate plates. Five-fold dilution series were used, with the first spot in each row corresponding to 2.5 μl of 0.2 OD_600_. In Kennedy_OFF_ conditions, plates were supplemented with 10 mM ethanolamine and choline. Cells were allowed to grow at 30 °C (unless noted otherwise) for 2-3 days before imaging.

### Transposition library generation

Briefly, yeast strains were transformed with pBK626, the plasmid that contains a Maize Ac/Ds transposase under the control of a galactose promoter, TRP1 ORF disrupted by the miniDS transposon, and 600 bp sequence overlapping with TRP1 to facilitate homology-directed repair of TRP1 after transposition. Individual clones were streaked on SC–HIS –URA –LEU (SC –HUL) and used to inoculate a pre-culture in SC –HUL + 2% Raffinose + 0.2% Glucose at an OD_600_ of 0.2, which was grown until saturation. Saturated pre-cultures were diluted to OD_600_ 0.2 in 400-800 mL YP + 2% Galactose medium and grown for ~56 hours to induce transposition. After transposition induction, cultures were washed twice with fresh SC medium, inoculated at an OD_600_ ranging between 0.3 and 1 in 2L of SC –LEU –TRP– HIS (SC –LTH) + 2% Glucose medium with and without supplemented ethanolamine and choline (10mM each) and grown until saturation. Cells were reseeded for a second growth round by diluting the saturated cultures in 300 mL of the appropriate medium and allowed to grow from an OD_600_ of 0.2 till saturation. Cells were harvested and frozen at −20°C until processed for DNA extraction and sequencing. To estimate the number of transposed clones, appropriate dilutions were plated on SC - TRP and SC-URA media at t=0 and t=56 hours. Colony count at t_0_ served to estimate the background of TRP1+ clones due to spontaneous recombination of pBK626. The number of cycles that cells underwent after transposition was determined by counting the TRP1+ cells at the start and end of each re-seed by plating appropriate dilutions on SC –TRP plates. Typically, we estimated between ~2×10^6^ and ~15×10^6^ individual transposition events per library and cells underwent between 13 and 15 cycles of growth in SC medium in Kennedy_ON_ and _OFF_ conditions.

### Sequencing

DNA was extracted from 0.5 g yeast pellets and processed for sequencing as described previously (Michel *et al*, 2017). Briefly, 2 μg of extracted DNA was processed by restriction digestion with NlaIII and DpnII, circularization and PCR amplification. A distinct 8bp illumina barcode was introduced in the PCR reverse primer for each library. Barcoded libraries were pooled and sequenced using the Nextseq High Output kit (Illumina) according to the manufacturer’s guidelines.

### Sequencing data analysis

Reads were aligned to the yeast genome and processed further to detect individual transposon insertion sites as described previously (Michel *et al*, 2017). The *bed* and *wig* output files were uploaded to the UCSC genome browser to display transposition events and read counts, respectively. Transposon and read counts per gene were normalized to the total number of transposons/reads mapped for each library for subsequent analyses.

To generate volcano plots, two sets of libraries were defined: a test set and a reference set. For both sets, the mean of normalized transposon counts or read counts per gene were computed. The fold change for each gene was calculated as the mean of the test set divided by that of the reference set and a *p*-value associated with this difference was computed using a Student’s t-test.

Hierarchical clustering of the libraries was performed in R (version 4.0.4) on the normalized transposon count per gene and clustered using the hclust() function using the ward.D method. The heatmap was visualized using the heatmap.2() function, using the dendrogram generated by hclust() to order the libraries and using a color scheme depicting the log2 fold change (TN count) for each gene in a library with respect to the mean TN count for that gene across libraries.

To identify genes that were variable across libraries, the StdDev was calculated for the normalized transposon count per gene across libraries. For GO term analysis, the 200 and 500 most variable genes (based on StdDev) were entered in the yeastmine webserver (Balakrishnan *et al*, 2012) and Holm-Bonferroni corrected GO term enrichments were computed.

Correlation clusterogram was computed as follows. Pearson Correlation coefficients were calculated for the normalized transposon count per gene for most variable genes (determined by the StdDev as described above) using the cor() function in R (version 4.0.4). The resulting correlation matrix was clustered in R using the heatmaply_cor() function of the heatmaply package (Moreland, 2009). A subset comprising of variable genes was used to reduce noise generated by small, biologically meaningless fluctuations in transposon number for less variable genes.

### Homology searches

Hidden Markov model homology searches that revealed the Chorein-N domain in Csf1 were performed using HHpred server from the MPI Bioinformatics Toolkit (Zimmermann *et al*, 2018).

### Fluorescence microscopy

Cells were grown to mid-log phase in synthetic dextrose (SD) medium using the appropriate amino acid drop-out mix for the selection of the plasmids. Cells were supplemented with 10 mM ethanolamine and 10 mM choline for activation of the Kennedy pathway. Images were acquired using a DeltaVision MPX microscope (Applied Precision) equipped with a 100× 1.40 NA oil UplanS-Apo objective lens (Olympus), a multicolor illumination light source, and a CoolSNAPHQ2 camera (Roper Scientific). Image acquisition was done at RT. Images were deconvolved with SoftWoRx software using the manufacturer’s parameters. Images were processed further using the FIJI ImageJ bundle. Z-projections using MAX intensities are depicted in the figures unless mentioned otherwise.

### Fm4-64 staining

For vacuolar staining, cells were pulse labeled with FM4-64 (Molecular Probes) at a concentration of 5 μg/ml for 20 min in the dark at 30 °C. Cells were washed twice with ice-cold media without FM4-64 and imaged subsequently.

### Construction of ‘dark’ GFP plasmids

‘Dark’ (G65T/G67A-GFP) GFP variants were created by Quikchange mutagenesis using the original *PSD* and *PMT* plasmids as backbone.

### Thin Layer Chromatography (TLC)

25 OD_600_ units of cells from a culture grown to mid-log phase were harvested, snap-frozen in liquid nitrogen and stored at −80°C until lipid extraction. Lipids were extracted as described previously with minor modifications (da Silveira dos Santos *et al*, 2014). Briefly, cells were washed in ice-cold water and subsequently resuspended in 2 ml of extraction solvent containing 95% ethanol, water, diethyl ether, pyridine, and 4.2 N ammonium hydroxide (v/v 15:15:5:1:0.18). After the addition of 300μL glass beads, samples were vortexed vigorously for 5 minutes and incubated at 60 °C for 20 min. Cell debris were pelleted by centrifugation at 1,800 ×g for 10 min and the supernatant was dried under a stream of nitrogen. The dried extract was resuspended in 2 mL of water-saturated butanol and sonicated for 5 min using a water bath sonicator. 1 ml of water was added and vortexed further for 2 min. After centrifugation, 250 μL of the upper butanol phase was collected, dried under a stream of nitrogen and reconstituted in 25 μL of chloroform:methanol (2:1, vol/vol) and loaded on a TLC plate (Sigma Aldrich Cat #1118450001). The plate was activated using 1.8% (w/v) boric acid in 100% ethanol and placed in a pre-equilibrated TLC chamber containing the mobile phase (chloroform/ethanol/water/triethylamine (30/35/7/35, v/v)). The solvent was allowed to run till the mobile phase was ~2 cm before the end of the plate. After drying, lipids were visualized by spraying the plate with a primuline solution (5mg in 100ml acetone/water (80/20 v/v)). The plate was imaged using the ethidium bromide program on the BioRad ChemiDoc™ MP Imaging System.

### Quantification of lipid signal on TLC

Images of TLC plates were quantified using the FIJI ImageJ bundle. The ImageJ gels analysis tool was used to plot the lane profile for each sample and the peak area corresponding to individual lipid species was quantified. The amount of lipid species was expressed as the fraction of the total lipid species quantified in the sample/lane. Two biological replicates were used for quantification.

### ^15^N-serine pulse labeling

Precultures in synthetic minimal medium were diluted to 0.05 OD_600_/ml in 25 ml and grown overnight in the presence of 10 mM ethanolamine and choline. Next day, when the cultures reached an OD_600_ of ~3, they were pelleted, washed in minimal medium without ethanolamine and choline, and diluted to 0.8 OD_600_/ml in 20 ml. To start the pulse labeling, 3mM 15N-serine was added. At the indicated time points, 8 OD_600_ of cells were pelleted, snap-frozen and stored at −80°C.

### Cold stress experiment

Cultures of wild-type or *csf1ΔC* cells were grown until mid-log phase at 30 °C (t=0). Subsequently, cultures were diluted to OD_600_ 0.3 and grown for an additional 14.5 hours at 16 °C. At t=0 and t=14.5 h, 8 OD_600_ of cells were pelleted, snap-frozen and stored at −80°C until further analysis. The experiment was performed using two (wt) or three (*csf1ΔC*) biological replicates.

### Lipid extraction and lipidomics analysis

Lipids were extracted as described for the TLC analysis and resuspended in 50% methanol. LC analysis was performed as described previously with several modifications (Castro-Perez *et al*, 2010). Phospholipids were separated on a nanoAcquity UPLC (Waters) equipped with a HSS T3 capillary column (150 m ×30mm, 1.8 m particle size, Waters), applying a 10 min linear gradient of buffer A (5 mM ammonium acetate in acetonitrile/water 60:40) and B (5 mM ammonium acetate in isopropanol/acetonitrile 90:10) from 10% B to 100% B. Conditions were kept at 100% B for the next 7 min, followed by a 8 min re-equilibration to 10% B. The injection volume was 1 μL. The flow rate was constant at 2.5 μl/min.

The UPLC was coupled to QExactive mass spectrometer (Thermo) by a nanoESI source (New Objective Digital PicoView^®^ 550). The source was operated with a spray voltage of 2.9 kV in positive mode and 2.5 kV in negative mode. Sheath gas flow rate was set to 25 and 20 for positive and negative mode, respectively. MS data was acquired using either positive or negative polarization, alternating between full MS and all ion fragmentation (AIF) scans. Full scan MS spectra were acquired in profile mode from 107-1600 m/z with an automatic gain control target of 1e6, an Orbitrap resolution of 70`000, and a maximum injection time of 200 ms. AIF spectra were acquired from 107-1600 m/z with an automatic gain control value of 5e4, a resolution of 17`500, a maximum injection time of 50 ms and fragmented with a normalized collision energy of 20, 30 and 40 (arbitrary units). Generated fragment ions were scanned in the linear trap. Positive-ion-mode was employed for monitoring PC and negative-ion-mode was used for monitoring PS and PE. Lipid species were identified based on their m/z and elution time. We used a standard mixture comprising PS 10:0/10:0, PE 17:0/17:0 and PC 17:0/17:0 for deriving an estimate of specific elution times. Lipid intensities were quantified using the Skyline software (Adams *et al*, 2020).

## Acknowledgements

We are thankful to members of the Peter and Kornmann laboratories for discussions and helpful suggestions. Microscopy analysis was carried out at the ETH Zürich ScopeM facility, and lipidomic measurements and Illumina sequencing at the Functional Genomics Center Zurich (FGCZ). We especially thank Dr. Sebastian Streb and Dr. Endre Laczko of the FGCZ Metabolomics division for establishing and optimizing lipidomics workflows, and for excellent technical guidance. The Kornmann lab is supported by grants from the Swiss National Science Foundation (SNSF, 31003A_179549) and the Wellcome Trust (214291/A/18/Z). Work in the Peter laboratory was supported by the SNSF, the Synapsis Foundation and ETH Zürich. A.T.J.P was supported by ETH Zürich/Institute of Biochemistry and a Spark grant of the SNSF (CRSK-3_190364).

The authors declare no competing financial interests.

## Author contributions

A.T. John Peter, S.N.S. van Schie and B. Kornmann conceived the study. A.T. John Peter and S.N.S. van Schie designed and performed the experiments under the supervision of M. Peter and B. Kornmann. A.T. John Peter, S.N.S. van Schie and B. Kornmann analyzed data. S.N.S. van Schie and N.J. Cheung performed bioinformatics analyses. N.J. Cheung carried out work on making the SATAY data available on a web interface. A.H. Michel designed and constructed the plasmid for the SATAY screens. S.N.S. van Schie, A.T. John Peter and B. Kornmann wrote the paper with input from all the authors.

## References

Adams KJ, Pratt B, Bose N, Dubois LG, St. John-Williams L, Perrott KM, Ky K, Kapahi P, Sharma V, Maccoss MJ, et al (2020) Skyline for Small Molecules: A Unifying Software Package for Quantitative Metabolomics. J Proteome Res 19: 1447–1458

AhYoung AP, Lu B, Cascio D & Egea PF (2017) Crystal structure of Mdm12 and combinatorial reconstitution of Mdm12/Mmm1 ERMES complexes for structural studies. Biochem Biophys Res Commun 488: 129–135

Balakrishnan R, Park J, Karra K, Hitz BC, Binkley G, Hong EL, Sullivan J, Micklem G & Michael Cherry J (2012) YeastMine—an integrated data warehouse for Saccharomyces cerevisiae data as a multipurpose tool-kit. Database 2012

Barbosa AD, Lim K, Mari M, Edgar JR, Gal L, Sterk P, Jenkins BJ, Koulman A, Savage DB, Schuldiner M, et al (2019) Compartmentalized Synthesis of Triacylglycerol at the Inner Nuclear Membrane Regulates Nuclear Organization. Dev Cell 50: 755–766.e6

Carman GM & Henry SA (1999) Article in Progress in Lipid Research.

Castro-Perez JM, Kamphorst J, Degroot J, Lafeber F, Goshawk J, Yu K, Shockcor JP, Vreeken RJ & Hankemeier T (2010) Comprehensive LC-MSE lipidomic analysis using a shotgun approach and its application to biomarker detection and identification in osteoarthritis patients. J Proteome Res 9: 2377–2389

Choi JY, Augagneur Y, Mamoun C Ben & Voelker DR (2012) Identification of gene encoding Plasmodium knowlesi phosphatidylserine decarboxylase by genetic complementation in yeast and characterization of in vitro maturation of encoded enzyme. J Biol Chem 287: 222–232

Costanzo M, VanderSluis B, Koch EN, Baryshnikova A, Pons C, Tan G, Wang W, Usaj M, Hanchard J, Lee SD, et al (2016) A global genetic interaction network maps a wiring diagram of cellular function. Science (80-) 353

Covino R, Ballweg S, Stordeur C, Michaelis JB, Puth K, Wernig F, Bahrami A, Ernst AM, Hummer G & Ernst R (2016) A Eukaryotic Sensor for Membrane Lipid Saturation. Mol Cell 63: 49–59

DeMille D, Pape JA, Bikman BT, Ghassemian M & Grose JH (2019) The regulation of Cbf1 by PAS kinase is a pivotal control point for lipogenesis vs. respiration in saccharomyces cerevisiae. G3 Genes, Genomes, Genet 9: 33–46

Elbaz-Alon Y, Rosenfeld-Gur E, Shinder V, Futerman AH, Geiger T & Schuldiner M (2014) A dynamic interface between vacuoles and mitochondria in yeast. Dev Cell 30: 95–102

Friedman JR, Kannan M, Toulmay A, Jan CH, Weissman JS, Prinz WA & Nunnari J (2018) Lipid Homeostasis Is Maintained by Dual Targeting of the Mitochondrial PE Biosynthesis Enzyme to the ER. Dev Cell 44: 261–270.e6

Gatta AT & Levine TP (2017) Piecing Together the Patchwork of Contact Sites. Trends Cell Biol 27: 214–229 doi:10.1016/j.tcb.2016.08.010 [PREPRINT]

Gohil VM, Thompson MN & Greenberg ML (2005) Synthetic lethal interaction of the mitochondrial phosphatidylethanolamine and cardiolipin biosynthetic pathways in Saccharomyces cerevisiae. J Biol Chem 280: 35410–35416

Gulshan K, Shahi P & Moye-Rowley WS (2010) Compartment-specific Synthesis of Phosphatidylethanolamine Is Required for Normal Heavy Metal Resistance. Mol Biol Cell 21: 443–455

Hanada T, Kashima Y, Kosugi A, Koizumi Y, Yanagida F & Udaka S (2001) A gene encoding phosphatidylethanolamine N-methyltransferase from Acetobacter aceti and some properties of its disruptant. Biosci Biotechnol Biochem 65: 2741–2748

Henry SA, Kohlwein SD & Carman GM (2012) Metabolism and regulation of glycerolipids in the yeast Saccharomyces cerevisiae. Genetics 190: 317–349

Hönscher C, Mari M, Auffarth K, Bohnert M, Griffith J, Geerts W, van der Laan M, Cabrera M, Reggiori F & Ungermann C (2014) Cellular metabolism regulates contact sites between vacuoles and mitochondria. Dev Cell 30: 86–94

Janke C, Magiera MM, Rathfelder N, Taxis C, Reber S, Maekawa H, Moreno-Borchart A, Doenges G, Schwob E, Schiebel E, et al (2004) A versatile toolbox for PCR-based tagging of yeast genes: new fluorescent proteins, more markers and promoter substitution cassettes. Yeast 21: 947–962

Jeong H, Park J, Jun Y & Lee C (2017) Crystal structures of Mmm1 and Mdm12–Mmm1 reveal mechanistic insight into phospholipid trafficking at ER-mitochondria contact sites. Proc Natl Acad Sci U S A 114: E9502–E9511

Jeong H, Park J & Lee C (2016) Crystal structure of Mdm12 reveals the architecture and dynamic organization of the ERMES complex. EMBO Rep 17: 1857–1871

John Peter AT, Herrmann B, Antunes D, Rapaport D, Dimmer KS & Kornmann B (2017) Vps13-Mcp1 interact at vacuole-mitochondria interfaces and bypass ER-mitochondria contact sites. J Cell Biol 216: 3219–3229

Joshi AS, Thompson MN, Fei N, Ttemann MH & Greenberg ML (2012) Cardiolipin and mitochondrial phosphatidylethanolamine have overlapping functions in mitochondrial fusion in Saccharomyces cerevisiae. J Biol Chem 287: 17589–17597

Kane MS, Diamonstein CJ, Hauser N, Deeken JF, Niederhuber JE & Vilboux T (2019) Endosomal trafficking defects in patient cells with KIAA1109 biallelic variants. Genes Dis 6: 56–67

Kennedy EP & Weiss SB (1956) The function of cytidine coenzymes in the biosynthesis of phospholipides. J Biol Chem 222: 193–214

Klose C, Surma MA, Gerl MJ, Meyenhofer F, Shevchenko A & Simons K (2012) Flexibility of a eukaryotic lipidome - insights from yeast lipidomics. PLoS One 7: 35063

Kobayashi S, Mizuike A, Horiuchi H, Fukuda R & Ohta A (2014) Mitochondrially-targeted bacterial phosphatidylethanolamine methyltransferase sustained phosphatidylcholine synthesis of a Saccharomyces cerevisiae ∆pem1 ∆pem2 double mutant without exogenous choline supply. Biochim Biophys Acta - Mol Cell Biol Lipids 1841: 1264–1271

Kooijman EE, Carter KM, Van Laar EG, Chupin V, Burger KNJ & De Kruijff B (2005) What makes the bioactive lipids phosphatidic acid and lysophosphatidic acid so special? Biochemistry 44: 17007–17015

Kornmann B, Currie E, Collins SR, Schuldiner M, Nunnari J, Weissman JS & Walter P (2009) An ER-mitochondria tethering complex revealed by a synthetic biology screen. Science (80-) 325: 477–481

Krogan NJ, Dover J, Khorrami S, Greenblatt JF, Schneider J, Johnston M & Shilatifard A (2002) COMPASS, a histone H3 (lysine 4) methyltransferase required for telomeric silencing of gene expression. J Biol Chem 277: 10753–10755

Kumar N, Leonzino M, Hancock‑Cerutti W, Horenkamp FA, Li PQ, Lees JA, Wheeler H, Reinisch KM & De Camilli P (2018) VPS13A and VPS13C are lipid transport proteins differentially localized at ER contact sites. J Cell Biol 217: 3625–3639

Lang AB, John Peter AT, Walter P & Kornmann B (2015) ER-mitochondrial junctions can be bypassed by dominant mutations in the endosomal protein Vps13. J Cell Biol 210: 883–90

Laughery MF, Hunter T, Brown A, Hoopes J, Ostbye T, Shumaker T & Wyrick JJ (2015) New vectors for simple and streamlined CRISPR-Cas9 genome editing in *Saccharomyces cerevisiae*. Yeast 32: 711–720

Lees JA & Reinisch KM (2020) Inter-organelle lipid transfer: a channel model for Vps13 and chorein-N motif proteins. Curr Opin Cell Biol 65: 66–71 doi:10.1016/j.ceb.2020.02.008 [PREPRINT]

Levine TP (2019) Remote homology searches identify bacterial homologues of eukaryotic lipid transfer proteins, including Chorein-N domains in TamB and AsmA and Mdm31p. BMC Mol Cell Biol 20: 1–12

Li PQ, Lees JA, Patrick Lusk C & Reinisch KM (2020) Cryo-EM reconstruction of a VPS13 fragment reveals a long groove to channel lipids between membranes. J Cell Biol 219

Longtine MS, Mckenzie III A, Demarini DJ, Shah NG, Wach A, Brachat A, Philippsen P & Pringle JR (1998) Additional modules for versatile and economical PCR-based gene deletion and modification in Saccharomyces cerevisiae. Yeast 14: 953–961

Marobbio CMT, Agrimi G, Lasorsa FM & Palmieri F (2003) Identification and functional reconstitution of yeast mitochondrial carrier for S-adenosylmethionine. EMBO J 22: 5975–5982

McKay RM, McKay JP, Avery L & Graff JM (2003) C. elegans: A model for exploring the genetics of fat storage. Dev Cell 4: 131–142

Michel AH, Hatakeyama R, Kimmig P, Arter M, Peter M, Matos J, De Virgilio C & Kornmann BT (2017) Functional mapping of yeast genomes by saturated transposition. Elife 6

Michel AH, van Schie S, Mosbach A, Scalliet G & Kornmann B (2019) Exploiting homologous recombination increases SATAY efficiency for loss- And gain-of-function screening. bioRxiv: 866483 doi:10.1101/866483 [PREPRINT]

Miyata N, Goda N, Matsuo K, Hoketsu T & Kuge O (2017) Cooperative function of Fmp30, Mdm31, and Mdm32 in Ups1-independent cardiolipin accumulation in the yeast Saccharomyces cerevisiae. Sci Rep 7: 1–12

Mizuguchi G, Shen X, Landry J, Wu WH, Sen S & Wu C (2004) ATP-Driven Exchange of Histone H2AZ Variant Catalyzed by SWR1 Chromatin Remodeling Complex. Science (80-) 303: 343–348

Moreland K (2009) Diverging color maps for scientific visualization. In Lecture Notes in Computer Science (including subseries Lecture Notes in Artificial Intelligence and Lecture Notes in Bioinformatics) pp 92–103. Springer, Berlin, Heidelberg

Mumberg D, Müller R & Funk M (1995) Yeast vectors for the controlled expression of heterologous proteins in different genetic backgrounds. Gene 156: 119–122

Noble SM & Guthrie C (1996) Identification of Novel Genes Required for Yeast Pre-mRNA Splicing by Means of Cold-Sensitive Mutations. Genetics 143

Park J-S, Thorsness MK, Policastro R, McGoldrick LL, Hollingsworth NM, Thorsness PE & Neiman AM (2016) Yeast Vps13 promotes mitochondrial function and is localized at membrane contact sites. Mol Biol Cell 27: 2435–49

Quon E, Sere YY, Chauhan N, Johansen J, Sullivan DP, Dittman JS, Rice WJ, Chan RB, Di Paolo G, Beh CT, et al (2018) Endoplasmic reticulum-plasma membrane contact sites integrate sterol and phospholipid regulation. PLoS Biol 16: e2003864

Raychaudhuri S & Prinz WA (2008) Nonvesicular phospholipid transfer between peroxisomes and the endoplasmic reticulum. Proc Natl Acad Sci U S A 105: 15785–15790

Reinisch KM & Prinz WA (2021) Mechanisms of nonvesicular lipid transport.

Rosenberger S, Connerth M, Zellnig G & Daum G (2009) Phosphatidylethanolamine synthesized by three different pathways is supplied to peroxisomes of the yeast Saccharomyces cerevisiae. Biochim Biophys Acta - Mol Cell Biol Lipids 1791: 379–387

Schnabl M, Daum G & Pichler H (2005) Multiple lipid transport pathways to the plasma membrane in yeast. Biochim Biophys Acta - Mol Cell Biol Lipids 1687: 130–140

Schuldiner M & Bohnert M (2017) A different kind of love –lipid droplet contact sites. Biochim Biophys Acta - Mol Cell Biol Lipids 1862: 1188–1196 doi:10.1016/j.bbalip.2017.06.005 [PREPRINT]

Sebastian TT, Baldridge RD, Xu P & Graham TR (2012) Phospholipid flippases: Building asymmetric membranes and transport vesicles. Biochim Biophys Acta - Mol Cell Biol Lipids 1821: 1068–1077 doi:10.1016/j.bbalip.2011.12.007 [PREPRINT]

Shai N, Yifrach E, Van Roermund CWT, Cohen N, Bibi C, Ijlst L, Cavellini L, Meurisse J, Schuster R, Zada L, et al (2018) Systematic mapping of contact sites reveals tethers and a function for the peroxisome-mitochondria contact. Nat Commun 9: 1–13

Shetty A & Lopes JM (2010) Derepression of INO1 transcription requires cooperation between the Ino2p-Ino4p heterodimer and Cbf1p and recruitment of the ISW2 chromatin-remodeling complex. Eukaryot Cell 9: 1845–1855

Shiino H, Furuta S, Kojima R, Kimura K, Endo T & Tamura Y (2021) Phosphatidylserine flux into mitochondria unveiled by organelle-targeted Escherichia coli phosphatidylserine synthase PssA. FEBS J 288: 3285–3299

da Silveira dos Santos AX, Riezman I, Aguilera-Romero M-A, David F, Piccolis M, Loewith R, Schaad O & Riezman H (2014) Systematic lipidomic analysis of yeast protein kinase and phosphatase mutants reveals novel insights into regulation of lipid homeostasis. Mol Biol Cell 25: 3234–3246

Siniossoglou S, Santos-Rosa H, Rappsilber J, Mann M & Hurt E (1998) A novel complex of membrane proteins required for formation of a spherical nucleus. EMBO J 17: 6449–6464

Tanaka K, Nakafuku M, Satoh T, Marshall MS, Gibbs JB, Matsumoto K, Kaziro Y & Toh-e A (1990) S. cerevisiae genes IRA1 and IRA2 encode proteins that may be functionally equivalent to mammalian ras GTPase activating protein. Cell 60: 803–807

Tokai M, Kawasaki H, Kikuchi Y & Ouchi K (2000) Cloning and characterization of the CSF1 gene of Saccharomyces cerevisiae, which is required for nutrient uptake at low temperature. J Bacteriol 182: 2865–2868

Vance JE, Aasman EJ & Szarka R (1991) Brefeldin a does not inhibit the movement of phosphatidylethanolamine from its sites of synthesis to the cell surface. J Biol Chem 266: 8241–8247

Verstreken P, Ohyama T, Haueter C, Habets RLP, Lin YQ, Swan LE, Ly C V., Venken KJT, De Camilli P & Bellen HJ (2009) Tweek, an Evolutionarily Conserved Protein, Is Required for Synaptic Vesicle Recycling. Neuron 63: 203–215

Walter JM, Schubert MG, Kung SH, Hawkins K, Platt DM, Hernday AD, Mahatdejkul-Meadows T, Szeto W, Chandran SS, Newman JD, et al (2019) Method for Multiplexed Integration of Synergistic Alleles and Metabolic Pathways in Yeasts via CRISPR-Cas9. In Methods in Molecular Biology pp 39–72. Humana Press Inc.

Young BP, Shin JJH, Orij R, Chao JT, Li SC, Guan XL, Khong A, Jan E, Wenk MR, Prinz WA, et al (2010) Phosphatidic acid is a pH biosensor that links membrane biogenesis to metabolism. Science (80-) 329: 1085–1088

Zimmermann L, Stephens A, Nam SZ, Rau D, Kübler J, Lozajic M, Gabler F, Söding J, Lupas AN & Alva V (2018) A Completely Reimplemented MPI Bioinformatics Toolkit with a New HHpred Server at its Core. J Mol Biol 430: 2237–2243

