## Supplemental figures S1-S20 for "Rewiring phospholipid biosynthesis reveals robustness in membrane homeostasis and uncovers lipid regulatory players"

### Supplementary Figures

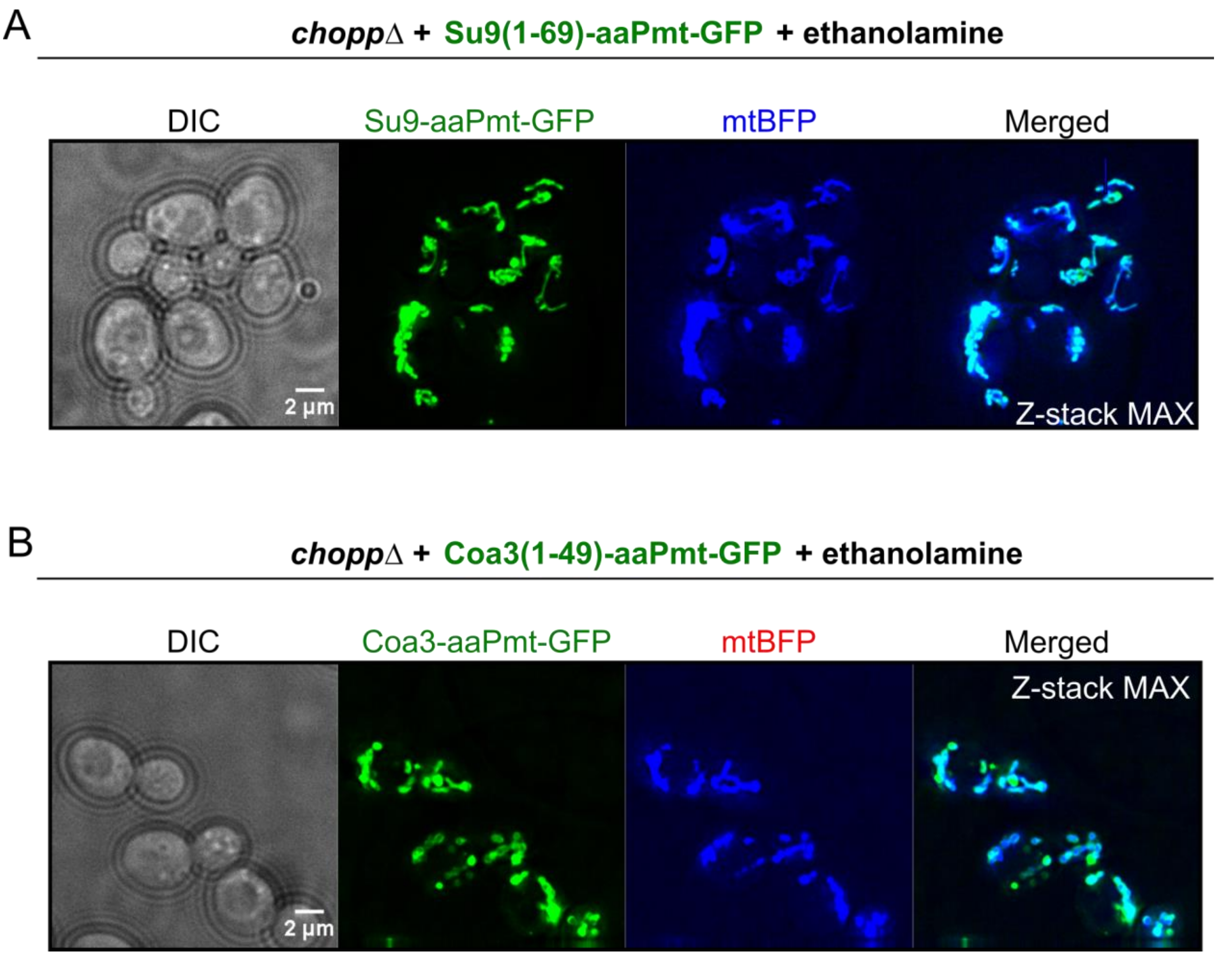


**Fig S1. Co-localization of the Pmt mitochondrial constructs with a mitochondrial marker.**

A) Co-localization of Su9-aaPmt-GFP (MM) with mtBFP (MM marker) expressed in *choppΔ* cells, grown in SD medium supplemented with 10 mM ethanolamine. Images shown are maximum intensity projections of several Z-sections.

B) Co-localization of Coa3-aaPmt-GFP (MIM) with mtBFP (MM marker) expressed in *choppΔ* cells, grown in SD medium supplemented with 10 mM ethanolamine. Images shown are maximum intensity projections of several Z-sections.

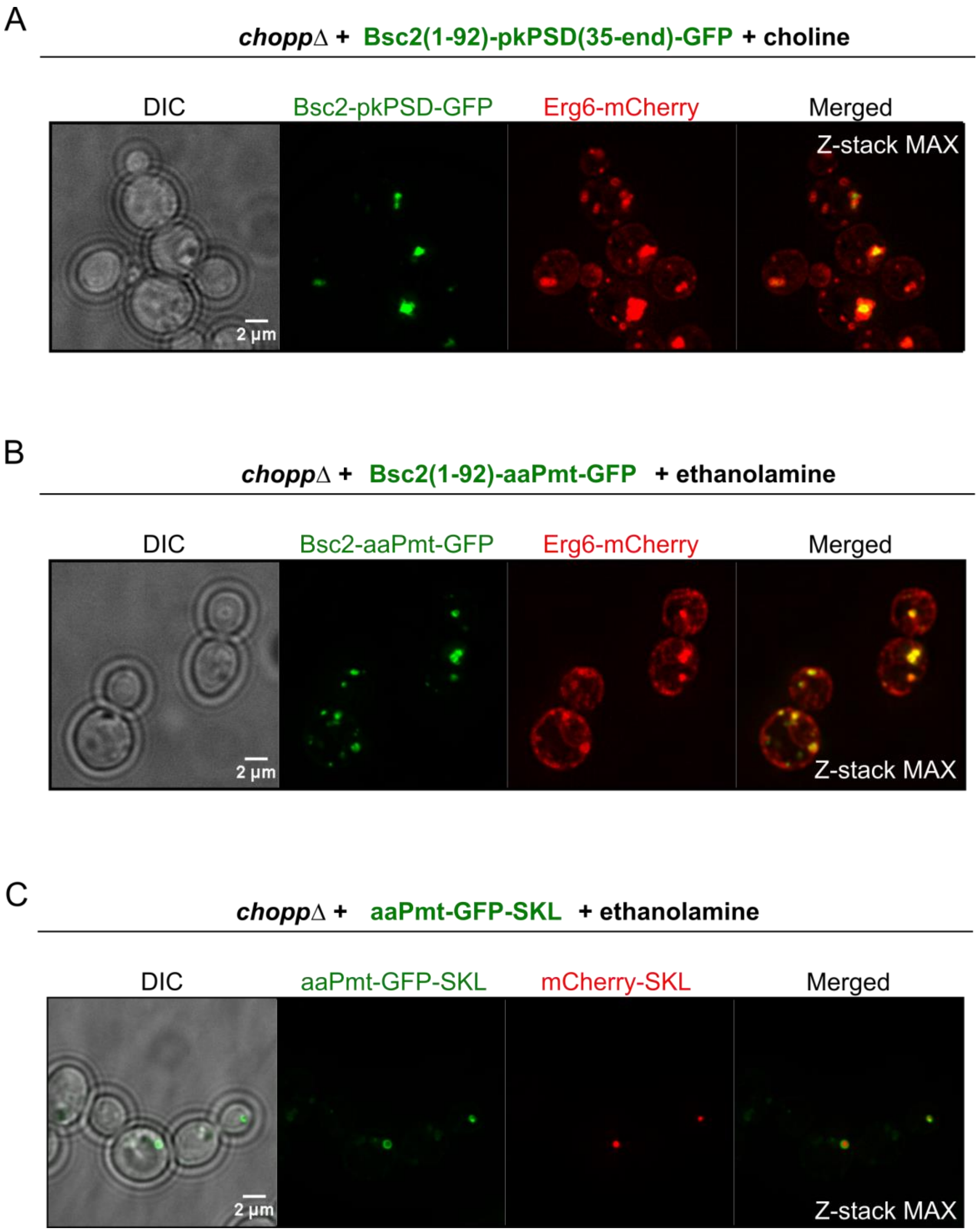


**Fig S2. Co-localization of chimeric constructs with a LD and peroxisome marker.**

A) Co-localization of Bsc2-pkPsd-GFP with Erg6-mCherry (LD marker) expressed in *choppΔ* cells, grown in SD medium supplemented with 10 mM choline. Images shown are maximum intensity projections of several Z-sections.

B) Co-localization of Bsc2-aaPmt-GFP with Erg6-mCherry (LD marker) expressed in *choppΔ* cells, grown in SD medium supplemented with 10 mM ethanolamine. Images shown are maximum intensity projections of several Z-sections.

C) Co-localization of aaPmt-GFP-SKL with mCherry-SKL (peroxisome marker) expressed in *choppΔ* cells, grown in SD medium supplemented with 10 mM ethanolamine. Images shown are maximum intensity projections of several Z-sections.

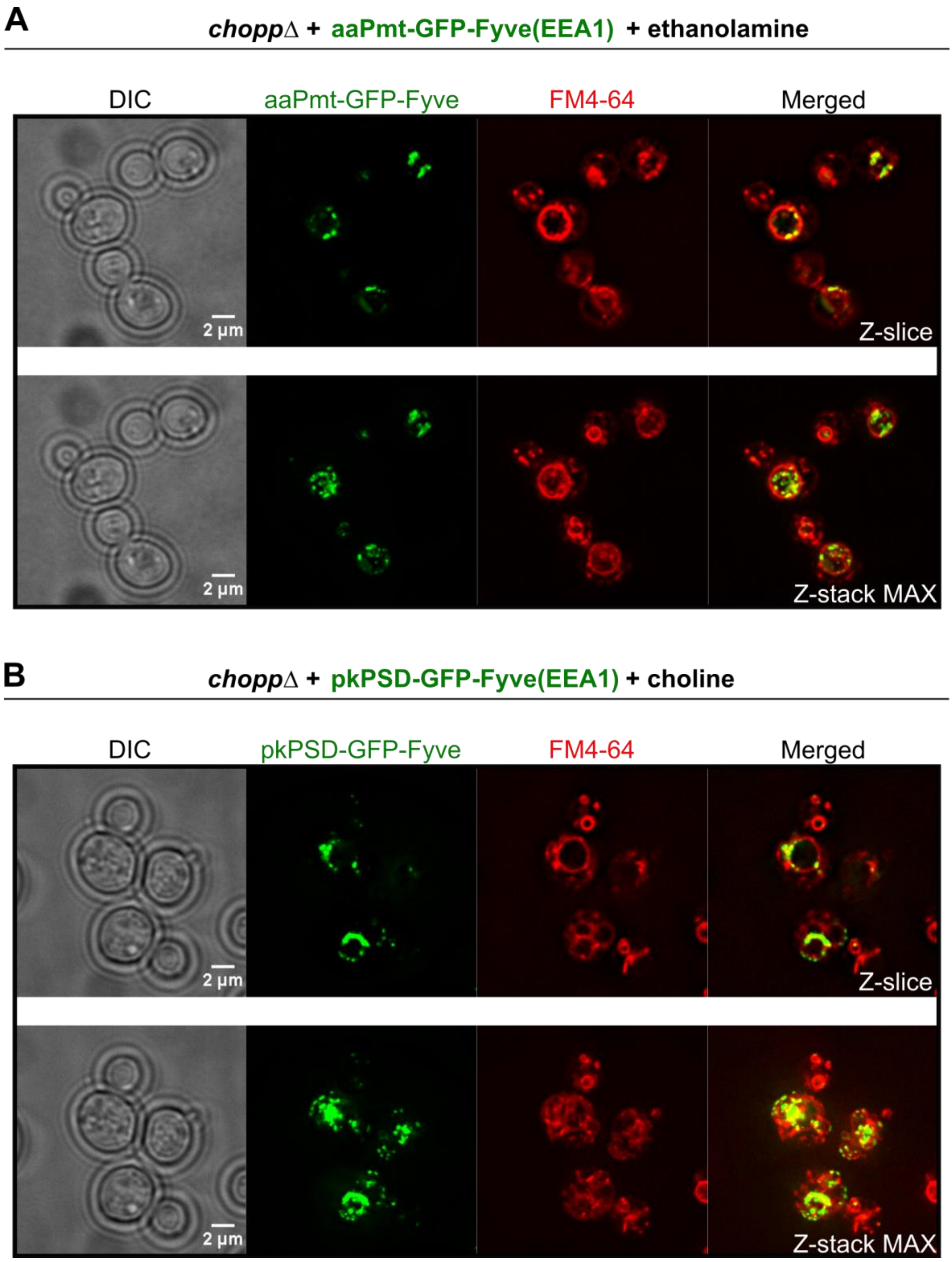


**Fig S3. Co-localization of FYVE-domain containing constructs with the endocytic compartment dye FM4-64.**

A) Co-localization of aaPmt -GFP-Fyve expressed in *choppΔ* cells with FM4-64, grown in SD medium supplemented with 10 mM ethanolamine. Images shown are either a single z-slice or maximum intensity projections of several Z-sections, as indicated.

B) Co-localization of pkPsd -GFP-Fyve expressed in *choppΔ* cells with FM4-64, grown in SD medium supplemented with 10 mM choline. Images shown are either a single z-slice or maximum intensity projections of several Z-sections, as indicated.

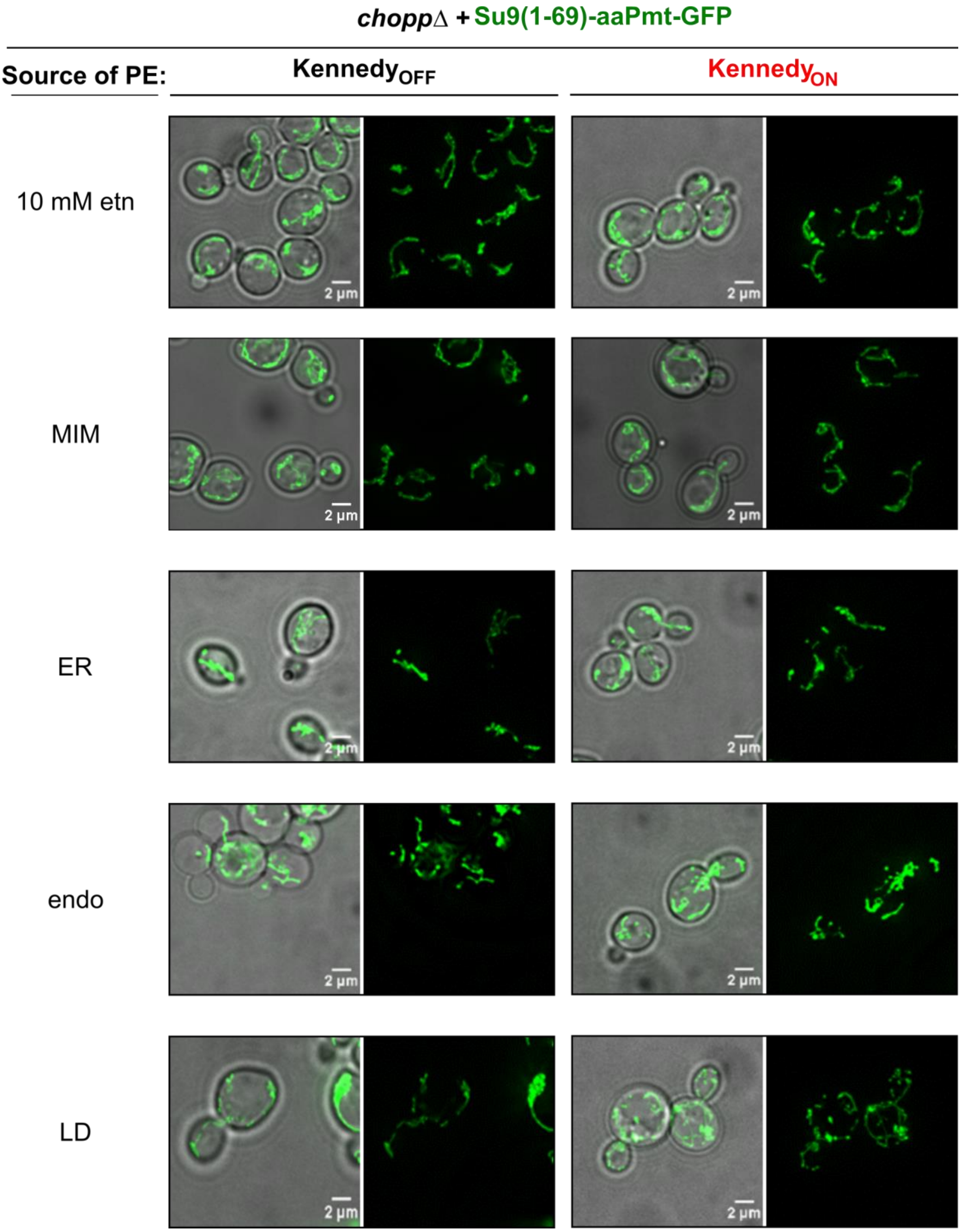


**Fig S4. Localization of Pmt (MM) in different rewired strains.**

Localization of the Su9(ss)-aaPmt-GFP (PC–MM) construct expressed in *choppΔ* cells with PE produced either by the Kennedy pathway (+10 mM ethanolamine) or a 'dark' version of one of the Psd constructs, as indicated. Images shown are maximum intensity projections of several Z-sections.

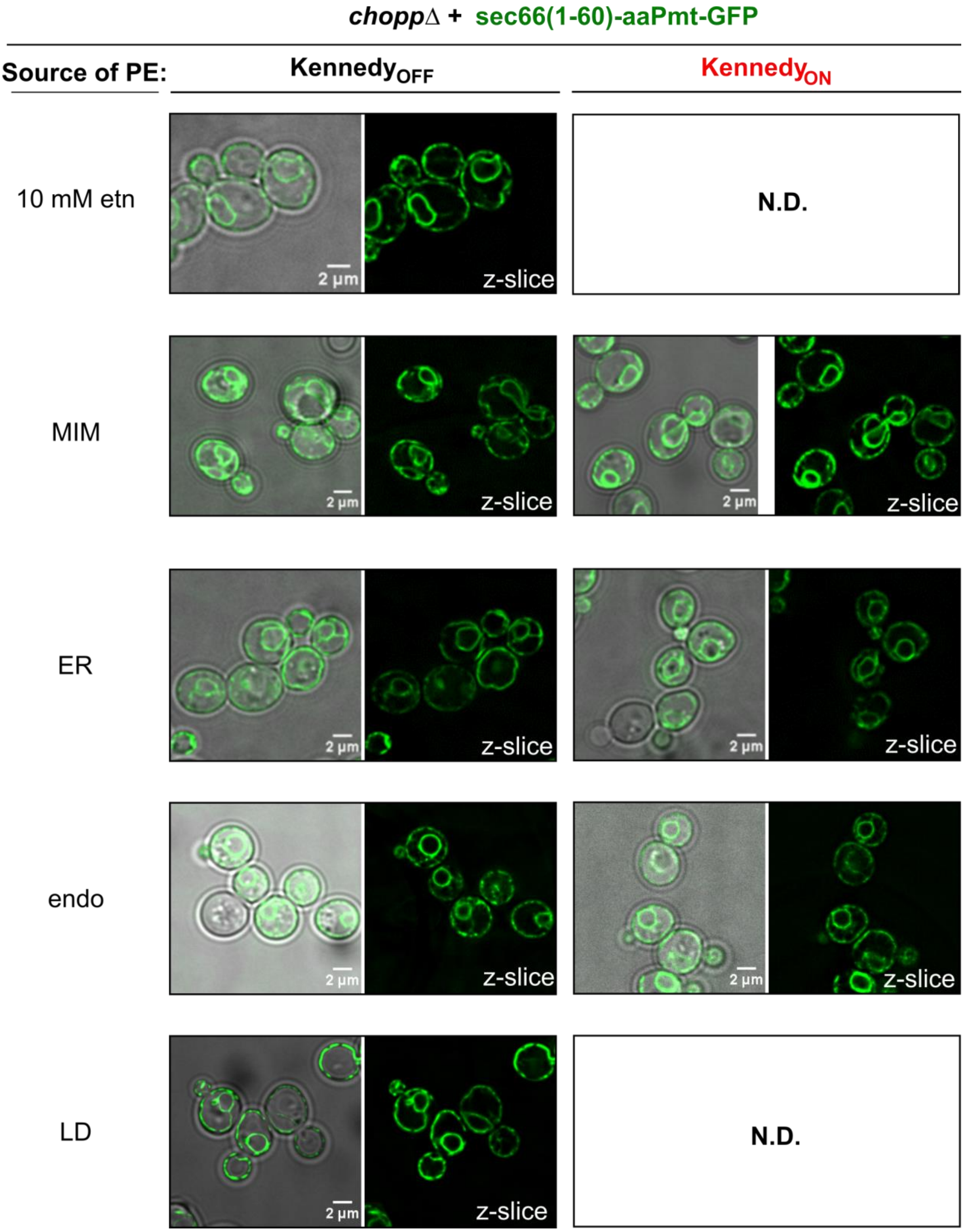


**Fig S5. Localization of Pmt (ER) in different rewired strains.**

Localization of sec66(1-60)-aaPmt-GFP (PC-ER) construct expressed in *choppΔ* cells with PE produced either by the Kennedy pathway (+10 mM ethanolamine) or a 'dark' version of one of the Psd constructs, as indicated. Images shown represent one Z-section.

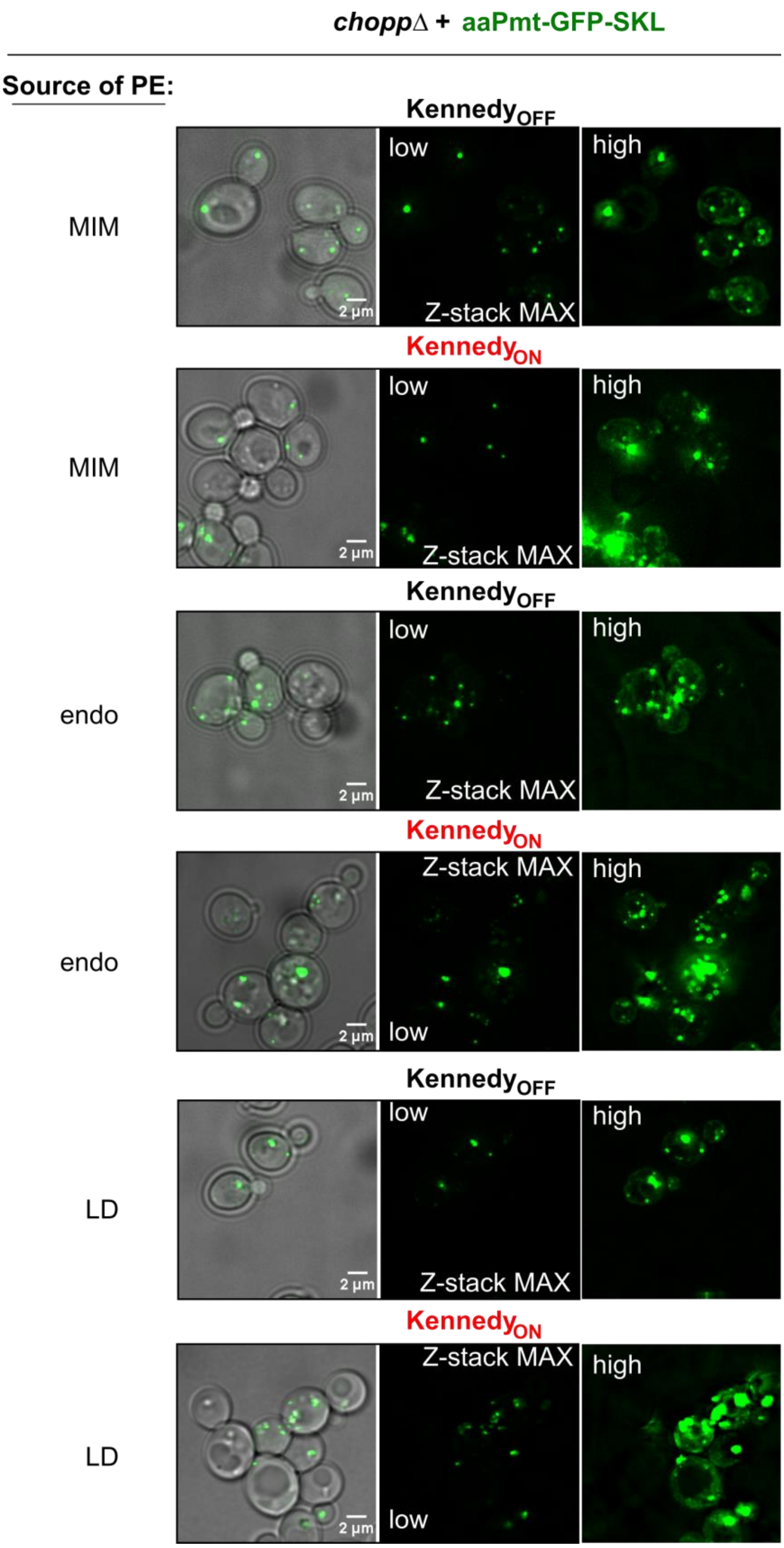


**Fig S6. Localization of Pmt (peroxisome) in different rewired strains.**

Localization of the aaPmt-GFP-SKL (PC-pex) construct expressed in *choppΔ* cells together with a 'dark' version of one of the Psd constructs, as indicated. Images shown are maximum intensity projections of several Z-sections.

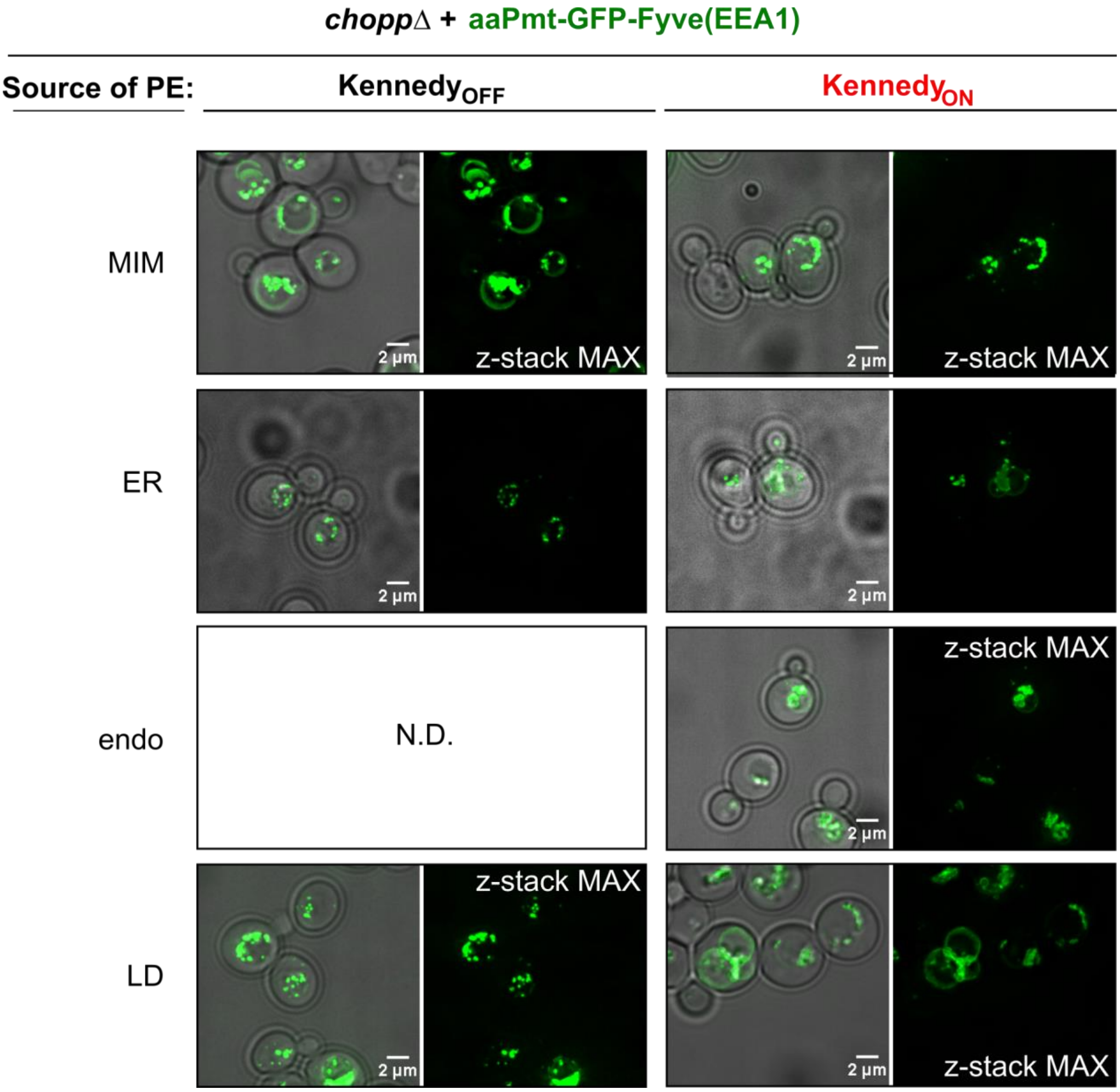


**Fig S7. Localization of Pmt (endosome) in different rewired strains.**

Localization of the aaPmt-GFP-Fyve (PC-endo) construct expressed in *choppΔ* cells together with a 'dark' version of one of the Psd constructs, as indicated. Images shown are maximum intensity projections of several Z-sections.

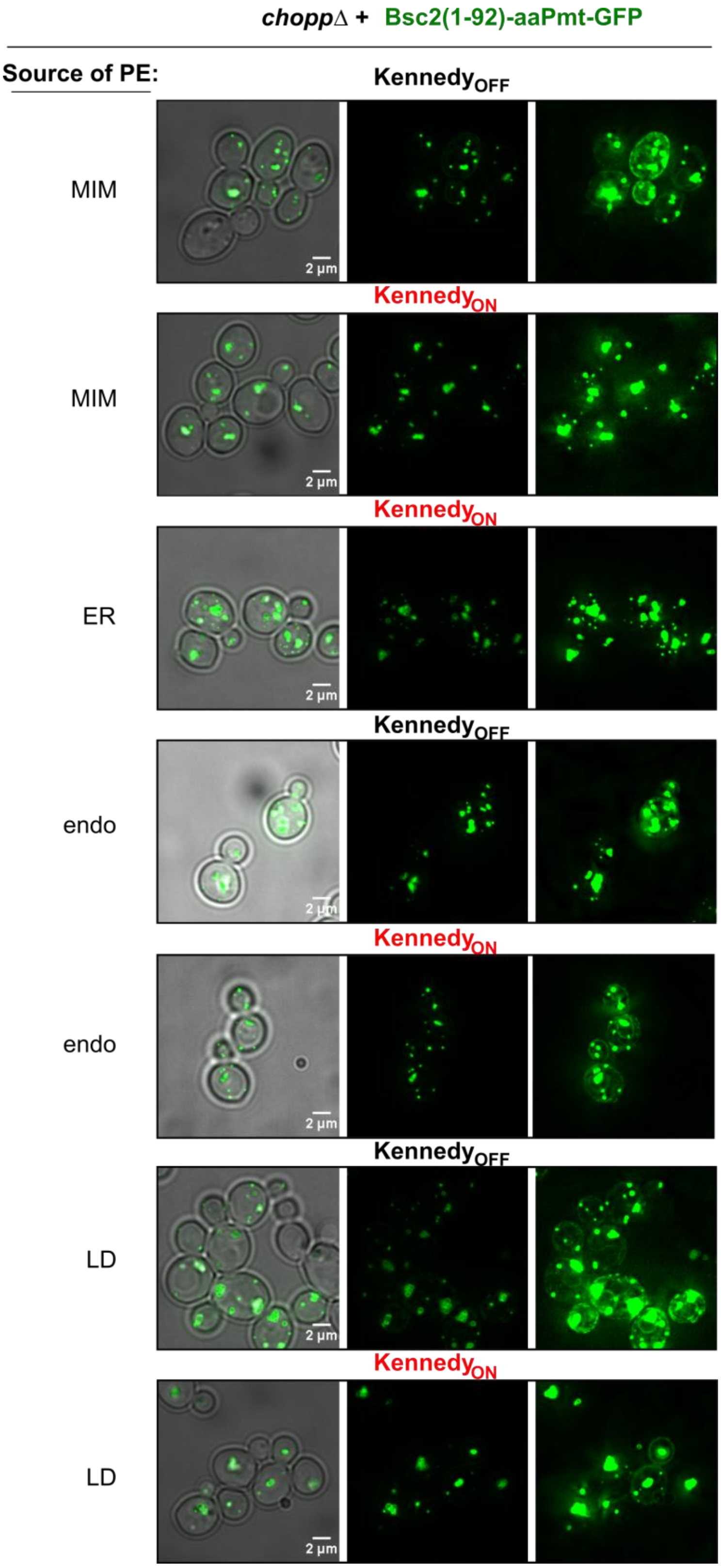


**Fig S8. Localization of Pmt (LD) in different rewired strains.**

Localization of the Bsc2(tm)-aaPmt-GFP (PC-LD) construct expressed in *choppΔ* cells together with a 'dark' version of one of the Psd constructs, as indicated. Images shown are maximum intensity projections of several Z-sections.

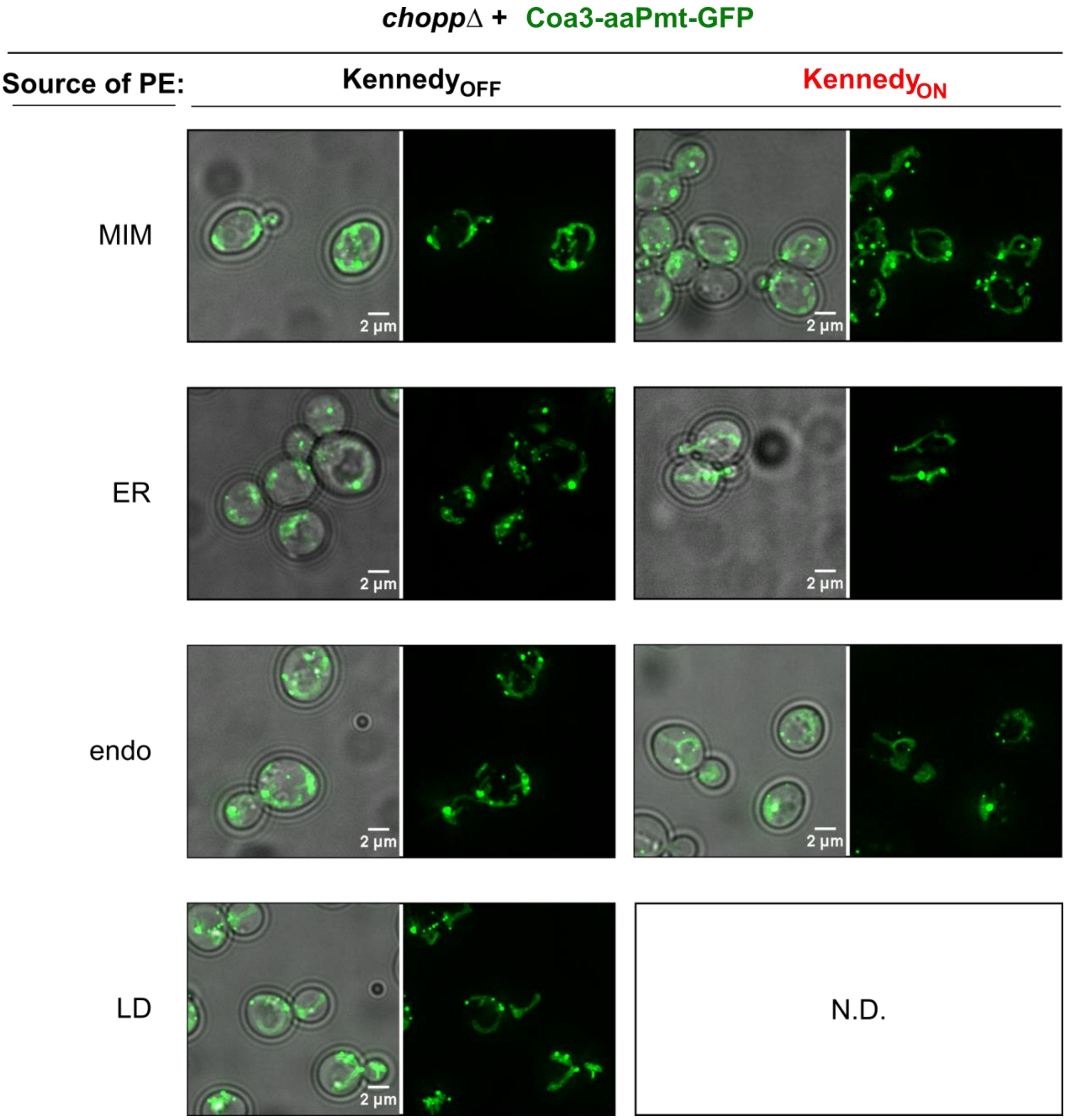


**Fig S9. Localization of Pmt (MIM) in different rewired strains.**

Localization of the Coa3(tm)-aaPmt-GFP (PC-MIM) construct expressed in *choppΔ* cells together with a 'dark' version of one of the Psd constructs, as indicated. Images shown are maximum intensity projections of several Z-sections.

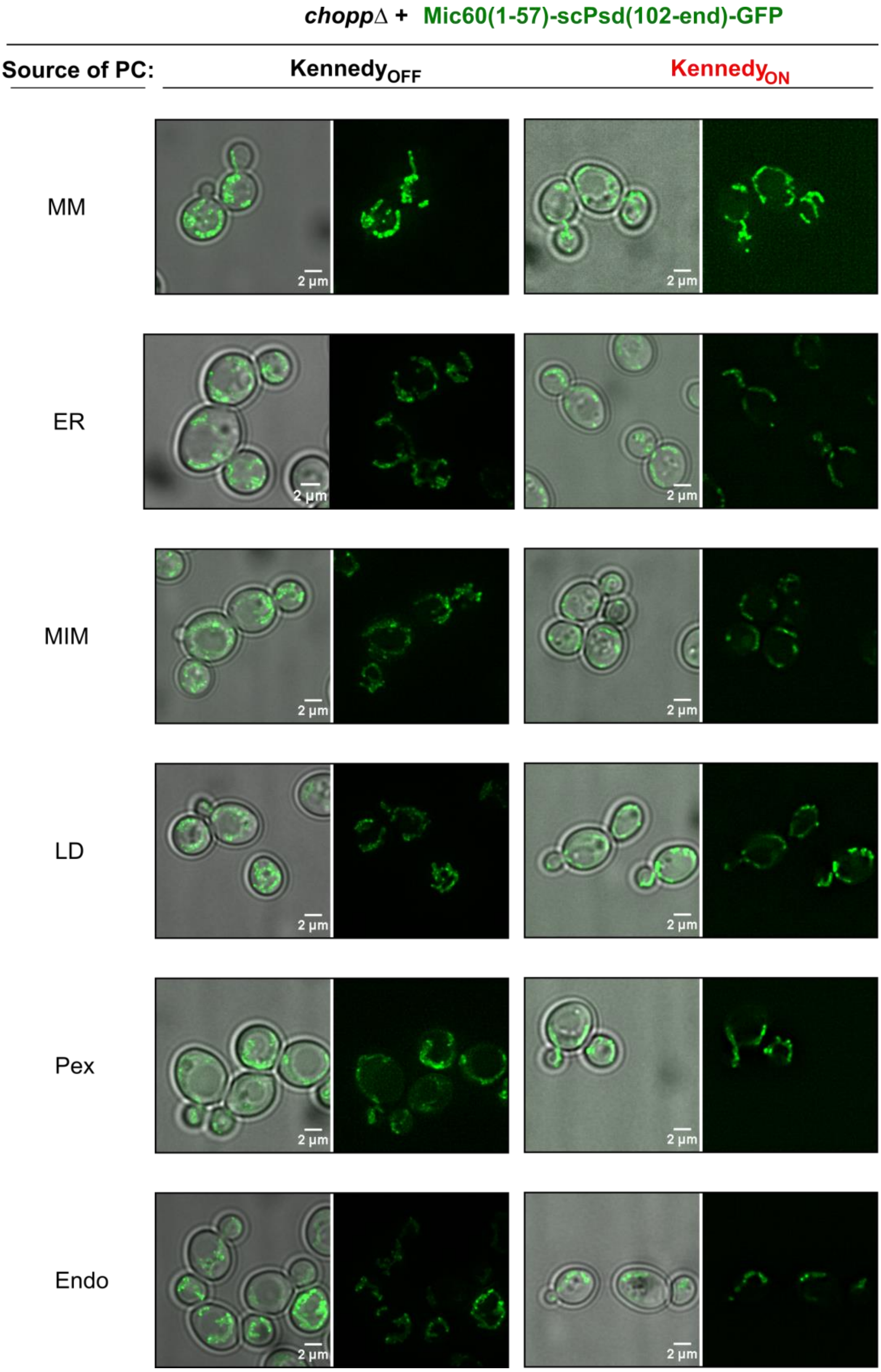


**Fig S10. Localization of Psd (MIM) in different rewired strains.**

Localization of the Mic60(tm)-scPsd-GFP (PE-MIM) construct expressed in *choppΔ* cells together with a 'dark' version of one of the Pmt constructs, as indicated. Images shown are maximum intensity projections of several Z-sections.

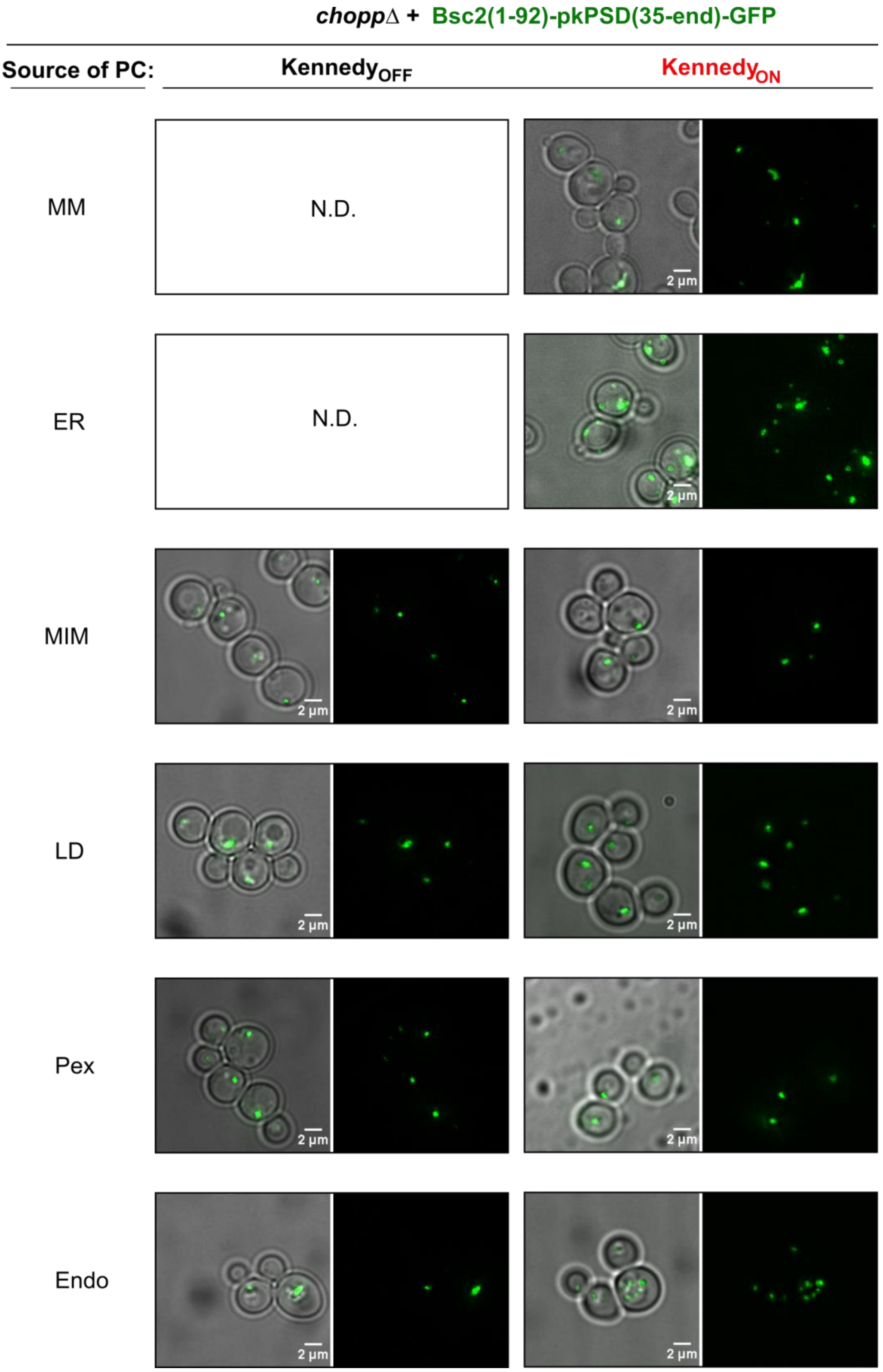


**Fig S11. Localization of Psd (LD) in different rewired strains.**

Localization of the Bsc2(tm)-pkPsd-GFP (PE-LD) construct expressed in *choppΔ* cells together with a 'dark' version of one of the Pmt constructs, as indicated. Images shown are maximum intensity projections of several Z-sections.

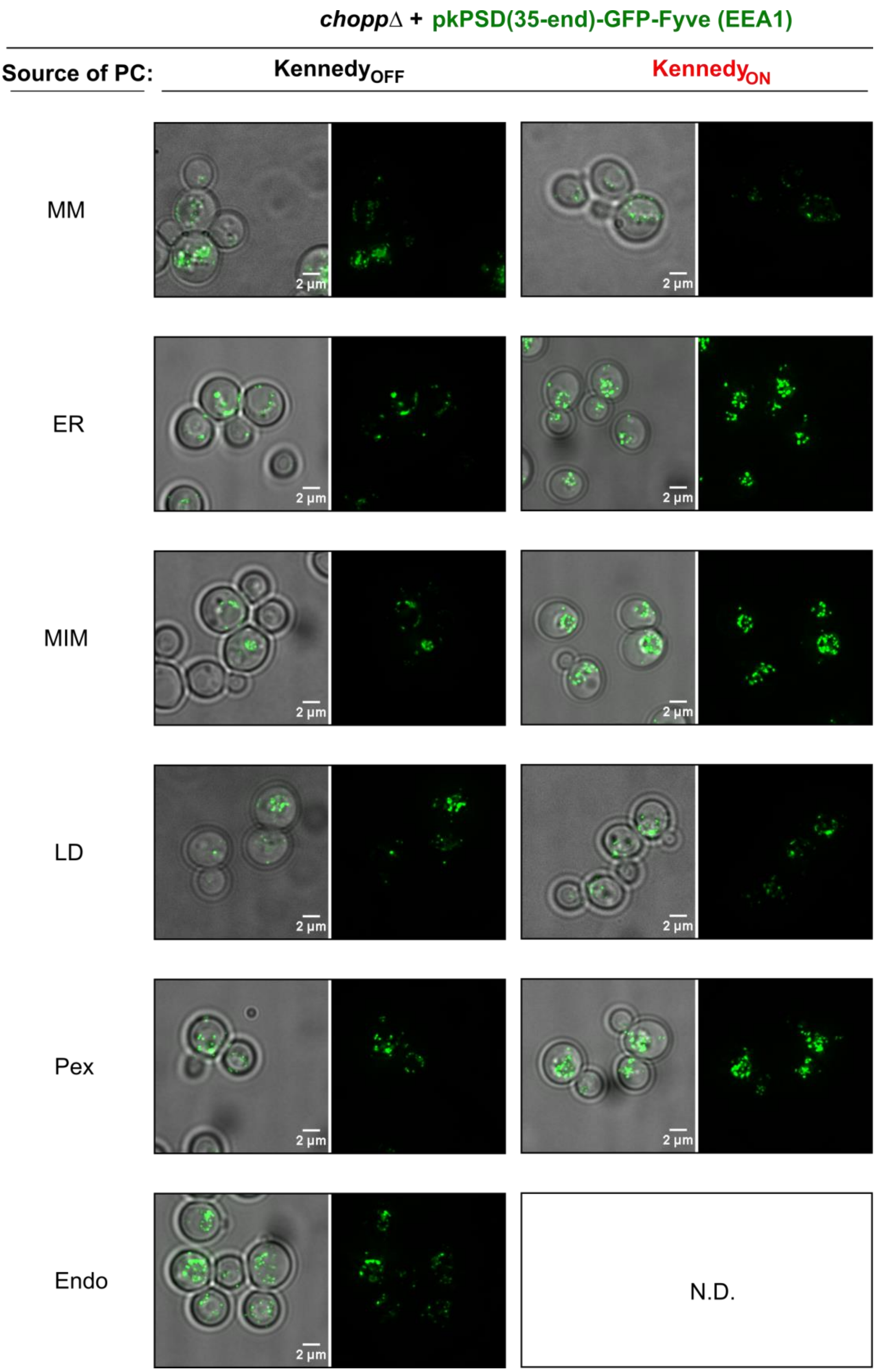


**Fig S12. Localization of Psd (endosome) in different rewired strains.**

Localization of the pkPsd-GFP-Fyve (PE-endo) construct expressed in *choppΔ* cells together with a 'dark' version of one of the Pmt constructs, as indicated. Images shown are maximum intensity projections of several Z-sections.

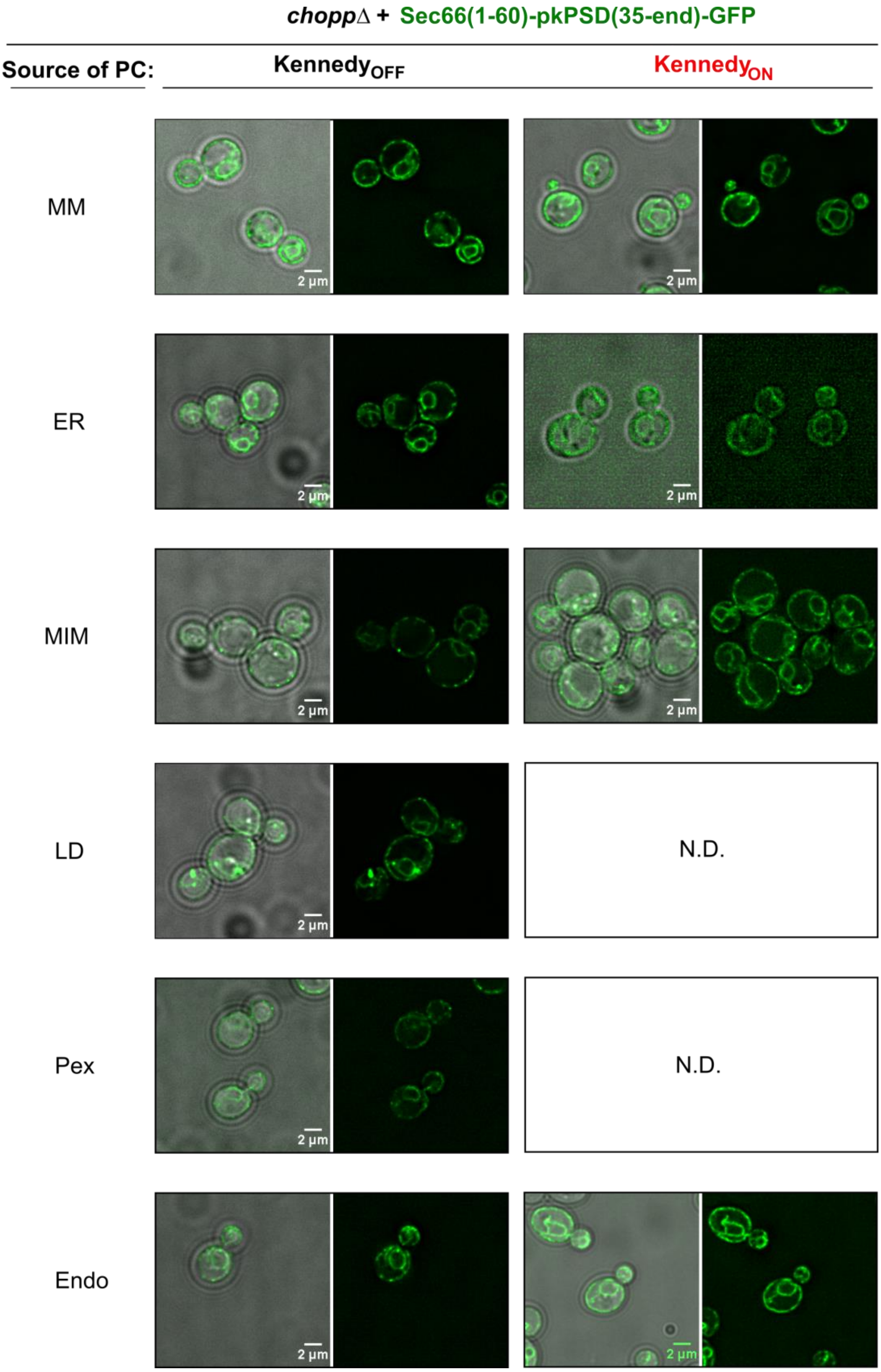


**Fig S13. Localization of Psd (ER) in different rewired strains.**

Localization of the Sec66(tm)-pkPsd-GFP (PE-ER) construct expressed in *choppΔ* cells together with a 'dark' version of one of the Pmt constructs, as indicated. Images shown are maximum intensity projections of several Z-sections.

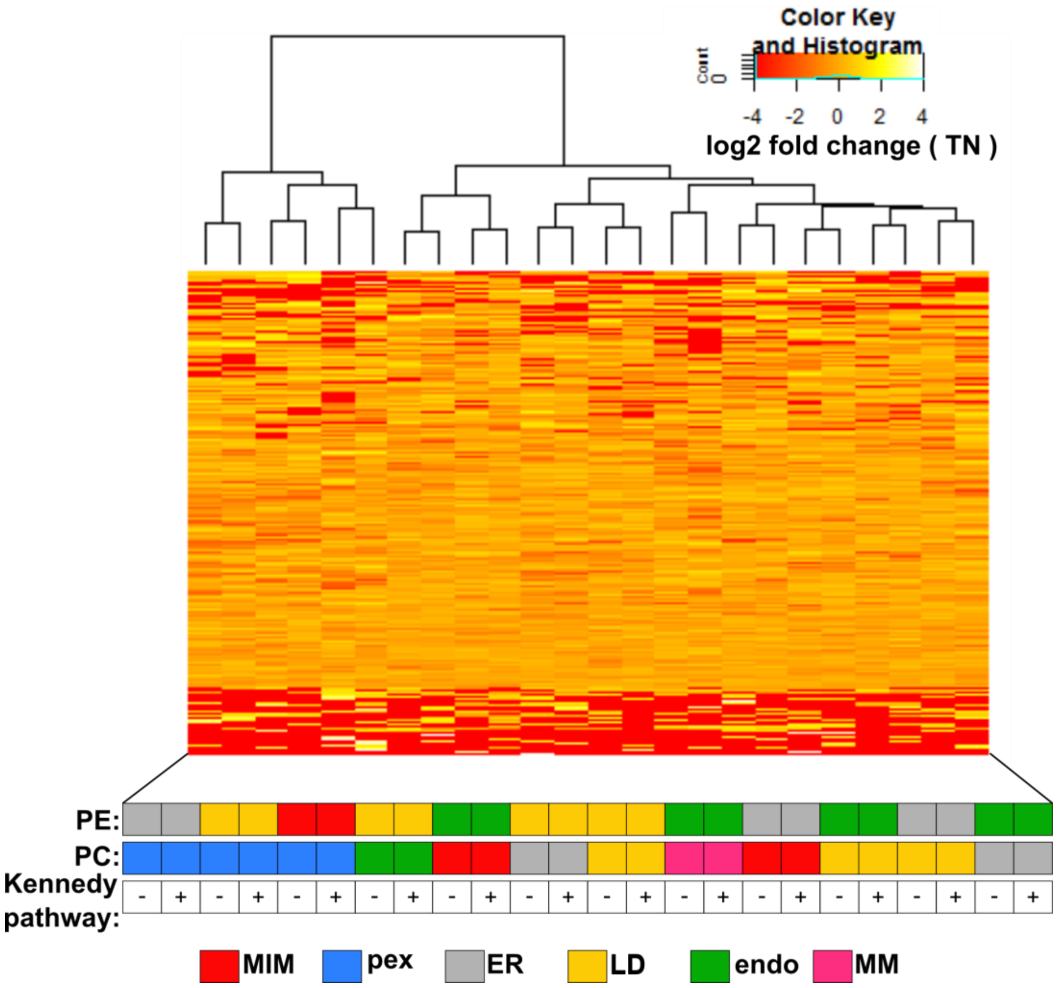


**Fig S14. Hierarchical clustering of gene transposon insertions profiles for all libraries.**

Hierarchical clustering of transposon numbers per gene (rows) computed for all libraries (columns). Color key depicts the log2 fold change of transposon count (TN) for each gene in a library with respect to the mean TN count per gene of all libraries. Dendrogram for library clustering is shown on top. The bottom panel depicts the localization of the Psd (PE) and Pmt (PC) enzymes and the growth conditions for each library. Libraries grown in $Kennedy_{OFF}$ and $_{ON}$ conditions are indicated with - and +, respectively.

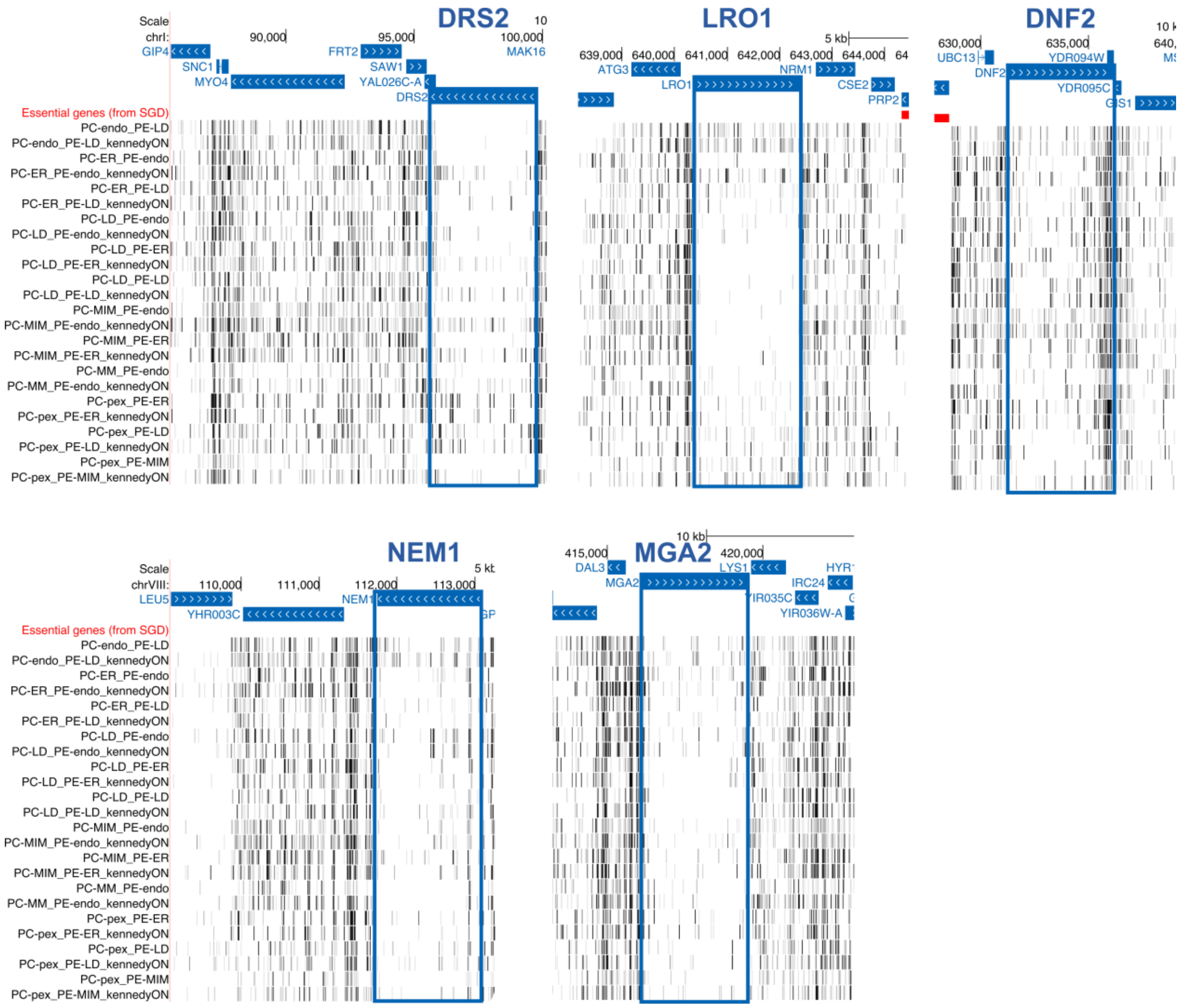


**Fig S15. Lipid metabolism and lipid flippases are required for adaptation in specific rewired conditions.**

Transposon insertion maps of the *DRS2, LRO1, DNF2, NEM1,* and *MGA2* genomic loci generated in the UCSC genome browser for libraries of strains expressing the chimeric Psd and Pmt enzymes to produce PE and PC at the indicated cellular locations in the *choppΔ* background. "KennedyON" refers to libraries that were grown in media supplemented with 10 mM ethanolamine and choline.

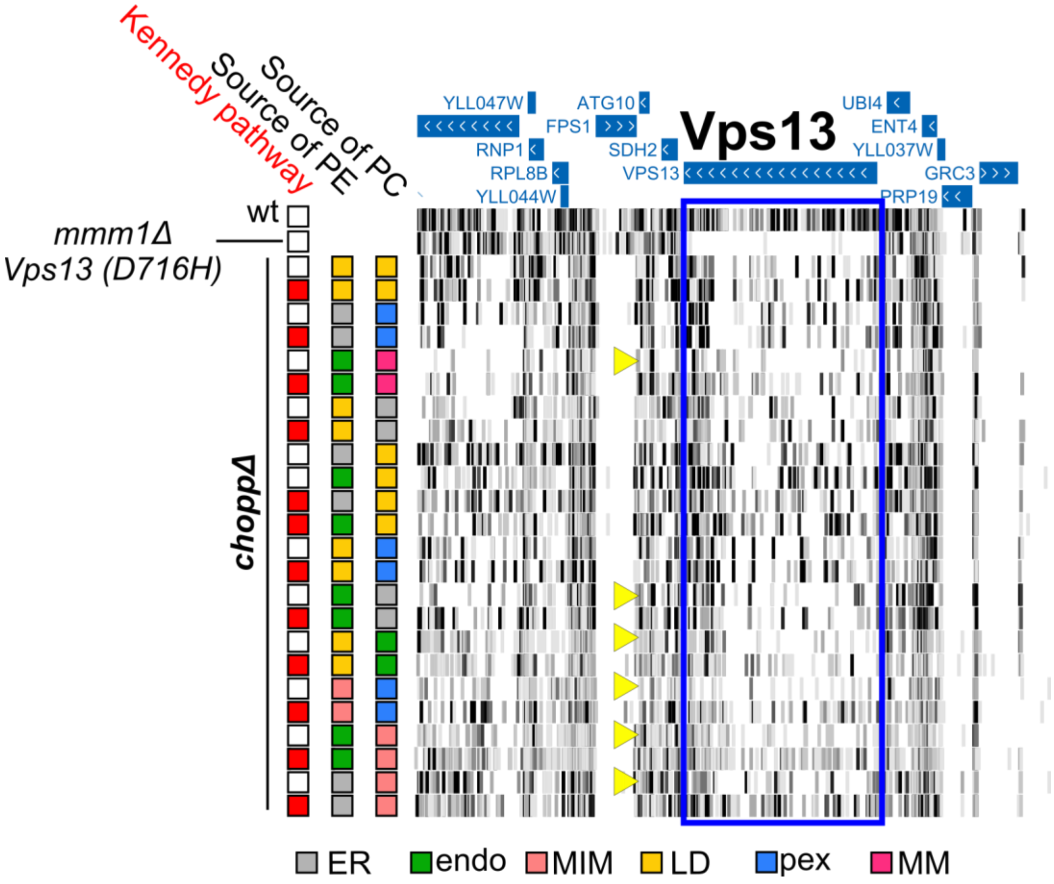


**Fig S16. Vps13 is required in libraries where lipid synthesis occurs at its native locations.**

Transposon insertion maps generated in the UCSC genome browser for the *VPS13* genomic locus for the indicated genotypes. Source of PE and PC corresponds to the Psd and Pmt constructs expressed in the *choppΔ* strain and targeted to the indicated locations. Red box = Kennedy$_{ON}$, white box = Kennedy$_{OFF}$. In addition, a library is displayed where Vps13 is not required (wt) and a library where Vps13 is expected to be essential (*mmm1*Δ (ERMESΔ) v*ps13*D176H (ERMES suppressor allele)). Yellow arrowheads highlight rewired libraries grown in Kennedy$_{OFF}$ conditions where the number of transposons is depleted with respect to the same library grown in Kennedy$_{ON}$ conditions.

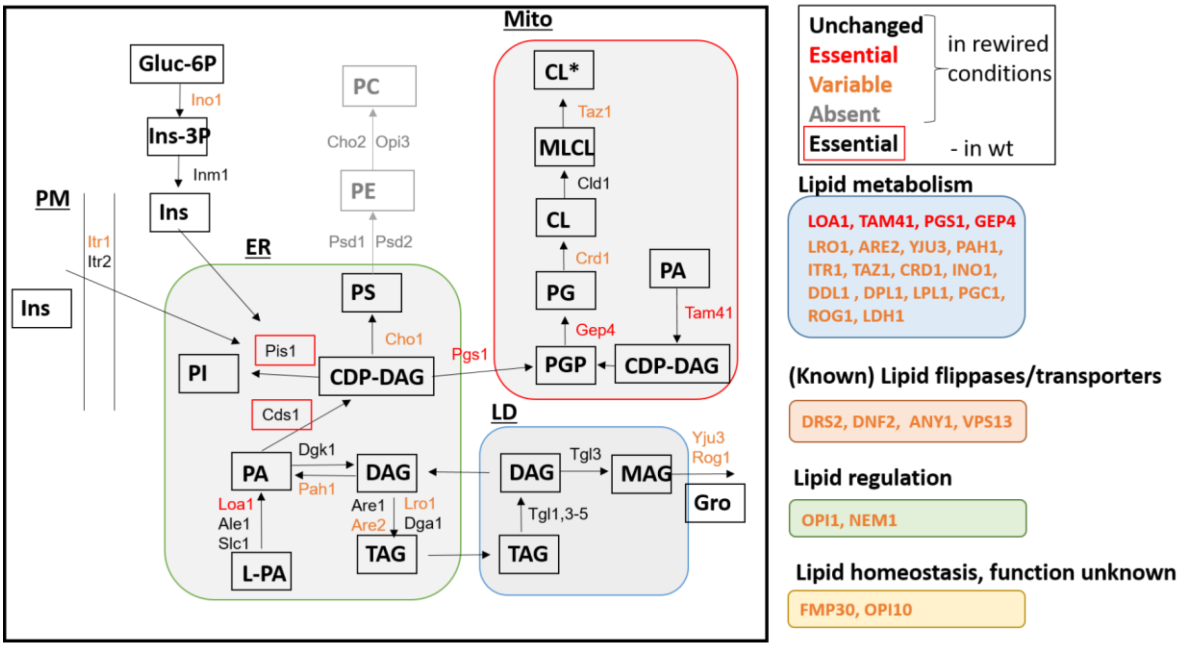


**Fig S17. Requirement for lipid metabolic genes in rewired yeast.**

Schematic illustration of lipid synthesis pathways in yeast. Genes that are essential or absent in all libraries are depicted in red or grey, respectively. Genes variably required in one or more rewired libraries, as assessed by manual inspection or volcano plots, are depicted in orange. Genes that are essential in wild-type conditions are boxed in red.

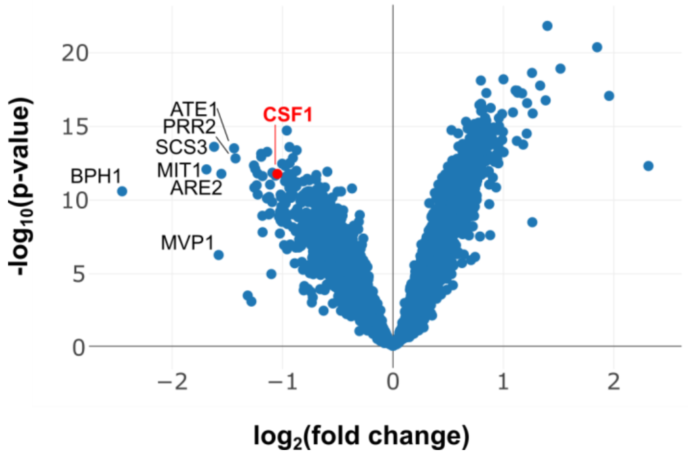


**Fig S18. The requirement for Csf1 is specific to the PE-MIM / PC-pex library grown in Kennedy$_{OFF}$ conditions.**

Volcano plot comparing the number of transposon insertions per gene in the PE-MIM / PC-pex Kennedy$_{OFF}$ condition versus all other libraries. *CSF1* is highlighted in red.

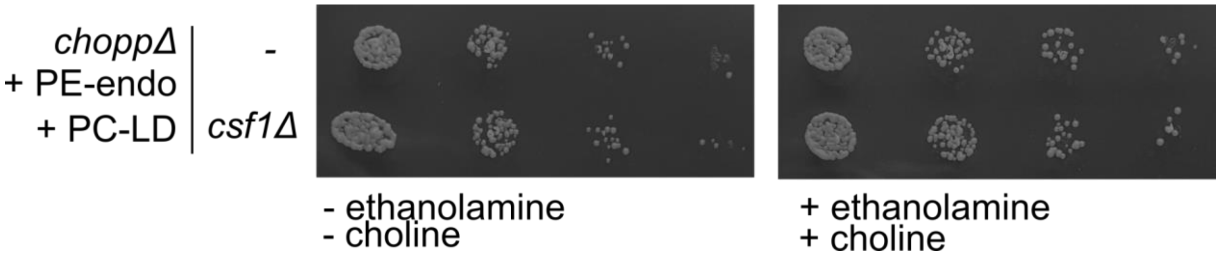


**Fig S19. CSF1 is not required when PE is targeted to endosomes and PC is targeted to LD.**

Five-fold serial dilutions of strains of the indicated genotypes on SD medium without ethanolamine and choline ($Kennedy_{OFF}$), or with ethanolamine and choline ($Kennedy_{ON}$).

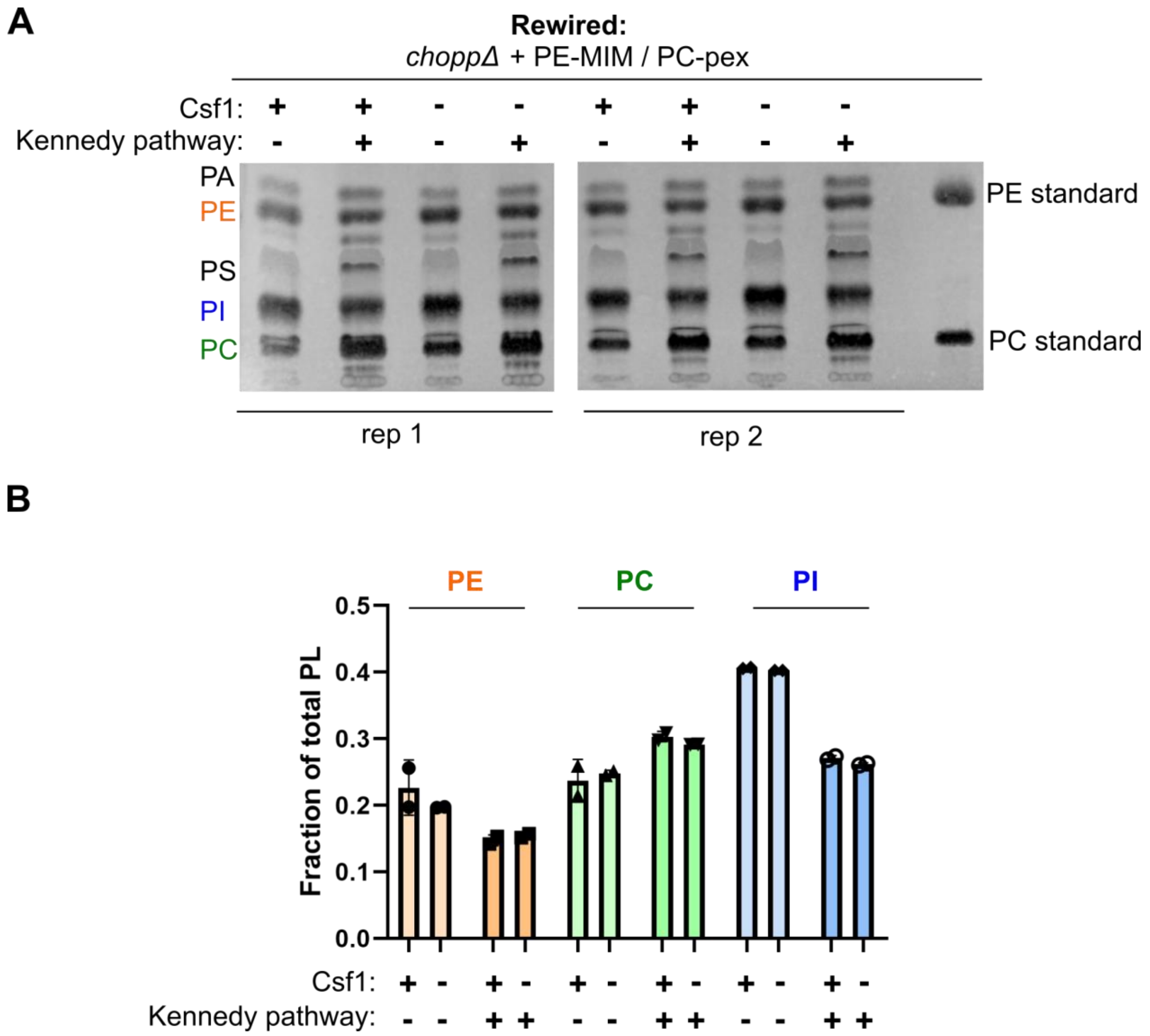


**Fig S20. The absence of Csf1 in rewired conditions does not affect steady state PE or PC production.**

A) Thin layer chromatography (TLC) analysis of the steady-state lipid profiles of the indicated strains grown in $Kennedy_{OFF}$ or $Kennedy_{ON}$ conditions (+ 10 mM ethanolamine and choline).

B) Quantification of the fraction of PE, PC and PI of total phospholipids measured by TLC analysis shown in A. Quantification was done using Fiji/ImageJ software as described in the Materials and Methods.
