## Supplemental Tables S1-S3 for "Rewiring phospholipid biosynthesis reveals robustness in membrane homeostasis and uncovers lipid regulatory players"

Table S1.	**Yeast strains used in this study**

**Strain	Genotype	Reference**

ByK45	BY4741 MATa his3Δ leu2Δ0 met15Δ0 ura3Δ0	Euroscarf

ByK830	w303 MATa	yDF126 - Peter Lab

ByK1148	ByK830 *psd1*Δ *psd2*Δ::KanMX	This study
*cho2Δ*::hphNT1 *opi3Δ*

ByK1149	ByK1148 *csf1Δ*::His3MX6	This study

ByK1418	ByK45 CSF1::GFP- KANMX	This study

ByK1419	ByK45 CSF1::GFP- KANMX PEX10::mCHERRY-NAT	This study

ByK1420	ByK45 *csf1(737aa-end∆)*	This study

Table S2.	**Plasmids used in this study**

**Plasmids	Genotype	Primers used	Reference**

*Plasmids for rewiring:*

pBK586	pRS415-TEFpr- Bsc2(1-92)-pkPSD(35-end)-GFP	This study

pBK588	pRS415-TEFpr- pkPSD(35-end)-GFP-fyve (EEA1)	This study

pBK590	pRS415-TEFpr- Sec66(1-60)-pkPSD(35-end)-GFP	This study

pBK785	pRS415-PSDpr-Mic60(1-57)-scPSD(102-end)-GFP	This study

pBK556	pRS415-TEFpr-Su9(1-69)-aaPmt-GFP	This study

pBK557	pRS415-TEFpr-aaPmt-GFP(skl)	This study

pBK558	pRS415-TEFpr-aaPmt-GFP-fyve (EEA1)	This study

pBK559	pRS415-TEFpr-Coa3(1-49)-aaPmt-GFP	This study

pBK561	pRS415-TEFpr- Bsc2(1-92)-aaPmt-GFP	This study

pBK562	pRS415-TEFpr-Sec66(1-60)-aaPmt-GFP	This study

pBK786	pBK586 *Leu2Δ*::HIS3	This study

pBK787	pBK588	*Leu2Δ*::HIS3	This study

pBK788	pBK590	*Leu2Δ*::HIS3	This study

pBK789	pBK590	*Leu2Δ*::HIS3	This study

pBK790	pBK586	*Leu2Δ*::URA3	This study

pBK792	pBK785 *Leu2Δ*::URA3	This study

pBK793	pBK786 GFP(G65T G67A)	This study

pBK794	pBK787 GFP(G65T G67A)	This study

pBK795	pBK788 GFP(G65T G67A)	This study

pBK796	pBK789 GFP(G65T G67A)	This study

pBK797	pBK556 GFP(G65T G67A)	This study

pBK798	pBK557 GFP(G65T G67A)	This study

pBK799	pBK558 GFP(G65T G67A)	This study

pBK800	pBK559 GFP(G65T G67A)	This study

Table S2. continued	**Plasmids used in this study**

**Plasmids	Genotype	Primers used	Reference**

pBK801	pBK561 GFP(G65T G67A)	This study

pBK802	pBK562 GFP(G65T G67A)	This study

*Other plasmids:*

pBK506	pRS415-TEFpr	Mumberg D, Müller R, Funk M.
Gene. 1995

pBK196	pRS424GFP-FYVE(EEA1)	Burd CG, Emr SD. Mol Cell. 1998
(Addgene plasmid # 36096)

pBK626	pRS316-*trp1*::miniDs GAL1pr-TPase	This study

pBK804	pRS413-ERG6pr-ERG6-mCHERRY	This study

pBK417	pRS413-TEFpr-mCHERRY-Ubc6(TA)	John Peter et al. 2017

pBK64	pVTU100-mtBFP	Westermann and Neupert 2000

Table S3.	**Primers used in this study**

| Name | Sequence (5' 3') |
| --- | --- |
| **Primers used for gene deletions** | |
| CSF1 C' tag pYM F | CAAAAGCTTGTTTATCTTGCAGAAAAGCAGTATGTCAAGATACTAGATGATACGCATcgtacgctgcaggtcgac |
| CSF1 C' tag pYM R | ACATAAACCAGAAATATGGTATCAAGGACTTTTGAATATAATTAGGAACGAGATTAatcgatgaattcgagctcg |
| CSF1 KO pYM F | CATTAAAGCCACCTGACTCAAGTCTTCTATTGACGGTAATAAGTTAGCAAGCATGcgtacgctgcaggtcgac |
| CSF1_C'tag_797aa_f | ACGTCAGAAGAGTACACAGGTGTCCTTGGCGCTAGGGAAGTCGGAGATGTCACcgtacgctgcaggtcgac |
| CSF1_KO_797aa_f | ACGTCAGAAGAGTACACAGGTGTCCTTGGCGCTAGGGAAGTCGGAGATGTCACTAAcgtacgctgcaggtcgac |
| Psd2_pringle_F | GATGCTGTATCAATTGGTAAAGAATCCTCGATTTTCAGGAGCATCCAACGcgtacgctgcaggtcgac |
| Psd2_pringle_R | CTTGTTTGTACACGCTATAGTCTATAATAAAGTCTGAGGGAGATTGTTCATGatcgatgaattcgagctcg |
| Cho2_S1_knop | CTGAATATTTCGAGTGATTTTCTTAGTGACAAAGCTTTTTCTTCATCTGTAGATGcgtacgctgcaggtcgac |
| Cho2_S2_knop | TAACTTGAATCCTAGTACTTTTTAAATATATATACTCAAAAAAAAAAAACTCAatcgatgaattcgagctcg |
| **Primers used for CRISPR-Cas9-mediated gene deletions** | |
| Psd1_gRNA_IntRev | ctagctctaaaacACCCCTTGATGTCTAAGAGTgatcatttatctttcactgcggagaag |
| Psd1_gRNA_Intfwd | ACTCTTAGACATCAAGGGGTgttttagagctagaaatagcaagttaaaataaggctagtccg |
| Psd1_link_IntRev | gtccgccggcgttggacgagcgGCTGGCTTTGCTTTTCCTTCTTCTTC |
| Psd1_link_IntFwd | ctcgtccaacgccggcggacctAAGCAATCATATGTAAAGTTAGCATTTATTTTGCTG |
| Psd1_-484up_f | GACTGGTACACCTGCAGGTGTAG |
| Psd1_-500dwSTOP_r | CACCTCTTCGCAACTGGTTGAAAG |
| Opi3_gRNA_fwd | gtttcggcgttcgaAACTTCTCCGCAGTGAAAGATAAATGATCcccgtgatcagagaacacgtGTTTTAGAGCTAGAAATAGCAAGTTAAAATAAGGCTAGTCCGTTATCAACTTGAAAAAG |
| Opi3_gRNA_rev | CTTTTTCAAGTTGATAACGGACTAGCCTTATTTTAACTTGCTATTTCTAGCTCTAAAACacgtgttctctgatcacgggGATCATTTATCTTTCACTGCGGAGAAGTTtcgaacgccgaaac |
| opi3_link_intRev | gtccgccggcgttggacgagcgTGCTTGACTTGCGCTATTCTTGTTG |
| opi3_link_intfwd | ctcgtccaacgccggcggacctCCTATGCTATTACCGTTTCTATATAGCTCC |
| opi3_407upATG_f | GGTGGCTAGTCCGTCTTCAAATTC |
| opi3_402dwSTP_r | CTGGTGACAATGGCACCGTTC |
| **Primers used for colony PCR** | |
| Psd2 up | ACGCATGTGCTACTTCAAGG |
| Psd2 down | AAGGCCGAGAAGTACCTTTG |
| Psd1_-584up_f | CGG ACA GTT GAG ACA AGA TGG TGG |
| Psd1_-500dwSTOP_r | CACCTCTTCGCAACTGGTTGAAAG |
| CSF1_152upATG_f | CCACTATAAAGCTGTTGGCACGG |
| CSF1_2537dnATG_r | GAGCACCATCCCAAACCGAAAATG |
| CSF1_2137dnATG_f | CCCCTTGGAATACATTGAACGAATTC |
| KanMX_twd_5prime | CATGTTGGAATTTAATCGCGGCCTC |
| natNT2_+26_rev | CTGGTGCGGTACCGGTAAG |
| Nat -249Rev | GGATGTATGGGCTAAATGTAC |
| **Primers used for plasmid construction** | |
| URA3_gr_fwd | GTGCGGTATTTCACACCGCATATCGACGGTCGAGGgagtgcaccataccacagcttttc |
| URA3_gr_rev | GTTTATGTACAAATATCATAAAAAAAGAGAATCTTTttagttttgctggccgcatcttc |
| HIS1_gr_fwd | GTGCGGTATTTCACACCGCATATCGACGGTCGAGGatgcgtacgctgcaggtcgac |
| HIS1_gr_rev | GTTTATGTACAAATATCATAAAAAAAGAGAATCTTTtaaatcgatgaattcgagctcg |
| G229A_G230C_G236C_f | gtatctcgcaaaacattgaacagcataagtgaaagtagtgactaaggttggc |
| G229A_G230C_G236C_r | gccaaccttagtcactactttcacttatgctgttcaatgttttgcgagatac |
| Psd1pr_gr_f | aaccctcactaaagggaacaaaagctggagctcTGAGACAAGATGGTGGTACTAACC |
| Psd1pr_Mic60_gr_r | aatttttcgtgaggcagtagttcttagcatcattctagaGCTGGCTTTGCTTTTCCTTCTTC |
| Mic60_f | atgatgctaagaactactgcctcac |
| mic60_yPSD_r | GATTTTTCTTGTCCTTCTCCCTTTTTTGCCCTCTGTAGCATCCTCCGAATATATGATACCTCCAGCGTAG |
| yPSD_AA102_f | GAGGATGCTACAGAGGGCAA |
| yPSD_NOSTOP_R | TTTTAAATCATTCTTTCCAATTATGCC |
| yPSD_HIND3-GFP-F | GGGACAGAAATTAGGCATAATTGGAAAGAATGATTTAAAAaagcttggagcaggtgctggtgctgg |
| pRS415_GFP_r | CTAATTACATGACTCGAGGTCGACGGTATCGATAAGCTTttatttgtacaattcatccataccatgggt |
| Aapmt_f | actagtATGAGTACCTCCAGACAAAGAGAAGATATG |
| Aapmt_Bsc2_r | CATATCTTCTCTTTGTCTGGAGGTACTCATactagtGATCGCGTCCAGTATGATAATAGGC |
| pRS415_Aapmt_r | CTAATTACATGACTCGAGGTCGACGGTATCGATAAGCTTtcagacaggcaagttttccaaagttac |
| Tef_Sec66_f | AAGCATAGCAATCTAATCTAAGTTTTCTAGAatgtccgaatttaatgaaacaaaattctcc |
| Aapmt_Sec66_r | CATATCTTCTCTTTGTCTGGAGGTACTCATactagtTGGTTGCTCACTAATTTTTTTGGCC |
| Tef_Aapmt_f | aagcatagcaatctaatctaagttttctagaATGAGTACCTCCAGACAAAGAGAAGATATG |
| p415_SKL_Aapmt_r | TTACATGACTCGAGGTCGACGGTATCGATaagcttTTAtaatttagaGACAGGCAAGTTTTCCAAAGTTACTAAG |
| Aapmt_rev | GACAGGCAAGTTTTCCAAAGTTACTAAGG |
| Aapmt-GFP_f | CCTTAGTAACTTTGGAAAACTTGCCTGTCactagtGGAGCAGGTGCTGGTGCTG |
| pYM25-GFP-r | tttgtacaattcatccataccatgggt |
| GFP_FYVE_f | acccatggtatggatgaattgtacaaaTGGCAATCTAGTCAACGGAGAGTTAG |
| FYVE_pRS415_r | ctaattacatgactcgaggtcgacggtatcgatTTATCCTTGCAAGTCATTGAAACATGCATC |
| Tef_Su9_f | AAGCATAGCAATCTAATCTAAGTTTTCTAGAatggcctccactcgtgtcctc |
| Aapmt_Su9_r | CATATCTTCTCTTTGTCTGGAGGTACTCATactagtGGAAGAGTAGGCGCGCTTCTG |
| Aapmt_Vac8_r | CATATCTTCTCTTTGTCTGGAGGTACTCATactagtATGTAAAAATTGTAAAATCTGTTGAGTAATATTATAC |
| GFP_AaPmt_r | AGCACCAGCACCAGCACCTGCTCCactagtGACAGGCAAGTTTTCCAAAGTTACTAAGG |
| **Primers used for library preparation** | |
| P5_MiniDs | AATGATACGGCGACCACCGAGATCTACtccgtcccgcaagttaaata |
| P7_indexed_N701 | CAA GCA GAA GAC GGC ATA CGA GAT TCG CCT TAA CGA AAA CGA ACG GGA TAA A |
| P7_indexed_N702 | CAA GCA GAA GAC GGC ATA CGA GAT CTA GTA CGA CGA AAA CGA ACG GGA TAA A |
| P7_indexed_N703 | CAA GCA GAA GAC GGC ATA CGA GAT TTC TGC CTA CGA AAA CGA ACG GGA TAA A |
| P7_indexed_N704 | CAA GCA GAA GAC GGC ATA CGA GAT GCT CAG GAA CGA AAA CGA ACG GGA TAA A |
| P7_indexed_N705 | CAA GCA GAA GAC GGC ATA CGA GAT AGG AGT CCA CGA AAA CGA ACG GGA TAA A |
| P7_indexed_N706 | CAA GCA GAA GAC GGC ATA CGA GAT CAT GCC TAA CGA AAA CGA ACG GGA TAA A |
| P7_indexed_N707 | CAA GCA GAA GAC GGC ATA CGA GAT GTA GAG AGA CGA AAA CGA ACG GGA TAA A |
| P7_indexed_N710 | CAA GCA GAA GAC GGC ATA CGA GAT CAG CCT CGA CGA AAA CGA ACG GGA TAA A |
| P7_indexed_N711 | CAA GCA GAA GAC GGC ATA CGA GAT TGC CTC TTA CGA AAA CGA ACG GGA TAA A |
| P7_indexed_N712 | CAA GCA GAA GAC GGC ATA CGA GAT TCC TCT ACA CGA AAA CGA ACG GGA TAA A |
| P7_indexed_N714 | CAA GCA GAA GAC GGC ATA CGA GAT TCA TGA GCA CGA AAA CGA ACG GGA TAA A |
| P7_indexed_N715 | CAA GCA GAA GAC GGC ATA CGA GAT CCT GAG ATA CGA AAA CGA ACG GGA TAA A |
| P7_indexed_N716 | CAA GCA GAA GAC GGC ATA CGA GAT TAG CGA GTA CGA AAA CGA ACG GGA TAA A |
| P7_indexed_N718 | CAA GCA GAA GAC GGC ATA CGA GAT GTA GCT CCA CGA AAA CGA ACG GGA TAA A |
| P7_indexed_N719 | CAA GCA GAA GAC GGC ATA CGA GAT TAC TAC GCA CGA AAA CGA ACG GGA TAA A |
| P7_indexed_N720 | CAA GCA GAA GAC GGC ATA CGA GAT AGG CTC CGA CGA AAA CGA ACG GGA TAA A |
| P7_indexed_N721 | CAA GCA GAA GAC GGC ATA CGA GAT GCA GCG TAA CGA AAA CGA ACG GGA TAA A |
| P7_indexed_N722 | CAA GCA GAA GAC GGC ATA CGA GAT CTG CGC ATA CGA AAA CGA ACG GGA TAA A |
| P7_indexed_N723 | CAA GCA GAA GAC GGC ATA CGA GAT GAG CGC TAA CGA AAA CGA ACG GGA TAA A |
| P7_indexed_N724 | CAA GCA GAA GAC GGC ATA CGA GAT CGC TCA GTA CGA AAA CGA ACG GGA TAA A |
| P7_indexed_N726 | CAA GCA GAA GAC GGC ATA CGA GAT GTC TTA GGA CGA AAA CGA ACG GGA TAA A |
| P7_indexed_N727 | CAA GCA GAA GAC GGC ATA CGA GAT ACT GAT CGA CGA AAA CGA ACG GGA TAA A |
| P7_indexed_N728 | CAA GCA GAA GAC GGC ATA CGA GAT TAG CTG CAA CGA AAA CGA ACG GGA TAA A |
| P7_indexed_N729 | CAA GCA GAA GAC GGC ATA CGA GAT GAC GTC GAA CGA AAA CGA ACG GGA TAA A |
| **Primers used for sequencing** | |
| 688_minidsSEQ1210 | tttaccgaccgttaccgaccgttttcatcccta |
| Custom_index1 | GGT TTT CGA TTA CCG TAT TTA TCC CGT TCG TTT TCG T |
