## Supplemental Data 2 for "Rewiring phospholipid biosynthesis reveals robustness in membrane homeostasis and uncovers lipid regulatory players": htmlwidgets.js

```
(function() {
  // If window.HTMLWidgets is already defined, then use it; otherwise create a
  // new object. This allows preceding code to set options that affect the
  // initialization process (though none currently exist).
  window.HTMLWidgets = window.HTMLWidgets || {};

  // See if we're running in a viewer pane. If not, we're in a web browser.
  var viewerMode = window.HTMLWidgets.viewerMode =
      /\bviewer_pane=1\b/.test(window.location);

  // See if we're running in Shiny mode. If not, it's a static document.
  // Note that static widgets can appear in both Shiny and static modes, but
  // obviously, Shiny widgets can only appear in Shiny apps/documents.
  var shinyMode = window.HTMLWidgets.shinyMode =
      typeof(window.Shiny) !== "undefined" && !!window.Shiny.outputBindings;

  // We can't count on jQuery being available, so we implement our own
  // version if necessary.
  function querySelectorAll(scope, selector) {
    if (typeof(jQuery) !== "undefined" && scope instanceof jQuery) {
      return scope.find(selector);
    }
    if (scope.querySelectorAll) {
      return scope.querySelectorAll(selector);
    }
  }

  function asArray(value) {
    if (value === null)
      return [];
    if ($.isArray(value))
      return value;
    return [value];
  }

  // Implement jQuery's extend
  function extend(target /*, ... */) {
    if (arguments.length == 1) {
      return target;
    }
    for (var i = 1; i < arguments.length; i++) {
      var source = arguments[i];
      for (var prop in source) {
        if (source.hasOwnProperty(prop)) {
          target[prop] = source[prop];
        }
      }
    }
    return target;
  }

  // IE8 doesn't support Array.forEach.
  function forEach(values, callback, thisArg) {
    if (values.forEach) {
      values.forEach(callback, thisArg);
    } else {
      for (var i = 0; i < values.length; i++) {
        callback.call(thisArg, values[i], i, values);
      }
    }
  }

  // Replaces the specified method with the return value of funcSource.
  //
  // Note that funcSource should not BE the new method, it should be a function
  // that RETURNS the new method. funcSource receives a single argument that is
  // the overridden method, it can be called from the new method. The overridden
  // method can be called like a regular function, it has the target permanently
  // bound to it so "this" will work correctly.
  function overrideMethod(target, methodName, funcSource) {
    var superFunc = target[methodName] || function() {};
    var superFuncBound = function() {
      return superFunc.apply(target, arguments);
    };
    target[methodName] = funcSource(superFuncBound);
  }

  // Add a method to delegator that, when invoked, calls
  // delegatee.methodName. If there is no such method on
  // the delegatee, but there was one on delegator before
  // delegateMethod was called, then the original version
  // is invoked instead.
  // For example:
  //
  // var a = {
  //   method1: function() { console.log('a1'); }
  //   method2: function() { console.log('a2'); }
  // };
  // var b = {
  //   method1: function() { console.log('b1'); }
  // };
  // delegateMethod(a, b, "method1");
  // delegateMethod(a, b, "method2");
  // a.method1();
  // a.method2();
  //
  // The output would be "b1", "a2".
  function delegateMethod(delegator, delegatee, methodName) {
    var inherited = delegator[methodName];
    delegator[methodName] = function() {
      var target = delegatee;
      var method = delegatee[methodName];

      // The method doesn't exist on the delegatee. Instead,
      // call the method on the delegator, if it exists.
      if (!method) {
        target = delegator;
        method = inherited;
      }

      if (method) {
        return method.apply(target, arguments);
      }
    };
  }

  // Implement a vague facsimilie of jQuery's data method
  function elementData(el, name, value) {
    if (arguments.length == 2) {
      return el["htmlwidget_data_" + name];
    } else if (arguments.length == 3) {
      el["htmlwidget_data_" + name] = value;
      return el;
    } else {
      throw new Error("Wrong number of arguments for elementData: " +
        arguments.length);
    }
  }

  // http://stackoverflow.com/questions/3446170/escape-string-for-use-in-javascript-regex
  function escapeRegExp(str) {
    return str.replace(/[\-\[\]\/\{\}\(\)\*\+\?\.\\\^\$\|]/g, "\\$&");
  }

  function hasClass(el, className) {
    var re = new RegExp("\\b" + escapeRegExp(className) + "\\b");
    return re.test(el.className);
  }

  // elements - array (or array-like object) of HTML elements
  // className - class name to test for
  // include - if true, only return elements with given className;
  //   if false, only return elements *without* given className
  function filterByClass(elements, className, include) {
    var results = [];
    for (var i = 0; i < elements.length; i++) {
      if (hasClass(elements[i], className) == include)
        results.push(elements[i]);
    }
    return results;
  }

  function on(obj, eventName, func) {
    if (obj.addEventListener) {
      obj.addEventListener(eventName, func, false);
    } else if (obj.attachEvent) {
      obj.attachEvent(eventName, func);
    }
  }

  function off(obj, eventName, func) {
    if (obj.removeEventListener)
      obj.removeEventListener(eventName, func, false);
    else if (obj.detachEvent) {
      obj.detachEvent(eventName, func);
    }
  }

  // Translate array of values to top/right/bottom/left, as usual with
  // the "padding" CSS property
  // https://developer.mozilla.org/en-US/docs/Web/CSS/padding
  function unpackPadding(value) {
    if (typeof(value) === "number")
      value = [value];
    if (value.length === 1) {
      return {top: value[0], right: value[0], bottom: value[0], left: value[0]};
    }
    if (value.length === 2) {
      return {top: value[0], right: value[1], bottom: value[0], left: value[1]};
    }
    if (value.length === 3) {
      return {top: value[0], right: value[1], bottom: value[2], left: value[1]};
    }
    if (value.length === 4) {
      return {top: value[0], right: value[1], bottom: value[2], left: value[3]};
    }
  }

  // Convert an unpacked padding object to a CSS value
  function paddingToCss(paddingObj) {
    return paddingObj.top + "px " + paddingObj.right + "px " + paddingObj.bottom + "px " + paddingObj.left + "px";
  }

  // Makes a number suitable for CSS
  function px(x) {
    if (typeof(x) === "number")
      return x + "px";
    else
      return x;
  }

  // Retrieves runtime widget sizing information for an element.
  // The return value is either null, or an object with fill, padding,
  // defaultWidth, defaultHeight fields.
  function sizingPolicy(el) {
    var sizingEl = document.querySelector("script[data-for='" + el.id + "'][type='application/htmlwidget-sizing']");
    if (!sizingEl)
      return null;
    var sp = JSON.parse(sizingEl.textContent || sizingEl.text || "{}");
    if (viewerMode) {
      return sp.viewer;
    } else {
      return sp.browser;
    }
  }

  // @param tasks Array of strings (or falsy value, in which case no-op).
  //   Each element must be a valid JavaScript expression that yields a
  //   function. Or, can be an array of objects with "code" and "data"
  //   properties; in this case, the "code" property should be a string
  //   of JS that's an expr that yields a function, and "data" should be
  //   an object that will be added as an additional argument when that
  //   function is called.
  // @param target The object that will be "this" for each function
  //   execution.
  // @param args Array of arguments to be passed to the functions. (The
  //   same arguments will be passed to all functions.)
  function evalAndRun(tasks, target, args) {
    if (tasks) {
      forEach(tasks, function(task) {
        var theseArgs = args;
        if (typeof(task) === "object") {
          theseArgs = theseArgs.concat([task.data]);
          task = task.code;
        }
        var taskFunc = tryEval(task);
        if (typeof(taskFunc) !== "function") {
          throw new Error("Task must be a function! Source:\n" + task);
        }
        taskFunc.apply(target, theseArgs);
      });
    }
  }

  // Attempt eval() both with and without enclosing in parentheses.
  // Note that enclosing coerces a function declaration into
  // an expression that eval() can parse
  // (otherwise, a SyntaxError is thrown)
  function tryEval(code) {
    var result = null;
    try {
      result = eval(code);
    } catch(error) {
      if (!error instanceof SyntaxError) {
        throw error;
      }
      try {
        result = eval("(" + code + ")");
      } catch(e) {
        if (e instanceof SyntaxError) {
          throw error;
        } else {
          throw e;
        }
      }
    }
    return result;
  }

  function initSizing(el) {
    var sizing = sizingPolicy(el);
    if (!sizing)
      return;

    var cel = document.getElementById("htmlwidget_container");
    if (!cel)
      return;

    if (typeof(sizing.padding) !== "undefined") {
      document.body.style.margin = "0";
      document.body.style.padding = paddingToCss(unpackPadding(sizing.padding));
    }

    if (sizing.fill) {
      document.body.style.overflow = "hidden";
      document.body.style.width = "100%";
      document.body.style.height = "100%";
      document.documentElement.style.width = "100%";
      document.documentElement.style.height = "100%";
      if (cel) {
        cel.style.position = "absolute";
        var pad = unpackPadding(sizing.padding);
        cel.style.top = pad.top + "px";
        cel.style.right = pad.right + "px";
        cel.style.bottom = pad.bottom + "px";
        cel.style.left = pad.left + "px";
        el.style.width = "100%";
        el.style.height = "100%";
      }

      return {
        getWidth: function() { return cel.offsetWidth; },
        getHeight: function() { return cel.offsetHeight; }
      };

    } else {
      el.style.width = px(sizing.width);
      el.style.height = px(sizing.height);

      return {
        getWidth: function() { return el.offsetWidth; },
        getHeight: function() { return el.offsetHeight; }
      };
    }
  }

  // Default implementations for methods
  var defaults = {
    find: function(scope) {
      return querySelectorAll(scope, "." + this.name);
    },
    renderError: function(el, err) {
      var $el = $(el);

      this.clearError(el);

      // Add all these error classes, as Shiny does
      var errClass = "shiny-output-error";
      if (err.type !== null) {
        // use the classes of the error condition as CSS class names
        errClass = errClass + " " + $.map(asArray(err.type), function(type) {
          return errClass + "-" + type;
        }).join(" ");
      }
      errClass = errClass + " htmlwidgets-error";

      // Is el inline or block? If inline or inline-block, just display:none it
      // and add an inline error.
      var display = $el.css("display");
      $el.data("restore-display-mode", display);

      if (display === "inline" || display === "inline-block") {
        $el.hide();
        if (err.message !== "") {
          var errorSpan = $("<span>").addClass(errClass);
          errorSpan.text(err.message);
          $el.after(errorSpan);
        }
      } else if (display === "block") {
        // If block, add an error just after the el, set visibility:none on the
        // el, and position the error to be on top of the el.
        // Mark it with a unique ID and CSS class so we can remove it later.
        $el.css("visibility", "hidden");
        if (err.message !== "") {
          var errorDiv = $("<div>").addClass(errClass).css("position", "absolute")
            .css("top", el.offsetTop)
            .css("left", el.offsetLeft)
            // setting width can push out the page size, forcing otherwise
            // unnecessary scrollbars to appear and making it impossible for
            // the element to shrink; so use max-width instead
            .css("maxWidth", el.offsetWidth)
            .css("height", el.offsetHeight);
          errorDiv.text(err.message);
          $el.after(errorDiv);

          // Really dumb way to keep the size/position of the error in sync with
          // the parent element as the window is resized or whatever.
          var intId = setInterval(function() {
            if (!errorDiv[0].parentElement) {
              clearInterval(intId);
              return;
            }
            errorDiv
              .css("top", el.offsetTop)
              .css("left", el.offsetLeft)
              .css("maxWidth", el.offsetWidth)
              .css("height", el.offsetHeight);
          }, 500);
        }
      }
    },
    clearError: function(el) {
      var $el = $(el);
      var display = $el.data("restore-display-mode");
      $el.data("restore-display-mode", null);

      if (display === "inline" || display === "inline-block") {
        if (display)
          $el.css("display", display);
        $(el.nextSibling).filter(".htmlwidgets-error").remove();
      } else if (display === "block"){
        $el.css("visibility", "inherit");
        $(el.nextSibling).filter(".htmlwidgets-error").remove();
      }
    },
    sizing: {}
  };

  // Called by widget bindings to register a new type of widget. The definition
  // object can contain the following properties:
  // - name (required) - A string indicating the binding name, which will be
  //   used by default as the CSS classname to look for.
  // - initialize (optional) - A function(el) that will be called once per
  //   widget element; if a value is returned, it will be passed as the third
  //   value to renderValue.
  // - renderValue (required) - A function(el, data, initValue) that will be
  //   called with data. Static contexts will cause this to be called once per
  //   element; Shiny apps will cause this to be called multiple times per
  //   element, as the data changes.
  window.HTMLWidgets.widget = function(definition) {
    if (!definition.name) {
      throw new Error("Widget must have a name");
    }
    if (!definition.type) {
      throw new Error("Widget must have a type");
    }
    // Currently we only support output widgets
    if (definition.type !== "output") {
      throw new Error("Unrecognized widget type '" + definition.type + "'");
    }
    // TODO: Verify that .name is a valid CSS classname

    // Support new-style instance-bound definitions. Old-style class-bound
    // definitions have one widget "object" per widget per type/class of
    // widget; the renderValue and resize methods on such widget objects
    // take el and instance arguments, because the widget object can't
    // store them. New-style instance-bound definitions have one widget
    // object per widget instance; the definition that's passed in doesn't
    // provide renderValue or resize methods at all, just the single method
    //   factory(el, width, height)
    // which returns an object that has renderValue(x) and resize(w, h).
    // This enables a far more natural programming style for the widget
    // author, who can store per-instance state using either OO-style
    // instance fields or functional-style closure variables (I guess this
    // is in contrast to what can only be called C-style pseudo-OO which is
    // what we required before).
    if (definition.factory) {
      definition = createLegacyDefinitionAdapter(definition);
    }

    if (!definition.renderValue) {
      throw new Error("Widget must have a renderValue function");
    }

    // For static rendering (non-Shiny), use a simple widget registration
    // scheme. We also use this scheme for Shiny apps/documents that also
    // contain static widgets.
    window.HTMLWidgets.widgets = window.HTMLWidgets.widgets || [];
    // Merge defaults into the definition; don't mutate the original definition.
    var staticBinding = extend({}, defaults, definition);
    overrideMethod(staticBinding, "find", function(superfunc) {
      return function(scope) {
        var results = superfunc(scope);
        // Filter out Shiny outputs, we only want the static kind
        return filterByClass(results, "html-widget-output", false);
      };
    });
    window.HTMLWidgets.widgets.push(staticBinding);

    if (shinyMode) {
      // Shiny is running. Register the definition with an output binding.
      // The definition itself will not be the output binding, instead
      // we will make an output binding object that delegates to the
      // definition. This is because we foolishly used the same method
      // name (renderValue) for htmlwidgets definition and Shiny bindings
      // but they actually have quite different semantics (the Shiny
      // bindings receive data that includes lots of metadata that it
      // strips off before calling htmlwidgets renderValue). We can't
      // just ignore the difference because in some widgets it's helpful
      // to call this.renderValue() from inside of resize(), and if
      // we're not delegating, then that call will go to the Shiny
      // version instead of the htmlwidgets version.

      // Merge defaults with definition, without mutating either.
      var bindingDef = extend({}, defaults, definition);

      // This object will be our actual Shiny binding.
      var shinyBinding = new Shiny.OutputBinding();

      // With a few exceptions, we'll want to simply use the bindingDef's
      // version of methods if they are available, otherwise fall back to
      // Shiny's defaults. NOTE: If Shiny's output bindings gain additional
      // methods in the future, and we want them to be overrideable by
      // HTMLWidget binding definitions, then we'll need to add them to this
      // list.
      delegateMethod(shinyBinding, bindingDef, "getId");
      delegateMethod(shinyBinding, bindingDef, "onValueChange");
      delegateMethod(shinyBinding, bindingDef, "onValueError");
      delegateMethod(shinyBinding, bindingDef, "renderError");
      delegateMethod(shinyBinding, bindingDef, "clearError");
      delegateMethod(shinyBinding, bindingDef, "showProgress");

      // The find, renderValue, and resize are handled differently, because we
      // want to actually decorate the behavior of the bindingDef methods.

      shinyBinding.find = function(scope) {
        var results = bindingDef.find(scope);

        // Only return elements that are Shiny outputs, not static ones
        var dynamicResults = results.filter(".html-widget-output");

        // It's possible that whatever caused Shiny to think there might be
        // new dynamic outputs, also caused there to be new static outputs.
        // Since there might be lots of different htmlwidgets bindings, we
        // schedule execution for later--no need to staticRender multiple
        // times.
        if (results.length !== dynamicResults.length)
          scheduleStaticRender();

        return dynamicResults;
      };

      // Wrap renderValue to handle initialization, which unfortunately isn't
      // supported natively by Shiny at the time of this writing.

      shinyBinding.renderValue = function(el, data) {
        Shiny.renderDependencies(data.deps);
        // Resolve strings marked as javascript literals to objects
        if (!(data.evals instanceof Array)) data.evals = [data.evals];
        for (var i = 0; data.evals && i < data.evals.length; i++) {
          window.HTMLWidgets.evaluateStringMember(data.x, data.evals[i]);
        }
        if (!bindingDef.renderOnNullValue) {
          if (data.x === null) {
            el.style.visibility = "hidden";
            return;
          } else {
            el.style.visibility = "inherit";
          }
        }
        if (!elementData(el, "initialized")) {
          initSizing(el);

          elementData(el, "initialized", true);
          if (bindingDef.initialize) {
            var result = bindingDef.initialize(el, el.offsetWidth,
              el.offsetHeight);
            elementData(el, "init_result", result);
          }
        }
        bindingDef.renderValue(el, data.x, elementData(el, "init_result"));
        evalAndRun(data.jsHooks.render, elementData(el, "init_result"), [el, data.x]);
      };

      // Only override resize if bindingDef implements it
      if (bindingDef.resize) {
        shinyBinding.resize = function(el, width, height) {
          // Shiny can call resize before initialize/renderValue have been
          // called, which doesn't make sense for widgets.
          if (elementData(el, "initialized")) {
            bindingDef.resize(el, width, height, elementData(el, "init_result"));
          }
        };
      }

      Shiny.outputBindings.register(shinyBinding, bindingDef.name);
    }
  };

  var scheduleStaticRenderTimerId = null;
  function scheduleStaticRender() {
    if (!scheduleStaticRenderTimerId) {
      scheduleStaticRenderTimerId = setTimeout(function() {
        scheduleStaticRenderTimerId = null;
        window.HTMLWidgets.staticRender();
      }, 1);
    }
  }

  // Render static widgets after the document finishes loading
  // Statically render all elements that are of this widget's class
  window.HTMLWidgets.staticRender = function() {
    var bindings = window.HTMLWidgets.widgets || [];
    forEach(bindings, function(binding) {
      var matches = binding.find(document.documentElement);
      forEach(matches, function(el) {
        var sizeObj = initSizing(el, binding);

        if (hasClass(el, "html-widget-static-bound"))
          return;
        el.className = el.className + " html-widget-static-bound";

        var initResult;
        if (binding.initialize) {
          initResult = binding.initialize(el,
            sizeObj ? sizeObj.getWidth() : el.offsetWidth,
            sizeObj ? sizeObj.getHeight() : el.offsetHeight
          );
          elementData(el, "init_result", initResult);
        }

        if (binding.resize) {
          var lastSize = {
            w: sizeObj ? sizeObj.getWidth() : el.offsetWidth,
            h: sizeObj ? sizeObj.getHeight() : el.offsetHeight
          };
          var resizeHandler = function(e) {
            var size = {
              w: sizeObj ? sizeObj.getWidth() : el.offsetWidth,
              h: sizeObj ? sizeObj.getHeight() : el.offsetHeight
            };
            if (size.w === 0 && size.h === 0)
              return;
            if (size.w === lastSize.w && size.h === lastSize.h)
              return;
            lastSize = size;
            binding.resize(el, size.w, size.h, initResult);
          };

          on(window, "resize", resizeHandler);

          // This is needed for cases where we're running in a Shiny
          // app, but the widget itself is not a Shiny output, but
          // rather a simple static widget. One example of this is
          // an rmarkdown document that has runtime:shiny and widget
          // that isn't in a render function. Shiny only knows to
          // call resize handlers for Shiny outputs, not for static
          // widgets, so we do it ourselves.
          if (window.jQuery) {
            window.jQuery(document).on(
              "shown.htmlwidgets shown.bs.tab.htmlwidgets shown.bs.collapse.htmlwidgets",
              resizeHandler
            );
            window.jQuery(document).on(
              "hidden.htmlwidgets hidden.bs.tab.htmlwidgets hidden.bs.collapse.htmlwidgets",
              resizeHandler
            );
          }

          // This is needed for the specific case of ioslides, which
          // flips slides between display:none and display:block.
          // Ideally we would not have to have ioslide-specific code
          // here, but rather have ioslides raise a generic event,
          // but the rmarkdown package just went to CRAN so the
          // window to getting that fixed may be long.
          if (window.addEventListener) {
            // It's OK to limit this to window.addEventListener
            // browsers because ioslides itself only supports
            // such browsers.
            on(document, "slideenter", resizeHandler);
            on(document, "slideleave", resizeHandler);
          }
        }

        var scriptData = document.querySelector("script[data-for='" + el.id + "'][type='application/json']");
        if (scriptData) {
          var data = JSON.parse(scriptData.textContent || scriptData.text);
          // Resolve strings marked as javascript literals to objects
          if (!(data.evals instanceof Array)) data.evals = [data.evals];
          for (var k = 0; data.evals && k < data.evals.length; k++) {
            window.HTMLWidgets.evaluateStringMember(data.x, data.evals[k]);
          }
          binding.renderValue(el, data.x, initResult);
          evalAndRun(data.jsHooks.render, initResult, [el, data.x]);
        }
      });
    });

    invokePostRenderHandlers();
  }


  function has_jQuery3() {
    if (!window.jQuery) {
      return false;
    }
    var $version = window.jQuery.fn.jquery;
    var $major_version = parseInt($version.split(".")[0]);
    return $major_version >= 3;
  }

  /*
  / Shiny 1.4 bumped jQuery from 1.x to 3.x which means jQuery's
  / on-ready handler (i.e., $(fn)) is now asyncronous (i.e., it now
  / really means $(setTimeout(fn)).
  / https://jquery.com/upgrade-guide/3.0/#breaking-change-document-ready-handlers-are-now-asynchronous
  /
  / Since Shiny uses $() to schedule initShiny, shiny>=1.4 calls initShiny
  / one tick later than it did before, which means staticRender() is
  / called renderValue() earlier than (advanced) widget authors might be expecting.
  / https://github.com/rstudio/shiny/issues/2630
  /
  / For a concrete example, leaflet has some methods (e.g., updateBounds)
  / which reference Shiny methods registered in initShiny (e.g., setInputValue).
  / Since leaflet is privy to this life-cycle, it knows to use setTimeout() to
  / delay execution of those methods (until Shiny methods are ready)
  / https://github.com/rstudio/leaflet/blob/18ec981/javascript/src/index.js#L266-L268
  /
  / Ideally widget authors wouldn't need to use this setTimeout() hack that
  / leaflet uses to call Shiny methods on a staticRender(). In the long run,
  / the logic initShiny should be broken up so that method registration happens
  / right away, but binding happens later.
  */
  function maybeStaticRenderLater() {
    if (shinyMode && has_jQuery3()) {
      window.jQuery(window.HTMLWidgets.staticRender);
    } else {
      window.HTMLWidgets.staticRender();
    }
  }

  if (document.addEventListener) {
    document.addEventListener("DOMContentLoaded", function() {
      document.removeEventListener("DOMContentLoaded", arguments.callee, false);
      maybeStaticRenderLater();
    }, false);
  } else if (document.attachEvent) {
    document.attachEvent("onreadystatechange", function() {
      if (document.readyState === "complete") {
        document.detachEvent("onreadystatechange", arguments.callee);
        maybeStaticRenderLater();
      }
    });
  }


  window.HTMLWidgets.getAttachmentUrl = function(depname, key) {
    // If no key, default to the first item
    if (typeof(key) === "undefined")
      key = 1;

    var link = document.getElementById(depname + "-" + key + "-attachment");
    if (!link) {
      throw new Error("Attachment " + depname + "/" + key + " not found in document");
    }
    return link.getAttribute("href");
  };

  window.HTMLWidgets.dataframeToD3 = function(df) {
    var names = [];
    var length;
    for (var name in df) {
        if (df.hasOwnProperty(name))
            names.push(name);
        if (typeof(df[name]) !== "object" || typeof(df[name].length) === "undefined") {
            throw new Error("All fields must be arrays");
        } else if (typeof(length) !== "undefined" && length !== df[name].length) {
            throw new Error("All fields must be arrays of the same length");
        }
        length = df[name].length;
    }
    var results = [];
    var item;
    for (var row = 0; row < length; row++) {
        item = {};
        for (var col = 0; col < names.length; col++) {
            item[names[col]] = df[names[col]][row];
        }
        results.push(item);
    }
    return results;
  };

  window.HTMLWidgets.transposeArray2D = function(array) {
      if (array.length === 0) return array;
      var newArray = array[0].map(function(col, i) {
          return array.map(function(row) {
              return row[i]
          })
      });
      return newArray;
  };
  // Split value at splitChar, but allow splitChar to be escaped
  // using escapeChar. Any other characters escaped by escapeChar
  // will be included as usual (including escapeChar itself).
  function splitWithEscape(value, splitChar, escapeChar) {
    var results = [];
    var escapeMode = false;
    var currentResult = "";
    for (var pos = 0; pos < value.length; pos++) {
      if (!escapeMode) {
        if (value[pos] === splitChar) {
          results.push(currentResult);
          currentResult = "";
        } else if (value[pos] === escapeChar) {
          escapeMode = true;
        } else {
          currentResult += value[pos];
        }
      } else {
        currentResult += value[pos];
        escapeMode = false;
      }
    }
    if (currentResult !== "") {
      results.push(currentResult);
    }
    return results;
  }
  // Function authored by Yihui/JJ Allaire
  window.HTMLWidgets.evaluateStringMember = function(o, member) {
    var parts = splitWithEscape(member, '.', '\\');
    for (var i = 0, l = parts.length; i < l; i++) {
      var part = parts[i];
      // part may be a character or 'numeric' member name
      if (o !== null && typeof o === "object" && part in o) {
        if (i == (l - 1)) { // if we are at the end of the line then evalulate
          if (typeof o[part] === "string")
            o[part] = tryEval(o[part]);
        } else { // otherwise continue to next embedded object
          o = o[part];
        }
      }
    }
  };

  // Retrieve the HTMLWidget instance (i.e. the return value of an
  // HTMLWidget binding's initialize() or factory() function)
  // associated with an element, or null if none.
  window.HTMLWidgets.getInstance = function(el) {
    return elementData(el, "init_result");
  };

  // Finds the first element in the scope that matches the selector,
  // and returns the HTMLWidget instance (i.e. the return value of
  // an HTMLWidget binding's initialize() or factory() function)
  // associated with that element, if any. If no element matches the
  // selector, or the first matching element has no HTMLWidget
  // instance associated with it, then null is returned.
  //
  // The scope argument is optional, and defaults to window.document.
  window.HTMLWidgets.find = function(scope, selector) {
    if (arguments.length == 1) {
      selector = scope;
      scope = document;
    }

    var el = scope.querySelector(selector);
    if (el === null) {
      return null;
    } else {
      return window.HTMLWidgets.getInstance(el);
    }
  };

  // Finds all elements in the scope that match the selector, and
  // returns the HTMLWidget instances (i.e. the return values of
  // an HTMLWidget binding's initialize() or factory() function)
  // associated with the elements, in an array. If elements that
  // match the selector don't have an associated HTMLWidget
  // instance, the returned array will contain nulls.
  //
  // The scope argument is optional, and defaults to window.document.
  window.HTMLWidgets.findAll = function(scope, selector) {
    if (arguments.length == 1) {
      selector = scope;
      scope = document;
    }

    var nodes = scope.querySelectorAll(selector);
    var results = [];
    for (var i = 0; i < nodes.length; i++) {
      results.push(window.HTMLWidgets.getInstance(nodes[i]));
    }
    return results;
  };

  var postRenderHandlers = [];
  function invokePostRenderHandlers() {
    while (postRenderHandlers.length) {
      var handler = postRenderHandlers.shift();
      if (handler) {
        handler();
      }
    }
  }

  // Register the given callback function to be invoked after the
  // next time static widgets are rendered.
  window.HTMLWidgets.addPostRenderHandler = function(callback) {
    postRenderHandlers.push(callback);
  };

  // Takes a new-style instance-bound definition, and returns an
  // old-style class-bound definition. This saves us from having
  // to rewrite all the logic in this file to accomodate both
  // types of definitions.
  function createLegacyDefinitionAdapter(defn) {
    var result = {
      name: defn.name,
      type: defn.type,
      initialize: function(el, width, height) {
        return defn.factory(el, width, height);
      },
      renderValue: function(el, x, instance) {
        return instance.renderValue(x);
      },
      resize: function(el, width, height, instance) {
        return instance.resize(width, height);
      }
    };

    if (defn.find)
      result.find = defn.find;
    if (defn.renderError)
      result.renderError = defn.renderError;
    if (defn.clearError)
      result.clearError = defn.clearError;

    return result;
  }
})();
```
