## Supplemental Data 2 for "Rewiring phospholipid biosynthesis reveals robustness in membrane homeostasis and uncovers lipid regulatory players": plotly-latest.min.js

```
/**
* plotly.js v1.49.4
* Copyright 2012-2019, Plotly, Inc.
* All rights reserved.
* Licensed under the MIT license
*/
!function(t){if("object"==typeof exports&&"undefined"!=typeof module)module.exports=t();else if("function"==typeof define&&define.amd)define([],t);else{("undefined"!=typeof window?window:"undefined"!=typeof global?global:"undefined"!=typeof self?self:this).Plotly=t()}}(function(){return function(){return function t(e,r,n){function i(o,s){if(!r[o]){if(!e[o]){var l="function"==typeof require&&require;if(!s&&l)return l(o,!0);if(a)return a(o,!0);var c=new Error("Cannot find module '"+o+"'");throw c.code="MODULE_NOT_FOUND",c}var u=r[o]={exports:{}};e[o][0].call(u.exports,function(t){return i(e[o][1][t]||t)},u,u.exports,t,e,r,n)}return r[o].exports}for(var a="function"==typeof require&&require,o=0;o<n.length;o++)i(n[o]);return i}}()({1:[function(t,e,r){"use strict";var n=t("../src/lib"),i={"X,X div":"direction:ltr;font-family:'Open Sans', verdana, arial, sans-serif;margin:0;padding:0;","X input,X button":"font-family:'Open Sans', verdana, arial, sans-serif;","X input:focus,X button:focus":"outline:none;","X a":"text-decoration:none;","X a:hover":"text-decoration:none;","X .crisp":"shape-rendering:crispEdges;","X .user-select-none":"-webkit-user-select:none;-moz-user-select:none;-ms-user-select:none;-o-user-select:none;user-select:none;","X svg":"overflow:hidden;","X svg a":"fill:#447adb;","X svg a:hover":"fill:#3c6dc5;","X .main-svg":"position:absolute;top:0;left:0;pointer-events:none;","X .main-svg .draglayer":"pointer-events:all;","X .cursor-default":"cursor:default;","X .cursor-pointer":"cursor:pointer;","X .cursor-crosshair":"cursor:crosshair;","X .cursor-move":"cursor:move;","X .cursor-col-resize":"cursor:col-resize;","X .cursor-row-resize":"cursor:row-resize;","X .cursor-ns-resize":"cursor:ns-resize;","X .cursor-ew-resize":"cursor:ew-resize;","X .cursor-sw-resize":"cursor:sw-resize;","X .cursor-s-resize":"cursor:s-resize;","X .cursor-se-resize":"cursor:se-resize;","X .cursor-w-resize":"cursor:w-resize;","X .cursor-e-resize":"cursor:e-resize;","X .cursor-nw-resize":"cursor:nw-resize;","X .cursor-n-resize":"cursor:n-resize;","X .cursor-ne-resize":"cursor:ne-resize;","X .cursor-grab":"cursor:-webkit-grab;cursor:grab;","X .modebar":"position:absolute;top:2px;right:2px;","X .ease-bg":"-webkit-transition:background-color 0.3s ease 0s;-moz-transition:background-color 0.3s ease 0s;-ms-transition:background-color 0.3s ease 0s;-o-transition:background-color 0.3s ease 0s;transition:background-color 0.3s ease 0s;","X .modebar--hover>:not(.watermark)":"opacity:0;-webkit-transition:opacity 0.3s ease 0s;-moz-transition:opacity 0.3s ease 0s;-ms-transition:opacity 0.3s ease 0s;-o-transition:opacity 0.3s ease 0s;transition:opacity 0.3s ease 0s;","X:hover .modebar--hover .modebar-group":"opacity:1;","X .modebar-group":"float:left;display:inline-block;box-sizing:border-box;padding-left:8px;position:relative;vertical-align:middle;white-space:nowrap;","X .modebar-btn":"position:relative;font-size:16px;padding:3px 4px;height:22px;cursor:pointer;line-height:normal;box-sizing:border-box;","X .modebar-btn svg":"position:relative;top:2px;","X .modebar.vertical":"display:flex;flex-direction:column;flex-wrap:wrap;align-content:flex-end;max-height:100%;","X .modebar.vertical svg":"top:-1px;","X .modebar.vertical .modebar-group":"display:block;float:none;padding-left:0px;padding-bottom:8px;","X .modebar.vertical .modebar-group .modebar-btn":"display:block;text-align:center;","X [data-title]:before,X [data-title]:after":"position:absolute;-webkit-transform:translate3d(0, 0, 0);-moz-transform:translate3d(0, 0, 0);-ms-transform:translate3d(0, 0, 0);-o-transform:translate3d(0, 0, 0);transform:translate3d(0, 0, 0);display:none;opacity:0;z-index:1001;pointer-events:none;top:110%;right:50%;","X [data-title]:hover:before,X [data-title]:hover:after":"display:block;opacity:1;","X [data-title]:before":"content:'';position:absolute;background:transparent;border:6px solid transparent;z-index:1002;margin-top:-12px;border-bottom-color:#69738a;margin-right:-6px;","X [data-title]:after":"content:attr(data-title);background:#69738a;color:white;padding:8px 10px;font-size:12px;line-height:12px;white-space:nowrap;margin-right:-18px;border-radius:2px;","X .vertical [data-title]:before,X .vertical [data-title]:after":"top:0%;right:200%;","X .vertical [data-title]:before":"border:6px solid transparent;border-left-color:#69738a;margin-top:8px;margin-right:-30px;","X .select-outline":"fill:none;stroke-width:1;shape-rendering:crispEdges;","X .select-outline-1":"stroke:white;","X .select-outline-2":"stroke:black;stroke-dasharray:2px 2px;",Y:"font-family:'Open Sans';position:fixed;top:50px;right:20px;z-index:10000;font-size:10pt;max-width:180px;","Y p":"margin:0;","Y .notifier-note":"min-width:180px;max-width:250px;border:1px solid #fff;z-index:3000;margin:0;background-color:#8c97af;background-color:rgba(140,151,175,0.9);color:#fff;padding:10px;overflow-wrap:break-word;word-wrap:break-word;-ms-hyphens:auto;-webkit-hyphens:auto;hyphens:auto;","Y .notifier-close":"color:#fff;opacity:0.8;float:right;padding:0 5px;background:none;border:none;font-size:20px;font-weight:bold;line-height:20px;","Y .notifier-close:hover":"color:#444;text-decoration:none;cursor:pointer;"};for(var a in i){var o=a.replace(/^,/," ,").replace(/X/g,".js-plotly-plot .plotly").replace(/Y/g,".plotly-notifier");n.addStyleRule(o,i[a])}},{"../src/lib":719}],2:[function(t,e,r){"use strict";e.exports=t("../src/transforms/aggregate")},{"../src/transforms/aggregate":1266}],3:[function(t,e,r){"use strict";e.exports=t("../src/traces/bar")},{"../src/traces/bar":864}],4:[function(t,e,r){"use strict";e.exports=t("../src/traces/barpolar")},{"../src/traces/barpolar":876}],5:[function(t,e,r){"use strict";e.exports=t("../src/traces/box")},{"../src/traces/box":886}],6:[function(t,e,r){"use strict";e.exports=t("../src/components/calendars")},{"../src/components/calendars":591}],7:[function(t,e,r){"use strict";e.exports=t("../src/traces/candlestick")},{"../src/traces/candlestick":895}],8:[function(t,e,r){"use strict";e.exports=t("../src/traces/carpet")},{"../src/traces/carpet":914}],9:[function(t,e,r){"use strict";e.exports=t("../src/traces/choropleth")},{"../src/traces/choropleth":928}],10:[function(t,e,r){"use strict";e.exports=t("../src/traces/choroplethmapbox")},{"../src/traces/choroplethmapbox":935}],11:[function(t,e,r){"use strict";e.exports=t("../src/traces/cone")},{"../src/traces/cone":941}],12:[function(t,e,r){"use strict";e.exports=t("../src/traces/contour")},{"../src/traces/contour":956}],13:[function(t,e,r){"use strict";e.exports=t("../src/traces/contourcarpet")},{"../src/traces/contourcarpet":967}],14:[function(t,e,r){"use strict";e.exports=t("../src/core")},{"../src/core":697}],15:[function(t,e,r){"use strict";e.exports=t("../src/traces/densitymapbox")},{"../src/traces/densitymapbox":977}],16:[function(t,e,r){"use strict";e.exports=t("../src/transforms/filter")},{"../src/transforms/filter":1267}],17:[function(t,e,r){"use strict";e.exports=t("../src/traces/funnel")},{"../src/traces/funnel":987}],18:[function(t,e,r){"use strict";e.exports=t("../src/traces/funnelarea")},{"../src/traces/funnelarea":996}],19:[function(t,e,r){"use strict";e.exports=t("../src/transforms/groupby")},{"../src/transforms/groupby":1268}],20:[function(t,e,r){"use strict";e.exports=t("../src/traces/heatmap")},{"../src/traces/heatmap":1009}],21:[function(t,e,r){"use strict";e.exports=t("../src/traces/heatmapgl")},{"../src/traces/heatmapgl":1018}],22:[function(t,e,r){"use strict";e.exports=t("../src/traces/histogram")},{"../src/traces/histogram":1030}],23:[function(t,e,r){"use strict";e.exports=t("../src/traces/histogram2d")},{"../src/traces/histogram2d":1036}],24:[function(t,e,r){"use strict";e.exports=t("../src/traces/histogram2dcontour")},{"../src/traces/histogram2dcontour":1040}],25:[function(t,e,r){"use strict";var n=t("./core");n.register([t("./bar"),t("./box"),t("./heatmap"),t("./histogram"),t("./histogram2d"),t("./histogram2dcontour"),t("./contour"),t("./scatterternary"),t("./violin"),t("./funnel"),t("./waterfall"),t("./pie"),t("./sunburst"),t("./funnelarea"),t("./scatter3d"),t("./surface"),t("./isosurface"),t("./volume"),t("./mesh3d"),t("./cone"),t("./streamtube"),t("./scattergeo"),t("./choropleth"),t("./scattergl"),t("./splom"),t("./pointcloud"),t("./heatmapgl"),t("./parcoords"),t("./parcats"),t("./scattermapbox"),t("./choroplethmapbox"),t("./densitymapbox"),t("./sankey"),t("./indicator"),t("./table"),t("./carpet"),t("./scattercarpet"),t("./contourcarpet"),t("./ohlc"),t("./candlestick"),t("./scatterpolar"),t("./scatterpolargl"),t("./barpolar")]),n.register([t("./aggregate"),t("./filter"),t("./groupby"),t("./sort")]),n.register([t("./calendars")]),e.exports=n},{"./aggregate":2,"./bar":3,"./barpolar":4,"./box":5,"./calendars":6,"./candlestick":7,"./carpet":8,"./choropleth":9,"./choroplethmapbox":10,"./cone":11,"./contour":12,"./contourcarpet":13,"./core":14,"./densitymapbox":15,"./filter":16,"./funnel":17,"./funnelarea":18,"./groupby":19,"./heatmap":20,"./heatmapgl":21,"./histogram":22,"./histogram2d":23,"./histogram2dcontour":24,"./indicator":26,"./isosurface":27,"./mesh3d":28,"./ohlc":29,"./parcats":30,"./parcoords":31,"./pie":32,"./pointcloud":33,"./sankey":34,"./scatter3d":35,"./scattercarpet":36,"./scattergeo":37,"./scattergl":38,"./scattermapbox":39,"./scatterpolar":40,"./scatterpolargl":41,"./scatterternary":42,"./sort":43,"./splom":44,"./streamtube":45,"./sunburst":46,"./surface":47,"./table":48,"./violin":49,"./volume":50,"./waterfall":51}],26:[function(t,e,r){"use strict";e.exports=t("../src/traces/indicator")},{"../src/traces/indicator":1046}],27:[function(t,e,r){"use strict";e.exports=t("../src/traces/isosurface")},{"../src/traces/isosurface":1052}],28:[function(t,e,r){"use strict";e.exports=t("../src/traces/mesh3d")},{"../src/traces/mesh3d":1057}],29:[function(t,e,r){"use strict";e.exports=t("../src/traces/ohlc")},{"../src/traces/ohlc":1062}],30:[function(t,e,r){"use strict";e.exports=t("../src/traces/parcats")},{"../src/traces/parcats":1071}],31:[function(t,e,r){"use strict";e.exports=t("../src/traces/parcoords")},{"../src/traces/parcoords":1081}],32:[function(t,e,r){"use strict";e.exports=t("../src/traces/pie")},{"../src/traces/pie":1092}],33:[function(t,e,r){"use strict";e.exports=t("../src/traces/pointcloud")},{"../src/traces/pointcloud":1101}],34:[function(t,e,r){"use strict";e.exports=t("../src/traces/sankey")},{"../src/traces/sankey":1107}],35:[function(t,e,r){"use strict";e.exports=t("../src/traces/scatter3d")},{"../src/traces/scatter3d":1143}],36:[function(t,e,r){"use strict";e.exports=t("../src/traces/scattercarpet")},{"../src/traces/scattercarpet":1149}],37:[function(t,e,r){"use strict";e.exports=t("../src/traces/scattergeo")},{"../src/traces/scattergeo":1156}],38:[function(t,e,r){"use strict";e.exports=t("../src/traces/scattergl")},{"../src/traces/scattergl":1167}],39:[function(t,e,r){"use strict";e.exports=t("../src/traces/scattermapbox")},{"../src/traces/scattermapbox":1176}],40:[function(t,e,r){"use strict";e.exports=t("../src/traces/scatterpolar")},{"../src/traces/scatterpolar":1183}],41:[function(t,e,r){"use strict";e.exports=t("../src/traces/scatterpolargl")},{"../src/traces/scatterpolargl":1189}],42:[function(t,e,r){"use strict";e.exports=t("../src/traces/scatterternary")},{"../src/traces/scatterternary":1196}],43:[function(t,e,r){"use strict";e.exports=t("../src/transforms/sort")},{"../src/transforms/sort":1270}],44:[function(t,e,r){"use strict";e.exports=t("../src/traces/splom")},{"../src/traces/splom":1205}],45:[function(t,e,r){"use strict";e.exports=t("../src/traces/streamtube")},{"../src/traces/streamtube":1213}],46:[function(t,e,r){"use strict";e.exports=t("../src/traces/sunburst")},{"../src/traces/sunburst":1220}],47:[function(t,e,r){"use strict";e.exports=t("../src/traces/surface")},{"../src/traces/surface":1229}],48:[function(t,e,r){"use strict";e.exports=t("../src/traces/table")},{"../src/traces/table":1237}],49:[function(t,e,r){"use strict";e.exports=t("../src/traces/violin")},{"../src/traces/violin":1245}],50:[function(t,e,r){"use strict";e.exports=t("../src/traces/volume")},{"../src/traces/volume":1253}],51:[function(t,e,r){"use strict";e.exports=t("../src/traces/waterfall")},{"../src/traces/waterfall":1261}],52:[function(t,e,r){"use strict";e.exports=function(t){var e=(t=t||{}).eye||[0,0,1],r=t.center||[0,0,0],s=t.up||[0,1,0],l=t.distanceLimits||[0,1/0],c=t.mode||"turntable",u=n(),h=i(),f=a();return u.setDistanceLimits(l[0],l[1]),u.lookAt(0,e,r,s),h.setDistanceLimits(l[0],l[1]),h.lookAt(0,e,r,s),f.setDistanceLimits(l[0],l[1]),f.lookAt(0,e,r,s),new o({turntable:u,orbit:h,matrix:f},c)};var n=t("turntable-camera-controller"),i=t("orbit-camera-controller"),a=t("matrix-camera-controller");function o(t,e){this._controllerNames=Object.keys(t),this._controllerList=this._controllerNames.map(function(e){return t[e]}),this._mode=e,this._active=t[e],this._active||(this._mode="turntable",this._active=t.turntable),this.modes=this._controllerNames,this.computedMatrix=this._active.computedMatrix,this.computedEye=this._active.computedEye,this.computedUp=this._active.computedUp,this.computedCenter=this._active.computedCenter,this.computedRadius=this._active.computedRadius}var s=o.prototype;[["flush",1],["idle",1],["lookAt",4],["rotate",4],["pan",4],["translate",4],["setMatrix",2],["setDistanceLimits",2],["setDistance",2]].forEach(function(t){for(var e=t[0],r=[],n=0;n<t[1];++n)r.push("a"+n);var i="var cc=this._controllerList;for(var i=0;i<cc.length;++i){cc[i]."+t[0]+"("+r.join()+")}";s[e]=Function.apply(null,r.concat(i))}),s.recalcMatrix=function(t){this._active.recalcMatrix(t)},s.getDistance=function(t){return this._active.getDistance(t)},s.getDistanceLimits=function(t){return this._active.getDistanceLimits(t)},s.lastT=function(){return this._active.lastT()},s.setMode=function(t){if(t!==this._mode){var e=this._controllerNames.indexOf(t);if(!(e<0)){var r=this._active,n=this._controllerList[e],i=Math.max(r.lastT(),n.lastT());r.recalcMatrix(i),n.setMatrix(i,r.computedMatrix),this._active=n,this._mode=t,this.computedMatrix=this._active.computedMatrix,this.computedEye=this._active.computedEye,this.computedUp=this._active.computedUp,this.computedCenter=this._active.computedCenter,this.computedRadius=this._active.computedRadius}}},s.getMode=function(){return this._mode}},{"matrix-camera-controller":433,"orbit-camera-controller":456,"turntable-camera-controller":542}],53:[function(t,e,r){var n,i;n=this,i=function(t,e,r,n,i){"use strict";function a(t){return t.target.depth}function o(t,e){return t.sourceLinks.length?t.depth:e-1}function s(t){return function(){return t}}i=i&&i.hasOwnProperty("default")?i.default:i;var l="function"==typeof Symbol&&"symbol"==typeof Symbol.iterator?function(t){return typeof t}:function(t){return t&&"function"==typeof Symbol&&t.constructor===Symbol&&t!==Symbol.prototype?"symbol":typeof t};function c(t,e){return h(t.source,e.source)||t.index-e.index}function u(t,e){return h(t.target,e.target)||t.index-e.index}function h(t,e){return t.partOfCycle===e.partOfCycle?t.y0-e.y0:"top"===t.circularLinkType||"bottom"===e.circularLinkType?-1:1}function f(t){return t.value}function p(t){return(t.y0+t.y1)/2}function d(t){return p(t.source)}function g(t){return p(t.target)}function v(t){return t.index}function m(t){return t.nodes}function y(t){return t.links}function x(t,e){var r=t.get(e);if(!r)throw new Error("missing: "+e);return r}function b(t,e){return e(t)}var _=25,w=10,k=.3;function T(t,e){var r=0,n=0;t.links.forEach(function(i){i.circular&&(i.source.circularLinkType||i.target.circularLinkType?i.circularLinkType=i.source.circularLinkType?i.source.circularLinkType:i.target.circularLinkType:i.circularLinkType=r<n?"top":"bottom","top"==i.circularLinkType?r+=1:n+=1,t.nodes.forEach(function(t){b(t,e)!=b(i.source,e)&&b(t,e)!=b(i.target,e)||(t.circularLinkType=i.circularLinkType)}))}),t.links.forEach(function(t){t.circular&&(t.source.circularLinkType==t.target.circularLinkType&&(t.circularLinkType=t.source.circularLinkType),Y(t,e)&&(t.circularLinkType=t.source.circularLinkType))})}function A(t){var e=Math.abs(t.y1-t.y0),r=Math.abs(t.target.x0-t.source.x1);return Math.atan(r/e)}function M(t,e){var r=0;t.sourceLinks.forEach(function(t){r=t.circular&&!Y(t,e)?r+1:r});var n=0;return t.targetLinks.forEach(function(t){n=t.circular&&!Y(t,e)?n+1:n}),r+n}function S(t){var e=t.source.sourceLinks,r=0;e.forEach(function(t){r=t.circular?r+1:r});var n=t.target.targetLinks,i=0;return n.forEach(function(t){i=t.circular?i+1:i}),!(r>1||i>1)}function E(t,e,r){return t.sort(L),t.forEach(function(n,i){var a,o,s=0;if(Y(n,r)&&S(n))n.circularPathData.verticalBuffer=s+n.width/2;else{for(var l=0;l<i;l++)if(a=t[i],o=t[l],!(a.source.column<o.target.column||a.target.column>o.source.column)){var c=t[l].circularPathData.verticalBuffer+t[l].width/2+e;s=c>s?c:s}n.circularPathData.verticalBuffer=s+n.width/2}}),t}function C(t,r,i,a){var o=e.min(t.links,function(t){return t.source.y0});t.links.forEach(function(t){t.circular&&(t.circularPathData={})}),E(t.links.filter(function(t){return"top"==t.circularLinkType}),r,a),E(t.links.filter(function(t){return"bottom"==t.circularLinkType}),r,a),t.links.forEach(function(e){if(e.circular){if(e.circularPathData.arcRadius=e.width+w,e.circularPathData.leftNodeBuffer=5,e.circularPathData.rightNodeBuffer=5,e.circularPathData.sourceWidth=e.source.x1-e.source.x0,e.circularPathData.sourceX=e.source.x0+e.circularPathData.sourceWidth,e.circularPathData.targetX=e.target.x0,e.circularPathData.sourceY=e.y0,e.circularPathData.targetY=e.y1,Y(e,a)&&S(e))e.circularPathData.leftSmallArcRadius=w+e.width/2,e.circularPathData.leftLargeArcRadius=w+e.width/2,e.circularPathData.rightSmallArcRadius=w+e.width/2,e.circularPathData.rightLargeArcRadius=w+e.width/2,"bottom"==e.circularLinkType?(e.circularPathData.verticalFullExtent=e.source.y1+_+e.circularPathData.verticalBuffer,e.circularPathData.verticalLeftInnerExtent=e.circularPathData.verticalFullExtent-e.circularPathData.leftLargeArcRadius,e.circularPathData.verticalRightInnerExtent=e.circularPathData.verticalFullExtent-e.circularPathData.rightLargeArcRadius):(e.circularPathData.verticalFullExtent=e.source.y0-_-e.circularPathData.verticalBuffer,e.circularPathData.verticalLeftInnerExtent=e.circularPathData.verticalFullExtent+e.circularPathData.leftLargeArcRadius,e.circularPathData.verticalRightInnerExtent=e.circularPathData.verticalFullExtent+e.circularPathData.rightLargeArcRadius);else{var s=e.source.column,l=e.circularLinkType,c=t.links.filter(function(t){return t.source.column==s&&t.circularLinkType==l});"bottom"==e.circularLinkType?c.sort(O):c.sort(P);var u=0;c.forEach(function(t,n){t.circularLinkID==e.circularLinkID&&(e.circularPathData.leftSmallArcRadius=w+e.width/2+u,e.circularPathData.leftLargeArcRadius=w+e.width/2+n*r+u),u+=t.width}),s=e.target.column,c=t.links.filter(function(t){return t.target.column==s&&t.circularLinkType==l}),"bottom"==e.circularLinkType?c.sort(I):c.sort(z),u=0,c.forEach(function(t,n){t.circularLinkID==e.circularLinkID&&(e.circularPathData.rightSmallArcRadius=w+e.width/2+u,e.circularPathData.rightLargeArcRadius=w+e.width/2+n*r+u),u+=t.width}),"bottom"==e.circularLinkType?(e.circularPathData.verticalFullExtent=Math.max(i,e.source.y1,e.target.y1)+_+e.circularPathData.verticalBuffer,e.circularPathData.verticalLeftInnerExtent=e.circularPathData.verticalFullExtent-e.circularPathData.leftLargeArcRadius,e.circularPathData.verticalRightInnerExtent=e.circularPathData.verticalFullExtent-e.circularPathData.rightLargeArcRadius):(e.circularPathData.verticalFullExtent=o-_-e.circularPathData.verticalBuffer,e.circularPathData.verticalLeftInnerExtent=e.circularPathData.verticalFullExtent+e.circularPathData.leftLargeArcRadius,e.circularPathData.verticalRightInnerExtent=e.circularPathData.verticalFullExtent+e.circularPathData.rightLargeArcRadius)}e.circularPathData.leftInnerExtent=e.circularPathData.sourceX+e.circularPathData.leftNodeBuffer,e.circularPathData.rightInnerExtent=e.circularPathData.targetX-e.circularPathData.rightNodeBuffer,e.circularPathData.leftFullExtent=e.circularPathData.sourceX+e.circularPathData.leftLargeArcRadius+e.circularPathData.leftNodeBuffer,e.circularPathData.rightFullExtent=e.circularPathData.targetX-e.circularPathData.rightLargeArcRadius-e.circularPathData.rightNodeBuffer}if(e.circular)e.path=function(t){var e="";e="top"==t.circularLinkType?"M"+t.circularPathData.sourceX+" "+t.circularPathData.sourceY+" L"+t.circularPathData.leftInnerExtent+" "+t.circularPathData.sourceY+" A"+t.circularPathData.leftLargeArcRadius+" "+t.circularPathData.leftSmallArcRadius+" 0 0 0 "+t.circularPathData.leftFullExtent+" "+(t.circularPathData.sourceY-t.circularPathData.leftSmallArcRadius)+" L"+t.circularPathData.leftFullExtent+" "+t.circularPathData.verticalLeftInnerExtent+" A"+t.circularPathData.leftLargeArcRadius+" "+t.circularPathData.leftLargeArcRadius+" 0 0 0 "+t.circularPathData.leftInnerExtent+" "+t.circularPathData.verticalFullExtent+" L"+t.circularPathData.rightInnerExtent+" "+t.circularPathData.verticalFullExtent+" A"+t.circularPathData.rightLargeArcRadius+" "+t.circularPathData.rightLargeArcRadius+" 0 0 0 "+t.circularPathData.rightFullExtent+" "+t.circularPathData.verticalRightInnerExtent+" L"+t.circularPathData.rightFullExtent+" "+(t.circularPathData.targetY-t.circularPathData.rightSmallArcRadius)+" A"+t.circularPathData.rightLargeArcRadius+" "+t.circularPathData.rightSmallArcRadius+" 0 0 0 "+t.circularPathData.rightInnerExtent+" "+t.circularPathData.targetY+" L"+t.circularPathData.targetX+" "+t.circularPathData.targetY:"M"+t.circularPathData.sourceX+" "+t.circularPathData.sourceY+" L"+t.circularPathData.leftInnerExtent+" "+t.circularPathData.sourceY+" A"+t.circularPathData.leftLargeArcRadius+" "+t.circularPathData.leftSmallArcRadius+" 0 0 1 "+t.circularPathData.leftFullExtent+" "+(t.circularPathData.sourceY+t.circularPathData.leftSmallArcRadius)+" L"+t.circularPathData.leftFullExtent+" "+t.circularPathData.verticalLeftInnerExtent+" A"+t.circularPathData.leftLargeArcRadius+" "+t.circularPathData.leftLargeArcRadius+" 0 0 1 "+t.circularPathData.leftInnerExtent+" "+t.circularPathData.verticalFullExtent+" L"+t.circularPathData.rightInnerExtent+" "+t.circularPathData.verticalFullExtent+" A"+t.circularPathData.rightLargeArcRadius+" "+t.circularPathData.rightLargeArcRadius+" 0 0 1 "+t.circularPathData.rightFullExtent+" "+t.circularPathData.verticalRightInnerExtent+" L"+t.circularPathData.rightFullExtent+" "+(t.circularPathData.targetY+t.circularPathData.rightSmallArcRadius)+" A"+t.circularPathData.rightLargeArcRadius+" "+t.circularPathData.rightSmallArcRadius+" 0 0 1 "+t.circularPathData.rightInnerExtent+" "+t.circularPathData.targetY+" L"+t.circularPathData.targetX+" "+t.circularPathData.targetY;return e}(e);else{var h=n.linkHorizontal().source(function(t){return[t.source.x0+(t.source.x1-t.source.x0),t.y0]}).target(function(t){return[t.target.x0,t.y1]});e.path=h(e)}})}function L(t,e){return D(t)==D(e)?"bottom"==t.circularLinkType?O(t,e):P(t,e):D(e)-D(t)}function P(t,e){return t.y0-e.y0}function O(t,e){return e.y0-t.y0}function z(t,e){return t.y1-e.y1}function I(t,e){return e.y1-t.y1}function D(t){return t.target.column-t.source.column}function R(t){return t.target.x0-t.source.x1}function F(t,e){var r=A(t),n=R(e)/Math.tan(r);return"up"==G(t)?t.y1+n:t.y1-n}function B(t,e){var r=A(t),n=R(e)/Math.tan(r);return"up"==G(t)?t.y1-n:t.y1+n}function N(t,e,r,n){t.links.forEach(function(i){if(!i.circular&&i.target.column-i.source.column>1){var a=i.source.column+1,o=i.target.column-1,s=1,l=o-a+1;for(s=1;a<=o;a++,s++)t.nodes.forEach(function(o){if(o.column==a){var c,u=s/(l+1),h=Math.pow(1-u,3),f=3*u*Math.pow(1-u,2),p=3*Math.pow(u,2)*(1-u),d=Math.pow(u,3),g=h*i.y0+f*i.y0+p*i.y1+d*i.y1,v=g-i.width/2,m=g+i.width/2;v>o.y0&&v<o.y1?(c=o.y1-v+10,c="bottom"==o.circularLinkType?c:-c,o=U(o,c,e,r),t.nodes.forEach(function(t){b(t,n)!=b(o,n)&&t.column==o.column&&j(o,t)&&U(t,c,e,r)})):m>o.y0&&m<o.y1?(c=m-o.y0+10,o=U(o,c,e,r),t.nodes.forEach(function(t){b(t,n)!=b(o,n)&&t.column==o.column&&t.y0<o.y1&&t.y1>o.y1&&U(t,c,e,r)})):v<o.y0&&m>o.y1&&(c=m-o.y0+10,o=U(o,c,e,r),t.nodes.forEach(function(t){b(t,n)!=b(o,n)&&t.column==o.column&&t.y0<o.y1&&t.y1>o.y1&&U(t,c,e,r)}))}})}})}function j(t,e){return t.y0>e.y0&&t.y0<e.y1||(t.y1>e.y0&&t.y1<e.y1||t.y0<e.y0&&t.y1>e.y1)}function U(t,e,r,n){return t.y0+e>=r&&t.y1+e<=n&&(t.y0=t.y0+e,t.y1=t.y1+e,t.targetLinks.forEach(function(t){t.y1=t.y1+e}),t.sourceLinks.forEach(function(t){t.y0=t.y0+e})),t}function V(t,e,r,n){t.nodes.forEach(function(i){n&&i.y+(i.y1-i.y0)>e&&(i.y=i.y-(i.y+(i.y1-i.y0)-e));var a=t.links.filter(function(t){return b(t.source,r)==b(i,r)}),o=a.length;o>1&&a.sort(function(t,e){if(!t.circular&&!e.circular){if(t.target.column==e.target.column)return t.y1-e.y1;if(!H(t,e))return t.y1-e.y1;if(t.target.column>e.target.column){var r=B(e,t);return t.y1-r}if(e.target.column>t.target.column)return B(t,e)-e.y1}return t.circular&&!e.circular?"top"==t.circularLinkType?-1:1:e.circular&&!t.circular?"top"==e.circularLinkType?1:-1:t.circular&&e.circular?t.circularLinkType===e.circularLinkType&&"top"==t.circularLinkType?t.target.column===e.target.column?t.target.y1-e.target.y1:e.target.column-t.target.column:t.circularLinkType===e.circularLinkType&&"bottom"==t.circularLinkType?t.target.column===e.target.column?e.target.y1-t.target.y1:t.target.column-e.target.column:"top"==t.circularLinkType?-1:1:void 0});var s=i.y0;a.forEach(function(t){t.y0=s+t.width/2,s+=t.width}),a.forEach(function(t,e){if("bottom"==t.circularLinkType){for(var r=e+1,n=0;r<o;r++)n+=a[r].width;t.y0=i.y1-n-t.width/2}})})}function q(t,e,r){t.nodes.forEach(function(e){var n=t.links.filter(function(t){return b(t.target,r)==b(e,r)}),i=n.length;i>1&&n.sort(function(t,e){if(!t.circular&&!e.circular){if(t.source.column==e.source.column)return t.y0-e.y0;if(!H(t,e))return t.y0-e.y0;if(e.source.column<t.source.column){var r=F(e,t);return t.y0-r}if(t.source.column<e.source.column)return F(t,e)-e.y0}return t.circular&&!e.circular?"top"==t.circularLinkType?-1:1:e.circular&&!t.circular?"top"==e.circularLinkType?1:-1:t.circular&&e.circular?t.circularLinkType===e.circularLinkType&&"top"==t.circularLinkType?t.source.column===e.source.column?t.source.y1-e.source.y1:t.source.column-e.source.column:t.circularLinkType===e.circularLinkType&&"bottom"==t.circularLinkType?t.source.column===e.source.column?t.source.y1-e.source.y1:e.source.column-t.source.column:"top"==t.circularLinkType?-1:1:void 0});var a=e.y0;n.forEach(function(t){t.y1=a+t.width/2,a+=t.width}),n.forEach(function(t,r){if("bottom"==t.circularLinkType){for(var a=r+1,o=0;a<i;a++)o+=n[a].width;t.y1=e.y1-o-t.width/2}})})}function H(t,e){return G(t)==G(e)}function G(t){return t.y0-t.y1>0?"up":"down"}function Y(t,e){return b(t.source,e)==b(t.target,e)}t.sankeyCircular=function(){var t,n,a=0,b=0,A=1,S=1,E=24,L=v,P=o,O=m,z=y,I=32,D=2,R=null;function F(){var o={nodes:O.apply(null,arguments),links:z.apply(null,arguments)};!function(t){t.nodes.forEach(function(t,e){t.index=e,t.sourceLinks=[],t.targetLinks=[]});var e=r.map(t.nodes,L);t.links.forEach(function(t,r){t.index=r;var n=t.source,i=t.target;"object"!==("undefined"==typeof n?"undefined":l(n))&&(n=t.source=x(e,n)),"object"!==("undefined"==typeof i?"undefined":l(i))&&(i=t.target=x(e,i)),n.sourceLinks.push(t),i.targetLinks.push(t)})}(o),function(t,e,r){var n=0;if(null===r){for(var a=[],o=0;o<t.links.length;o++){var s=t.links[o],l=s.source.index,c=s.target.index;a[l]||(a[l]=[]),a[c]||(a[c]=[]),-1===a[l].indexOf(c)&&a[l].push(c)}var u=i(a);u.sort(function(t,e){return t.length-e.length});var h={};for(o=0;o<u.length;o++){var f=u[o],p=f.slice(-2);h[p[0]]||(h[p[0]]={}),h[p[0]][p[1]]=!0}t.links.forEach(function(t){var e=t.target.index,r=t.source.index;e===r||h[r]&&h[r][e]?(t.circular=!0,t.circularLinkID=n,n+=1):t.circular=!1})}else t.links.forEach(function(t){t.source[r]<t.target[r]?t.circular=!1:(t.circular=!0,t.circularLinkID=n,n+=1)})}(o,0,R),function(t){t.nodes.forEach(function(t){t.partOfCycle=!1,t.value=Math.max(e.sum(t.sourceLinks,f),e.sum(t.targetLinks,f)),t.sourceLinks.forEach(function(e){e.circular&&(t.partOfCycle=!0,t.circularLinkType=e.circularLinkType)}),t.targetLinks.forEach(function(e){e.circular&&(t.partOfCycle=!0,t.circularLinkType=e.circularLinkType)})})}(o),function(t){var e,r,n;for(e=t.nodes,r=[],n=0;e.length;++n,e=r,r=[])e.forEach(function(t){t.depth=n,t.sourceLinks.forEach(function(t){r.indexOf(t.target)<0&&!t.circular&&r.push(t.target)})});for(e=t.nodes,r=[],n=0;e.length;++n,e=r,r=[])e.forEach(function(t){t.height=n,t.targetLinks.forEach(function(t){r.indexOf(t.source)<0&&!t.circular&&r.push(t.source)})});t.nodes.forEach(function(t){t.column=Math.floor(P.call(null,t,n))})}(o),T(o,L),function(i,o,s){var l=r.nest().key(function(t){return t.column}).sortKeys(e.ascending).entries(i.nodes).map(function(t){return t.values});(function(r){if(n){var o=1/0;l.forEach(function(t){var e=S*n/(t.length+1);o=e<o?e:o}),t=o}var s=e.min(l,function(r){return(S-b-(r.length-1)*t)/e.sum(r,f)});s*=k,i.links.forEach(function(t){t.width=t.value*s});var c=function(t){var r=0,n=0,i=0,a=0,o=e.max(t.nodes,function(t){return t.column});return t.links.forEach(function(t){t.circular&&("top"==t.circularLinkType?r+=t.width:n+=t.width,0==t.target.column&&(a+=t.width),t.source.column==o&&(i+=t.width))}),{top:r=r>0?r+_+w:r,bottom:n=n>0?n+_+w:n,left:a=a>0?a+_+w:a,right:i=i>0?i+_+w:i}}(i),u=function(t,r){var n=e.max(t.nodes,function(t){return t.column}),i=A-a,o=S-b,s=i+r.right+r.left,l=o+r.top+r.bottom,c=i/s,u=o/l;return a=a*c+r.left,A=0==r.right?A:A*c,b=b*u+r.top,S*=u,t.nodes.forEach(function(t){t.x0=a+t.column*((A-a-E)/n),t.x1=t.x0+E}),u}(i,c);s*=u,i.links.forEach(function(t){t.width=t.value*s}),l.forEach(function(t){var e=t.length;t.forEach(function(t,n){t.depth==l.length-1&&1==e?(t.y0=S/2-t.value*s,t.y1=t.y0+t.value*s):0==t.depth&&1==e?(t.y0=S/2-t.value*s,t.y1=t.y0+t.value*s):t.partOfCycle?0==M(t,r)?(t.y0=S/2+n,t.y1=t.y0+t.value*s):"top"==t.circularLinkType?(t.y0=b+n,t.y1=t.y0+t.value*s):(t.y0=S-t.value*s-n,t.y1=t.y0+t.value*s):0==c.top||0==c.bottom?(t.y0=(S-b)/e*n,t.y1=t.y0+t.value*s):(t.y0=(S-b)/2-e/2+n,t.y1=t.y0+t.value*s)})})})(s),m();for(var c=1,u=o;u>0;--u)v(c*=.99,s),m();function v(t,r){var n=l.length;l.forEach(function(i){var a=i.length,o=i[0].depth;i.forEach(function(i){var s;if(i.sourceLinks.length||i.targetLinks.length)if(i.partOfCycle&&M(i,r)>0);else if(0==o&&1==a)s=i.y1-i.y0,i.y0=S/2-s/2,i.y1=S/2+s/2;else if(o==n-1&&1==a)s=i.y1-i.y0,i.y0=S/2-s/2,i.y1=S/2+s/2;else{var l=e.mean(i.sourceLinks,g),c=e.mean(i.targetLinks,d),u=((l&&c?(l+c)/2:l||c)-p(i))*t;i.y0+=u,i.y1+=u}})})}function m(){l.forEach(function(e){var r,n,i,a=b,o=e.length;for(e.sort(h),i=0;i<o;++i)r=e[i],(n=a-r.y0)>0&&(r.y0+=n,r.y1+=n),a=r.y1+t;if((n=a-t-S)>0)for(a=r.y0-=n,r.y1-=n,i=o-2;i>=0;--i)r=e[i],(n=r.y1+t-a)>0&&(r.y0-=n,r.y1-=n),a=r.y0})}}(o,I,L),B(o);for(var s=0;s<4;s++)V(o,S,L),q(o,0,L),N(o,b,S,L),V(o,S,L),q(o,0,L);return function(t,r,n){var i=t.nodes,a=t.links,o=!1,s=!1;if(a.forEach(function(t){"top"==t.circularLinkType?o=!0:"bottom"==t.circularLinkType&&(s=!0)}),0==o||0==s){var l=e.min(i,function(t){return t.y0}),c=e.max(i,function(t){return t.y1}),u=c-l,h=n-r,f=h/u;i.forEach(function(t){var e=(t.y1-t.y0)*f;t.y0=(t.y0-l)*f,t.y1=t.y0+e}),a.forEach(function(t){t.y0=(t.y0-l)*f,t.y1=(t.y1-l)*f,t.width=t.width*f})}}(o,b,S),C(o,D,S,L),o}function B(t){t.nodes.forEach(function(t){t.sourceLinks.sort(u),t.targetLinks.sort(c)}),t.nodes.forEach(function(t){var e=t.y0,r=e,n=t.y1,i=n;t.sourceLinks.forEach(function(t){t.circular?(t.y0=n-t.width/2,n-=t.width):(t.y0=e+t.width/2,e+=t.width)}),t.targetLinks.forEach(function(t){t.circular?(t.y1=i-t.width/2,i-=t.width):(t.y1=r+t.width/2,r+=t.width)})})}return F.nodeId=function(t){return arguments.length?(L="function"==typeof t?t:s(t),F):L},F.nodeAlign=function(t){return arguments.length?(P="function"==typeof t?t:s(t),F):P},F.nodeWidth=function(t){return arguments.length?(E=+t,F):E},F.nodePadding=function(e){return arguments.length?(t=+e,F):t},F.nodes=function(t){return arguments.length?(O="function"==typeof t?t:s(t),F):O},F.links=function(t){return arguments.length?(z="function"==typeof t?t:s(t),F):z},F.size=function(t){return arguments.length?(a=b=0,A=+t[0],S=+t[1],F):[A-a,S-b]},F.extent=function(t){return arguments.length?(a=+t[0][0],A=+t[1][0],b=+t[0][1],S=+t[1][1],F):[[a,b],[A,S]]},F.iterations=function(t){return arguments.length?(I=+t,F):I},F.circularLinkGap=function(t){return arguments.length?(D=+t,F):D},F.nodePaddingRatio=function(t){return arguments.length?(n=+t,F):n},F.sortNodes=function(t){return arguments.length?(R=t,F):R},F.update=function(t){return T(t,L),B(t),t.links.forEach(function(t){t.circular&&(t.circularLinkType=t.y0+t.y1<S?"top":"bottom",t.source.circularLinkType=t.circularLinkType,t.target.circularLinkType=t.circularLinkType)}),V(t,S,L,!1),q(t,0,L),C(t,D,S,L),t},F},t.sankeyCenter=function(t){return t.targetLinks.length?t.depth:t.sourceLinks.length?e.min(t.sourceLinks,a)-1:0},t.sankeyLeft=function(t){return t.depth},t.sankeyRight=function(t,e){return e-1-t.height},t.sankeyJustify=o,Object.defineProperty(t,"__esModule",{value:!0})},"object"==typeof r&&"undefined"!=typeof e?i(r,t("d3-array"),t("d3-collection"),t("d3-shape"),t("elementary-circuits-directed-graph")):i(n.d3=n.d3||{},n.d3,n.d3,n.d3,null)},{"d3-array":152,"d3-collection":153,"d3-shape":161,"elementary-circuits-directed-graph":172}],54:[function(t,e,r){var n,i;n=this,i=function(t,e,r,n){"use strict";function i(t){return t.target.depth}function a(t,e){return t.sourceLinks.length?t.depth:e-1}function o(t){return function(){return t}}function s(t,e){return c(t.source,e.source)||t.index-e.index}function l(t,e){return c(t.target,e.target)||t.index-e.index}function c(t,e){return t.y0-e.y0}function u(t){return t.value}function h(t){return(t.y0+t.y1)/2}function f(t){return h(t.source)*t.value}function p(t){return h(t.target)*t.value}function d(t){return t.index}function g(t){return t.nodes}function v(t){return t.links}function m(t,e){var r=t.get(e);if(!r)throw new Error("missing: "+e);return r}function y(t){return[t.source.x1,t.y0]}function x(t){return[t.target.x0,t.y1]}t.sankey=function(){var t=0,n=0,i=1,y=1,x=24,b=8,_=d,w=a,k=g,T=v,A=32,M=2/3;function S(){var a={nodes:k.apply(null,arguments),links:T.apply(null,arguments)};return function(t){t.nodes.forEach(function(t,e){t.index=e,t.sourceLinks=[],t.targetLinks=[]});var e=r.map(t.nodes,_);t.links.forEach(function(t,r){t.index=r;var n=t.source,i=t.target;"object"!=typeof n&&(n=t.source=m(e,n)),"object"!=typeof i&&(i=t.target=m(e,i)),n.sourceLinks.push(t),i.targetLinks.push(t)})}(a),function(t){t.nodes.forEach(function(t){t.value=Math.max(e.sum(t.sourceLinks,u),e.sum(t.targetLinks,u))})}(a),function(e){var r,n,a;for(r=e.nodes,n=[],a=0;r.length;++a,r=n,n=[])r.forEach(function(t){t.depth=a,t.sourceLinks.forEach(function(t){n.indexOf(t.target)<0&&n.push(t.target)})});for(r=e.nodes,n=[],a=0;r.length;++a,r=n,n=[])r.forEach(function(t){t.height=a,t.targetLinks.forEach(function(t){n.indexOf(t.source)<0&&n.push(t.source)})});var o=(i-t-x)/(a-1);e.nodes.forEach(function(e){e.x1=(e.x0=t+Math.max(0,Math.min(a-1,Math.floor(w.call(null,e,a))))*o)+x})}(a),function(t){var i=r.nest().key(function(t){return t.x0}).sortKeys(e.ascending).entries(t.nodes).map(function(t){return t.values});(function(){var r=e.max(i,function(t){return t.length}),a=M*(y-n)/(r-1);b>a&&(b=a);var o=e.min(i,function(t){return(y-n-(t.length-1)*b)/e.sum(t,u)});i.forEach(function(t){t.forEach(function(t,e){t.y1=(t.y0=e)+t.value*o})}),t.links.forEach(function(t){t.width=t.value*o})})(),d();for(var a=1,o=A;o>0;--o)l(a*=.99),d(),s(a),d();function s(t){i.forEach(function(r){r.forEach(function(r){if(r.targetLinks.length){var n=(e.sum(r.targetLinks,f)/e.sum(r.targetLinks,u)-h(r))*t;r.y0+=n,r.y1+=n}})})}function l(t){i.slice().reverse().forEach(function(r){r.forEach(function(r){if(r.sourceLinks.length){var n=(e.sum(r.sourceLinks,p)/e.sum(r.sourceLinks,u)-h(r))*t;r.y0+=n,r.y1+=n}})})}function d(){i.forEach(function(t){var e,r,i,a=n,o=t.length;for(t.sort(c),i=0;i<o;++i)e=t[i],(r=a-e.y0)>0&&(e.y0+=r,e.y1+=r),a=e.y1+b;if((r=a-b-y)>0)for(a=e.y0-=r,e.y1-=r,i=o-2;i>=0;--i)e=t[i],(r=e.y1+b-a)>0&&(e.y0-=r,e.y1-=r),a=e.y0})}}(a),E(a),a}function E(t){t.nodes.forEach(function(t){t.sourceLinks.sort(l),t.targetLinks.sort(s)}),t.nodes.forEach(function(t){var e=t.y0,r=e;t.sourceLinks.forEach(function(t){t.y0=e+t.width/2,e+=t.width}),t.targetLinks.forEach(function(t){t.y1=r+t.width/2,r+=t.width})})}return S.update=function(t){return E(t),t},S.nodeId=function(t){return arguments.length?(_="function"==typeof t?t:o(t),S):_},S.nodeAlign=function(t){return arguments.length?(w="function"==typeof t?t:o(t),S):w},S.nodeWidth=function(t){return arguments.length?(x=+t,S):x},S.nodePadding=function(t){return arguments.length?(b=+t,S):b},S.nodes=function(t){return arguments.length?(k="function"==typeof t?t:o(t),S):k},S.links=function(t){return arguments.length?(T="function"==typeof t?t:o(t),S):T},S.size=function(e){return arguments.length?(t=n=0,i=+e[0],y=+e[1],S):[i-t,y-n]},S.extent=function(e){return arguments.length?(t=+e[0][0],i=+e[1][0],n=+e[0][1],y=+e[1][1],S):[[t,n],[i,y]]},S.iterations=function(t){return arguments.length?(A=+t,S):A},S},t.sankeyCenter=function(t){return t.targetLinks.length?t.depth:t.sourceLinks.length?e.min(t.sourceLinks,i)-1:0},t.sankeyLeft=function(t){return t.depth},t.sankeyRight=function(t,e){return e-1-t.height},t.sankeyJustify=a,t.sankeyLinkHorizontal=function(){return n.linkHorizontal().source(y).target(x)},Object.defineProperty(t,"__esModule",{value:!0})},"object"==typeof r&&"undefined"!=typeof e?i(r,t("d3-array"),t("d3-collection"),t("d3-shape")):i(n.d3=n.d3||{},n.d3,n.d3,n.d3)},{"d3-array":152,"d3-collection":153,"d3-shape":161}],55:[function(t,e,r){"use strict";Object.defineProperty(r,"__esModule",{value:!0});var n=t("@turf/meta"),i=6378137;function a(t){var e=0;if(t&&t.length>0){e+=Math.abs(o(t[0]));for(var r=1;r<t.length;r++)e-=Math.abs(o(t[r]))}return e}function o(t){var e,r,n,a,o,l,c=0,u=t.length;if(u>2){for(l=0;l<u;l++)l===u-2?(n=u-2,a=u-1,o=0):l===u-1?(n=u-1,a=0,o=1):(n=l,a=l+1,o=l+2),e=t[n],r=t[a],c+=(s(t[o][0])-s(e[0]))*Math.sin(s(r[1]));c=c*i*i/2}return c}function s(t){return t*Math.PI/180}r.default=function(t){return n.geomReduce(t,function(t,e){return t+function(t){var e,r=0;switch(t.type){case"Polygon":return a(t.coordinates);case"MultiPolygon":for(e=0;e<t.coordinates.length;e++)r+=a(t.coordinates[e]);return r;case"Point":case"MultiPoint":case"LineString":case"MultiLineString":return 0}return 0}(e)},0)}},{"@turf/meta":58}],56:[function(t,e,r){"use strict";Object.defineProperty(r,"__esModule",{value:!0});var n=t("@turf/meta"),i=t("@turf/helpers");r.default=function(t,e){void 0===e&&(e={});var r=0,a=0,o=0;return n.coordEach(t,function(t){r+=t[0],a+=t[1],o++}),i.point([r/o,a/o],e.properties)}},{"@turf/helpers":57,"@turf/meta":58}],57:[function(t,e,r){"use strict";function n(t,e,r){void 0===r&&(r={});var n={type:"Feature"};return(0===r.id||r.id)&&(n.id=r.id),r.bbox&&(n.bbox=r.bbox),n.properties=e||{},n.geometry=t,n}function i(t,e,r){return void 0===r&&(r={}),n({type:"Point",coordinates:t},e,r)}function a(t,e,r){void 0===r&&(r={});for(var i=0,a=t;i<a.length;i++){var o=a[i];if(o.length<4)throw new Error("Each LinearRing of a Polygon must have 4 or more Positions.");for(var s=0;s<o[o.length-1].length;s++)if(o[o.length-1][s]!==o[0][s])throw new Error("First and last Position are not equivalent.")}return n({type:"Polygon",coordinates:t},e,r)}function o(t,e,r){if(void 0===r&&(r={}),t.length<2)throw new Error("coordinates must be an array of two or more positions");return n({type:"LineString",coordinates:t},e,r)}function s(t,e){void 0===e&&(e={});var r={type:"FeatureCollection"};return e.id&&(r.id=e.id),e.bbox&&(r.bbox=e.bbox),r.features=t,r}function l(t,e,r){return void 0===r&&(r={}),n({type:"MultiLineString",coordinates:t},e,r)}function c(t,e,r){return void 0===r&&(r={}),n({type:"MultiPoint",coordinates:t},e,r)}function u(t,e,r){return void 0===r&&(r={}),n({type:"MultiPolygon",coordinates:t},e,r)}function h(t,e){void 0===e&&(e="kilometers");var n=r.factors[e];if(!n)throw new Error(e+" units is invalid");return t*n}function f(t,e){void 0===e&&(e="kilometers");var n=r.factors[e];if(!n)throw new Error(e+" units is invalid");return t/n}function p(t){return 180*(t%(2*Math.PI))/Math.PI}function d(t){return!isNaN(t)&&null!==t&&!Array.isArray(t)&&!/^\s*$/.test(t)}Object.defineProperty(r,"__esModule",{value:!0}),r.earthRadius=6371008.8,r.factors={centimeters:100*r.earthRadius,centimetres:100*r.earthRadius,degrees:r.earthRadius/111325,feet:3.28084*r.earthRadius,inches:39.37*r.earthRadius,kilometers:r.earthRadius/1e3,kilometres:r.earthRadius/1e3,meters:r.earthRadius,metres:r.earthRadius,miles:r.earthRadius/1609.344,millimeters:1e3*r.earthRadius,millimetres:1e3*r.earthRadius,nauticalmiles:r.earthRadius/1852,radians:1,yards:r.earthRadius/1.0936},r.unitsFactors={centimeters:100,centimetres:100,degrees:1/111325,feet:3.28084,inches:39.37,kilometers:.001,kilometres:.001,meters:1,metres:1,miles:1/1609.344,millimeters:1e3,millimetres:1e3,nauticalmiles:1/1852,radians:1/r.earthRadius,yards:1/1.0936},r.areaFactors={acres:247105e-9,centimeters:1e4,centimetres:1e4,feet:10.763910417,inches:1550.003100006,kilometers:1e-6,kilometres:1e-6,meters:1,metres:1,miles:3.86e-7,millimeters:1e6,millimetres:1e6,yards:1.195990046},r.feature=n,r.geometry=function(t,e,r){switch(void 0===r&&(r={}),t){case"Point":return i(e).geometry;case"LineString":return o(e).geometry;case"Polygon":return a(e).geometry;case"MultiPoint":return c(e).geometry;case"MultiLineString":return l(e).geometry;case"MultiPolygon":return u(e).geometry;default:throw new Error(t+" is invalid")}},r.point=i,r.points=function(t,e,r){return void 0===r&&(r={}),s(t.map(function(t){return i(t,e)}),r)},r.polygon=a,r.polygons=function(t,e,r){return void 0===r&&(r={}),s(t.map(function(t){return a(t,e)}),r)},r.lineString=o,r.lineStrings=function(t,e,r){return void 0===r&&(r={}),s(t.map(function(t){return o(t,e)}),r)},r.featureCollection=s,r.multiLineString=l,r.multiPoint=c,r.multiPolygon=u,r.geometryCollection=function(t,e,r){return void 0===r&&(r={}),n({type:"GeometryCollection",geometries:t},e,r)},r.round=function(t,e){if(void 0===e&&(e=0),e&&!(e>=0))throw new Error("precision must be a positive number");var r=Math.pow(10,e||0);return Math.round(t*r)/r},r.radiansToLength=h,r.lengthToRadians=f,r.lengthToDegrees=function(t,e){return p(f(t,e))},r.bearingToAzimuth=function(t){var e=t%360;return e<0&&(e+=360),e},r.radiansToDegrees=p,r.degreesToRadians=function(t){return t%360*Math.PI/180},r.convertLength=function(t,e,r){if(void 0===e&&(e="kilometers"),void 0===r&&(r="kilometers"),!(t>=0))throw new Error("length must be a positive number");return h(f(t,e),r)},r.convertArea=function(t,e,n){if(void 0===e&&(e="meters"),void 0===n&&(n="kilometers"),!(t>=0))throw new Error("area must be a positive number");var i=r.areaFactors[e];if(!i)throw new Error("invalid original units");var a=r.areaFactors[n];if(!a)throw new Error("invalid final units");return t/i*a},r.isNumber=d,r.isObject=function(t){return!!t&&t.constructor===Object},r.validateBBox=function(t){if(!t)throw new Error("bbox is required");if(!Array.isArray(t))throw new Error("bbox must be an Array");if(4!==t.length&&6!==t.length)throw new Error("bbox must be an Array of 4 or 6 numbers");t.forEach(function(t){if(!d(t))throw new Error("bbox must only contain numbers")})},r.validateId=function(t){if(!t)throw new Error("id is required");if(-1===["string","number"].indexOf(typeof t))throw new Error("id must be a number or a string")},r.radians2degrees=function(){throw new Error("method has been renamed to `radiansToDegrees`")},r.degrees2radians=function(){throw new Error("method has been renamed to `degreesToRadians`")},r.distanceToDegrees=function(){throw new Error("method has been renamed to `lengthToDegrees`")},r.distanceToRadians=function(){throw new Error("method has been renamed to `lengthToRadians`")},r.radiansToDistance=function(){throw new Error("method has been renamed to `radiansToLength`")},r.bearingToAngle=function(){throw new Error("method has been renamed to `bearingToAzimuth`")},r.convertDistance=function(){throw new Error("method has been renamed to `convertLength`")}},{}],58:[function(t,e,r){"use strict";Object.defineProperty(r,"__esModule",{value:!0});var n=t("@turf/helpers");function i(t,e,r){if(null!==t)for(var n,a,o,s,l,c,u,h,f=0,p=0,d=t.type,g="FeatureCollection"===d,v="Feature"===d,m=g?t.features.length:1,y=0;y<m;y++){l=(h=!!(u=g?t.features[y].geometry:v?t.geometry:t)&&"GeometryCollection"===u.type)?u.geometries.length:1;for(var x=0;x<l;x++){var b=0,_=0;if(null!==(s=h?u.geometries[x]:u)){c=s.coordinates;var w=s.type;switch(f=!r||"Polygon"!==w&&"MultiPolygon"!==w?0:1,w){case null:break;case"Point":if(!1===e(c,p,y,b,_))return!1;p++,b++;break;case"LineString":case"MultiPoint":for(n=0;n<c.length;n++){if(!1===e(c[n],p,y,b,_))return!1;p++,"MultiPoint"===w&&b++}"LineString"===w&&b++;break;case"Polygon":case"MultiLineString":for(n=0;n<c.length;n++){for(a=0;a<c[n].length-f;a++){if(!1===e(c[n][a],p,y,b,_))return!1;p++}"MultiLineString"===w&&b++,"Polygon"===w&&_++}"Polygon"===w&&b++;break;case"MultiPolygon":for(n=0;n<c.length;n++){for(_=0,a=0;a<c[n].length;a++){for(o=0;o<c[n][a].length-f;o++){if(!1===e(c[n][a][o],p,y,b,_))return!1;p++}_++}b++}break;case"GeometryCollection":for(n=0;n<s.geometries.length;n++)if(!1===i(s.geometries[n],e,r))return!1;break;default:throw new Error("Unknown Geometry Type")}}}}}function a(t,e){var r;switch(t.type){case"FeatureCollection":for(r=0;r<t.features.length&&!1!==e(t.features[r].properties,r);r++);break;case"Feature":e(t.properties,0)}}function o(t,e){if("Feature"===t.type)e(t,0);else if("FeatureCollection"===t.type)for(var r=0;r<t.features.length&&!1!==e(t.features[r],r);r++);}function s(t,e){var r,n,i,a,o,s,l,c,u,h,f=0,p="FeatureCollection"===t.type,d="Feature"===t.type,g=p?t.features.length:1;for(r=0;r<g;r++){for(s=p?t.features[r].geometry:d?t.geometry:t,c=p?t.features[r].properties:d?t.properties:{},u=p?t.features[r].bbox:d?t.bbox:void 0,h=p?t.features[r].id:d?t.id:void 0,o=(l=!!s&&"GeometryCollection"===s.type)?s.geometries.length:1,i=0;i<o;i++)if(null!==(a=l?s.geometries[i]:s))switch(a.type){case"Point":case"LineString":case"MultiPoint":case"Polygon":case"MultiLineString":case"MultiPolygon":if(!1===e(a,f,c,u,h))return!1;break;case"GeometryCollection":for(n=0;n<a.geometries.length;n++)if(!1===e(a.geometries[n],f,c,u,h))return!1;break;default:throw new Error("Unknown Geometry Type")}else if(!1===e(null,f,c,u,h))return!1;f++}}function l(t,e){s(t,function(t,r,i,a,o){var s,l=null===t?null:t.type;switch(l){case null:case"Point":case"LineString":case"Polygon":return!1!==e(n.feature(t,i,{bbox:a,id:o}),r,0)&&void 0}switch(l){case"MultiPoint":s="Point";break;case"MultiLineString":s="LineString";break;case"MultiPolygon":s="Polygon"}for(var c=0;c<t.coordinates.length;c++){var u={type:s,coordinates:t.coordinates[c]};if(!1===e(n.feature(u,i),r,c))return!1}})}function c(t,e){l(t,function(t,r,a){var o=0;if(t.geometry){var s=t.geometry.type;if("Point"!==s&&"MultiPoint"!==s){var l,c=0,u=0,h=0;return!1!==i(t,function(i,s,f,p,d){if(void 0===l||r>c||p>u||d>h)return l=i,c=r,u=p,h=d,void(o=0);var g=n.lineString([l,i],t.properties);if(!1===e(g,r,a,d,o))return!1;o++,l=i})&&void 0}}})}function u(t,e){if(!t)throw new Error("geojson is required");l(t,function(t,r,i){if(null!==t.geometry){var a=t.geometry.type,o=t.geometry.coordinates;switch(a){case"LineString":if(!1===e(t,r,i,0,0))return!1;break;case"Polygon":for(var s=0;s<o.length;s++)if(!1===e(n.lineString(o[s],t.properties),r,i,s))return!1}}})}r.coordEach=i,r.coordReduce=function(t,e,r,n){var a=r;return i(t,function(t,n,i,o,s){a=0===n&&void 0===r?t:e(a,t,n,i,o,s)},n),a},r.propEach=a,r.propReduce=function(t,e,r){var n=r;return a(t,function(t,i){n=0===i&&void 0===r?t:e(n,t,i)}),n},r.featureEach=o,r.featureReduce=function(t,e,r){var n=r;return o(t,function(t,i){n=0===i&&void 0===r?t:e(n,t,i)}),n},r.coordAll=function(t){var e=[];return i(t,function(t){e.push(t)}),e},r.geomEach=s,r.geomReduce=function(t,e,r){var n=r;return s(t,function(t,i,a,o,s){n=0===i&&void 0===r?t:e(n,t,i,a,o,s)}),n},r.flattenEach=l,r.flattenReduce=function(t,e,r){var n=r;return l(t,function(t,i,a){n=0===i&&0===a&&void 0===r?t:e(n,t,i,a)}),n},r.segmentEach=c,r.segmentReduce=function(t,e,r){var n=r,i=!1;return c(t,function(t,a,o,s,l){n=!1===i&&void 0===r?t:e(n,t,a,o,s,l),i=!0}),n},r.lineEach=u,r.lineReduce=function(t,e,r){var n=r;return u(t,function(t,i,a,o){n=0===i&&void 0===r?t:e(n,t,i,a,o)}),n},r.findSegment=function(t,e){if(e=e||{},!n.isObject(e))throw new Error("options is invalid");var r,i=e.featureIndex||0,a=e.multiFeatureIndex||0,o=e.geometryIndex||0,s=e.segmentIndex||0,l=e.properties;switch(t.type){case"FeatureCollection":i<0&&(i=t.features.length+i),l=l||t.features[i].properties,r=t.features[i].geometry;break;case"Feature":l=l||t.properties,r=t.geometry;break;case"Point":case"MultiPoint":return null;case"LineString":case"Polygon":case"MultiLineString":case"MultiPolygon":r=t;break;default:throw new Error("geojson is invalid")}if(null===r)return null;var c=r.coordinates;switch(r.type){case"Point":case"MultiPoint":return null;case"LineString":return s<0&&(s=c.length+s-1),n.lineString([c[s],c[s+1]],l,e);case"Polygon":return o<0&&(o=c.length+o),s<0&&(s=c[o].length+s-1),n.lineString([c[o][s],c[o][s+1]],l,e);case"MultiLineString":return a<0&&(a=c.length+a),s<0&&(s=c[a].length+s-1),n.lineString([c[a][s],c[a][s+1]],l,e);case"MultiPolygon":return a<0&&(a=c.length+a),o<0&&(o=c[a].length+o),s<0&&(s=c[a][o].length-s-1),n.lineString([c[a][o][s],c[a][o][s+1]],l,e)}throw new Error("geojson is invalid")},r.findPoint=function(t,e){if(e=e||{},!n.isObject(e))throw new Error("options is invalid");var r,i=e.featureIndex||0,a=e.multiFeatureIndex||0,o=e.geometryIndex||0,s=e.coordIndex||0,l=e.properties;switch(t.type){case"FeatureCollection":i<0&&(i=t.features.length+i),l=l||t.features[i].properties,r=t.features[i].geometry;break;case"Feature":l=l||t.properties,r=t.geometry;break;case"Point":case"MultiPoint":return null;case"LineString":case"Polygon":case"MultiLineString":case"MultiPolygon":r=t;break;default:throw new Error("geojson is invalid")}if(null===r)return null;var c=r.coordinates;switch(r.type){case"Point":return n.point(c,l,e);case"MultiPoint":return a<0&&(a=c.length+a),n.point(c[a],l,e);case"LineString":return s<0&&(s=c.length+s),n.point(c[s],l,e);case"Polygon":return o<0&&(o=c.length+o),s<0&&(s=c[o].length+s),n.point(c[o][s],l,e);case"MultiLineString":return a<0&&(a=c.length+a),s<0&&(s=c[a].length+s),n.point(c[a][s],l,e);case"MultiPolygon":return a<0&&(a=c.length+a),o<0&&(o=c[a].length+o),s<0&&(s=c[a][o].length-s),n.point(c[a][o][s],l,e)}throw new Error("geojson is invalid")}},{"@turf/helpers":57}],59:[function(t,e,r){"use strict";var n="undefined"==typeof WeakMap?t("weak-map"):WeakMap,i=t("gl-buffer"),a=t("gl-vao"),o=new n;e.exports=function(t){var e=o.get(t),r=e&&(e._triangleBuffer.handle||e._triangleBuffer.buffer);if(!r||!t.isBuffer(r)){var n=i(t,new Float32Array([-1,-1,-1,4,4,-1]));(e=a(t,[{buffer:n,type:t.FLOAT,size:2}]))._triangleBuffer=n,o.set(t,e)}e.bind(),t.drawArrays(t.TRIANGLES,0,3),e.unbind()}},{"gl-buffer":241,"gl-vao":327,"weak-map":552}],60:[function(t,e,r){e.exports=function(t){var e=0,r=0,n=0,i=0;return t.map(function(t){var a=(t=t.slice())[0],o=a.toUpperCase();if(a!=o)switch(t[0]=o,a){case"a":t[6]+=n,t[7]+=i;break;case"v":t[1]+=i;break;case"h":t[1]+=n;break;default:for(var s=1;s<t.length;)t[s++]+=n,t[s++]+=i}switch(o){case"Z":n=e,i=r;break;case"H":n=t[1];break;case"V":i=t[1];break;case"M":n=e=t[1],i=r=t[2];break;default:n=t[t.length-2],i=t[t.length-1]}return t})}},{}],61:[function(t,e,r){var n=t("pad-left");e.exports=function(t,e,r){e="number"==typeof e?e:1,r=r||": ";var i=t.split(/\r?\n/),a=String(i.length+e-1).length;return i.map(function(t,i){var o=i+e,s=String(o).length,l=n(o,a-s);return l+r+t}).join("\n")}},{"pad-left":457}],62:[function(t,e,r){"use strict";e.exports=function(t){var e=t.length;if(0===e)return[];if(1===e)return[0];for(var r=t[0].length,n=[t[0]],a=[0],o=1;o<e;++o)if(n.push(t[o]),i(n,r)){if(a.push(o),a.length===r+1)return a}else n.pop();return a};var n=t("robust-orientation");function i(t,e){for(var r=new Array(e+1),i=0;i<t.length;++i)r[i]=t[i];for(i=0;i<=t.length;++i){for(var a=t.length;a<=e;++a){for(var o=new Array(e),s=0;s<e;++s)o[s]=Math.pow(a+1-i,s);r[a]=o}if(n.apply(void 0,r))return!0}return!1}},{"robust-orientation":510}],63:[function(t,e,r){"use strict";e.exports=function(t,e){return n(e).filter(function(r){for(var n=new Array(r.length),a=0;a<r.length;++a)n[a]=e[r[a]];return i(n)*t<1})};var n=t("delaunay-triangulate"),i=t("circumradius")},{circumradius:114,"delaunay-triangulate":165}],64:[function(t,e,r){e.exports=function(t,e){return i(n(t,e))};var n=t("alpha-complex"),i=t("simplicial-complex-boundary")},{"alpha-complex":63,"simplicial-complex-boundary":517}],65:[function(t,e,r){"use strict";e.exports=function(t,e){if(!t||null==t.length)throw Error("Argument should be an array");e=null==e?1:Math.floor(e);for(var r=Array(2*e),n=0;n<e;n++){for(var i=-1/0,a=1/0,o=n,s=t.length;o<s;o+=e)t[o]>i&&(i=t[o]),t[o]<a&&(a=t[o]);r[n]=a,r[e+n]=i}return r}},{}],66:[function(t,e,r){"use strict";var n=t("array-bounds");e.exports=function(t,e,r){if(!t||null==t.length)throw Error("Argument should be an array");null==e&&(e=1);null==r&&(r=n(t,e));for(var i=0;i<e;i++){var a=r[e+i],o=r[i],s=i,l=t.length;if(a===1/0&&o===-1/0)for(s=i;s<l;s+=e)t[s]=t[s]===a?1:t[s]===o?0:.5;else if(a===1/0)for(s=i;s<l;s+=e)t[s]=t[s]===a?1:0;else if(o===-1/0)for(s=i;s<l;s+=e)t[s]=t[s]===o?0:1;else{var c=a-o;for(s=i;s<l;s+=e)t[s]=0===c?.5:(t[s]-o)/c}}return t}},{"array-bounds":65}],67:[function(t,e,r){e.exports=function(t,e){var r="number"==typeof t,n="number"==typeof e;r&&!n?(e=t,t=0):r||n||(t=0,e=0);var i=(e|=0)-(t|=0);if(i<0)throw new Error("array length must be positive");for(var a=new Array(i),o=0,s=t;o<i;o++,s++)a[o]=s;return a}},{}],68:[function(t,e,r){(function(r){"use strict";var n=t("object-assign");function i(t,e){if(t===e)return 0;for(var r=t.length,n=e.length,i=0,a=Math.min(r,n);i<a;++i)if(t[i]!==e[i]){r=t[i],n=e[i];break}return r<n?-1:n<r?1:0}function a(t){return r.Buffer&&"function"==typeof r.Buffer.isBuffer?r.Buffer.isBuffer(t):!(null==t||!t._isBuffer)}var o=t("util/"),s=Object.prototype.hasOwnProperty,l=Array.prototype.slice,c="foo"===function(){}.name;function u(t){return Object.prototype.toString.call(t)}function h(t){return!a(t)&&("function"==typeof r.ArrayBuffer&&("function"==typeof ArrayBuffer.isView?ArrayBuffer.isView(t):!!t&&(t instanceof DataView||!!(t.buffer&&t.buffer instanceof ArrayBuffer))))}var f=e.exports=y,p=/\s*function\s+([^\(\s]*)\s*/;function d(t){if(o.isFunction(t)){if(c)return t.name;var e=t.toString().match(p);return e&&e[1]}}function g(t,e){return"string"==typeof t?t.length<e?t:t.slice(0,e):t}function v(t){if(c||!o.isFunction(t))return o.inspect(t);var e=d(t);return"[Function"+(e?": "+e:"")+"]"}function m(t,e,r,n,i){throw new f.AssertionError({message:r,actual:t,expected:e,operator:n,stackStartFunction:i})}function y(t,e){t||m(t,!0,e,"==",f.ok)}function x(t,e,r,n){if(t===e)return!0;if(a(t)&&a(e))return 0===i(t,e);if(o.isDate(t)&&o.isDate(e))return t.getTime()===e.getTime();if(o.isRegExp(t)&&o.isRegExp(e))return t.source===e.source&&t.global===e.global&&t.multiline===e.multiline&&t.lastIndex===e.lastIndex&&t.ignoreCase===e.ignoreCase;if(null!==t&&"object"==typeof t||null!==e&&"object"==typeof e){if(h(t)&&h(e)&&u(t)===u(e)&&!(t instanceof Float32Array||t instanceof Float64Array))return 0===i(new Uint8Array(t.buffer),new Uint8Array(e.buffer));if(a(t)!==a(e))return!1;var s=(n=n||{actual:[],expected:[]}).actual.indexOf(t);return-1!==s&&s===n.expected.indexOf(e)||(n.actual.push(t),n.expected.push(e),function(t,e,r,n){if(null==t||null==e)return!1;if(o.isPrimitive(t)||o.isPrimitive(e))return t===e;if(r&&Object.getPrototypeOf(t)!==Object.getPrototypeOf(e))return!1;var i=b(t),a=b(e);if(i&&!a||!i&&a)return!1;if(i)return t=l.call(t),e=l.call(e),x(t,e,r);var s,c,u=k(t),h=k(e);if(u.length!==h.length)return!1;for(u.sort(),h.sort(),c=u.length-1;c>=0;c--)if(u[c]!==h[c])return!1;for(c=u.length-1;c>=0;c--)if(s=u[c],!x(t[s],e[s],r,n))return!1;return!0}(t,e,r,n))}return r?t===e:t==e}function b(t){return"[object Arguments]"==Object.prototype.toString.call(t)}function _(t,e){if(!t||!e)return!1;if("[object RegExp]"==Object.prototype.toString.call(e))return e.test(t);try{if(t instanceof e)return!0}catch(t){}return!Error.isPrototypeOf(e)&&!0===e.call({},t)}function w(t,e,r,n){var i;if("function"!=typeof e)throw new TypeError('"block" argument must be a function');"string"==typeof r&&(n=r,r=null),i=function(t){var e;try{t()}catch(t){e=t}return e}(e),n=(r&&r.name?" ("+r.name+").":".")+(n?" "+n:"."),t&&!i&&m(i,r,"Missing expected exception"+n);var a="string"==typeof n,s=!t&&i&&!r;if((!t&&o.isError(i)&&a&&_(i,r)||s)&&m(i,r,"Got unwanted exception"+n),t&&i&&r&&!_(i,r)||!t&&i)throw i}f.AssertionError=function(t){var e;this.name="AssertionError",this.actual=t.actual,this.expected=t.expected,this.operator=t.operator,t.message?(this.message=t.message,this.generatedMessage=!1):(this.message=g(v((e=this).actual),128)+" "+e.operator+" "+g(v(e.expected),128),this.generatedMessage=!0);var r=t.stackStartFunction||m;if(Error.captureStackTrace)Error.captureStackTrace(this,r);else{var n=new Error;if(n.stack){var i=n.stack,a=d(r),o=i.indexOf("\n"+a);if(o>=0){var s=i.indexOf("\n",o+1);i=i.substring(s+1)}this.stack=i}}},o.inherits(f.AssertionError,Error),f.fail=m,f.ok=y,f.equal=function(t,e,r){t!=e&&m(t,e,r,"==",f.equal)},f.notEqual=function(t,e,r){t==e&&m(t,e,r,"!=",f.notEqual)},f.deepEqual=function(t,e,r){x(t,e,!1)||m(t,e,r,"deepEqual",f.deepEqual)},f.deepStrictEqual=function(t,e,r){x(t,e,!0)||m(t,e,r,"deepStrictEqual",f.deepStrictEqual)},f.notDeepEqual=function(t,e,r){x(t,e,!1)&&m(t,e,r,"notDeepEqual",f.notDeepEqual)},f.notDeepStrictEqual=function t(e,r,n){x(e,r,!0)&&m(e,r,n,"notDeepStrictEqual",t)},f.strictEqual=function(t,e,r){t!==e&&m(t,e,r,"===",f.strictEqual)},f.notStrictEqual=function(t,e,r){t===e&&m(t,e,r,"!==",f.notStrictEqual)},f.throws=function(t,e,r){w(!0,t,e,r)},f.doesNotThrow=function(t,e,r){w(!1,t,e,r)},f.ifError=function(t){if(t)throw t},f.strict=n(function t(e,r){e||m(e,!0,r,"==",t)},f,{equal:f.strictEqual,deepEqual:f.deepStrictEqual,notEqual:f.notStrictEqual,notDeepEqual:f.notDeepStrictEqual}),f.strict.strict=f.strict;var k=Object.keys||function(t){var e=[];for(var r in t)s.call(t,r)&&e.push(r);return e}}).call(this,"undefined"!=typeof global?global:"undefined"!=typeof self?self:"undefined"!=typeof window?window:{})},{"object-assign":454,"util/":71}],69:[function(t,e,r){"function"==typeof Object.create?e.exports=function(t,e){t.super_=e,t.prototype=Object.create(e.prototype,{constructor:{value:t,enumerable:!1,writable:!0,configurable:!0}})}:e.exports=function(t,e){t.super_=e;var r=function(){};r.prototype=e.prototype,t.prototype=new r,t.prototype.constructor=t}},{}],70:[function(t,e,r){e.exports=function(t){return t&&"object"==typeof t&&"function"==typeof t.copy&&"function"==typeof t.fill&&"function"==typeof t.readUInt8}},{}],71:[function(t,e,r){(function(e,n){var i=/%[sdj%]/g;r.format=function(t){if(!m(t)){for(var e=[],r=0;r<arguments.length;r++)e.push(s(arguments[r]));return e.join(" ")}r=1;for(var n=arguments,a=n.length,o=String(t).replace(i,function(t){if("%%"===t)return"%";if(r>=a)return t;switch(t){case"%s":return String(n[r++]);case"%d":return Number(n[r++]);case"%j":try{return JSON.stringify(n[r++])}catch(t){return"[Circular]"}default:return t}}),l=n[r];r<a;l=n[++r])g(l)||!b(l)?o+=" "+l:o+=" "+s(l);return o},r.deprecate=function(t,i){if(y(n.process))return function(){return r.deprecate(t,i).apply(this,arguments)};if(!0===e.noDeprecation)return t;var a=!1;return function(){if(!a){if(e.throwDeprecation)throw new Error(i);e.traceDeprecation?console.trace(i):console.error(i),a=!0}return t.apply(this,arguments)}};var a,o={};function s(t,e){var n={seen:[],stylize:c};return arguments.length>=3&&(n.depth=arguments[2]),arguments.length>=4&&(n.colors=arguments[3]),d(e)?n.showHidden=e:e&&r._extend(n,e),y(n.showHidden)&&(n.showHidden=!1),y(n.depth)&&(n.depth=2),y(n.colors)&&(n.colors=!1),y(n.customInspect)&&(n.customInspect=!0),n.colors&&(n.stylize=l),u(n,t,n.depth)}function l(t,e){var r=s.styles[e];return r?"\x1b["+s.colors[r][0]+"m"+t+"\x1b["+s.colors[r][1]+"m":t}function c(t,e){return t}function u(t,e,n){if(t.customInspect&&e&&k(e.inspect)&&e.inspect!==r.inspect&&(!e.constructor||e.constructor.prototype!==e)){var i=e.inspect(n,t);return m(i)||(i=u(t,i,n)),i}var a=function(t,e){if(y(e))return t.stylize("undefined","undefined");if(m(e)){var r="'"+JSON.stringify(e).replace(/^"|"$/g,"").replace(/'/g,"\\'").replace(/\\"/g,'"')+"'";return t.stylize(r,"string")}if(v(e))return t.stylize(""+e,"number");if(d(e))return t.stylize(""+e,"boolean");if(g(e))return t.stylize("null","null")}(t,e);if(a)return a;var o=Object.keys(e),s=function(t){var e={};return t.forEach(function(t,r){e[t]=!0}),e}(o);if(t.showHidden&&(o=Object.getOwnPropertyNames(e)),w(e)&&(o.indexOf("message")>=0||o.indexOf("description")>=0))return h(e);if(0===o.length){if(k(e)){var l=e.name?": "+e.name:"";return t.stylize("[Function"+l+"]","special")}if(x(e))return t.stylize(RegExp.prototype.toString.call(e),"regexp");if(_(e))return t.stylize(Date.prototype.toString.call(e),"date");if(w(e))return h(e)}var c,b="",T=!1,A=["{","}"];(p(e)&&(T=!0,A=["[","]"]),k(e))&&(b=" [Function"+(e.name?": "+e.name:"")+"]");return x(e)&&(b=" "+RegExp.prototype.toString.call(e)),_(e)&&(b=" "+Date.prototype.toUTCString.call(e)),w(e)&&(b=" "+h(e)),0!==o.length||T&&0!=e.length?n<0?x(e)?t.stylize(RegExp.prototype.toString.call(e),"regexp"):t.stylize("[Object]","special"):(t.seen.push(e),c=T?function(t,e,r,n,i){for(var a=[],o=0,s=e.length;o<s;++o)S(e,String(o))?a.push(f(t,e,r,n,String(o),!0)):a.push("");return i.forEach(function(i){i.match(/^\d+$/)||a.push(f(t,e,r,n,i,!0))}),a}(t,e,n,s,o):o.map(function(r){return f(t,e,n,s,r,T)}),t.seen.pop(),function(t,e,r){if(t.reduce(function(t,e){return 0,e.indexOf("\n")>=0&&0,t+e.replace(/\u001b\[\d\d?m/g,"").length+1},0)>60)return r[0]+(""===e?"":e+"\n ")+" "+t.join(",\n  ")+" "+r[1];return r[0]+e+" "+t.join(", ")+" "+r[1]}(c,b,A)):A[0]+b+A[1]}function h(t){return"["+Error.prototype.toString.call(t)+"]"}function f(t,e,r,n,i,a){var o,s,l;if((l=Object.getOwnPropertyDescriptor(e,i)||{value:e[i]}).get?s=l.set?t.stylize("[Getter/Setter]","special"):t.stylize("[Getter]","special"):l.set&&(s=t.stylize("[Setter]","special")),S(n,i)||(o="["+i+"]"),s||(t.seen.indexOf(l.value)<0?(s=g(r)?u(t,l.value,null):u(t,l.value,r-1)).indexOf("\n")>-1&&(s=a?s.split("\n").map(function(t){return"  "+t}).join("\n").substr(2):"\n"+s.split("\n").map(function(t){return"   "+t}).join("\n")):s=t.stylize("[Circular]","special")),y(o)){if(a&&i.match(/^\d+$/))return s;(o=JSON.stringify(""+i)).match(/^"([a-zA-Z_][a-zA-Z_0-9]*)"$/)?(o=o.substr(1,o.length-2),o=t.stylize(o,"name")):(o=o.replace(/'/g,"\\'").replace(/\\"/g,'"').replace(/(^"|"$)/g,"'"),o=t.stylize(o,"string"))}return o+": "+s}function p(t){return Array.isArray(t)}function d(t){return"boolean"==typeof t}function g(t){return null===t}function v(t){return"number"==typeof t}function m(t){return"string"==typeof t}function y(t){return void 0===t}function x(t){return b(t)&&"[object RegExp]"===T(t)}function b(t){return"object"==typeof t&&null!==t}function _(t){return b(t)&&"[object Date]"===T(t)}function w(t){return b(t)&&("[object Error]"===T(t)||t instanceof Error)}function k(t){return"function"==typeof t}function T(t){return Object.prototype.toString.call(t)}function A(t){return t<10?"0"+t.toString(10):t.toString(10)}r.debuglog=function(t){if(y(a)&&(a=e.env.NODE_DEBUG||""),t=t.toUpperCase(),!o[t])if(new RegExp("\\b"+t+"\\b","i").test(a)){var n=e.pid;o[t]=function(){var e=r.format.apply(r,arguments);console.error("%s %d: %s",t,n,e)}}else o[t]=function(){};return o[t]},r.inspect=s,s.colors={bold:[1,22],italic:[3,23],underline:[4,24],inverse:[7,27],white:[37,39],grey:[90,39],black:[30,39],blue:[34,39],cyan:[36,39],green:[32,39],magenta:[35,39],red:[31,39],yellow:[33,39]},s.styles={special:"cyan",number:"yellow",boolean:"yellow",undefined:"grey",null:"bold",string:"green",date:"magenta",regexp:"red"},r.isArray=p,r.isBoolean=d,r.isNull=g,r.isNullOrUndefined=function(t){return null==t},r.isNumber=v,r.isString=m,r.isSymbol=function(t){return"symbol"==typeof t},r.isUndefined=y,r.isRegExp=x,r.isObject=b,r.isDate=_,r.isError=w,r.isFunction=k,r.isPrimitive=function(t){return null===t||"boolean"==typeof t||"number"==typeof t||"string"==typeof t||"symbol"==typeof t||"undefined"==typeof t},r.isBuffer=t("./support/isBuffer");var M=["Jan","Feb","Mar","Apr","May","Jun","Jul","Aug","Sep","Oct","Nov","Dec"];function S(t,e){return Object.prototype.hasOwnProperty.call(t,e)}r.log=function(){var t,e;console.log("%s - %s",(t=new Date,e=[A(t.getHours()),A(t.getMinutes()),A(t.getSeconds())].join(":"),[t.getDate(),M[t.getMonth()],e].join(" ")),r.format.apply(r,arguments))},r.inherits=t("inherits"),r._extend=function(t,e){if(!e||!b(e))return t;for(var r=Object.keys(e),n=r.length;n--;)t[r[n]]=e[r[n]];return t}}).call(this,t("_process"),"undefined"!=typeof global?global:"undefined"!=typeof self?self:"undefined"!=typeof window?window:{})},{"./support/isBuffer":70,_process:482,inherits:69}],72:[function(t,e,r){e.exports=function(t){return atob(t)}},{}],73:[function(t,e,r){"use strict";e.exports=function(t,e){for(var r=e.length,a=new Array(r+1),o=0;o<r;++o){for(var s=new Array(r+1),l=0;l<=r;++l)s[l]=t[l][o];a[o]=s}a[r]=new Array(r+1);for(var o=0;o<=r;++o)a[r][o]=1;for(var c=new Array(r+1),o=0;o<r;++o)c[o]=e[o];c[r]=1;var u=n(a,c),h=i(u[r+1]);0===h&&(h=1);for(var f=new Array(r+1),o=0;o<=r;++o)f[o]=i(u[o])/h;return f};var n=t("robust-linear-solve");function i(t){for(var e=0,r=0;r<t.length;++r)e+=t[r];return e}},{"robust-linear-solve":509}],74:[function(t,e,r){"use strict";r.byteLength=function(t){var e=c(t),r=e[0],n=e[1];return 3*(r+n)/4-n},r.toByteArray=function(t){for(var e,r=c(t),n=r[0],o=r[1],s=new a(function(t,e,r){return 3*(e+r)/4-r}(0,n,o)),l=0,u=o>0?n-4:n,h=0;h<u;h+=4)e=i[t.charCodeAt(h)]<<18|i[t.charCodeAt(h+1)]<<12|i[t.charCodeAt(h+2)]<<6|i[t.charCodeAt(h+3)],s[l++]=e>>16&255,s[l++]=e>>8&255,s[l++]=255&e;2===o&&(e=i[t.charCodeAt(h)]<<2|i[t.charCodeAt(h+1)]>>4,s[l++]=255&e);1===o&&(e=i[t.charCodeAt(h)]<<10|i[t.charCodeAt(h+1)]<<4|i[t.charCodeAt(h+2)]>>2,s[l++]=e>>8&255,s[l++]=255&e);return s},r.fromByteArray=function(t){for(var e,r=t.length,i=r%3,a=[],o=0,s=r-i;o<s;o+=16383)a.push(u(t,o,o+16383>s?s:o+16383));1===i?(e=t[r-1],a.push(n[e>>2]+n[e<<4&63]+"==")):2===i&&(e=(t[r-2]<<8)+t[r-1],a.push(n[e>>10]+n[e>>4&63]+n[e<<2&63]+"="));return a.join("")};for(var n=[],i=[],a="undefined"!=typeof Uint8Array?Uint8Array:Array,o="ABCDEFGHIJKLMNOPQRSTUVWXYZabcdefghijklmnopqrstuvwxyz0123456789+/",s=0,l=o.length;s<l;++s)n[s]=o[s],i[o.charCodeAt(s)]=s;function c(t){var e=t.length;if(e%4>0)throw new Error("Invalid string. Length must be a multiple of 4");var r=t.indexOf("=");return-1===r&&(r=e),[r,r===e?0:4-r%4]}function u(t,e,r){for(var i,a,o=[],s=e;s<r;s+=3)i=(t[s]<<16&16711680)+(t[s+1]<<8&65280)+(255&t[s+2]),o.push(n[(a=i)>>18&63]+n[a>>12&63]+n[a>>6&63]+n[63&a]);return o.join("")}i["-".charCodeAt(0)]=62,i["_".charCodeAt(0)]=63},{}],75:[function(t,e,r){"use strict";var n=t("./lib/rationalize");e.exports=function(t,e){return n(t[0].mul(e[1]).add(e[0].mul(t[1])),t[1].mul(e[1]))}},{"./lib/rationalize":85}],76:[function(t,e,r){"use strict";e.exports=function(t,e){return t[0].mul(e[1]).cmp(e[0].mul(t[1]))}},{}],77:[function(t,e,r){"use strict";var n=t("./lib/rationalize");e.exports=function(t,e){return n(t[0].mul(e[1]),t[1].mul(e[0]))}},{"./lib/rationalize":85}],78:[function(t,e,r){"use strict";var n=t("./is-rat"),i=t("./lib/is-bn"),a=t("./lib/num-to-bn"),o=t("./lib/str-to-bn"),s=t("./lib/rationalize"),l=t("./div");e.exports=function t(e,r){if(n(e))return r?l(e,t(r)):[e[0].clone(),e[1].clone()];var c=0;var u,h;if(i(e))u=e.clone();else if("string"==typeof e)u=o(e);else{if(0===e)return[a(0),a(1)];if(e===Math.floor(e))u=a(e);else{for(;e!==Math.floor(e);)e*=Math.pow(2,256),c-=256;u=a(e)}}if(n(r))u.mul(r[1]),h=r[0].clone();else if(i(r))h=r.clone();else if("string"==typeof r)h=o(r);else if(r)if(r===Math.floor(r))h=a(r);else{for(;r!==Math.floor(r);)r*=Math.pow(2,256),c+=256;h=a(r)}else h=a(1);c>0?u=u.ushln(c):c<0&&(h=h.ushln(-c));return s(u,h)}},{"./div":77,"./is-rat":79,"./lib/is-bn":83,"./lib/num-to-bn":84,"./lib/rationalize":85,"./lib/str-to-bn":86}],79:[function(t,e,r){"use strict";var n=t("./lib/is-bn");e.exports=function(t){return Array.isArray(t)&&2===t.length&&n(t[0])&&n(t[1])}},{"./lib/is-bn":83}],80:[function(t,e,r){"use strict";var n=t("bn.js");e.exports=function(t){return t.cmp(new n(0))}},{"bn.js":94}],81:[function(t,e,r){"use strict";var n=t("./bn-sign");e.exports=function(t){var e=t.length,r=t.words,i=0;if(1===e)i=r[0];else if(2===e)i=r[0]+67108864*r[1];else for(var a=0;a<e;a++){var o=r[a];i+=o*Math.pow(67108864,a)}return n(t)*i}},{"./bn-sign":80}],82:[function(t,e,r){"use strict";var n=t("double-bits"),i=t("bit-twiddle").countTrailingZeros;e.exports=function(t){var e=i(n.lo(t));if(e<32)return e;var r=i(n.hi(t));if(r>20)return 52;return r+32}},{"bit-twiddle":92,"double-bits":167}],83:[function(t,e,r){"use strict";t("bn.js");e.exports=function(t){return t&&"object"==typeof t&&Boolean(t.words)}},{"bn.js":94}],84:[function(t,e,r){"use strict";var n=t("bn.js"),i=t("double-bits");e.exports=function(t){var e=i.exponent(t);return e<52?new n(t):new n(t*Math.pow(2,52-e)).ushln(e-52)}},{"bn.js":94,"double-bits":167}],85:[function(t,e,r){"use strict";var n=t("./num-to-bn"),i=t("./bn-sign");e.exports=function(t,e){var r=i(t),a=i(e);if(0===r)return[n(0),n(1)];if(0===a)return[n(0),n(0)];a<0&&(t=t.neg(),e=e.neg());var o=t.gcd(e);if(o.cmpn(1))return[t.div(o),e.div(o)];return[t,e]}},{"./bn-sign":80,"./num-to-bn":84}],86:[function(t,e,r){"use strict";var n=t("bn.js");e.exports=function(t){return new n(t)}},{"bn.js":94}],87:[function(t,e,r){"use strict";var n=t("./lib/rationalize");e.exports=function(t,e){return n(t[0].mul(e[0]),t[1].mul(e[1]))}},{"./lib/rationalize":85}],88:[function(t,e,r){"use strict";var n=t("./lib/bn-sign");e.exports=function(t){return n(t[0])*n(t[1])}},{"./lib/bn-sign":80}],89:[function(t,e,r){"use strict";var n=t("./lib/rationalize");e.exports=function(t,e){return n(t[0].mul(e[1]).sub(t[1].mul(e[0])),t[1].mul(e[1]))}},{"./lib/rationalize":85}],90:[function(t,e,r){"use strict";var n=t("./lib/bn-to-num"),i=t("./lib/ctz");e.exports=function(t){var e=t[0],r=t[1];if(0===e.cmpn(0))return 0;var a=e.abs().divmod(r.abs()),o=a.div,s=n(o),l=a.mod,c=e.negative!==r.negative?-1:1;if(0===l.cmpn(0))return c*s;if(s){var u=i(s)+4,h=n(l.ushln(u).divRound(r));return c*(s+h*Math.pow(2,-u))}var f=r.bitLength()-l.bitLength()+53,h=n(l.ushln(f).divRound(r));return f<1023?c*h*Math.pow(2,-f):(h*=Math.pow(2,-1023),c*h*Math.pow(2,1023-f))}},{"./lib/bn-to-num":81,"./lib/ctz":82}],91:[function(t,e,r){"use strict";function n(t,e,r,n,i,a){var o=["function ",t,"(a,l,h,",n.join(","),"){",a?"":"var i=",r?"l-1":"h+1",";while(l<=h){var m=(l+h)>>>1,x=a",i?".get(m)":"[m]"];return a?e.indexOf("c")<0?o.push(";if(x===y){return m}else if(x<=y){"):o.push(";var p=c(x,y);if(p===0){return m}else if(p<=0){"):o.push(";if(",e,"){i=m;"),r?o.push("l=m+1}else{h=m-1}"):o.push("h=m-1}else{l=m+1}"),o.push("}"),a?o.push("return -1};"):o.push("return i};"),o.join("")}function i(t,e,r,i){return new Function([n("A","x"+t+"y",e,["y"],!1,i),n("B","x"+t+"y",e,["y"],!0,i),n("P","c(x,y)"+t+"0",e,["y","c"],!1,i),n("Q","c(x,y)"+t+"0",e,["y","c"],!0,i),"function dispatchBsearch",r,"(a,y,c,l,h){if(a.shape){if(typeof(c)==='function'){return Q(a,(l===undefined)?0:l|0,(h===undefined)?a.shape[0]-1:h|0,y,c)}else{return B(a,(c===undefined)?0:c|0,(l===undefined)?a.shape[0]-1:l|0,y)}}else{if(typeof(c)==='function'){return P(a,(l===undefined)?0:l|0,(h===undefined)?a.length-1:h|0,y,c)}else{return A(a,(c===undefined)?0:c|0,(l===undefined)?a.length-1:l|0,y)}}}return dispatchBsearch",r].join(""))()}e.exports={ge:i(">=",!1,"GE"),gt:i(">",!1,"GT"),lt:i("<",!0,"LT"),le:i("<=",!0,"LE"),eq:i("-",!0,"EQ",!0)}},{}],92:[function(t,e,r){"use strict";function n(t){var e=32;return(t&=-t)&&e--,65535&t&&(e-=16),16711935&t&&(e-=8),252645135&t&&(e-=4),858993459&t&&(e-=2),1431655765&t&&(e-=1),e}r.INT_BITS=32,r.INT_MAX=2147483647,r.INT_MIN=-1<<31,r.sign=function(t){return(t>0)-(t<0)},r.abs=function(t){var e=t>>31;return(t^e)-e},r.min=function(t,e){return e^(t^e)&-(t<e)},r.max=function(t,e){return t^(t^e)&-(t<e)},r.isPow2=function(t){return!(t&t-1||!t)},r.log2=function(t){var e,r;return e=(t>65535)<<4,e|=r=((t>>>=e)>255)<<3,e|=r=((t>>>=r)>15)<<2,(e|=r=((t>>>=r)>3)<<1)|(t>>>=r)>>1},r.log10=function(t){return t>=1e9?9:t>=1e8?8:t>=1e7?7:t>=1e6?6:t>=1e5?5:t>=1e4?4:t>=1e3?3:t>=100?2:t>=10?1:0},r.popCount=function(t){return 16843009*((t=(858993459&(t-=t>>>1&1431655765))+(t>>>2&858993459))+(t>>>4)&252645135)>>>24},r.countTrailingZeros=n,r.nextPow2=function(t){return t+=0===t,--t,t|=t>>>1,t|=t>>>2,t|=t>>>4,t|=t>>>8,(t|=t>>>16)+1},r.prevPow2=function(t){return t|=t>>>1,t|=t>>>2,t|=t>>>4,t|=t>>>8,(t|=t>>>16)-(t>>>1)},r.parity=function(t){return t^=t>>>16,t^=t>>>8,t^=t>>>4,27030>>>(t&=15)&1};var i=new Array(256);!function(t){for(var e=0;e<256;++e){var r=e,n=e,i=7;for(r>>>=1;r;r>>>=1)n<<=1,n|=1&r,--i;t[e]=n<<i&255}}(i),r.reverse=function(t){return i[255&t]<<24|i[t>>>8&255]<<16|i[t>>>16&255]<<8|i[t>>>24&255]},r.interleave2=function(t,e){return(t=1431655765&((t=858993459&((t=252645135&((t=16711935&((t&=65535)|t<<8))|t<<4))|t<<2))|t<<1))|(e=1431655765&((e=858993459&((e=252645135&((e=16711935&((e&=65535)|e<<8))|e<<4))|e<<2))|e<<1))<<1},r.deinterleave2=function(t,e){return(t=65535&((t=16711935&((t=252645135&((t=858993459&((t=t>>>e&1431655765)|t>>>1))|t>>>2))|t>>>4))|t>>>16))<<16>>16},r.interleave3=function(t,e,r){return t=1227133513&((t=3272356035&((t=251719695&((t=4278190335&((t&=1023)|t<<16))|t<<8))|t<<4))|t<<2),(t|=(e=1227133513&((e=3272356035&((e=251719695&((e=4278190335&((e&=1023)|e<<16))|e<<8))|e<<4))|e<<2))<<1)|(r=1227133513&((r=3272356035&((r=251719695&((r=4278190335&((r&=1023)|r<<16))|r<<8))|r<<4))|r<<2))<<2},r.deinterleave3=function(t,e){return(t=1023&((t=4278190335&((t=251719695&((t=3272356035&((t=t>>>e&1227133513)|t>>>2))|t>>>4))|t>>>8))|t>>>16))<<22>>22},r.nextCombination=function(t){var e=t|t-1;return e+1|(~e&-~e)-1>>>n(t)+1}},{}],93:[function(t,e,r){"use strict";var n=t("clamp");e.exports=function(t,e){e||(e={});var r,o,s,l,c,u,h,f,p,d,g,v=null==e.cutoff?.25:e.cutoff,m=null==e.radius?8:e.radius,y=e.channel||0;if(ArrayBuffer.isView(t)||Array.isArray(t)){if(!e.width||!e.height)throw Error("For raw data width and height should be provided by options");r=e.width,o=e.height,l=t,u=e.stride?e.stride:Math.floor(t.length/r/o)}else window.HTMLCanvasElement&&t instanceof window.HTMLCanvasElement?(h=(f=t).getContext("2d"),r=f.width,o=f.height,p=h.getImageData(0,0,r,o),l=p.data,u=4):window.CanvasRenderingContext2D&&t instanceof window.CanvasRenderingContext2D?(f=t.canvas,h=t,r=f.width,o=f.height,p=h.getImageData(0,0,r,o),l=p.data,u=4):window.ImageData&&t instanceof window.ImageData&&(p=t,r=t.width,o=t.height,l=p.data,u=4);if(s=Math.max(r,o),window.Uint8ClampedArray&&l instanceof window.Uint8ClampedArray||window.Uint8Array&&l instanceof window.Uint8Array)for(c=l,l=Array(r*o),d=0,g=c.length;d<g;d++)l[d]=c[d*u+y]/255;else if(1!==u)throw Error("Raw data can have only 1 value per pixel");var x=Array(r*o),b=Array(r*o),_=Array(s),w=Array(s),k=Array(s+1),T=Array(s);for(d=0,g=r*o;d<g;d++){var A=l[d];x[d]=1===A?0:0===A?i:Math.pow(Math.max(0,.5-A),2),b[d]=1===A?i:0===A?0:Math.pow(Math.max(0,A-.5),2)}a(x,r,o,_,w,T,k),a(b,r,o,_,w,T,k);var M=window.Float32Array?new Float32Array(r*o):new Array(r*o);for(d=0,g=r*o;d<g;d++)M[d]=n(1-((x[d]-b[d])/m+v),0,1);return M};var i=1e20;function a(t,e,r,n,i,a,s){for(var l=0;l<e;l++){for(var c=0;c<r;c++)n[c]=t[c*e+l];for(o(n,i,a,s,r),c=0;c<r;c++)t[c*e+l]=i[c]}for(c=0;c<r;c++){for(l=0;l<e;l++)n[l]=t[c*e+l];for(o(n,i,a,s,e),l=0;l<e;l++)t[c*e+l]=Math.sqrt(i[l])}}function o(t,e,r,n,a){r[0]=0,n[0]=-i,n[1]=+i;for(var o=1,s=0;o<a;o++){for(var l=(t[o]+o*o-(t[r[s]]+r[s]*r[s]))/(2*o-2*r[s]);l<=n[s];)s--,l=(t[o]+o*o-(t[r[s]]+r[s]*r[s]))/(2*o-2*r[s]);r[++s]=o,n[s]=l,n[s+1]=+i}for(o=0,s=0;o<a;o++){for(;n[s+1]<o;)s++;e[o]=(o-r[s])*(o-r[s])+t[r[s]]}}},{clamp:115}],94:[function(t,e,r){!function(e,r){"use strict";function n(t,e){if(!t)throw new Error(e||"Assertion failed")}function i(t,e){t.super_=e;var r=function(){};r.prototype=e.prototype,t.prototype=new r,t.prototype.constructor=t}function a(t,e,r){if(a.isBN(t))return t;this.negative=0,this.words=null,this.length=0,this.red=null,null!==t&&("le"!==e&&"be"!==e||(r=e,e=10),this._init(t||0,e||10,r||"be"))}var o;"object"==typeof e?e.exports=a:r.BN=a,a.BN=a,a.wordSize=26;try{o=t("buffer").Buffer}catch(t){}function s(t,e,r){for(var n=0,i=Math.min(t.length,r),a=e;a<i;a++){var o=t.charCodeAt(a)-48;n<<=4,n|=o>=49&&o<=54?o-49+10:o>=17&&o<=22?o-17+10:15&o}return n}function l(t,e,r,n){for(var i=0,a=Math.min(t.length,r),o=e;o<a;o++){var s=t.charCodeAt(o)-48;i*=n,i+=s>=49?s-49+10:s>=17?s-17+10:s}return i}a.isBN=function(t){return t instanceof a||null!==t&&"object"==typeof t&&t.constructor.wordSize===a.wordSize&&Array.isArray(t.words)},a.max=function(t,e){return t.cmp(e)>0?t:e},a.min=function(t,e){return t.cmp(e)<0?t:e},a.prototype._init=function(t,e,r){if("number"==typeof t)return this._initNumber(t,e,r);if("object"==typeof t)return this._initArray(t,e,r);"hex"===e&&(e=16),n(e===(0|e)&&e>=2&&e<=36);var i=0;"-"===(t=t.toString().replace(/\s+/g,""))[0]&&i++,16===e?this._parseHex(t,i):this._parseBase(t,e,i),"-"===t[0]&&(this.negative=1),this.strip(),"le"===r&&this._initArray(this.toArray(),e,r)},a.prototype._initNumber=function(t,e,r){t<0&&(this.negative=1,t=-t),t<67108864?(this.words=[67108863&t],this.length=1):t<4503599627370496?(this.words=[67108863&t,t/67108864&67108863],this.length=2):(n(t<9007199254740992),this.words=[67108863&t,t/67108864&67108863,1],this.length=3),"le"===r&&this._initArray(this.toArray(),e,r)},a.prototype._initArray=function(t,e,r){if(n("number"==typeof t.length),t.length<=0)return this.words=[0],this.length=1,this;this.length=Math.ceil(t.length/3),this.words=new Array(this.length);for(var i=0;i<this.length;i++)this.words[i]=0;var a,o,s=0;if("be"===r)for(i=t.length-1,a=0;i>=0;i-=3)o=t[i]|t[i-1]<<8|t[i-2]<<16,this.words[a]|=o<<s&67108863,this.words[a+1]=o>>>26-s&67108863,(s+=24)>=26&&(s-=26,a++);else if("le"===r)for(i=0,a=0;i<t.length;i+=3)o=t[i]|t[i+1]<<8|t[i+2]<<16,this.words[a]|=o<<s&67108863,this.words[a+1]=o>>>26-s&67108863,(s+=24)>=26&&(s-=26,a++);return this.strip()},a.prototype._parseHex=function(t,e){this.length=Math.ceil((t.length-e)/6),this.words=new Array(this.length);for(var r=0;r<this.length;r++)this.words[r]=0;var n,i,a=0;for(r=t.length-6,n=0;r>=e;r-=6)i=s(t,r,r+6),this.words[n]|=i<<a&67108863,this.words[n+1]|=i>>>26-a&4194303,(a+=24)>=26&&(a-=26,n++);r+6!==e&&(i=s(t,e,r+6),this.words[n]|=i<<a&67108863,this.words[n+1]|=i>>>26-a&4194303),this.strip()},a.prototype._parseBase=function(t,e,r){this.words=[0],this.length=1;for(var n=0,i=1;i<=67108863;i*=e)n++;n--,i=i/e|0;for(var a=t.length-r,o=a%n,s=Math.min(a,a-o)+r,c=0,u=r;u<s;u+=n)c=l(t,u,u+n,e),this.imuln(i),this.words[0]+c<67108864?this.words[0]+=c:this._iaddn(c);if(0!==o){var h=1;for(c=l(t,u,t.length,e),u=0;u<o;u++)h*=e;this.imuln(h),this.words[0]+c<67108864?this.words[0]+=c:this._iaddn(c)}},a.prototype.copy=function(t){t.words=new Array(this.length);for(var e=0;e<this.length;e++)t.words[e]=this.words[e];t.length=this.length,t.negative=this.negative,t.red=this.red},a.prototype.clone=function(){var t=new a(null);return this.copy(t),t},a.prototype._expand=function(t){for(;this.length<t;)this.words[this.length++]=0;return this},a.prototype.strip=function(){for(;this.length>1&&0===this.words[this.length-1];)this.length--;return this._normSign()},a.prototype._normSign=function(){return 1===this.length&&0===this.words[0]&&(this.negative=0),this},a.prototype.inspect=function(){return(this.red?"<BN-R: ":"<BN: ")+this.toString(16)+">"};var c=["","0","00","000","0000","00000","000000","0000000","00000000","000000000","0000000000","00000000000","000000000000","0000000000000","00000000000000","000000000000000","0000000000000000","00000000000000000","000000000000000000","0000000000000000000","00000000000000000000","000000000000000000000","0000000000000000000000","00000000000000000000000","000000000000000000000000","0000000000000000000000000"],u=[0,0,25,16,12,11,10,9,8,8,7,7,7,7,6,6,6,6,6,6,6,5,5,5,5,5,5,5,5,5,5,5,5,5,5,5,5],h=[0,0,33554432,43046721,16777216,48828125,60466176,40353607,16777216,43046721,1e7,19487171,35831808,62748517,7529536,11390625,16777216,24137569,34012224,47045881,64e6,4084101,5153632,6436343,7962624,9765625,11881376,14348907,17210368,20511149,243e5,28629151,33554432,39135393,45435424,52521875,60466176];function f(t,e,r){r.negative=e.negative^t.negative;var n=t.length+e.length|0;r.length=n,n=n-1|0;var i=0|t.words[0],a=0|e.words[0],o=i*a,s=67108863&o,l=o/67108864|0;r.words[0]=s;for(var c=1;c<n;c++){for(var u=l>>>26,h=67108863&l,f=Math.min(c,e.length-1),p=Math.max(0,c-t.length+1);p<=f;p++){var d=c-p|0;u+=(o=(i=0|t.words[d])*(a=0|e.words[p])+h)/67108864|0,h=67108863&o}r.words[c]=0|h,l=0|u}return 0!==l?r.words[c]=0|l:r.length--,r.strip()}a.prototype.toString=function(t,e){var r;if(e=0|e||1,16===(t=t||10)||"hex"===t){r="";for(var i=0,a=0,o=0;o<this.length;o++){var s=this.words[o],l=(16777215&(s<<i|a)).toString(16);r=0!==(a=s>>>24-i&16777215)||o!==this.length-1?c[6-l.length]+l+r:l+r,(i+=2)>=26&&(i-=26,o--)}for(0!==a&&(r=a.toString(16)+r);r.length%e!=0;)r="0"+r;return 0!==this.negative&&(r="-"+r),r}if(t===(0|t)&&t>=2&&t<=36){var f=u[t],p=h[t];r="";var d=this.clone();for(d.negative=0;!d.isZero();){var g=d.modn(p).toString(t);r=(d=d.idivn(p)).isZero()?g+r:c[f-g.length]+g+r}for(this.isZero()&&(r="0"+r);r.length%e!=0;)r="0"+r;return 0!==this.negative&&(r="-"+r),r}n(!1,"Base should be between 2 and 36")},a.prototype.toNumber=function(){var t=this.words[0];return 2===this.length?t+=67108864*this.words[1]:3===this.length&&1===this.words[2]?t+=4503599627370496+67108864*this.words[1]:this.length>2&&n(!1,"Number can only safely store up to 53 bits"),0!==this.negative?-t:t},a.prototype.toJSON=function(){return this.toString(16)},a.prototype.toBuffer=function(t,e){return n("undefined"!=typeof o),this.toArrayLike(o,t,e)},a.prototype.toArray=function(t,e){return this.toArrayLike(Array,t,e)},a.prototype.toArrayLike=function(t,e,r){var i=this.byteLength(),a=r||Math.max(1,i);n(i<=a,"byte array longer than desired length"),n(a>0,"Requested array length <= 0"),this.strip();var o,s,l="le"===e,c=new t(a),u=this.clone();if(l){for(s=0;!u.isZero();s++)o=u.andln(255),u.iushrn(8),c[s]=o;for(;s<a;s++)c[s]=0}else{for(s=0;s<a-i;s++)c[s]=0;for(s=0;!u.isZero();s++)o=u.andln(255),u.iushrn(8),c[a-s-1]=o}return c},Math.clz32?a.prototype._countBits=function(t){return 32-Math.clz32(t)}:a.prototype._countBits=function(t){var e=t,r=0;return e>=4096&&(r+=13,e>>>=13),e>=64&&(r+=7,e>>>=7),e>=8&&(r+=4,e>>>=4),e>=2&&(r+=2,e>>>=2),r+e},a.prototype._zeroBits=function(t){if(0===t)return 26;var e=t,r=0;return 0==(8191&e)&&(r+=13,e>>>=13),0==(127&e)&&(r+=7,e>>>=7),0==(15&e)&&(r+=4,e>>>=4),0==(3&e)&&(r+=2,e>>>=2),0==(1&e)&&r++,r},a.prototype.bitLength=function(){var t=this.words[this.length-1],e=this._countBits(t);return 26*(this.length-1)+e},a.prototype.zeroBits=function(){if(this.isZero())return 0;for(var t=0,e=0;e<this.length;e++){var r=this._zeroBits(this.words[e]);if(t+=r,26!==r)break}return t},a.prototype.byteLength=function(){return Math.ceil(this.bitLength()/8)},a.prototype.toTwos=function(t){return 0!==this.negative?this.abs().inotn(t).iaddn(1):this.clone()},a.prototype.fromTwos=function(t){return this.testn(t-1)?this.notn(t).iaddn(1).ineg():this.clone()},a.prototype.isNeg=function(){return 0!==this.negative},a.prototype.neg=function(){return this.clone().ineg()},a.prototype.ineg=function(){return this.isZero()||(this.negative^=1),this},a.prototype.iuor=function(t){for(;this.length<t.length;)this.words[this.length++]=0;for(var e=0;e<t.length;e++)this.words[e]=this.words[e]|t.words[e];return this.strip()},a.prototype.ior=function(t){return n(0==(this.negative|t.negative)),this.iuor(t)},a.prototype.or=function(t){return this.length>t.length?this.clone().ior(t):t.clone().ior(this)},a.prototype.uor=function(t){return this.length>t.length?this.clone().iuor(t):t.clone().iuor(this)},a.prototype.iuand=function(t){var e;e=this.length>t.length?t:this;for(var r=0;r<e.length;r++)this.words[r]=this.words[r]&t.words[r];return this.length=e.length,this.strip()},a.prototype.iand=function(t){return n(0==(this.negative|t.negative)),this.iuand(t)},a.prototype.and=function(t){return this.length>t.length?this.clone().iand(t):t.clone().iand(this)},a.prototype.uand=function(t){return this.length>t.length?this.clone().iuand(t):t.clone().iuand(this)},a.prototype.iuxor=function(t){var e,r;this.length>t.length?(e=this,r=t):(e=t,r=this);for(var n=0;n<r.length;n++)this.words[n]=e.words[n]^r.words[n];if(this!==e)for(;n<e.length;n++)this.words[n]=e.words[n];return this.length=e.length,this.strip()},a.prototype.ixor=function(t){return n(0==(this.negative|t.negative)),this.iuxor(t)},a.prototype.xor=function(t){return this.length>t.length?this.clone().ixor(t):t.clone().ixor(this)},a.prototype.uxor=function(t){return this.length>t.length?this.clone().iuxor(t):t.clone().iuxor(this)},a.prototype.inotn=function(t){n("number"==typeof t&&t>=0);var e=0|Math.ceil(t/26),r=t%26;this._expand(e),r>0&&e--;for(var i=0;i<e;i++)this.words[i]=67108863&~this.words[i];return r>0&&(this.words[i]=~this.words[i]&67108863>>26-r),this.strip()},a.prototype.notn=function(t){return this.clone().inotn(t)},a.prototype.setn=function(t,e){n("number"==typeof t&&t>=0);var r=t/26|0,i=t%26;return this._expand(r+1),this.words[r]=e?this.words[r]|1<<i:this.words[r]&~(1<<i),this.strip()},a.prototype.iadd=function(t){var e,r,n;if(0!==this.negative&&0===t.negative)return this.negative=0,e=this.isub(t),this.negative^=1,this._normSign();if(0===this.negative&&0!==t.negative)return t.negative=0,e=this.isub(t),t.negative=1,e._normSign();this.length>t.length?(r=this,n=t):(r=t,n=this);for(var i=0,a=0;a<n.length;a++)e=(0|r.words[a])+(0|n.words[a])+i,this.words[a]=67108863&e,i=e>>>26;for(;0!==i&&a<r.length;a++)e=(0|r.words[a])+i,this.words[a]=67108863&e,i=e>>>26;if(this.length=r.length,0!==i)this.words[this.length]=i,this.length++;else if(r!==this)for(;a<r.length;a++)this.words[a]=r.words[a];return this},a.prototype.add=function(t){var e;return 0!==t.negative&&0===this.negative?(t.negative=0,e=this.sub(t),t.negative^=1,e):0===t.negative&&0!==this.negative?(this.negative=0,e=t.sub(this),this.negative=1,e):this.length>t.length?this.clone().iadd(t):t.clone().iadd(this)},a.prototype.isub=function(t){if(0!==t.negative){t.negative=0;var e=this.iadd(t);return t.negative=1,e._normSign()}if(0!==this.negative)return this.negative=0,this.iadd(t),this.negative=1,this._normSign();var r,n,i=this.cmp(t);if(0===i)return this.negative=0,this.length=1,this.words[0]=0,this;i>0?(r=this,n=t):(r=t,n=this);for(var a=0,o=0;o<n.length;o++)a=(e=(0|r.words[o])-(0|n.words[o])+a)>>26,this.words[o]=67108863&e;for(;0!==a&&o<r.length;o++)a=(e=(0|r.words[o])+a)>>26,this.words[o]=67108863&e;if(0===a&&o<r.length&&r!==this)for(;o<r.length;o++)this.words[o]=r.words[o];return this.length=Math.max(this.length,o),r!==this&&(this.negative=1),this.strip()},a.prototype.sub=function(t){return this.clone().isub(t)};var p=function(t,e,r){var n,i,a,o=t.words,s=e.words,l=r.words,c=0,u=0|o[0],h=8191&u,f=u>>>13,p=0|o[1],d=8191&p,g=p>>>13,v=0|o[2],m=8191&v,y=v>>>13,x=0|o[3],b=8191&x,_=x>>>13,w=0|o[4],k=8191&w,T=w>>>13,A=0|o[5],M=8191&A,S=A>>>13,E=0|o[6],C=8191&E,L=E>>>13,P=0|o[7],O=8191&P,z=P>>>13,I=0|o[8],D=8191&I,R=I>>>13,F=0|o[9],B=8191&F,N=F>>>13,j=0|s[0],U=8191&j,V=j>>>13,q=0|s[1],H=8191&q,G=q>>>13,Y=0|s[2],W=8191&Y,X=Y>>>13,Z=0|s[3],J=8191&Z,K=Z>>>13,$=0|s[4],Q=8191&$,tt=$>>>13,et=0|s[5],rt=8191&et,nt=et>>>13,it=0|s[6],at=8191&it,ot=it>>>13,st=0|s[7],lt=8191&st,ct=st>>>13,ut=0|s[8],ht=8191&ut,ft=ut>>>13,pt=0|s[9],dt=8191&pt,gt=pt>>>13;r.negative=t.negative^e.negative,r.length=19;var vt=(c+(n=Math.imul(h,U))|0)+((8191&(i=(i=Math.imul(h,V))+Math.imul(f,U)|0))<<13)|0;c=((a=Math.imul(f,V))+(i>>>13)|0)+(vt>>>26)|0,vt&=67108863,n=Math.imul(d,U),i=(i=Math.imul(d,V))+Math.imul(g,U)|0,a=Math.imul(g,V);var mt=(c+(n=n+Math.imul(h,H)|0)|0)+((8191&(i=(i=i+Math.imul(h,G)|0)+Math.imul(f,H)|0))<<13)|0;c=((a=a+Math.imul(f,G)|0)+(i>>>13)|0)+(mt>>>26)|0,mt&=67108863,n=Math.imul(m,U),i=(i=Math.imul(m,V))+Math.imul(y,U)|0,a=Math.imul(y,V),n=n+Math.imul(d,H)|0,i=(i=i+Math.imul(d,G)|0)+Math.imul(g,H)|0,a=a+Math.imul(g,G)|0;var yt=(c+(n=n+Math.imul(h,W)|0)|0)+((8191&(i=(i=i+Math.imul(h,X)|0)+Math.imul(f,W)|0))<<13)|0;c=((a=a+Math.imul(f,X)|0)+(i>>>13)|0)+(yt>>>26)|0,yt&=67108863,n=Math.imul(b,U),i=(i=Math.imul(b,V))+Math.imul(_,U)|0,a=Math.imul(_,V),n=n+Math.imul(m,H)|0,i=(i=i+Math.imul(m,G)|0)+Math.imul(y,H)|0,a=a+Math.imul(y,G)|0,n=n+Math.imul(d,W)|0,i=(i=i+Math.imul(d,X)|0)+Math.imul(g,W)|0,a=a+Math.imul(g,X)|0;var xt=(c+(n=n+Math.imul(h,J)|0)|0)+((8191&(i=(i=i+Math.imul(h,K)|0)+Math.imul(f,J)|0))<<13)|0;c=((a=a+Math.imul(f,K)|0)+(i>>>13)|0)+(xt>>>26)|0,xt&=67108863,n=Math.imul(k,U),i=(i=Math.imul(k,V))+Math.imul(T,U)|0,a=Math.imul(T,V),n=n+Math.imul(b,H)|0,i=(i=i+Math.imul(b,G)|0)+Math.imul(_,H)|0,a=a+Math.imul(_,G)|0,n=n+Math.imul(m,W)|0,i=(i=i+Math.imul(m,X)|0)+Math.imul(y,W)|0,a=a+Math.imul(y,X)|0,n=n+Math.imul(d,J)|0,i=(i=i+Math.imul(d,K)|0)+Math.imul(g,J)|0,a=a+Math.imul(g,K)|0;var bt=(c+(n=n+Math.imul(h,Q)|0)|0)+((8191&(i=(i=i+Math.imul(h,tt)|0)+Math.imul(f,Q)|0))<<13)|0;c=((a=a+Math.imul(f,tt)|0)+(i>>>13)|0)+(bt>>>26)|0,bt&=67108863,n=Math.imul(M,U),i=(i=Math.imul(M,V))+Math.imul(S,U)|0,a=Math.imul(S,V),n=n+Math.imul(k,H)|0,i=(i=i+Math.imul(k,G)|0)+Math.imul(T,H)|0,a=a+Math.imul(T,G)|0,n=n+Math.imul(b,W)|0,i=(i=i+Math.imul(b,X)|0)+Math.imul(_,W)|0,a=a+Math.imul(_,X)|0,n=n+Math.imul(m,J)|0,i=(i=i+Math.imul(m,K)|0)+Math.imul(y,J)|0,a=a+Math.imul(y,K)|0,n=n+Math.imul(d,Q)|0,i=(i=i+Math.imul(d,tt)|0)+Math.imul(g,Q)|0,a=a+Math.imul(g,tt)|0;var _t=(c+(n=n+Math.imul(h,rt)|0)|0)+((8191&(i=(i=i+Math.imul(h,nt)|0)+Math.imul(f,rt)|0))<<13)|0;c=((a=a+Math.imul(f,nt)|0)+(i>>>13)|0)+(_t>>>26)|0,_t&=67108863,n=Math.imul(C,U),i=(i=Math.imul(C,V))+Math.imul(L,U)|0,a=Math.imul(L,V),n=n+Math.imul(M,H)|0,i=(i=i+Math.imul(M,G)|0)+Math.imul(S,H)|0,a=a+Math.imul(S,G)|0,n=n+Math.imul(k,W)|0,i=(i=i+Math.imul(k,X)|0)+Math.imul(T,W)|0,a=a+Math.imul(T,X)|0,n=n+Math.imul(b,J)|0,i=(i=i+Math.imul(b,K)|0)+Math.imul(_,J)|0,a=a+Math.imul(_,K)|0,n=n+Math.imul(m,Q)|0,i=(i=i+Math.imul(m,tt)|0)+Math.imul(y,Q)|0,a=a+Math.imul(y,tt)|0,n=n+Math.imul(d,rt)|0,i=(i=i+Math.imul(d,nt)|0)+Math.imul(g,rt)|0,a=a+Math.imul(g,nt)|0;var wt=(c+(n=n+Math.imul(h,at)|0)|0)+((8191&(i=(i=i+Math.imul(h,ot)|0)+Math.imul(f,at)|0))<<13)|0;c=((a=a+Math.imul(f,ot)|0)+(i>>>13)|0)+(wt>>>26)|0,wt&=67108863,n=Math.imul(O,U),i=(i=Math.imul(O,V))+Math.imul(z,U)|0,a=Math.imul(z,V),n=n+Math.imul(C,H)|0,i=(i=i+Math.imul(C,G)|0)+Math.imul(L,H)|0,a=a+Math.imul(L,G)|0,n=n+Math.imul(M,W)|0,i=(i=i+Math.imul(M,X)|0)+Math.imul(S,W)|0,a=a+Math.imul(S,X)|0,n=n+Math.imul(k,J)|0,i=(i=i+Math.imul(k,K)|0)+Math.imul(T,J)|0,a=a+Math.imul(T,K)|0,n=n+Math.imul(b,Q)|0,i=(i=i+Math.imul(b,tt)|0)+Math.imul(_,Q)|0,a=a+Math.imul(_,tt)|0,n=n+Math.imul(m,rt)|0,i=(i=i+Math.imul(m,nt)|0)+Math.imul(y,rt)|0,a=a+Math.imul(y,nt)|0,n=n+Math.imul(d,at)|0,i=(i=i+Math.imul(d,ot)|0)+Math.imul(g,at)|0,a=a+Math.imul(g,ot)|0;var kt=(c+(n=n+Math.imul(h,lt)|0)|0)+((8191&(i=(i=i+Math.imul(h,ct)|0)+Math.imul(f,lt)|0))<<13)|0;c=((a=a+Math.imul(f,ct)|0)+(i>>>13)|0)+(kt>>>26)|0,kt&=67108863,n=Math.imul(D,U),i=(i=Math.imul(D,V))+Math.imul(R,U)|0,a=Math.imul(R,V),n=n+Math.imul(O,H)|0,i=(i=i+Math.imul(O,G)|0)+Math.imul(z,H)|0,a=a+Math.imul(z,G)|0,n=n+Math.imul(C,W)|0,i=(i=i+Math.imul(C,X)|0)+Math.imul(L,W)|0,a=a+Math.imul(L,X)|0,n=n+Math.imul(M,J)|0,i=(i=i+Math.imul(M,K)|0)+Math.imul(S,J)|0,a=a+Math.imul(S,K)|0,n=n+Math.imul(k,Q)|0,i=(i=i+Math.imul(k,tt)|0)+Math.imul(T,Q)|0,a=a+Math.imul(T,tt)|0,n=n+Math.imul(b,rt)|0,i=(i=i+Math.imul(b,nt)|0)+Math.imul(_,rt)|0,a=a+Math.imul(_,nt)|0,n=n+Math.imul(m,at)|0,i=(i=i+Math.imul(m,ot)|0)+Math.imul(y,at)|0,a=a+Math.imul(y,ot)|0,n=n+Math.imul(d,lt)|0,i=(i=i+Math.imul(d,ct)|0)+Math.imul(g,lt)|0,a=a+Math.imul(g,ct)|0;var Tt=(c+(n=n+Math.imul(h,ht)|0)|0)+((8191&(i=(i=i+Math.imul(h,ft)|0)+Math.imul(f,ht)|0))<<13)|0;c=((a=a+Math.imul(f,ft)|0)+(i>>>13)|0)+(Tt>>>26)|0,Tt&=67108863,n=Math.imul(B,U),i=(i=Math.imul(B,V))+Math.imul(N,U)|0,a=Math.imul(N,V),n=n+Math.imul(D,H)|0,i=(i=i+Math.imul(D,G)|0)+Math.imul(R,H)|0,a=a+Math.imul(R,G)|0,n=n+Math.imul(O,W)|0,i=(i=i+Math.imul(O,X)|0)+Math.imul(z,W)|0,a=a+Math.imul(z,X)|0,n=n+Math.imul(C,J)|0,i=(i=i+Math.imul(C,K)|0)+Math.imul(L,J)|0,a=a+Math.imul(L,K)|0,n=n+Math.imul(M,Q)|0,i=(i=i+Math.imul(M,tt)|0)+Math.imul(S,Q)|0,a=a+Math.imul(S,tt)|0,n=n+Math.imul(k,rt)|0,i=(i=i+Math.imul(k,nt)|0)+Math.imul(T,rt)|0,a=a+Math.imul(T,nt)|0,n=n+Math.imul(b,at)|0,i=(i=i+Math.imul(b,ot)|0)+Math.imul(_,at)|0,a=a+Math.imul(_,ot)|0,n=n+Math.imul(m,lt)|0,i=(i=i+Math.imul(m,ct)|0)+Math.imul(y,lt)|0,a=a+Math.imul(y,ct)|0,n=n+Math.imul(d,ht)|0,i=(i=i+Math.imul(d,ft)|0)+Math.imul(g,ht)|0,a=a+Math.imul(g,ft)|0;var At=(c+(n=n+Math.imul(h,dt)|0)|0)+((8191&(i=(i=i+Math.imul(h,gt)|0)+Math.imul(f,dt)|0))<<13)|0;c=((a=a+Math.imul(f,gt)|0)+(i>>>13)|0)+(At>>>26)|0,At&=67108863,n=Math.imul(B,H),i=(i=Math.imul(B,G))+Math.imul(N,H)|0,a=Math.imul(N,G),n=n+Math.imul(D,W)|0,i=(i=i+Math.imul(D,X)|0)+Math.imul(R,W)|0,a=a+Math.imul(R,X)|0,n=n+Math.imul(O,J)|0,i=(i=i+Math.imul(O,K)|0)+Math.imul(z,J)|0,a=a+Math.imul(z,K)|0,n=n+Math.imul(C,Q)|0,i=(i=i+Math.imul(C,tt)|0)+Math.imul(L,Q)|0,a=a+Math.imul(L,tt)|0,n=n+Math.imul(M,rt)|0,i=(i=i+Math.imul(M,nt)|0)+Math.imul(S,rt)|0,a=a+Math.imul(S,nt)|0,n=n+Math.imul(k,at)|0,i=(i=i+Math.imul(k,ot)|0)+Math.imul(T,at)|0,a=a+Math.imul(T,ot)|0,n=n+Math.imul(b,lt)|0,i=(i=i+Math.imul(b,ct)|0)+Math.imul(_,lt)|0,a=a+Math.imul(_,ct)|0,n=n+Math.imul(m,ht)|0,i=(i=i+Math.imul(m,ft)|0)+Math.imul(y,ht)|0,a=a+Math.imul(y,ft)|0;var Mt=(c+(n=n+Math.imul(d,dt)|0)|0)+((8191&(i=(i=i+Math.imul(d,gt)|0)+Math.imul(g,dt)|0))<<13)|0;c=((a=a+Math.imul(g,gt)|0)+(i>>>13)|0)+(Mt>>>26)|0,Mt&=67108863,n=Math.imul(B,W),i=(i=Math.imul(B,X))+Math.imul(N,W)|0,a=Math.imul(N,X),n=n+Math.imul(D,J)|0,i=(i=i+Math.imul(D,K)|0)+Math.imul(R,J)|0,a=a+Math.imul(R,K)|0,n=n+Math.imul(O,Q)|0,i=(i=i+Math.imul(O,tt)|0)+Math.imul(z,Q)|0,a=a+Math.imul(z,tt)|0,n=n+Math.imul(C,rt)|0,i=(i=i+Math.imul(C,nt)|0)+Math.imul(L,rt)|0,a=a+Math.imul(L,nt)|0,n=n+Math.imul(M,at)|0,i=(i=i+Math.imul(M,ot)|0)+Math.imul(S,at)|0,a=a+Math.imul(S,ot)|0,n=n+Math.imul(k,lt)|0,i=(i=i+Math.imul(k,ct)|0)+Math.imul(T,lt)|0,a=a+Math.imul(T,ct)|0,n=n+Math.imul(b,ht)|0,i=(i=i+Math.imul(b,ft)|0)+Math.imul(_,ht)|0,a=a+Math.imul(_,ft)|0;var St=(c+(n=n+Math.imul(m,dt)|0)|0)+((8191&(i=(i=i+Math.imul(m,gt)|0)+Math.imul(y,dt)|0))<<13)|0;c=((a=a+Math.imul(y,gt)|0)+(i>>>13)|0)+(St>>>26)|0,St&=67108863,n=Math.imul(B,J),i=(i=Math.imul(B,K))+Math.imul(N,J)|0,a=Math.imul(N,K),n=n+Math.imul(D,Q)|0,i=(i=i+Math.imul(D,tt)|0)+Math.imul(R,Q)|0,a=a+Math.imul(R,tt)|0,n=n+Math.imul(O,rt)|0,i=(i=i+Math.imul(O,nt)|0)+Math.imul(z,rt)|0,a=a+Math.imul(z,nt)|0,n=n+Math.imul(C,at)|0,i=(i=i+Math.imul(C,ot)|0)+Math.imul(L,at)|0,a=a+Math.imul(L,ot)|0,n=n+Math.imul(M,lt)|0,i=(i=i+Math.imul(M,ct)|0)+Math.imul(S,lt)|0,a=a+Math.imul(S,ct)|0,n=n+Math.imul(k,ht)|0,i=(i=i+Math.imul(k,ft)|0)+Math.imul(T,ht)|0,a=a+Math.imul(T,ft)|0;var Et=(c+(n=n+Math.imul(b,dt)|0)|0)+((8191&(i=(i=i+Math.imul(b,gt)|0)+Math.imul(_,dt)|0))<<13)|0;c=((a=a+Math.imul(_,gt)|0)+(i>>>13)|0)+(Et>>>26)|0,Et&=67108863,n=Math.imul(B,Q),i=(i=Math.imul(B,tt))+Math.imul(N,Q)|0,a=Math.imul(N,tt),n=n+Math.imul(D,rt)|0,i=(i=i+Math.imul(D,nt)|0)+Math.imul(R,rt)|0,a=a+Math.imul(R,nt)|0,n=n+Math.imul(O,at)|0,i=(i=i+Math.imul(O,ot)|0)+Math.imul(z,at)|0,a=a+Math.imul(z,ot)|0,n=n+Math.imul(C,lt)|0,i=(i=i+Math.imul(C,ct)|0)+Math.imul(L,lt)|0,a=a+Math.imul(L,ct)|0,n=n+Math.imul(M,ht)|0,i=(i=i+Math.imul(M,ft)|0)+Math.imul(S,ht)|0,a=a+Math.imul(S,ft)|0;var Ct=(c+(n=n+Math.imul(k,dt)|0)|0)+((8191&(i=(i=i+Math.imul(k,gt)|0)+Math.imul(T,dt)|0))<<13)|0;c=((a=a+Math.imul(T,gt)|0)+(i>>>13)|0)+(Ct>>>26)|0,Ct&=67108863,n=Math.imul(B,rt),i=(i=Math.imul(B,nt))+Math.imul(N,rt)|0,a=Math.imul(N,nt),n=n+Math.imul(D,at)|0,i=(i=i+Math.imul(D,ot)|0)+Math.imul(R,at)|0,a=a+Math.imul(R,ot)|0,n=n+Math.imul(O,lt)|0,i=(i=i+Math.imul(O,ct)|0)+Math.imul(z,lt)|0,a=a+Math.imul(z,ct)|0,n=n+Math.imul(C,ht)|0,i=(i=i+Math.imul(C,ft)|0)+Math.imul(L,ht)|0,a=a+Math.imul(L,ft)|0;var Lt=(c+(n=n+Math.imul(M,dt)|0)|0)+((8191&(i=(i=i+Math.imul(M,gt)|0)+Math.imul(S,dt)|0))<<13)|0;c=((a=a+Math.imul(S,gt)|0)+(i>>>13)|0)+(Lt>>>26)|0,Lt&=67108863,n=Math.imul(B,at),i=(i=Math.imul(B,ot))+Math.imul(N,at)|0,a=Math.imul(N,ot),n=n+Math.imul(D,lt)|0,i=(i=i+Math.imul(D,ct)|0)+Math.imul(R,lt)|0,a=a+Math.imul(R,ct)|0,n=n+Math.imul(O,ht)|0,i=(i=i+Math.imul(O,ft)|0)+Math.imul(z,ht)|0,a=a+Math.imul(z,ft)|0;var Pt=(c+(n=n+Math.imul(C,dt)|0)|0)+((8191&(i=(i=i+Math.imul(C,gt)|0)+Math.imul(L,dt)|0))<<13)|0;c=((a=a+Math.imul(L,gt)|0)+(i>>>13)|0)+(Pt>>>26)|0,Pt&=67108863,n=Math.imul(B,lt),i=(i=Math.imul(B,ct))+Math.imul(N,lt)|0,a=Math.imul(N,ct),n=n+Math.imul(D,ht)|0,i=(i=i+Math.imul(D,ft)|0)+Math.imul(R,ht)|0,a=a+Math.imul(R,ft)|0;var Ot=(c+(n=n+Math.imul(O,dt)|0)|0)+((8191&(i=(i=i+Math.imul(O,gt)|0)+Math.imul(z,dt)|0))<<13)|0;c=((a=a+Math.imul(z,gt)|0)+(i>>>13)|0)+(Ot>>>26)|0,Ot&=67108863,n=Math.imul(B,ht),i=(i=Math.imul(B,ft))+Math.imul(N,ht)|0,a=Math.imul(N,ft);var zt=(c+(n=n+Math.imul(D,dt)|0)|0)+((8191&(i=(i=i+Math.imul(D,gt)|0)+Math.imul(R,dt)|0))<<13)|0;c=((a=a+Math.imul(R,gt)|0)+(i>>>13)|0)+(zt>>>26)|0,zt&=67108863;var It=(c+(n=Math.imul(B,dt))|0)+((8191&(i=(i=Math.imul(B,gt))+Math.imul(N,dt)|0))<<13)|0;return c=((a=Math.imul(N,gt))+(i>>>13)|0)+(It>>>26)|0,It&=67108863,l[0]=vt,l[1]=mt,l[2]=yt,l[3]=xt,l[4]=bt,l[5]=_t,l[6]=wt,l[7]=kt,l[8]=Tt,l[9]=At,l[10]=Mt,l[11]=St,l[12]=Et,l[13]=Ct,l[14]=Lt,l[15]=Pt,l[16]=Ot,l[17]=zt,l[18]=It,0!==c&&(l[19]=c,r.length++),r};function d(t,e,r){return(new g).mulp(t,e,r)}function g(t,e){this.x=t,this.y=e}Math.imul||(p=f),a.prototype.mulTo=function(t,e){var r=this.length+t.length;return 10===this.length&&10===t.length?p(this,t,e):r<63?f(this,t,e):r<1024?function(t,e,r){r.negative=e.negative^t.negative,r.length=t.length+e.length;for(var n=0,i=0,a=0;a<r.length-1;a++){var o=i;i=0;for(var s=67108863&n,l=Math.min(a,e.length-1),c=Math.max(0,a-t.length+1);c<=l;c++){var u=a-c,h=(0|t.words[u])*(0|e.words[c]),f=67108863&h;s=67108863&(f=f+s|0),i+=(o=(o=o+(h/67108864|0)|0)+(f>>>26)|0)>>>26,o&=67108863}r.words[a]=s,n=o,o=i}return 0!==n?r.words[a]=n:r.length--,r.strip()}(this,t,e):d(this,t,e)},g.prototype.makeRBT=function(t){for(var e=new Array(t),r=a.prototype._countBits(t)-1,n=0;n<t;n++)e[n]=this.revBin(n,r,t);return e},g.prototype.revBin=function(t,e,r){if(0===t||t===r-1)return t;for(var n=0,i=0;i<e;i++)n|=(1&t)<<e-i-1,t>>=1;return n},g.prototype.permute=function(t,e,r,n,i,a){for(var o=0;o<a;o++)n[o]=e[t[o]],i[o]=r[t[o]]},g.prototype.transform=function(t,e,r,n,i,a){this.permute(a,t,e,r,n,i);for(var o=1;o<i;o<<=1)for(var s=o<<1,l=Math.cos(2*Math.PI/s),c=Math.sin(2*Math.PI/s),u=0;u<i;u+=s)for(var h=l,f=c,p=0;p<o;p++){var d=r[u+p],g=n[u+p],v=r[u+p+o],m=n[u+p+o],y=h*v-f*m;m=h*m+f*v,v=y,r[u+p]=d+v,n[u+p]=g+m,r[u+p+o]=d-v,n[u+p+o]=g-m,p!==s&&(y=l*h-c*f,f=l*f+c*h,h=y)}},g.prototype.guessLen13b=function(t,e){var r=1|Math.max(e,t),n=1&r,i=0;for(r=r/2|0;r;r>>>=1)i++;return 1<<i+1+n},g.prototype.conjugate=function(t,e,r){if(!(r<=1))for(var n=0;n<r/2;n++){var i=t[n];t[n]=t[r-n-1],t[r-n-1]=i,i=e[n],e[n]=-e[r-n-1],e[r-n-1]=-i}},g.prototype.normalize13b=function(t,e){for(var r=0,n=0;n<e/2;n++){var i=8192*Math.round(t[2*n+1]/e)+Math.round(t[2*n]/e)+r;t[n]=67108863&i,r=i<67108864?0:i/67108864|0}return t},g.prototype.convert13b=function(t,e,r,i){for(var a=0,o=0;o<e;o++)a+=0|t[o],r[2*o]=8191&a,a>>>=13,r[2*o+1]=8191&a,a>>>=13;for(o=2*e;o<i;++o)r[o]=0;n(0===a),n(0==(-8192&a))},g.prototype.stub=function(t){for(var e=new Array(t),r=0;r<t;r++)e[r]=0;return e},g.prototype.mulp=function(t,e,r){var n=2*this.guessLen13b(t.length,e.length),i=this.makeRBT(n),a=this.stub(n),o=new Array(n),s=new Array(n),l=new Array(n),c=new Array(n),u=new Array(n),h=new Array(n),f=r.words;f.length=n,this.convert13b(t.words,t.length,o,n),this.convert13b(e.words,e.length,c,n),this.transform(o,a,s,l,n,i),this.transform(c,a,u,h,n,i);for(var p=0;p<n;p++){var d=s[p]*u[p]-l[p]*h[p];l[p]=s[p]*h[p]+l[p]*u[p],s[p]=d}return this.conjugate(s,l,n),this.transform(s,l,f,a,n,i),this.conjugate(f,a,n),this.normalize13b(f,n),r.negative=t.negative^e.negative,r.length=t.length+e.length,r.strip()},a.prototype.mul=function(t){var e=new a(null);return e.words=new Array(this.length+t.length),this.mulTo(t,e)},a.prototype.mulf=function(t){var e=new a(null);return e.words=new Array(this.length+t.length),d(this,t,e)},a.prototype.imul=function(t){return this.clone().mulTo(t,this)},a.prototype.imuln=function(t){n("number"==typeof t),n(t<67108864);for(var e=0,r=0;r<this.length;r++){var i=(0|this.words[r])*t,a=(67108863&i)+(67108863&e);e>>=26,e+=i/67108864|0,e+=a>>>26,this.words[r]=67108863&a}return 0!==e&&(this.words[r]=e,this.length++),this},a.prototype.muln=function(t){return this.clone().imuln(t)},a.prototype.sqr=function(){return this.mul(this)},a.prototype.isqr=function(){return this.imul(this.clone())},a.prototype.pow=function(t){var e=function(t){for(var e=new Array(t.bitLength()),r=0;r<e.length;r++){var n=r/26|0,i=r%26;e[r]=(t.words[n]&1<<i)>>>i}return e}(t);if(0===e.length)return new a(1);for(var r=this,n=0;n<e.length&&0===e[n];n++,r=r.sqr());if(++n<e.length)for(var i=r.sqr();n<e.length;n++,i=i.sqr())0!==e[n]&&(r=r.mul(i));return r},a.prototype.iushln=function(t){n("number"==typeof t&&t>=0);var e,r=t%26,i=(t-r)/26,a=67108863>>>26-r<<26-r;if(0!==r){var o=0;for(e=0;e<this.length;e++){var s=this.words[e]&a,l=(0|this.words[e])-s<<r;this.words[e]=l|o,o=s>>>26-r}o&&(this.words[e]=o,this.length++)}if(0!==i){for(e=this.length-1;e>=0;e--)this.words[e+i]=this.words[e];for(e=0;e<i;e++)this.words[e]=0;this.length+=i}return this.strip()},a.prototype.ishln=function(t){return n(0===this.negative),this.iushln(t)},a.prototype.iushrn=function(t,e,r){var i;n("number"==typeof t&&t>=0),i=e?(e-e%26)/26:0;var a=t%26,o=Math.min((t-a)/26,this.length),s=67108863^67108863>>>a<<a,l=r;if(i-=o,i=Math.max(0,i),l){for(var c=0;c<o;c++)l.words[c]=this.words[c];l.length=o}if(0===o);else if(this.length>o)for(this.length-=o,c=0;c<this.length;c++)this.words[c]=this.words[c+o];else this.words[0]=0,this.length=1;var u=0;for(c=this.length-1;c>=0&&(0!==u||c>=i);c--){var h=0|this.words[c];this.words[c]=u<<26-a|h>>>a,u=h&s}return l&&0!==u&&(l.words[l.length++]=u),0===this.length&&(this.words[0]=0,this.length=1),this.strip()},a.prototype.ishrn=function(t,e,r){return n(0===this.negative),this.iushrn(t,e,r)},a.prototype.shln=function(t){return this.clone().ishln(t)},a.prototype.ushln=function(t){return this.clone().iushln(t)},a.prototype.shrn=function(t){return this.clone().ishrn(t)},a.prototype.ushrn=function(t){return this.clone().iushrn(t)},a.prototype.testn=function(t){n("number"==typeof t&&t>=0);var e=t%26,r=(t-e)/26,i=1<<e;return!(this.length<=r)&&!!(this.words[r]&i)},a.prototype.imaskn=function(t){n("number"==typeof t&&t>=0);var e=t%26,r=(t-e)/26;if(n(0===this.negative,"imaskn works only with positive numbers"),this.length<=r)return this;if(0!==e&&r++,this.length=Math.min(r,this.length),0!==e){var i=67108863^67108863>>>e<<e;this.words[this.length-1]&=i}return this.strip()},a.prototype.maskn=function(t){return this.clone().imaskn(t)},a.prototype.iaddn=function(t){return n("number"==typeof t),n(t<67108864),t<0?this.isubn(-t):0!==this.negative?1===this.length&&(0|this.words[0])<t?(this.words[0]=t-(0|this.words[0]),this.negative=0,this):(this.negative=0,this.isubn(t),this.negative=1,this):this._iaddn(t)},a.prototype._iaddn=function(t){this.words[0]+=t;for(var e=0;e<this.length&&this.words[e]>=67108864;e++)this.words[e]-=67108864,e===this.length-1?this.words[e+1]=1:this.words[e+1]++;return this.length=Math.max(this.length,e+1),this},a.prototype.isubn=function(t){if(n("number"==typeof t),n(t<67108864),t<0)return this.iaddn(-t);if(0!==this.negative)return this.negative=0,this.iaddn(t),this.negative=1,this;if(this.words[0]-=t,1===this.length&&this.words[0]<0)this.words[0]=-this.words[0],this.negative=1;else for(var e=0;e<this.length&&this.words[e]<0;e++)this.words[e]+=67108864,this.words[e+1]-=1;return this.strip()},a.prototype.addn=function(t){return this.clone().iaddn(t)},a.prototype.subn=function(t){return this.clone().isubn(t)},a.prototype.iabs=function(){return this.negative=0,this},a.prototype.abs=function(){return this.clone().iabs()},a.prototype._ishlnsubmul=function(t,e,r){var i,a,o=t.length+r;this._expand(o);var s=0;for(i=0;i<t.length;i++){a=(0|this.words[i+r])+s;var l=(0|t.words[i])*e;s=((a-=67108863&l)>>26)-(l/67108864|0),this.words[i+r]=67108863&a}for(;i<this.length-r;i++)s=(a=(0|this.words[i+r])+s)>>26,this.words[i+r]=67108863&a;if(0===s)return this.strip();for(n(-1===s),s=0,i=0;i<this.length;i++)s=(a=-(0|this.words[i])+s)>>26,this.words[i]=67108863&a;return this.negative=1,this.strip()},a.prototype._wordDiv=function(t,e){var r=(this.length,t.length),n=this.clone(),i=t,o=0|i.words[i.length-1];0!==(r=26-this._countBits(o))&&(i=i.ushln(r),n.iushln(r),o=0|i.words[i.length-1]);var s,l=n.length-i.length;if("mod"!==e){(s=new a(null)).length=l+1,s.words=new Array(s.length);for(var c=0;c<s.length;c++)s.words[c]=0}var u=n.clone()._ishlnsubmul(i,1,l);0===u.negative&&(n=u,s&&(s.words[l]=1));for(var h=l-1;h>=0;h--){var f=67108864*(0|n.words[i.length+h])+(0|n.words[i.length+h-1]);for(f=Math.min(f/o|0,67108863),n._ishlnsubmul(i,f,h);0!==n.negative;)f--,n.negative=0,n._ishlnsubmul(i,1,h),n.isZero()||(n.negative^=1);s&&(s.words[h]=f)}return s&&s.strip(),n.strip(),"div"!==e&&0!==r&&n.iushrn(r),{div:s||null,mod:n}},a.prototype.divmod=function(t,e,r){return n(!t.isZero()),this.isZero()?{div:new a(0),mod:new a(0)}:0!==this.negative&&0===t.negative?(s=this.neg().divmod(t,e),"mod"!==e&&(i=s.div.neg()),"div"!==e&&(o=s.mod.neg(),r&&0!==o.negative&&o.iadd(t)),{div:i,mod:o}):0===this.negative&&0!==t.negative?(s=this.divmod(t.neg(),e),"mod"!==e&&(i=s.div.neg()),{div:i,mod:s.mod}):0!=(this.negative&t.negative)?(s=this.neg().divmod(t.neg(),e),"div"!==e&&(o=s.mod.neg(),r&&0!==o.negative&&o.isub(t)),{div:s.div,mod:o}):t.length>this.length||this.cmp(t)<0?{div:new a(0),mod:this}:1===t.length?"div"===e?{div:this.divn(t.words[0]),mod:null}:"mod"===e?{div:null,mod:new a(this.modn(t.words[0]))}:{div:this.divn(t.words[0]),mod:new a(this.modn(t.words[0]))}:this._wordDiv(t,e);var i,o,s},a.prototype.div=function(t){return this.divmod(t,"div",!1).div},a.prototype.mod=function(t){return this.divmod(t,"mod",!1).mod},a.prototype.umod=function(t){return this.divmod(t,"mod",!0).mod},a.prototype.divRound=function(t){var e=this.divmod(t);if(e.mod.isZero())return e.div;var r=0!==e.div.negative?e.mod.isub(t):e.mod,n=t.ushrn(1),i=t.andln(1),a=r.cmp(n);return a<0||1===i&&0===a?e.div:0!==e.div.negative?e.div.isubn(1):e.div.iaddn(1)},a.prototype.modn=function(t){n(t<=67108863);for(var e=(1<<26)%t,r=0,i=this.length-1;i>=0;i--)r=(e*r+(0|this.words[i]))%t;return r},a.prototype.idivn=function(t){n(t<=67108863);for(var e=0,r=this.length-1;r>=0;r--){var i=(0|this.words[r])+67108864*e;this.words[r]=i/t|0,e=i%t}return this.strip()},a.prototype.divn=function(t){return this.clone().idivn(t)},a.prototype.egcd=function(t){n(0===t.negative),n(!t.isZero());var e=this,r=t.clone();e=0!==e.negative?e.umod(t):e.clone();for(var i=new a(1),o=new a(0),s=new a(0),l=new a(1),c=0;e.isEven()&&r.isEven();)e.iushrn(1),r.iushrn(1),++c;for(var u=r.clone(),h=e.clone();!e.isZero();){for(var f=0,p=1;0==(e.words[0]&p)&&f<26;++f,p<<=1);if(f>0)for(e.iushrn(f);f-- >0;)(i.isOdd()||o.isOdd())&&(i.iadd(u),o.isub(h)),i.iushrn(1),o.iushrn(1);for(var d=0,g=1;0==(r.words[0]&g)&&d<26;++d,g<<=1);if(d>0)for(r.iushrn(d);d-- >0;)(s.isOdd()||l.isOdd())&&(s.iadd(u),l.isub(h)),s.iushrn(1),l.iushrn(1);e.cmp(r)>=0?(e.isub(r),i.isub(s),o.isub(l)):(r.isub(e),s.isub(i),l.isub(o))}return{a:s,b:l,gcd:r.iushln(c)}},a.prototype._invmp=function(t){n(0===t.negative),n(!t.isZero());var e=this,r=t.clone();e=0!==e.negative?e.umod(t):e.clone();for(var i,o=new a(1),s=new a(0),l=r.clone();e.cmpn(1)>0&&r.cmpn(1)>0;){for(var c=0,u=1;0==(e.words[0]&u)&&c<26;++c,u<<=1);if(c>0)for(e.iushrn(c);c-- >0;)o.isOdd()&&o.iadd(l),o.iushrn(1);for(var h=0,f=1;0==(r.words[0]&f)&&h<26;++h,f<<=1);if(h>0)for(r.iushrn(h);h-- >0;)s.isOdd()&&s.iadd(l),s.iushrn(1);e.cmp(r)>=0?(e.isub(r),o.isub(s)):(r.isub(e),s.isub(o))}return(i=0===e.cmpn(1)?o:s).cmpn(0)<0&&i.iadd(t),i},a.prototype.gcd=function(t){if(this.isZero())return t.abs();if(t.isZero())return this.abs();var e=this.clone(),r=t.clone();e.negative=0,r.negative=0;for(var n=0;e.isEven()&&r.isEven();n++)e.iushrn(1),r.iushrn(1);for(;;){for(;e.isEven();)e.iushrn(1);for(;r.isEven();)r.iushrn(1);var i=e.cmp(r);if(i<0){var a=e;e=r,r=a}else if(0===i||0===r.cmpn(1))break;e.isub(r)}return r.iushln(n)},a.prototype.invm=function(t){return this.egcd(t).a.umod(t)},a.prototype.isEven=function(){return 0==(1&this.words[0])},a.prototype.isOdd=function(){return 1==(1&this.words[0])},a.prototype.andln=function(t){return this.words[0]&t},a.prototype.bincn=function(t){n("number"==typeof t);var e=t%26,r=(t-e)/26,i=1<<e;if(this.length<=r)return this._expand(r+1),this.words[r]|=i,this;for(var a=i,o=r;0!==a&&o<this.length;o++){var s=0|this.words[o];a=(s+=a)>>>26,s&=67108863,this.words[o]=s}return 0!==a&&(this.words[o]=a,this.length++),this},a.prototype.isZero=function(){return 1===this.length&&0===this.words[0]},a.prototype.cmpn=function(t){var e,r=t<0;if(0!==this.negative&&!r)return-1;if(0===this.negative&&r)return 1;if(this.strip(),this.length>1)e=1;else{r&&(t=-t),n(t<=67108863,"Number is too big");var i=0|this.words[0];e=i===t?0:i<t?-1:1}return 0!==this.negative?0|-e:e},a.prototype.cmp=function(t){if(0!==this.negative&&0===t.negative)return-1;if(0===this.negative&&0!==t.negative)return 1;var e=this.ucmp(t);return 0!==this.negative?0|-e:e},a.prototype.ucmp=function(t){if(this.length>t.length)return 1;if(this.length<t.length)return-1;for(var e=0,r=this.length-1;r>=0;r--){var n=0|this.words[r],i=0|t.words[r];if(n!==i){n<i?e=-1:n>i&&(e=1);break}}return e},a.prototype.gtn=function(t){return 1===this.cmpn(t)},a.prototype.gt=function(t){return 1===this.cmp(t)},a.prototype.gten=function(t){return this.cmpn(t)>=0},a.prototype.gte=function(t){return this.cmp(t)>=0},a.prototype.ltn=function(t){return-1===this.cmpn(t)},a.prototype.lt=function(t){return-1===this.cmp(t)},a.prototype.lten=function(t){return this.cmpn(t)<=0},a.prototype.lte=function(t){return this.cmp(t)<=0},a.prototype.eqn=function(t){return 0===this.cmpn(t)},a.prototype.eq=function(t){return 0===this.cmp(t)},a.red=function(t){return new w(t)},a.prototype.toRed=function(t){return n(!this.red,"Already a number in reduction context"),n(0===this.negative,"red works only with positives"),t.convertTo(this)._forceRed(t)},a.prototype.fromRed=function(){return n(this.red,"fromRed works only with numbers in reduction context"),this.red.convertFrom(this)},a.prototype._forceRed=function(t){return this.red=t,this},a.prototype.forceRed=function(t){return n(!this.red,"Already a number in reduction context"),this._forceRed(t)},a.prototype.redAdd=function(t){return n(this.red,"redAdd works only with red numbers"),this.red.add(this,t)},a.prototype.redIAdd=function(t){return n(this.red,"redIAdd works only with red numbers"),this.red.iadd(this,t)},a.prototype.redSub=function(t){return n(this.red,"redSub works only with red numbers"),this.red.sub(this,t)},a.prototype.redISub=function(t){return n(this.red,"redISub works only with red numbers"),this.red.isub(this,t)},a.prototype.redShl=function(t){return n(this.red,"redShl works only with red numbers"),this.red.shl(this,t)},a.prototype.redMul=function(t){return n(this.red,"redMul works only with red numbers"),this.red._verify2(this,t),this.red.mul(this,t)},a.prototype.redIMul=function(t){return n(this.red,"redMul works only with red numbers"),this.red._verify2(this,t),this.red.imul(this,t)},a.prototype.redSqr=function(){return n(this.red,"redSqr works only with red numbers"),this.red._verify1(this),this.red.sqr(this)},a.prototype.redISqr=function(){return n(this.red,"redISqr works only with red numbers"),this.red._verify1(this),this.red.isqr(this)},a.prototype.redSqrt=function(){return n(this.red,"redSqrt works only with red numbers"),this.red._verify1(this),this.red.sqrt(this)},a.prototype.redInvm=function(){return n(this.red,"redInvm works only with red numbers"),this.red._verify1(this),this.red.invm(this)},a.prototype.redNeg=function(){return n(this.red,"redNeg works only with red numbers"),this.red._verify1(this),this.red.neg(this)},a.prototype.redPow=function(t){return n(this.red&&!t.red,"redPow(normalNum)"),this.red._verify1(this),this.red.pow(this,t)};var v={k256:null,p224:null,p192:null,p25519:null};function m(t,e){this.name=t,this.p=new a(e,16),this.n=this.p.bitLength(),this.k=new a(1).iushln(this.n).isub(this.p),this.tmp=this._tmp()}function y(){m.call(this,"k256","ffffffff ffffffff ffffffff ffffffff ffffffff ffffffff fffffffe fffffc2f")}function x(){m.call(this,"p224","ffffffff ffffffff ffffffff ffffffff 00000000 00000000 00000001")}function b(){m.call(this,"p192","ffffffff ffffffff ffffffff fffffffe ffffffff ffffffff")}function _(){m.call(this,"25519","7fffffffffffffff ffffffffffffffff ffffffffffffffff ffffffffffffffed")}function w(t){if("string"==typeof t){var e=a._prime(t);this.m=e.p,this.prime=e}else n(t.gtn(1),"modulus must be greater than 1"),this.m=t,this.prime=null}function k(t){w.call(this,t),this.shift=this.m.bitLength(),this.shift%26!=0&&(this.shift+=26-this.shift%26),this.r=new a(1).iushln(this.shift),this.r2=this.imod(this.r.sqr()),this.rinv=this.r._invmp(this.m),this.minv=this.rinv.mul(this.r).isubn(1).div(this.m),this.minv=this.minv.umod(this.r),this.minv=this.r.sub(this.minv)}m.prototype._tmp=function(){var t=new a(null);return t.words=new Array(Math.ceil(this.n/13)),t},m.prototype.ireduce=function(t){var e,r=t;do{this.split(r,this.tmp),e=(r=(r=this.imulK(r)).iadd(this.tmp)).bitLength()}while(e>this.n);var n=e<this.n?-1:r.ucmp(this.p);return 0===n?(r.words[0]=0,r.length=1):n>0?r.isub(this.p):r.strip(),r},m.prototype.split=function(t,e){t.iushrn(this.n,0,e)},m.prototype.imulK=function(t){return t.imul(this.k)},i(y,m),y.prototype.split=function(t,e){for(var r=Math.min(t.length,9),n=0;n<r;n++)e.words[n]=t.words[n];if(e.length=r,t.length<=9)return t.words[0]=0,void(t.length=1);var i=t.words[9];for(e.words[e.length++]=4194303&i,n=10;n<t.length;n++){var a=0|t.words[n];t.words[n-10]=(4194303&a)<<4|i>>>22,i=a}i>>>=22,t.words[n-10]=i,0===i&&t.length>10?t.length-=10:t.length-=9},y.prototype.imulK=function(t){t.words[t.length]=0,t.words[t.length+1]=0,t.length+=2;for(var e=0,r=0;r<t.length;r++){var n=0|t.words[r];e+=977*n,t.words[r]=67108863&e,e=64*n+(e/67108864|0)}return 0===t.words[t.length-1]&&(t.length--,0===t.words[t.length-1]&&t.length--),t},i(x,m),i(b,m),i(_,m),_.prototype.imulK=function(t){for(var e=0,r=0;r<t.length;r++){var n=19*(0|t.words[r])+e,i=67108863&n;n>>>=26,t.words[r]=i,e=n}return 0!==e&&(t.words[t.length++]=e),t},a._prime=function(t){if(v[t])return v[t];var e;if("k256"===t)e=new y;else if("p224"===t)e=new x;else if("p192"===t)e=new b;else{if("p25519"!==t)throw new Error("Unknown prime "+t);e=new _}return v[t]=e,e},w.prototype._verify1=function(t){n(0===t.negative,"red works only with positives"),n(t.red,"red works only with red numbers")},w.prototype._verify2=function(t,e){n(0==(t.negative|e.negative),"red works only with positives"),n(t.red&&t.red===e.red,"red works only with red numbers")},w.prototype.imod=function(t){return this.prime?this.prime.ireduce(t)._forceRed(this):t.umod(this.m)._forceRed(this)},w.prototype.neg=function(t){return t.isZero()?t.clone():this.m.sub(t)._forceRed(this)},w.prototype.add=function(t,e){this._verify2(t,e);var r=t.add(e);return r.cmp(this.m)>=0&&r.isub(this.m),r._forceRed(this)},w.prototype.iadd=function(t,e){this._verify2(t,e);var r=t.iadd(e);return r.cmp(this.m)>=0&&r.isub(this.m),r},w.prototype.sub=function(t,e){this._verify2(t,e);var r=t.sub(e);return r.cmpn(0)<0&&r.iadd(this.m),r._forceRed(this)},w.prototype.isub=function(t,e){this._verify2(t,e);var r=t.isub(e);return r.cmpn(0)<0&&r.iadd(this.m),r},w.prototype.shl=function(t,e){return this._verify1(t),this.imod(t.ushln(e))},w.prototype.imul=function(t,e){return this._verify2(t,e),this.imod(t.imul(e))},w.prototype.mul=function(t,e){return this._verify2(t,e),this.imod(t.mul(e))},w.prototype.isqr=function(t){return this.imul(t,t.clone())},w.prototype.sqr=function(t){return this.mul(t,t)},w.prototype.sqrt=function(t){if(t.isZero())return t.clone();var e=this.m.andln(3);if(n(e%2==1),3===e){var r=this.m.add(new a(1)).iushrn(2);return this.pow(t,r)}for(var i=this.m.subn(1),o=0;!i.isZero()&&0===i.andln(1);)o++,i.iushrn(1);n(!i.isZero());var s=new a(1).toRed(this),l=s.redNeg(),c=this.m.subn(1).iushrn(1),u=this.m.bitLength();for(u=new a(2*u*u).toRed(this);0!==this.pow(u,c).cmp(l);)u.redIAdd(l);for(var h=this.pow(u,i),f=this.pow(t,i.addn(1).iushrn(1)),p=this.pow(t,i),d=o;0!==p.cmp(s);){for(var g=p,v=0;0!==g.cmp(s);v++)g=g.redSqr();n(v<d);var m=this.pow(h,new a(1).iushln(d-v-1));f=f.redMul(m),h=m.redSqr(),p=p.redMul(h),d=v}return f},w.prototype.invm=function(t){var e=t._invmp(this.m);return 0!==e.negative?(e.negative=0,this.imod(e).redNeg()):this.imod(e)},w.prototype.pow=function(t,e){if(e.isZero())return new a(1).toRed(this);if(0===e.cmpn(1))return t.clone();var r=new Array(16);r[0]=new a(1).toRed(this),r[1]=t;for(var n=2;n<r.length;n++)r[n]=this.mul(r[n-1],t);var i=r[0],o=0,s=0,l=e.bitLength()%26;for(0===l&&(l=26),n=e.length-1;n>=0;n--){for(var c=e.words[n],u=l-1;u>=0;u--){var h=c>>u&1;i!==r[0]&&(i=this.sqr(i)),0!==h||0!==o?(o<<=1,o|=h,(4===++s||0===n&&0===u)&&(i=this.mul(i,r[o]),s=0,o=0)):s=0}l=26}return i},w.prototype.convertTo=function(t){var e=t.umod(this.m);return e===t?e.clone():e},w.prototype.convertFrom=function(t){var e=t.clone();return e.red=null,e},a.mont=function(t){return new k(t)},i(k,w),k.prototype.convertTo=function(t){return this.imod(t.ushln(this.shift))},k.prototype.convertFrom=function(t){var e=this.imod(t.mul(this.rinv));return e.red=null,e},k.prototype.imul=function(t,e){if(t.isZero()||e.isZero())return t.words[0]=0,t.length=1,t;var r=t.imul(e),n=r.maskn(this.shift).mul(this.minv).imaskn(this.shift).mul(this.m),i=r.isub(n).iushrn(this.shift),a=i;return i.cmp(this.m)>=0?a=i.isub(this.m):i.cmpn(0)<0&&(a=i.iadd(this.m)),a._forceRed(this)},k.prototype.mul=function(t,e){if(t.isZero()||e.isZero())return new a(0)._forceRed(this);var r=t.mul(e),n=r.maskn(this.shift).mul(this.minv).imaskn(this.shift).mul(this.m),i=r.isub(n).iushrn(this.shift),o=i;return i.cmp(this.m)>=0?o=i.isub(this.m):i.cmpn(0)<0&&(o=i.iadd(this.m)),o._forceRed(this)},k.prototype.invm=function(t){return this.imod(t._invmp(this.m).mul(this.r2))._forceRed(this)}}("undefined"==typeof e||e,this)},{buffer:103}],95:[function(t,e,r){"use strict";e.exports=function(t){var e,r,n,i=t.length,a=0;for(e=0;e<i;++e)a+=t[e].length;var o=new Array(a),s=0;for(e=0;e<i;++e){var l=t[e],c=l.length;for(r=0;r<c;++r){var u=o[s++]=new Array(c-1),h=0;for(n=0;n<c;++n)n!==r&&(u[h++]=l[n]);if(1&r){var f=u[1];u[1]=u[0],u[0]=f}}}return o}},{}],96:[function(t,e,r){"use strict";e.exports=function(t,e,r){switch(arguments.length){case 1:return n=[],c(i=t,i,u,!0),n;case 2:return"function"==typeof e?c(t,t,e,!0):function(t,e){return n=[],c(t,e,u,!1),n}(t,e);case 3:return c(t,e,r,!1);default:throw new Error("box-intersect: Invalid arguments")}var i};var n,i=t("typedarray-pool"),a=t("./lib/sweep"),o=t("./lib/intersect");function s(t,e){for(var r=0;r<t;++r)if(!(e[r]<=e[r+t]))return!0;return!1}function l(t,e,r,n){for(var i=0,a=0,o=0,l=t.length;o<l;++o){var c=t[o];if(!s(e,c)){for(var u=0;u<2*e;++u)r[i++]=c[u];n[a++]=o}}return a}function c(t,e,r,n){var s=t.length,c=e.length;if(!(s<=0||c<=0)){var u=t[0].length>>>1;if(!(u<=0)){var h,f=i.mallocDouble(2*u*s),p=i.mallocInt32(s);if((s=l(t,u,f,p))>0){if(1===u&&n)a.init(s),h=a.sweepComplete(u,r,0,s,f,p,0,s,f,p);else{var d=i.mallocDouble(2*u*c),g=i.mallocInt32(c);(c=l(e,u,d,g))>0&&(a.init(s+c),h=1===u?a.sweepBipartite(u,r,0,s,f,p,0,c,d,g):o(u,r,n,s,f,p,c,d,g),i.free(d),i.free(g))}i.free(f),i.free(p)}return h}}}function u(t,e){n.push([t,e])}},{"./lib/intersect":98,"./lib/sweep":102,"typedarray-pool":545}],97:[function(t,e,r){"use strict";var n="d",i="ax",a="vv",o="fp",s="es",l="rs",c="re",u="rb",h="ri",f="rp",p="bs",d="be",g="bb",v="bi",m="bp",y="rv",x="Q",b=[n,i,a,l,c,u,h,p,d,g,v];function _(t){var e="bruteForce"+(t?"Full":"Partial"),r=[],_=b.slice();t||_.splice(3,0,o);var w=["function "+e+"("+_.join()+"){"];function k(e,o){var _=function(t,e,r){var o="bruteForce"+(t?"Red":"Blue")+(e?"Flip":"")+(r?"Full":""),_=["function ",o,"(",b.join(),"){","var ",s,"=2*",n,";"],w="for(var i="+l+","+f+"="+s+"*"+l+";i<"+c+";++i,"+f+"+="+s+"){var x0="+u+"["+i+"+"+f+"],x1="+u+"["+i+"+"+f+"+"+n+"],xi="+h+"[i];",k="for(var j="+p+","+m+"="+s+"*"+p+";j<"+d+";++j,"+m+"+="+s+"){var y0="+g+"["+i+"+"+m+"],"+(r?"y1="+g+"["+i+"+"+m+"+"+n+"],":"")+"yi="+v+"[j];";return t?_.push(w,x,":",k):_.push(k,x,":",w),r?_.push("if(y1<x0||x1<y0)continue;"):e?_.push("if(y0<=x0||x1<y0)continue;"):_.push("if(y0<x0||x1<y0)continue;"),_.push("for(var k="+i+"+1;k<"+n+";++k){var r0="+u+"[k+"+f+"],r1="+u+"[k+"+n+"+"+f+"],b0="+g+"[k+"+m+"],b1="+g+"[k+"+n+"+"+m+"];if(r1<b0||b1<r0)continue "+x+";}var "+y+"="+a+"("),e?_.push("yi,xi"):_.push("xi,yi"),_.push(");if("+y+"!==void 0)return "+y+";}}}"),{name:o,code:_.join("")}}(e,o,t);r.push(_.code),w.push("return "+_.name+"("+b.join()+");")}w.push("if("+c+"-"+l+">"+d+"-"+p+"){"),t?(k(!0,!1),w.push("}else{"),k(!1,!1)):(w.push("if("+o+"){"),k(!0,!0),w.push("}else{"),k(!0,!1),w.push("}}else{if("+o+"){"),k(!1,!0),w.push("}else{"),k(!1,!1),w.push("}")),w.push("}}return "+e);var T=r.join("")+w.join("");return new Function(T)()}r.partial=_(!1),r.full=_(!0)},{}],98:[function(t,e,r){"use strict";e.exports=function(t,e,r,a,u,S,E,C,L){!function(t,e){var r=8*i.log2(e+1)*(t+1)|0,a=i.nextPow2(b*r);w.length<a&&(n.free(w),w=n.mallocInt32(a));var o=i.nextPow2(_*r);k<o&&(n.free(k),k=n.mallocDouble(o))}(t,a+E);var P,O=0,z=2*t;T(O++,0,0,a,0,E,r?16:0,-1/0,1/0),r||T(O++,0,0,E,0,a,1,-1/0,1/0);for(;O>0;){var I=(O-=1)*b,D=w[I],R=w[I+1],F=w[I+2],B=w[I+3],N=w[I+4],j=w[I+5],U=O*_,V=k[U],q=k[U+1],H=1&j,G=!!(16&j),Y=u,W=S,X=C,Z=L;if(H&&(Y=C,W=L,X=u,Z=S),!(2&j&&(F=v(t,D,R,F,Y,W,q),R>=F)||4&j&&(R=m(t,D,R,F,Y,W,V))>=F)){var J=F-R,K=N-B;if(G){if(t*J*(J+K)<p){if(void 0!==(P=l.scanComplete(t,D,e,R,F,Y,W,B,N,X,Z)))return P;continue}}else{if(t*Math.min(J,K)<h){if(void 0!==(P=o(t,D,e,H,R,F,Y,W,B,N,X,Z)))return P;continue}if(t*J*K<f){if(void 0!==(P=l.scanBipartite(t,D,e,H,R,F,Y,W,B,N,X,Z)))return P;continue}}var $=d(t,D,R,F,Y,W,V,q);if(R<$)if(t*($-R)<h){if(void 0!==(P=s(t,D+1,e,R,$,Y,W,B,N,X,Z)))return P}else if(D===t-2){if(void 0!==(P=H?l.sweepBipartite(t,e,B,N,X,Z,R,$,Y,W):l.sweepBipartite(t,e,R,$,Y,W,B,N,X,Z)))return P}else T(O++,D+1,R,$,B,N,H,-1/0,1/0),T(O++,D+1,B,N,R,$,1^H,-1/0,1/0);if($<F){var Q=c(t,D,B,N,X,Z),tt=X[z*Q+D],et=g(t,D,Q,N,X,Z,tt);if(et<N&&T(O++,D,$,F,et,N,(4|H)+(G?16:0),tt,q),B<Q&&T(O++,D,$,F,B,Q,(2|H)+(G?16:0),V,tt),Q+1===et){if(void 0!==(P=G?M(t,D,e,$,F,Y,W,Q,X,Z[Q]):A(t,D,e,H,$,F,Y,W,Q,X,Z[Q])))return P}else if(Q<et){var rt;if(G){if(rt=y(t,D,$,F,Y,W,tt),$<rt){var nt=g(t,D,$,rt,Y,W,tt);if(D===t-2){if($<nt&&void 0!==(P=l.sweepComplete(t,e,$,nt,Y,W,Q,et,X,Z)))return P;if(nt<rt&&void 0!==(P=l.sweepBipartite(t,e,nt,rt,Y,W,Q,et,X,Z)))return P}else $<nt&&T(O++,D+1,$,nt,Q,et,16,-1/0,1/0),nt<rt&&(T(O++,D+1,nt,rt,Q,et,0,-1/0,1/0),T(O++,D+1,Q,et,nt,rt,1,-1/0,1/0))}}else rt=H?x(t,D,$,F,Y,W,tt):y(t,D,$,F,Y,W,tt),$<rt&&(D===t-2?P=H?l.sweepBipartite(t,e,Q,et,X,Z,$,rt,Y,W):l.sweepBipartite(t,e,$,rt,Y,W,Q,et,X,Z):(T(O++,D+1,$,rt,Q,et,H,-1/0,1/0),T(O++,D+1,Q,et,$,rt,1^H,-1/0,1/0)))}}}}};var n=t("typedarray-pool"),i=t("bit-twiddle"),a=t("./brute"),o=a.partial,s=a.full,l=t("./sweep"),c=t("./median"),u=t("./partition"),h=128,f=1<<22,p=1<<22,d=u("!(lo>=p0)&&!(p1>=hi)",["p0","p1"]),g=u("lo===p0",["p0"]),v=u("lo<p0",["p0"]),m=u("hi<=p0",["p0"]),y=u("lo<=p0&&p0<=hi",["p0"]),x=u("lo<p0&&p0<=hi",["p0"]),b=6,_=2,w=n.mallocInt32(1024),k=n.mallocDouble(1024);function T(t,e,r,n,i,a,o,s,l){var c=b*t;w[c]=e,w[c+1]=r,w[c+2]=n,w[c+3]=i,w[c+4]=a,w[c+5]=o;var u=_*t;k[u]=s,k[u+1]=l}function A(t,e,r,n,i,a,o,s,l,c,u){var h=2*t,f=l*h,p=c[f+e];t:for(var d=i,g=i*h;d<a;++d,g+=h){var v=o[g+e],m=o[g+e+t];if(!(p<v||m<p)&&(!n||p!==v)){for(var y,x=s[d],b=e+1;b<t;++b){v=o[g+b],m=o[g+b+t];var _=c[f+b],w=c[f+b+t];if(m<_||w<v)continue t}if(void 0!==(y=n?r(u,x):r(x,u)))return y}}}function M(t,e,r,n,i,a,o,s,l,c){var u=2*t,h=s*u,f=l[h+e];t:for(var p=n,d=n*u;p<i;++p,d+=u){var g=o[p];if(g!==c){var v=a[d+e],m=a[d+e+t];if(!(f<v||m<f)){for(var y=e+1;y<t;++y){v=a[d+y],m=a[d+y+t];var x=l[h+y],b=l[h+y+t];if(m<x||b<v)continue t}var _=r(g,c);if(void 0!==_)return _}}}}},{"./brute":97,"./median":99,"./partition":100,"./sweep":102,"bit-twiddle":92,"typedarray-pool":545}],99:[function(t,e,r){"use strict";e.exports=function(t,e,r,o,s,l){if(o<=r+1)return r;var c=r,u=o,h=o+r>>>1,f=2*t,p=h,d=s[f*h+e];for(;c<u;){if(u-c<i){a(t,e,c,u,s,l),d=s[f*h+e];break}var g=u-c,v=Math.random()*g+c|0,m=s[f*v+e],y=Math.random()*g+c|0,x=s[f*y+e],b=Math.random()*g+c|0,_=s[f*b+e];m<=x?_>=x?(p=y,d=x):m>=_?(p=v,d=m):(p=b,d=_):x>=_?(p=y,d=x):_>=m?(p=v,d=m):(p=b,d=_);for(var w=f*(u-1),k=f*p,T=0;T<f;++T,++w,++k){var A=s[w];s[w]=s[k],s[k]=A}var M=l[u-1];l[u-1]=l[p],l[p]=M,p=n(t,e,c,u-1,s,l,d);for(var w=f*(u-1),k=f*p,T=0;T<f;++T,++w,++k){var A=s[w];s[w]=s[k],s[k]=A}var M=l[u-1];if(l[u-1]=l[p],l[p]=M,h<p){for(u=p-1;c<u&&s[f*(u-1)+e]===d;)u-=1;u+=1}else{if(!(p<h))break;for(c=p+1;c<u&&s[f*c+e]===d;)c+=1}}return n(t,e,r,h,s,l,s[f*h+e])};var n=t("./partition")("lo<p0",["p0"]),i=8;function a(t,e,r,n,i,a){for(var o=2*t,s=o*(r+1)+e,l=r+1;l<n;++l,s+=o)for(var c=i[s],u=l,h=o*(l-1);u>r&&i[h+e]>c;--u,h-=o){for(var f=h,p=h+o,d=0;d<o;++d,++f,++p){var g=i[f];i[f]=i[p],i[p]=g}var v=a[u];a[u]=a[u-1],a[u-1]=v}}},{"./partition":100}],100:[function(t,e,r){"use strict";e.exports=function(t,e){var r="abcdef".split("").concat(e),i=[];t.indexOf("lo")>=0&&i.push("lo=e[k+n]");t.indexOf("hi")>=0&&i.push("hi=e[k+o]");return r.push(n.replace("_",i.join()).replace("$",t)),Function.apply(void 0,r)};var n="for(var j=2*a,k=j*c,l=k,m=c,n=b,o=a+b,p=c;d>p;++p,k+=j){var _;if($)if(m===p)m+=1,l+=j;else{for(var s=0;j>s;++s){var t=e[k+s];e[k+s]=e[l],e[l++]=t}var u=f[p];f[p]=f[m],f[m++]=u}}return m"},{}],101:[function(t,e,r){"use strict";e.exports=function(t,e){e<=4*n?i(0,e-1,t):function t(e,r,h){var f=(r-e+1)/6|0,p=e+f,d=r-f,g=e+r>>1,v=g-f,m=g+f,y=p,x=v,b=g,_=m,w=d,k=e+1,T=r-1,A=0;c(y,x,h)&&(A=y,y=x,x=A);c(_,w,h)&&(A=_,_=w,w=A);c(y,b,h)&&(A=y,y=b,b=A);c(x,b,h)&&(A=x,x=b,b=A);c(y,_,h)&&(A=y,y=_,_=A);c(b,_,h)&&(A=b,b=_,_=A);c(x,w,h)&&(A=x,x=w,w=A);c(x,b,h)&&(A=x,x=b,b=A);c(_,w,h)&&(A=_,_=w,w=A);var M=h[2*x];var S=h[2*x+1];var E=h[2*_];var C=h[2*_+1];var L=2*y;var P=2*b;var O=2*w;var z=2*p;var I=2*g;var D=2*d;for(var R=0;R<2;++R){var F=h[L+R],B=h[P+R],N=h[O+R];h[z+R]=F,h[I+R]=B,h[D+R]=N}o(v,e,h);o(m,r,h);for(var j=k;j<=T;++j)if(u(j,M,S,h))j!==k&&a(j,k,h),++k;else if(!u(j,E,C,h))for(;;){if(u(T,E,C,h)){u(T,M,S,h)?(s(j,k,T,h),++k,--T):(a(j,T,h),--T);break}if(--T<j)break}l(e,k-1,M,S,h);l(r,T+1,E,C,h);k-2-e<=n?i(e,k-2,h):t(e,k-2,h);r-(T+2)<=n?i(T+2,r,h):t(T+2,r,h);T-k<=n?i(k,T,h):t(k,T,h)}(0,e-1,t)};var n=32;function i(t,e,r){for(var n=2*(t+1),i=t+1;i<=e;++i){for(var a=r[n++],o=r[n++],s=i,l=n-2;s-- >t;){var c=r[l-2],u=r[l-1];if(c<a)break;if(c===a&&u<o)break;r[l]=c,r[l+1]=u,l-=2}r[l]=a,r[l+1]=o}}function a(t,e,r){e*=2;var n=r[t*=2],i=r[t+1];r[t]=r[e],r[t+1]=r[e+1],r[e]=n,r[e+1]=i}function o(t,e,r){e*=2,r[t*=2]=r[e],r[t+1]=r[e+1]}function s(t,e,r,n){e*=2,r*=2;var i=n[t*=2],a=n[t+1];n[t]=n[e],n[t+1]=n[e+1],n[e]=n[r],n[e+1]=n[r+1],n[r]=i,n[r+1]=a}function l(t,e,r,n,i){e*=2,i[t*=2]=i[e],i[e]=r,i[t+1]=i[e+1],i[e+1]=n}function c(t,e,r){e*=2;var n=r[t*=2],i=r[e];return!(n<i)&&(n!==i||r[t+1]>r[e+1])}function u(t,e,r,n){var i=n[t*=2];return i<e||i===e&&n[t+1]<r}},{}],102:[function(t,e,r){"use strict";e.exports={init:function(t){var e=i.nextPow2(t);s.length<e&&(n.free(s),s=n.mallocInt32(e));l.length<e&&(n.free(l),l=n.mallocInt32(e));c.length<e&&(n.free(c),c=n.mallocInt32(e));u.length<e&&(n.free(u),u=n.mallocInt32(e));h.length<e&&(n.free(h),h=n.mallocInt32(e));f.length<e&&(n.free(f),f=n.mallocInt32(e));var r=8*e;p.length<r&&(n.free(p),p=n.mallocDouble(r))},sweepBipartite:function(t,e,r,n,i,h,f,v,m,y){for(var x=0,b=2*t,_=t-1,w=b-1,k=r;k<n;++k){var T=h[k],A=b*k;p[x++]=i[A+_],p[x++]=-(T+1),p[x++]=i[A+w],p[x++]=T}for(var k=f;k<v;++k){var T=y[k]+o,M=b*k;p[x++]=m[M+_],p[x++]=-T,p[x++]=m[M+w],p[x++]=T}var S=x>>>1;a(p,S);for(var E=0,C=0,k=0;k<S;++k){var L=0|p[2*k+1];if(L>=o)d(c,u,C--,L=L-o|0);else if(L>=0)d(s,l,E--,L);else if(L<=-o){L=-L-o|0;for(var P=0;P<E;++P){var O=e(s[P],L);if(void 0!==O)return O}g(c,u,C++,L)}else{L=-L-1|0;for(var P=0;P<C;++P){var O=e(L,c[P]);if(void 0!==O)return O}g(s,l,E++,L)}}},sweepComplete:function(t,e,r,n,i,o,v,m,y,x){for(var b=0,_=2*t,w=t-1,k=_-1,T=r;T<n;++T){var A=o[T]+1<<1,M=_*T;p[b++]=i[M+w],p[b++]=-A,p[b++]=i[M+k],p[b++]=A}for(var T=v;T<m;++T){var A=x[T]+1<<1,S=_*T;p[b++]=y[S+w],p[b++]=1|-A,p[b++]=y[S+k],p[b++]=1|A}var E=b>>>1;a(p,E);for(var C=0,L=0,P=0,T=0;T<E;++T){var O=0|p[2*T+1],z=1&O;if(T<E-1&&O>>1==p[2*T+3]>>1&&(z=2,T+=1),O<0){for(var I=-(O>>1)-1,D=0;D<P;++D){var R=e(h[D],I);if(void 0!==R)return R}if(0!==z)for(var D=0;D<C;++D){var R=e(s[D],I);if(void 0!==R)return R}if(1!==z)for(var D=0;D<L;++D){var R=e(c[D],I);if(void 0!==R)return R}0===z?g(s,l,C++,I):1===z?g(c,u,L++,I):2===z&&g(h,f,P++,I)}else{var I=(O>>1)-1;0===z?d(s,l,C--,I):1===z?d(c,u,L--,I):2===z&&d(h,f,P--,I)}}},scanBipartite:function(t,e,r,n,i,c,u,h,f,v,m,y){var x=0,b=2*t,_=e,w=e+t,k=1,T=1;n?T=o:k=o;for(var A=i;A<c;++A){var M=A+k,S=b*A;p[x++]=u[S+_],p[x++]=-M,p[x++]=u[S+w],p[x++]=M}for(var A=f;A<v;++A){var M=A+T,E=b*A;p[x++]=m[E+_],p[x++]=-M}var C=x>>>1;a(p,C);for(var L=0,A=0;A<C;++A){var P=0|p[2*A+1];if(P<0){var M=-P,O=!1;if(M>=o?(O=!n,M-=o):(O=!!n,M-=1),O)g(s,l,L++,M);else{var z=y[M],I=b*M,D=m[I+e+1],R=m[I+e+1+t];t:for(var F=0;F<L;++F){var B=s[F],N=b*B;if(!(R<u[N+e+1]||u[N+e+1+t]<D)){for(var j=e+2;j<t;++j)if(m[I+j+t]<u[N+j]||u[N+j+t]<m[I+j])continue t;var U,V=h[B];if(void 0!==(U=n?r(z,V):r(V,z)))return U}}}}else d(s,l,L--,P-k)}},scanComplete:function(t,e,r,n,i,l,c,u,h,f,d){for(var g=0,v=2*t,m=e,y=e+t,x=n;x<i;++x){var b=x+o,_=v*x;p[g++]=l[_+m],p[g++]=-b,p[g++]=l[_+y],p[g++]=b}for(var x=u;x<h;++x){var b=x+1,w=v*x;p[g++]=f[w+m],p[g++]=-b}var k=g>>>1;a(p,k);for(var T=0,x=0;x<k;++x){var A=0|p[2*x+1];if(A<0){var b=-A;if(b>=o)s[T++]=b-o;else{var M=d[b-=1],S=v*b,E=f[S+e+1],C=f[S+e+1+t];t:for(var L=0;L<T;++L){var P=s[L],O=c[P];if(O===M)break;var z=v*P;if(!(C<l[z+e+1]||l[z+e+1+t]<E)){for(var I=e+2;I<t;++I)if(f[S+I+t]<l[z+I]||l[z+I+t]<f[S+I])continue t;var D=r(O,M);if(void 0!==D)return D}}}}else{for(var b=A-o,L=T-1;L>=0;--L)if(s[L]===b){for(var I=L+1;I<T;++I)s[I-1]=s[I];break}--T}}}};var n=t("typedarray-pool"),i=t("bit-twiddle"),a=t("./sort"),o=1<<28,s=n.mallocInt32(1024),l=n.mallocInt32(1024),c=n.mallocInt32(1024),u=n.mallocInt32(1024),h=n.mallocInt32(1024),f=n.mallocInt32(1024),p=n.mallocDouble(8192);function d(t,e,r,n){var i=e[n],a=t[r-1];t[i]=a,e[a]=i}function g(t,e,r,n){t[r]=n,e[n]=r}},{"./sort":101,"bit-twiddle":92,"typedarray-pool":545}],103:[function(t,e,r){},{}],104:[function(t,e,r){var n=Object.create||function(t){var e=function(){};return e.prototype=t,new e},i=Object.keys||function(t){var e=[];for(var r in t)Object.prototype.hasOwnProperty.call(t,r)&&e.push(r);return r},a=Function.prototype.bind||function(t){var e=this;return function(){return e.apply(t,arguments)}};function o(){this._events&&Object.prototype.hasOwnProperty.call(this,"_events")||(this._events=n(null),this._eventsCount=0),this._maxListeners=this._maxListeners||void 0}e.exports=o,o.EventEmitter=o,o.prototype._events=void 0,o.prototype._maxListeners=void 0;var s,l=10;try{var c={};Object.defineProperty&&Object.defineProperty(c,"x",{value:0}),s=0===c.x}catch(t){s=!1}function u(t){return void 0===t._maxListeners?o.defaultMaxListeners:t._maxListeners}function h(t,e,r,i){var a,o,s;if("function"!=typeof r)throw new TypeError('"listener" argument must be a function');if((o=t._events)?(o.newListener&&(t.emit("newListener",e,r.listener?r.listener:r),o=t._events),s=o[e]):(o=t._events=n(null),t._eventsCount=0),s){if("function"==typeof s?s=o[e]=i?[r,s]:[s,r]:i?s.unshift(r):s.push(r),!s.warned&&(a=u(t))&&a>0&&s.length>a){s.warned=!0;var l=new Error("Possible EventEmitter memory leak detected. "+s.length+' "'+String(e)+'" listeners added. Use emitter.setMaxListeners() to increase limit.');l.name="MaxListenersExceededWarning",l.emitter=t,l.type=e,l.count=s.length,"object"==typeof console&&console.warn&&console.warn("%s: %s",l.name,l.message)}}else s=o[e]=r,++t._eventsCount;return t}function f(){if(!this.fired)switch(this.target.removeListener(this.type,this.wrapFn),this.fired=!0,arguments.length){case 0:return this.listener.call(this.target);case 1:return this.listener.call(this.target,arguments[0]);case 2:return this.listener.call(this.target,arguments[0],arguments[1]);case 3:return this.listener.call(this.target,arguments[0],arguments[1],arguments[2]);default:for(var t=new Array(arguments.length),e=0;e<t.length;++e)t[e]=arguments[e];this.listener.apply(this.target,t)}}function p(t,e,r){var n={fired:!1,wrapFn:void 0,target:t,type:e,listener:r},i=a.call(f,n);return i.listener=r,n.wrapFn=i,i}function d(t,e,r){var n=t._events;if(!n)return[];var i=n[e];return i?"function"==typeof i?r?[i.listener||i]:[i]:r?function(t){for(var e=new Array(t.length),r=0;r<e.length;++r)e[r]=t[r].listener||t[r];return e}(i):v(i,i.length):[]}function g(t){var e=this._events;if(e){var r=e[t];if("function"==typeof r)return 1;if(r)return r.length}return 0}function v(t,e){for(var r=new Array(e),n=0;n<e;++n)r[n]=t[n];return r}s?Object.defineProperty(o,"defaultMaxListeners",{enumerable:!0,get:function(){return l},set:function(t){if("number"!=typeof t||t<0||t!=t)throw new TypeError('"defaultMaxListeners" must be a positive number');l=t}}):o.defaultMaxListeners=l,o.prototype.setMaxListeners=function(t){if("number"!=typeof t||t<0||isNaN(t))throw new TypeError('"n" argument must be a positive number');return this._maxListeners=t,this},o.prototype.getMaxListeners=function(){return u(this)},o.prototype.emit=function(t){var e,r,n,i,a,o,s="error"===t;if(o=this._events)s=s&&null==o.error;else if(!s)return!1;if(s){if(arguments.length>1&&(e=arguments[1]),e instanceof Error)throw e;var l=new Error('Unhandled "error" event. ('+e+")");throw l.context=e,l}if(!(r=o[t]))return!1;var c="function"==typeof r;switch(n=arguments.length){case 1:!function(t,e,r){if(e)t.call(r);else for(var n=t.length,i=v(t,n),a=0;a<n;++a)i[a].call(r)}(r,c,this);break;case 2:!function(t,e,r,n){if(e)t.call(r,n);else for(var i=t.length,a=v(t,i),o=0;o<i;++o)a[o].call(r,n)}(r,c,this,arguments[1]);break;case 3:!function(t,e,r,n,i){if(e)t.call(r,n,i);else for(var a=t.length,o=v(t,a),s=0;s<a;++s)o[s].call(r,n,i)}(r,c,this,arguments[1],arguments[2]);break;case 4:!function(t,e,r,n,i,a){if(e)t.call(r,n,i,a);else for(var o=t.length,s=v(t,o),l=0;l<o;++l)s[l].call(r,n,i,a)}(r,c,this,arguments[1],arguments[2],arguments[3]);break;default:for(i=new Array(n-1),a=1;a<n;a++)i[a-1]=arguments[a];!function(t,e,r,n){if(e)t.apply(r,n);else for(var i=t.length,a=v(t,i),o=0;o<i;++o)a[o].apply(r,n)}(r,c,this,i)}return!0},o.prototype.addListener=function(t,e){return h(this,t,e,!1)},o.prototype.on=o.prototype.addListener,o.prototype.prependListener=function(t,e){return h(this,t,e,!0)},o.prototype.once=function(t,e){if("function"!=typeof e)throw new TypeError('"listener" argument must be a function');return this.on(t,p(this,t,e)),this},o.prototype.prependOnceListener=function(t,e){if("function"!=typeof e)throw new TypeError('"listener" argument must be a function');return this.prependListener(t,p(this,t,e)),this},o.prototype.removeListener=function(t,e){var r,i,a,o,s;if("function"!=typeof e)throw new TypeError('"listener" argument must be a function');if(!(i=this._events))return this;if(!(r=i[t]))return this;if(r===e||r.listener===e)0==--this._eventsCount?this._events=n(null):(delete i[t],i.removeListener&&this.emit("removeListener",t,r.listener||e));else if("function"!=typeof r){for(a=-1,o=r.length-1;o>=0;o--)if(r[o]===e||r[o].listener===e){s=r[o].listener,a=o;break}if(a<0)return this;0===a?r.shift():function(t,e){for(var r=e,n=r+1,i=t.length;n<i;r+=1,n+=1)t[r]=t[n];t.pop()}(r,a),1===r.length&&(i[t]=r[0]),i.removeListener&&this.emit("removeListener",t,s||e)}return this},o.prototype.removeAllListeners=function(t){var e,r,a;if(!(r=this._events))return this;if(!r.removeListener)return 0===arguments.length?(this._events=n(null),this._eventsCount=0):r[t]&&(0==--this._eventsCount?this._events=n(null):delete r[t]),this;if(0===arguments.length){var o,s=i(r);for(a=0;a<s.length;++a)"removeListener"!==(o=s[a])&&this.removeAllListeners(o);return this.removeAllListeners("removeListener"),this._events=n(null),this._eventsCount=0,this}if("function"==typeof(e=r[t]))this.removeListener(t,e);else if(e)for(a=e.length-1;a>=0;a--)this.removeListener(t,e[a]);return this},o.prototype.listeners=function(t){return d(this,t,!0)},o.prototype.rawListeners=function(t){return d(this,t,!1)},o.listenerCount=function(t,e){return"function"==typeof t.listenerCount?t.listenerCount(e):g.call(t,e)},o.prototype.listenerCount=g,o.prototype.eventNames=function(){return this._eventsCount>0?Reflect.ownKeys(this._events):[]}},{}],105:[function(t,e,r){(function(e){"use strict";var n=t("base64-js"),i=t("ieee754");r.Buffer=e,r.SlowBuffer=function(t){+t!=t&&(t=0);return e.alloc(+t)},r.INSPECT_MAX_BYTES=50;var a=2147483647;function o(t){if(t>a)throw new RangeError('The value "'+t+'" is invalid for option "size"');var r=new Uint8Array(t);return r.__proto__=e.prototype,r}function e(t,e,r){if("number"==typeof t){if("string"==typeof e)throw new TypeError('The "string" argument must be of type string. Received type number');return c(t)}return s(t,e,r)}function s(t,r,n){if("string"==typeof t)return function(t,r){"string"==typeof r&&""!==r||(r="utf8");if(!e.isEncoding(r))throw new TypeError("Unknown encoding: "+r);var n=0|f(t,r),i=o(n),a=i.write(t,r);a!==n&&(i=i.slice(0,a));return i}(t,r);if(ArrayBuffer.isView(t))return u(t);if(null==t)throw TypeError("The first argument must be one of type string, Buffer, ArrayBuffer, Array, or Array-like Object. Received type "+typeof t);if(N(t,ArrayBuffer)||t&&N(t.buffer,ArrayBuffer))return function(t,r,n){if(r<0||t.byteLength<r)throw new RangeError('"offset" is outside of buffer bounds');if(t.byteLength<r+(n||0))throw new RangeError('"length" is outside of buffer bounds');var i;i=void 0===r&&void 0===n?new Uint8Array(t):void 0===n?new Uint8Array(t,r):new Uint8Array(t,r,n);return i.__proto__=e.prototype,i}(t,r,n);if("number"==typeof t)throw new TypeError('The "value" argument must not be of type number. Received type number');var i=t.valueOf&&t.valueOf();if(null!=i&&i!==t)return e.from(i,r,n);var a=function(t){if(e.isBuffer(t)){var r=0|h(t.length),n=o(r);return 0===n.length?n:(t.copy(n,0,0,r),n)}if(void 0!==t.length)return"number"!=typeof t.length||j(t.length)?o(0):u(t);if("Buffer"===t.type&&Array.isArray(t.data))return u(t.data)}(t);if(a)return a;if("undefined"!=typeof Symbol&&null!=Symbol.toPrimitive&&"function"==typeof t[Symbol.toPrimitive])return e.from(t[Symbol.toPrimitive]("string"),r,n);throw new TypeError("The first argument must be one of type string, Buffer, ArrayBuffer, Array, or Array-like Object. Received type "+typeof t)}function l(t){if("number"!=typeof t)throw new TypeError('"size" argument must be of type number');if(t<0)throw new RangeError('The value "'+t+'" is invalid for option "size"')}function c(t){return l(t),o(t<0?0:0|h(t))}function u(t){for(var e=t.length<0?0:0|h(t.length),r=o(e),n=0;n<e;n+=1)r[n]=255&t[n];return r}function h(t){if(t>=a)throw new RangeError("Attempt to allocate Buffer larger than maximum size: 0x"+a.toString(16)+" bytes");return 0|t}function f(t,r){if(e.isBuffer(t))return t.length;if(ArrayBuffer.isView(t)||N(t,ArrayBuffer))return t.byteLength;if("string"!=typeof t)throw new TypeError('The "string" argument must be one of type string, Buffer, or ArrayBuffer. Received type '+typeof t);var n=t.length,i=arguments.length>2&&!0===arguments[2];if(!i&&0===n)return 0;for(var a=!1;;)switch(r){case"ascii":case"latin1":case"binary":return n;case"utf8":case"utf-8":return R(t).length;case"ucs2":case"ucs-2":case"utf16le":case"utf-16le":return 2*n;case"hex":return n>>>1;case"base64":return F(t).length;default:if(a)return i?-1:R(t).length;r=(""+r).toLowerCase(),a=!0}}function p(t,e,r){var n=t[e];t[e]=t[r],t[r]=n}function d(t,r,n,i,a){if(0===t.length)return-1;if("string"==typeof n?(i=n,n=0):n>2147483647?n=2147483647:n<-2147483648&&(n=-2147483648),j(n=+n)&&(n=a?0:t.length-1),n<0&&(n=t.length+n),n>=t.length){if(a)return-1;n=t.length-1}else if(n<0){if(!a)return-1;n=0}if("string"==typeof r&&(r=e.from(r,i)),e.isBuffer(r))return 0===r.length?-1:g(t,r,n,i,a);if("number"==typeof r)return r&=255,"function"==typeof Uint8Array.prototype.indexOf?a?Uint8Array.prototype.indexOf.call(t,r,n):Uint8Array.prototype.lastIndexOf.call(t,r,n):g(t,[r],n,i,a);throw new TypeError("val must be string, number or Buffer")}function g(t,e,r,n,i){var a,o=1,s=t.length,l=e.length;if(void 0!==n&&("ucs2"===(n=String(n).toLowerCase())||"ucs-2"===n||"utf16le"===n||"utf-16le"===n)){if(t.length<2||e.length<2)return-1;o=2,s/=2,l/=2,r/=2}function c(t,e){return 1===o?t[e]:t.readUInt16BE(e*o)}if(i){var u=-1;for(a=r;a<s;a++)if(c(t,a)===c(e,-1===u?0:a-u)){if(-1===u&&(u=a),a-u+1===l)return u*o}else-1!==u&&(a-=a-u),u=-1}else for(r+l>s&&(r=s-l),a=r;a>=0;a--){for(var h=!0,f=0;f<l;f++)if(c(t,a+f)!==c(e,f)){h=!1;break}if(h)return a}return-1}function v(t,e,r,n){r=Number(r)||0;var i=t.length-r;n?(n=Number(n))>i&&(n=i):n=i;var a=e.length;n>a/2&&(n=a/2);for(var o=0;o<n;++o){var s=parseInt(e.substr(2*o,2),16);if(j(s))return o;t[r+o]=s}return o}function m(t,e,r,n){return B(R(e,t.length-r),t,r,n)}function y(t,e,r,n){return B(function(t){for(var e=[],r=0;r<t.length;++r)e.push(255&t.charCodeAt(r));return e}(e),t,r,n)}function x(t,e,r,n){return y(t,e,r,n)}function b(t,e,r,n){return B(F(e),t,r,n)}function _(t,e,r,n){return B(function(t,e){for(var r,n,i,a=[],o=0;o<t.length&&!((e-=2)<0);++o)r=t.charCodeAt(o),n=r>>8,i=r%256,a.push(i),a.push(n);return a}(e,t.length-r),t,r,n)}function w(t,e,r){return 0===e&&r===t.length?n.fromByteArray(t):n.fromByteArray(t.slice(e,r))}function k(t,e,r){r=Math.min(t.length,r);for(var n=[],i=e;i<r;){var a,o,s,l,c=t[i],u=null,h=c>239?4:c>223?3:c>191?2:1;if(i+h<=r)switch(h){case 1:c<128&&(u=c);break;case 2:128==(192&(a=t[i+1]))&&(l=(31&c)<<6|63&a)>127&&(u=l);break;case 3:a=t[i+1],o=t[i+2],128==(192&a)&&128==(192&o)&&(l=(15&c)<<12|(63&a)<<6|63&o)>2047&&(l<55296||l>57343)&&(u=l);break;case 4:a=t[i+1],o=t[i+2],s=t[i+3],128==(192&a)&&128==(192&o)&&128==(192&s)&&(l=(15&c)<<18|(63&a)<<12|(63&o)<<6|63&s)>65535&&l<1114112&&(u=l)}null===u?(u=65533,h=1):u>65535&&(u-=65536,n.push(u>>>10&1023|55296),u=56320|1023&u),n.push(u),i+=h}return function(t){var e=t.length;if(e<=T)return String.fromCharCode.apply(String,t);var r="",n=0;for(;n<e;)r+=String.fromCharCode.apply(String,t.slice(n,n+=T));return r}(n)}r.kMaxLength=a,e.TYPED_ARRAY_SUPPORT=function(){try{var t=new Uint8Array(1);return t.__proto__={__proto__:Uint8Array.prototype,foo:function(){return 42}},42===t.foo()}catch(t){return!1}}(),e.TYPED_ARRAY_SUPPORT||"undefined"==typeof console||"function"!=typeof console.error||console.error("This browser lacks typed array (Uint8Array) support which is required by `buffer` v5.x. Use `buffer` v4.x if you require old browser support."),Object.defineProperty(e.prototype,"parent",{enumerable:!0,get:function(){if(e.isBuffer(this))return this.buffer}}),Object.defineProperty(e.prototype,"offset",{enumerable:!0,get:function(){if(e.isBuffer(this))return this.byteOffset}}),"undefined"!=typeof Symbol&&null!=Symbol.species&&e[Symbol.species]===e&&Object.defineProperty(e,Symbol.species,{value:null,configurable:!0,enumerable:!1,writable:!1}),e.poolSize=8192,e.from=function(t,e,r){return s(t,e,r)},e.prototype.__proto__=Uint8Array.prototype,e.__proto__=Uint8Array,e.alloc=function(t,e,r){return function(t,e,r){return l(t),t<=0?o(t):void 0!==e?"string"==typeof r?o(t).fill(e,r):o(t).fill(e):o(t)}(t,e,r)},e.allocUnsafe=function(t){return c(t)},e.allocUnsafeSlow=function(t){return c(t)},e.isBuffer=function(t){return null!=t&&!0===t._isBuffer&&t!==e.prototype},e.compare=function(t,r){if(N(t,Uint8Array)&&(t=e.from(t,t.offset,t.byteLength)),N(r,Uint8Array)&&(r=e.from(r,r.offset,r.byteLength)),!e.isBuffer(t)||!e.isBuffer(r))throw new TypeError('The "buf1", "buf2" arguments must be one of type Buffer or Uint8Array');if(t===r)return 0;for(var n=t.length,i=r.length,a=0,o=Math.min(n,i);a<o;++a)if(t[a]!==r[a]){n=t[a],i=r[a];break}return n<i?-1:i<n?1:0},e.isEncoding=function(t){switch(String(t).toLowerCase()){case"hex":case"utf8":case"utf-8":case"ascii":case"latin1":case"binary":case"base64":case"ucs2":case"ucs-2":case"utf16le":case"utf-16le":return!0;default:return!1}},e.concat=function(t,r){if(!Array.isArray(t))throw new TypeError('"list" argument must be an Array of Buffers');if(0===t.length)return e.alloc(0);var n;if(void 0===r)for(r=0,n=0;n<t.length;++n)r+=t[n].length;var i=e.allocUnsafe(r),a=0;for(n=0;n<t.length;++n){var o=t[n];if(N(o,Uint8Array)&&(o=e.from(o)),!e.isBuffer(o))throw new TypeError('"list" argument must be an Array of Buffers');o.copy(i,a),a+=o.length}return i},e.byteLength=f,e.prototype._isBuffer=!0,e.prototype.swap16=function(){var t=this.length;if(t%2!=0)throw new RangeError("Buffer size must be a multiple of 16-bits");for(var e=0;e<t;e+=2)p(this,e,e+1);return this},e.prototype.swap32=function(){var t=this.length;if(t%4!=0)throw new RangeError("Buffer size must be a multiple of 32-bits");for(var e=0;e<t;e+=4)p(this,e,e+3),p(this,e+1,e+2);return this},e.prototype.swap64=function(){var t=this.length;if(t%8!=0)throw new RangeError("Buffer size must be a multiple of 64-bits");for(var e=0;e<t;e+=8)p(this,e,e+7),p(this,e+1,e+6),p(this,e+2,e+5),p(this,e+3,e+4);return this},e.prototype.toString=function(){var t=this.length;return 0===t?"":0===arguments.length?k(this,0,t):function(t,e,r){var n=!1;if((void 0===e||e<0)&&(e=0),e>this.length)return"";if((void 0===r||r>this.length)&&(r=this.length),r<=0)return"";if((r>>>=0)<=(e>>>=0))return"";for(t||(t="utf8");;)switch(t){case"hex":return S(this,e,r);case"utf8":case"utf-8":return k(this,e,r);case"ascii":return A(this,e,r);case"latin1":case"binary":return M(this,e,r);case"base64":return w(this,e,r);case"ucs2":case"ucs-2":case"utf16le":case"utf-16le":return E(this,e,r);default:if(n)throw new TypeError("Unknown encoding: "+t);t=(t+"").toLowerCase(),n=!0}}.apply(this,arguments)},e.prototype.toLocaleString=e.prototype.toString,e.prototype.equals=function(t){if(!e.isBuffer(t))throw new TypeError("Argument must be a Buffer");return this===t||0===e.compare(this,t)},e.prototype.inspect=function(){var t="",e=r.INSPECT_MAX_BYTES;return t=this.toString("hex",0,e).replace(/(.{2})/g,"$1 ").trim(),this.length>e&&(t+=" ... "),"<Buffer "+t+">"},e.prototype.compare=function(t,r,n,i,a){if(N(t,Uint8Array)&&(t=e.from(t,t.offset,t.byteLength)),!e.isBuffer(t))throw new TypeError('The "target" argument must be one of type Buffer or Uint8Array. Received type '+typeof t);if(void 0===r&&(r=0),void 0===n&&(n=t?t.length:0),void 0===i&&(i=0),void 0===a&&(a=this.length),r<0||n>t.length||i<0||a>this.length)throw new RangeError("out of range index");if(i>=a&&r>=n)return 0;if(i>=a)return-1;if(r>=n)return 1;if(this===t)return 0;for(var o=(a>>>=0)-(i>>>=0),s=(n>>>=0)-(r>>>=0),l=Math.min(o,s),c=this.slice(i,a),u=t.slice(r,n),h=0;h<l;++h)if(c[h]!==u[h]){o=c[h],s=u[h];break}return o<s?-1:s<o?1:0},e.prototype.includes=function(t,e,r){return-1!==this.indexOf(t,e,r)},e.prototype.indexOf=function(t,e,r){return d(this,t,e,r,!0)},e.prototype.lastIndexOf=function(t,e,r){return d(this,t,e,r,!1)},e.prototype.write=function(t,e,r,n){if(void 0===e)n="utf8",r=this.length,e=0;else if(void 0===r&&"string"==typeof e)n=e,r=this.length,e=0;else{if(!isFinite(e))throw new Error("Buffer.write(string, encoding, offset[, length]) is no longer supported");e>>>=0,isFinite(r)?(r>>>=0,void 0===n&&(n="utf8")):(n=r,r=void 0)}var i=this.length-e;if((void 0===r||r>i)&&(r=i),t.length>0&&(r<0||e<0)||e>this.length)throw new RangeError("Attempt to write outside buffer bounds");n||(n="utf8");for(var a=!1;;)switch(n){case"hex":return v(this,t,e,r);case"utf8":case"utf-8":return m(this,t,e,r);case"ascii":return y(this,t,e,r);case"latin1":case"binary":return x(this,t,e,r);case"base64":return b(this,t,e,r);case"ucs2":case"ucs-2":case"utf16le":case"utf-16le":return _(this,t,e,r);default:if(a)throw new TypeError("Unknown encoding: "+n);n=(""+n).toLowerCase(),a=!0}},e.prototype.toJSON=function(){return{type:"Buffer",data:Array.prototype.slice.call(this._arr||this,0)}};var T=4096;function A(t,e,r){var n="";r=Math.min(t.length,r);for(var i=e;i<r;++i)n+=String.fromCharCode(127&t[i]);return n}function M(t,e,r){var n="";r=Math.min(t.length,r);for(var i=e;i<r;++i)n+=String.fromCharCode(t[i]);return n}function S(t,e,r){var n=t.length;(!e||e<0)&&(e=0),(!r||r<0||r>n)&&(r=n);for(var i="",a=e;a<r;++a)i+=D(t[a]);return i}function E(t,e,r){for(var n=t.slice(e,r),i="",a=0;a<n.length;a+=2)i+=String.fromCharCode(n[a]+256*n[a+1]);return i}function C(t,e,r){if(t%1!=0||t<0)throw new RangeError("offset is not uint");if(t+e>r)throw new RangeError("Trying to access beyond buffer length")}function L(t,r,n,i,a,o){if(!e.isBuffer(t))throw new TypeError('"buffer" argument must be a Buffer instance');if(r>a||r<o)throw new RangeError('"value" argument is out of bounds');if(n+i>t.length)throw new RangeError("Index out of range")}function P(t,e,r,n,i,a){if(r+n>t.length)throw new RangeError("Index out of range");if(r<0)throw new RangeError("Index out of range")}function O(t,e,r,n,a){return e=+e,r>>>=0,a||P(t,0,r,4),i.write(t,e,r,n,23,4),r+4}function z(t,e,r,n,a){return e=+e,r>>>=0,a||P(t,0,r,8),i.write(t,e,r,n,52,8),r+8}e.prototype.slice=function(t,r){var n=this.length;(t=~~t)<0?(t+=n)<0&&(t=0):t>n&&(t=n),(r=void 0===r?n:~~r)<0?(r+=n)<0&&(r=0):r>n&&(r=n),r<t&&(r=t);var i=this.subarray(t,r);return i.__proto__=e.prototype,i},e.prototype.readUIntLE=function(t,e,r){t>>>=0,e>>>=0,r||C(t,e,this.length);for(var n=this[t],i=1,a=0;++a<e&&(i*=256);)n+=this[t+a]*i;return n},e.prototype.readUIntBE=function(t,e,r){t>>>=0,e>>>=0,r||C(t,e,this.length);for(var n=this[t+--e],i=1;e>0&&(i*=256);)n+=this[t+--e]*i;return n},e.prototype.readUInt8=function(t,e){return t>>>=0,e||C(t,1,this.length),this[t]},e.prototype.readUInt16LE=function(t,e){return t>>>=0,e||C(t,2,this.length),this[t]|this[t+1]<<8},e.prototype.readUInt16BE=function(t,e){return t>>>=0,e||C(t,2,this.length),this[t]<<8|this[t+1]},e.prototype.readUInt32LE=function(t,e){return t>>>=0,e||C(t,4,this.length),(this[t]|this[t+1]<<8|this[t+2]<<16)+16777216*this[t+3]},e.prototype.readUInt32BE=function(t,e){return t>>>=0,e||C(t,4,this.length),16777216*this[t]+(this[t+1]<<16|this[t+2]<<8|this[t+3])},e.prototype.readIntLE=function(t,e,r){t>>>=0,e>>>=0,r||C(t,e,this.length);for(var n=this[t],i=1,a=0;++a<e&&(i*=256);)n+=this[t+a]*i;return n>=(i*=128)&&(n-=Math.pow(2,8*e)),n},e.prototype.readIntBE=function(t,e,r){t>>>=0,e>>>=0,r||C(t,e,this.length);for(var n=e,i=1,a=this[t+--n];n>0&&(i*=256);)a+=this[t+--n]*i;return a>=(i*=128)&&(a-=Math.pow(2,8*e)),a},e.prototype.readInt8=function(t,e){return t>>>=0,e||C(t,1,this.length),128&this[t]?-1*(255-this[t]+1):this[t]},e.prototype.readInt16LE=function(t,e){t>>>=0,e||C(t,2,this.length);var r=this[t]|this[t+1]<<8;return 32768&r?4294901760|r:r},e.prototype.readInt16BE=function(t,e){t>>>=0,e||C(t,2,this.length);var r=this[t+1]|this[t]<<8;return 32768&r?4294901760|r:r},e.prototype.readInt32LE=function(t,e){return t>>>=0,e||C(t,4,this.length),this[t]|this[t+1]<<8|this[t+2]<<16|this[t+3]<<24},e.prototype.readInt32BE=function(t,e){return t>>>=0,e||C(t,4,this.length),this[t]<<24|this[t+1]<<16|this[t+2]<<8|this[t+3]},e.prototype.readFloatLE=function(t,e){return t>>>=0,e||C(t,4,this.length),i.read(this,t,!0,23,4)},e.prototype.readFloatBE=function(t,e){return t>>>=0,e||C(t,4,this.length),i.read(this,t,!1,23,4)},e.prototype.readDoubleLE=function(t,e){return t>>>=0,e||C(t,8,this.length),i.read(this,t,!0,52,8)},e.prototype.readDoubleBE=function(t,e){return t>>>=0,e||C(t,8,this.length),i.read(this,t,!1,52,8)},e.prototype.writeUIntLE=function(t,e,r,n){(t=+t,e>>>=0,r>>>=0,n)||L(this,t,e,r,Math.pow(2,8*r)-1,0);var i=1,a=0;for(this[e]=255&t;++a<r&&(i*=256);)this[e+a]=t/i&255;return e+r},e.prototype.writeUIntBE=function(t,e,r,n){(t=+t,e>>>=0,r>>>=0,n)||L(this,t,e,r,Math.pow(2,8*r)-1,0);var i=r-1,a=1;for(this[e+i]=255&t;--i>=0&&(a*=256);)this[e+i]=t/a&255;return e+r},e.prototype.writeUInt8=function(t,e,r){return t=+t,e>>>=0,r||L(this,t,e,1,255,0),this[e]=255&t,e+1},e.prototype.writeUInt16LE=function(t,e,r){return t=+t,e>>>=0,r||L(this,t,e,2,65535,0),this[e]=255&t,this[e+1]=t>>>8,e+2},e.prototype.writeUInt16BE=function(t,e,r){return t=+t,e>>>=0,r||L(this,t,e,2,65535,0),this[e]=t>>>8,this[e+1]=255&t,e+2},e.prototype.writeUInt32LE=function(t,e,r){return t=+t,e>>>=0,r||L(this,t,e,4,4294967295,0),this[e+3]=t>>>24,this[e+2]=t>>>16,this[e+1]=t>>>8,this[e]=255&t,e+4},e.prototype.writeUInt32BE=function(t,e,r){return t=+t,e>>>=0,r||L(this,t,e,4,4294967295,0),this[e]=t>>>24,this[e+1]=t>>>16,this[e+2]=t>>>8,this[e+3]=255&t,e+4},e.prototype.writeIntLE=function(t,e,r,n){if(t=+t,e>>>=0,!n){var i=Math.pow(2,8*r-1);L(this,t,e,r,i-1,-i)}var a=0,o=1,s=0;for(this[e]=255&t;++a<r&&(o*=256);)t<0&&0===s&&0!==this[e+a-1]&&(s=1),this[e+a]=(t/o>>0)-s&255;return e+r},e.prototype.writeIntBE=function(t,e,r,n){if(t=+t,e>>>=0,!n){var i=Math.pow(2,8*r-1);L(this,t,e,r,i-1,-i)}var a=r-1,o=1,s=0;for(this[e+a]=255&t;--a>=0&&(o*=256);)t<0&&0===s&&0!==this[e+a+1]&&(s=1),this[e+a]=(t/o>>0)-s&255;return e+r},e.prototype.writeInt8=function(t,e,r){return t=+t,e>>>=0,r||L(this,t,e,1,127,-128),t<0&&(t=255+t+1),this[e]=255&t,e+1},e.prototype.writeInt16LE=function(t,e,r){return t=+t,e>>>=0,r||L(this,t,e,2,32767,-32768),this[e]=255&t,this[e+1]=t>>>8,e+2},e.prototype.writeInt16BE=function(t,e,r){return t=+t,e>>>=0,r||L(this,t,e,2,32767,-32768),this[e]=t>>>8,this[e+1]=255&t,e+2},e.prototype.writeInt32LE=function(t,e,r){return t=+t,e>>>=0,r||L(this,t,e,4,2147483647,-2147483648),this[e]=255&t,this[e+1]=t>>>8,this[e+2]=t>>>16,this[e+3]=t>>>24,e+4},e.prototype.writeInt32BE=function(t,e,r){return t=+t,e>>>=0,r||L(this,t,e,4,2147483647,-2147483648),t<0&&(t=4294967295+t+1),this[e]=t>>>24,this[e+1]=t>>>16,this[e+2]=t>>>8,this[e+3]=255&t,e+4},e.prototype.writeFloatLE=function(t,e,r){return O(this,t,e,!0,r)},e.prototype.writeFloatBE=function(t,e,r){return O(this,t,e,!1,r)},e.prototype.writeDoubleLE=function(t,e,r){return z(this,t,e,!0,r)},e.prototype.writeDoubleBE=function(t,e,r){return z(this,t,e,!1,r)},e.prototype.copy=function(t,r,n,i){if(!e.isBuffer(t))throw new TypeError("argument should be a Buffer");if(n||(n=0),i||0===i||(i=this.length),r>=t.length&&(r=t.length),r||(r=0),i>0&&i<n&&(i=n),i===n)return 0;if(0===t.length||0===this.length)return 0;if(r<0)throw new RangeError("targetStart out of bounds");if(n<0||n>=this.length)throw new RangeError("Index out of range");if(i<0)throw new RangeError("sourceEnd out of bounds");i>this.length&&(i=this.length),t.length-r<i-n&&(i=t.length-r+n);var a=i-n;if(this===t&&"function"==typeof Uint8Array.prototype.copyWithin)this.copyWithin(r,n,i);else if(this===t&&n<r&&r<i)for(var o=a-1;o>=0;--o)t[o+r]=this[o+n];else Uint8Array.prototype.set.call(t,this.subarray(n,i),r);return a},e.prototype.fill=function(t,r,n,i){if("string"==typeof t){if("string"==typeof r?(i=r,r=0,n=this.length):"string"==typeof n&&(i=n,n=this.length),void 0!==i&&"string"!=typeof i)throw new TypeError("encoding must be a string");if("string"==typeof i&&!e.isEncoding(i))throw new TypeError("Unknown encoding: "+i);if(1===t.length){var a=t.charCodeAt(0);("utf8"===i&&a<128||"latin1"===i)&&(t=a)}}else"number"==typeof t&&(t&=255);if(r<0||this.length<r||this.length<n)throw new RangeError("Out of range index");if(n<=r)return this;var o;if(r>>>=0,n=void 0===n?this.length:n>>>0,t||(t=0),"number"==typeof t)for(o=r;o<n;++o)this[o]=t;else{var s=e.isBuffer(t)?t:e.from(t,i),l=s.length;if(0===l)throw new TypeError('The value "'+t+'" is invalid for argument "value"');for(o=0;o<n-r;++o)this[o+r]=s[o%l]}return this};var I=/[^+\/0-9A-Za-z-_]/g;function D(t){return t<16?"0"+t.toString(16):t.toString(16)}function R(t,e){var r;e=e||1/0;for(var n=t.length,i=null,a=[],o=0;o<n;++o){if((r=t.charCodeAt(o))>55295&&r<57344){if(!i){if(r>56319){(e-=3)>-1&&a.push(239,191,189);continue}if(o+1===n){(e-=3)>-1&&a.push(239,191,189);continue}i=r;continue}if(r<56320){(e-=3)>-1&&a.push(239,191,189),i=r;continue}r=65536+(i-55296<<10|r-56320)}else i&&(e-=3)>-1&&a.push(239,191,189);if(i=null,r<128){if((e-=1)<0)break;a.push(r)}else if(r<2048){if((e-=2)<0)break;a.push(r>>6|192,63&r|128)}else if(r<65536){if((e-=3)<0)break;a.push(r>>12|224,r>>6&63|128,63&r|128)}else{if(!(r<1114112))throw new Error("Invalid code point");if((e-=4)<0)break;a.push(r>>18|240,r>>12&63|128,r>>6&63|128,63&r|128)}}return a}function F(t){return n.toByteArray(function(t){if((t=(t=t.split("=")[0]).trim().replace(I,"")).length<2)return"";for(;t.length%4!=0;)t+="=";return t}(t))}function B(t,e,r,n){for(var i=0;i<n&&!(i+r>=e.length||i>=t.length);++i)e[i+r]=t[i];return i}function N(t,e){return t instanceof e||null!=t&&null!=t.constructor&&null!=t.constructor.name&&t.constructor.name===e.name}function j(t){return t!=t}}).call(this,t("buffer").Buffer)},{"base64-js":74,buffer:105,ieee754:412}],106:[function(t,e,r){"use strict";var n=t("./lib/monotone"),i=t("./lib/triangulation"),a=t("./lib/delaunay"),o=t("./lib/filter");function s(t){return[Math.min(t[0],t[1]),Math.max(t[0],t[1])]}function l(t,e){return t[0]-e[0]||t[1]-e[1]}function c(t,e,r){return e in t?t[e]:r}e.exports=function(t,e,r){Array.isArray(e)?(r=r||{},e=e||[]):(r=e||{},e=[]);var u=!!c(r,"delaunay",!0),h=!!c(r,"interior",!0),f=!!c(r,"exterior",!0),p=!!c(r,"infinity",!1);if(!h&&!f||0===t.length)return[];var d=n(t,e);if(u||h!==f||p){for(var g=i(t.length,function(t){return t.map(s).sort(l)}(e)),v=0;v<d.length;++v){var m=d[v];g.addTriangle(m[0],m[1],m[2])}return u&&a(t,g),f?h?p?o(g,0,p):g.cells():o(g,1,p):o(g,-1)}return d}},{"./lib/delaunay":107,"./lib/filter":108,"./lib/monotone":109,"./lib/triangulation":110}],107:[function(t,e,r){"use strict";var n=t("robust-in-sphere")[4];t("binary-search-bounds");function i(t,e,r,i,a,o){var s=e.opposite(i,a);if(!(s<0)){if(a<i){var l=i;i=a,a=l,l=o,o=s,s=l}e.isConstraint(i,a)||n(t[i],t[a],t[o],t[s])<0&&r.push(i,a)}}e.exports=function(t,e){for(var r=[],a=t.length,o=e.stars,s=0;s<a;++s)for(var l=o[s],c=1;c<l.length;c+=2){var u=l[c];if(!(u<s)&&!e.isConstraint(s,u)){for(var h=l[c-1],f=-1,p=1;p<l.length;p+=2)if(l[p-1]===u){f=l[p];break}f<0||n(t[s],t[u],t[h],t[f])<0&&r.push(s,u)}}for(;r.length>0;){for(var u=r.pop(),s=r.pop(),h=-1,f=-1,l=o[s],d=1;d<l.length;d+=2){var g=l[d-1],v=l[d];g===u?f=v:v===u&&(h=g)}h<0||f<0||(n(t[s],t[u],t[h],t[f])>=0||(e.flip(s,u),i(t,e,r,h,s,f),i(t,e,r,s,f,h),i(t,e,r,f,u,h),i(t,e,r,u,h,f)))}}},{"binary-search-bounds":111,"robust-in-sphere":508}],108:[function(t,e,r){"use strict";var n,i=t("binary-search-bounds");function a(t,e,r,n,i,a,o){this.cells=t,this.neighbor=e,this.flags=n,this.constraint=r,this.active=i,this.next=a,this.boundary=o}function o(t,e){return t[0]-e[0]||t[1]-e[1]||t[2]-e[2]}e.exports=function(t,e,r){var n=function(t,e){for(var r=t.cells(),n=r.length,i=0;i<n;++i){var s=r[i],l=s[0],c=s[1],u=s[2];c<u?c<l&&(s[0]=c,s[1]=u,s[2]=l):u<l&&(s[0]=u,s[1]=l,s[2]=c)}r.sort(o);for(var h=new Array(n),i=0;i<h.length;++i)h[i]=0;var f=[],p=[],d=new Array(3*n),g=new Array(3*n),v=null;e&&(v=[]);for(var m=new a(r,d,g,h,f,p,v),i=0;i<n;++i)for(var s=r[i],y=0;y<3;++y){var l=s[y],c=s[(y+1)%3],x=d[3*i+y]=m.locate(c,l,t.opposite(c,l)),b=g[3*i+y]=t.isConstraint(l,c);x<0&&(b?p.push(i):(f.push(i),h[i]=1),e&&v.push([c,l,-1]))}return m}(t,r);if(0===e)return r?n.cells.concat(n.boundary):n.cells;var i=1,s=n.active,l=n.next,c=n.flags,u=n.cells,h=n.constraint,f=n.neighbor;for(;s.length>0||l.length>0;){for(;s.length>0;){var p=s.pop();if(c[p]!==-i){c[p]=i;u[p];for(var d=0;d<3;++d){var g=f[3*p+d];g>=0&&0===c[g]&&(h[3*p+d]?l.push(g):(s.push(g),c[g]=i))}}}var v=l;l=s,s=v,l.length=0,i=-i}var m=function(t,e,r){for(var n=0,i=0;i<t.length;++i)e[i]===r&&(t[n++]=t[i]);return t.length=n,t}(u,c,e);if(r)return m.concat(n.boundary);return m},a.prototype.locate=(n=[0,0,0],function(t,e,r){var a=t,s=e,l=r;return e<r?e<t&&(a=e,s=r,l=t):r<t&&(a=r,s=t,l=e),a<0?-1:(n[0]=a,n[1]=s,n[2]=l,i.eq(this.cells,n,o))})},{"binary-search-bounds":111}],109:[function(t,e,r){"use strict";var n=t("binary-search-bounds"),i=t("robust-orientation")[3],a=0,o=1,s=2;function l(t,e,r,n,i){this.a=t,this.b=e,this.idx=r,this.lowerIds=n,this.upperIds=i}function c(t,e,r,n){this.a=t,this.b=e,this.type=r,this.idx=n}function u(t,e){var r=t.a[0]-e.a[0]||t.a[1]-e.a[1]||t.type-e.type;return r||(t.type!==a&&(r=i(t.a,t.b,e.b))?r:t.idx-e.idx)}function h(t,e){return i(t.a,t.b,e)}function f(t,e,r,a,o){for(var s=n.lt(e,a,h),l=n.gt(e,a,h),c=s;c<l;++c){for(var u=e[c],f=u.lowerIds,p=f.length;p>1&&i(r[f[p-2]],r[f[p-1]],a)>0;)t.push([f[p-1],f[p-2],o]),p-=1;f.length=p,f.push(o);var d=u.upperIds;for(p=d.length;p>1&&i(r[d[p-2]],r[d[p-1]],a)<0;)t.push([d[p-2],d[p-1],o]),p-=1;d.length=p,d.push(o)}}function p(t,e){var r;return(r=t.a[0]<e.a[0]?i(t.a,t.b,e.a):i(e.b,e.a,t.a))?r:(r=e.b[0]<t.b[0]?i(t.a,t.b,e.b):i(e.b,e.a,t.b))||t.idx-e.idx}function d(t,e,r){var i=n.le(t,r,p),a=t[i],o=a.upperIds,s=o[o.length-1];a.upperIds=[s],t.splice(i+1,0,new l(r.a,r.b,r.idx,[s],o))}function g(t,e,r){var i=r.a;r.a=r.b,r.b=i;var a=n.eq(t,r,p),o=t[a];t[a-1].upperIds=o.upperIds,t.splice(a,1)}e.exports=function(t,e){for(var r=t.length,n=e.length,i=[],h=0;h<r;++h)i.push(new c(t[h],null,a,h));for(var h=0;h<n;++h){var p=e[h],v=t[p[0]],m=t[p[1]];v[0]<m[0]?i.push(new c(v,m,s,h),new c(m,v,o,h)):v[0]>m[0]&&i.push(new c(m,v,s,h),new c(v,m,o,h))}i.sort(u);for(var y=i[0].a[0]-(1+Math.abs(i[0].a[0]))*Math.pow(2,-52),x=[new l([y,1],[y,0],-1,[],[],[],[])],b=[],h=0,_=i.length;h<_;++h){var w=i[h],k=w.type;k===a?f(b,x,t,w.a,w.idx):k===s?d(x,t,w):g(x,t,w)}return b}},{"binary-search-bounds":111,"robust-orientation":510}],110:[function(t,e,r){"use strict";var n=t("binary-search-bounds");function i(t,e){this.stars=t,this.edges=e}e.exports=function(t,e){for(var r=new Array(t),n=0;n<t;++n)r[n]=[];return new i(r,e)};var a=i.prototype;function o(t,e,r){for(var n=1,i=t.length;n<i;n+=2)if(t[n-1]===e&&t[n]===r)return t[n-1]=t[i-2],t[n]=t[i-1],void(t.length=i-2)}a.isConstraint=function(){var t=[0,0];function e(t,e){return t[0]-e[0]||t[1]-e[1]}return function(r,i){return t[0]=Math.min(r,i),t[1]=Math.max(r,i),n.eq(this.edges,t,e)>=0}}(),a.removeTriangle=function(t,e,r){var n=this.stars;o(n[t],e,r),o(n[e],r,t),o(n[r],t,e)},a.addTriangle=function(t,e,r){var n=this.stars;n[t].push(e,r),n[e].push(r,t),n[r].push(t,e)},a.opposite=function(t,e){for(var r=this.stars[e],n=1,i=r.length;n<i;n+=2)if(r[n]===t)return r[n-1];return-1},a.flip=function(t,e){var r=this.opposite(t,e),n=this.opposite(e,t);this.removeTriangle(t,e,r),this.removeTriangle(e,t,n),this.addTriangle(t,n,r),this.addTriangle(e,r,n)},a.edges=function(){for(var t=this.stars,e=[],r=0,n=t.length;r<n;++r)for(var i=t[r],a=0,o=i.length;a<o;a+=2)e.push([i[a],i[a+1]]);return e},a.cells=function(){for(var t=this.stars,e=[],r=0,n=t.length;r<n;++r)for(var i=t[r],a=0,o=i.length;a<o;a+=2){var s=i[a],l=i[a+1];r<Math.min(s,l)&&e.push([r,s,l])}return e}},{"binary-search-bounds":111}],111:[function(t,e,r){"use strict";function n(t,e,r,n,i){var a=["function ",t,"(a,l,h,",n.join(","),"){",i?"":"var i=",r?"l-1":"h+1",";while(l<=h){var m=(l+h)>>>1,x=a[m]"];return i?e.indexOf("c")<0?a.push(";if(x===y){return m}else if(x<=y){"):a.push(";var p=c(x,y);if(p===0){return m}else if(p<=0){"):a.push(";if(",e,"){i=m;"),r?a.push("l=m+1}else{h=m-1}"):a.push("h=m-1}else{l=m+1}"),a.push("}"),i?a.push("return -1};"):a.push("return i};"),a.join("")}function i(t,e,r,i){return new Function([n("A","x"+t+"y",e,["y"],i),n("P","c(x,y)"+t+"0",e,["y","c"],i),"function dispatchBsearch",r,"(a,y,c,l,h){if(typeof(c)==='function'){return P(a,(l===void 0)?0:l|0,(h===void 0)?a.length-1:h|0,y,c)}else{return A(a,(c===void 0)?0:c|0,(l===void 0)?a.length-1:l|0,y)}}return dispatchBsearch",r].join(""))()}e.exports={ge:i(">=",!1,"GE"),gt:i(">",!1,"GT"),lt:i("<",!0,"LT"),le:i("<=",!0,"LE"),eq:i("-",!0,"EQ",!0)}},{}],112:[function(t,e,r){"use strict";e.exports=function(t){for(var e=1,r=1;r<t.length;++r)for(var n=0;n<r;++n)if(t[r]<t[n])e=-e;else if(t[n]===t[r])return 0;return e}},{}],113:[function(t,e,r){"use strict";var n=t("dup"),i=t("robust-linear-solve");function a(t,e){for(var r=0,n=t.length,i=0;i<n;++i)r+=t[i]*e[i];return r}function o(t){var e=t.length;if(0===e)return[];t[0].length;var r=n([t.length+1,t.length+1],1),o=n([t.length+1],1);r[e][e]=0;for(var s=0;s<e;++s){for(var l=0;l<=s;++l)r[l][s]=r[s][l]=2*a(t[s],t[l]);o[s]=a(t[s],t[s])}var c=i(r,o),u=0,h=c[e+1];for(s=0;s<h.length;++s)u+=h[s];var f=new Array(e);for(s=0;s<e;++s){h=c[s];var p=0;for(l=0;l<h.length;++l)p+=h[l];f[s]=p/u}return f}function s(t){if(0===t.length)return[];for(var e=t[0].length,r=n([e]),i=o(t),a=0;a<t.length;++a)for(var s=0;s<e;++s)r[s]+=t[a][s]*i[a];return r}s.barycenetric=o,e.exports=s},{dup:170,"robust-linear-solve":509}],114:[function(t,e,r){e.exports=function(t){for(var e=n(t),r=0,i=0;i<t.length;++i)for(var a=t[i],o=0;o<e.length;++o)r+=Math.pow(a[o]-e[o],2);return Math.sqrt(r/t.length)};var n=t("circumcenter")},{circumcenter:113}],115:[function(t,e,r){e.exports=function(t,e,r){return e<r?t<e?e:t>r?r:t:t<r?r:t>e?e:t}},{}],116:[function(t,e,r){"use strict";e.exports=function(t,e,r){var n;if(r){n=e;for(var i=new Array(e.length),a=0;a<e.length;++a){var o=e[a];i[a]=[o[0],o[1],r[a]]}e=i}var s=function(t,e,r){var n=d(t,[],p(t));return m(e,n,r),!!n}(t,e,!!r);for(;y(t,e,!!r);)s=!0;if(r&&s){n.length=0,r.length=0;for(var a=0;a<e.length;++a){var o=e[a];n.push([o[0],o[1]]),r.push(o[2])}}return s};var n=t("union-find"),i=t("box-intersect"),a=t("robust-segment-intersect"),o=t("big-rat"),s=t("big-rat/cmp"),l=t("big-rat/to-float"),c=t("rat-vec"),u=t("nextafter"),h=t("./lib/rat-seg-intersect");function f(t){var e=l(t);return[u(e,-1/0),u(e,1/0)]}function p(t){for(var e=new Array(t.length),r=0;r<t.length;++r){var n=t[r];e[r]=[u(n[0],-1/0),u(n[1],-1/0),u(n[0],1/0),u(n[1],1/0)]}return e}function d(t,e,r){for(var a=e.length,o=new n(a),s=[],l=0;l<e.length;++l){var c=e[l],h=f(c[0]),p=f(c[1]);s.push([u(h[0],-1/0),u(p[0],-1/0),u(h[1],1/0),u(p[1],1/0)])}i(s,function(t,e){o.link(t,e)});var d=!0,g=new Array(a);for(l=0;l<a;++l){(m=o.find(l))!==l&&(d=!1,t[m]=[Math.min(t[l][0],t[m][0]),Math.min(t[l][1],t[m][1])])}if(d)return null;var v=0;for(l=0;l<a;++l){var m;(m=o.find(l))===l?(g[l]=v,t[v++]=t[l]):g[l]=-1}t.length=v;for(l=0;l<a;++l)g[l]<0&&(g[l]=g[o.find(l)]);return g}function g(t,e){return t[0]-e[0]||t[1]-e[1]}function v(t,e){var r=t[0]-e[0]||t[1]-e[1];return r||(t[2]<e[2]?-1:t[2]>e[2]?1:0)}function m(t,e,r){if(0!==t.length){if(e)for(var n=0;n<t.length;++n){var i=e[(o=t[n])[0]],a=e[o[1]];o[0]=Math.min(i,a),o[1]=Math.max(i,a)}else for(n=0;n<t.length;++n){var o;i=(o=t[n])[0],a=o[1];o[0]=Math.min(i,a),o[1]=Math.max(i,a)}r?t.sort(v):t.sort(g);var s=1;for(n=1;n<t.length;++n){var l=t[n-1],c=t[n];(c[0]!==l[0]||c[1]!==l[1]||r&&c[2]!==l[2])&&(t[s++]=c)}t.length=s}}function y(t,e,r){var n=function(t,e){for(var r=new Array(e.length),n=0;n<e.length;++n){var i=e[n],a=t[i[0]],o=t[i[1]];r[n]=[u(Math.min(a[0],o[0]),-1/0),u(Math.min(a[1],o[1]),-1/0),u(Math.max(a[0],o[0]),1/0),u(Math.max(a[1],o[1]),1/0)]}return r}(t,e),f=function(t,e,r){var n=[];return i(r,function(r,i){var o=e[r],s=e[i];if(o[0]!==s[0]&&o[0]!==s[1]&&o[1]!==s[0]&&o[1]!==s[1]){var l=t[o[0]],c=t[o[1]],u=t[s[0]],h=t[s[1]];a(l,c,u,h)&&n.push([r,i])}}),n}(t,e,n),g=p(t),v=function(t,e,r,n){var o=[];return i(r,n,function(r,n){var i=e[r];if(i[0]!==n&&i[1]!==n){var s=t[n],l=t[i[0]],c=t[i[1]];a(l,c,s,s)&&o.push([r,n])}}),o}(t,e,n,g),y=d(t,function(t,e,r,n,i){var a,u,f=t.map(function(t){return[o(t[0]),o(t[1])]});for(a=0;a<r.length;++a){var p=r[a];u=p[0];var d=p[1],g=e[u],v=e[d],m=h(c(t[g[0]]),c(t[g[1]]),c(t[v[0]]),c(t[v[1]]));if(m){var y=t.length;t.push([l(m[0]),l(m[1])]),f.push(m),n.push([u,y],[d,y])}}for(n.sort(function(t,e){if(t[0]!==e[0])return t[0]-e[0];var r=f[t[1]],n=f[e[1]];return s(r[0],n[0])||s(r[1],n[1])}),a=n.length-1;a>=0;--a){var x=e[u=(S=n[a])[0]],b=x[0],_=x[1],w=t[b],k=t[_];if((w[0]-k[0]||w[1]-k[1])<0){var T=b;b=_,_=T}x[0]=b;var A,M=x[1]=S[1];for(i&&(A=x[2]);a>0&&n[a-1][0]===u;){var S,E=(S=n[--a])[1];i?e.push([M,E,A]):e.push([M,E]),M=E}i?e.push([M,_,A]):e.push([M,_])}return f}(t,e,f,v,r));return m(e,y,r),!!y||(f.length>0||v.length>0)}},{"./lib/rat-seg-intersect":117,"big-rat":78,"big-rat/cmp":76,"big-rat/to-float":90,"box-intersect":96,nextafter:451,"rat-vec":486,"robust-segment-intersect":513,"union-find":546}],117:[function(t,e,r){"use strict";e.exports=function(t,e,r,n){var a=s(e,t),h=s(n,r),f=u(a,h);if(0===o(f))return null;var p=s(t,r),d=u(h,p),g=i(d,f),v=c(a,g);return l(t,v)};var n=t("big-rat/mul"),i=t("big-rat/div"),a=t("big-rat/sub"),o=t("big-rat/sign"),s=t("rat-vec/sub"),l=t("rat-vec/add"),c=t("rat-vec/muls");function u(t,e){return a(n(t[0],e[1]),n(t[1],e[0]))}},{"big-rat/div":77,"big-rat/mul":87,"big-rat/sign":88,"big-rat/sub":89,"rat-vec/add":485,"rat-vec/muls":487,"rat-vec/sub":488}],118:[function(t,e,r){"use strict";var n=t("clamp");function i(t,e){null==e&&(e=!0);var r=t[0],i=t[1],a=t[2],o=t[3];return null==o&&(o=e?1:255),e&&(r*=255,i*=255,a*=255,o*=255),16777216*(r=255&n(r,0,255))+((i=255&n(i,0,255))<<16)+((a=255&n(a,0,255))<<8)+(o=255&n(o,0,255))}e.exports=i,e.exports.to=i,e.exports.from=function(t,e){var r=(t=+t)>>>24,n=(16711680&t)>>>16,i=(65280&t)>>>8,a=255&t;return!1===e?[r,n,i,a]:[r/255,n/255,i/255,a/255]}},{clamp:115}],119:[function(t,e,r){"use strict";e.exports={aliceblue:[240,248,255],antiquewhite:[250,235,215],aqua:[0,255,255],aquamarine:[127,255,212],azure:[240,255,255],beige:[245,245,220],bisque:[255,228,196],black:[0,0,0],blanchedalmond:[255,235,205],blue:[0,0,255],blueviolet:[138,43,226],brown:[165,42,42],burlywood:[222,184,135],cadetblue:[95,158,160],chartreuse:[127,255,0],chocolate:[210,105,30],coral:[255,127,80],cornflowerblue:[100,149,237],cornsilk:[255,248,220],crimson:[220,20,60],cyan:[0,255,255],darkblue:[0,0,139],darkcyan:[0,139,139],darkgoldenrod:[184,134,11],darkgray:[169,169,169],darkgreen:[0,100,0],darkgrey:[169,169,169],darkkhaki:[189,183,107],darkmagenta:[139,0,139],darkolivegreen:[85,107,47],darkorange:[255,140,0],darkorchid:[153,50,204],darkred:[139,0,0],darksalmon:[233,150,122],darkseagreen:[143,188,143],darkslateblue:[72,61,139],darkslategray:[47,79,79],darkslategrey:[47,79,79],darkturquoise:[0,206,209],darkviolet:[148,0,211],deeppink:[255,20,147],deepskyblue:[0,191,255],dimgray:[105,105,105],dimgrey:[105,105,105],dodgerblue:[30,144,255],firebrick:[178,34,34],floralwhite:[255,250,240],forestgreen:[34,139,34],fuchsia:[255,0,255],gainsboro:[220,220,220],ghostwhite:[248,248,255],gold:[255,215,0],goldenrod:[218,165,32],gray:[128,128,128],green:[0,128,0],greenyellow:[173,255,47],grey:[128,128,128],honeydew:[240,255,240],hotpink:[255,105,180],indianred:[205,92,92],indigo:[75,0,130],ivory:[255,255,240],khaki:[240,230,140],lavender:[230,230,250],lavenderblush:[255,240,245],lawngreen:[124,252,0],lemonchiffon:[255,250,205],lightblue:[173,216,230],lightcoral:[240,128,128],lightcyan:[224,255,255],lightgoldenrodyellow:[250,250,210],lightgray:[211,211,211],lightgreen:[144,238,144],lightgrey:[211,211,211],lightpink:[255,182,193],lightsalmon:[255,160,122],lightseagreen:[32,178,170],lightskyblue:[135,206,250],lightslategray:[119,136,153],lightslategrey:[119,136,153],lightsteelblue:[176,196,222],lightyellow:[255,255,224],lime:[0,255,0],limegreen:[50,205,50],linen:[250,240,230],magenta:[255,0,255],maroon:[128,0,0],mediumaquamarine:[102,205,170],mediumblue:[0,0,205],mediumorchid:[186,85,211],mediumpurple:[147,112,219],mediumseagreen:[60,179,113],mediumslateblue:[123,104,238],mediumspringgreen:[0,250,154],mediumturquoise:[72,209,204],mediumvioletred:[199,21,133],midnightblue:[25,25,112],mintcream:[245,255,250],mistyrose:[255,228,225],moccasin:[255,228,181],navajowhite:[255,222,173],navy:[0,0,128],oldlace:[253,245,230],olive:[128,128,0],olivedrab:[107,142,35],orange:[255,165,0],orangered:[255,69,0],orchid:[218,112,214],palegoldenrod:[238,232,170],palegreen:[152,251,152],paleturquoise:[175,238,238],palevioletred:[219,112,147],papayawhip:[255,239,213],peachpuff:[255,218,185],peru:[205,133,63],pink:[255,192,203],plum:[221,160,221],powderblue:[176,224,230],purple:[128,0,128],rebeccapurple:[102,51,153],red:[255,0,0],rosybrown:[188,143,143],royalblue:[65,105,225],saddlebrown:[139,69,19],salmon:[250,128,114],sandybrown:[244,164,96],seagreen:[46,139,87],seashell:[255,245,238],sienna:[160,82,45],silver:[192,192,192],skyblue:[135,206,235],slateblue:[106,90,205],slategray:[112,128,144],slategrey:[112,128,144],snow:[255,250,250],springgreen:[0,255,127],steelblue:[70,130,180],tan:[210,180,140],teal:[0,128,128],thistle:[216,191,216],tomato:[255,99,71],turquoise:[64,224,208],violet:[238,130,238],wheat:[245,222,179],white:[255,255,255],whitesmoke:[245,245,245],yellow:[255,255,0],yellowgreen:[154,205,50]}},{}],120:[function(t,e,r){"use strict";var n=t("color-rgba"),i=t("clamp"),a=t("dtype");e.exports=function(t,e){"float"!==e&&e||(e="array"),"uint"===e&&(e="uint8"),"uint_clamped"===e&&(e="uint8_clamped");var r=new(a(e))(4),o="uint8"!==e&&"uint8_clamped"!==e;return t.length&&"string"!=typeof t||((t=n(t))[0]/=255,t[1]/=255,t[2]/=255),function(t){return t instanceof Uint8Array||t instanceof Uint8ClampedArray||!!(Array.isArray(t)&&(t[0]>1||0===t[0])&&(t[1]>1||0===t[1])&&(t[2]>1||0===t[2])&&(!t[3]||t[3]>1))}(t)?(r[0]=t[0],r[1]=t[1],r[2]=t[2],r[3]=null!=t[3]?t[3]:255,o&&(r[0]/=255,r[1]/=255,r[2]/=255,r[3]/=255),r):(o?(r[0]=t[0],r[1]=t[1],r[2]=t[2],r[3]=null!=t[3]?t[3]:1):(r[0]=i(Math.floor(255*t[0]),0,255),r[1]=i(Math.floor(255*t[1]),0,255),r[2]=i(Math.floor(255*t[2]),0,255),r[3]=null==t[3]?255:i(Math.floor(255*t[3]),0,255)),r)}},{clamp:115,"color-rgba":122,dtype:169}],121:[function(t,e,r){(function(r){"use strict";var n=t("color-name"),i=t("is-plain-obj"),a=t("defined");e.exports=function(t){var e,s,l=[],c=1;if("string"==typeof t)if(n[t])l=n[t].slice(),s="rgb";else if("transparent"===t)c=0,s="rgb",l=[0,0,0];else if(/^#[A-Fa-f0-9]+$/.test(t)){var u=t.slice(1),h=u.length,f=h<=4;c=1,f?(l=[parseInt(u[0]+u[0],16),parseInt(u[1]+u[1],16),parseInt(u[2]+u[2],16)],4===h&&(c=parseInt(u[3]+u[3],16)/255)):(l=[parseInt(u[0]+u[1],16),parseInt(u[2]+u[3],16),parseInt(u[4]+u[5],16)],8===h&&(c=parseInt(u[6]+u[7],16)/255)),l[0]||(l[0]=0),l[1]||(l[1]=0),l[2]||(l[2]=0),s="rgb"}else if(e=/^((?:rgb|hs[lvb]|hwb|cmyk?|xy[zy]|gray|lab|lchu?v?|[ly]uv|lms)a?)\s*\(([^\)]*)\)/.exec(t)){var p=e[1],d="rgb"===p,u=p.replace(/a$/,"");s=u;var h="cmyk"===u?4:"gray"===u?1:3;l=e[2].trim().split(/\s*,\s*/).map(function(t,e){if(/%$/.test(t))return e===h?parseFloat(t)/100:"rgb"===u?255*parseFloat(t)/100:parseFloat(t);if("h"===u[e]){if(/deg$/.test(t))return parseFloat(t);if(void 0!==o[t])return o[t]}return parseFloat(t)}),p===u&&l.push(1),c=d?1:void 0===l[h]?1:l[h],l=l.slice(0,h)}else t.length>10&&/[0-9](?:\s|\/)/.test(t)&&(l=t.match(/([0-9]+)/g).map(function(t){return parseFloat(t)}),s=t.match(/([a-z])/gi).join("").toLowerCase());else if(isNaN(t))if(i(t)){var g=a(t.r,t.red,t.R,null);null!==g?(s="rgb",l=[g,a(t.g,t.green,t.G),a(t.b,t.blue,t.B)]):(s="hsl",l=[a(t.h,t.hue,t.H),a(t.s,t.saturation,t.S),a(t.l,t.lightness,t.L,t.b,t.brightness)]),c=a(t.a,t.alpha,t.opacity,1),null!=t.opacity&&(c/=100)}else(Array.isArray(t)||r.ArrayBuffer&&ArrayBuffer.isView&&ArrayBuffer.isView(t))&&(l=[t[0],t[1],t[2]],s="rgb",c=4===t.length?t[3]:1);else s="rgb",l=[t>>>16,(65280&t)>>>8,255&t];return{space:s,values:l,alpha:c}};var o={red:0,orange:60,yellow:120,green:180,blue:240,purple:300}}).call(this,"undefined"!=typeof global?global:"undefined"!=typeof self?self:"undefined"!=typeof window?window:{})},{"color-name":119,defined:164,"is-plain-obj":422}],122:[function(t,e,r){"use strict";var n=t("color-parse"),i=t("color-space/hsl"),a=t("clamp");e.exports=function(t){var e,r=n(t);return r.space?((e=Array(3))[0]=a(r.values[0],0,255),e[1]=a(r.values[1],0,255),e[2]=a(r.values[2],0,255),"h"===r.space[0]&&(e=i.rgb(e)),e.push(a(r.alpha,0,1)),e):[]}},{clamp:115,"color-parse":121,"color-space/hsl":123}],123:[function(t,e,r){"use strict";var n=t("./rgb");e.exports={name:"hsl",min:[0,0,0],max:[360,100,100],channel:["hue","saturation","lightness"],alias:["HSL"],rgb:function(t){var e,r,n,i,a,o=t[0]/360,s=t[1]/100,l=t[2]/100;if(0===s)return[a=255*l,a,a];e=2*l-(r=l<.5?l*(1+s):l+s-l*s),i=[0,0,0];for(var c=0;c<3;c++)(n=o+1/3*-(c-1))<0?n++:n>1&&n--,a=6*n<1?e+6*(r-e)*n:2*n<1?r:3*n<2?e+(r-e)*(2/3-n)*6:e,i[c]=255*a;return i}},n.hsl=function(t){var e,r,n=t[0]/255,i=t[1]/255,a=t[2]/255,o=Math.min(n,i,a),s=Math.max(n,i,a),l=s-o;return s===o?e=0:n===s?e=(i-a)/l:i===s?e=2+(a-n)/l:a===s&&(e=4+(n-i)/l),(e=Math.min(60*e,360))<0&&(e+=360),r=(o+s)/2,[e,100*(s===o?0:r<=.5?l/(s+o):l/(2-s-o)),100*r]}},{"./rgb":124}],124:[function(t,e,r){"use strict";e.exports={name:"rgb",min:[0,0,0],max:[255,255,255],channel:["red","green","blue"],alias:["RGB"]}},{}],125:[function(t,e,r){e.exports={jet:[{index:0,rgb:[0,0,131]},{index:.125,rgb:[0,60,170]},{index:.375,rgb:[5,255,255]},{index:.625,rgb:[255,255,0]},{index:.875,rgb:[250,0,0]},{index:1,rgb:[128,0,0]}],hsv:[{index:0,rgb:[255,0,0]},{index:.169,rgb:[253,255,2]},{index:.173,rgb:[247,255,2]},{index:.337,rgb:[0,252,4]},{index:.341,rgb:[0,252,10]},{index:.506,rgb:[1,249,255]},{index:.671,rgb:[2,0,253]},{index:.675,rgb:[8,0,253]},{index:.839,rgb:[255,0,251]},{index:.843,rgb:[255,0,245]},{index:1,rgb:[255,0,6]}],hot:[{index:0,rgb:[0,0,0]},{index:.3,rgb:[230,0,0]},{index:.6,rgb:[255,210,0]},{index:1,rgb:[255,255,255]}],cool:[{index:0,rgb:[0,255,255]},{index:1,rgb:[255,0,255]}],spring:[{index:0,rgb:[255,0,255]},{index:1,rgb:[255,255,0]}],summer:[{index:0,rgb:[0,128,102]},{index:1,rgb:[255,255,102]}],autumn:[{index:0,rgb:[255,0,0]},{index:1,rgb:[255,255,0]}],winter:[{index:0,rgb:[0,0,255]},{index:1,rgb:[0,255,128]}],bone:[{index:0,rgb:[0,0,0]},{index:.376,rgb:[84,84,116]},{index:.753,rgb:[169,200,200]},{index:1,rgb:[255,255,255]}],copper:[{index:0,rgb:[0,0,0]},{index:.804,rgb:[255,160,102]},{index:1,rgb:[255,199,127]}],greys:[{index:0,rgb:[0,0,0]},{index:1,rgb:[255,255,255]}],yignbu:[{index:0,rgb:[8,29,88]},{index:.125,rgb:[37,52,148]},{index:.25,rgb:[34,94,168]},{index:.375,rgb:[29,145,192]},{index:.5,rgb:[65,182,196]},{index:.625,rgb:[127,205,187]},{index:.75,rgb:[199,233,180]},{index:.875,rgb:[237,248,217]},{index:1,rgb:[255,255,217]}],greens:[{index:0,rgb:[0,68,27]},{index:.125,rgb:[0,109,44]},{index:.25,rgb:[35,139,69]},{index:.375,rgb:[65,171,93]},{index:.5,rgb:[116,196,118]},{index:.625,rgb:[161,217,155]},{index:.75,rgb:[199,233,192]},{index:.875,rgb:[229,245,224]},{index:1,rgb:[247,252,245]}],yiorrd:[{index:0,rgb:[128,0,38]},{index:.125,rgb:[189,0,38]},{index:.25,rgb:[227,26,28]},{index:.375,rgb:[252,78,42]},{index:.5,rgb:[253,141,60]},{index:.625,rgb:[254,178,76]},{index:.75,rgb:[254,217,118]},{index:.875,rgb:[255,237,160]},{index:1,rgb:[255,255,204]}],bluered:[{index:0,rgb:[0,0,255]},{index:1,rgb:[255,0,0]}],rdbu:[{index:0,rgb:[5,10,172]},{index:.35,rgb:[106,137,247]},{index:.5,rgb:[190,190,190]},{index:.6,rgb:[220,170,132]},{index:.7,rgb:[230,145,90]},{index:1,rgb:[178,10,28]}],picnic:[{index:0,rgb:[0,0,255]},{index:.1,rgb:[51,153,255]},{index:.2,rgb:[102,204,255]},{index:.3,rgb:[153,204,255]},{index:.4,rgb:[204,204,255]},{index:.5,rgb:[255,255,255]},{index:.6,rgb:[255,204,255]},{index:.7,rgb:[255,153,255]},{index:.8,rgb:[255,102,204]},{index:.9,rgb:[255,102,102]},{index:1,rgb:[255,0,0]}],rainbow:[{index:0,rgb:[150,0,90]},{index:.125,rgb:[0,0,200]},{index:.25,rgb:[0,25,255]},{index:.375,rgb:[0,152,255]},{index:.5,rgb:[44,255,150]},{index:.625,rgb:[151,255,0]},{index:.75,rgb:[255,234,0]},{index:.875,rgb:[255,111,0]},{index:1,rgb:[255,0,0]}],portland:[{index:0,rgb:[12,51,131]},{index:.25,rgb:[10,136,186]},{index:.5,rgb:[242,211,56]},{index:.75,rgb:[242,143,56]},{index:1,rgb:[217,30,30]}],blackbody:[{index:0,rgb:[0,0,0]},{index:.2,rgb:[230,0,0]},{index:.4,rgb:[230,210,0]},{index:.7,rgb:[255,255,255]},{index:1,rgb:[160,200,255]}],earth:[{index:0,rgb:[0,0,130]},{index:.1,rgb:[0,180,180]},{index:.2,rgb:[40,210,40]},{index:.4,rgb:[230,230,50]},{index:.6,rgb:[120,70,20]},{index:1,rgb:[255,255,255]}],electric:[{index:0,rgb:[0,0,0]},{index:.15,rgb:[30,0,100]},{index:.4,rgb:[120,0,100]},{index:.6,rgb:[160,90,0]},{index:.8,rgb:[230,200,0]},{index:1,rgb:[255,250,220]}],alpha:[{index:0,rgb:[255,255,255,0]},{index:1,rgb:[255,255,255,1]}],viridis:[{index:0,rgb:[68,1,84]},{index:.13,rgb:[71,44,122]},{index:.25,rgb:[59,81,139]},{index:.38,rgb:[44,113,142]},{index:.5,rgb:[33,144,141]},{index:.63,rgb:[39,173,129]},{index:.75,rgb:[92,200,99]},{index:.88,rgb:[170,220,50]},{index:1,rgb:[253,231,37]}],inferno:[{index:0,rgb:[0,0,4]},{index:.13,rgb:[31,12,72]},{index:.25,rgb:[85,15,109]},{index:.38,rgb:[136,34,106]},{index:.5,rgb:[186,54,85]},{index:.63,rgb:[227,89,51]},{index:.75,rgb:[249,140,10]},{index:.88,rgb:[249,201,50]},{index:1,rgb:[252,255,164]}],magma:[{index:0,rgb:[0,0,4]},{index:.13,rgb:[28,16,68]},{index:.25,rgb:[79,18,123]},{index:.38,rgb:[129,37,129]},{index:.5,rgb:[181,54,122]},{index:.63,rgb:[229,80,100]},{index:.75,rgb:[251,135,97]},{index:.88,rgb:[254,194,135]},{index:1,rgb:[252,253,191]}],plasma:[{index:0,rgb:[13,8,135]},{index:.13,rgb:[75,3,161]},{index:.25,rgb:[125,3,168]},{index:.38,rgb:[168,34,150]},{index:.5,rgb:[203,70,121]},{index:.63,rgb:[229,107,93]},{index:.75,rgb:[248,148,65]},{index:.88,rgb:[253,195,40]},{index:1,rgb:[240,249,33]}],warm:[{index:0,rgb:[125,0,179]},{index:.13,rgb:[172,0,187]},{index:.25,rgb:[219,0,170]},{index:.38,rgb:[255,0,130]},{index:.5,rgb:[255,63,74]},{index:.63,rgb:[255,123,0]},{index:.75,rgb:[234,176,0]},{index:.88,rgb:[190,228,0]},{index:1,rgb:[147,255,0]}],cool:[{index:0,rgb:[125,0,179]},{index:.13,rgb:[116,0,218]},{index:.25,rgb:[98,74,237]},{index:.38,rgb:[68,146,231]},{index:.5,rgb:[0,204,197]},{index:.63,rgb:[0,247,146]},{index:.75,rgb:[0,255,88]},{index:.88,rgb:[40,255,8]},{index:1,rgb:[147,255,0]}],"rainbow-soft":[{index:0,rgb:[125,0,179]},{index:.1,rgb:[199,0,180]},{index:.2,rgb:[255,0,121]},{index:.3,rgb:[255,108,0]},{index:.4,rgb:[222,194,0]},{index:.5,rgb:[150,255,0]},{index:.6,rgb:[0,255,55]},{index:.7,rgb:[0,246,150]},{index:.8,rgb:[50,167,222]},{index:.9,rgb:[103,51,235]},{index:1,rgb:[124,0,186]}],bathymetry:[{index:0,rgb:[40,26,44]},{index:.13,rgb:[59,49,90]},{index:.25,rgb:[64,76,139]},{index:.38,rgb:[63,110,151]},{index:.5,rgb:[72,142,158]},{index:.63,rgb:[85,174,163]},{index:.75,rgb:[120,206,163]},{index:.88,rgb:[187,230,172]},{index:1,rgb:[253,254,204]}],cdom:[{index:0,rgb:[47,15,62]},{index:.13,rgb:[87,23,86]},{index:.25,rgb:[130,28,99]},{index:.38,rgb:[171,41,96]},{index:.5,rgb:[206,67,86]},{index:.63,rgb:[230,106,84]},{index:.75,rgb:[242,149,103]},{index:.88,rgb:[249,193,135]},{index:1,rgb:[254,237,176]}],chlorophyll:[{index:0,rgb:[18,36,20]},{index:.13,rgb:[25,63,41]},{index:.25,rgb:[24,91,59]},{index:.38,rgb:[13,119,72]},{index:.5,rgb:[18,148,80]},{index:.63,rgb:[80,173,89]},{index:.75,rgb:[132,196,122]},{index:.88,rgb:[175,221,162]},{index:1,rgb:[215,249,208]}],density:[{index:0,rgb:[54,14,36]},{index:.13,rgb:[89,23,80]},{index:.25,rgb:[110,45,132]},{index:.38,rgb:[120,77,178]},{index:.5,rgb:[120,113,213]},{index:.63,rgb:[115,151,228]},{index:.75,rgb:[134,185,227]},{index:.88,rgb:[177,214,227]},{index:1,rgb:[230,241,241]}],"freesurface-blue":[{index:0,rgb:[30,4,110]},{index:.13,rgb:[47,14,176]},{index:.25,rgb:[41,45,236]},{index:.38,rgb:[25,99,212]},{index:.5,rgb:[68,131,200]},{index:.63,rgb:[114,156,197]},{index:.75,rgb:[157,181,203]},{index:.88,rgb:[200,208,216]},{index:1,rgb:[241,237,236]}],"freesurface-red":[{index:0,rgb:[60,9,18]},{index:.13,rgb:[100,17,27]},{index:.25,rgb:[142,20,29]},{index:.38,rgb:[177,43,27]},{index:.5,rgb:[192,87,63]},{index:.63,rgb:[205,125,105]},{index:.75,rgb:[216,162,148]},{index:.88,rgb:[227,199,193]},{index:1,rgb:[241,237,236]}],oxygen:[{index:0,rgb:[64,5,5]},{index:.13,rgb:[106,6,15]},{index:.25,rgb:[144,26,7]},{index:.38,rgb:[168,64,3]},{index:.5,rgb:[188,100,4]},{index:.63,rgb:[206,136,11]},{index:.75,rgb:[220,174,25]},{index:.88,rgb:[231,215,44]},{index:1,rgb:[248,254,105]}],par:[{index:0,rgb:[51,20,24]},{index:.13,rgb:[90,32,35]},{index:.25,rgb:[129,44,34]},{index:.38,rgb:[159,68,25]},{index:.5,rgb:[182,99,19]},{index:.63,rgb:[199,134,22]},{index:.75,rgb:[212,171,35]},{index:.88,rgb:[221,210,54]},{index:1,rgb:[225,253,75]}],phase:[{index:0,rgb:[145,105,18]},{index:.13,rgb:[184,71,38]},{index:.25,rgb:[186,58,115]},{index:.38,rgb:[160,71,185]},{index:.5,rgb:[110,97,218]},{index:.63,rgb:[50,123,164]},{index:.75,rgb:[31,131,110]},{index:.88,rgb:[77,129,34]},{index:1,rgb:[145,105,18]}],salinity:[{index:0,rgb:[42,24,108]},{index:.13,rgb:[33,50,162]},{index:.25,rgb:[15,90,145]},{index:.38,rgb:[40,118,137]},{index:.5,rgb:[59,146,135]},{index:.63,rgb:[79,175,126]},{index:.75,rgb:[120,203,104]},{index:.88,rgb:[193,221,100]},{index:1,rgb:[253,239,154]}],temperature:[{index:0,rgb:[4,35,51]},{index:.13,rgb:[23,51,122]},{index:.25,rgb:[85,59,157]},{index:.38,rgb:[129,79,143]},{index:.5,rgb:[175,95,130]},{index:.63,rgb:[222,112,101]},{index:.75,rgb:[249,146,66]},{index:.88,rgb:[249,196,65]},{index:1,rgb:[232,250,91]}],turbidity:[{index:0,rgb:[34,31,27]},{index:.13,rgb:[65,50,41]},{index:.25,rgb:[98,69,52]},{index:.38,rgb:[131,89,57]},{index:.5,rgb:[161,112,59]},{index:.63,rgb:[185,140,66]},{index:.75,rgb:[202,174,88]},{index:.88,rgb:[216,209,126]},{index:1,rgb:[233,246,171]}],"velocity-blue":[{index:0,rgb:[17,32,64]},{index:.13,rgb:[35,52,116]},{index:.25,rgb:[29,81,156]},{index:.38,rgb:[31,113,162]},{index:.5,rgb:[50,144,169]},{index:.63,rgb:[87,173,176]},{index:.75,rgb:[149,196,189]},{index:.88,rgb:[203,221,211]},{index:1,rgb:[254,251,230]}],"velocity-green":[{index:0,rgb:[23,35,19]},{index:.13,rgb:[24,64,38]},{index:.25,rgb:[11,95,45]},{index:.38,rgb:[39,123,35]},{index:.5,rgb:[95,146,12]},{index:.63,rgb:[152,165,18]},{index:.75,rgb:[201,186,69]},{index:.88,rgb:[233,216,137]},{index:1,rgb:[255,253,205]}],cubehelix:[{index:0,rgb:[0,0,0]},{index:.07,rgb:[22,5,59]},{index:.13,rgb:[60,4,105]},{index:.2,rgb:[109,1,135]},{index:.27,rgb:[161,0,147]},{index:.33,rgb:[210,2,142]},{index:.4,rgb:[251,11,123]},{index:.47,rgb:[255,29,97]},{index:.53,rgb:[255,54,69]},{index:.6,rgb:[255,85,46]},{index:.67,rgb:[255,120,34]},{index:.73,rgb:[255,157,37]},{index:.8,rgb:[241,191,57]},{index:.87,rgb:[224,220,93]},{index:.93,rgb:[218,241,142]},{index:1,rgb:[227,253,198]}]}},{}],126:[function(t,e,r){"use strict";var n=t("./colorScale"),i=t("lerp");function a(t){return[t[0]/255,t[1]/255,t[2]/255,t[3]]}function o(t){for(var e,r="#",n=0;n<3;++n)r+=("00"+(e=(e=t[n]).toString(16))).substr(e.length);return r}function s(t){return"rgba("+t.join(",")+")"}e.exports=function(t){var e,r,l,c,u,h,f,p,d,g;t||(t={});p=(t.nshades||72)-1,f=t.format||"hex",(h=t.colormap)||(h="jet");if("string"==typeof h){if(h=h.toLowerCase(),!n[h])throw Error(h+" not a supported colorscale");u=n[h]}else{if(!Array.isArray(h))throw Error("unsupported colormap option",h);u=h.slice()}if(u.length>p+1)throw new Error(h+" map requires nshades to be at least size "+u.length);d=Array.isArray(t.alpha)?2!==t.alpha.length?[1,1]:t.alpha.slice():"number"==typeof t.alpha?[t.alpha,t.alpha]:[1,1];e=u.map(function(t){return Math.round(t.index*p)}),d[0]=Math.min(Math.max(d[0],0),1),d[1]=Math.min(Math.max(d[1],0),1);var v=u.map(function(t,e){var r=u[e].index,n=u[e].rgb.slice();return 4===n.length&&n[3]>=0&&n[3]<=1?n:(n[3]=d[0]+(d[1]-d[0])*r,n)}),m=[];for(g=0;g<e.length-1;++g){c=e[g+1]-e[g],r=v[g],l=v[g+1];for(var y=0;y<c;y++){var x=y/c;m.push([Math.round(i(r[0],l[0],x)),Math.round(i(r[1],l[1],x)),Math.round(i(r[2],l[2],x)),i(r[3],l[3],x)])}}m.push(u[u.length-1].rgb.concat(d[1])),"hex"===f?m=m.map(o):"rgbaString"===f?m=m.map(s):"float"===f&&(m=m.map(a));return m}},{"./colorScale":125,lerp:425}],127:[function(t,e,r){"use strict";e.exports=function(t,e,r,a){var o=n(e,r,a);if(0===o){var s=i(n(t,e,r)),c=i(n(t,e,a));if(s===c){if(0===s){var u=l(t,e,r),h=l(t,e,a);return u===h?0:u?1:-1}return 0}return 0===c?s>0?-1:l(t,e,a)?-1:1:0===s?c>0?1:l(t,e,r)?1:-1:i(c-s)}var f=n(t,e,r);if(f>0)return o>0&&n(t,e,a)>0?1:-1;if(f<0)return o>0||n(t,e,a)>0?1:-1;var p=n(t,e,a);return p>0?1:l(t,e,r)?1:-1};var n=t("robust-orientation"),i=t("signum"),a=t("two-sum"),o=t("robust-product"),s=t("robust-sum");function l(t,e,r){var n=a(t[0],-e[0]),i=a(t[1],-e[1]),l=a(r[0],-e[0]),c=a(r[1],-e[1]),u=s(o(n,l),o(i,c));return u[u.length-1]>=0}},{"robust-orientation":510,"robust-product":511,"robust-sum":515,signum:516,"two-sum":544}],128:[function(t,e,r){e.exports=function(t,e){var r=t.length,a=t.length-e.length;if(a)return a;switch(r){case 0:return 0;case 1:return t[0]-e[0];case 2:return t[0]+t[1]-e[0]-e[1]||n(t[0],t[1])-n(e[0],e[1]);case 3:var o=t[0]+t[1],s=e[0]+e[1];if(a=o+t[2]-(s+e[2]))return a;var l=n(t[0],t[1]),c=n(e[0],e[1]);return n(l,t[2])-n(c,e[2])||n(l+t[2],o)-n(c+e[2],s);case 4:var u=t[0],h=t[1],f=t[2],p=t[3],d=e[0],g=e[1],v=e[2],m=e[3];return u+h+f+p-(d+g+v+m)||n(u,h,f,p)-n(d,g,v,m,d)||n(u+h,u+f,u+p,h+f,h+p,f+p)-n(d+g,d+v,d+m,g+v,g+m,v+m)||n(u+h+f,u+h+p,u+f+p,h+f+p)-n(d+g+v,d+g+m,d+v+m,g+v+m);default:for(var y=t.slice().sort(i),x=e.slice().sort(i),b=0;b<r;++b)if(a=y[b]-x[b])return a;return 0}};var n=Math.min;function i(t,e){return t-e}},{}],129:[function(t,e,r){"use strict";var n=t("compare-cell"),i=t("cell-orientation");e.exports=function(t,e){return n(t,e)||i(t)-i(e)}},{"cell-orientation":112,"compare-cell":128}],130:[function(t,e,r){"use strict";var n=t("./lib/ch1d"),i=t("./lib/ch2d"),a=t("./lib/chnd");e.exports=function(t){var e=t.length;if(0===e)return[];if(1===e)return[[0]];var r=t[0].length;if(0===r)return[];if(1===r)return n(t);if(2===r)return i(t);return a(t,r)}},{"./lib/ch1d":131,"./lib/ch2d":132,"./lib/chnd":133}],131:[function(t,e,r){"use strict";e.exports=function(t){for(var e=0,r=0,n=1;n<t.length;++n)t[n][0]<t[e][0]&&(e=n),t[n][0]>t[r][0]&&(r=n);return e<r?[[e],[r]]:e>r?[[r],[e]]:[[e]]}},{}],132:[function(t,e,r){"use strict";e.exports=function(t){var e=n(t),r=e.length;if(r<=2)return[];for(var i=new Array(r),a=e[r-1],o=0;o<r;++o){var s=e[o];i[o]=[a,s],a=s}return i};var n=t("monotone-convex-hull-2d")},{"monotone-convex-hull-2d":434}],133:[function(t,e,r){"use strict";e.exports=function(t,e){try{return n(t,!0)}catch(s){var r=i(t);if(r.length<=e)return[];var a=function(t,e){for(var r=t.length,n=new Array(r),i=0;i<e.length;++i)n[i]=t[e[i]];for(var a=e.length,i=0;i<r;++i)e.indexOf(i)<0&&(n[a++]=t[i]);return n}(t,r),o=n(a,!0);return function(t,e){for(var r=t.length,n=e.length,i=0;i<r;++i)for(var a=t[i],o=0;o<a.length;++o){var s=a[o];if(s<n)a[o]=e[s];else{s-=n;for(var l=0;l<n;++l)s>=e[l]&&(s+=1);a[o]=s}}return t}(o,r)}};var n=t("incremental-convex-hull"),i=t("affine-hull")},{"affine-hull":62,"incremental-convex-hull":413}],134:[function(t,e,r){e.exports={AFG:"afghan",ALA:"\\b\\wland",ALB:"albania",DZA:"algeria",ASM:"^(?=.*americ).*samoa",AND:"andorra",AGO:"angola",AIA:"anguill?a",ATA:"antarctica",ATG:"antigua",ARG:"argentin",ARM:"armenia",ABW:"^(?!.*bonaire).*\\baruba",AUS:"australia",AUT:"^(?!.*hungary).*austria|\\baustri.*\\bemp",AZE:"azerbaijan",BHS:"bahamas",BHR:"bahrain",BGD:"bangladesh|^(?=.*east).*paki?stan",BRB:"barbados",BLR:"belarus|byelo",BEL:"^(?!.*luxem).*belgium",BLZ:"belize|^(?=.*british).*honduras",BEN:"benin|dahome",BMU:"bermuda",BTN:"bhutan",BOL:"bolivia",BES:"^(?=.*bonaire).*eustatius|^(?=.*carib).*netherlands|\\bbes.?islands",BIH:"herzegovina|bosnia",BWA:"botswana|bechuana",BVT:"bouvet",BRA:"brazil",IOT:"british.?indian.?ocean",BRN:"brunei",BGR:"bulgaria",BFA:"burkina|\\bfaso|upper.?volta",BDI:"burundi",CPV:"verde",KHM:"cambodia|kampuchea|khmer",CMR:"cameroon",CAN:"canada",CYM:"cayman",CAF:"\\bcentral.african.republic",TCD:"\\bchad",CHL:"\\bchile",CHN:"^(?!.*\\bmac)(?!.*\\bhong)(?!.*\\btai)(?!.*\\brep).*china|^(?=.*peo)(?=.*rep).*china",CXR:"christmas",CCK:"\\bcocos|keeling",COL:"colombia",COM:"comoro",COG:"^(?!.*\\bdem)(?!.*\\bd[\\.]?r)(?!.*kinshasa)(?!.*zaire)(?!.*belg)(?!.*l.opoldville)(?!.*free).*\\bcongo",COK:"\\bcook",CRI:"costa.?rica",CIV:"ivoire|ivory",HRV:"croatia",CUB:"\\bcuba",CUW:"^(?!.*bonaire).*\\bcura(c|\xe7)ao",CYP:"cyprus",CSK:"czechoslovakia",CZE:"^(?=.*rep).*czech|czechia|bohemia",COD:"\\bdem.*congo|congo.*\\bdem|congo.*\\bd[\\.]?r|\\bd[\\.]?r.*congo|belgian.?congo|congo.?free.?state|kinshasa|zaire|l.opoldville|drc|droc|rdc",DNK:"denmark",DJI:"djibouti",DMA:"dominica(?!n)",DOM:"dominican.rep",ECU:"ecuador",EGY:"egypt",SLV:"el.?salvador",GNQ:"guine.*eq|eq.*guine|^(?=.*span).*guinea",ERI:"eritrea",EST:"estonia",ETH:"ethiopia|abyssinia",FLK:"falkland|malvinas",FRO:"faroe|faeroe",FJI:"fiji",FIN:"finland",FRA:"^(?!.*\\bdep)(?!.*martinique).*france|french.?republic|\\bgaul",GUF:"^(?=.*french).*guiana",PYF:"french.?polynesia|tahiti",ATF:"french.?southern",GAB:"gabon",GMB:"gambia",GEO:"^(?!.*south).*georgia",DDR:"german.?democratic.?republic|democratic.?republic.*germany|east.germany",DEU:"^(?!.*east).*germany|^(?=.*\\bfed.*\\brep).*german",GHA:"ghana|gold.?coast",GIB:"gibraltar",GRC:"greece|hellenic|hellas",GRL:"greenland",GRD:"grenada",GLP:"guadeloupe",GUM:"\\bguam",GTM:"guatemala",GGY:"guernsey",GIN:"^(?!.*eq)(?!.*span)(?!.*bissau)(?!.*portu)(?!.*new).*guinea",GNB:"bissau|^(?=.*portu).*guinea",GUY:"guyana|british.?guiana",HTI:"haiti",HMD:"heard.*mcdonald",VAT:"holy.?see|vatican|papal.?st",HND:"^(?!.*brit).*honduras",HKG:"hong.?kong",HUN:"^(?!.*austr).*hungary",ISL:"iceland",IND:"india(?!.*ocea)",IDN:"indonesia",IRN:"\\biran|persia",IRQ:"\\biraq|mesopotamia",IRL:"(^ireland)|(^republic.*ireland)",IMN:"^(?=.*isle).*\\bman",ISR:"israel",ITA:"italy",JAM:"jamaica",JPN:"japan",JEY:"jersey",JOR:"jordan",KAZ:"kazak",KEN:"kenya|british.?east.?africa|east.?africa.?prot",KIR:"kiribati",PRK:"^(?=.*democrat|people|north|d.*p.*.r).*\\bkorea|dprk|korea.*(d.*p.*r)",KWT:"kuwait",KGZ:"kyrgyz|kirghiz",LAO:"\\blaos?\\b",LVA:"latvia",LBN:"lebanon",LSO:"lesotho|basuto",LBR:"liberia",LBY:"libya",LIE:"liechtenstein",LTU:"lithuania",LUX:"^(?!.*belg).*luxem",MAC:"maca(o|u)",MDG:"madagascar|malagasy",MWI:"malawi|nyasa",MYS:"malaysia",MDV:"maldive",MLI:"\\bmali\\b",MLT:"\\bmalta",MHL:"marshall",MTQ:"martinique",MRT:"mauritania",MUS:"mauritius",MYT:"\\bmayotte",MEX:"\\bmexic",FSM:"fed.*micronesia|micronesia.*fed",MCO:"monaco",MNG:"mongolia",MNE:"^(?!.*serbia).*montenegro",MSR:"montserrat",MAR:"morocco|\\bmaroc",MOZ:"mozambique",MMR:"myanmar|burma",NAM:"namibia",NRU:"nauru",NPL:"nepal",NLD:"^(?!.*\\bant)(?!.*\\bcarib).*netherlands",ANT:"^(?=.*\\bant).*(nether|dutch)",NCL:"new.?caledonia",NZL:"new.?zealand",NIC:"nicaragua",NER:"\\bniger(?!ia)",NGA:"nigeria",NIU:"niue",NFK:"norfolk",MNP:"mariana",NOR:"norway",OMN:"\\boman|trucial",PAK:"^(?!.*east).*paki?stan",PLW:"palau",PSE:"palestin|\\bgaza|west.?bank",PAN:"panama",PNG:"papua|new.?guinea",PRY:"paraguay",PER:"peru",PHL:"philippines",PCN:"pitcairn",POL:"poland",PRT:"portugal",PRI:"puerto.?rico",QAT:"qatar",KOR:"^(?!.*d.*p.*r)(?!.*democrat)(?!.*people)(?!.*north).*\\bkorea(?!.*d.*p.*r)",MDA:"moldov|b(a|e)ssarabia",REU:"r(e|\xe9)union",ROU:"r(o|u|ou)mania",RUS:"\\brussia|soviet.?union|u\\.?s\\.?s\\.?r|socialist.?republics",RWA:"rwanda",BLM:"barth(e|\xe9)lemy",SHN:"helena",KNA:"kitts|\\bnevis",LCA:"\\blucia",MAF:"^(?=.*collectivity).*martin|^(?=.*france).*martin(?!ique)|^(?=.*french).*martin(?!ique)",SPM:"miquelon",VCT:"vincent",WSM:"^(?!.*amer).*samoa",SMR:"san.?marino",STP:"\\bs(a|\xe3)o.?tom(e|\xe9)",SAU:"\\bsa\\w*.?arabia",SEN:"senegal",SRB:"^(?!.*monte).*serbia",SYC:"seychell",SLE:"sierra",SGP:"singapore",SXM:"^(?!.*martin)(?!.*saba).*maarten",SVK:"^(?!.*cze).*slovak",SVN:"slovenia",SLB:"solomon",SOM:"somali",ZAF:"south.africa|s\\\\..?africa",SGS:"south.?georgia|sandwich",SSD:"\\bs\\w*.?sudan",ESP:"spain",LKA:"sri.?lanka|ceylon",SDN:"^(?!.*\\bs(?!u)).*sudan",SUR:"surinam|dutch.?guiana",SJM:"svalbard",SWZ:"swaziland",SWE:"sweden",CHE:"switz|swiss",SYR:"syria",TWN:"taiwan|taipei|formosa|^(?!.*peo)(?=.*rep).*china",TJK:"tajik",THA:"thailand|\\bsiam",MKD:"macedonia|fyrom",TLS:"^(?=.*leste).*timor|^(?=.*east).*timor",TGO:"togo",TKL:"tokelau",TON:"tonga",TTO:"trinidad|tobago",TUN:"tunisia",TUR:"turkey",TKM:"turkmen",TCA:"turks",TUV:"tuvalu",UGA:"uganda",UKR:"ukrain",ARE:"emirates|^u\\.?a\\.?e\\.?$|united.?arab.?em",GBR:"united.?kingdom|britain|^u\\.?k\\.?$",TZA:"tanzania",USA:"united.?states\\b(?!.*islands)|\\bu\\.?s\\.?a\\.?\\b|^\\s*u\\.?s\\.?\\b(?!.*islands)",UMI:"minor.?outlying.?is",URY:"uruguay",UZB:"uzbek",VUT:"vanuatu|new.?hebrides",VEN:"venezuela",VNM:"^(?!.*republic).*viet.?nam|^(?=.*socialist).*viet.?nam",VGB:"^(?=.*\\bu\\.?\\s?k).*virgin|^(?=.*brit).*virgin|^(?=.*kingdom).*virgin",VIR:"^(?=.*\\bu\\.?\\s?s).*virgin|^(?=.*states).*virgin",WLF:"futuna|wallis",ESH:"western.sahara",YEM:"^(?!.*arab)(?!.*north)(?!.*sana)(?!.*peo)(?!.*dem)(?!.*south)(?!.*aden)(?!.*\\bp\\.?d\\.?r).*yemen",YMD:"^(?=.*peo).*yemen|^(?!.*rep)(?=.*dem).*yemen|^(?=.*south).*yemen|^(?=.*aden).*yemen|^(?=.*\\bp\\.?d\\.?r).*yemen",YUG:"yugoslavia",ZMB:"zambia|northern.?rhodesia",EAZ:"zanzibar",ZWE:"zimbabwe|^(?!.*northern).*rhodesia"}},{}],135:[function(t,e,r){e.exports=["xx-small","x-small","small","medium","large","x-large","xx-large","larger","smaller"]},{}],136:[function(t,e,r){e.exports=["normal","condensed","semi-condensed","extra-condensed","ultra-condensed","expanded","semi-expanded","extra-expanded","ultra-expanded"]},{}],137:[function(t,e,r){e.exports=["normal","italic","oblique"]},{}],138:[function(t,e,r){e.exports=["normal","bold","bolder","lighter","100","200","300","400","500","600","700","800","900"]},{}],139:[function(t,e,r){"use strict";e.exports={parse:t("./parse"),stringify:t("./stringify")}},{"./parse":141,"./stringify":142}],140:[function(t,e,r){"use strict";var n=t("css-font-size-keywords");e.exports={isSize:function(t){return/^[\d\.]/.test(t)||-1!==t.indexOf("/")||-1!==n.indexOf(t)}}},{"css-font-size-keywords":135}],141:[function(t,e,r){"use strict";var n=t("unquote"),i=t("css-global-keywords"),a=t("css-system-font-keywords"),o=t("css-font-weight-keywords"),s=t("css-font-style-keywords"),l=t("css-font-stretch-keywords"),c=t("string-split-by"),u=t("./lib/util").isSize;e.exports=f;var h=f.cache={};function f(t){if("string"!=typeof t)throw new Error("Font argument must be a string.");if(h[t])return h[t];if(""===t)throw new Error("Cannot parse an empty string.");if(-1!==a.indexOf(t))return h[t]={system:t};for(var e,r={style:"normal",variant:"normal",weight:"normal",stretch:"normal",lineHeight:"normal",size:"1rem",family:["serif"]},f=c(t,/\s+/);e=f.shift();){if(-1!==i.indexOf(e))return["style","variant","weight","stretch"].forEach(function(t){r[t]=e}),h[t]=r;if(-1===s.indexOf(e))if("normal"!==e&&"small-caps"!==e)if(-1===l.indexOf(e)){if(-1===o.indexOf(e)){if(u(e)){var d=c(e,"/");if(r.size=d[0],null!=d[1]?r.lineHeight=p(d[1]):"/"===f[0]&&(f.shift(),r.lineHeight=p(f.shift())),!f.length)throw new Error("Missing required font-family.");return r.family=c(f.join(" "),/\s*,\s*/).map(n),h[t]=r}throw new Error("Unknown or unsupported font token: "+e)}r.weight=e}else r.stretch=e;else r.variant=e;else r.style=e}throw new Error("Missing required font-size.")}function p(t){var e=parseFloat(t);return e.toString()===t?e:t}},{"./lib/util":140,"css-font-stretch-keywords":136,"css-font-style-keywords":137,"css-font-weight-keywords":138,"css-global-keywords":143,"css-system-font-keywords":144,"string-split-by":529,unquote:548}],142:[function(t,e,r){"use strict";var n=t("pick-by-alias"),i=t("./lib/util").isSize,a=g(t("css-global-keywords")),o=g(t("css-system-font-keywords")),s=g(t("css-font-weight-keywords")),l=g(t("css-font-style-keywords")),c=g(t("css-font-stretch-keywords")),u={normal:1,"small-caps":1},h={serif:1,"sans-serif":1,monospace:1,cursive:1,fantasy:1,"system-ui":1},f="1rem",p="serif";function d(t,e){if(t&&!e[t]&&!a[t])throw Error("Unknown keyword `"+t+"`");return t}function g(t){for(var e={},r=0;r<t.length;r++)e[t[r]]=1;return e}e.exports=function(t){if((t=n(t,{style:"style fontstyle fontStyle font-style slope distinction",variant:"variant font-variant fontVariant fontvariant var capitalization",weight:"weight w font-weight fontWeight fontweight",stretch:"stretch font-stretch fontStretch fontstretch width",size:"size s font-size fontSize fontsize height em emSize",lineHeight:"lh line-height lineHeight lineheight leading",family:"font family fontFamily font-family fontfamily type typeface face",system:"system reserved default global"})).system)return t.system&&d(t.system,o),t.system;if(d(t.style,l),d(t.variant,u),d(t.weight,s),d(t.stretch,c),null==t.size&&(t.size=f),"number"==typeof t.size&&(t.size+="px"),!i)throw Error("Bad size value `"+t.size+"`");t.family||(t.family=p),Array.isArray(t.family)&&(t.family.length||(t.family=[p]),t.family=t.family.map(function(t){return h[t]?t:'"'+t+'"'}).join(", "));var e=[];return e.push(t.style),t.variant!==t.style&&e.push(t.variant),t.weight!==t.variant&&t.weight!==t.style&&e.push(t.weight),t.stretch!==t.weight&&t.stretch!==t.variant&&t.stretch!==t.style&&e.push(t.stretch),e.push(t.size+(null==t.lineHeight||"normal"===t.lineHeight||t.lineHeight+""=="1"?"":"/"+t.lineHeight)),e.push(t.family),e.filter(Boolean).join(" ")}},{"./lib/util":140,"css-font-stretch-keywords":136,"css-font-style-keywords":137,"css-font-weight-keywords":138,"css-global-keywords":143,"css-system-font-keywords":144,"pick-by-alias":465}],143:[function(t,e,r){e.exports=["inherit","initial","unset"]},{}],144:[function(t,e,r){e.exports=["caption","icon","menu","message-box","small-caption","status-bar"]},{}],145:[function(t,e,r){"use strict";e.exports=function(t,e,r,n,i,a){var o=i-1,s=i*i,l=o*o,c=(1+2*i)*l,u=i*l,h=s*(3-2*i),f=s*o;if(t.length){a||(a=new Array(t.length));for(var p=t.length-1;p>=0;--p)a[p]=c*t[p]+u*e[p]+h*r[p]+f*n[p];return a}return c*t+u*e+h*r+f*n},e.exports.derivative=function(t,e,r,n,i,a){var o=6*i*i-6*i,s=3*i*i-4*i+1,l=-6*i*i+6*i,c=3*i*i-2*i;if(t.length){a||(a=new Array(t.length));for(var u=t.length-1;u>=0;--u)a[u]=o*t[u]+s*e[u]+l*r[u]+c*n[u];return a}return o*t+s*e+l*r[u]+c*n}},{}],146:[function(t,e,r){"use strict";var n=t("./lib/thunk.js");function i(){this.argTypes=[],this.shimArgs=[],this.arrayArgs=[],this.arrayBlockIndices=[],this.scalarArgs=[],this.offsetArgs=[],this.offsetArgIndex=[],this.indexArgs=[],this.shapeArgs=[],this.funcName="",this.pre=null,this.body=null,this.post=null,this.debug=!1}e.exports=function(t){var e=new i;e.pre=t.pre,e.body=t.body,e.post=t.post;var r=t.args.slice(0);e.argTypes=r;for(var a=0;a<r.length;++a){var o=r[a];if("array"===o||"object"==typeof o&&o.blockIndices){if(e.argTypes[a]="array",e.arrayArgs.push(a),e.arrayBlockIndices.push(o.blockIndices?o.blockIndices:0),e.shimArgs.push("array"+a),a<e.pre.args.length&&e.pre.args[a].count>0)throw new Error("cwise: pre() block may not reference array args");if(a<e.post.args.length&&e.post.args[a].count>0)throw new Error("cwise: post() block may not reference array args")}else if("scalar"===o)e.scalarArgs.push(a),e.shimArgs.push("scalar"+a);else if("index"===o){if(e.indexArgs.push(a),a<e.pre.args.length&&e.pre.args[a].count>0)throw new Error("cwise: pre() block may not reference array index");if(a<e.body.args.length&&e.body.args[a].lvalue)throw new Error("cwise: body() block may not write to array index");if(a<e.post.args.length&&e.post.args[a].count>0)throw new Error("cwise: post() block may not reference array index")}else if("shape"===o){if(e.shapeArgs.push(a),a<e.pre.args.length&&e.pre.args[a].lvalue)throw new Error("cwise: pre() block may not write to array shape");if(a<e.body.args.length&&e.body.args[a].lvalue)throw new Error("cwise: body() block may not write to array shape");if(a<e.post.args.length&&e.post.args[a].lvalue)throw new Error("cwise: post() block may not write to array shape")}else{if("object"!=typeof o||!o.offset)throw new Error("cwise: Unknown argument type "+r[a]);e.argTypes[a]="offset",e.offsetArgs.push({array:o.array,offset:o.offset}),e.offsetArgIndex.push(a)}}if(e.arrayArgs.length<=0)throw new Error("cwise: No array arguments specified");if(e.pre.args.length>r.length)throw new Error("cwise: Too many arguments in pre() block");if(e.body.args.length>r.length)throw new Error("cwise: Too many arguments in body() block");if(e.post.args.length>r.length)throw new Error("cwise: Too many arguments in post() block");return e.debug=!!t.printCode||!!t.debug,e.funcName=t.funcName||"cwise",e.blockSize=t.blockSize||64,n(e)}},{"./lib/thunk.js":148}],147:[function(t,e,r){"use strict";var n=t("uniq");function i(t,e,r){var n,i,a=t.length,o=e.arrayArgs.length,s=e.indexArgs.length>0,l=[],c=[],u=0,h=0;for(n=0;n<a;++n)c.push(["i",n,"=0"].join(""));for(i=0;i<o;++i)for(n=0;n<a;++n)h=u,u=t[n],0===n?c.push(["d",i,"s",n,"=t",i,"p",u].join("")):c.push(["d",i,"s",n,"=(t",i,"p",u,"-s",h,"*t",i,"p",h,")"].join(""));for(c.length>0&&l.push("var "+c.join(",")),n=a-1;n>=0;--n)u=t[n],l.push(["for(i",n,"=0;i",n,"<s",u,";++i",n,"){"].join(""));for(l.push(r),n=0;n<a;++n){for(h=u,u=t[n],i=0;i<o;++i)l.push(["p",i,"+=d",i,"s",n].join(""));s&&(n>0&&l.push(["index[",h,"]-=s",h].join("")),l.push(["++index[",u,"]"].join(""))),l.push("}")}return l.join("\n")}function a(t,e,r){for(var n=t.body,i=[],a=[],o=0;o<t.args.length;++o){var s=t.args[o];if(!(s.count<=0)){var l=new RegExp(s.name,"g"),c="",u=e.arrayArgs.indexOf(o);switch(e.argTypes[o]){case"offset":var h=e.offsetArgIndex.indexOf(o);u=e.offsetArgs[h].array,c="+q"+h;case"array":c="p"+u+c;var f="l"+o,p="a"+u;if(0===e.arrayBlockIndices[u])1===s.count?"generic"===r[u]?s.lvalue?(i.push(["var ",f,"=",p,".get(",c,")"].join("")),n=n.replace(l,f),a.push([p,".set(",c,",",f,")"].join(""))):n=n.replace(l,[p,".get(",c,")"].join("")):n=n.replace(l,[p,"[",c,"]"].join("")):"generic"===r[u]?(i.push(["var ",f,"=",p,".get(",c,")"].join("")),n=n.replace(l,f),s.lvalue&&a.push([p,".set(",c,",",f,")"].join(""))):(i.push(["var ",f,"=",p,"[",c,"]"].join("")),n=n.replace(l,f),s.lvalue&&a.push([p,"[",c,"]=",f].join("")));else{for(var d=[s.name],g=[c],v=0;v<Math.abs(e.arrayBlockIndices[u]);v++)d.push("\\s*\\[([^\\]]+)\\]"),g.push("$"+(v+1)+"*t"+u+"b"+v);if(l=new RegExp(d.join(""),"g"),c=g.join("+"),"generic"===r[u])throw new Error("cwise: Generic arrays not supported in combination with blocks!");n=n.replace(l,[p,"[",c,"]"].join(""))}break;case"scalar":n=n.replace(l,"Y"+e.scalarArgs.indexOf(o));break;case"index":n=n.replace(l,"index");break;case"shape":n=n.replace(l,"shape")}}}return[i.join("\n"),n,a.join("\n")].join("\n").trim()}e.exports=function(t,e){for(var r=e[1].length-Math.abs(t.arrayBlockIndices[0])|0,o=new Array(t.arrayArgs.length),s=new Array(t.arrayArgs.length),l=0;l<t.arrayArgs.length;++l)s[l]=e[2*l],o[l]=e[2*l+1];var c=[],u=[],h=[],f=[],p=[];for(l=0;l<t.arrayArgs.length;++l){t.arrayBlockIndices[l]<0?(h.push(0),f.push(r),c.push(r),u.push(r+t.arrayBlockIndices[l])):(h.push(t.arrayBlockIndices[l]),f.push(t.arrayBlockIndices[l]+r),c.push(0),u.push(t.arrayBlockIndices[l]));for(var d=[],g=0;g<o[l].length;g++)h[l]<=o[l][g]&&o[l][g]<f[l]&&d.push(o[l][g]-h[l]);p.push(d)}var v=["SS"],m=["'use strict'"],y=[];for(g=0;g<r;++g)y.push(["s",g,"=SS[",g,"]"].join(""));for(l=0;l<t.arrayArgs.length;++l){for(v.push("a"+l),v.push("t"+l),v.push("p"+l),g=0;g<r;++g)y.push(["t",l,"p",g,"=t",l,"[",h[l]+g,"]"].join(""));for(g=0;g<Math.abs(t.arrayBlockIndices[l]);++g)y.push(["t",l,"b",g,"=t",l,"[",c[l]+g,"]"].join(""))}for(l=0;l<t.scalarArgs.length;++l)v.push("Y"+l);if(t.shapeArgs.length>0&&y.push("shape=SS.slice(0)"),t.indexArgs.length>0){var x=new Array(r);for(l=0;l<r;++l)x[l]="0";y.push(["index=[",x.join(","),"]"].join(""))}for(l=0;l<t.offsetArgs.length;++l){var b=t.offsetArgs[l],_=[];for(g=0;g<b.offset.length;++g)0!==b.offset[g]&&(1===b.offset[g]?_.push(["t",b.array,"p",g].join("")):_.push([b.offset[g],"*t",b.array,"p",g].join("")));0===_.length?y.push("q"+l+"=0"):y.push(["q",l,"=",_.join("+")].join(""))}var w=n([].concat(t.pre.thisVars).concat(t.body.thisVars).concat(t.post.thisVars));for((y=y.concat(w)).length>0&&m.push("var "+y.join(",")),l=0;l<t.arrayArgs.length;++l)m.push("p"+l+"|=0");t.pre.body.length>3&&m.push(a(t.pre,t,s));var k=a(t.body,t,s),T=function(t){for(var e=0,r=t[0].length;e<r;){for(var n=1;n<t.length;++n)if(t[n][e]!==t[0][e])return e;++e}return e}(p);T<r?m.push(function(t,e,r,n){for(var a=e.length,o=r.arrayArgs.length,s=r.blockSize,l=r.indexArgs.length>0,c=[],u=0;u<o;++u)c.push(["var offset",u,"=p",u].join(""));for(u=t;u<a;++u)c.push(["for(var j"+u+"=SS[",e[u],"]|0;j",u,">0;){"].join("")),c.push(["if(j",u,"<",s,"){"].join("")),c.push(["s",e[u],"=j",u].join("")),c.push(["j",u,"=0"].join("")),c.push(["}else{s",e[u],"=",s].join("")),c.push(["j",u,"-=",s,"}"].join("")),l&&c.push(["index[",e[u],"]=j",u].join(""));for(u=0;u<o;++u){for(var h=["offset"+u],f=t;f<a;++f)h.push(["j",f,"*t",u,"p",e[f]].join(""));c.push(["p",u,"=(",h.join("+"),")"].join(""))}for(c.push(i(e,r,n)),u=t;u<a;++u)c.push("}");return c.join("\n")}(T,p[0],t,k)):m.push(i(p[0],t,k)),t.post.body.length>3&&m.push(a(t.post,t,s)),t.debug&&console.log("-----Generated cwise routine for ",e,":\n"+m.join("\n")+"\n----------");var A=[t.funcName||"unnamed","_cwise_loop_",o[0].join("s"),"m",T,function(t){for(var e=new Array(t.length),r=!0,n=0;n<t.length;++n){var i=t[n],a=i.match(/\d+/);a=a?a[0]:"",0===i.charAt(0)?e[n]="u"+i.charAt(1)+a:e[n]=i.charAt(0)+a,n>0&&(r=r&&e[n]===e[n-1])}return r?e[0]:e.join("")}(s)].join("");return new Function(["function ",A,"(",v.join(","),"){",m.join("\n"),"} return ",A].join(""))()}},{uniq:547}],148:[function(t,e,r){"use strict";var n=t("./compile.js");e.exports=function(t){var e=["'use strict'","var CACHED={}"],r=[],i=t.funcName+"_cwise_thunk";e.push(["return function ",i,"(",t.shimArgs.join(","),"){"].join(""));for(var a=[],o=[],s=[["array",t.arrayArgs[0],".shape.slice(",Math.max(0,t.arrayBlockIndices[0]),t.arrayBlockIndices[0]<0?","+t.arrayBlockIndices[0]+")":")"].join("")],l=[],c=[],u=0;u<t.arrayArgs.length;++u){var h=t.arrayArgs[u];r.push(["t",h,"=array",h,".dtype,","r",h,"=array",h,".order"].join("")),a.push("t"+h),a.push("r"+h),o.push("t"+h),o.push("r"+h+".join()"),s.push("array"+h+".data"),s.push("array"+h+".stride"),s.push("array"+h+".offset|0"),u>0&&(l.push("array"+t.arrayArgs[0]+".shape.length===array"+h+".shape.length+"+(Math.abs(t.arrayBlockIndices[0])-Math.abs(t.arrayBlockIndices[u]))),c.push("array"+t.arrayArgs[0]+".shape[shapeIndex+"+Math.max(0,t.arrayBlockIndices[0])+"]===array"+h+".shape[shapeIndex+"+Math.max(0,t.arrayBlockIndices[u])+"]"))}for(t.arrayArgs.length>1&&(e.push("if (!("+l.join(" && ")+")) throw new Error('cwise: Arrays do not all have the same dimensionality!')"),e.push("for(var shapeIndex=array"+t.arrayArgs[0]+".shape.length-"+Math.abs(t.arrayBlockIndices[0])+"; shapeIndex--\x3e0;) {"),e.push("if (!("+c.join(" && ")+")) throw new Error('cwise: Arrays do not all have the same shape!')"),e.push("}")),u=0;u<t.scalarArgs.length;++u)s.push("scalar"+t.scalarArgs[u]);return r.push(["type=[",o.join(","),"].join()"].join("")),r.push("proc=CACHED[type]"),e.push("var "+r.join(",")),e.push(["if(!proc){","CACHED[type]=proc=compile([",a.join(","),"])}","return proc(",s.join(","),")}"].join("")),t.debug&&console.log("-----Generated thunk:\n"+e.join("\n")+"\n----------"),new Function("compile",e.join("\n"))(n.bind(void 0,t))}},{"./compile.js":147}],149:[function(t,e,r){e.exports=t("cwise-compiler")},{"cwise-compiler":146}],150:[function(t,e,r){"use strict";var n,i=t("es5-ext/object/copy"),a=t("es5-ext/object/normalize-options"),o=t("es5-ext/object/valid-callable"),s=t("es5-ext/object/map"),l=t("es5-ext/object/valid-callable"),c=t("es5-ext/object/valid-value"),u=Function.prototype.bind,h=Object.defineProperty,f=Object.prototype.hasOwnProperty;n=function(t,e,r){var n,a=c(e)&&l(e.value);return delete(n=i(e)).writable,delete n.value,n.get=function(){return!r.overwriteDefinition&&f.call(this,t)?a:(e.value=u.call(a,r.resolveContext?r.resolveContext(this):this),h(this,t,e),this[t])},n},e.exports=function(t){var e=a(arguments[1]);return null!=e.resolveContext&&o(e.resolveContext),s(t,function(t,r){return n(r,t,e)})}},{"es5-ext/object/copy":189,"es5-ext/object/map":198,"es5-ext/object/normalize-options":199,"es5-ext/object/valid-callable":203,"es5-ext/object/valid-value":205}],151:[function(t,e,r){"use strict";var n=t("es5-ext/object/assign"),i=t("es5-ext/object/normalize-options"),a=t("es5-ext/object/is-callable"),o=t("es5-ext/string/#/contains");(e.exports=function(t,e){var r,a,s,l,c;return arguments.length<2||"string"!=typeof t?(l=e,e=t,t=null):l=arguments[2],null==t?(r=s=!0,a=!1):(r=o.call(t,"c"),a=o.call(t,"e"),s=o.call(t,"w")),c={value:e,configurable:r,enumerable:a,writable:s},l?n(i(l),c):c}).gs=function(t,e,r){var s,l,c,u;return"string"!=typeof t?(c=r,r=e,e=t,t=null):c=arguments[3],null==e?e=void 0:a(e)?null==r?r=void 0:a(r)||(c=r,r=void 0):(c=e,e=r=void 0),null==t?(s=!0,l=!1):(s=o.call(t,"c"),l=o.call(t,"e")),u={get:e,set:r,configurable:s,enumerable:l},c?n(i(c),u):u}},{"es5-ext/object/assign":186,"es5-ext/object/is-callable":192,"es5-ext/object/normalize-options":199,"es5-ext/string/#/contains":206}],152:[function(t,e,r){var n;n=this,function(t){"use strict";function e(t,e){return t<e?-1:t>e?1:t>=e?0:NaN}function r(t){var r;return 1===t.length&&(r=t,t=function(t,n){return e(r(t),n)}),{left:function(e,r,n,i){for(null==n&&(n=0),null==i&&(i=e.length);n<i;){var a=n+i>>>1;t(e[a],r)<0?n=a+1:i=a}return n},right:function(e,r,n,i){for(null==n&&(n=0),null==i&&(i=e.length);n<i;){var a=n+i>>>1;t(e[a],r)>0?i=a:n=a+1}return n}}}var n=r(e),i=n.right,a=n.left;function o(t,e){return[t,e]}function s(t){return null===t?NaN:+t}function l(t,e){var r,n,i=t.length,a=0,o=-1,l=0,c=0;if(null==e)for(;++o<i;)isNaN(r=s(t[o]))||(c+=(n=r-l)*(r-(l+=n/++a)));else for(;++o<i;)isNaN(r=s(e(t[o],o,t)))||(c+=(n=r-l)*(r-(l+=n/++a)));if(a>1)return c/(a-1)}function c(t,e){var r=l(t,e);return r?Math.sqrt(r):r}function u(t,e){var r,n,i,a=t.length,o=-1;if(null==e){for(;++o<a;)if(null!=(r=t[o])&&r>=r)for(n=i=r;++o<a;)null!=(r=t[o])&&(n>r&&(n=r),i<r&&(i=r))}else for(;++o<a;)if(null!=(r=e(t[o],o,t))&&r>=r)for(n=i=r;++o<a;)null!=(r=e(t[o],o,t))&&(n>r&&(n=r),i<r&&(i=r));return[n,i]}var h=Array.prototype,f=h.slice,p=h.map;function d(t){return function(){return t}}function g(t){return t}function v(t,e,r){t=+t,e=+e,r=(i=arguments.length)<2?(e=t,t=0,1):i<3?1:+r;for(var n=-1,i=0|Math.max(0,Math.ceil((e-t)/r)),a=new Array(i);++n<i;)a[n]=t+n*r;return a}var m=Math.sqrt(50),y=Math.sqrt(10),x=Math.sqrt(2);function b(t,e,r){var n=(e-t)/Math.max(0,r),i=Math.floor(Math.log(n)/Math.LN10),a=n/Math.pow(10,i);return i>=0?(a>=m?10:a>=y?5:a>=x?2:1)*Math.pow(10,i):-Math.pow(10,-i)/(a>=m?10:a>=y?5:a>=x?2:1)}function _(t,e,r){var n=Math.abs(e-t)/Math.max(0,r),i=Math.pow(10,Math.floor(Math.log(n)/Math.LN10)),a=n/i;return a>=m?i*=10:a>=y?i*=5:a>=x&&(i*=2),e<t?-i:i}function w(t){return Math.ceil(Math.log(t.length)/Math.LN2)+1}function k(t,e,r){if(null==r&&(r=s),n=t.length){if((e=+e)<=0||n<2)return+r(t[0],0,t);if(e>=1)return+r(t[n-1],n-1,t);var n,i=(n-1)*e,a=Math.floor(i),o=+r(t[a],a,t);return o+(+r(t[a+1],a+1,t)-o)*(i-a)}}function T(t,e){var r,n,i=t.length,a=-1;if(null==e){for(;++a<i;)if(null!=(r=t[a])&&r>=r)for(n=r;++a<i;)null!=(r=t[a])&&n>r&&(n=r)}else for(;++a<i;)if(null!=(r=e(t[a],a,t))&&r>=r)for(n=r;++a<i;)null!=(r=e(t[a],a,t))&&n>r&&(n=r);return n}function A(t){if(!(i=t.length))return[];for(var e=-1,r=T(t,M),n=new Array(r);++e<r;)for(var i,a=-1,o=n[e]=new Array(i);++a<i;)o[a]=t[a][e];return n}function M(t){return t.length}t.bisect=i,t.bisectRight=i,t.bisectLeft=a,t.ascending=e,t.bisector=r,t.cross=function(t,e,r){var n,i,a,s,l=t.length,c=e.length,u=new Array(l*c);for(null==r&&(r=o),n=a=0;n<l;++n)for(s=t[n],i=0;i<c;++i,++a)u[a]=r(s,e[i]);return u},t.descending=function(t,e){return e<t?-1:e>t?1:e>=t?0:NaN},t.deviation=c,t.extent=u,t.histogram=function(){var t=g,e=u,r=w;function n(n){var a,o,s=n.length,l=new Array(s);for(a=0;a<s;++a)l[a]=t(n[a],a,n);var c=e(l),u=c[0],h=c[1],f=r(l,u,h);Array.isArray(f)||(f=_(u,h,f),f=v(Math.ceil(u/f)*f,h,f));for(var p=f.length;f[0]<=u;)f.shift(),--p;for(;f[p-1]>h;)f.pop(),--p;var d,g=new Array(p+1);for(a=0;a<=p;++a)(d=g[a]=[]).x0=a>0?f[a-1]:u,d.x1=a<p?f[a]:h;for(a=0;a<s;++a)u<=(o=l[a])&&o<=h&&g[i(f,o,0,p)].push(n[a]);return g}return n.value=function(e){return arguments.length?(t="function"==typeof e?e:d(e),n):t},n.domain=function(t){return arguments.length?(e="function"==typeof t?t:d([t[0],t[1]]),n):e},n.thresholds=function(t){return arguments.length?(r="function"==typeof t?t:Array.isArray(t)?d(f.call(t)):d(t),n):r},n},t.thresholdFreedmanDiaconis=function(t,r,n){return t=p.call(t,s).sort(e),Math.ceil((n-r)/(2*(k(t,.75)-k(t,.25))*Math.pow(t.length,-1/3)))},t.thresholdScott=function(t,e,r){return Math.ceil((r-e)/(3.5*c(t)*Math.pow(t.length,-1/3)))},t.thresholdSturges=w,t.max=function(t,e){var r,n,i=t.length,a=-1;if(null==e){for(;++a<i;)if(null!=(r=t[a])&&r>=r)for(n=r;++a<i;)null!=(r=t[a])&&r>n&&(n=r)}else for(;++a<i;)if(null!=(r=e(t[a],a,t))&&r>=r)for(n=r;++a<i;)null!=(r=e(t[a],a,t))&&r>n&&(n=r);return n},t.mean=function(t,e){var r,n=t.length,i=n,a=-1,o=0;if(null==e)for(;++a<n;)isNaN(r=s(t[a]))?--i:o+=r;else for(;++a<n;)isNaN(r=s(e(t[a],a,t)))?--i:o+=r;if(i)return o/i},t.median=function(t,r){var n,i=t.length,a=-1,o=[];if(null==r)for(;++a<i;)isNaN(n=s(t[a]))||o.push(n);else for(;++a<i;)isNaN(n=s(r(t[a],a,t)))||o.push(n);return k(o.sort(e),.5)},t.merge=function(t){for(var e,r,n,i=t.length,a=-1,o=0;++a<i;)o+=t[a].length;for(r=new Array(o);--i>=0;)for(e=(n=t[i]).length;--e>=0;)r[--o]=n[e];return r},t.min=T,t.pairs=function(t,e){null==e&&(e=o);for(var r=0,n=t.length-1,i=t[0],a=new Array(n<0?0:n);r<n;)a[r]=e(i,i=t[++r]);return a},t.permute=function(t,e){for(var r=e.length,n=new Array(r);r--;)n[r]=t[e[r]];return n},t.quantile=k,t.range=v,t.scan=function(t,r){if(n=t.length){var n,i,a=0,o=0,s=t[o];for(null==r&&(r=e);++a<n;)(r(i=t[a],s)<0||0!==r(s,s))&&(s=i,o=a);return 0===r(s,s)?o:void 0}},t.shuffle=function(t,e,r){for(var n,i,a=(null==r?t.length:r)-(e=null==e?0:+e);a;)i=Math.random()*a--|0,n=t[a+e],t[a+e]=t[i+e],t[i+e]=n;return t},t.sum=function(t,e){var r,n=t.length,i=-1,a=0;if(null==e)for(;++i<n;)(r=+t[i])&&(a+=r);else for(;++i<n;)(r=+e(t[i],i,t))&&(a+=r);return a},t.ticks=function(t,e,r){var n,i,a,o,s=-1;if(r=+r,(t=+t)==(e=+e)&&r>0)return[t];if((n=e<t)&&(i=t,t=e,e=i),0===(o=b(t,e,r))||!isFinite(o))return[];if(o>0)for(t=Math.ceil(t/o),e=Math.floor(e/o),a=new Array(i=Math.ceil(e-t+1));++s<i;)a[s]=(t+s)*o;else for(t=Math.floor(t*o),e=Math.ceil(e*o),a=new Array(i=Math.ceil(t-e+1));++s<i;)a[s]=(t-s)/o;return n&&a.reverse(),a},t.tickIncrement=b,t.tickStep=_,t.transpose=A,t.variance=l,t.zip=function(){return A(arguments)},Object.defineProperty(t,"__esModule",{value:!0})}("object"==typeof r&&"undefined"!=typeof e?r:n.d3=n.d3||{})},{}],153:[function(t,e,r){var n;n=this,function(t){"use strict";function e(){}function r(t,r){var n=new e;if(t instanceof e)t.each(function(t,e){n.set(e,t)});else if(Array.isArray(t)){var i,a=-1,o=t.length;if(null==r)for(;++a<o;)n.set(a,t[a]);else for(;++a<o;)n.set(r(i=t[a],a,t),i)}else if(t)for(var s in t)n.set(s,t[s]);return n}e.prototype=r.prototype={constructor:e,has:function(t){return"$"+t in this},get:function(t){return this["$"+t]},set:function(t,e){return this["$"+t]=e,this},remove:function(t){var e="$"+t;return e in this&&delete this[e]},clear:function(){for(var t in this)"$"===t[0]&&delete this[t]},keys:function(){var t=[];for(var e in this)"$"===e[0]&&t.push(e.slice(1));return t},values:function(){var t=[];for(var e in this)"$"===e[0]&&t.push(this[e]);return t},entries:function(){var t=[];for(var e in this)"$"===e[0]&&t.push({key:e.slice(1),value:this[e]});return t},size:function(){var t=0;for(var e in this)"$"===e[0]&&++t;return t},empty:function(){for(var t in this)if("$"===t[0])return!1;return!0},each:function(t){for(var e in this)"$"===e[0]&&t(this[e],e.slice(1),this)}};function n(){return{}}function i(t,e,r){t[e]=r}function a(){return r()}function o(t,e,r){t.set(e,r)}function s(){}var l=r.prototype;function c(t,e){var r=new s;if(t instanceof s)t.each(function(t){r.add(t)});else if(t){var n=-1,i=t.length;if(null==e)for(;++n<i;)r.add(t[n]);else for(;++n<i;)r.add(e(t[n],n,t))}return r}s.prototype=c.prototype={constructor:s,has:l.has,add:function(t){return this["$"+(t+="")]=t,this},remove:l.remove,clear:l.clear,values:l.keys,size:l.size,empty:l.empty,each:l.each};t.nest=function(){var t,e,s,l=[],c=[];function u(n,i,a,o){if(i>=l.length)return null!=t&&n.sort(t),null!=e?e(n):n;for(var s,c,h,f=-1,p=n.length,d=l[i++],g=r(),v=a();++f<p;)(h=g.get(s=d(c=n[f])+""))?h.push(c):g.set(s,[c]);return g.each(function(t,e){o(v,e,u(t,i,a,o))}),v}return s={object:function(t){return u(t,0,n,i)},map:function(t){return u(t,0,a,o)},entries:function(t){return function t(r,n){if(++n>l.length)return r;var i,a=c[n-1];return null!=e&&n>=l.length?i=r.entries():(i=[],r.each(function(e,r){i.push({key:r,values:t(e,n)})})),null!=a?i.sort(function(t,e){return a(t.key,e.key)}):i}(u(t,0,a,o),0)},key:function(t){return l.push(t),s},sortKeys:function(t){return c[l.length-1]=t,s},sortValues:function(e){return t=e,s},rollup:function(t){return e=t,s}}},t.set=c,t.map=r,t.keys=function(t){var e=[];for(var r in t)e.push(r);return e},t.values=function(t){var e=[];for(var r in t)e.push(t[r]);return e},t.entries=function(t){var e=[];for(var r in t)e.push({key:r,value:t[r]});return e},Object.defineProperty(t,"__esModule",{value:!0})}("object"==typeof r&&"undefined"!=typeof e?r:n.d3=n.d3||{})},{}],154:[function(t,e,r){var n;n=this,function(t){"use strict";function e(t,e,r){t.prototype=e.prototype=r,r.constructor=t}function r(t,e){var r=Object.create(t.prototype);for(var n in e)r[n]=e[n];return r}function n(){}var i="\\s*([+-]?\\d+)\\s*",a="\\s*([+-]?\\d*\\.?\\d+(?:[eE][+-]?\\d+)?)\\s*",o="\\s*([+-]?\\d*\\.?\\d+(?:[eE][+-]?\\d+)?)%\\s*",s=/^#([0-9a-f]{3})$/,l=/^#([0-9a-f]{6})$/,c=new RegExp("^rgb\\("+[i,i,i]+"\\)$"),u=new RegExp("^rgb\\("+[o,o,o]+"\\)$"),h=new RegExp("^rgba\\("+[i,i,i,a]+"\\)$"),f=new RegExp("^rgba\\("+[o,o,o,a]+"\\)$"),p=new RegExp("^hsl\\("+[a,o,o]+"\\)$"),d=new RegExp("^hsla\\("+[a,o,o,a]+"\\)$"),g={aliceblue:15792383,antiquewhite:16444375,aqua:65535,aquamarine:8388564,azure:15794175,beige:16119260,bisque:16770244,black:0,blanchedalmond:16772045,blue:255,blueviolet:9055202,brown:10824234,burlywood:14596231,cadetblue:6266528,chartreuse:8388352,chocolate:13789470,coral:16744272,cornflowerblue:6591981,cornsilk:16775388,crimson:14423100,cyan:65535,darkblue:139,darkcyan:35723,darkgoldenrod:12092939,darkgray:11119017,darkgreen:25600,darkgrey:11119017,darkkhaki:12433259,darkmagenta:9109643,darkolivegreen:5597999,darkorange:16747520,darkorchid:10040012,darkred:9109504,darksalmon:15308410,darkseagreen:9419919,darkslateblue:4734347,darkslategray:3100495,darkslategrey:3100495,darkturquoise:52945,darkviolet:9699539,deeppink:16716947,deepskyblue:49151,dimgray:6908265,dimgrey:6908265,dodgerblue:2003199,firebrick:11674146,floralwhite:16775920,forestgreen:2263842,fuchsia:16711935,gainsboro:14474460,ghostwhite:16316671,gold:16766720,goldenrod:14329120,gray:8421504,green:32768,greenyellow:11403055,grey:8421504,honeydew:15794160,hotpink:16738740,indianred:13458524,indigo:4915330,ivory:16777200,khaki:15787660,lavender:15132410,lavenderblush:16773365,lawngreen:8190976,lemonchiffon:16775885,lightblue:11393254,lightcoral:15761536,lightcyan:14745599,lightgoldenrodyellow:16448210,lightgray:13882323,lightgreen:9498256,lightgrey:13882323,lightpink:16758465,lightsalmon:16752762,lightseagreen:2142890,lightskyblue:8900346,lightslategray:7833753,lightslategrey:7833753,lightsteelblue:11584734,lightyellow:16777184,lime:65280,limegreen:3329330,linen:16445670,magenta:16711935,maroon:8388608,mediumaquamarine:6737322,mediumblue:205,mediumorchid:12211667,mediumpurple:9662683,mediumseagreen:3978097,mediumslateblue:8087790,mediumspringgreen:64154,mediumturquoise:4772300,mediumvioletred:13047173,midnightblue:1644912,mintcream:16121850,mistyrose:16770273,moccasin:16770229,navajowhite:16768685,navy:128,oldlace:16643558,olive:8421376,olivedrab:7048739,orange:16753920,orangered:16729344,orchid:14315734,palegoldenrod:15657130,palegreen:10025880,paleturquoise:11529966,palevioletred:14381203,papayawhip:16773077,peachpuff:16767673,peru:13468991,pink:16761035,plum:14524637,powderblue:11591910,purple:8388736,rebeccapurple:6697881,red:16711680,rosybrown:12357519,royalblue:4286945,saddlebrown:9127187,salmon:16416882,sandybrown:16032864,seagreen:3050327,seashell:16774638,sienna:10506797,silver:12632256,skyblue:8900331,slateblue:6970061,slategray:7372944,slategrey:7372944,snow:16775930,springgreen:65407,steelblue:4620980,tan:13808780,teal:32896,thistle:14204888,tomato:16737095,turquoise:4251856,violet:15631086,wheat:16113331,white:16777215,whitesmoke:16119285,yellow:16776960,yellowgreen:10145074};function v(t){var e;return t=(t+"").trim().toLowerCase(),(e=s.exec(t))?new _((e=parseInt(e[1],16))>>8&15|e>>4&240,e>>4&15|240&e,(15&e)<<4|15&e,1):(e=l.exec(t))?m(parseInt(e[1],16)):(e=c.exec(t))?new _(e[1],e[2],e[3],1):(e=u.exec(t))?new _(255*e[1]/100,255*e[2]/100,255*e[3]/100,1):(e=h.exec(t))?y(e[1],e[2],e[3],e[4]):(e=f.exec(t))?y(255*e[1]/100,255*e[2]/100,255*e[3]/100,e[4]):(e=p.exec(t))?k(e[1],e[2]/100,e[3]/100,1):(e=d.exec(t))?k(e[1],e[2]/100,e[3]/100,e[4]):g.hasOwnProperty(t)?m(g[t]):"transparent"===t?new _(NaN,NaN,NaN,0):null}function m(t){return new _(t>>16&255,t>>8&255,255&t,1)}function y(t,e,r,n){return n<=0&&(t=e=r=NaN),new _(t,e,r,n)}function x(t){return t instanceof n||(t=v(t)),t?new _((t=t.rgb()).r,t.g,t.b,t.opacity):new _}function b(t,e,r,n){return 1===arguments.length?x(t):new _(t,e,r,null==n?1:n)}function _(t,e,r,n){this.r=+t,this.g=+e,this.b=+r,this.opacity=+n}function w(t){return((t=Math.max(0,Math.min(255,Math.round(t)||0)))<16?"0":"")+t.toString(16)}function k(t,e,r,n){return n<=0?t=e=r=NaN:r<=0||r>=1?t=e=NaN:e<=0&&(t=NaN),new A(t,e,r,n)}function T(t,e,r,i){return 1===arguments.length?function(t){if(t instanceof A)return new A(t.h,t.s,t.l,t.opacity);if(t instanceof n||(t=v(t)),!t)return new A;if(t instanceof A)return t;var e=(t=t.rgb()).r/255,r=t.g/255,i=t.b/255,a=Math.min(e,r,i),o=Math.max(e,r,i),s=NaN,l=o-a,c=(o+a)/2;return l?(s=e===o?(r-i)/l+6*(r<i):r===o?(i-e)/l+2:(e-r)/l+4,l/=c<.5?o+a:2-o-a,s*=60):l=c>0&&c<1?0:s,new A(s,l,c,t.opacity)}(t):new A(t,e,r,null==i?1:i)}function A(t,e,r,n){this.h=+t,this.s=+e,this.l=+r,this.opacity=+n}function M(t,e,r){return 255*(t<60?e+(r-e)*t/60:t<180?r:t<240?e+(r-e)*(240-t)/60:e)}e(n,v,{displayable:function(){return this.rgb().displayable()},hex:function(){return this.rgb().hex()},toString:function(){return this.rgb()+""}}),e(_,b,r(n,{brighter:function(t){return t=null==t?1/.7:Math.pow(1/.7,t),new _(this.r*t,this.g*t,this.b*t,this.opacity)},darker:function(t){return t=null==t?.7:Math.pow(.7,t),new _(this.r*t,this.g*t,this.b*t,this.opacity)},rgb:function(){return this},displayable:function(){return 0<=this.r&&this.r<=255&&0<=this.g&&this.g<=255&&0<=this.b&&this.b<=255&&0<=this.opacity&&this.opacity<=1},hex:function(){return"#"+w(this.r)+w(this.g)+w(this.b)},toString:function(){var t=this.opacity;return(1===(t=isNaN(t)?1:Math.max(0,Math.min(1,t)))?"rgb(":"rgba(")+Math.max(0,Math.min(255,Math.round(this.r)||0))+", "+Math.max(0,Math.min(255,Math.round(this.g)||0))+", "+Math.max(0,Math.min(255,Math.round(this.b)||0))+(1===t?")":", "+t+")")}})),e(A,T,r(n,{brighter:function(t){return t=null==t?1/.7:Math.pow(1/.7,t),new A(this.h,this.s,this.l*t,this.opacity)},darker:function(t){return t=null==t?.7:Math.pow(.7,t),new A(this.h,this.s,this.l*t,this.opacity)},rgb:function(){var t=this.h%360+360*(this.h<0),e=isNaN(t)||isNaN(this.s)?0:this.s,r=this.l,n=r+(r<.5?r:1-r)*e,i=2*r-n;return new _(M(t>=240?t-240:t+120,i,n),M(t,i,n),M(t<120?t+240:t-120,i,n),this.opacity)},displayable:function(){return(0<=this.s&&this.s<=1||isNaN(this.s))&&0<=this.l&&this.l<=1&&0<=this.opacity&&this.opacity<=1}}));var S=Math.PI/180,E=180/Math.PI,C=.96422,L=1,P=.82521,O=4/29,z=6/29,I=3*z*z,D=z*z*z;function R(t){if(t instanceof B)return new B(t.l,t.a,t.b,t.opacity);if(t instanceof G){if(isNaN(t.h))return new B(t.l,0,0,t.opacity);var e=t.h*S;return new B(t.l,Math.cos(e)*t.c,Math.sin(e)*t.c,t.opacity)}t instanceof _||(t=x(t));var r,n,i=V(t.r),a=V(t.g),o=V(t.b),s=N((.2225045*i+.7168786*a+.0606169*o)/L);return i===a&&a===o?r=n=s:(r=N((.4360747*i+.3850649*a+.1430804*o)/C),n=N((.0139322*i+.0971045*a+.7141733*o)/P)),new B(116*s-16,500*(r-s),200*(s-n),t.opacity)}function F(t,e,r,n){return 1===arguments.length?R(t):new B(t,e,r,null==n?1:n)}function B(t,e,r,n){this.l=+t,this.a=+e,this.b=+r,this.opacity=+n}function N(t){return t>D?Math.pow(t,1/3):t/I+O}function j(t){return t>z?t*t*t:I*(t-O)}function U(t){return 255*(t<=.0031308?12.92*t:1.055*Math.pow(t,1/2.4)-.055)}function V(t){return(t/=255)<=.04045?t/12.92:Math.pow((t+.055)/1.055,2.4)}function q(t){if(t instanceof G)return new G(t.h,t.c,t.l,t.opacity);if(t instanceof B||(t=R(t)),0===t.a&&0===t.b)return new G(NaN,0,t.l,t.opacity);var e=Math.atan2(t.b,t.a)*E;return new G(e<0?e+360:e,Math.sqrt(t.a*t.a+t.b*t.b),t.l,t.opacity)}function H(t,e,r,n){return 1===arguments.length?q(t):new G(t,e,r,null==n?1:n)}function G(t,e,r,n){this.h=+t,this.c=+e,this.l=+r,this.opacity=+n}e(B,F,r(n,{brighter:function(t){return new B(this.l+18*(null==t?1:t),this.a,this.b,this.opacity)},darker:function(t){return new B(this.l-18*(null==t?1:t),this.a,this.b,this.opacity)},rgb:function(){var t=(this.l+16)/116,e=isNaN(this.a)?t:t+this.a/500,r=isNaN(this.b)?t:t-this.b/200;return new _(U(3.1338561*(e=C*j(e))-1.6168667*(t=L*j(t))-.4906146*(r=P*j(r))),U(-.9787684*e+1.9161415*t+.033454*r),U(.0719453*e-.2289914*t+1.4052427*r),this.opacity)}})),e(G,H,r(n,{brighter:function(t){return new G(this.h,this.c,this.l+18*(null==t?1:t),this.opacity)},darker:function(t){return new G(this.h,this.c,this.l-18*(null==t?1:t),this.opacity)},rgb:function(){return R(this).rgb()}}));var Y=-.14861,W=1.78277,X=-.29227,Z=-.90649,J=1.97294,K=J*Z,$=J*W,Q=W*X-Z*Y;function tt(t,e,r,n){return 1===arguments.length?function(t){if(t instanceof et)return new et(t.h,t.s,t.l,t.opacity);t instanceof _||(t=x(t));var e=t.r/255,r=t.g/255,n=t.b/255,i=(Q*n+K*e-$*r)/(Q+K-$),a=n-i,o=(J*(r-i)-X*a)/Z,s=Math.sqrt(o*o+a*a)/(J*i*(1-i)),l=s?Math.atan2(o,a)*E-120:NaN;return new et(l<0?l+360:l,s,i,t.opacity)}(t):new et(t,e,r,null==n?1:n)}function et(t,e,r,n){this.h=+t,this.s=+e,this.l=+r,this.opacity=+n}e(et,tt,r(n,{brighter:function(t){return t=null==t?1/.7:Math.pow(1/.7,t),new et(this.h,this.s,this.l*t,this.opacity)},darker:function(t){return t=null==t?.7:Math.pow(.7,t),new et(this.h,this.s,this.l*t,this.opacity)},rgb:function(){var t=isNaN(this.h)?0:(this.h+120)*S,e=+this.l,r=isNaN(this.s)?0:this.s*e*(1-e),n=Math.cos(t),i=Math.sin(t);return new _(255*(e+r*(Y*n+W*i)),255*(e+r*(X*n+Z*i)),255*(e+r*(J*n)),this.opacity)}})),t.color=v,t.rgb=b,t.hsl=T,t.lab=F,t.hcl=H,t.lch=function(t,e,r,n){return 1===arguments.length?q(t):new G(r,e,t,null==n?1:n)},t.gray=function(t,e){return new B(t,0,0,null==e?1:e)},t.cubehelix=tt,Object.defineProperty(t,"__esModule",{value:!0})}("object"==typeof r&&"undefined"!=typeof e?r:n.d3=n.d3||{})},{}],155:[function(t,e,r){var n;n=this,function(t){"use strict";var e={value:function(){}};function r(){for(var t,e=0,r=arguments.length,i={};e<r;++e){if(!(t=arguments[e]+"")||t in i)throw new Error("illegal type: "+t);i[t]=[]}return new n(i)}function n(t){this._=t}function i(t,e){for(var r,n=0,i=t.length;n<i;++n)if((r=t[n]).name===e)return r.value}function a(t,r,n){for(var i=0,a=t.length;i<a;++i)if(t[i].name===r){t[i]=e,t=t.slice(0,i).concat(t.slice(i+1));break}return null!=n&&t.push({name:r,value:n}),t}n.prototype=r.prototype={constructor:n,on:function(t,e){var r,n,o=this._,s=(n=o,(t+"").trim().split(/^|\s+/).map(function(t){var e="",r=t.indexOf(".");if(r>=0&&(e=t.slice(r+1),t=t.slice(0,r)),t&&!n.hasOwnProperty(t))throw new Error("unknown type: "+t);return{type:t,name:e}})),l=-1,c=s.length;if(!(arguments.length<2)){if(null!=e&&"function"!=typeof e)throw new Error("invalid callback: "+e);for(;++l<c;)if(r=(t=s[l]).type)o[r]=a(o[r],t.name,e);else if(null==e)for(r in o)o[r]=a(o[r],t.name,null);return this}for(;++l<c;)if((r=(t=s[l]).type)&&(r=i(o[r],t.name)))return r},copy:function(){var t={},e=this._;for(var r in e)t[r]=e[r].slice();return new n(t)},call:function(t,e){if((r=arguments.length-2)>0)for(var r,n,i=new Array(r),a=0;a<r;++a)i[a]=arguments[a+2];if(!this._.hasOwnProperty(t))throw new Error("unknown type: "+t);for(a=0,r=(n=this._[t]).length;a<r;++a)n[a].value.apply(e,i)},apply:function(t,e,r){if(!this._.hasOwnProperty(t))throw new Error("unknown type: "+t);for(var n=this._[t],i=0,a=n.length;i<a;++i)n[i].value.apply(e,r)}},t.dispatch=r,Object.defineProperty(t,"__esModule",{value:!0})}("object"==typeof r&&"undefined"!=typeof e?r:n.d3=n.d3||{})},{}],156:[function(t,e,r){var n,i;n=this,i=function(t,e,r,n,i){"use strict";var a=function(t){return function(){return t}},o=function(){return 1e-6*(Math.random()-.5)};function s(t){return t.x+t.vx}function l(t){return t.y+t.vy}function c(t){return t.index}function u(t,e){var r=t.get(e);if(!r)throw new Error("missing: "+e);return r}function h(t){return t.x}function f(t){return t.y}var p=10,d=Math.PI*(3-Math.sqrt(5));t.forceCenter=function(t,e){var r;function n(){var n,i,a=r.length,o=0,s=0;for(n=0;n<a;++n)o+=(i=r[n]).x,s+=i.y;for(o=o/a-t,s=s/a-e,n=0;n<a;++n)(i=r[n]).x-=o,i.y-=s}return null==t&&(t=0),null==e&&(e=0),n.initialize=function(t){r=t},n.x=function(e){return arguments.length?(t=+e,n):t},n.y=function(t){return arguments.length?(e=+t,n):e},n},t.forceCollide=function(t){var r,n,i=1,c=1;function u(){for(var t,a,u,f,p,d,g,v=r.length,m=0;m<c;++m)for(a=e.quadtree(r,s,l).visitAfter(h),t=0;t<v;++t)u=r[t],d=n[u.index],g=d*d,f=u.x+u.vx,p=u.y+u.vy,a.visit(y);function y(t,e,r,n,a){var s=t.data,l=t.r,c=d+l;if(!s)return e>f+c||n<f-c||r>p+c||a<p-c;if(s.index>u.index){var h=f-s.x-s.vx,v=p-s.y-s.vy,m=h*h+v*v;m<c*c&&(0===h&&(m+=(h=o())*h),0===v&&(m+=(v=o())*v),m=(c-(m=Math.sqrt(m)))/m*i,u.vx+=(h*=m)*(c=(l*=l)/(g+l)),u.vy+=(v*=m)*c,s.vx-=h*(c=1-c),s.vy-=v*c)}}}function h(t){if(t.data)return t.r=n[t.data.index];for(var e=t.r=0;e<4;++e)t[e]&&t[e].r>t.r&&(t.r=t[e].r)}function f(){if(r){var e,i,a=r.length;for(n=new Array(a),e=0;e<a;++e)i=r[e],n[i.index]=+t(i,e,r)}}return"function"!=typeof t&&(t=a(null==t?1:+t)),u.initialize=function(t){r=t,f()},u.iterations=function(t){return arguments.length?(c=+t,u):c},u.strength=function(t){return arguments.length?(i=+t,u):i},u.radius=function(e){return arguments.length?(t="function"==typeof e?e:a(+e),f(),u):t},u},t.forceLink=function(t){var e,n,i,s,l,h=c,f=function(t){return 1/Math.min(s[t.source.index],s[t.target.index])},p=a(30),d=1;function g(r){for(var i=0,a=t.length;i<d;++i)for(var s,c,u,h,f,p,g,v=0;v<a;++v)c=(s=t[v]).source,h=(u=s.target).x+u.vx-c.x-c.vx||o(),f=u.y+u.vy-c.y-c.vy||o(),h*=p=((p=Math.sqrt(h*h+f*f))-n[v])/p*r*e[v],f*=p,u.vx-=h*(g=l[v]),u.vy-=f*g,c.vx+=h*(g=1-g),c.vy+=f*g}function v(){if(i){var a,o,c=i.length,f=t.length,p=r.map(i,h);for(a=0,s=new Array(c);a<f;++a)(o=t[a]).index=a,"object"!=typeof o.source&&(o.source=u(p,o.source)),"object"!=typeof o.target&&(o.target=u(p,o.target)),s[o.source.index]=(s[o.source.index]||0)+1,s[o.target.index]=(s[o.target.index]||0)+1;for(a=0,l=new Array(f);a<f;++a)o=t[a],l[a]=s[o.source.index]/(s[o.source.index]+s[o.target.index]);e=new Array(f),m(),n=new Array(f),y()}}function m(){if(i)for(var r=0,n=t.length;r<n;++r)e[r]=+f(t[r],r,t)}function y(){if(i)for(var e=0,r=t.length;e<r;++e)n[e]=+p(t[e],e,t)}return null==t&&(t=[]),g.initialize=function(t){i=t,v()},g.links=function(e){return arguments.length?(t=e,v(),g):t},g.id=function(t){return arguments.length?(h=t,g):h},g.iterations=function(t){return arguments.length?(d=+t,g):d},g.strength=function(t){return arguments.length?(f="function"==typeof t?t:a(+t),m(),g):f},g.distance=function(t){return arguments.length?(p="function"==typeof t?t:a(+t),y(),g):p},g},t.forceManyBody=function(){var t,r,n,i,s=a(-30),l=1,c=1/0,u=.81;function p(i){var a,o=t.length,s=e.quadtree(t,h,f).visitAfter(g);for(n=i,a=0;a<o;++a)r=t[a],s.visit(v)}function d(){if(t){var e,r,n=t.length;for(i=new Array(n),e=0;e<n;++e)r=t[e],i[r.index]=+s(r,e,t)}}function g(t){var e,r,n,a,o,s=0,l=0;if(t.length){for(n=a=o=0;o<4;++o)(e=t[o])&&(r=Math.abs(e.value))&&(s+=e.value,l+=r,n+=r*e.x,a+=r*e.y);t.x=n/l,t.y=a/l}else{(e=t).x=e.data.x,e.y=e.data.y;do{s+=i[e.data.index]}while(e=e.next)}t.value=s}function v(t,e,a,s){if(!t.value)return!0;var h=t.x-r.x,f=t.y-r.y,p=s-e,d=h*h+f*f;if(p*p/u<d)return d<c&&(0===h&&(d+=(h=o())*h),0===f&&(d+=(f=o())*f),d<l&&(d=Math.sqrt(l*d)),r.vx+=h*t.value*n/d,r.vy+=f*t.value*n/d),!0;if(!(t.length||d>=c)){(t.data!==r||t.next)&&(0===h&&(d+=(h=o())*h),0===f&&(d+=(f=o())*f),d<l&&(d=Math.sqrt(l*d)));do{t.data!==r&&(p=i[t.data.index]*n/d,r.vx+=h*p,r.vy+=f*p)}while(t=t.next)}}return p.initialize=function(e){t=e,d()},p.strength=function(t){return arguments.length?(s="function"==typeof t?t:a(+t),d(),p):s},p.distanceMin=function(t){return arguments.length?(l=t*t,p):Math.sqrt(l)},p.distanceMax=function(t){return arguments.length?(c=t*t,p):Math.sqrt(c)},p.theta=function(t){return arguments.length?(u=t*t,p):Math.sqrt(u)},p},t.forceRadial=function(t,e,r){var n,i,o,s=a(.1);function l(t){for(var a=0,s=n.length;a<s;++a){var l=n[a],c=l.x-e||1e-6,u=l.y-r||1e-6,h=Math.sqrt(c*c+u*u),f=(o[a]-h)*i[a]*t/h;l.vx+=c*f,l.vy+=u*f}}function c(){if(n){var e,r=n.length;for(i=new Array(r),o=new Array(r),e=0;e<r;++e)o[e]=+t(n[e],e,n),i[e]=isNaN(o[e])?0:+s(n[e],e,n)}}return"function"!=typeof t&&(t=a(+t)),null==e&&(e=0),null==r&&(r=0),l.initialize=function(t){n=t,c()},l.strength=function(t){return arguments.length?(s="function"==typeof t?t:a(+t),c(),l):s},l.radius=function(e){return arguments.length?(t="function"==typeof e?e:a(+e),c(),l):t},l.x=function(t){return arguments.length?(e=+t,l):e},l.y=function(t){return arguments.length?(r=+t,l):r},l},t.forceSimulation=function(t){var e,a=1,o=.001,s=1-Math.pow(o,1/300),l=0,c=.6,u=r.map(),h=i.timer(g),f=n.dispatch("tick","end");function g(){v(),f.call("tick",e),a<o&&(h.stop(),f.call("end",e))}function v(){var e,r,n=t.length;for(a+=(l-a)*s,u.each(function(t){t(a)}),e=0;e<n;++e)null==(r=t[e]).fx?r.x+=r.vx*=c:(r.x=r.fx,r.vx=0),null==r.fy?r.y+=r.vy*=c:(r.y=r.fy,r.vy=0)}function m(){for(var e,r=0,n=t.length;r<n;++r){if((e=t[r]).index=r,isNaN(e.x)||isNaN(e.y)){var i=p*Math.sqrt(r),a=r*d;e.x=i*Math.cos(a),e.y=i*Math.sin(a)}(isNaN(e.vx)||isNaN(e.vy))&&(e.vx=e.vy=0)}}function y(e){return e.initialize&&e.initialize(t),e}return null==t&&(t=[]),m(),e={tick:v,restart:function(){return h.restart(g),e},stop:function(){return h.stop(),e},nodes:function(r){return arguments.length?(t=r,m(),u.each(y),e):t},alpha:function(t){return arguments.length?(a=+t,e):a},alphaMin:function(t){return arguments.length?(o=+t,e):o},alphaDecay:function(t){return arguments.length?(s=+t,e):+s},alphaTarget:function(t){return arguments.length?(l=+t,e):l},velocityDecay:function(t){return arguments.length?(c=1-t,e):1-c},force:function(t,r){return arguments.length>1?(null==r?u.remove(t):u.set(t,y(r)),e):u.get(t)},find:function(e,r,n){var i,a,o,s,l,c=0,u=t.length;for(null==n?n=1/0:n*=n,c=0;c<u;++c)(o=(i=e-(s=t[c]).x)*i+(a=r-s.y)*a)<n&&(l=s,n=o);return l},on:function(t,r){return arguments.length>1?(f.on(t,r),e):f.on(t)}}},t.forceX=function(t){var e,r,n,i=a(.1);function o(t){for(var i,a=0,o=e.length;a<o;++a)(i=e[a]).vx+=(n[a]-i.x)*r[a]*t}function s(){if(e){var a,o=e.length;for(r=new Array(o),n=new Array(o),a=0;a<o;++a)r[a]=isNaN(n[a]=+t(e[a],a,e))?0:+i(e[a],a,e)}}return"function"!=typeof t&&(t=a(null==t?0:+t)),o.initialize=function(t){e=t,s()},o.strength=function(t){return arguments.length?(i="function"==typeof t?t:a(+t),s(),o):i},o.x=function(e){return arguments.length?(t="function"==typeof e?e:a(+e),s(),o):t},o},t.forceY=function(t){var e,r,n,i=a(.1);function o(t){for(var i,a=0,o=e.length;a<o;++a)(i=e[a]).vy+=(n[a]-i.y)*r[a]*t}function s(){if(e){var a,o=e.length;for(r=new Array(o),n=new Array(o),a=0;a<o;++a)r[a]=isNaN(n[a]=+t(e[a],a,e))?0:+i(e[a],a,e)}}return"function"!=typeof t&&(t=a(null==t?0:+t)),o.initialize=function(t){e=t,s()},o.strength=function(t){return arguments.length?(i="function"==typeof t?t:a(+t),s(),o):i},o.y=function(e){return arguments.length?(t="function"==typeof e?e:a(+e),s(),o):t},o},Object.defineProperty(t,"__esModule",{value:!0})},"object"==typeof r&&"undefined"!=typeof e?i(r,t("d3-quadtree"),t("d3-collection"),t("d3-dispatch"),t("d3-timer")):i(n.d3=n.d3||{},n.d3,n.d3,n.d3,n.d3)},{"d3-collection":153,"d3-dispatch":155,"d3-quadtree":160,"d3-timer":162}],157:[function(t,e,r){var n;n=this,function(t){"use strict";function e(t,e){return t.parent===e.parent?1:2}function r(t,e){return t+e.x}function n(t,e){return Math.max(t,e.y)}function i(t){var e=0,r=t.children,n=r&&r.length;if(n)for(;--n>=0;)e+=r[n].value;else e=1;t.value=e}function a(t,e){var r,n,i,a,s,u=new c(t),h=+t.value&&(u.value=t.value),f=[u];for(null==e&&(e=o);r=f.pop();)if(h&&(r.value=+r.data.value),(i=e(r.data))&&(s=i.length))for(r.children=new Array(s),a=s-1;a>=0;--a)f.push(n=r.children[a]=new c(i[a])),n.parent=r,n.depth=r.depth+1;return u.eachBefore(l)}function o(t){return t.children}function s(t){t.data=t.data.data}function l(t){var e=0;do{t.height=e}while((t=t.parent)&&t.height<++e)}function c(t){this.data=t,this.depth=this.height=0,this.parent=null}c.prototype=a.prototype={constructor:c,count:function(){return this.eachAfter(i)},each:function(t){var e,r,n,i,a=this,o=[a];do{for(e=o.reverse(),o=[];a=e.pop();)if(t(a),r=a.children)for(n=0,i=r.length;n<i;++n)o.push(r[n])}while(o.length);return this},eachAfter:function(t){for(var e,r,n,i=this,a=[i],o=[];i=a.pop();)if(o.push(i),e=i.children)for(r=0,n=e.length;r<n;++r)a.push(e[r]);for(;i=o.pop();)t(i);return this},eachBefore:function(t){for(var e,r,n=this,i=[n];n=i.pop();)if(t(n),e=n.children)for(r=e.length-1;r>=0;--r)i.push(e[r]);return this},sum:function(t){return this.eachAfter(function(e){for(var r=+t(e.data)||0,n=e.children,i=n&&n.length;--i>=0;)r+=n[i].value;e.value=r})},sort:function(t){return this.eachBefore(function(e){e.children&&e.children.sort(t)})},path:function(t){for(var e=this,r=function(t,e){if(t===e)return t;var r=t.ancestors(),n=e.ancestors(),i=null;for(t=r.pop(),e=n.pop();t===e;)i=t,t=r.pop(),e=n.pop();return i}(e,t),n=[e];e!==r;)e=e.parent,n.push(e);for(var i=n.length;t!==r;)n.splice(i,0,t),t=t.parent;return n},ancestors:function(){for(var t=this,e=[t];t=t.parent;)e.push(t);return e},descendants:function(){var t=[];return this.each(function(e){t.push(e)}),t},leaves:function(){var t=[];return this.eachBefore(function(e){e.children||t.push(e)}),t},links:function(){var t=this,e=[];return t.each(function(r){r!==t&&e.push({source:r.parent,target:r})}),e},copy:function(){return a(this).eachBefore(s)}};var u=Array.prototype.slice;function h(t){for(var e,r,n=0,i=(t=function(t){for(var e,r,n=t.length;n;)r=Math.random()*n--|0,e=t[n],t[n]=t[r],t[r]=e;return t}(u.call(t))).length,a=[];n<i;)e=t[n],r&&d(r,e)?++n:(r=v(a=f(a,e)),n=0);return r}function f(t,e){var r,n;if(g(e,t))return[e];for(r=0;r<t.length;++r)if(p(e,t[r])&&g(m(t[r],e),t))return[t[r],e];for(r=0;r<t.length-1;++r)for(n=r+1;n<t.length;++n)if(p(m(t[r],t[n]),e)&&p(m(t[r],e),t[n])&&p(m(t[n],e),t[r])&&g(y(t[r],t[n],e),t))return[t[r],t[n],e];throw new Error}function p(t,e){var r=t.r-e.r,n=e.x-t.x,i=e.y-t.y;return r<0||r*r<n*n+i*i}function d(t,e){var r=t.r-e.r+1e-6,n=e.x-t.x,i=e.y-t.y;return r>0&&r*r>n*n+i*i}function g(t,e){for(var r=0;r<e.length;++r)if(!d(t,e[r]))return!1;return!0}function v(t){switch(t.length){case 1:return{x:(e=t[0]).x,y:e.y,r:e.r};case 2:return m(t[0],t[1]);case 3:return y(t[0],t[1],t[2])}var e}function m(t,e){var r=t.x,n=t.y,i=t.r,a=e.x,o=e.y,s=e.r,l=a-r,c=o-n,u=s-i,h=Math.sqrt(l*l+c*c);return{x:(r+a+l/h*u)/2,y:(n+o+c/h*u)/2,r:(h+i+s)/2}}function y(t,e,r){var n=t.x,i=t.y,a=t.r,o=e.x,s=e.y,l=e.r,c=r.x,u=r.y,h=r.r,f=n-o,p=n-c,d=i-s,g=i-u,v=l-a,m=h-a,y=n*n+i*i-a*a,x=y-o*o-s*s+l*l,b=y-c*c-u*u+h*h,_=p*d-f*g,w=(d*b-g*x)/(2*_)-n,k=(g*v-d*m)/_,T=(p*x-f*b)/(2*_)-i,A=(f*m-p*v)/_,M=k*k+A*A-1,S=2*(a+w*k+T*A),E=w*w+T*T-a*a,C=-(M?(S+Math.sqrt(S*S-4*M*E))/(2*M):E/S);return{x:n+w+k*C,y:i+T+A*C,r:C}}function x(t,e,r){var n,i,a,o,s=t.x-e.x,l=t.y-e.y,c=s*s+l*l;c?(i=e.r+r.r,i*=i,o=t.r+r.r,i>(o*=o)?(n=(c+o-i)/(2*c),a=Math.sqrt(Math.max(0,o/c-n*n)),r.x=t.x-n*s-a*l,r.y=t.y-n*l+a*s):(n=(c+i-o)/(2*c),a=Math.sqrt(Math.max(0,i/c-n*n)),r.x=e.x+n*s-a*l,r.y=e.y+n*l+a*s)):(r.x=e.x+r.r,r.y=e.y)}function b(t,e){var r=t.r+e.r-1e-6,n=e.x-t.x,i=e.y-t.y;return r>0&&r*r>n*n+i*i}function _(t){var e=t._,r=t.next._,n=e.r+r.r,i=(e.x*r.r+r.x*e.r)/n,a=(e.y*r.r+r.y*e.r)/n;return i*i+a*a}function w(t){this._=t,this.next=null,this.previous=null}function k(t){if(!(i=t.length))return 0;var e,r,n,i,a,o,s,l,c,u,f;if((e=t[0]).x=0,e.y=0,!(i>1))return e.r;if(r=t[1],e.x=-r.r,r.x=e.r,r.y=0,!(i>2))return e.r+r.r;x(r,e,n=t[2]),e=new w(e),r=new w(r),n=new w(n),e.next=n.previous=r,r.next=e.previous=n,n.next=r.previous=e;t:for(s=3;s<i;++s){x(e._,r._,n=t[s]),n=new w(n),l=r.next,c=e.previous,u=r._.r,f=e._.r;do{if(u<=f){if(b(l._,n._)){r=l,e.next=r,r.previous=e,--s;continue t}u+=l._.r,l=l.next}else{if(b(c._,n._)){(e=c).next=r,r.previous=e,--s;continue t}f+=c._.r,c=c.previous}}while(l!==c.next);for(n.previous=e,n.next=r,e.next=r.previous=r=n,a=_(e);(n=n.next)!==r;)(o=_(n))<a&&(e=n,a=o);r=e.next}for(e=[r._],n=r;(n=n.next)!==r;)e.push(n._);for(n=h(e),s=0;s<i;++s)(e=t[s]).x-=n.x,e.y-=n.y;return n.r}function T(t){if("function"!=typeof t)throw new Error;return t}function A(){return 0}function M(t){return function(){return t}}function S(t){return Math.sqrt(t.value)}function E(t){return function(e){e.children||(e.r=Math.max(0,+t(e)||0))}}function C(t,e){return function(r){if(n=r.children){var n,i,a,o=n.length,s=t(r)*e||0;if(s)for(i=0;i<o;++i)n[i].r+=s;if(a=k(n),s)for(i=0;i<o;++i)n[i].r-=s;r.r=a+s}}}function L(t){return function(e){var r=e.parent;e.r*=t,r&&(e.x=r.x+t*e.x,e.y=r.y+t*e.y)}}function P(t){t.x0=Math.round(t.x0),t.y0=Math.round(t.y0),t.x1=Math.round(t.x1),t.y1=Math.round(t.y1)}function O(t,e,r,n,i){for(var a,o=t.children,s=-1,l=o.length,c=t.value&&(n-e)/t.value;++s<l;)(a=o[s]).y0=r,a.y1=i,a.x0=e,a.x1=e+=a.value*c}var z="$",I={depth:-1},D={};function R(t){return t.id}function F(t){return t.parentId}function B(t,e){return t.parent===e.parent?1:2}function N(t){var e=t.children;return e?e[0]:t.t}function j(t){var e=t.children;return e?e[e.length-1]:t.t}function U(t,e,r){var n=r/(e.i-t.i);e.c-=n,e.s+=r,t.c+=n,e.z+=r,e.m+=r}function V(t,e,r){return t.a.parent===e.parent?t.a:r}function q(t,e){this._=t,this.parent=null,this.children=null,this.A=null,this.a=this,this.z=0,this.m=0,this.c=0,this.s=0,this.t=null,this.i=e}function H(t,e,r,n,i){for(var a,o=t.children,s=-1,l=o.length,c=t.value&&(i-r)/t.value;++s<l;)(a=o[s]).x0=e,a.x1=n,a.y0=r,a.y1=r+=a.value*c}q.prototype=Object.create(c.prototype);var G=(1+Math.sqrt(5))/2;function Y(t,e,r,n,i,a){for(var o,s,l,c,u,h,f,p,d,g,v,m=[],y=e.children,x=0,b=0,_=y.length,w=e.value;x<_;){l=i-r,c=a-n;do{u=y[b++].value}while(!u&&b<_);for(h=f=u,v=u*u*(g=Math.max(c/l,l/c)/(w*t)),d=Math.max(f/v,v/h);b<_;++b){if(u+=s=y[b].value,s<h&&(h=s),s>f&&(f=s),v=u*u*g,(p=Math.max(f/v,v/h))>d){u-=s;break}d=p}m.push(o={value:u,dice:l<c,children:y.slice(x,b)}),o.dice?O(o,r,n,i,w?n+=c*u/w:a):H(o,r,n,w?r+=l*u/w:i,a),w-=u,x=b}return m}var W=function t(e){function r(t,r,n,i,a){Y(e,t,r,n,i,a)}return r.ratio=function(e){return t((e=+e)>1?e:1)},r}(G);var X=function t(e){function r(t,r,n,i,a){if((o=t._squarify)&&o.ratio===e)for(var o,s,l,c,u,h=-1,f=o.length,p=t.value;++h<f;){for(l=(s=o[h]).children,c=s.value=0,u=l.length;c<u;++c)s.value+=l[c].value;s.dice?O(s,r,n,i,n+=(a-n)*s.value/p):H(s,r,n,r+=(i-r)*s.value/p,a),p-=s.value}else t._squarify=o=Y(e,t,r,n,i,a),o.ratio=e}return r.ratio=function(e){return t((e=+e)>1?e:1)},r}(G);t.cluster=function(){var t=e,i=1,a=1,o=!1;function s(e){var s,l=0;e.eachAfter(function(e){var i=e.children;i?(e.x=function(t){return t.reduce(r,0)/t.length}(i),e.y=function(t){return 1+t.reduce(n,0)}(i)):(e.x=s?l+=t(e,s):0,e.y=0,s=e)});var c=function(t){for(var e;e=t.children;)t=e[0];return t}(e),u=function(t){for(var e;e=t.children;)t=e[e.length-1];return t}(e),h=c.x-t(c,u)/2,f=u.x+t(u,c)/2;return e.eachAfter(o?function(t){t.x=(t.x-e.x)*i,t.y=(e.y-t.y)*a}:function(t){t.x=(t.x-h)/(f-h)*i,t.y=(1-(e.y?t.y/e.y:1))*a})}return s.separation=function(e){return arguments.length?(t=e,s):t},s.size=function(t){return arguments.length?(o=!1,i=+t[0],a=+t[1],s):o?null:[i,a]},s.nodeSize=function(t){return arguments.length?(o=!0,i=+t[0],a=+t[1],s):o?[i,a]:null},s},t.hierarchy=a,t.pack=function(){var t=null,e=1,r=1,n=A;function i(i){return i.x=e/2,i.y=r/2,t?i.eachBefore(E(t)).eachAfter(C(n,.5)).eachBefore(L(1)):i.eachBefore(E(S)).eachAfter(C(A,1)).eachAfter(C(n,i.r/Math.min(e,r))).eachBefore(L(Math.min(e,r)/(2*i.r))),i}return i.radius=function(e){return arguments.length?(t=null==(r=e)?null:T(r),i):t;var r},i.size=function(t){return arguments.length?(e=+t[0],r=+t[1],i):[e,r]},i.padding=function(t){return arguments.length?(n="function"==typeof t?t:M(+t),i):n},i},t.packSiblings=function(t){return k(t),t},t.packEnclose=h,t.partition=function(){var t=1,e=1,r=0,n=!1;function i(i){var a=i.height+1;return i.x0=i.y0=r,i.x1=t,i.y1=e/a,i.eachBefore(function(t,e){return function(n){n.children&&O(n,n.x0,t*(n.depth+1)/e,n.x1,t*(n.depth+2)/e);var i=n.x0,a=n.y0,o=n.x1-r,s=n.y1-r;o<i&&(i=o=(i+o)/2),s<a&&(a=s=(a+s)/2),n.x0=i,n.y0=a,n.x1=o,n.y1=s}}(e,a)),n&&i.eachBefore(P),i}return i.round=function(t){return arguments.length?(n=!!t,i):n},i.size=function(r){return arguments.length?(t=+r[0],e=+r[1],i):[t,e]},i.padding=function(t){return arguments.length?(r=+t,i):r},i},t.stratify=function(){var t=R,e=F;function r(r){var n,i,a,o,s,u,h,f=r.length,p=new Array(f),d={};for(i=0;i<f;++i)n=r[i],s=p[i]=new c(n),null!=(u=t(n,i,r))&&(u+="")&&(d[h=z+(s.id=u)]=h in d?D:s);for(i=0;i<f;++i)if(s=p[i],null!=(u=e(r[i],i,r))&&(u+="")){if(!(o=d[z+u]))throw new Error("missing: "+u);if(o===D)throw new Error("ambiguous: "+u);o.children?o.children.push(s):o.children=[s],s.parent=o}else{if(a)throw new Error("multiple roots");a=s}if(!a)throw new Error("no root");if(a.parent=I,a.eachBefore(function(t){t.depth=t.parent.depth+1,--f}).eachBefore(l),a.parent=null,f>0)throw new Error("cycle");return a}return r.id=function(e){return arguments.length?(t=T(e),r):t},r.parentId=function(t){return arguments.length?(e=T(t),r):e},r},t.tree=function(){var t=B,e=1,r=1,n=null;function i(i){var l=function(t){for(var e,r,n,i,a,o=new q(t,0),s=[o];e=s.pop();)if(n=e._.children)for(e.children=new Array(a=n.length),i=a-1;i>=0;--i)s.push(r=e.children[i]=new q(n[i],i)),r.parent=e;return(o.parent=new q(null,0)).children=[o],o}(i);if(l.eachAfter(a),l.parent.m=-l.z,l.eachBefore(o),n)i.eachBefore(s);else{var c=i,u=i,h=i;i.eachBefore(function(t){t.x<c.x&&(c=t),t.x>u.x&&(u=t),t.depth>h.depth&&(h=t)});var f=c===u?1:t(c,u)/2,p=f-c.x,d=e/(u.x+f+p),g=r/(h.depth||1);i.eachBefore(function(t){t.x=(t.x+p)*d,t.y=t.depth*g})}return i}function a(e){var r=e.children,n=e.parent.children,i=e.i?n[e.i-1]:null;if(r){!function(t){for(var e,r=0,n=0,i=t.children,a=i.length;--a>=0;)(e=i[a]).z+=r,e.m+=r,r+=e.s+(n+=e.c)}(e);var a=(r[0].z+r[r.length-1].z)/2;i?(e.z=i.z+t(e._,i._),e.m=e.z-a):e.z=a}else i&&(e.z=i.z+t(e._,i._));e.parent.A=function(e,r,n){if(r){for(var i,a=e,o=e,s=r,l=a.parent.children[0],c=a.m,u=o.m,h=s.m,f=l.m;s=j(s),a=N(a),s&&a;)l=N(l),(o=j(o)).a=e,(i=s.z+h-a.z-c+t(s._,a._))>0&&(U(V(s,e,n),e,i),c+=i,u+=i),h+=s.m,c+=a.m,f+=l.m,u+=o.m;s&&!j(o)&&(o.t=s,o.m+=h-u),a&&!N(l)&&(l.t=a,l.m+=c-f,n=e)}return n}(e,i,e.parent.A||n[0])}function o(t){t._.x=t.z+t.parent.m,t.m+=t.parent.m}function s(t){t.x*=e,t.y=t.depth*r}return i.separation=function(e){return arguments.length?(t=e,i):t},i.size=function(t){return arguments.length?(n=!1,e=+t[0],r=+t[1],i):n?null:[e,r]},i.nodeSize=function(t){return arguments.length?(n=!0,e=+t[0],r=+t[1],i):n?[e,r]:null},i},t.treemap=function(){var t=W,e=!1,r=1,n=1,i=[0],a=A,o=A,s=A,l=A,c=A;function u(t){return t.x0=t.y0=0,t.x1=r,t.y1=n,t.eachBefore(h),i=[0],e&&t.eachBefore(P),t}function h(e){var r=i[e.depth],n=e.x0+r,u=e.y0+r,h=e.x1-r,f=e.y1-r;h<n&&(n=h=(n+h)/2),f<u&&(u=f=(u+f)/2),e.x0=n,e.y0=u,e.x1=h,e.y1=f,e.children&&(r=i[e.depth+1]=a(e)/2,n+=c(e)-r,u+=o(e)-r,(h-=s(e)-r)<n&&(n=h=(n+h)/2),(f-=l(e)-r)<u&&(u=f=(u+f)/2),t(e,n,u,h,f))}return u.round=function(t){return arguments.length?(e=!!t,u):e},u.size=function(t){return arguments.length?(r=+t[0],n=+t[1],u):[r,n]},u.tile=function(e){return arguments.length?(t=T(e),u):t},u.padding=function(t){return arguments.length?u.paddingInner(t).paddingOuter(t):u.paddingInner()},u.paddingInner=function(t){return arguments.length?(a="function"==typeof t?t:M(+t),u):a},u.paddingOuter=function(t){return arguments.length?u.paddingTop(t).paddingRight(t).paddingBottom(t).paddingLeft(t):u.paddingTop()},u.paddingTop=function(t){return arguments.length?(o="function"==typeof t?t:M(+t),u):o},u.paddingRight=function(t){return arguments.length?(s="function"==typeof t?t:M(+t),u):s},u.paddingBottom=function(t){return arguments.length?(l="function"==typeof t?t:M(+t),u):l},u.paddingLeft=function(t){return arguments.length?(c="function"==typeof t?t:M(+t),u):c},u},t.treemapBinary=function(t,e,r,n,i){var a,o,s=t.children,l=s.length,c=new Array(l+1);for(c[0]=o=a=0;a<l;++a)c[a+1]=o+=s[a].value;!function t(e,r,n,i,a,o,l){if(e>=r-1){var u=s[e];return u.x0=i,u.y0=a,u.x1=o,void(u.y1=l)}for(var h=c[e],f=n/2+h,p=e+1,d=r-1;p<d;){var g=p+d>>>1;c[g]<f?p=g+1:d=g}f-c[p-1]<c[p]-f&&e+1<p&&--p;var v=c[p]-h,m=n-v;if(o-i>l-a){var y=(i*m+o*v)/n;t(e,p,v,i,a,y,l),t(p,r,m,y,a,o,l)}else{var x=(a*m+l*v)/n;t(e,p,v,i,a,o,x),t(p,r,m,i,x,o,l)}}(0,l,t.value,e,r,n,i)},t.treemapDice=O,t.treemapSlice=H,t.treemapSliceDice=function(t,e,r,n,i){(1&t.depth?H:O)(t,e,r,n,i)},t.treemapSquarify=W,t.treemapResquarify=X,Object.defineProperty(t,"__esModule",{value:!0})}("object"==typeof r&&"undefined"!=typeof e?r:n.d3=n.d3||{})},{}],158:[function(t,e,r){var n,i;n=this,i=function(t,e){"use strict";function r(t,e,r,n,i){var a=t*t,o=a*t;return((1-3*t+3*a-o)*e+(4-6*a+3*o)*r+(1+3*t+3*a-3*o)*n+o*i)/6}function n(t){var e=t.length-1;return function(n){var i=n<=0?n=0:n>=1?(n=1,e-1):Math.floor(n*e),a=t[i],o=t[i+1],s=i>0?t[i-1]:2*a-o,l=i<e-1?t[i+2]:2*o-a;return r((n-i/e)*e,s,a,o,l)}}function i(t){var e=t.length;return function(n){var i=Math.floor(((n%=1)<0?++n:n)*e),a=t[(i+e-1)%e],o=t[i%e],s=t[(i+1)%e],l=t[(i+2)%e];return r((n-i/e)*e,a,o,s,l)}}function a(t){return function(){return t}}function o(t,e){return function(r){return t+r*e}}function s(t,e){var r=e-t;return r?o(t,r>180||r<-180?r-360*Math.round(r/360):r):a(isNaN(t)?e:t)}function l(t){return 1==(t=+t)?c:function(e,r){return r-e?function(t,e,r){return t=Math.pow(t,r),e=Math.pow(e,r)-t,r=1/r,function(n){return Math.pow(t+n*e,r)}}(e,r,t):a(isNaN(e)?r:e)}}function c(t,e){var r=e-t;return r?o(t,r):a(isNaN(t)?e:t)}var u=function t(r){var n=l(r);function i(t,r){var i=n((t=e.rgb(t)).r,(r=e.rgb(r)).r),a=n(t.g,r.g),o=n(t.b,r.b),s=c(t.opacity,r.opacity);return function(e){return t.r=i(e),t.g=a(e),t.b=o(e),t.opacity=s(e),t+""}}return i.gamma=t,i}(1);function h(t){return function(r){var n,i,a=r.length,o=new Array(a),s=new Array(a),l=new Array(a);for(n=0;n<a;++n)i=e.rgb(r[n]),o[n]=i.r||0,s[n]=i.g||0,l[n]=i.b||0;return o=t(o),s=t(s),l=t(l),i.opacity=1,function(t){return i.r=o(t),i.g=s(t),i.b=l(t),i+""}}}var f=h(n),p=h(i);function d(t,e){var r,n=e?e.length:0,i=t?Math.min(n,t.length):0,a=new Array(i),o=new Array(n);for(r=0;r<i;++r)a[r]=_(t[r],e[r]);for(;r<n;++r)o[r]=e[r];return function(t){for(r=0;r<i;++r)o[r]=a[r](t);return o}}function g(t,e){var r=new Date;return e-=t=+t,function(n){return r.setTime(t+e*n),r}}function v(t,e){return e-=t=+t,function(r){return t+e*r}}function m(t,e){var r,n={},i={};for(r in null!==t&&"object"==typeof t||(t={}),null!==e&&"object"==typeof e||(e={}),e)r in t?n[r]=_(t[r],e[r]):i[r]=e[r];return function(t){for(r in n)i[r]=n[r](t);return i}}var y=/[-+]?(?:\d+\.?\d*|\.?\d+)(?:[eE][-+]?\d+)?/g,x=new RegExp(y.source,"g");function b(t,e){var r,n,i,a=y.lastIndex=x.lastIndex=0,o=-1,s=[],l=[];for(t+="",e+="";(r=y.exec(t))&&(n=x.exec(e));)(i=n.index)>a&&(i=e.slice(a,i),s[o]?s[o]+=i:s[++o]=i),(r=r[0])===(n=n[0])?s[o]?s[o]+=n:s[++o]=n:(s[++o]=null,l.push({i:o,x:v(r,n)})),a=x.lastIndex;return a<e.length&&(i=e.slice(a),s[o]?s[o]+=i:s[++o]=i),s.length<2?l[0]?function(t){return function(e){return t(e)+""}}(l[0].x):function(t){return function(){return t}}(e):(e=l.length,function(t){for(var r,n=0;n<e;++n)s[(r=l[n]).i]=r.x(t);return s.join("")})}function _(t,r){var n,i=typeof r;return null==r||"boolean"===i?a(r):("number"===i?v:"string"===i?(n=e.color(r))?(r=n,u):b:r instanceof e.color?u:r instanceof Date?g:Array.isArray(r)?d:"function"!=typeof r.valueOf&&"function"!=typeof r.toString||isNaN(r)?m:v)(t,r)}var w,k,T,A,M=180/Math.PI,S={translateX:0,translateY:0,rotate:0,skewX:0,scaleX:1,scaleY:1};function E(t,e,r,n,i,a){var o,s,l;return(o=Math.sqrt(t*t+e*e))&&(t/=o,e/=o),(l=t*r+e*n)&&(r-=t*l,n-=e*l),(s=Math.sqrt(r*r+n*n))&&(r/=s,n/=s,l/=s),t*n<e*r&&(t=-t,e=-e,l=-l,o=-o),{translateX:i,translateY:a,rotate:Math.atan2(e,t)*M,skewX:Math.atan(l)*M,scaleX:o,scaleY:s}}function C(t,e,r,n){function i(t){return t.length?t.pop()+" ":""}return function(a,o){var s=[],l=[];return a=t(a),o=t(o),function(t,n,i,a,o,s){if(t!==i||n!==a){var l=o.push("translate(",null,e,null,r);s.push({i:l-4,x:v(t,i)},{i:l-2,x:v(n,a)})}else(i||a)&&o.push("translate("+i+e+a+r)}(a.translateX,a.translateY,o.translateX,o.translateY,s,l),function(t,e,r,a){t!==e?(t-e>180?e+=360:e-t>180&&(t+=360),a.push({i:r.push(i(r)+"rotate(",null,n)-2,x:v(t,e)})):e&&r.push(i(r)+"rotate("+e+n)}(a.rotate,o.rotate,s,l),function(t,e,r,a){t!==e?a.push({i:r.push(i(r)+"skewX(",null,n)-2,x:v(t,e)}):e&&r.push(i(r)+"skewX("+e+n)}(a.skewX,o.skewX,s,l),function(t,e,r,n,a,o){if(t!==r||e!==n){var s=a.push(i(a)+"scale(",null,",",null,")");o.push({i:s-4,x:v(t,r)},{i:s-2,x:v(e,n)})}else 1===r&&1===n||a.push(i(a)+"scale("+r+","+n+")")}(a.scaleX,a.scaleY,o.scaleX,o.scaleY,s,l),a=o=null,function(t){for(var e,r=-1,n=l.length;++r<n;)s[(e=l[r]).i]=e.x(t);return s.join("")}}}var L=C(function(t){return"none"===t?S:(w||(w=document.createElement("DIV"),k=document.documentElement,T=document.defaultView),w.style.transform=t,t=T.getComputedStyle(k.appendChild(w),null).getPropertyValue("transform"),k.removeChild(w),E(+(t=t.slice(7,-1).split(","))[0],+t[1],+t[2],+t[3],+t[4],+t[5]))},"px, ","px)","deg)"),P=C(function(t){return null==t?S:(A||(A=document.createElementNS("http://www.w3.org/2000/svg","g")),A.setAttribute("transform",t),(t=A.transform.baseVal.consolidate())?E((t=t.matrix).a,t.b,t.c,t.d,t.e,t.f):S)},", ",")",")"),O=Math.SQRT2,z=2,I=4,D=1e-12;function R(t){return((t=Math.exp(t))+1/t)/2}function F(t){return function(r,n){var i=t((r=e.hsl(r)).h,(n=e.hsl(n)).h),a=c(r.s,n.s),o=c(r.l,n.l),s=c(r.opacity,n.opacity);return function(t){return r.h=i(t),r.s=a(t),r.l=o(t),r.opacity=s(t),r+""}}}var B=F(s),N=F(c);function j(t){return function(r,n){var i=t((r=e.hcl(r)).h,(n=e.hcl(n)).h),a=c(r.c,n.c),o=c(r.l,n.l),s=c(r.opacity,n.opacity);return function(t){return r.h=i(t),r.c=a(t),r.l=o(t),r.opacity=s(t),r+""}}}var U=j(s),V=j(c);function q(t){return function r(n){function i(r,i){var a=t((r=e.cubehelix(r)).h,(i=e.cubehelix(i)).h),o=c(r.s,i.s),s=c(r.l,i.l),l=c(r.opacity,i.opacity);return function(t){return r.h=a(t),r.s=o(t),r.l=s(Math.pow(t,n)),r.opacity=l(t),r+""}}return n=+n,i.gamma=r,i}(1)}var H=q(s),G=q(c);t.interpolate=_,t.interpolateArray=d,t.interpolateBasis=n,t.interpolateBasisClosed=i,t.interpolateDate=g,t.interpolateDiscrete=function(t){var e=t.length;return function(r){return t[Math.max(0,Math.min(e-1,Math.floor(r*e)))]}},t.interpolateHue=function(t,e){var r=s(+t,+e);return function(t){var e=r(t);return e-360*Math.floor(e/360)}},t.interpolateNumber=v,t.interpolateObject=m,t.interpolateRound=function(t,e){return e-=t=+t,function(r){return Math.round(t+e*r)}},t.interpolateString=b,t.interpolateTransformCss=L,t.interpolateTransformSvg=P,t.interpolateZoom=function(t,e){var r,n,i=t[0],a=t[1],o=t[2],s=e[0],l=e[1],c=e[2],u=s-i,h=l-a,f=u*u+h*h;if(f<D)n=Math.log(c/o)/O,r=function(t){return[i+t*u,a+t*h,o*Math.exp(O*t*n)]};else{var p=Math.sqrt(f),d=(c*c-o*o+I*f)/(2*o*z*p),g=(c*c-o*o-I*f)/(2*c*z*p),v=Math.log(Math.sqrt(d*d+1)-d),m=Math.log(Math.sqrt(g*g+1)-g);n=(m-v)/O,r=function(t){var e,r=t*n,s=R(v),l=o/(z*p)*(s*(e=O*r+v,((e=Math.exp(2*e))-1)/(e+1))-function(t){return((t=Math.exp(t))-1/t)/2}(v));return[i+l*u,a+l*h,o*s/R(O*r+v)]}}return r.duration=1e3*n,r},t.interpolateRgb=u,t.interpolateRgbBasis=f,t.interpolateRgbBasisClosed=p,t.interpolateHsl=B,t.interpolateHslLong=N,t.interpolateLab=function(t,r){var n=c((t=e.lab(t)).l,(r=e.lab(r)).l),i=c(t.a,r.a),a=c(t.b,r.b),o=c(t.opacity,r.opacity);return function(e){return t.l=n(e),t.a=i(e),t.b=a(e),t.opacity=o(e),t+""}},t.interpolateHcl=U,t.interpolateHclLong=V,t.interpolateCubehelix=H,t.interpolateCubehelixLong=G,t.piecewise=function(t,e){for(var r=0,n=e.length-1,i=e[0],a=new Array(n<0?0:n);r<n;)a[r]=t(i,i=e[++r]);return function(t){var e=Math.max(0,Math.min(n-1,Math.floor(t*=n)));return a[e](t-e)}},t.quantize=function(t,e){for(var r=new Array(e),n=0;n<e;++n)r[n]=t(n/(e-1));return r},Object.defineProperty(t,"__esModule",{value:!0})},"object"==typeof r&&"undefined"!=typeof e?i(r,t("d3-color")):i(n.d3=n.d3||{},n.d3)},{"d3-color":154}],159:[function(t,e,r){var n;n=this,function(t){"use strict";var e=Math.PI,r=2*e,n=r-1e-6;function i(){this._x0=this._y0=this._x1=this._y1=null,this._=""}function a(){return new i}i.prototype=a.prototype={constructor:i,moveTo:function(t,e){this._+="M"+(this._x0=this._x1=+t)+","+(this._y0=this._y1=+e)},closePath:function(){null!==this._x1&&(this._x1=this._x0,this._y1=this._y0,this._+="Z")},lineTo:function(t,e){this._+="L"+(this._x1=+t)+","+(this._y1=+e)},quadraticCurveTo:function(t,e,r,n){this._+="Q"+ +t+","+ +e+","+(this._x1=+r)+","+(this._y1=+n)},bezierCurveTo:function(t,e,r,n,i,a){this._+="C"+ +t+","+ +e+","+ +r+","+ +n+","+(this._x1=+i)+","+(this._y1=+a)},arcTo:function(t,r,n,i,a){t=+t,r=+r,n=+n,i=+i,a=+a;var o=this._x1,s=this._y1,l=n-t,c=i-r,u=o-t,h=s-r,f=u*u+h*h;if(a<0)throw new Error("negative radius: "+a);if(null===this._x1)this._+="M"+(this._x1=t)+","+(this._y1=r);else if(f>1e-6)if(Math.abs(h*l-c*u)>1e-6&&a){var p=n-o,d=i-s,g=l*l+c*c,v=p*p+d*d,m=Math.sqrt(g),y=Math.sqrt(f),x=a*Math.tan((e-Math.acos((g+f-v)/(2*m*y)))/2),b=x/y,_=x/m;Math.abs(b-1)>1e-6&&(this._+="L"+(t+b*u)+","+(r+b*h)),this._+="A"+a+","+a+",0,0,"+ +(h*p>u*d)+","+(this._x1=t+_*l)+","+(this._y1=r+_*c)}else this._+="L"+(this._x1=t)+","+(this._y1=r);else;},arc:function(t,i,a,o,s,l){t=+t,i=+i;var c=(a=+a)*Math.cos(o),u=a*Math.sin(o),h=t+c,f=i+u,p=1^l,d=l?o-s:s-o;if(a<0)throw new Error("negative radius: "+a);null===this._x1?this._+="M"+h+","+f:(Math.abs(this._x1-h)>1e-6||Math.abs(this._y1-f)>1e-6)&&(this._+="L"+h+","+f),a&&(d<0&&(d=d%r+r),d>n?this._+="A"+a+","+a+",0,1,"+p+","+(t-c)+","+(i-u)+"A"+a+","+a+",0,1,"+p+","+(this._x1=h)+","+(this._y1=f):d>1e-6&&(this._+="A"+a+","+a+",0,"+ +(d>=e)+","+p+","+(this._x1=t+a*Math.cos(s))+","+(this._y1=i+a*Math.sin(s))))},rect:function(t,e,r,n){this._+="M"+(this._x0=this._x1=+t)+","+(this._y0=this._y1=+e)+"h"+ +r+"v"+ +n+"h"+-r+"Z"},toString:function(){return this._}},t.path=a,Object.defineProperty(t,"__esModule",{value:!0})}("object"==typeof r&&"undefined"!=typeof e?r:n.d3=n.d3||{})},{}],160:[function(t,e,r){var n;n=this,function(t){"use strict";function e(t,e,r,n){if(isNaN(e)||isNaN(r))return t;var i,a,o,s,l,c,u,h,f,p=t._root,d={data:n},g=t._x0,v=t._y0,m=t._x1,y=t._y1;if(!p)return t._root=d,t;for(;p.length;)if((c=e>=(a=(g+m)/2))?g=a:m=a,(u=r>=(o=(v+y)/2))?v=o:y=o,i=p,!(p=p[h=u<<1|c]))return i[h]=d,t;if(s=+t._x.call(null,p.data),l=+t._y.call(null,p.data),e===s&&r===l)return d.next=p,i?i[h]=d:t._root=d,t;do{i=i?i[h]=new Array(4):t._root=new Array(4),(c=e>=(a=(g+m)/2))?g=a:m=a,(u=r>=(o=(v+y)/2))?v=o:y=o}while((h=u<<1|c)==(f=(l>=o)<<1|s>=a));return i[f]=p,i[h]=d,t}var r=function(t,e,r,n,i){this.node=t,this.x0=e,this.y0=r,this.x1=n,this.y1=i};function n(t){return t[0]}function i(t){return t[1]}function a(t,e,r){var a=new o(null==e?n:e,null==r?i:r,NaN,NaN,NaN,NaN);return null==t?a:a.addAll(t)}function o(t,e,r,n,i,a){this._x=t,this._y=e,this._x0=r,this._y0=n,this._x1=i,this._y1=a,this._root=void 0}function s(t){for(var e={data:t.data},r=e;t=t.next;)r=r.next={data:t.data};return e}var l=a.prototype=o.prototype;l.copy=function(){var t,e,r=new o(this._x,this._y,this._x0,this._y0,this._x1,this._y1),n=this._root;if(!n)return r;if(!n.length)return r._root=s(n),r;for(t=[{source:n,target:r._root=new Array(4)}];n=t.pop();)for(var i=0;i<4;++i)(e=n.source[i])&&(e.length?t.push({source:e,target:n.target[i]=new Array(4)}):n.target[i]=s(e));return r},l.add=function(t){var r=+this._x.call(null,t),n=+this._y.call(null,t);return e(this.cover(r,n),r,n,t)},l.addAll=function(t){var r,n,i,a,o=t.length,s=new Array(o),l=new Array(o),c=1/0,u=1/0,h=-1/0,f=-1/0;for(n=0;n<o;++n)isNaN(i=+this._x.call(null,r=t[n]))||isNaN(a=+this._y.call(null,r))||(s[n]=i,l[n]=a,i<c&&(c=i),i>h&&(h=i),a<u&&(u=a),a>f&&(f=a));for(h<c&&(c=this._x0,h=this._x1),f<u&&(u=this._y0,f=this._y1),this.cover(c,u).cover(h,f),n=0;n<o;++n)e(this,s[n],l[n],t[n]);return this},l.cover=function(t,e){if(isNaN(t=+t)||isNaN(e=+e))return this;var r=this._x0,n=this._y0,i=this._x1,a=this._y1;if(isNaN(r))i=(r=Math.floor(t))+1,a=(n=Math.floor(e))+1;else{if(!(r>t||t>i||n>e||e>a))return this;var o,s,l=i-r,c=this._root;switch(s=(e<(n+a)/2)<<1|t<(r+i)/2){case 0:do{(o=new Array(4))[s]=c,c=o}while(a=n+(l*=2),t>(i=r+l)||e>a);break;case 1:do{(o=new Array(4))[s]=c,c=o}while(a=n+(l*=2),(r=i-l)>t||e>a);break;case 2:do{(o=new Array(4))[s]=c,c=o}while(n=a-(l*=2),t>(i=r+l)||n>e);break;case 3:do{(o=new Array(4))[s]=c,c=o}while(n=a-(l*=2),(r=i-l)>t||n>e)}this._root&&this._root.length&&(this._root=c)}return this._x0=r,this._y0=n,this._x1=i,this._y1=a,this},l.data=function(){var t=[];return this.visit(function(e){if(!e.length)do{t.push(e.data)}while(e=e.next)}),t},l.extent=function(t){return arguments.length?this.cover(+t[0][0],+t[0][1]).cover(+t[1][0],+t[1][1]):isNaN(this._x0)?void 0:[[this._x0,this._y0],[this._x1,this._y1]]},l.find=function(t,e,n){var i,a,o,s,l,c,u,h=this._x0,f=this._y0,p=this._x1,d=this._y1,g=[],v=this._root;for(v&&g.push(new r(v,h,f,p,d)),null==n?n=1/0:(h=t-n,f=e-n,p=t+n,d=e+n,n*=n);c=g.pop();)if(!(!(v=c.node)||(a=c.x0)>p||(o=c.y0)>d||(s=c.x1)<h||(l=c.y1)<f))if(v.length){var m=(a+s)/2,y=(o+l)/2;g.push(new r(v[3],m,y,s,l),new r(v[2],a,y,m,l),new r(v[1],m,o,s,y),new r(v[0],a,o,m,y)),(u=(e>=y)<<1|t>=m)&&(c=g[g.length-1],g[g.length-1]=g[g.length-1-u],g[g.length-1-u]=c)}else{var x=t-+this._x.call(null,v.data),b=e-+this._y.call(null,v.data),_=x*x+b*b;if(_<n){var w=Math.sqrt(n=_);h=t-w,f=e-w,p=t+w,d=e+w,i=v.data}}return i},l.remove=function(t){if(isNaN(a=+this._x.call(null,t))||isNaN(o=+this._y.call(null,t)))return this;var e,r,n,i,a,o,s,l,c,u,h,f,p=this._root,d=this._x0,g=this._y0,v=this._x1,m=this._y1;if(!p)return this;if(p.length)for(;;){if((c=a>=(s=(d+v)/2))?d=s:v=s,(u=o>=(l=(g+m)/2))?g=l:m=l,e=p,!(p=p[h=u<<1|c]))return this;if(!p.length)break;(e[h+1&3]||e[h+2&3]||e[h+3&3])&&(r=e,f=h)}for(;p.data!==t;)if(n=p,!(p=p.next))return this;return(i=p.next)&&delete p.next,n?(i?n.next=i:delete n.next,this):e?(i?e[h]=i:delete e[h],(p=e[0]||e[1]||e[2]||e[3])&&p===(e[3]||e[2]||e[1]||e[0])&&!p.length&&(r?r[f]=p:this._root=p),this):(this._root=i,this)},l.removeAll=function(t){for(var e=0,r=t.length;e<r;++e)this.remove(t[e]);return this},l.root=function(){return this._root},l.size=function(){var t=0;return this.visit(function(e){if(!e.length)do{++t}while(e=e.next)}),t},l.visit=function(t){var e,n,i,a,o,s,l=[],c=this._root;for(c&&l.push(new r(c,this._x0,this._y0,this._x1,this._y1));e=l.pop();)if(!t(c=e.node,i=e.x0,a=e.y0,o=e.x1,s=e.y1)&&c.length){var u=(i+o)/2,h=(a+s)/2;(n=c[3])&&l.push(new r(n,u,h,o,s)),(n=c[2])&&l.push(new r(n,i,h,u,s)),(n=c[1])&&l.push(new r(n,u,a,o,h)),(n=c[0])&&l.push(new r(n,i,a,u,h))}return this},l.visitAfter=function(t){var e,n=[],i=[];for(this._root&&n.push(new r(this._root,this._x0,this._y0,this._x1,this._y1));e=n.pop();){var a=e.node;if(a.length){var o,s=e.x0,l=e.y0,c=e.x1,u=e.y1,h=(s+c)/2,f=(l+u)/2;(o=a[0])&&n.push(new r(o,s,l,h,f)),(o=a[1])&&n.push(new r(o,h,l,c,f)),(o=a[2])&&n.push(new r(o,s,f,h,u)),(o=a[3])&&n.push(new r(o,h,f,c,u))}i.push(e)}for(;e=i.pop();)t(e.node,e.x0,e.y0,e.x1,e.y1);return this},l.x=function(t){return arguments.length?(this._x=t,this):this._x},l.y=function(t){return arguments.length?(this._y=t,this):this._y},t.quadtree=a,Object.defineProperty(t,"__esModule",{value:!0})}("object"==typeof r&&"undefined"!=typeof e?r:n.d3=n.d3||{})},{}],161:[function(t,e,r){var n,i;n=this,i=function(t,e){"use strict";function r(t){return function(){return t}}var n=Math.abs,i=Math.atan2,a=Math.cos,o=Math.max,s=Math.min,l=Math.sin,c=Math.sqrt,u=1e-12,h=Math.PI,f=h/2,p=2*h;function d(t){return t>=1?f:t<=-1?-f:Math.asin(t)}function g(t){return t.innerRadius}function v(t){return t.outerRadius}function m(t){return t.startAngle}function y(t){return t.endAngle}function x(t){return t&&t.padAngle}function b(t,e,r,n,i,a,s){var l=t-r,u=e-n,h=(s?a:-a)/c(l*l+u*u),f=h*u,p=-h*l,d=t+f,g=e+p,v=r+f,m=n+p,y=(d+v)/2,x=(g+m)/2,b=v-d,_=m-g,w=b*b+_*_,k=i-a,T=d*m-v*g,A=(_<0?-1:1)*c(o(0,k*k*w-T*T)),M=(T*_-b*A)/w,S=(-T*b-_*A)/w,E=(T*_+b*A)/w,C=(-T*b+_*A)/w,L=M-y,P=S-x,O=E-y,z=C-x;return L*L+P*P>O*O+z*z&&(M=E,S=C),{cx:M,cy:S,x01:-f,y01:-p,x11:M*(i/k-1),y11:S*(i/k-1)}}function _(t){this._context=t}function w(t){return new _(t)}function k(t){return t[0]}function T(t){return t[1]}function A(){var t=k,n=T,i=r(!0),a=null,o=w,s=null;function l(r){var l,c,u,h=r.length,f=!1;for(null==a&&(s=o(u=e.path())),l=0;l<=h;++l)!(l<h&&i(c=r[l],l,r))===f&&((f=!f)?s.lineStart():s.lineEnd()),f&&s.point(+t(c,l,r),+n(c,l,r));if(u)return s=null,u+""||null}return l.x=function(e){return arguments.length?(t="function"==typeof e?e:r(+e),l):t},l.y=function(t){return arguments.length?(n="function"==typeof t?t:r(+t),l):n},l.defined=function(t){return arguments.length?(i="function"==typeof t?t:r(!!t),l):i},l.curve=function(t){return arguments.length?(o=t,null!=a&&(s=o(a)),l):o},l.context=function(t){return arguments.length?(null==t?a=s=null:s=o(a=t),l):a},l}function M(){var t=k,n=null,i=r(0),a=T,o=r(!0),s=null,l=w,c=null;function u(r){var u,h,f,p,d,g=r.length,v=!1,m=new Array(g),y=new Array(g);for(null==s&&(c=l(d=e.path())),u=0;u<=g;++u){if(!(u<g&&o(p=r[u],u,r))===v)if(v=!v)h=u,c.areaStart(),c.lineStart();else{for(c.lineEnd(),c.lineStart(),f=u-1;f>=h;--f)c.point(m[f],y[f]);c.lineEnd(),c.areaEnd()}v&&(m[u]=+t(p,u,r),y[u]=+i(p,u,r),c.point(n?+n(p,u,r):m[u],a?+a(p,u,r):y[u]))}if(d)return c=null,d+""||null}function h(){return A().defined(o).curve(l).context(s)}return u.x=function(e){return arguments.length?(t="function"==typeof e?e:r(+e),n=null,u):t},u.x0=function(e){return arguments.length?(t="function"==typeof e?e:r(+e),u):t},u.x1=function(t){return arguments.length?(n=null==t?null:"function"==typeof t?t:r(+t),u):n},u.y=function(t){return arguments.length?(i="function"==typeof t?t:r(+t),a=null,u):i},u.y0=function(t){return arguments.length?(i="function"==typeof t?t:r(+t),u):i},u.y1=function(t){return arguments.length?(a=null==t?null:"function"==typeof t?t:r(+t),u):a},u.lineX0=u.lineY0=function(){return h().x(t).y(i)},u.lineY1=function(){return h().x(t).y(a)},u.lineX1=function(){return h().x(n).y(i)},u.defined=function(t){return arguments.length?(o="function"==typeof t?t:r(!!t),u):o},u.curve=function(t){return arguments.length?(l=t,null!=s&&(c=l(s)),u):l},u.context=function(t){return arguments.length?(null==t?s=c=null:c=l(s=t),u):s},u}function S(t,e){return e<t?-1:e>t?1:e>=t?0:NaN}function E(t){return t}_.prototype={areaStart:function(){this._line=0},areaEnd:function(){this._line=NaN},lineStart:function(){this._point=0},lineEnd:function(){(this._line||0!==this._line&&1===this._point)&&this._context.closePath(),this._line=1-this._line},point:function(t,e){switch(t=+t,e=+e,this._point){case 0:this._point=1,this._line?this._context.lineTo(t,e):this._context.moveTo(t,e);break;case 1:this._point=2;default:this._context.lineTo(t,e)}}};var C=P(w);function L(t){this._curve=t}function P(t){function e(e){return new L(t(e))}return e._curve=t,e}function O(t){var e=t.curve;return t.angle=t.x,delete t.x,t.radius=t.y,delete t.y,t.curve=function(t){return arguments.length?e(P(t)):e()._curve},t}function z(){return O(A().curve(C))}function I(){var t=M().curve(C),e=t.curve,r=t.lineX0,n=t.lineX1,i=t.lineY0,a=t.lineY1;return t.angle=t.x,delete t.x,t.startAngle=t.x0,delete t.x0,t.endAngle=t.x1,delete t.x1,t.radius=t.y,delete t.y,t.innerRadius=t.y0,delete t.y0,t.outerRadius=t.y1,delete t.y1,t.lineStartAngle=function(){return O(r())},delete t.lineX0,t.lineEndAngle=function(){return O(n())},delete t.lineX1,t.lineInnerRadius=function(){return O(i())},delete t.lineY0,t.lineOuterRadius=function(){return O(a())},delete t.lineY1,t.curve=function(t){return arguments.length?e(P(t)):e()._curve},t}function D(t,e){return[(e=+e)*Math.cos(t-=Math.PI/2),e*Math.sin(t)]}L.prototype={areaStart:function(){this._curve.areaStart()},areaEnd:function(){this._curve.areaEnd()},lineStart:function(){this._curve.lineStart()},lineEnd:function(){this._curve.lineEnd()},point:function(t,e){this._curve.point(e*Math.sin(t),e*-Math.cos(t))}};var R=Array.prototype.slice;function F(t){return t.source}function B(t){return t.target}function N(t){var n=F,i=B,a=k,o=T,s=null;function l(){var r,l=R.call(arguments),c=n.apply(this,l),u=i.apply(this,l);if(s||(s=r=e.path()),t(s,+a.apply(this,(l[0]=c,l)),+o.apply(this,l),+a.apply(this,(l[0]=u,l)),+o.apply(this,l)),r)return s=null,r+""||null}return l.source=function(t){return arguments.length?(n=t,l):n},l.target=function(t){return arguments.length?(i=t,l):i},l.x=function(t){return arguments.length?(a="function"==typeof t?t:r(+t),l):a},l.y=function(t){return arguments.length?(o="function"==typeof t?t:r(+t),l):o},l.context=function(t){return arguments.length?(s=null==t?null:t,l):s},l}function j(t,e,r,n,i){t.moveTo(e,r),t.bezierCurveTo(e=(e+n)/2,r,e,i,n,i)}function U(t,e,r,n,i){t.moveTo(e,r),t.bezierCurveTo(e,r=(r+i)/2,n,r,n,i)}function V(t,e,r,n,i){var a=D(e,r),o=D(e,r=(r+i)/2),s=D(n,r),l=D(n,i);t.moveTo(a[0],a[1]),t.bezierCurveTo(o[0],o[1],s[0],s[1],l[0],l[1])}var q={draw:function(t,e){var r=Math.sqrt(e/h);t.moveTo(r,0),t.arc(0,0,r,0,p)}},H={draw:function(t,e){var r=Math.sqrt(e/5)/2;t.moveTo(-3*r,-r),t.lineTo(-r,-r),t.lineTo(-r,-3*r),t.lineTo(r,-3*r),t.lineTo(r,-r),t.lineTo(3*r,-r),t.lineTo(3*r,r),t.lineTo(r,r),t.lineTo(r,3*r),t.lineTo(-r,3*r),t.lineTo(-r,r),t.lineTo(-3*r,r),t.closePath()}},G=Math.sqrt(1/3),Y=2*G,W={draw:function(t,e){var r=Math.sqrt(e/Y),n=r*G;t.moveTo(0,-r),t.lineTo(n,0),t.lineTo(0,r),t.lineTo(-n,0),t.closePath()}},X=Math.sin(h/10)/Math.sin(7*h/10),Z=Math.sin(p/10)*X,J=-Math.cos(p/10)*X,K={draw:function(t,e){var r=Math.sqrt(.8908130915292852*e),n=Z*r,i=J*r;t.moveTo(0,-r),t.lineTo(n,i);for(var a=1;a<5;++a){var o=p*a/5,s=Math.cos(o),l=Math.sin(o);t.lineTo(l*r,-s*r),t.lineTo(s*n-l*i,l*n+s*i)}t.closePath()}},$={draw:function(t,e){var r=Math.sqrt(e),n=-r/2;t.rect(n,n,r,r)}},Q=Math.sqrt(3),tt={draw:function(t,e){var r=-Math.sqrt(e/(3*Q));t.moveTo(0,2*r),t.lineTo(-Q*r,-r),t.lineTo(Q*r,-r),t.closePath()}},et=-.5,rt=Math.sqrt(3)/2,nt=1/Math.sqrt(12),it=3*(nt/2+1),at={draw:function(t,e){var r=Math.sqrt(e/it),n=r/2,i=r*nt,a=n,o=r*nt+r,s=-a,l=o;t.moveTo(n,i),t.lineTo(a,o),t.lineTo(s,l),t.lineTo(et*n-rt*i,rt*n+et*i),t.lineTo(et*a-rt*o,rt*a+et*o),t.lineTo(et*s-rt*l,rt*s+et*l),t.lineTo(et*n+rt*i,et*i-rt*n),t.lineTo(et*a+rt*o,et*o-rt*a),t.lineTo(et*s+rt*l,et*l-rt*s),t.closePath()}},ot=[q,H,W,$,K,tt,at];function st(){}function lt(t,e,r){t._context.bezierCurveTo((2*t._x0+t._x1)/3,(2*t._y0+t._y1)/3,(t._x0+2*t._x1)/3,(t._y0+2*t._y1)/3,(t._x0+4*t._x1+e)/6,(t._y0+4*t._y1+r)/6)}function ct(t){this._context=t}function ut(t){this._context=t}function ht(t){this._context=t}function ft(t,e){this._basis=new ct(t),this._beta=e}ct.prototype={areaStart:function(){this._line=0},areaEnd:function(){this._line=NaN},lineStart:function(){this._x0=this._x1=this._y0=this._y1=NaN,this._point=0},lineEnd:function(){switch(this._point){case 3:lt(this,this._x1,this._y1);case 2:this._context.lineTo(this._x1,this._y1)}(this._line||0!==this._line&&1===this._point)&&this._context.closePath(),this._line=1-this._line},point:function(t,e){switch(t=+t,e=+e,this._point){case 0:this._point=1,this._line?this._context.lineTo(t,e):this._context.moveTo(t,e);break;case 1:this._point=2;break;case 2:this._point=3,this._context.lineTo((5*this._x0+this._x1)/6,(5*this._y0+this._y1)/6);default:lt(this,t,e)}this._x0=this._x1,this._x1=t,this._y0=this._y1,this._y1=e}},ut.prototype={areaStart:st,areaEnd:st,lineStart:function(){this._x0=this._x1=this._x2=this._x3=this._x4=this._y0=this._y1=this._y2=this._y3=this._y4=NaN,this._point=0},lineEnd:function(){switch(this._point){case 1:this._context.moveTo(this._x2,this._y2),this._context.closePath();break;case 2:this._context.moveTo((this._x2+2*this._x3)/3,(this._y2+2*this._y3)/3),this._context.lineTo((this._x3+2*this._x2)/3,(this._y3+2*this._y2)/3),this._context.closePath();break;case 3:this.point(this._x2,this._y2),this.point(this._x3,this._y3),this.point(this._x4,this._y4)}},point:function(t,e){switch(t=+t,e=+e,this._point){case 0:this._point=1,this._x2=t,this._y2=e;break;case 1:this._point=2,this._x3=t,this._y3=e;break;case 2:this._point=3,this._x4=t,this._y4=e,this._context.moveTo((this._x0+4*this._x1+t)/6,(this._y0+4*this._y1+e)/6);break;default:lt(this,t,e)}this._x0=this._x1,this._x1=t,this._y0=this._y1,this._y1=e}},ht.prototype={areaStart:function(){this._line=0},areaEnd:function(){this._line=NaN},lineStart:function(){this._x0=this._x1=this._y0=this._y1=NaN,this._point=0},lineEnd:function(){(this._line||0!==this._line&&3===this._point)&&this._context.closePath(),this._line=1-this._line},point:function(t,e){switch(t=+t,e=+e,this._point){case 0:this._point=1;break;case 1:this._point=2;break;case 2:this._point=3;var r=(this._x0+4*this._x1+t)/6,n=(this._y0+4*this._y1+e)/6;this._line?this._context.lineTo(r,n):this._context.moveTo(r,n);break;case 3:this._point=4;default:lt(this,t,e)}this._x0=this._x1,this._x1=t,this._y0=this._y1,this._y1=e}},ft.prototype={lineStart:function(){this._x=[],this._y=[],this._basis.lineStart()},lineEnd:function(){var t=this._x,e=this._y,r=t.length-1;if(r>0)for(var n,i=t[0],a=e[0],o=t[r]-i,s=e[r]-a,l=-1;++l<=r;)n=l/r,this._basis.point(this._beta*t[l]+(1-this._beta)*(i+n*o),this._beta*e[l]+(1-this._beta)*(a+n*s));this._x=this._y=null,this._basis.lineEnd()},point:function(t,e){this._x.push(+t),this._y.push(+e)}};var pt=function t(e){function r(t){return 1===e?new ct(t):new ft(t,e)}return r.beta=function(e){return t(+e)},r}(.85);function dt(t,e,r){t._context.bezierCurveTo(t._x1+t._k*(t._x2-t._x0),t._y1+t._k*(t._y2-t._y0),t._x2+t._k*(t._x1-e),t._y2+t._k*(t._y1-r),t._x2,t._y2)}function gt(t,e){this._context=t,this._k=(1-e)/6}gt.prototype={areaStart:function(){this._line=0},areaEnd:function(){this._line=NaN},lineStart:function(){this._x0=this._x1=this._x2=this._y0=this._y1=this._y2=NaN,this._point=0},lineEnd:function(){switch(this._point){case 2:this._context.lineTo(this._x2,this._y2);break;case 3:dt(this,this._x1,this._y1)}(this._line||0!==this._line&&1===this._point)&&this._context.closePath(),this._line=1-this._line},point:function(t,e){switch(t=+t,e=+e,this._point){case 0:this._point=1,this._line?this._context.lineTo(t,e):this._context.moveTo(t,e);break;case 1:this._point=2,this._x1=t,this._y1=e;break;case 2:this._point=3;default:dt(this,t,e)}this._x0=this._x1,this._x1=this._x2,this._x2=t,this._y0=this._y1,this._y1=this._y2,this._y2=e}};var vt=function t(e){function r(t){return new gt(t,e)}return r.tension=function(e){return t(+e)},r}(0);function mt(t,e){this._context=t,this._k=(1-e)/6}mt.prototype={areaStart:st,areaEnd:st,lineStart:function(){this._x0=this._x1=this._x2=this._x3=this._x4=this._x5=this._y0=this._y1=this._y2=this._y3=this._y4=this._y5=NaN,this._point=0},lineEnd:function(){switch(this._point){case 1:this._context.moveTo(this._x3,this._y3),this._context.closePath();break;case 2:this._context.lineTo(this._x3,this._y3),this._context.closePath();break;case 3:this.point(this._x3,this._y3),this.point(this._x4,this._y4),this.point(this._x5,this._y5)}},point:function(t,e){switch(t=+t,e=+e,this._point){case 0:this._point=1,this._x3=t,this._y3=e;break;case 1:this._point=2,this._context.moveTo(this._x4=t,this._y4=e);break;case 2:this._point=3,this._x5=t,this._y5=e;break;default:dt(this,t,e)}this._x0=this._x1,this._x1=this._x2,this._x2=t,this._y0=this._y1,this._y1=this._y2,this._y2=e}};var yt=function t(e){function r(t){return new mt(t,e)}return r.tension=function(e){return t(+e)},r}(0);function xt(t,e){this._context=t,this._k=(1-e)/6}xt.prototype={areaStart:function(){this._line=0},areaEnd:function(){this._line=NaN},lineStart:function(){this._x0=this._x1=this._x2=this._y0=this._y1=this._y2=NaN,this._point=0},lineEnd:function(){(this._line||0!==this._line&&3===this._point)&&this._context.closePath(),this._line=1-this._line},point:function(t,e){switch(t=+t,e=+e,this._point){case 0:this._point=1;break;case 1:this._point=2;break;case 2:this._point=3,this._line?this._context.lineTo(this._x2,this._y2):this._context.moveTo(this._x2,this._y2);break;case 3:this._point=4;default:dt(this,t,e)}this._x0=this._x1,this._x1=this._x2,this._x2=t,this._y0=this._y1,this._y1=this._y2,this._y2=e}};var bt=function t(e){function r(t){return new xt(t,e)}return r.tension=function(e){return t(+e)},r}(0);function _t(t,e,r){var n=t._x1,i=t._y1,a=t._x2,o=t._y2;if(t._l01_a>u){var s=2*t._l01_2a+3*t._l01_a*t._l12_a+t._l12_2a,l=3*t._l01_a*(t._l01_a+t._l12_a);n=(n*s-t._x0*t._l12_2a+t._x2*t._l01_2a)/l,i=(i*s-t._y0*t._l12_2a+t._y2*t._l01_2a)/l}if(t._l23_a>u){var c=2*t._l23_2a+3*t._l23_a*t._l12_a+t._l12_2a,h=3*t._l23_a*(t._l23_a+t._l12_a);a=(a*c+t._x1*t._l23_2a-e*t._l12_2a)/h,o=(o*c+t._y1*t._l23_2a-r*t._l12_2a)/h}t._context.bezierCurveTo(n,i,a,o,t._x2,t._y2)}function wt(t,e){this._context=t,this._alpha=e}wt.prototype={areaStart:function(){this._line=0},areaEnd:function(){this._line=NaN},lineStart:function(){this._x0=this._x1=this._x2=this._y0=this._y1=this._y2=NaN,this._l01_a=this._l12_a=this._l23_a=this._l01_2a=this._l12_2a=this._l23_2a=this._point=0},lineEnd:function(){switch(this._point){case 2:this._context.lineTo(this._x2,this._y2);break;case 3:this.point(this._x2,this._y2)}(this._line||0!==this._line&&1===this._point)&&this._context.closePath(),this._line=1-this._line},point:function(t,e){if(t=+t,e=+e,this._point){var r=this._x2-t,n=this._y2-e;this._l23_a=Math.sqrt(this._l23_2a=Math.pow(r*r+n*n,this._alpha))}switch(this._point){case 0:this._point=1,this._line?this._context.lineTo(t,e):this._context.moveTo(t,e);break;case 1:this._point=2;break;case 2:this._point=3;default:_t(this,t,e)}this._l01_a=this._l12_a,this._l12_a=this._l23_a,this._l01_2a=this._l12_2a,this._l12_2a=this._l23_2a,this._x0=this._x1,this._x1=this._x2,this._x2=t,this._y0=this._y1,this._y1=this._y2,this._y2=e}};var kt=function t(e){function r(t){return e?new wt(t,e):new gt(t,0)}return r.alpha=function(e){return t(+e)},r}(.5);function Tt(t,e){this._context=t,this._alpha=e}Tt.prototype={areaStart:st,areaEnd:st,lineStart:function(){this._x0=this._x1=this._x2=this._x3=this._x4=this._x5=this._y0=this._y1=this._y2=this._y3=this._y4=this._y5=NaN,this._l01_a=this._l12_a=this._l23_a=this._l01_2a=this._l12_2a=this._l23_2a=this._point=0},lineEnd:function(){switch(this._point){case 1:this._context.moveTo(this._x3,this._y3),this._context.closePath();break;case 2:this._context.lineTo(this._x3,this._y3),this._context.closePath();break;case 3:this.point(this._x3,this._y3),this.point(this._x4,this._y4),this.point(this._x5,this._y5)}},point:function(t,e){if(t=+t,e=+e,this._point){var r=this._x2-t,n=this._y2-e;this._l23_a=Math.sqrt(this._l23_2a=Math.pow(r*r+n*n,this._alpha))}switch(this._point){case 0:this._point=1,this._x3=t,this._y3=e;break;case 1:this._point=2,this._context.moveTo(this._x4=t,this._y4=e);break;case 2:this._point=3,this._x5=t,this._y5=e;break;default:_t(this,t,e)}this._l01_a=this._l12_a,this._l12_a=this._l23_a,this._l01_2a=this._l12_2a,this._l12_2a=this._l23_2a,this._x0=this._x1,this._x1=this._x2,this._x2=t,this._y0=this._y1,this._y1=this._y2,this._y2=e}};var At=function t(e){function r(t){return e?new Tt(t,e):new mt(t,0)}return r.alpha=function(e){return t(+e)},r}(.5);function Mt(t,e){this._context=t,this._alpha=e}Mt.prototype={areaStart:function(){this._line=0},areaEnd:function(){this._line=NaN},lineStart:function(){this._x0=this._x1=this._x2=this._y0=this._y1=this._y2=NaN,this._l01_a=this._l12_a=this._l23_a=this._l01_2a=this._l12_2a=this._l23_2a=this._point=0},lineEnd:function(){(this._line||0!==this._line&&3===this._point)&&this._context.closePath(),this._line=1-this._line},point:function(t,e){if(t=+t,e=+e,this._point){var r=this._x2-t,n=this._y2-e;this._l23_a=Math.sqrt(this._l23_2a=Math.pow(r*r+n*n,this._alpha))}switch(this._point){case 0:this._point=1;break;case 1:this._point=2;break;case 2:this._point=3,this._line?this._context.lineTo(this._x2,this._y2):this._context.moveTo(this._x2,this._y2);break;case 3:this._point=4;default:_t(this,t,e)}this._l01_a=this._l12_a,this._l12_a=this._l23_a,this._l01_2a=this._l12_2a,this._l12_2a=this._l23_2a,this._x0=this._x1,this._x1=this._x2,this._x2=t,this._y0=this._y1,this._y1=this._y2,this._y2=e}};var St=function t(e){function r(t){return e?new Mt(t,e):new xt(t,0)}return r.alpha=function(e){return t(+e)},r}(.5);function Et(t){this._context=t}function Ct(t){return t<0?-1:1}function Lt(t,e,r){var n=t._x1-t._x0,i=e-t._x1,a=(t._y1-t._y0)/(n||i<0&&-0),o=(r-t._y1)/(i||n<0&&-0),s=(a*i+o*n)/(n+i);return(Ct(a)+Ct(o))*Math.min(Math.abs(a),Math.abs(o),.5*Math.abs(s))||0}function Pt(t,e){var r=t._x1-t._x0;return r?(3*(t._y1-t._y0)/r-e)/2:e}function Ot(t,e,r){var n=t._x0,i=t._y0,a=t._x1,o=t._y1,s=(a-n)/3;t._context.bezierCurveTo(n+s,i+s*e,a-s,o-s*r,a,o)}function zt(t){this._context=t}function It(t){this._context=new Dt(t)}function Dt(t){this._context=t}function Rt(t){this._context=t}function Ft(t){var e,r,n=t.length-1,i=new Array(n),a=new Array(n),o=new Array(n);for(i[0]=0,a[0]=2,o[0]=t[0]+2*t[1],e=1;e<n-1;++e)i[e]=1,a[e]=4,o[e]=4*t[e]+2*t[e+1];for(i[n-1]=2,a[n-1]=7,o[n-1]=8*t[n-1]+t[n],e=1;e<n;++e)r=i[e]/a[e-1],a[e]-=r,o[e]-=r*o[e-1];for(i[n-1]=o[n-1]/a[n-1],e=n-2;e>=0;--e)i[e]=(o[e]-i[e+1])/a[e];for(a[n-1]=(t[n]+i[n-1])/2,e=0;e<n-1;++e)a[e]=2*t[e+1]-i[e+1];return[i,a]}function Bt(t,e){this._context=t,this._t=e}function Nt(t,e){if((i=t.length)>1)for(var r,n,i,a=1,o=t[e[0]],s=o.length;a<i;++a)for(n=o,o=t[e[a]],r=0;r<s;++r)o[r][1]+=o[r][0]=isNaN(n[r][1])?n[r][0]:n[r][1]}function jt(t){for(var e=t.length,r=new Array(e);--e>=0;)r[e]=e;return r}function Ut(t,e){return t[e]}function Vt(t){var e=t.map(qt);return jt(t).sort(function(t,r){return e[t]-e[r]})}function qt(t){for(var e,r=-1,n=0,i=t.length,a=-1/0;++r<i;)(e=+t[r][1])>a&&(a=e,n=r);return n}function Ht(t){var e=t.map(Gt);return jt(t).sort(function(t,r){return e[t]-e[r]})}function Gt(t){for(var e,r=0,n=-1,i=t.length;++n<i;)(e=+t[n][1])&&(r+=e);return r}Et.prototype={areaStart:st,areaEnd:st,lineStart:function(){this._point=0},lineEnd:function(){this._point&&this._context.closePath()},point:function(t,e){t=+t,e=+e,this._point?this._context.lineTo(t,e):(this._point=1,this._context.moveTo(t,e))}},zt.prototype={areaStart:function(){this._line=0},areaEnd:function(){this._line=NaN},lineStart:function(){this._x0=this._x1=this._y0=this._y1=this._t0=NaN,this._point=0},lineEnd:function(){switch(this._point){case 2:this._context.lineTo(this._x1,this._y1);break;case 3:Ot(this,this._t0,Pt(this,this._t0))}(this._line||0!==this._line&&1===this._point)&&this._context.closePath(),this._line=1-this._line},point:function(t,e){var r=NaN;if(e=+e,(t=+t)!==this._x1||e!==this._y1){switch(this._point){case 0:this._point=1,this._line?this._context.lineTo(t,e):this._context.moveTo(t,e);break;case 1:this._point=2;break;case 2:this._point=3,Ot(this,Pt(this,r=Lt(this,t,e)),r);break;default:Ot(this,this._t0,r=Lt(this,t,e))}this._x0=this._x1,this._x1=t,this._y0=this._y1,this._y1=e,this._t0=r}}},(It.prototype=Object.create(zt.prototype)).point=function(t,e){zt.prototype.point.call(this,e,t)},Dt.prototype={moveTo:function(t,e){this._context.moveTo(e,t)},closePath:function(){this._context.closePath()},lineTo:function(t,e){this._context.lineTo(e,t)},bezierCurveTo:function(t,e,r,n,i,a){this._context.bezierCurveTo(e,t,n,r,a,i)}},Rt.prototype={areaStart:function(){this._line=0},areaEnd:function(){this._line=NaN},lineStart:function(){this._x=[],this._y=[]},lineEnd:function(){var t=this._x,e=this._y,r=t.length;if(r)if(this._line?this._context.lineTo(t[0],e[0]):this._context.moveTo(t[0],e[0]),2===r)this._context.lineTo(t[1],e[1]);else for(var n=Ft(t),i=Ft(e),a=0,o=1;o<r;++a,++o)this._context.bezierCurveTo(n[0][a],i[0][a],n[1][a],i[1][a],t[o],e[o]);(this._line||0!==this._line&&1===r)&&this._context.closePath(),this._line=1-this._line,this._x=this._y=null},point:function(t,e){this._x.push(+t),this._y.push(+e)}},Bt.prototype={areaStart:function(){this._line=0},areaEnd:function(){this._line=NaN},lineStart:function(){this._x=this._y=NaN,this._point=0},lineEnd:function(){0<this._t&&this._t<1&&2===this._point&&this._context.lineTo(this._x,this._y),(this._line||0!==this._line&&1===this._point)&&this._context.closePath(),this._line>=0&&(this._t=1-this._t,this._line=1-this._line)},point:function(t,e){switch(t=+t,e=+e,this._point){case 0:this._point=1,this._line?this._context.lineTo(t,e):this._context.moveTo(t,e);break;case 1:this._point=2;default:if(this._t<=0)this._context.lineTo(this._x,e),this._context.lineTo(t,e);else{var r=this._x*(1-this._t)+t*this._t;this._context.lineTo(r,this._y),this._context.lineTo(r,e)}}this._x=t,this._y=e}},t.arc=function(){var t=g,o=v,_=r(0),w=null,k=m,T=y,A=x,M=null;function S(){var r,g,v,m=+t.apply(this,arguments),y=+o.apply(this,arguments),x=k.apply(this,arguments)-f,S=T.apply(this,arguments)-f,E=n(S-x),C=S>x;if(M||(M=r=e.path()),y<m&&(g=y,y=m,m=g),y>u)if(E>p-u)M.moveTo(y*a(x),y*l(x)),M.arc(0,0,y,x,S,!C),m>u&&(M.moveTo(m*a(S),m*l(S)),M.arc(0,0,m,S,x,C));else{var L,P,O=x,z=S,I=x,D=S,R=E,F=E,B=A.apply(this,arguments)/2,N=B>u&&(w?+w.apply(this,arguments):c(m*m+y*y)),j=s(n(y-m)/2,+_.apply(this,arguments)),U=j,V=j;if(N>u){var q=d(N/m*l(B)),H=d(N/y*l(B));(R-=2*q)>u?(I+=q*=C?1:-1,D-=q):(R=0,I=D=(x+S)/2),(F-=2*H)>u?(O+=H*=C?1:-1,z-=H):(F=0,O=z=(x+S)/2)}var G=y*a(O),Y=y*l(O),W=m*a(D),X=m*l(D);if(j>u){var Z,J=y*a(z),K=y*l(z),$=m*a(I),Q=m*l(I);if(E<h&&(Z=function(t,e,r,n,i,a,o,s){var l=r-t,c=n-e,h=o-i,f=s-a,p=f*l-h*c;if(!(p*p<u))return[t+(p=(h*(e-a)-f*(t-i))/p)*l,e+p*c]}(G,Y,$,Q,J,K,W,X))){var tt=G-Z[0],et=Y-Z[1],rt=J-Z[0],nt=K-Z[1],it=1/l(((v=(tt*rt+et*nt)/(c(tt*tt+et*et)*c(rt*rt+nt*nt)))>1?0:v<-1?h:Math.acos(v))/2),at=c(Z[0]*Z[0]+Z[1]*Z[1]);U=s(j,(m-at)/(it-1)),V=s(j,(y-at)/(it+1))}}F>u?V>u?(L=b($,Q,G,Y,y,V,C),P=b(J,K,W,X,y,V,C),M.moveTo(L.cx+L.x01,L.cy+L.y01),V<j?M.arc(L.cx,L.cy,V,i(L.y01,L.x01),i(P.y01,P.x01),!C):(M.arc(L.cx,L.cy,V,i(L.y01,L.x01),i(L.y11,L.x11),!C),M.arc(0,0,y,i(L.cy+L.y11,L.cx+L.x11),i(P.cy+P.y11,P.cx+P.x11),!C),M.arc(P.cx,P.cy,V,i(P.y11,P.x11),i(P.y01,P.x01),!C))):(M.moveTo(G,Y),M.arc(0,0,y,O,z,!C)):M.moveTo(G,Y),m>u&&R>u?U>u?(L=b(W,X,J,K,m,-U,C),P=b(G,Y,$,Q,m,-U,C),M.lineTo(L.cx+L.x01,L.cy+L.y01),U<j?M.arc(L.cx,L.cy,U,i(L.y01,L.x01),i(P.y01,P.x01),!C):(M.arc(L.cx,L.cy,U,i(L.y01,L.x01),i(L.y11,L.x11),!C),M.arc(0,0,m,i(L.cy+L.y11,L.cx+L.x11),i(P.cy+P.y11,P.cx+P.x11),C),M.arc(P.cx,P.cy,U,i(P.y11,P.x11),i(P.y01,P.x01),!C))):M.arc(0,0,m,D,I,C):M.lineTo(W,X)}else M.moveTo(0,0);if(M.closePath(),r)return M=null,r+""||null}return S.centroid=function(){var e=(+t.apply(this,arguments)+ +o.apply(this,arguments))/2,r=(+k.apply(this,arguments)+ +T.apply(this,arguments))/2-h/2;return[a(r)*e,l(r)*e]},S.innerRadius=function(e){return arguments.length?(t="function"==typeof e?e:r(+e),S):t},S.outerRadius=function(t){return arguments.length?(o="function"==typeof t?t:r(+t),S):o},S.cornerRadius=function(t){return arguments.length?(_="function"==typeof t?t:r(+t),S):_},S.padRadius=function(t){return arguments.length?(w=null==t?null:"function"==typeof t?t:r(+t),S):w},S.startAngle=function(t){return arguments.length?(k="function"==typeof t?t:r(+t),S):k},S.endAngle=function(t){return arguments.length?(T="function"==typeof t?t:r(+t),S):T},S.padAngle=function(t){return arguments.length?(A="function"==typeof t?t:r(+t),S):A},S.context=function(t){return arguments.length?(M=null==t?null:t,S):M},S},t.area=M,t.line=A,t.pie=function(){var t=E,e=S,n=null,i=r(0),a=r(p),o=r(0);function s(r){var s,l,c,u,h,f=r.length,d=0,g=new Array(f),v=new Array(f),m=+i.apply(this,arguments),y=Math.min(p,Math.max(-p,a.apply(this,arguments)-m)),x=Math.min(Math.abs(y)/f,o.apply(this,arguments)),b=x*(y<0?-1:1);for(s=0;s<f;++s)(h=v[g[s]=s]=+t(r[s],s,r))>0&&(d+=h);for(null!=e?g.sort(function(t,r){return e(v[t],v[r])}):null!=n&&g.sort(function(t,e){return n(r[t],r[e])}),s=0,c=d?(y-f*b)/d:0;s<f;++s,m=u)l=g[s],u=m+((h=v[l])>0?h*c:0)+b,v[l]={data:r[l],index:s,value:h,startAngle:m,endAngle:u,padAngle:x};return v}return s.value=function(e){return arguments.length?(t="function"==typeof e?e:r(+e),s):t},s.sortValues=function(t){return arguments.length?(e=t,n=null,s):e},s.sort=function(t){return arguments.length?(n=t,e=null,s):n},s.startAngle=function(t){return arguments.length?(i="function"==typeof t?t:r(+t),s):i},s.endAngle=function(t){return arguments.length?(a="function"==typeof t?t:r(+t),s):a},s.padAngle=function(t){return arguments.length?(o="function"==typeof t?t:r(+t),s):o},s},t.areaRadial=I,t.radialArea=I,t.lineRadial=z,t.radialLine=z,t.pointRadial=D,t.linkHorizontal=function(){return N(j)},t.linkVertical=function(){return N(U)},t.linkRadial=function(){var t=N(V);return t.angle=t.x,delete t.x,t.radius=t.y,delete t.y,t},t.symbol=function(){var t=r(q),n=r(64),i=null;function a(){var r;if(i||(i=r=e.path()),t.apply(this,arguments).draw(i,+n.apply(this,arguments)),r)return i=null,r+""||null}return a.type=function(e){return arguments.length?(t="function"==typeof e?e:r(e),a):t},a.size=function(t){return arguments.length?(n="function"==typeof t?t:r(+t),a):n},a.context=function(t){return arguments.length?(i=null==t?null:t,a):i},a},t.symbols=ot,t.symbolCircle=q,t.symbolCross=H,t.symbolDiamond=W,t.symbolSquare=$,t.symbolStar=K,t.symbolTriangle=tt,t.symbolWye=at,t.curveBasisClosed=function(t){return new ut(t)},t.curveBasisOpen=function(t){return new ht(t)},t.curveBasis=function(t){return new ct(t)},t.curveBundle=pt,t.curveCardinalClosed=yt,t.curveCardinalOpen=bt,t.curveCardinal=vt,t.curveCatmullRomClosed=At,t.curveCatmullRomOpen=St,t.curveCatmullRom=kt,t.curveLinearClosed=function(t){return new Et(t)},t.curveLinear=w,t.curveMonotoneX=function(t){return new zt(t)},t.curveMonotoneY=function(t){return new It(t)},t.curveNatural=function(t){return new Rt(t)},t.curveStep=function(t){return new Bt(t,.5)},t.curveStepAfter=function(t){return new Bt(t,1)},t.curveStepBefore=function(t){return new Bt(t,0)},t.stack=function(){var t=r([]),e=jt,n=Nt,i=Ut;function a(r){var a,o,s=t.apply(this,arguments),l=r.length,c=s.length,u=new Array(c);for(a=0;a<c;++a){for(var h,f=s[a],p=u[a]=new Array(l),d=0;d<l;++d)p[d]=h=[0,+i(r[d],f,d,r)],h.data=r[d];p.key=f}for(a=0,o=e(u);a<c;++a)u[o[a]].index=a;return n(u,o),u}return a.keys=function(e){return arguments.length?(t="function"==typeof e?e:r(R.call(e)),a):t},a.value=function(t){return arguments.length?(i="function"==typeof t?t:r(+t),a):i},a.order=function(t){return arguments.length?(e=null==t?jt:"function"==typeof t?t:r(R.call(t)),a):e},a.offset=function(t){return arguments.length?(n=null==t?Nt:t,a):n},a},t.stackOffsetExpand=function(t,e){if((n=t.length)>0){for(var r,n,i,a=0,o=t[0].length;a<o;++a){for(i=r=0;r<n;++r)i+=t[r][a][1]||0;if(i)for(r=0;r<n;++r)t[r][a][1]/=i}Nt(t,e)}},t.stackOffsetDiverging=function(t,e){if((s=t.length)>1)for(var r,n,i,a,o,s,l=0,c=t[e[0]].length;l<c;++l)for(a=o=0,r=0;r<s;++r)(i=(n=t[e[r]][l])[1]-n[0])>=0?(n[0]=a,n[1]=a+=i):i<0?(n[1]=o,n[0]=o+=i):n[0]=a},t.stackOffsetNone=Nt,t.stackOffsetSilhouette=function(t,e){if((r=t.length)>0){for(var r,n=0,i=t[e[0]],a=i.length;n<a;++n){for(var o=0,s=0;o<r;++o)s+=t[o][n][1]||0;i[n][1]+=i[n][0]=-s/2}Nt(t,e)}},t.stackOffsetWiggle=function(t,e){if((i=t.length)>0&&(n=(r=t[e[0]]).length)>0){for(var r,n,i,a=0,o=1;o<n;++o){for(var s=0,l=0,c=0;s<i;++s){for(var u=t[e[s]],h=u[o][1]||0,f=(h-(u[o-1][1]||0))/2,p=0;p<s;++p){var d=t[e[p]];f+=(d[o][1]||0)-(d[o-1][1]||0)}l+=h,c+=f*h}r[o-1][1]+=r[o-1][0]=a,l&&(a-=c/l)}r[o-1][1]+=r[o-1][0]=a,Nt(t,e)}},t.stackOrderAppearance=Vt,t.stackOrderAscending=Ht,t.stackOrderDescending=function(t){return Ht(t).reverse()},t.stackOrderInsideOut=function(t){var e,r,n=t.length,i=t.map(Gt),a=Vt(t),o=0,s=0,l=[],c=[];for(e=0;e<n;++e)r=a[e],o<s?(o+=i[r],l.push(r)):(s+=i[r],c.push(r));return c.reverse().concat(l)},t.stackOrderNone=jt,t.stackOrderReverse=function(t){return jt(t).reverse()},Object.defineProperty(t,"__esModule",{value:!0})},"object"==typeof r&&"undefined"!=typeof e?i(r,t("d3-path")):i(n.d3=n.d3||{},n.d3)},{"d3-path":159}],162:[function(t,e,r){var n;n=this,function(t){"use strict";var e,r,n=0,i=0,a=0,o=1e3,s=0,l=0,c=0,u="object"==typeof performance&&performance.now?performance:Date,h="object"==typeof window&&window.requestAnimationFrame?window.requestAnimationFrame.bind(window):function(t){setTimeout(t,17)};function f(){return l||(h(p),l=u.now()+c)}function p(){l=0}function d(){this._call=this._time=this._next=null}function g(t,e,r){var n=new d;return n.restart(t,e,r),n}function v(){f(),++n;for(var t,r=e;r;)(t=l-r._time)>=0&&r._call.call(null,t),r=r._next;--n}function m(){l=(s=u.now())+c,n=i=0;try{v()}finally{n=0,function(){var t,n,i=e,a=1/0;for(;i;)i._call?(a>i._time&&(a=i._time),t=i,i=i._next):(n=i._next,i._next=null,i=t?t._next=n:e=n);r=t,x(a)}(),l=0}}function y(){var t=u.now(),e=t-s;e>o&&(c-=e,s=t)}function x(t){n||(i&&(i=clearTimeout(i)),t-l>24?(t<1/0&&(i=setTimeout(m,t-u.now()-c)),a&&(a=clearInterval(a))):(a||(s=u.now(),a=setInterval(y,o)),n=1,h(m)))}d.prototype=g.prototype={constructor:d,restart:function(t,n,i){if("function"!=typeof t)throw new TypeError("callback is not a function");i=(null==i?f():+i)+(null==n?0:+n),this._next||r===this||(r?r._next=this:e=this,r=this),this._call=t,this._time=i,x()},stop:function(){this._call&&(this._call=null,this._time=1/0,x())}};t.now=f,t.timer=g,t.timerFlush=v,t.timeout=function(t,e,r){var n=new d;return e=null==e?0:+e,n.restart(function(r){n.stop(),t(r+e)},e,r),n},t.interval=function(t,e,r){var n=new d,i=e;return null==e?(n.restart(t,e,r),n):(e=+e,r=null==r?f():+r,n.restart(function a(o){o+=i,n.restart(a,i+=e,r),t(o)},e,r),n)},Object.defineProperty(t,"__esModule",{value:!0})}("object"==typeof r&&"undefined"!=typeof e?r:n.d3=n.d3||{})},{}],163:[function(t,e,r){!function(){var t={version:"3.5.17"},r=[].slice,n=function(t){return r.call(t)},i=this.document;function a(t){return t&&(t.ownerDocument||t.document||t).documentElement}function o(t){return t&&(t.ownerDocument&&t.ownerDocument.defaultView||t.document&&t||t.defaultView)}if(i)try{n(i.documentElement.childNodes)[0].nodeType}catch(t){n=function(t){for(var e=t.length,r=new Array(e);e--;)r[e]=t[e];return r}}if(Date.now||(Date.now=function(){return+new Date}),i)try{i.createElement("DIV").style.setProperty("opacity",0,"")}catch(t){var s=this.Element.prototype,l=s.setAttribute,c=s.setAttributeNS,u=this.CSSStyleDeclaration.prototype,h=u.setProperty;s.setAttribute=function(t,e){l.call(this,t,e+"")},s.setAttributeNS=function(t,e,r){c.call(this,t,e,r+"")},u.setProperty=function(t,e,r){h.call(this,t,e+"",r)}}function f(t,e){return t<e?-1:t>e?1:t>=e?0:NaN}function p(t){return null===t?NaN:+t}function d(t){return!isNaN(t)}function g(t){return{left:function(e,r,n,i){for(arguments.length<3&&(n=0),arguments.length<4&&(i=e.length);n<i;){var a=n+i>>>1;t(e[a],r)<0?n=a+1:i=a}return n},right:function(e,r,n,i){for(arguments.length<3&&(n=0),arguments.length<4&&(i=e.length);n<i;){var a=n+i>>>1;t(e[a],r)>0?i=a:n=a+1}return n}}}t.ascending=f,t.descending=function(t,e){return e<t?-1:e>t?1:e>=t?0:NaN},t.min=function(t,e){var r,n,i=-1,a=t.length;if(1===arguments.length){for(;++i<a;)if(null!=(n=t[i])&&n>=n){r=n;break}for(;++i<a;)null!=(n=t[i])&&r>n&&(r=n)}else{for(;++i<a;)if(null!=(n=e.call(t,t[i],i))&&n>=n){r=n;break}for(;++i<a;)null!=(n=e.call(t,t[i],i))&&r>n&&(r=n)}return r},t.max=function(t,e){var r,n,i=-1,a=t.length;if(1===arguments.length){for(;++i<a;)if(null!=(n=t[i])&&n>=n){r=n;break}for(;++i<a;)null!=(n=t[i])&&n>r&&(r=n)}else{for(;++i<a;)if(null!=(n=e.call(t,t[i],i))&&n>=n){r=n;break}for(;++i<a;)null!=(n=e.call(t,t[i],i))&&n>r&&(r=n)}return r},t.extent=function(t,e){var r,n,i,a=-1,o=t.length;if(1===arguments.length){for(;++a<o;)if(null!=(n=t[a])&&n>=n){r=i=n;break}for(;++a<o;)null!=(n=t[a])&&(r>n&&(r=n),i<n&&(i=n))}else{for(;++a<o;)if(null!=(n=e.call(t,t[a],a))&&n>=n){r=i=n;break}for(;++a<o;)null!=(n=e.call(t,t[a],a))&&(r>n&&(r=n),i<n&&(i=n))}return[r,i]},t.sum=function(t,e){var r,n=0,i=t.length,a=-1;if(1===arguments.length)for(;++a<i;)d(r=+t[a])&&(n+=r);else for(;++a<i;)d(r=+e.call(t,t[a],a))&&(n+=r);return n},t.mean=function(t,e){var r,n=0,i=t.length,a=-1,o=i;if(1===arguments.length)for(;++a<i;)d(r=p(t[a]))?n+=r:--o;else for(;++a<i;)d(r=p(e.call(t,t[a],a)))?n+=r:--o;if(o)return n/o},t.quantile=function(t,e){var r=(t.length-1)*e+1,n=Math.floor(r),i=+t[n-1],a=r-n;return a?i+a*(t[n]-i):i},t.median=function(e,r){var n,i=[],a=e.length,o=-1;if(1===arguments.length)for(;++o<a;)d(n=p(e[o]))&&i.push(n);else for(;++o<a;)d(n=p(r.call(e,e[o],o)))&&i.push(n);if(i.length)return t.quantile(i.sort(f),.5)},t.variance=function(t,e){var r,n,i=t.length,a=0,o=0,s=-1,l=0;if(1===arguments.length)for(;++s<i;)d(r=p(t[s]))&&(o+=(n=r-a)*(r-(a+=n/++l)));else for(;++s<i;)d(r=p(e.call(t,t[s],s)))&&(o+=(n=r-a)*(r-(a+=n/++l)));if(l>1)return o/(l-1)},t.deviation=function(){var e=t.variance.apply(this,arguments);return e?Math.sqrt(e):e};var v=g(f);function m(t){return t.length}t.bisectLeft=v.left,t.bisect=t.bisectRight=v.right,t.bisector=function(t){return g(1===t.length?function(e,r){return f(t(e),r)}:t)},t.shuffle=function(t,e,r){(a=arguments.length)<3&&(r=t.length,a<2&&(e=0));for(var n,i,a=r-e;a;)i=Math.random()*a--|0,n=t[a+e],t[a+e]=t[i+e],t[i+e]=n;return t},t.permute=function(t,e){for(var r=e.length,n=new Array(r);r--;)n[r]=t[e[r]];return n},t.pairs=function(t){for(var e=0,r=t.length-1,n=t[0],i=new Array(r<0?0:r);e<r;)i[e]=[n,n=t[++e]];return i},t.transpose=function(e){if(!(a=e.length))return[];for(var r=-1,n=t.min(e,m),i=new Array(n);++r<n;)for(var a,o=-1,s=i[r]=new Array(a);++o<a;)s[o]=e[o][r];return i},t.zip=function(){return t.transpose(arguments)},t.keys=function(t){var e=[];for(var r in t)e.push(r);return e},t.values=function(t){var e=[];for(var r in t)e.push(t[r]);return e},t.entries=function(t){var e=[];for(var r in t)e.push({key:r,value:t[r]});return e},t.merge=function(t){for(var e,r,n,i=t.length,a=-1,o=0;++a<i;)o+=t[a].length;for(r=new Array(o);--i>=0;)for(e=(n=t[i]).length;--e>=0;)r[--o]=n[e];return r};var y=Math.abs;function x(t,e){for(var r in e)Object.defineProperty(t.prototype,r,{value:e[r],enumerable:!1})}function b(){this._=Object.create(null)}t.range=function(t,e,r){if(arguments.length<3&&(r=1,arguments.length<2&&(e=t,t=0)),(e-t)/r==1/0)throw new Error("infinite range");var n,i=[],a=function(t){var e=1;for(;t*e%1;)e*=10;return e}(y(r)),o=-1;if(t*=a,e*=a,(r*=a)<0)for(;(n=t+r*++o)>e;)i.push(n/a);else for(;(n=t+r*++o)<e;)i.push(n/a);return i},t.map=function(t,e){var r=new b;if(t instanceof b)t.forEach(function(t,e){r.set(t,e)});else if(Array.isArray(t)){var n,i=-1,a=t.length;if(1===arguments.length)for(;++i<a;)r.set(i,t[i]);else for(;++i<a;)r.set(e.call(t,n=t[i],i),n)}else for(var o in t)r.set(o,t[o]);return r};var _="__proto__",w="\0";function k(t){return(t+="")===_||t[0]===w?w+t:t}function T(t){return(t+="")[0]===w?t.slice(1):t}function A(t){return k(t)in this._}function M(t){return(t=k(t))in this._&&delete this._[t]}function S(){var t=[];for(var e in this._)t.push(T(e));return t}function E(){var t=0;for(var e in this._)++t;return t}function C(){for(var t in this._)return!1;return!0}function L(){this._=Object.create(null)}function P(t){return t}function O(t,e,r){return function(){var n=r.apply(e,arguments);return n===e?t:n}}function z(t,e){if(e in t)return e;e=e.charAt(0).toUpperCase()+e.slice(1);for(var r=0,n=I.length;r<n;++r){var i=I[r]+e;if(i in t)return i}}x(b,{has:A,get:function(t){return this._[k(t)]},set:function(t,e){return this._[k(t)]=e},remove:M,keys:S,values:function(){var t=[];for(var e in this._)t.push(this._[e]);return t},entries:function(){var t=[];for(var e in this._)t.push({key:T(e),value:this._[e]});return t},size:E,empty:C,forEach:function(t){for(var e in this._)t.call(this,T(e),this._[e])}}),t.nest=function(){var e,r,n={},i=[],a=[];function o(t,a,s){if(s>=i.length)return r?r.call(n,a):e?a.sort(e):a;for(var l,c,u,h,f=-1,p=a.length,d=i[s++],g=new b;++f<p;)(h=g.get(l=d(c=a[f])))?h.push(c):g.set(l,[c]);return t?(c=t(),u=function(e,r){c.set(e,o(t,r,s))}):(c={},u=function(e,r){c[e]=o(t,r,s)}),g.forEach(u),c}return n.map=function(t,e){return o(e,t,0)},n.entries=function(e){return function t(e,r){if(r>=i.length)return e;var n=[],o=a[r++];return e.forEach(function(e,i){n.push({key:e,values:t(i,r)})}),o?n.sort(function(t,e){return o(t.key,e.key)}):n}(o(t.map,e,0),0)},n.key=function(t){return i.push(t),n},n.sortKeys=function(t){return a[i.length-1]=t,n},n.sortValues=function(t){return e=t,n},n.rollup=function(t){return r=t,n},n},t.set=function(t){var e=new L;if(t)for(var r=0,n=t.length;r<n;++r)e.add(t[r]);return e},x(L,{has:A,add:function(t){return this._[k(t+="")]=!0,t},remove:M,values:S,size:E,empty:C,forEach:function(t){for(var e in this._)t.call(this,T(e))}}),t.behavior={},t.rebind=function(t,e){for(var r,n=1,i=arguments.length;++n<i;)t[r=arguments[n]]=O(t,e,e[r]);return t};var I=["webkit","ms","moz","Moz","o","O"];function D(){}function R(){}function F(t){var e=[],r=new b;function n(){for(var r,n=e,i=-1,a=n.length;++i<a;)(r=n[i].on)&&r.apply(this,arguments);return t}return n.on=function(n,i){var a,o=r.get(n);return arguments.length<2?o&&o.on:(o&&(o.on=null,e=e.slice(0,a=e.indexOf(o)).concat(e.slice(a+1)),r.remove(n)),i&&e.push(r.set(n,{on:i})),t)},n}function B(){t.event.preventDefault()}function N(){for(var e,r=t.event;e=r.sourceEvent;)r=e;return r}function j(e){for(var r=new R,n=0,i=arguments.length;++n<i;)r[arguments[n]]=F(r);return r.of=function(n,i){return function(a){try{var o=a.sourceEvent=t.event;a.target=e,t.event=a,r[a.type].apply(n,i)}finally{t.event=o}}},r}t.dispatch=function(){for(var t=new R,e=-1,r=arguments.length;++e<r;)t[arguments[e]]=F(t);return t},R.prototype.on=function(t,e){var r=t.indexOf("."),n="";if(r>=0&&(n=t.slice(r+1),t=t.slice(0,r)),t)return arguments.length<2?this[t].on(n):this[t].on(n,e);if(2===arguments.length){if(null==e)for(t in this)this.hasOwnProperty(t)&&this[t].on(n,null);return this}},t.event=null,t.requote=function(t){return t.replace(U,"\\$&")};var U=/[\\\^\$\*\+\?\|\[\]\(\)\.\{\}]/g,V={}.__proto__?function(t,e){t.__proto__=e}:function(t,e){for(var r in e)t[r]=e[r]};function q(t){return V(t,W),t}var H=function(t,e){return e.querySelector(t)},G=function(t,e){return e.querySelectorAll(t)},Y=function(t,e){var r=t.matches||t[z(t,"matchesSelector")];return(Y=function(t,e){return r.call(t,e)})(t,e)};"function"==typeof Sizzle&&(H=function(t,e){return Sizzle(t,e)[0]||null},G=Sizzle,Y=Sizzle.matchesSelector),t.selection=function(){return t.select(i.documentElement)};var W=t.selection.prototype=[];function X(t){return"function"==typeof t?t:function(){return H(t,this)}}function Z(t){return"function"==typeof t?t:function(){return G(t,this)}}W.select=function(t){var e,r,n,i,a=[];t=X(t);for(var o=-1,s=this.length;++o<s;){a.push(e=[]),e.parentNode=(n=this[o]).parentNode;for(var l=-1,c=n.length;++l<c;)(i=n[l])?(e.push(r=t.call(i,i.__data__,l,o)),r&&"__data__"in i&&(r.__data__=i.__data__)):e.push(null)}return q(a)},W.selectAll=function(t){var e,r,i=[];t=Z(t);for(var a=-1,o=this.length;++a<o;)for(var s=this[a],l=-1,c=s.length;++l<c;)(r=s[l])&&(i.push(e=n(t.call(r,r.__data__,l,a))),e.parentNode=r);return q(i)};var J="http://www.w3.org/1999/xhtml",K={svg:"http://www.w3.org/2000/svg",xhtml:J,xlink:"http://www.w3.org/1999/xlink",xml:"http://www.w3.org/XML/1998/namespace",xmlns:"http://www.w3.org/2000/xmlns/"};function $(e,r){return e=t.ns.qualify(e),null==r?e.local?function(){this.removeAttributeNS(e.space,e.local)}:function(){this.removeAttribute(e)}:"function"==typeof r?e.local?function(){var t=r.apply(this,arguments);null==t?this.removeAttributeNS(e.space,e.local):this.setAttributeNS(e.space,e.local,t)}:function(){var t=r.apply(this,arguments);null==t?this.removeAttribute(e):this.setAttribute(e,t)}:e.local?function(){this.setAttributeNS(e.space,e.local,r)}:function(){this.setAttribute(e,r)}}function Q(t){return t.trim().replace(/\s+/g," ")}function tt(e){return new RegExp("(?:^|\\s+)"+t.requote(e)+"(?:\\s+|$)","g")}function et(t){return(t+"").trim().split(/^|\s+/)}function rt(t,e){var r=(t=et(t).map(nt)).length;return"function"==typeof e?function(){for(var n=-1,i=e.apply(this,arguments);++n<r;)t[n](this,i)}:function(){for(var n=-1;++n<r;)t[n](this,e)}}function nt(t){var e=tt(t);return function(r,n){if(i=r.classList)return n?i.add(t):i.remove(t);var i=r.getAttribute("class")||"";n?(e.lastIndex=0,e.test(i)||r.setAttribute("class",Q(i+" "+t))):r.setAttribute("class",Q(i.replace(e," ")))}}function it(t,e,r){return null==e?function(){this.style.removeProperty(t)}:"function"==typeof e?function(){var n=e.apply(this,arguments);null==n?this.style.removeProperty(t):this.style.setProperty(t,n,r)}:function(){this.style.setProperty(t,e,r)}}function at(t,e){return null==e?function(){delete this[t]}:"function"==typeof e?function(){var r=e.apply(this,arguments);null==r?delete this[t]:this[t]=r}:function(){this[t]=e}}function ot(e){return"function"==typeof e?e:(e=t.ns.qualify(e)).local?function(){return this.ownerDocument.createElementNS(e.space,e.local)}:function(){var t=this.ownerDocument,r=this.namespaceURI;return r===J&&t.documentElement.namespaceURI===J?t.createElement(e):t.createElementNS(r,e)}}function st(){var t=this.parentNode;t&&t.removeChild(this)}function lt(t){return{__data__:t}}function ct(t){return function(){return Y(this,t)}}function ut(t,e){for(var r=0,n=t.length;r<n;r++)for(var i,a=t[r],o=0,s=a.length;o<s;o++)(i=a[o])&&e(i,o,r);return t}function ht(t){return V(t,ft),t}t.ns={prefix:K,qualify:function(t){var e=t.indexOf(":"),r=t;return e>=0&&"xmlns"!==(r=t.slice(0,e))&&(t=t.slice(e+1)),K.hasOwnProperty(r)?{space:K[r],local:t}:t}},W.attr=function(e,r){if(arguments.length<2){if("string"==typeof e){var n=this.node();return(e=t.ns.qualify(e)).local?n.getAttributeNS(e.space,e.local):n.getAttribute(e)}for(r in e)this.each($(r,e[r]));return this}return this.each($(e,r))},W.classed=function(t,e){if(arguments.length<2){if("string"==typeof t){var r=this.node(),n=(t=et(t)).length,i=-1;if(e=r.classList){for(;++i<n;)if(!e.contains(t[i]))return!1}else for(e=r.getAttribute("class");++i<n;)if(!tt(t[i]).test(e))return!1;return!0}for(e in t)this.each(rt(e,t[e]));return this}return this.each(rt(t,e))},W.style=function(t,e,r){var n=arguments.length;if(n<3){if("string"!=typeof t){for(r in n<2&&(e=""),t)this.each(it(r,t[r],e));return this}if(n<2){var i=this.node();return o(i).getComputedStyle(i,null).getPropertyValue(t)}r=""}return this.each(it(t,e,r))},W.property=function(t,e){if(arguments.length<2){if("string"==typeof t)return this.node()[t];for(e in t)this.each(at(e,t[e]));return this}return this.each(at(t,e))},W.text=function(t){return arguments.length?this.each("function"==typeof t?function(){var e=t.apply(this,arguments);this.textContent=null==e?"":e}:null==t?function(){this.textContent=""}:function(){this.textContent=t}):this.node().textContent},W.html=function(t){return arguments.length?this.each("function"==typeof t?function(){var e=t.apply(this,arguments);this.innerHTML=null==e?"":e}:null==t?function(){this.innerHTML=""}:function(){this.innerHTML=t}):this.node().innerHTML},W.append=function(t){return t=ot(t),this.select(function(){return this.appendChild(t.apply(this,arguments))})},W.insert=function(t,e){return t=ot(t),e=X(e),this.select(function(){return this.insertBefore(t.apply(this,arguments),e.apply(this,arguments)||null)})},W.remove=function(){return this.each(st)},W.data=function(t,e){var r,n,i=-1,a=this.length;if(!arguments.length){for(t=new Array(a=(r=this[0]).length);++i<a;)(n=r[i])&&(t[i]=n.__data__);return t}function o(t,r){var n,i,a,o=t.length,u=r.length,h=Math.min(o,u),f=new Array(u),p=new Array(u),d=new Array(o);if(e){var g,v=new b,m=new Array(o);for(n=-1;++n<o;)(i=t[n])&&(v.has(g=e.call(i,i.__data__,n))?d[n]=i:v.set(g,i),m[n]=g);for(n=-1;++n<u;)(i=v.get(g=e.call(r,a=r[n],n)))?!0!==i&&(f[n]=i,i.__data__=a):p[n]=lt(a),v.set(g,!0);for(n=-1;++n<o;)n in m&&!0!==v.get(m[n])&&(d[n]=t[n])}else{for(n=-1;++n<h;)i=t[n],a=r[n],i?(i.__data__=a,f[n]=i):p[n]=lt(a);for(;n<u;++n)p[n]=lt(r[n]);for(;n<o;++n)d[n]=t[n]}p.update=f,p.parentNode=f.parentNode=d.parentNode=t.parentNode,s.push(p),l.push(f),c.push(d)}var s=ht([]),l=q([]),c=q([]);if("function"==typeof t)for(;++i<a;)o(r=this[i],t.call(r,r.parentNode.__data__,i));else for(;++i<a;)o(r=this[i],t);return l.enter=function(){return s},l.exit=function(){return c},l},W.datum=function(t){return arguments.length?this.property("__data__",t):this.property("__data__")},W.filter=function(t){var e,r,n,i=[];"function"!=typeof t&&(t=ct(t));for(var a=0,o=this.length;a<o;a++){i.push(e=[]),e.parentNode=(r=this[a]).parentNode;for(var s=0,l=r.length;s<l;s++)(n=r[s])&&t.call(n,n.__data__,s,a)&&e.push(n)}return q(i)},W.order=function(){for(var t=-1,e=this.length;++t<e;)for(var r,n=this[t],i=n.length-1,a=n[i];--i>=0;)(r=n[i])&&(a&&a!==r.nextSibling&&a.parentNode.insertBefore(r,a),a=r);return this},W.sort=function(t){t=function(t){arguments.length||(t=f);return function(e,r){return e&&r?t(e.__data__,r.__data__):!e-!r}}.apply(this,arguments);for(var e=-1,r=this.length;++e<r;)this[e].sort(t);return this.order()},W.each=function(t){return ut(this,function(e,r,n){t.call(e,e.__data__,r,n)})},W.call=function(t){var e=n(arguments);return t.apply(e[0]=this,e),this},W.empty=function(){return!this.node()},W.node=function(){for(var t=0,e=this.length;t<e;t++)for(var r=this[t],n=0,i=r.length;n<i;n++){var a=r[n];if(a)return a}return null},W.size=function(){var t=0;return ut(this,function(){++t}),t};var ft=[];function pt(e,r,i){var a="__on"+e,o=e.indexOf("."),s=gt;o>0&&(e=e.slice(0,o));var l=dt.get(e);function c(){var t=this[a];t&&(this.removeEventListener(e,t,t.$),delete this[a])}return l&&(e=l,s=vt),o?r?function(){var t=s(r,n(arguments));c.call(this),this.addEventListener(e,this[a]=t,t.$=i),t._=r}:c:r?D:function(){var r,n=new RegExp("^__on([^.]+)"+t.requote(e)+"$");for(var i in this)if(r=i.match(n)){var a=this[i];this.removeEventListener(r[1],a,a.$),delete this[i]}}}t.selection.enter=ht,t.selection.enter.prototype=ft,ft.append=W.append,ft.empty=W.empty,ft.node=W.node,ft.call=W.call,ft.size=W.size,ft.select=function(t){for(var e,r,n,i,a,o=[],s=-1,l=this.length;++s<l;){n=(i=this[s]).update,o.push(e=[]),e.parentNode=i.parentNode;for(var c=-1,u=i.length;++c<u;)(a=i[c])?(e.push(n[c]=r=t.call(i.parentNode,a.__data__,c,s)),r.__data__=a.__data__):e.push(null)}return q(o)},ft.insert=function(t,e){var r,n,i;return arguments.length<2&&(r=this,e=function(t,e,a){var o,s=r[a].update,l=s.length;for(a!=i&&(i=a,n=0),e>=n&&(n=e+1);!(o=s[n])&&++n<l;);return o}),W.insert.call(this,t,e)},t.select=function(t){var e;return"string"==typeof t?(e=[H(t,i)]).parentNode=i.documentElement:(e=[t]).parentNode=a(t),q([e])},t.selectAll=function(t){var e;return"string"==typeof t?(e=n(G(t,i))).parentNode=i.documentElement:(e=n(t)).parentNode=null,q([e])},W.on=function(t,e,r){var n=arguments.length;if(n<3){if("string"!=typeof t){for(r in n<2&&(e=!1),t)this.each(pt(r,t[r],e));return this}if(n<2)return(n=this.node()["__on"+t])&&n._;r=!1}return this.each(pt(t,e,r))};var dt=t.map({mouseenter:"mouseover",mouseleave:"mouseout"});function gt(e,r){return function(n){var i=t.event;t.event=n,r[0]=this.__data__;try{e.apply(this,r)}finally{t.event=i}}}function vt(t,e){var r=gt(t,e);return function(t){var e=t.relatedTarget;e&&(e===this||8&e.compareDocumentPosition(this))||r.call(this,t)}}i&&dt.forEach(function(t){"on"+t in i&&dt.remove(t)});var mt,yt=0;function xt(e){var r=".dragsuppress-"+ ++yt,n="click"+r,i=t.select(o(e)).on("touchmove"+r,B).on("dragstart"+r,B).on("selectstart"+r,B);if(null==mt&&(mt=!("onselectstart"in e)&&z(e.style,"userSelect")),mt){var s=a(e).style,l=s[mt];s[mt]="none"}return function(t){if(i.on(r,null),mt&&(s[mt]=l),t){var e=function(){i.on(n,null)};i.on(n,function(){B(),e()},!0),setTimeout(e,0)}}}t.mouse=function(t){return _t(t,N())};var bt=this.navigator&&/WebKit/.test(this.navigator.userAgent)?-1:0;function _t(e,r){r.changedTouches&&(r=r.changedTouches[0]);var n=e.ownerSVGElement||e;if(n.createSVGPoint){var i=n.createSVGPoint();if(bt<0){var a=o(e);if(a.scrollX||a.scrollY){var s=(n=t.select("body").append("svg").style({position:"absolute",top:0,left:0,margin:0,padding:0,border:"none"},"important"))[0][0].getScreenCTM();bt=!(s.f||s.e),n.remove()}}return bt?(i.x=r.pageX,i.y=r.pageY):(i.x=r.clientX,i.y=r.clientY),[(i=i.matrixTransform(e.getScreenCTM().inverse())).x,i.y]}var l=e.getBoundingClientRect();return[r.clientX-l.left-e.clientLeft,r.clientY-l.top-e.clientTop]}function wt(){return t.event.changedTouches[0].identifier}t.touch=function(t,e,r){if(arguments.length<3&&(r=e,e=N().changedTouches),e)for(var n,i=0,a=e.length;i<a;++i)if((n=e[i]).identifier===r)return _t(t,n)},t.behavior.drag=function(){var e=j(a,"drag","dragstart","dragend"),r=null,n=s(D,t.mouse,o,"mousemove","mouseup"),i=s(wt,t.touch,P,"touchmove","touchend");function a(){this.on("mousedown.drag",n).on("touchstart.drag",i)}function s(n,i,a,o,s){return function(){var l,c=t.event.target.correspondingElement||t.event.target,u=this.parentNode,h=e.of(this,arguments),f=0,p=n(),d=".drag"+(null==p?"":"-"+p),g=t.select(a(c)).on(o+d,function(){var t,e,r=i(u,p);if(!r)return;t=r[0]-m[0],e=r[1]-m[1],f|=t|e,m=r,h({type:"drag",x:r[0]+l[0],y:r[1]+l[1],dx:t,dy:e})}).on(s+d,function(){if(!i(u,p))return;g.on(o+d,null).on(s+d,null),v(f),h({type:"dragend"})}),v=xt(c),m=i(u,p);l=r?[(l=r.apply(this,arguments)).x-m[0],l.y-m[1]]:[0,0],h({type:"dragstart"})}}return a.origin=function(t){return arguments.length?(r=t,a):r},t.rebind(a,e,"on")},t.touches=function(t,e){return arguments.length<2&&(e=N().touches),e?n(e).map(function(e){var r=_t(t,e);return r.identifier=e.identifier,r}):[]};var kt=1e-6,Tt=kt*kt,At=Math.PI,Mt=2*At,St=Mt-kt,Et=At/2,Ct=At/180,Lt=180/At;function Pt(t){return t>0?1:t<0?-1:0}function Ot(t,e,r){return(e[0]-t[0])*(r[1]-t[1])-(e[1]-t[1])*(r[0]-t[0])}function zt(t){return t>1?0:t<-1?At:Math.acos(t)}function It(t){return t>1?Et:t<-1?-Et:Math.asin(t)}function Dt(t){return((t=Math.exp(t))+1/t)/2}function Rt(t){return(t=Math.sin(t/2))*t}var Ft=Math.SQRT2;t.interpolateZoom=function(t,e){var r,n,i=t[0],a=t[1],o=t[2],s=e[0],l=e[1],c=e[2],u=s-i,h=l-a,f=u*u+h*h;if(f<Tt)n=Math.log(c/o)/Ft,r=function(t){return[i+t*u,a+t*h,o*Math.exp(Ft*t*n)]};else{var p=Math.sqrt(f),d=(c*c-o*o+4*f)/(2*o*2*p),g=(c*c-o*o-4*f)/(2*c*2*p),v=Math.log(Math.sqrt(d*d+1)-d),m=Math.log(Math.sqrt(g*g+1)-g);n=(m-v)/Ft,r=function(t){var e,r=t*n,s=Dt(v),l=o/(2*p)*(s*(e=Ft*r+v,((e=Math.exp(2*e))-1)/(e+1))-function(t){return((t=Math.exp(t))-1/t)/2}(v));return[i+l*u,a+l*h,o*s/Dt(Ft*r+v)]}}return r.duration=1e3*n,r},t.behavior.zoom=function(){var e,r,n,a,s,l,c,u,h,f={x:0,y:0,k:1},p=[960,500],d=jt,g=250,v=0,m="mousedown.zoom",y="mousemove.zoom",x="mouseup.zoom",b="touchstart.zoom",_=j(w,"zoomstart","zoom","zoomend");function w(t){t.on(m,P).on(Nt+".zoom",z).on("dblclick.zoom",I).on(b,O)}function k(t){return[(t[0]-f.x)/f.k,(t[1]-f.y)/f.k]}function T(t){f.k=Math.max(d[0],Math.min(d[1],t))}function A(t,e){e=function(t){return[t[0]*f.k+f.x,t[1]*f.k+f.y]}(e),f.x+=t[0]-e[0],f.y+=t[1]-e[1]}function M(e,n,i,a){e.__chart__={x:f.x,y:f.y,k:f.k},T(Math.pow(2,a)),A(r=n,i),e=t.select(e),g>0&&(e=e.transition().duration(g)),e.call(w.event)}function S(){c&&c.domain(l.range().map(function(t){return(t-f.x)/f.k}).map(l.invert)),h&&h.domain(u.range().map(function(t){return(t-f.y)/f.k}).map(u.invert))}function E(t){v++||t({type:"zoomstart"})}function C(t){S(),t({type:"zoom",scale:f.k,translate:[f.x,f.y]})}function L(t){--v||(t({type:"zoomend"}),r=null)}function P(){var e=this,r=_.of(e,arguments),n=0,i=t.select(o(e)).on(y,function(){n=1,A(t.mouse(e),a),C(r)}).on(x,function(){i.on(y,null).on(x,null),s(n),L(r)}),a=k(t.mouse(e)),s=xt(e);hs.call(e),E(r)}function O(){var e,r=this,n=_.of(r,arguments),i={},a=0,o=".zoom-"+t.event.changedTouches[0].identifier,l="touchmove"+o,c="touchend"+o,u=[],h=t.select(r),p=xt(r);function d(){var n=t.touches(r);return e=f.k,n.forEach(function(t){t.identifier in i&&(i[t.identifier]=k(t))}),n}function g(){var e=t.event.target;t.select(e).on(l,v).on(c,y),u.push(e);for(var n=t.event.changedTouches,o=0,h=n.length;o<h;++o)i[n[o].identifier]=null;var p=d(),g=Date.now();if(1===p.length){if(g-s<500){var m=p[0];M(r,m,i[m.identifier],Math.floor(Math.log(f.k)/Math.LN2)+1),B()}s=g}else if(p.length>1){m=p[0];var x=p[1],b=m[0]-x[0],_=m[1]-x[1];a=b*b+_*_}}function v(){var o,l,c,u,h=t.touches(r);hs.call(r);for(var f=0,p=h.length;f<p;++f,u=null)if(c=h[f],u=i[c.identifier]){if(l)break;o=c,l=u}if(u){var d=(d=c[0]-o[0])*d+(d=c[1]-o[1])*d,g=a&&Math.sqrt(d/a);o=[(o[0]+c[0])/2,(o[1]+c[1])/2],l=[(l[0]+u[0])/2,(l[1]+u[1])/2],T(g*e)}s=null,A(o,l),C(n)}function y(){if(t.event.touches.length){for(var e=t.event.changedTouches,r=0,a=e.length;r<a;++r)delete i[e[r].identifier];for(var s in i)return void d()}t.selectAll(u).on(o,null),h.on(m,P).on(b,O),p(),L(n)}g(),E(n),h.on(m,null).on(b,g)}function z(){var i=_.of(this,arguments);a?clearTimeout(a):(hs.call(this),e=k(r=n||t.mouse(this)),E(i)),a=setTimeout(function(){a=null,L(i)},50),B(),T(Math.pow(2,.002*Bt())*f.k),A(r,e),C(i)}function I(){var e=t.mouse(this),r=Math.log(f.k)/Math.LN2;M(this,e,k(e),t.event.shiftKey?Math.ceil(r)-1:Math.floor(r)+1)}return Nt||(Nt="onwheel"in i?(Bt=function(){return-t.event.deltaY*(t.event.deltaMode?120:1)},"wheel"):"onmousewheel"in i?(Bt=function(){return t.event.wheelDelta},"mousewheel"):(Bt=function(){return-t.event.detail},"MozMousePixelScroll")),w.event=function(e){e.each(function(){var e=_.of(this,arguments),n=f;ds?t.select(this).transition().each("start.zoom",function(){f=this.__chart__||{x:0,y:0,k:1},E(e)}).tween("zoom:zoom",function(){var i=p[0],a=p[1],o=r?r[0]:i/2,s=r?r[1]:a/2,l=t.interpolateZoom([(o-f.x)/f.k,(s-f.y)/f.k,i/f.k],[(o-n.x)/n.k,(s-n.y)/n.k,i/n.k]);return function(t){var r=l(t),n=i/r[2];this.__chart__=f={x:o-r[0]*n,y:s-r[1]*n,k:n},C(e)}}).each("interrupt.zoom",function(){L(e)}).each("end.zoom",function(){L(e)}):(this.__chart__=f,E(e),C(e),L(e))})},w.translate=function(t){return arguments.length?(f={x:+t[0],y:+t[1],k:f.k},S(),w):[f.x,f.y]},w.scale=function(t){return arguments.length?(f={x:f.x,y:f.y,k:null},T(+t),S(),w):f.k},w.scaleExtent=function(t){return arguments.length?(d=null==t?jt:[+t[0],+t[1]],w):d},w.center=function(t){return arguments.length?(n=t&&[+t[0],+t[1]],w):n},w.size=function(t){return arguments.length?(p=t&&[+t[0],+t[1]],w):p},w.duration=function(t){return arguments.length?(g=+t,w):g},w.x=function(t){return arguments.length?(c=t,l=t.copy(),f={x:0,y:0,k:1},w):c},w.y=function(t){return arguments.length?(h=t,u=t.copy(),f={x:0,y:0,k:1},w):h},t.rebind(w,_,"on")};var Bt,Nt,jt=[0,1/0];function Ut(){}function Vt(t,e,r){return this instanceof Vt?(this.h=+t,this.s=+e,void(this.l=+r)):arguments.length<2?t instanceof Vt?new Vt(t.h,t.s,t.l):ue(""+t,he,Vt):new Vt(t,e,r)}t.color=Ut,Ut.prototype.toString=function(){return this.rgb()+""},t.hsl=Vt;var qt=Vt.prototype=new Ut;function Ht(t,e,r){var n,i;function a(t){return Math.round(255*function(t){return t>360?t-=360:t<0&&(t+=360),t<60?n+(i-n)*t/60:t<180?i:t<240?n+(i-n)*(240-t)/60:n}(t))}return t=isNaN(t)?0:(t%=360)<0?t+360:t,e=isNaN(e)?0:e<0?0:e>1?1:e,n=2*(r=r<0?0:r>1?1:r)-(i=r<=.5?r*(1+e):r+e-r*e),new ae(a(t+120),a(t),a(t-120))}function Gt(e,r,n){return this instanceof Gt?(this.h=+e,this.c=+r,void(this.l=+n)):arguments.length<2?e instanceof Gt?new Gt(e.h,e.c,e.l):ee(e instanceof Xt?e.l:(e=fe((e=t.rgb(e)).r,e.g,e.b)).l,e.a,e.b):new Gt(e,r,n)}qt.brighter=function(t){return t=Math.pow(.7,arguments.length?t:1),new Vt(this.h,this.s,this.l/t)},qt.darker=function(t){return t=Math.pow(.7,arguments.length?t:1),new Vt(this.h,this.s,t*this.l)},qt.rgb=function(){return Ht(this.h,this.s,this.l)},t.hcl=Gt;var Yt=Gt.prototype=new Ut;function Wt(t,e,r){return isNaN(t)&&(t=0),isNaN(e)&&(e=0),new Xt(r,Math.cos(t*=Ct)*e,Math.sin(t)*e)}function Xt(t,e,r){return this instanceof Xt?(this.l=+t,this.a=+e,void(this.b=+r)):arguments.length<2?t instanceof Xt?new Xt(t.l,t.a,t.b):t instanceof Gt?Wt(t.h,t.c,t.l):fe((t=ae(t)).r,t.g,t.b):new Xt(t,e,r)}Yt.brighter=function(t){return new Gt(this.h,this.c,Math.min(100,this.l+Zt*(arguments.length?t:1)))},Yt.darker=function(t){return new Gt(this.h,this.c,Math.max(0,this.l-Zt*(arguments.length?t:1)))},Yt.rgb=function(){return Wt(this.h,this.c,this.l).rgb()},t.lab=Xt;var Zt=18,Jt=.95047,Kt=1,$t=1.08883,Qt=Xt.prototype=new Ut;function te(t,e,r){var n=(t+16)/116,i=n+e/500,a=n-r/200;return new ae(ie(3.2404542*(i=re(i)*Jt)-1.5371385*(n=re(n)*Kt)-.4985314*(a=re(a)*$t)),ie(-.969266*i+1.8760108*n+.041556*a),ie(.0556434*i-.2040259*n+1.0572252*a))}function ee(t,e,r){return t>0?new Gt(Math.atan2(r,e)*Lt,Math.sqrt(e*e+r*r),t):new Gt(NaN,NaN,t)}function re(t){return t>.206893034?t*t*t:(t-4/29)/7.787037}function ne(t){return t>.008856?Math.pow(t,1/3):7.787037*t+4/29}function ie(t){return Math.round(255*(t<=.00304?12.92*t:1.055*Math.pow(t,1/2.4)-.055))}function ae(t,e,r){return this instanceof ae?(this.r=~~t,this.g=~~e,void(this.b=~~r)):arguments.length<2?t instanceof ae?new ae(t.r,t.g,t.b):ue(""+t,ae,Ht):new ae(t,e,r)}function oe(t){return new ae(t>>16,t>>8&255,255&t)}function se(t){return oe(t)+""}Qt.brighter=function(t){return new Xt(Math.min(100,this.l+Zt*(arguments.length?t:1)),this.a,this.b)},Qt.darker=function(t){return new Xt(Math.max(0,this.l-Zt*(arguments.length?t:1)),this.a,this.b)},Qt.rgb=function(){return te(this.l,this.a,this.b)},t.rgb=ae;var le=ae.prototype=new Ut;function ce(t){return t<16?"0"+Math.max(0,t).toString(16):Math.min(255,t).toString(16)}function ue(t,e,r){var n,i,a,o=0,s=0,l=0;if(n=/([a-z]+)\((.*)\)/.exec(t=t.toLowerCase()))switch(i=n[2].split(","),n[1]){case"hsl":return r(parseFloat(i[0]),parseFloat(i[1])/100,parseFloat(i[2])/100);case"rgb":return e(de(i[0]),de(i[1]),de(i[2]))}return(a=ge.get(t))?e(a.r,a.g,a.b):(null==t||"#"!==t.charAt(0)||isNaN(a=parseInt(t.slice(1),16))||(4===t.length?(o=(3840&a)>>4,o|=o>>4,s=240&a,s|=s>>4,l=15&a,l|=l<<4):7===t.length&&(o=(16711680&a)>>16,s=(65280&a)>>8,l=255&a)),e(o,s,l))}function he(t,e,r){var n,i,a=Math.min(t/=255,e/=255,r/=255),o=Math.max(t,e,r),s=o-a,l=(o+a)/2;return s?(i=l<.5?s/(o+a):s/(2-o-a),n=t==o?(e-r)/s+(e<r?6:0):e==o?(r-t)/s+2:(t-e)/s+4,n*=60):(n=NaN,i=l>0&&l<1?0:n),new Vt(n,i,l)}function fe(t,e,r){var n=ne((.4124564*(t=pe(t))+.3575761*(e=pe(e))+.1804375*(r=pe(r)))/Jt),i=ne((.2126729*t+.7151522*e+.072175*r)/Kt);return Xt(116*i-16,500*(n-i),200*(i-ne((.0193339*t+.119192*e+.9503041*r)/$t)))}function pe(t){return(t/=255)<=.04045?t/12.92:Math.pow((t+.055)/1.055,2.4)}function de(t){var e=parseFloat(t);return"%"===t.charAt(t.length-1)?Math.round(2.55*e):e}le.brighter=function(t){t=Math.pow(.7,arguments.length?t:1);var e=this.r,r=this.g,n=this.b,i=30;return e||r||n?(e&&e<i&&(e=i),r&&r<i&&(r=i),n&&n<i&&(n=i),new ae(Math.min(255,e/t),Math.min(255,r/t),Math.min(255,n/t))):new ae(i,i,i)},le.darker=function(t){return new ae((t=Math.pow(.7,arguments.length?t:1))*this.r,t*this.g,t*this.b)},le.hsl=function(){return he(this.r,this.g,this.b)},le.toString=function(){return"#"+ce(this.r)+ce(this.g)+ce(this.b)};var ge=t.map({aliceblue:15792383,antiquewhite:16444375,aqua:65535,aquamarine:8388564,azure:15794175,beige:16119260,bisque:16770244,black:0,blanchedalmond:16772045,blue:255,blueviolet:9055202,brown:10824234,burlywood:14596231,cadetblue:6266528,chartreuse:8388352,chocolate:13789470,coral:16744272,cornflowerblue:6591981,cornsilk:16775388,crimson:14423100,cyan:65535,darkblue:139,darkcyan:35723,darkgoldenrod:12092939,darkgray:11119017,darkgreen:25600,darkgrey:11119017,darkkhaki:12433259,darkmagenta:9109643,darkolivegreen:5597999,darkorange:16747520,darkorchid:10040012,darkred:9109504,darksalmon:15308410,darkseagreen:9419919,darkslateblue:4734347,darkslategray:3100495,darkslategrey:3100495,darkturquoise:52945,darkviolet:9699539,deeppink:16716947,deepskyblue:49151,dimgray:6908265,dimgrey:6908265,dodgerblue:2003199,firebrick:11674146,floralwhite:16775920,forestgreen:2263842,fuchsia:16711935,gainsboro:14474460,ghostwhite:16316671,gold:16766720,goldenrod:14329120,gray:8421504,green:32768,greenyellow:11403055,grey:8421504,honeydew:15794160,hotpink:16738740,indianred:13458524,indigo:4915330,ivory:16777200,khaki:15787660,lavender:15132410,lavenderblush:16773365,lawngreen:8190976,lemonchiffon:16775885,lightblue:11393254,lightcoral:15761536,lightcyan:14745599,lightgoldenrodyellow:16448210,lightgray:13882323,lightgreen:9498256,lightgrey:13882323,lightpink:16758465,lightsalmon:16752762,lightseagreen:2142890,lightskyblue:8900346,lightslategray:7833753,lightslategrey:7833753,lightsteelblue:11584734,lightyellow:16777184,lime:65280,limegreen:3329330,linen:16445670,magenta:16711935,maroon:8388608,mediumaquamarine:6737322,mediumblue:205,mediumorchid:12211667,mediumpurple:9662683,mediumseagreen:3978097,mediumslateblue:8087790,mediumspringgreen:64154,mediumturquoise:4772300,mediumvioletred:13047173,midnightblue:1644912,mintcream:16121850,mistyrose:16770273,moccasin:16770229,navajowhite:16768685,navy:128,oldlace:16643558,olive:8421376,olivedrab:7048739,orange:16753920,orangered:16729344,orchid:14315734,palegoldenrod:15657130,palegreen:10025880,paleturquoise:11529966,palevioletred:14381203,papayawhip:16773077,peachpuff:16767673,peru:13468991,pink:16761035,plum:14524637,powderblue:11591910,purple:8388736,rebeccapurple:6697881,red:16711680,rosybrown:12357519,royalblue:4286945,saddlebrown:9127187,salmon:16416882,sandybrown:16032864,seagreen:3050327,seashell:16774638,sienna:10506797,silver:12632256,skyblue:8900331,slateblue:6970061,slategray:7372944,slategrey:7372944,snow:16775930,springgreen:65407,steelblue:4620980,tan:13808780,teal:32896,thistle:14204888,tomato:16737095,turquoise:4251856,violet:15631086,wheat:16113331,white:16777215,whitesmoke:16119285,yellow:16776960,yellowgreen:10145074});function ve(t){return"function"==typeof t?t:function(){return t}}function me(t){return function(e,r,n){return 2===arguments.length&&"function"==typeof r&&(n=r,r=null),ye(e,r,t,n)}}function ye(e,r,i,a){var o={},s=t.dispatch("beforesend","progress","load","error"),l={},c=new XMLHttpRequest,u=null;function h(){var t,e=c.status;if(!e&&function(t){var e=t.responseType;return e&&"text"!==e?t.response:t.responseText}(c)||e>=200&&e<300||304===e){try{t=i.call(o,c)}catch(t){return void s.error.call(o,t)}s.load.call(o,t)}else s.error.call(o,c)}return!this.XDomainRequest||"withCredentials"in c||!/^(http(s)?:)?\/\//.test(e)||(c=new XDomainRequest),"onload"in c?c.onload=c.onerror=h:c.onreadystatechange=function(){c.readyState>3&&h()},c.onprogress=function(e){var r=t.event;t.event=e;try{s.progress.call(o,c)}finally{t.event=r}},o.header=function(t,e){return t=(t+"").toLowerCase(),arguments.length<2?l[t]:(null==e?delete l[t]:l[t]=e+"",o)},o.mimeType=function(t){return arguments.length?(r=null==t?null:t+"",o):r},o.responseType=function(t){return arguments.length?(u=t,o):u},o.response=function(t){return i=t,o},["get","post"].forEach(function(t){o[t]=function(){return o.send.apply(o,[t].concat(n(arguments)))}}),o.send=function(t,n,i){if(2===arguments.length&&"function"==typeof n&&(i=n,n=null),c.open(t,e,!0),null==r||"accept"in l||(l.accept=r+",*/*"),c.setRequestHeader)for(var a in l)c.setRequestHeader(a,l[a]);return null!=r&&c.overrideMimeType&&c.overrideMimeType(r),null!=u&&(c.responseType=u),null!=i&&o.on("error",i).on("load",function(t){i(null,t)}),s.beforesend.call(o,c),c.send(null==n?null:n),o},o.abort=function(){return c.abort(),o},t.rebind(o,s,"on"),null==a?o:o.get(function(t){return 1===t.length?function(e,r){t(null==e?r:null)}:t}(a))}ge.forEach(function(t,e){ge.set(t,oe(e))}),t.functor=ve,t.xhr=me(P),t.dsv=function(t,e){var r=new RegExp('["'+t+"\n]"),n=t.charCodeAt(0);function i(t,r,n){arguments.length<3&&(n=r,r=null);var i=ye(t,e,null==r?a:o(r),n);return i.row=function(t){return arguments.length?i.response(null==(r=t)?a:o(t)):r},i}function a(t){return i.parse(t.responseText)}function o(t){return function(e){return i.parse(e.responseText,t)}}function s(e){return e.map(l).join(t)}function l(t){return r.test(t)?'"'+t.replace(/\"/g,'""')+'"':t}return i.parse=function(t,e){var r;return i.parseRows(t,function(t,n){if(r)return r(t,n-1);var i=new Function("d","return {"+t.map(function(t,e){return JSON.stringify(t)+": d["+e+"]"}).join(",")+"}");r=e?function(t,r){return e(i(t),r)}:i})},i.parseRows=function(t,e){var r,i,a={},o={},s=[],l=t.length,c=0,u=0;function h(){if(c>=l)return o;if(i)return i=!1,a;var e=c;if(34===t.charCodeAt(e)){for(var r=e;r++<l;)if(34===t.charCodeAt(r)){if(34!==t.charCodeAt(r+1))break;++r}return c=r+2,13===(s=t.charCodeAt(r+1))?(i=!0,10===t.charCodeAt(r+2)&&++c):10===s&&(i=!0),t.slice(e+1,r).replace(/""/g,'"')}for(;c<l;){var s,u=1;if(10===(s=t.charCodeAt(c++)))i=!0;else if(13===s)i=!0,10===t.charCodeAt(c)&&(++c,++u);else if(s!==n)continue;return t.slice(e,c-u)}return t.slice(e)}for(;(r=h())!==o;){for(var f=[];r!==a&&r!==o;)f.push(r),r=h();e&&null==(f=e(f,u++))||s.push(f)}return s},i.format=function(e){if(Array.isArray(e[0]))return i.formatRows(e);var r=new L,n=[];return e.forEach(function(t){for(var e in t)r.has(e)||n.push(r.add(e))}),[n.map(l).join(t)].concat(e.map(function(e){return n.map(function(t){return l(e[t])}).join(t)})).join("\n")},i.formatRows=function(t){return t.map(s).join("\n")},i},t.csv=t.dsv(",","text/csv"),t.tsv=t.dsv("\t","text/tab-separated-values");var xe,be,_e,we,ke=this[z(this,"requestAnimationFrame")]||function(t){setTimeout(t,17)};function Te(t,e,r){var n=arguments.length;n<2&&(e=0),n<3&&(r=Date.now());var i={c:t,t:r+e,n:null};return be?be.n=i:xe=i,be=i,_e||(we=clearTimeout(we),_e=1,ke(Ae)),i}function Ae(){var t=Me(),e=Se()-t;e>24?(isFinite(e)&&(clearTimeout(we),we=setTimeout(Ae,e)),_e=0):(_e=1,ke(Ae))}function Me(){for(var t=Date.now(),e=xe;e;)t>=e.t&&e.c(t-e.t)&&(e.c=null),e=e.n;return t}function Se(){for(var t,e=xe,r=1/0;e;)e.c?(e.t<r&&(r=e.t),e=(t=e).n):e=t?t.n=e.n:xe=e.n;return be=t,r}function Ee(t,e){return e-(t?Math.ceil(Math.log(t)/Math.LN10):1)}t.timer=function(){Te.apply(this,arguments)},t.timer.flush=function(){Me(),Se()},t.round=function(t,e){return e?Math.round(t*(e=Math.pow(10,e)))/e:Math.round(t)};var Ce=["y","z","a","f","p","n","\xb5","m","","k","M","G","T","P","E","Z","Y"].map(function(t,e){var r=Math.pow(10,3*y(8-e));return{scale:e>8?function(t){return t/r}:function(t){return t*r},symbol:t}});t.formatPrefix=function(e,r){var n=0;return(e=+e)&&(e<0&&(e*=-1),r&&(e=t.round(e,Ee(e,r))),n=1+Math.floor(1e-12+Math.log(e)/Math.LN10),n=Math.max(-24,Math.min(24,3*Math.floor((n-1)/3)))),Ce[8+n/3]};var Le=/(?:([^{])?([<>=^]))?([+\- ])?([$#])?(0)?(\d+)?(,)?(\.-?\d+)?([a-z%])?/i,Pe=t.map({b:function(t){return t.toString(2)},c:function(t){return String.fromCharCode(t)},o:function(t){return t.toString(8)},x:function(t){return t.toString(16)},X:function(t){return t.toString(16).toUpperCase()},g:function(t,e){return t.toPrecision(e)},e:function(t,e){return t.toExponential(e)},f:function(t,e){return t.toFixed(e)},r:function(e,r){return(e=t.round(e,Ee(e,r))).toFixed(Math.max(0,Math.min(20,Ee(e*(1+1e-15),r))))}});function Oe(t){return t+""}var ze=t.time={},Ie=Date;function De(){this._=new Date(arguments.length>1?Date.UTC.apply(this,arguments):arguments[0])}De.prototype={getDate:function(){return this._.getUTCDate()},getDay:function(){return this._.getUTCDay()},getFullYear:function(){return this._.getUTCFullYear()},getHours:function(){return this._.getUTCHours()},getMilliseconds:function(){return this._.getUTCMilliseconds()},getMinutes:function(){return this._.getUTCMinutes()},getMonth:function(){return this._.getUTCMonth()},getSeconds:function(){return this._.getUTCSeconds()},getTime:function(){return this._.getTime()},getTimezoneOffset:function(){return 0},valueOf:function(){return this._.valueOf()},setDate:function(){Re.setUTCDate.apply(this._,arguments)},setDay:function(){Re.setUTCDay.apply(this._,arguments)},setFullYear:function(){Re.setUTCFullYear.apply(this._,arguments)},setHours:function(){Re.setUTCHours.apply(this._,arguments)},setMilliseconds:function(){Re.setUTCMilliseconds.apply(this._,arguments)},setMinutes:function(){Re.setUTCMinutes.apply(this._,arguments)},setMonth:function(){Re.setUTCMonth.apply(this._,arguments)},setSeconds:function(){Re.setUTCSeconds.apply(this._,arguments)},setTime:function(){Re.setTime.apply(this._,arguments)}};var Re=Date.prototype;function Fe(t,e,r){function n(e){var r=t(e),n=a(r,1);return e-r<n-e?r:n}function i(r){return e(r=t(new Ie(r-1)),1),r}function a(t,r){return e(t=new Ie(+t),r),t}function o(t,n,a){var o=i(t),s=[];if(a>1)for(;o<n;)r(o)%a||s.push(new Date(+o)),e(o,1);else for(;o<n;)s.push(new Date(+o)),e(o,1);return s}t.floor=t,t.round=n,t.ceil=i,t.offset=a,t.range=o;var s=t.utc=Be(t);return s.floor=s,s.round=Be(n),s.ceil=Be(i),s.offset=Be(a),s.range=function(t,e,r){try{Ie=De;var n=new De;return n._=t,o(n,e,r)}finally{Ie=Date}},t}function Be(t){return function(e,r){try{Ie=De;var n=new De;return n._=e,t(n,r)._}finally{Ie=Date}}}ze.year=Fe(function(t){return(t=ze.day(t)).setMonth(0,1),t},function(t,e){t.setFullYear(t.getFullYear()+e)},function(t){return t.getFullYear()}),ze.years=ze.year.range,ze.years.utc=ze.year.utc.range,ze.day=Fe(function(t){var e=new Ie(2e3,0);return e.setFullYear(t.getFullYear(),t.getMonth(),t.getDate()),e},function(t,e){t.setDate(t.getDate()+e)},function(t){return t.getDate()-1}),ze.days=ze.day.range,ze.days.utc=ze.day.utc.range,ze.dayOfYear=function(t){var e=ze.year(t);return Math.floor((t-e-6e4*(t.getTimezoneOffset()-e.getTimezoneOffset()))/864e5)},["sunday","monday","tuesday","wednesday","thursday","friday","saturday"].forEach(function(t,e){e=7-e;var r=ze[t]=Fe(function(t){return(t=ze.day(t)).setDate(t.getDate()-(t.getDay()+e)%7),t},function(t,e){t.setDate(t.getDate()+7*Math.floor(e))},function(t){var r=ze.year(t).getDay();return Math.floor((ze.dayOfYear(t)+(r+e)%7)/7)-(r!==e)});ze[t+"s"]=r.range,ze[t+"s"].utc=r.utc.range,ze[t+"OfYear"]=function(t){var r=ze.year(t).getDay();return Math.floor((ze.dayOfYear(t)+(r+e)%7)/7)}}),ze.week=ze.sunday,ze.weeks=ze.sunday.range,ze.weeks.utc=ze.sunday.utc.range,ze.weekOfYear=ze.sundayOfYear;var Ne={"-":"",_:" ",0:"0"},je=/^\s*\d+/,Ue=/^%/;function Ve(t,e,r){var n=t<0?"-":"",i=(n?-t:t)+"",a=i.length;return n+(a<r?new Array(r-a+1).join(e)+i:i)}function qe(e){return new RegExp("^(?:"+e.map(t.requote).join("|")+")","i")}function He(t){for(var e=new b,r=-1,n=t.length;++r<n;)e.set(t[r].toLowerCase(),r);return e}function Ge(t,e,r){je.lastIndex=0;var n=je.exec(e.slice(r,r+1));return n?(t.w=+n[0],r+n[0].length):-1}function Ye(t,e,r){je.lastIndex=0;var n=je.exec(e.slice(r));return n?(t.U=+n[0],r+n[0].length):-1}function We(t,e,r){je.lastIndex=0;var n=je.exec(e.slice(r));return n?(t.W=+n[0],r+n[0].length):-1}function Xe(t,e,r){je.lastIndex=0;var n=je.exec(e.slice(r,r+4));return n?(t.y=+n[0],r+n[0].length):-1}function Ze(t,e,r){je.lastIndex=0;var n,i=je.exec(e.slice(r,r+2));return i?(t.y=(n=+i[0])+(n>68?1900:2e3),r+i[0].length):-1}function Je(t,e,r){return/^[+-]\d{4}$/.test(e=e.slice(r,r+5))?(t.Z=-e,r+5):-1}function Ke(t,e,r){je.lastIndex=0;var n=je.exec(e.slice(r,r+2));return n?(t.m=n[0]-1,r+n[0].length):-1}function $e(t,e,r){je.lastIndex=0;var n=je.exec(e.slice(r,r+2));return n?(t.d=+n[0],r+n[0].length):-1}function Qe(t,e,r){je.lastIndex=0;var n=je.exec(e.slice(r,r+3));return n?(t.j=+n[0],r+n[0].length):-1}function tr(t,e,r){je.lastIndex=0;var n=je.exec(e.slice(r,r+2));return n?(t.H=+n[0],r+n[0].length):-1}function er(t,e,r){je.lastIndex=0;var n=je.exec(e.slice(r,r+2));return n?(t.M=+n[0],r+n[0].length):-1}function rr(t,e,r){je.lastIndex=0;var n=je.exec(e.slice(r,r+2));return n?(t.S=+n[0],r+n[0].length):-1}function nr(t,e,r){je.lastIndex=0;var n=je.exec(e.slice(r,r+3));return n?(t.L=+n[0],r+n[0].length):-1}function ir(t){var e=t.getTimezoneOffset(),r=e>0?"-":"+",n=y(e)/60|0,i=y(e)%60;return r+Ve(n,"0",2)+Ve(i,"0",2)}function ar(t,e,r){Ue.lastIndex=0;var n=Ue.exec(e.slice(r,r+1));return n?r+n[0].length:-1}function or(t){for(var e=t.length,r=-1;++r<e;)t[r][0]=this(t[r][0]);return function(e){for(var r=0,n=t[r];!n[1](e);)n=t[++r];return n[0](e)}}t.locale=function(e){return{numberFormat:function(e){var r=e.decimal,n=e.thousands,i=e.grouping,a=e.currency,o=i&&n?function(t,e){for(var r=t.length,a=[],o=0,s=i[0],l=0;r>0&&s>0&&(l+s+1>e&&(s=Math.max(1,e-l)),a.push(t.substring(r-=s,r+s)),!((l+=s+1)>e));)s=i[o=(o+1)%i.length];return a.reverse().join(n)}:P;return function(e){var n=Le.exec(e),i=n[1]||" ",s=n[2]||">",l=n[3]||"-",c=n[4]||"",u=n[5],h=+n[6],f=n[7],p=n[8],d=n[9],g=1,v="",m="",y=!1,x=!0;switch(p&&(p=+p.substring(1)),(u||"0"===i&&"="===s)&&(u=i="0",s="="),d){case"n":f=!0,d="g";break;case"%":g=100,m="%",d="f";break;case"p":g=100,m="%",d="r";break;case"b":case"o":case"x":case"X":"#"===c&&(v="0"+d.toLowerCase());case"c":x=!1;case"d":y=!0,p=0;break;case"s":g=-1,d="r"}"$"===c&&(v=a[0],m=a[1]),"r"!=d||p||(d="g"),null!=p&&("g"==d?p=Math.max(1,Math.min(21,p)):"e"!=d&&"f"!=d||(p=Math.max(0,Math.min(20,p)))),d=Pe.get(d)||Oe;var b=u&&f;return function(e){var n=m;if(y&&e%1)return"";var a=e<0||0===e&&1/e<0?(e=-e,"-"):"-"===l?"":l;if(g<0){var c=t.formatPrefix(e,p);e=c.scale(e),n=c.symbol+m}else e*=g;var _,w,k=(e=d(e,p)).lastIndexOf(".");if(k<0){var T=x?e.lastIndexOf("e"):-1;T<0?(_=e,w=""):(_=e.substring(0,T),w=e.substring(T))}else _=e.substring(0,k),w=r+e.substring(k+1);!u&&f&&(_=o(_,1/0));var A=v.length+_.length+w.length+(b?0:a.length),M=A<h?new Array(A=h-A+1).join(i):"";return b&&(_=o(M+_,M.length?h-w.length:1/0)),a+=v,e=_+w,("<"===s?a+e+M:">"===s?M+a+e:"^"===s?M.substring(0,A>>=1)+a+e+M.substring(A):a+(b?e:M+e))+n}}}(e),timeFormat:function(e){var r=e.dateTime,n=e.date,i=e.time,a=e.periods,o=e.days,s=e.shortDays,l=e.months,c=e.shortMonths;function u(t){var e=t.length;function r(r){for(var n,i,a,o=[],s=-1,l=0;++s<e;)37===t.charCodeAt(s)&&(o.push(t.slice(l,s)),null!=(i=Ne[n=t.charAt(++s)])&&(n=t.charAt(++s)),(a=_[n])&&(n=a(r,null==i?"e"===n?" ":"0":i)),o.push(n),l=s+1);return o.push(t.slice(l,s)),o.join("")}return r.parse=function(e){var r={y:1900,m:0,d:1,H:0,M:0,S:0,L:0,Z:null};if(h(r,t,e,0)!=e.length)return null;"p"in r&&(r.H=r.H%12+12*r.p);var n=null!=r.Z&&Ie!==De,i=new(n?De:Ie);return"j"in r?i.setFullYear(r.y,0,r.j):"W"in r||"U"in r?("w"in r||(r.w="W"in r?1:0),i.setFullYear(r.y,0,1),i.setFullYear(r.y,0,"W"in r?(r.w+6)%7+7*r.W-(i.getDay()+5)%7:r.w+7*r.U-(i.getDay()+6)%7)):i.setFullYear(r.y,r.m,r.d),i.setHours(r.H+(r.Z/100|0),r.M+r.Z%100,r.S,r.L),n?i._:i},r.toString=function(){return t},r}function h(t,e,r,n){for(var i,a,o,s=0,l=e.length,c=r.length;s<l;){if(n>=c)return-1;if(37===(i=e.charCodeAt(s++))){if(o=e.charAt(s++),!(a=w[o in Ne?e.charAt(s++):o])||(n=a(t,r,n))<0)return-1}else if(i!=r.charCodeAt(n++))return-1}return n}u.utc=function(t){var e=u(t);function r(t){try{var r=new(Ie=De);return r._=t,e(r)}finally{Ie=Date}}return r.parse=function(t){try{Ie=De;var r=e.parse(t);return r&&r._}finally{Ie=Date}},r.toString=e.toString,r},u.multi=u.utc.multi=or;var f=t.map(),p=qe(o),d=He(o),g=qe(s),v=He(s),m=qe(l),y=He(l),x=qe(c),b=He(c);a.forEach(function(t,e){f.set(t.toLowerCase(),e)});var _={a:function(t){return s[t.getDay()]},A:function(t){return o[t.getDay()]},b:function(t){return c[t.getMonth()]},B:function(t){return l[t.getMonth()]},c:u(r),d:function(t,e){return Ve(t.getDate(),e,2)},e:function(t,e){return Ve(t.getDate(),e,2)},H:function(t,e){return Ve(t.getHours(),e,2)},I:function(t,e){return Ve(t.getHours()%12||12,e,2)},j:function(t,e){return Ve(1+ze.dayOfYear(t),e,3)},L:function(t,e){return Ve(t.getMilliseconds(),e,3)},m:function(t,e){return Ve(t.getMonth()+1,e,2)},M:function(t,e){return Ve(t.getMinutes(),e,2)},p:function(t){return a[+(t.getHours()>=12)]},S:function(t,e){return Ve(t.getSeconds(),e,2)},U:function(t,e){return Ve(ze.sundayOfYear(t),e,2)},w:function(t){return t.getDay()},W:function(t,e){return Ve(ze.mondayOfYear(t),e,2)},x:u(n),X:u(i),y:function(t,e){return Ve(t.getFullYear()%100,e,2)},Y:function(t,e){return Ve(t.getFullYear()%1e4,e,4)},Z:ir,"%":function(){return"%"}},w={a:function(t,e,r){g.lastIndex=0;var n=g.exec(e.slice(r));return n?(t.w=v.get(n[0].toLowerCase()),r+n[0].length):-1},A:function(t,e,r){p.lastIndex=0;var n=p.exec(e.slice(r));return n?(t.w=d.get(n[0].toLowerCase()),r+n[0].length):-1},b:function(t,e,r){x.lastIndex=0;var n=x.exec(e.slice(r));return n?(t.m=b.get(n[0].toLowerCase()),r+n[0].length):-1},B:function(t,e,r){m.lastIndex=0;var n=m.exec(e.slice(r));return n?(t.m=y.get(n[0].toLowerCase()),r+n[0].length):-1},c:function(t,e,r){return h(t,_.c.toString(),e,r)},d:$e,e:$e,H:tr,I:tr,j:Qe,L:nr,m:Ke,M:er,p:function(t,e,r){var n=f.get(e.slice(r,r+=2).toLowerCase());return null==n?-1:(t.p=n,r)},S:rr,U:Ye,w:Ge,W:We,x:function(t,e,r){return h(t,_.x.toString(),e,r)},X:function(t,e,r){return h(t,_.X.toString(),e,r)},y:Ze,Y:Xe,Z:Je,"%":ar};return u}(e)}};var sr=t.locale({decimal:".",thousands:",",grouping:[3],currency:["$",""],dateTime:"%a %b %e %X %Y",date:"%m/%d/%Y",time:"%H:%M:%S",periods:["AM","PM"],days:["Sunday","Monday","Tuesday","Wednesday","Thursday","Friday","Saturday"],shortDays:["Sun","Mon","Tue","Wed","Thu","Fri","Sat"],months:["January","February","March","April","May","June","July","August","September","October","November","December"],shortMonths:["Jan","Feb","Mar","Apr","May","Jun","Jul","Aug","Sep","Oct","Nov","Dec"]});function lr(){}t.format=sr.numberFormat,t.geo={},lr.prototype={s:0,t:0,add:function(t){ur(t,this.t,cr),ur(cr.s,this.s,this),this.s?this.t+=cr.t:this.s=cr.t},reset:function(){this.s=this.t=0},valueOf:function(){return this.s}};var cr=new lr;function ur(t,e,r){var n=r.s=t+e,i=n-t,a=n-i;r.t=t-a+(e-i)}function hr(t,e){t&&pr.hasOwnProperty(t.type)&&pr[t.type](t,e)}t.geo.stream=function(t,e){t&&fr.hasOwnProperty(t.type)?fr[t.type](t,e):hr(t,e)};var fr={Feature:function(t,e){hr(t.geometry,e)},FeatureCollection:function(t,e){for(var r=t.features,n=-1,i=r.length;++n<i;)hr(r[n].geometry,e)}},pr={Sphere:function(t,e){e.sphere()},Point:function(t,e){t=t.coordinates,e.point(t[0],t[1],t[2])},MultiPoint:function(t,e){for(var r=t.coordinates,n=-1,i=r.length;++n<i;)t=r[n],e.point(t[0],t[1],t[2])},LineString:function(t,e){dr(t.coordinates,e,0)},MultiLineString:function(t,e){for(var r=t.coordinates,n=-1,i=r.length;++n<i;)dr(r[n],e,0)},Polygon:function(t,e){gr(t.coordinates,e)},MultiPolygon:function(t,e){for(var r=t.coordinates,n=-1,i=r.length;++n<i;)gr(r[n],e)},GeometryCollection:function(t,e){for(var r=t.geometries,n=-1,i=r.length;++n<i;)hr(r[n],e)}};function dr(t,e,r){var n,i=-1,a=t.length-r;for(e.lineStart();++i<a;)n=t[i],e.point(n[0],n[1],n[2]);e.lineEnd()}function gr(t,e){var r=-1,n=t.length;for(e.polygonStart();++r<n;)dr(t[r],e,1);e.polygonEnd()}t.geo.area=function(e){return vr=0,t.geo.stream(e,Cr),vr};var vr,mr,yr,xr,br,_r,wr,kr,Tr,Ar,Mr,Sr,Er=new lr,Cr={sphere:function(){vr+=4*At},point:D,lineStart:D,lineEnd:D,polygonStart:function(){Er.reset(),Cr.lineStart=Lr},polygonEnd:function(){var t=2*Er;vr+=t<0?4*At+t:t,Cr.lineStart=Cr.lineEnd=Cr.point=D}};function Lr(){var t,e,r,n,i;function a(t,e){e=e*Ct/2+At/4;var a=(t*=Ct)-r,o=a>=0?1:-1,s=o*a,l=Math.cos(e),c=Math.sin(e),u=i*c,h=n*l+u*Math.cos(s),f=u*o*Math.sin(s);Er.add(Math.atan2(f,h)),r=t,n=l,i=c}Cr.point=function(o,s){Cr.point=a,r=(t=o)*Ct,n=Math.cos(s=(e=s)*Ct/2+At/4),i=Math.sin(s)},Cr.lineEnd=function(){a(t,e)}}function Pr(t){var e=t[0],r=t[1],n=Math.cos(r);return[n*Math.cos(e),n*Math.sin(e),Math.sin(r)]}function Or(t,e){return t[0]*e[0]+t[1]*e[1]+t[2]*e[2]}function zr(t,e){return[t[1]*e[2]-t[2]*e[1],t[2]*e[0]-t[0]*e[2],t[0]*e[1]-t[1]*e[0]]}function Ir(t,e){t[0]+=e[0],t[1]+=e[1],t[2]+=e[2]}function Dr(t,e){return[t[0]*e,t[1]*e,t[2]*e]}function Rr(t){var e=Math.sqrt(t[0]*t[0]+t[1]*t[1]+t[2]*t[2]);t[0]/=e,t[1]/=e,t[2]/=e}function Fr(t){return[Math.atan2(t[1],t[0]),It(t[2])]}function Br(t,e){return y(t[0]-e[0])<kt&&y(t[1]-e[1])<kt}t.geo.bounds=function(){var e,r,n,i,a,o,s,l,c,u,h,f={point:p,lineStart:g,lineEnd:v,polygonStart:function(){f.point=m,f.lineStart=x,f.lineEnd=b,c=0,Cr.polygonStart()},polygonEnd:function(){Cr.polygonEnd(),f.point=p,f.lineStart=g,f.lineEnd=v,Er<0?(e=-(n=180),r=-(i=90)):c>kt?i=90:c<-kt&&(r=-90),h[0]=e,h[1]=n}};function p(t,a){u.push(h=[e=t,n=t]),a<r&&(r=a),a>i&&(i=a)}function d(t,o){var s=Pr([t*Ct,o*Ct]);if(l){var c=zr(l,s),u=zr([c[1],-c[0],0],c);Rr(u),u=Fr(u);var h=t-a,f=h>0?1:-1,d=u[0]*Lt*f,g=y(h)>180;if(g^(f*a<d&&d<f*t))(v=u[1]*Lt)>i&&(i=v);else if(g^(f*a<(d=(d+360)%360-180)&&d<f*t)){var v;(v=-u[1]*Lt)<r&&(r=v)}else o<r&&(r=o),o>i&&(i=o);g?t<a?_(e,t)>_(e,n)&&(n=t):_(t,n)>_(e,n)&&(e=t):n>=e?(t<e&&(e=t),t>n&&(n=t)):t>a?_(e,t)>_(e,n)&&(n=t):_(t,n)>_(e,n)&&(e=t)}else p(t,o);l=s,a=t}function g(){f.point=d}function v(){h[0]=e,h[1]=n,f.point=p,l=null}function m(t,e){if(l){var r=t-a;c+=y(r)>180?r+(r>0?360:-360):r}else o=t,s=e;Cr.point(t,e),d(t,e)}function x(){Cr.lineStart()}function b(){m(o,s),Cr.lineEnd(),y(c)>kt&&(e=-(n=180)),h[0]=e,h[1]=n,l=null}function _(t,e){return(e-=t)<0?e+360:e}function w(t,e){return t[0]-e[0]}function k(t,e){return e[0]<=e[1]?e[0]<=t&&t<=e[1]:t<e[0]||e[1]<t}return function(a){if(i=n=-(e=r=1/0),u=[],t.geo.stream(a,f),c=u.length){u.sort(w);for(var o=1,s=[g=u[0]];o<c;++o)k((p=u[o])[0],g)||k(p[1],g)?(_(g[0],p[1])>_(g[0],g[1])&&(g[1]=p[1]),_(p[0],g[1])>_(g[0],g[1])&&(g[0]=p[0])):s.push(g=p);for(var l,c,p,d=-1/0,g=(o=0,s[c=s.length-1]);o<=c;g=p,++o)p=s[o],(l=_(g[1],p[0]))>d&&(d=l,e=p[0],n=g[1])}return u=h=null,e===1/0||r===1/0?[[NaN,NaN],[NaN,NaN]]:[[e,r],[n,i]]}}(),t.geo.centroid=function(e){mr=yr=xr=br=_r=wr=kr=Tr=Ar=Mr=Sr=0,t.geo.stream(e,Nr);var r=Ar,n=Mr,i=Sr,a=r*r+n*n+i*i;return a<Tt&&(r=wr,n=kr,i=Tr,yr<kt&&(r=xr,n=br,i=_r),(a=r*r+n*n+i*i)<Tt)?[NaN,NaN]:[Math.atan2(n,r)*Lt,It(i/Math.sqrt(a))*Lt]};var Nr={sphere:D,point:jr,lineStart:Vr,lineEnd:qr,polygonStart:function(){Nr.lineStart=Hr},polygonEnd:function(){Nr.lineStart=Vr}};function jr(t,e){t*=Ct;var r=Math.cos(e*=Ct);Ur(r*Math.cos(t),r*Math.sin(t),Math.sin(e))}function Ur(t,e,r){xr+=(t-xr)/++mr,br+=(e-br)/mr,_r+=(r-_r)/mr}function Vr(){var t,e,r;function n(n,i){n*=Ct;var a=Math.cos(i*=Ct),o=a*Math.cos(n),s=a*Math.sin(n),l=Math.sin(i),c=Math.atan2(Math.sqrt((c=e*l-r*s)*c+(c=r*o-t*l)*c+(c=t*s-e*o)*c),t*o+e*s+r*l);yr+=c,wr+=c*(t+(t=o)),kr+=c*(e+(e=s)),Tr+=c*(r+(r=l)),Ur(t,e,r)}Nr.point=function(i,a){i*=Ct;var o=Math.cos(a*=Ct);t=o*Math.cos(i),e=o*Math.sin(i),r=Math.sin(a),Nr.point=n,Ur(t,e,r)}}function qr(){Nr.point=jr}function Hr(){var t,e,r,n,i;function a(t,e){t*=Ct;var a=Math.cos(e*=Ct),o=a*Math.cos(t),s=a*Math.sin(t),l=Math.sin(e),c=n*l-i*s,u=i*o-r*l,h=r*s-n*o,f=Math.sqrt(c*c+u*u+h*h),p=r*o+n*s+i*l,d=f&&-zt(p)/f,g=Math.atan2(f,p);Ar+=d*c,Mr+=d*u,Sr+=d*h,yr+=g,wr+=g*(r+(r=o)),kr+=g*(n+(n=s)),Tr+=g*(i+(i=l)),Ur(r,n,i)}Nr.point=function(o,s){t=o,e=s,Nr.point=a,o*=Ct;var l=Math.cos(s*=Ct);r=l*Math.cos(o),n=l*Math.sin(o),i=Math.sin(s),Ur(r,n,i)},Nr.lineEnd=function(){a(t,e),Nr.lineEnd=qr,Nr.point=jr}}function Gr(t,e){function r(r,n){return r=t(r,n),e(r[0],r[1])}return t.invert&&e.invert&&(r.invert=function(r,n){return(r=e.invert(r,n))&&t.invert(r[0],r[1])}),r}function Yr(){return!0}function Wr(t,e,r,n,i){var a=[],o=[];if(t.forEach(function(t){if(!((e=t.length-1)<=0)){var e,r=t[0],n=t[e];if(Br(r,n)){i.lineStart();for(var s=0;s<e;++s)i.point((r=t[s])[0],r[1]);i.lineEnd()}else{var l=new Zr(r,t,null,!0),c=new Zr(r,null,l,!1);l.o=c,a.push(l),o.push(c),l=new Zr(n,t,null,!1),c=new Zr(n,null,l,!0),l.o=c,a.push(l),o.push(c)}}}),o.sort(e),Xr(a),Xr(o),a.length){for(var s=0,l=r,c=o.length;s<c;++s)o[s].e=l=!l;for(var u,h,f=a[0];;){for(var p=f,d=!0;p.v;)if((p=p.n)===f)return;u=p.z,i.lineStart();do{if(p.v=p.o.v=!0,p.e){if(d)for(s=0,c=u.length;s<c;++s)i.point((h=u[s])[0],h[1]);else n(p.x,p.n.x,1,i);p=p.n}else{if(d)for(s=(u=p.p.z).length-1;s>=0;--s)i.point((h=u[s])[0],h[1]);else n(p.x,p.p.x,-1,i);p=p.p}u=(p=p.o).z,d=!d}while(!p.v);i.lineEnd()}}}function Xr(t){if(e=t.length){for(var e,r,n=0,i=t[0];++n<e;)i.n=r=t[n],r.p=i,i=r;i.n=r=t[0],r.p=i}}function Zr(t,e,r,n){this.x=t,this.z=e,this.o=r,this.e=n,this.v=!1,this.n=this.p=null}function Jr(e,r,n,i){return function(a,o){var s,l=r(o),c=a.invert(i[0],i[1]),u={point:h,lineStart:p,lineEnd:d,polygonStart:function(){u.point=b,u.lineStart=_,u.lineEnd=w,s=[],g=[]},polygonEnd:function(){u.point=h,u.lineStart=p,u.lineEnd=d,s=t.merge(s);var e=function(t,e){var r=t[0],n=t[1],i=[Math.sin(r),-Math.cos(r),0],a=0,o=0;Er.reset();for(var s=0,l=e.length;s<l;++s){var c=e[s],u=c.length;if(u)for(var h=c[0],f=h[0],p=h[1]/2+At/4,d=Math.sin(p),g=Math.cos(p),v=1;;){v===u&&(v=0);var m=(t=c[v])[0],y=t[1]/2+At/4,x=Math.sin(y),b=Math.cos(y),_=m-f,w=_>=0?1:-1,k=w*_,T=k>At,A=d*x;if(Er.add(Math.atan2(A*w*Math.sin(k),g*b+A*Math.cos(k))),a+=T?_+w*Mt:_,T^f>=r^m>=r){var M=zr(Pr(h),Pr(t));Rr(M);var S=zr(i,M);Rr(S);var E=(T^_>=0?-1:1)*It(S[2]);(n>E||n===E&&(M[0]||M[1]))&&(o+=T^_>=0?1:-1)}if(!v++)break;f=m,d=x,g=b,h=t}}return(a<-kt||a<kt&&Er<-kt)^1&o}(c,g);s.length?(x||(o.polygonStart(),x=!0),Wr(s,Qr,e,n,o)):e&&(x||(o.polygonStart(),x=!0),o.lineStart(),n(null,null,1,o),o.lineEnd()),x&&(o.polygonEnd(),x=!1),s=g=null},sphere:function(){o.polygonStart(),o.lineStart(),n(null,null,1,o),o.lineEnd(),o.polygonEnd()}};function h(t,r){var n=a(t,r);e(t=n[0],r=n[1])&&o.point(t,r)}function f(t,e){var r=a(t,e);l.point(r[0],r[1])}function p(){u.point=f,l.lineStart()}function d(){u.point=h,l.lineEnd()}var g,v,m=$r(),y=r(m),x=!1;function b(t,e){v.push([t,e]);var r=a(t,e);y.point(r[0],r[1])}function _(){y.lineStart(),v=[]}function w(){b(v[0][0],v[0][1]),y.lineEnd();var t,e=y.clean(),r=m.buffer(),n=r.length;if(v.pop(),g.push(v),v=null,n)if(1&e){var i,a=-1;if((n=(t=r[0]).length-1)>0){for(x||(o.polygonStart(),x=!0),o.lineStart();++a<n;)o.point((i=t[a])[0],i[1]);o.lineEnd()}}else n>1&&2&e&&r.push(r.pop().concat(r.shift())),s.push(r.filter(Kr))}return u}}function Kr(t){return t.length>1}function $r(){var t,e=[];return{lineStart:function(){e.push(t=[])},point:function(e,r){t.push([e,r])},lineEnd:D,buffer:function(){var r=e;return e=[],t=null,r},rejoin:function(){e.length>1&&e.push(e.pop().concat(e.shift()))}}}function Qr(t,e){return((t=t.x)[0]<0?t[1]-Et-kt:Et-t[1])-((e=e.x)[0]<0?e[1]-Et-kt:Et-e[1])}var tn=Jr(Yr,function(t){var e,r=NaN,n=NaN,i=NaN;return{lineStart:function(){t.lineStart(),e=1},point:function(a,o){var s=a>0?At:-At,l=y(a-r);y(l-At)<kt?(t.point(r,n=(n+o)/2>0?Et:-Et),t.point(i,n),t.lineEnd(),t.lineStart(),t.point(s,n),t.point(a,n),e=0):i!==s&&l>=At&&(y(r-i)<kt&&(r-=i*kt),y(a-s)<kt&&(a-=s*kt),n=function(t,e,r,n){var i,a,o=Math.sin(t-r);return y(o)>kt?Math.atan((Math.sin(e)*(a=Math.cos(n))*Math.sin(r)-Math.sin(n)*(i=Math.cos(e))*Math.sin(t))/(i*a*o)):(e+n)/2}(r,n,a,o),t.point(i,n),t.lineEnd(),t.lineStart(),t.point(s,n),e=0),t.point(r=a,n=o),i=s},lineEnd:function(){t.lineEnd(),r=n=NaN},clean:function(){return 2-e}}},function(t,e,r,n){var i;if(null==t)i=r*Et,n.point(-At,i),n.point(0,i),n.point(At,i),n.point(At,0),n.point(At,-i),n.point(0,-i),n.point(-At,-i),n.point(-At,0),n.point(-At,i);else if(y(t[0]-e[0])>kt){var a=t[0]<e[0]?At:-At;i=r*a/2,n.point(-a,i),n.point(0,i),n.point(a,i)}else n.point(e[0],e[1])},[-At,-At/2]);function en(t,e,r,n){return function(i){var a,o=i.a,s=i.b,l=o.x,c=o.y,u=0,h=1,f=s.x-l,p=s.y-c;if(a=t-l,f||!(a>0)){if(a/=f,f<0){if(a<u)return;a<h&&(h=a)}else if(f>0){if(a>h)return;a>u&&(u=a)}if(a=r-l,f||!(a<0)){if(a/=f,f<0){if(a>h)return;a>u&&(u=a)}else if(f>0){if(a<u)return;a<h&&(h=a)}if(a=e-c,p||!(a>0)){if(a/=p,p<0){if(a<u)return;a<h&&(h=a)}else if(p>0){if(a>h)return;a>u&&(u=a)}if(a=n-c,p||!(a<0)){if(a/=p,p<0){if(a>h)return;a>u&&(u=a)}else if(p>0){if(a<u)return;a<h&&(h=a)}return u>0&&(i.a={x:l+u*f,y:c+u*p}),h<1&&(i.b={x:l+h*f,y:c+h*p}),i}}}}}}var rn=1e9;function nn(e,r,n,i){return function(l){var c,u,h,f,p,d,g,v,m,y,x,b=l,_=$r(),w=en(e,r,n,i),k={point:M,lineStart:function(){k.point=S,u&&u.push(h=[]);y=!0,m=!1,g=v=NaN},lineEnd:function(){c&&(S(f,p),d&&m&&_.rejoin(),c.push(_.buffer()));k.point=M,m&&l.lineEnd()},polygonStart:function(){l=_,c=[],u=[],x=!0},polygonEnd:function(){l=b,c=t.merge(c);var r=function(t){for(var e=0,r=u.length,n=t[1],i=0;i<r;++i)for(var a,o=1,s=u[i],l=s.length,c=s[0];o<l;++o)a=s[o],c[1]<=n?a[1]>n&&Ot(c,a,t)>0&&++e:a[1]<=n&&Ot(c,a,t)<0&&--e,c=a;return 0!==e}([e,i]),n=x&&r,a=c.length;(n||a)&&(l.polygonStart(),n&&(l.lineStart(),T(null,null,1,l),l.lineEnd()),a&&Wr(c,o,r,T,l),l.polygonEnd()),c=u=h=null}};function T(t,o,l,c){var u=0,h=0;if(null==t||(u=a(t,l))!==(h=a(o,l))||s(t,o)<0^l>0)do{c.point(0===u||3===u?e:n,u>1?i:r)}while((u=(u+l+4)%4)!==h);else c.point(o[0],o[1])}function A(t,a){return e<=t&&t<=n&&r<=a&&a<=i}function M(t,e){A(t,e)&&l.point(t,e)}function S(t,e){var r=A(t=Math.max(-rn,Math.min(rn,t)),e=Math.max(-rn,Math.min(rn,e)));if(u&&h.push([t,e]),y)f=t,p=e,d=r,y=!1,r&&(l.lineStart(),l.point(t,e));else if(r&&m)l.point(t,e);else{var n={a:{x:g,y:v},b:{x:t,y:e}};w(n)?(m||(l.lineStart(),l.point(n.a.x,n.a.y)),l.point(n.b.x,n.b.y),r||l.lineEnd(),x=!1):r&&(l.lineStart(),l.point(t,e),x=!1)}g=t,v=e,m=r}return k};function a(t,i){return y(t[0]-e)<kt?i>0?0:3:y(t[0]-n)<kt?i>0?2:1:y(t[1]-r)<kt?i>0?1:0:i>0?3:2}function o(t,e){return s(t.x,e.x)}function s(t,e){var r=a(t,1),n=a(e,1);return r!==n?r-n:0===r?e[1]-t[1]:1===r?t[0]-e[0]:2===r?t[1]-e[1]:e[0]-t[0]}}function an(t){var e=0,r=At/3,n=Cn(t),i=n(e,r);return i.parallels=function(t){return arguments.length?n(e=t[0]*At/180,r=t[1]*At/180):[e/At*180,r/At*180]},i}function on(t,e){var r=Math.sin(t),n=(r+Math.sin(e))/2,i=1+r*(2*n-r),a=Math.sqrt(i)/n;function o(t,e){var r=Math.sqrt(i-2*n*Math.sin(e))/n;return[r*Math.sin(t*=n),a-r*Math.cos(t)]}return o.invert=function(t,e){var r=a-e;return[Math.atan2(t,r)/n,It((i-(t*t+r*r)*n*n)/(2*n))]},o}t.geo.clipExtent=function(){var t,e,r,n,i,a,o={stream:function(t){return i&&(i.valid=!1),(i=a(t)).valid=!0,i},extent:function(s){return arguments.length?(a=nn(t=+s[0][0],e=+s[0][1],r=+s[1][0],n=+s[1][1]),i&&(i.valid=!1,i=null),o):[[t,e],[r,n]]}};return o.extent([[0,0],[960,500]])},(t.geo.conicEqualArea=function(){return an(on)}).raw=on,t.geo.albers=function(){return t.geo.conicEqualArea().rotate([96,0]).center([-.6,38.7]).parallels([29.5,45.5]).scale(1070)},t.geo.albersUsa=function(){var e,r,n,i,a=t.geo.albers(),o=t.geo.conicEqualArea().rotate([154,0]).center([-2,58.5]).parallels([55,65]),s=t.geo.conicEqualArea().rotate([157,0]).center([-3,19.9]).parallels([8,18]),l={point:function(t,r){e=[t,r]}};function c(t){var a=t[0],o=t[1];return e=null,r(a,o),e||(n(a,o),e)||i(a,o),e}return c.invert=function(t){var e=a.scale(),r=a.translate(),n=(t[0]-r[0])/e,i=(t[1]-r[1])/e;return(i>=.12&&i<.234&&n>=-.425&&n<-.214?o:i>=.166&&i<.234&&n>=-.214&&n<-.115?s:a).invert(t)},c.stream=function(t){var e=a.stream(t),r=o.stream(t),n=s.stream(t);return{point:function(t,i){e.point(t,i),r.point(t,i),n.point(t,i)},sphere:function(){e.sphere(),r.sphere(),n.sphere()},lineStart:function(){e.lineStart(),r.lineStart(),n.lineStart()},lineEnd:function(){e.lineEnd(),r.lineEnd(),n.lineEnd()},polygonStart:function(){e.polygonStart(),r.polygonStart(),n.polygonStart()},polygonEnd:function(){e.polygonEnd(),r.polygonEnd(),n.polygonEnd()}}},c.precision=function(t){return arguments.length?(a.precision(t),o.precision(t),s.precision(t),c):a.precision()},c.scale=function(t){return arguments.length?(a.scale(t),o.scale(.35*t),s.scale(t),c.translate(a.translate())):a.scale()},c.translate=function(t){if(!arguments.length)return a.translate();var e=a.scale(),u=+t[0],h=+t[1];return r=a.translate(t).clipExtent([[u-.455*e,h-.238*e],[u+.455*e,h+.238*e]]).stream(l).point,n=o.translate([u-.307*e,h+.201*e]).clipExtent([[u-.425*e+kt,h+.12*e+kt],[u-.214*e-kt,h+.234*e-kt]]).stream(l).point,i=s.translate([u-.205*e,h+.212*e]).clipExtent([[u-.214*e+kt,h+.166*e+kt],[u-.115*e-kt,h+.234*e-kt]]).stream(l).point,c},c.scale(1070)};var sn,ln,cn,un,hn,fn,pn={point:D,lineStart:D,lineEnd:D,polygonStart:function(){ln=0,pn.lineStart=dn},polygonEnd:function(){pn.lineStart=pn.lineEnd=pn.point=D,sn+=y(ln/2)}};function dn(){var t,e,r,n;function i(t,e){ln+=n*t-r*e,r=t,n=e}pn.point=function(a,o){pn.point=i,t=r=a,e=n=o},pn.lineEnd=function(){i(t,e)}}var gn={point:function(t,e){t<cn&&(cn=t);t>hn&&(hn=t);e<un&&(un=e);e>fn&&(fn=e)},lineStart:D,lineEnd:D,polygonStart:D,polygonEnd:D};function vn(){var t=mn(4.5),e=[],r={point:n,lineStart:function(){r.point=i},lineEnd:o,polygonStart:function(){r.lineEnd=s},polygonEnd:function(){r.lineEnd=o,r.point=n},pointRadius:function(e){return t=mn(e),r},result:function(){if(e.length){var t=e.join("");return e=[],t}}};function n(r,n){e.push("M",r,",",n,t)}function i(t,n){e.push("M",t,",",n),r.point=a}function a(t,r){e.push("L",t,",",r)}function o(){r.point=n}function s(){e.push("Z")}return r}function mn(t){return"m0,"+t+"a"+t+","+t+" 0 1,1 0,"+-2*t+"a"+t+","+t+" 0 1,1 0,"+2*t+"z"}var yn,xn={point:bn,lineStart:_n,lineEnd:wn,polygonStart:function(){xn.lineStart=kn},polygonEnd:function(){xn.point=bn,xn.lineStart=_n,xn.lineEnd=wn}};function bn(t,e){xr+=t,br+=e,++_r}function _n(){var t,e;function r(r,n){var i=r-t,a=n-e,o=Math.sqrt(i*i+a*a);wr+=o*(t+r)/2,kr+=o*(e+n)/2,Tr+=o,bn(t=r,e=n)}xn.point=function(n,i){xn.point=r,bn(t=n,e=i)}}function wn(){xn.point=bn}function kn(){var t,e,r,n;function i(t,e){var i=t-r,a=e-n,o=Math.sqrt(i*i+a*a);wr+=o*(r+t)/2,kr+=o*(n+e)/2,Tr+=o,Ar+=(o=n*t-r*e)*(r+t),Mr+=o*(n+e),Sr+=3*o,bn(r=t,n=e)}xn.point=function(a,o){xn.point=i,bn(t=r=a,e=n=o)},xn.lineEnd=function(){i(t,e)}}function Tn(t){var e=4.5,r={point:n,lineStart:function(){r.point=i},lineEnd:o,polygonStart:function(){r.lineEnd=s},polygonEnd:function(){r.lineEnd=o,r.point=n},pointRadius:function(t){return e=t,r},result:D};function n(r,n){t.moveTo(r+e,n),t.arc(r,n,e,0,Mt)}function i(e,n){t.moveTo(e,n),r.point=a}function a(e,r){t.lineTo(e,r)}function o(){r.point=n}function s(){t.closePath()}return r}function An(t){var e=.5,r=Math.cos(30*Ct),n=16;function i(e){return(n?function(e){var r,i,o,s,l,c,u,h,f,p,d,g,v={point:m,lineStart:y,lineEnd:b,polygonStart:function(){e.polygonStart(),v.lineStart=_},polygonEnd:function(){e.polygonEnd(),v.lineStart=y}};function m(r,n){r=t(r,n),e.point(r[0],r[1])}function y(){h=NaN,v.point=x,e.lineStart()}function x(r,i){var o=Pr([r,i]),s=t(r,i);a(h,f,u,p,d,g,h=s[0],f=s[1],u=r,p=o[0],d=o[1],g=o[2],n,e),e.point(h,f)}function b(){v.point=m,e.lineEnd()}function _(){y(),v.point=w,v.lineEnd=k}function w(t,e){x(r=t,e),i=h,o=f,s=p,l=d,c=g,v.point=x}function k(){a(h,f,u,p,d,g,i,o,r,s,l,c,n,e),v.lineEnd=b,b()}return v}:function(e){return Sn(e,function(r,n){r=t(r,n),e.point(r[0],r[1])})})(e)}function a(n,i,o,s,l,c,u,h,f,p,d,g,v,m){var x=u-n,b=h-i,_=x*x+b*b;if(_>4*e&&v--){var w=s+p,k=l+d,T=c+g,A=Math.sqrt(w*w+k*k+T*T),M=Math.asin(T/=A),S=y(y(T)-1)<kt||y(o-f)<kt?(o+f)/2:Math.atan2(k,w),E=t(S,M),C=E[0],L=E[1],P=C-n,O=L-i,z=b*P-x*O;(z*z/_>e||y((x*P+b*O)/_-.5)>.3||s*p+l*d+c*g<r)&&(a(n,i,o,s,l,c,C,L,S,w/=A,k/=A,T,v,m),m.point(C,L),a(C,L,S,w,k,T,u,h,f,p,d,g,v,m))}}return i.precision=function(t){return arguments.length?(n=(e=t*t)>0&&16,i):Math.sqrt(e)},i}function Mn(t){this.stream=t}function Sn(t,e){return{point:e,sphere:function(){t.sphere()},lineStart:function(){t.lineStart()},lineEnd:function(){t.lineEnd()},polygonStart:function(){t.polygonStart()},polygonEnd:function(){t.polygonEnd()}}}function En(t){return Cn(function(){return t})()}function Cn(e){var r,n,i,a,o,s,l=An(function(t,e){return[(t=r(t,e))[0]*c+a,o-t[1]*c]}),c=150,u=480,h=250,f=0,p=0,d=0,g=0,v=0,m=tn,x=P,b=null,_=null;function w(t){return[(t=i(t[0]*Ct,t[1]*Ct))[0]*c+a,o-t[1]*c]}function k(t){return(t=i.invert((t[0]-a)/c,(o-t[1])/c))&&[t[0]*Lt,t[1]*Lt]}function T(){i=Gr(n=zn(d,g,v),r);var t=r(f,p);return a=u-t[0]*c,o=h+t[1]*c,A()}function A(){return s&&(s.valid=!1,s=null),w}return w.stream=function(t){return s&&(s.valid=!1),(s=Ln(m(n,l(x(t))))).valid=!0,s},w.clipAngle=function(t){return arguments.length?(m=null==t?(b=t,tn):function(t){var e=Math.cos(t),r=e>0,n=y(e)>kt;return Jr(i,function(t){var e,s,l,c,u;return{lineStart:function(){c=l=!1,u=1},point:function(h,f){var p,d=[h,f],g=i(h,f),v=r?g?0:o(h,f):g?o(h+(h<0?At:-At),f):0;if(!e&&(c=l=g)&&t.lineStart(),g!==l&&(p=a(e,d),(Br(e,p)||Br(d,p))&&(d[0]+=kt,d[1]+=kt,g=i(d[0],d[1]))),g!==l)u=0,g?(t.lineStart(),p=a(d,e),t.point(p[0],p[1])):(p=a(e,d),t.point(p[0],p[1]),t.lineEnd()),e=p;else if(n&&e&&r^g){var m;v&s||!(m=a(d,e,!0))||(u=0,r?(t.lineStart(),t.point(m[0][0],m[0][1]),t.point(m[1][0],m[1][1]),t.lineEnd()):(t.point(m[1][0],m[1][1]),t.lineEnd(),t.lineStart(),t.point(m[0][0],m[0][1])))}!g||e&&Br(e,d)||t.point(d[0],d[1]),e=d,l=g,s=v},lineEnd:function(){l&&t.lineEnd(),e=null},clean:function(){return u|(c&&l)<<1}}},Fn(t,6*Ct),r?[0,-t]:[-At,t-At]);function i(t,r){return Math.cos(t)*Math.cos(r)>e}function a(t,r,n){var i=[1,0,0],a=zr(Pr(t),Pr(r)),o=Or(a,a),s=a[0],l=o-s*s;if(!l)return!n&&t;var c=e*o/l,u=-e*s/l,h=zr(i,a),f=Dr(i,c);Ir(f,Dr(a,u));var p=h,d=Or(f,p),g=Or(p,p),v=d*d-g*(Or(f,f)-1);if(!(v<0)){var m=Math.sqrt(v),x=Dr(p,(-d-m)/g);if(Ir(x,f),x=Fr(x),!n)return x;var b,_=t[0],w=r[0],k=t[1],T=r[1];w<_&&(b=_,_=w,w=b);var A=w-_,M=y(A-At)<kt;if(!M&&T<k&&(b=k,k=T,T=b),M||A<kt?M?k+T>0^x[1]<(y(x[0]-_)<kt?k:T):k<=x[1]&&x[1]<=T:A>At^(_<=x[0]&&x[0]<=w)){var S=Dr(p,(-d+m)/g);return Ir(S,f),[x,Fr(S)]}}}function o(e,n){var i=r?t:At-t,a=0;return e<-i?a|=1:e>i&&(a|=2),n<-i?a|=4:n>i&&(a|=8),a}}((b=+t)*Ct),A()):b},w.clipExtent=function(t){return arguments.length?(_=t,x=t?nn(t[0][0],t[0][1],t[1][0],t[1][1]):P,A()):_},w.scale=function(t){return arguments.length?(c=+t,T()):c},w.translate=function(t){return arguments.length?(u=+t[0],h=+t[1],T()):[u,h]},w.center=function(t){return arguments.length?(f=t[0]%360*Ct,p=t[1]%360*Ct,T()):[f*Lt,p*Lt]},w.rotate=function(t){return arguments.length?(d=t[0]%360*Ct,g=t[1]%360*Ct,v=t.length>2?t[2]%360*Ct:0,T()):[d*Lt,g*Lt,v*Lt]},t.rebind(w,l,"precision"),function(){return r=e.apply(this,arguments),w.invert=r.invert&&k,T()}}function Ln(t){return Sn(t,function(e,r){t.point(e*Ct,r*Ct)})}function Pn(t,e){return[t,e]}function On(t,e){return[t>At?t-Mt:t<-At?t+Mt:t,e]}function zn(t,e,r){return t?e||r?Gr(Dn(t),Rn(e,r)):Dn(t):e||r?Rn(e,r):On}function In(t){return function(e,r){return[(e+=t)>At?e-Mt:e<-At?e+Mt:e,r]}}function Dn(t){var e=In(t);return e.invert=In(-t),e}function Rn(t,e){var r=Math.cos(t),n=Math.sin(t),i=Math.cos(e),a=Math.sin(e);function o(t,e){var o=Math.cos(e),s=Math.cos(t)*o,l=Math.sin(t)*o,c=Math.sin(e),u=c*r+s*n;return[Math.atan2(l*i-u*a,s*r-c*n),It(u*i+l*a)]}return o.invert=function(t,e){var o=Math.cos(e),s=Math.cos(t)*o,l=Math.sin(t)*o,c=Math.sin(e),u=c*i-l*a;return[Math.atan2(l*i+c*a,s*r+u*n),It(u*r-s*n)]},o}function Fn(t,e){var r=Math.cos(t),n=Math.sin(t);return function(i,a,o,s){var l=o*e;null!=i?(i=Bn(r,i),a=Bn(r,a),(o>0?i<a:i>a)&&(i+=o*Mt)):(i=t+o*Mt,a=t-.5*l);for(var c,u=i;o>0?u>a:u<a;u-=l)s.point((c=Fr([r,-n*Math.cos(u),-n*Math.sin(u)]))[0],c[1])}}function Bn(t,e){var r=Pr(e);r[0]-=t,Rr(r);var n=zt(-r[1]);return((-r[2]<0?-n:n)+2*Math.PI-kt)%(2*Math.PI)}function Nn(e,r,n){var i=t.range(e,r-kt,n).concat(r);return function(t){return i.map(function(e){return[t,e]})}}function jn(e,r,n){var i=t.range(e,r-kt,n).concat(r);return function(t){return i.map(function(e){return[e,t]})}}function Un(t){return t.source}function Vn(t){return t.target}t.geo.path=function(){var e,r,n,i,a,o=4.5;function s(e){return e&&("function"==typeof o&&i.pointRadius(+o.apply(this,arguments)),a&&a.valid||(a=n(i)),t.geo.stream(e,a)),i.result()}function l(){return a=null,s}return s.area=function(e){return sn=0,t.geo.stream(e,n(pn)),sn},s.centroid=function(e){return xr=br=_r=wr=kr=Tr=Ar=Mr=Sr=0,t.geo.stream(e,n(xn)),Sr?[Ar/Sr,Mr/Sr]:Tr?[wr/Tr,kr/Tr]:_r?[xr/_r,br/_r]:[NaN,NaN]},s.bounds=function(e){return hn=fn=-(cn=un=1/0),t.geo.stream(e,n(gn)),[[cn,un],[hn,fn]]},s.projection=function(t){return arguments.length?(n=(e=t)?t.stream||(r=t,i=An(function(t,e){return r([t*Lt,e*Lt])}),function(t){return Ln(i(t))}):P,l()):e;var r,i},s.context=function(t){return arguments.length?(i=null==(r=t)?new vn:new Tn(t),"function"!=typeof o&&i.pointRadius(o),l()):r},s.pointRadius=function(t){return arguments.length?(o="function"==typeof t?t:(i.pointRadius(+t),+t),s):o},s.projection(t.geo.albersUsa()).context(null)},t.geo.transform=function(t){return{stream:function(e){var r=new Mn(e);for(var n in t)r[n]=t[n];return r}}},Mn.prototype={point:function(t,e){this.stream.point(t,e)},sphere:function(){this.stream.sphere()},lineStart:function(){this.stream.lineStart()},lineEnd:function(){this.stream.lineEnd()},polygonStart:function(){this.stream.polygonStart()},polygonEnd:function(){this.stream.polygonEnd()}},t.geo.projection=En,t.geo.projectionMutator=Cn,(t.geo.equirectangular=function(){return En(Pn)}).raw=Pn.invert=Pn,t.geo.rotation=function(t){function e(e){return(e=t(e[0]*Ct,e[1]*Ct))[0]*=Lt,e[1]*=Lt,e}return t=zn(t[0]%360*Ct,t[1]*Ct,t.length>2?t[2]*Ct:0),e.invert=function(e){return(e=t.invert(e[0]*Ct,e[1]*Ct))[0]*=Lt,e[1]*=Lt,e},e},On.invert=Pn,t.geo.circle=function(){var t,e,r=[0,0],n=6;function i(){var t="function"==typeof r?r.apply(this,arguments):r,n=zn(-t[0]*Ct,-t[1]*Ct,0).invert,i=[];return e(null,null,1,{point:function(t,e){i.push(t=n(t,e)),t[0]*=Lt,t[1]*=Lt}}),{type:"Polygon",coordinates:[i]}}return i.origin=function(t){return arguments.length?(r=t,i):r},i.angle=function(r){return arguments.length?(e=Fn((t=+r)*Ct,n*Ct),i):t},i.precision=function(r){return arguments.length?(e=Fn(t*Ct,(n=+r)*Ct),i):n},i.angle(90)},t.geo.distance=function(t,e){var r,n=(e[0]-t[0])*Ct,i=t[1]*Ct,a=e[1]*Ct,o=Math.sin(n),s=Math.cos(n),l=Math.sin(i),c=Math.cos(i),u=Math.sin(a),h=Math.cos(a);return Math.atan2(Math.sqrt((r=h*o)*r+(r=c*u-l*h*s)*r),l*u+c*h*s)},t.geo.graticule=function(){var e,r,n,i,a,o,s,l,c,u,h,f,p=10,d=p,g=90,v=360,m=2.5;function x(){return{type:"MultiLineString",coordinates:b()}}function b(){return t.range(Math.ceil(i/g)*g,n,g).map(h).concat(t.range(Math.ceil(l/v)*v,s,v).map(f)).concat(t.range(Math.ceil(r/p)*p,e,p).filter(function(t){return y(t%g)>kt}).map(c)).concat(t.range(Math.ceil(o/d)*d,a,d).filter(function(t){return y(t%v)>kt}).map(u))}return x.lines=function(){return b().map(function(t){return{type:"LineString",coordinates:t}})},x.outline=function(){return{type:"Polygon",coordinates:[h(i).concat(f(s).slice(1),h(n).reverse().slice(1),f(l).reverse().slice(1))]}},x.extent=function(t){return arguments.length?x.majorExtent(t).minorExtent(t):x.minorExtent()},x.majorExtent=function(t){return arguments.length?(i=+t[0][0],n=+t[1][0],l=+t[0][1],s=+t[1][1],i>n&&(t=i,i=n,n=t),l>s&&(t=l,l=s,s=t),x.precision(m)):[[i,l],[n,s]]},x.minorExtent=function(t){return arguments.length?(r=+t[0][0],e=+t[1][0],o=+t[0][1],a=+t[1][1],r>e&&(t=r,r=e,e=t),o>a&&(t=o,o=a,a=t),x.precision(m)):[[r,o],[e,a]]},x.step=function(t){return arguments.length?x.majorStep(t).minorStep(t):x.minorStep()},x.majorStep=function(t){return arguments.length?(g=+t[0],v=+t[1],x):[g,v]},x.minorStep=function(t){return arguments.length?(p=+t[0],d=+t[1],x):[p,d]},x.precision=function(t){return arguments.length?(m=+t,c=Nn(o,a,90),u=jn(r,e,m),h=Nn(l,s,90),f=jn(i,n,m),x):m},x.majorExtent([[-180,-90+kt],[180,90-kt]]).minorExtent([[-180,-80-kt],[180,80+kt]])},t.geo.greatArc=function(){var e,r,n=Un,i=Vn;function a(){return{type:"LineString",coordinates:[e||n.apply(this,arguments),r||i.apply(this,arguments)]}}return a.distance=function(){return t.geo.distance(e||n.apply(this,arguments),r||i.apply(this,arguments))},a.source=function(t){return arguments.length?(n=t,e="function"==typeof t?null:t,a):n},a.target=function(t){return arguments.length?(i=t,r="function"==typeof t?null:t,a):i},a.precision=function(){return arguments.length?a:0},a},t.geo.interpolate=function(t,e){return r=t[0]*Ct,n=t[1]*Ct,i=e[0]*Ct,a=e[1]*Ct,o=Math.cos(n),s=Math.sin(n),l=Math.cos(a),c=Math.sin(a),u=o*Math.cos(r),h=o*Math.sin(r),f=l*Math.cos(i),p=l*Math.sin(i),d=2*Math.asin(Math.sqrt(Rt(a-n)+o*l*Rt(i-r))),g=1/Math.sin(d),(v=d?function(t){var e=Math.sin(t*=d)*g,r=Math.sin(d-t)*g,n=r*u+e*f,i=r*h+e*p,a=r*s+e*c;return[Math.atan2(i,n)*Lt,Math.atan2(a,Math.sqrt(n*n+i*i))*Lt]}:function(){return[r*Lt,n*Lt]}).distance=d,v;var r,n,i,a,o,s,l,c,u,h,f,p,d,g,v},t.geo.length=function(e){return yn=0,t.geo.stream(e,qn),yn};var qn={sphere:D,point:D,lineStart:function(){var t,e,r;function n(n,i){var a=Math.sin(i*=Ct),o=Math.cos(i),s=y((n*=Ct)-t),l=Math.cos(s);yn+=Math.atan2(Math.sqrt((s=o*Math.sin(s))*s+(s=r*a-e*o*l)*s),e*a+r*o*l),t=n,e=a,r=o}qn.point=function(i,a){t=i*Ct,e=Math.sin(a*=Ct),r=Math.cos(a),qn.point=n},qn.lineEnd=function(){qn.point=qn.lineEnd=D}},lineEnd:D,polygonStart:D,polygonEnd:D};function Hn(t,e){function r(e,r){var n=Math.cos(e),i=Math.cos(r),a=t(n*i);return[a*i*Math.sin(e),a*Math.sin(r)]}return r.invert=function(t,r){var n=Math.sqrt(t*t+r*r),i=e(n),a=Math.sin(i),o=Math.cos(i);return[Math.atan2(t*a,n*o),Math.asin(n&&r*a/n)]},r}var Gn=Hn(function(t){return Math.sqrt(2/(1+t))},function(t){return 2*Math.asin(t/2)});(t.geo.azimuthalEqualArea=function(){return En(Gn)}).raw=Gn;var Yn=Hn(function(t){var e=Math.acos(t);return e&&e/Math.sin(e)},P);function Wn(t,e){var r=Math.cos(t),n=function(t){return Math.tan(At/4+t/2)},i=t===e?Math.sin(t):Math.log(r/Math.cos(e))/Math.log(n(e)/n(t)),a=r*Math.pow(n(t),i)/i;if(!i)return Jn;function o(t,e){a>0?e<-Et+kt&&(e=-Et+kt):e>Et-kt&&(e=Et-kt);var r=a/Math.pow(n(e),i);return[r*Math.sin(i*t),a-r*Math.cos(i*t)]}return o.invert=function(t,e){var r=a-e,n=Pt(i)*Math.sqrt(t*t+r*r);return[Math.atan2(t,r)/i,2*Math.atan(Math.pow(a/n,1/i))-Et]},o}function Xn(t,e){var r=Math.cos(t),n=t===e?Math.sin(t):(r-Math.cos(e))/(e-t),i=r/n+t;if(y(n)<kt)return Pn;function a(t,e){var r=i-e;return[r*Math.sin(n*t),i-r*Math.cos(n*t)]}return a.invert=function(t,e){var r=i-e;return[Math.atan2(t,r)/n,i-Pt(n)*Math.sqrt(t*t+r*r)]},a}(t.geo.azimuthalEquidistant=function(){return En(Yn)}).raw=Yn,(t.geo.conicConformal=function(){return an(Wn)}).raw=Wn,(t.geo.conicEquidistant=function(){return an(Xn)}).raw=Xn;var Zn=Hn(function(t){return 1/t},Math.atan);function Jn(t,e){return[t,Math.log(Math.tan(At/4+e/2))]}function Kn(t){var e,r=En(t),n=r.scale,i=r.translate,a=r.clipExtent;return r.scale=function(){var t=n.apply(r,arguments);return t===r?e?r.clipExtent(null):r:t},r.translate=function(){var t=i.apply(r,arguments);return t===r?e?r.clipExtent(null):r:t},r.clipExtent=function(t){var o=a.apply(r,arguments);if(o===r){if(e=null==t){var s=At*n(),l=i();a([[l[0]-s,l[1]-s],[l[0]+s,l[1]+s]])}}else e&&(o=null);return o},r.clipExtent(null)}(t.geo.gnomonic=function(){return En(Zn)}).raw=Zn,Jn.invert=function(t,e){return[t,2*Math.atan(Math.exp(e))-Et]},(t.geo.mercator=function(){return Kn(Jn)}).raw=Jn;var $n=Hn(function(){return 1},Math.asin);(t.geo.orthographic=function(){return En($n)}).raw=$n;var Qn=Hn(function(t){return 1/(1+t)},function(t){return 2*Math.atan(t)});function ti(t,e){return[Math.log(Math.tan(At/4+e/2)),-t]}function ei(t){return t[0]}function ri(t){return t[1]}function ni(t){for(var e=t.length,r=[0,1],n=2,i=2;i<e;i++){for(;n>1&&Ot(t[r[n-2]],t[r[n-1]],t[i])<=0;)--n;r[n++]=i}return r.slice(0,n)}function ii(t,e){return t[0]-e[0]||t[1]-e[1]}(t.geo.stereographic=function(){return En(Qn)}).raw=Qn,ti.invert=function(t,e){return[-e,2*Math.atan(Math.exp(t))-Et]},(t.geo.transverseMercator=function(){var t=Kn(ti),e=t.center,r=t.rotate;return t.center=function(t){return t?e([-t[1],t[0]]):[(t=e())[1],-t[0]]},t.rotate=function(t){return t?r([t[0],t[1],t.length>2?t[2]+90:90]):[(t=r())[0],t[1],t[2]-90]},r([0,0,90])}).raw=ti,t.geom={},t.geom.hull=function(t){var e=ei,r=ri;if(arguments.length)return n(t);function n(t){if(t.length<3)return[];var n,i=ve(e),a=ve(r),o=t.length,s=[],l=[];for(n=0;n<o;n++)s.push([+i.call(this,t[n],n),+a.call(this,t[n],n),n]);for(s.sort(ii),n=0;n<o;n++)l.push([s[n][0],-s[n][1]]);var c=ni(s),u=ni(l),h=u[0]===c[0],f=u[u.length-1]===c[c.length-1],p=[];for(n=c.length-1;n>=0;--n)p.push(t[s[c[n]][2]]);for(n=+h;n<u.length-f;++n)p.push(t[s[u[n]][2]]);return p}return n.x=function(t){return arguments.length?(e=t,n):e},n.y=function(t){return arguments.length?(r=t,n):r},n},t.geom.polygon=function(t){return V(t,ai),t};var ai=t.geom.polygon.prototype=[];function oi(t,e,r){return(r[0]-e[0])*(t[1]-e[1])<(r[1]-e[1])*(t[0]-e[0])}function si(t,e,r,n){var i=t[0],a=r[0],o=e[0]-i,s=n[0]-a,l=t[1],c=r[1],u=e[1]-l,h=n[1]-c,f=(s*(l-c)-h*(i-a))/(h*o-s*u);return[i+f*o,l+f*u]}function li(t){var e=t[0],r=t[t.length-1];return!(e[0]-r[0]||e[1]-r[1])}ai.area=function(){for(var t,e=-1,r=this.length,n=this[r-1],i=0;++e<r;)t=n,n=this[e],i+=t[1]*n[0]-t[0]*n[1];return.5*i},ai.centroid=function(t){var e,r,n=-1,i=this.length,a=0,o=0,s=this[i-1];for(arguments.length||(t=-1/(6*this.area()));++n<i;)e=s,s=this[n],r=e[0]*s[1]-s[0]*e[1],a+=(e[0]+s[0])*r,o+=(e[1]+s[1])*r;return[a*t,o*t]},ai.clip=function(t){for(var e,r,n,i,a,o,s=li(t),l=-1,c=this.length-li(this),u=this[c-1];++l<c;){for(e=t.slice(),t.length=0,i=this[l],a=e[(n=e.length-s)-1],r=-1;++r<n;)oi(o=e[r],u,i)?(oi(a,u,i)||t.push(si(a,o,u,i)),t.push(o)):oi(a,u,i)&&t.push(si(a,o,u,i)),a=o;s&&t.push(t[0]),u=i}return t};var ci,ui,hi,fi,pi,di=[],gi=[];function vi(){Ii(this),this.edge=this.site=this.circle=null}function mi(t){var e=di.pop()||new vi;return e.site=t,e}function yi(t){Si(t),hi.remove(t),di.push(t),Ii(t)}function xi(t){var e=t.circle,r=e.x,n=e.cy,i={x:r,y:n},a=t.P,o=t.N,s=[t];yi(t);for(var l=a;l.circle&&y(r-l.circle.x)<kt&&y(n-l.circle.cy)<kt;)a=l.P,s.unshift(l),yi(l),l=a;s.unshift(l),Si(l);for(var c=o;c.circle&&y(r-c.circle.x)<kt&&y(n-c.circle.cy)<kt;)o=c.N,s.push(c),yi(c),c=o;s.push(c),Si(c);var u,h=s.length;for(u=1;u<h;++u)c=s[u],l=s[u-1],Pi(c.edge,l.site,c.site,i);l=s[0],(c=s[h-1]).edge=Li(l.site,c.site,null,i),Mi(l),Mi(c)}function bi(t){for(var e,r,n,i,a=t.x,o=t.y,s=hi._;s;)if((n=_i(s,o)-a)>kt)s=s.L;else{if(!((i=a-wi(s,o))>kt)){n>-kt?(e=s.P,r=s):i>-kt?(e=s,r=s.N):e=r=s;break}if(!s.R){e=s;break}s=s.R}var l=mi(t);if(hi.insert(e,l),e||r){if(e===r)return Si(e),r=mi(e.site),hi.insert(l,r),l.edge=r.edge=Li(e.site,l.site),Mi(e),void Mi(r);if(r){Si(e),Si(r);var c=e.site,u=c.x,h=c.y,f=t.x-u,p=t.y-h,d=r.site,g=d.x-u,v=d.y-h,m=2*(f*v-p*g),y=f*f+p*p,x=g*g+v*v,b={x:(v*y-p*x)/m+u,y:(f*x-g*y)/m+h};Pi(r.edge,c,d,b),l.edge=Li(c,t,null,b),r.edge=Li(t,d,null,b),Mi(e),Mi(r)}else l.edge=Li(e.site,l.site)}}function _i(t,e){var r=t.site,n=r.x,i=r.y,a=i-e;if(!a)return n;var o=t.P;if(!o)return-1/0;var s=(r=o.site).x,l=r.y,c=l-e;if(!c)return s;var u=s-n,h=1/a-1/c,f=u/c;return h?(-f+Math.sqrt(f*f-2*h*(u*u/(-2*c)-l+c/2+i-a/2)))/h+n:(n+s)/2}function wi(t,e){var r=t.N;if(r)return _i(r,e);var n=t.site;return n.y===e?n.x:1/0}function ki(t){this.site=t,this.edges=[]}function Ti(t,e){return e.angle-t.angle}function Ai(){Ii(this),this.x=this.y=this.arc=this.site=this.cy=null}function Mi(t){var e=t.P,r=t.N;if(e&&r){var n=e.site,i=t.site,a=r.site;if(n!==a){var o=i.x,s=i.y,l=n.x-o,c=n.y-s,u=a.x-o,h=2*(l*(v=a.y-s)-c*u);if(!(h>=-Tt)){var f=l*l+c*c,p=u*u+v*v,d=(v*f-c*p)/h,g=(l*p-u*f)/h,v=g+s,m=gi.pop()||new Ai;m.arc=t,m.site=i,m.x=d+o,m.y=v+Math.sqrt(d*d+g*g),m.cy=v,t.circle=m;for(var y=null,x=pi._;x;)if(m.y<x.y||m.y===x.y&&m.x<=x.x){if(!x.L){y=x.P;break}x=x.L}else{if(!x.R){y=x;break}x=x.R}pi.insert(y,m),y||(fi=m)}}}}function Si(t){var e=t.circle;e&&(e.P||(fi=e.N),pi.remove(e),gi.push(e),Ii(e),t.circle=null)}function Ei(t,e){var r=t.b;if(r)return!0;var n,i,a=t.a,o=e[0][0],s=e[1][0],l=e[0][1],c=e[1][1],u=t.l,h=t.r,f=u.x,p=u.y,d=h.x,g=h.y,v=(f+d)/2,m=(p+g)/2;if(g===p){if(v<o||v>=s)return;if(f>d){if(a){if(a.y>=c)return}else a={x:v,y:l};r={x:v,y:c}}else{if(a){if(a.y<l)return}else a={x:v,y:c};r={x:v,y:l}}}else if(i=m-(n=(f-d)/(g-p))*v,n<-1||n>1)if(f>d){if(a){if(a.y>=c)return}else a={x:(l-i)/n,y:l};r={x:(c-i)/n,y:c}}else{if(a){if(a.y<l)return}else a={x:(c-i)/n,y:c};r={x:(l-i)/n,y:l}}else if(p<g){if(a){if(a.x>=s)return}else a={x:o,y:n*o+i};r={x:s,y:n*s+i}}else{if(a){if(a.x<o)return}else a={x:s,y:n*s+i};r={x:o,y:n*o+i}}return t.a=a,t.b=r,!0}function Ci(t,e){this.l=t,this.r=e,this.a=this.b=null}function Li(t,e,r,n){var i=new Ci(t,e);return ci.push(i),r&&Pi(i,t,e,r),n&&Pi(i,e,t,n),ui[t.i].edges.push(new Oi(i,t,e)),ui[e.i].edges.push(new Oi(i,e,t)),i}function Pi(t,e,r,n){t.a||t.b?t.l===r?t.b=n:t.a=n:(t.a=n,t.l=e,t.r=r)}function Oi(t,e,r){var n=t.a,i=t.b;this.edge=t,this.site=e,this.angle=r?Math.atan2(r.y-e.y,r.x-e.x):t.l===e?Math.atan2(i.x-n.x,n.y-i.y):Math.atan2(n.x-i.x,i.y-n.y)}function zi(){this._=null}function Ii(t){t.U=t.C=t.L=t.R=t.P=t.N=null}function Di(t,e){var r=e,n=e.R,i=r.U;i?i.L===r?i.L=n:i.R=n:t._=n,n.U=i,r.U=n,r.R=n.L,r.R&&(r.R.U=r),n.L=r}function Ri(t,e){var r=e,n=e.L,i=r.U;i?i.L===r?i.L=n:i.R=n:t._=n,n.U=i,r.U=n,r.L=n.R,r.L&&(r.L.U=r),n.R=r}function Fi(t){for(;t.L;)t=t.L;return t}function Bi(t,e){var r,n,i,a=t.sort(Ni).pop();for(ci=[],ui=new Array(t.length),hi=new zi,pi=new zi;;)if(i=fi,a&&(!i||a.y<i.y||a.y===i.y&&a.x<i.x))a.x===r&&a.y===n||(ui[a.i]=new ki(a),bi(a),r=a.x,n=a.y),a=t.pop();else{if(!i)break;xi(i.arc)}e&&(function(t){for(var e,r=ci,n=en(t[0][0],t[0][1],t[1][0],t[1][1]),i=r.length;i--;)(!Ei(e=r[i],t)||!n(e)||y(e.a.x-e.b.x)<kt&&y(e.a.y-e.b.y)<kt)&&(e.a=e.b=null,r.splice(i,1))}(e),function(t){for(var e,r,n,i,a,o,s,l,c,u,h=t[0][0],f=t[1][0],p=t[0][1],d=t[1][1],g=ui,v=g.length;v--;)if((a=g[v])&&a.prepare())for(l=(s=a.edges).length,o=0;o<l;)n=(u=s[o].end()).x,i=u.y,e=(c=s[++o%l].start()).x,r=c.y,(y(n-e)>kt||y(i-r)>kt)&&(s.splice(o,0,new Oi((m=a.site,x=u,b=y(n-h)<kt&&d-i>kt?{x:h,y:y(e-h)<kt?r:d}:y(i-d)<kt&&f-n>kt?{x:y(r-d)<kt?e:f,y:d}:y(n-f)<kt&&i-p>kt?{x:f,y:y(e-f)<kt?r:p}:y(i-p)<kt&&n-h>kt?{x:y(r-p)<kt?e:h,y:p}:null,_=void 0,_=new Ci(m,null),_.a=x,_.b=b,ci.push(_),_),a.site,null)),++l);var m,x,b,_}(e));var o={cells:ui,edges:ci};return hi=pi=ci=ui=null,o}function Ni(t,e){return e.y-t.y||e.x-t.x}ki.prototype.prepare=function(){for(var t,e=this.edges,r=e.length;r--;)(t=e[r].edge).b&&t.a||e.splice(r,1);return e.sort(Ti),e.length},Oi.prototype={start:function(){return this.edge.l===this.site?this.edge.a:this.edge.b},end:function(){return this.edge.l===this.site?this.edge.b:this.edge.a}},zi.prototype={insert:function(t,e){var r,n,i;if(t){if(e.P=t,e.N=t.N,t.N&&(t.N.P=e),t.N=e,t.R){for(t=t.R;t.L;)t=t.L;t.L=e}else t.R=e;r=t}else this._?(t=Fi(this._),e.P=null,e.N=t,t.P=t.L=e,r=t):(e.P=e.N=null,this._=e,r=null);for(e.L=e.R=null,e.U=r,e.C=!0,t=e;r&&r.C;)r===(n=r.U).L?(i=n.R)&&i.C?(r.C=i.C=!1,n.C=!0,t=n):(t===r.R&&(Di(this,r),r=(t=r).U),r.C=!1,n.C=!0,Ri(this,n)):(i=n.L)&&i.C?(r.C=i.C=!1,n.C=!0,t=n):(t===r.L&&(Ri(this,r),r=(t=r).U),r.C=!1,n.C=!0,Di(this,n)),r=t.U;this._.C=!1},remove:function(t){t.N&&(t.N.P=t.P),t.P&&(t.P.N=t.N),t.N=t.P=null;var e,r,n,i=t.U,a=t.L,o=t.R;if(r=a?o?Fi(o):a:o,i?i.L===t?i.L=r:i.R=r:this._=r,a&&o?(n=r.C,r.C=t.C,r.L=a,a.U=r,r!==o?(i=r.U,r.U=t.U,t=r.R,i.L=t,r.R=o,o.U=r):(r.U=i,i=r,t=r.R)):(n=t.C,t=r),t&&(t.U=i),!n)if(t&&t.C)t.C=!1;else{do{if(t===this._)break;if(t===i.L){if((e=i.R).C&&(e.C=!1,i.C=!0,Di(this,i),e=i.R),e.L&&e.L.C||e.R&&e.R.C){e.R&&e.R.C||(e.L.C=!1,e.C=!0,Ri(this,e),e=i.R),e.C=i.C,i.C=e.R.C=!1,Di(this,i),t=this._;break}}else if((e=i.L).C&&(e.C=!1,i.C=!0,Ri(this,i),e=i.L),e.L&&e.L.C||e.R&&e.R.C){e.L&&e.L.C||(e.R.C=!1,e.C=!0,Di(this,e),e=i.L),e.C=i.C,i.C=e.L.C=!1,Ri(this,i),t=this._;break}e.C=!0,t=i,i=i.U}while(!t.C);t&&(t.C=!1)}}},t.geom.voronoi=function(t){var e=ei,r=ri,n=e,i=r,a=ji;if(t)return o(t);function o(t){var e=new Array(t.length),r=a[0][0],n=a[0][1],i=a[1][0],o=a[1][1];return Bi(s(t),a).cells.forEach(function(a,s){var l=a.edges,c=a.site;(e[s]=l.length?l.map(function(t){var e=t.start();return[e.x,e.y]}):c.x>=r&&c.x<=i&&c.y>=n&&c.y<=o?[[r,o],[i,o],[i,n],[r,n]]:[]).point=t[s]}),e}function s(t){return t.map(function(t,e){return{x:Math.round(n(t,e)/kt)*kt,y:Math.round(i(t,e)/kt)*kt,i:e}})}return o.links=function(t){return Bi(s(t)).edges.filter(function(t){return t.l&&t.r}).map(function(e){return{source:t[e.l.i],target:t[e.r.i]}})},o.triangles=function(t){var e=[];return Bi(s(t)).cells.forEach(function(r,n){for(var i,a,o,s,l=r.site,c=r.edges.sort(Ti),u=-1,h=c.length,f=c[h-1].edge,p=f.l===l?f.r:f.l;++u<h;)f,i=p,p=(f=c[u].edge).l===l?f.r:f.l,n<i.i&&n<p.i&&(o=i,s=p,((a=l).x-s.x)*(o.y-a.y)-(a.x-o.x)*(s.y-a.y)<0)&&e.push([t[n],t[i.i],t[p.i]])}),e},o.x=function(t){return arguments.length?(n=ve(e=t),o):e},o.y=function(t){return arguments.length?(i=ve(r=t),o):r},o.clipExtent=function(t){return arguments.length?(a=null==t?ji:t,o):a===ji?null:a},o.size=function(t){return arguments.length?o.clipExtent(t&&[[0,0],t]):a===ji?null:a&&a[1]},o};var ji=[[-1e6,-1e6],[1e6,1e6]];function Ui(t){return t.x}function Vi(t){return t.y}function qi(e,r){e=t.rgb(e),r=t.rgb(r);var n=e.r,i=e.g,a=e.b,o=r.r-n,s=r.g-i,l=r.b-a;return function(t){return"#"+ce(Math.round(n+o*t))+ce(Math.round(i+s*t))+ce(Math.round(a+l*t))}}function Hi(t,e){var r,n={},i={};for(r in t)r in e?n[r]=Zi(t[r],e[r]):i[r]=t[r];for(r in e)r in t||(i[r]=e[r]);return function(t){for(r in n)i[r]=n[r](t);return i}}function Gi(t,e){return t=+t,e=+e,function(r){return t*(1-r)+e*r}}function Yi(t,e){var r,n,i,a=Wi.lastIndex=Xi.lastIndex=0,o=-1,s=[],l=[];for(t+="",e+="";(r=Wi.exec(t))&&(n=Xi.exec(e));)(i=n.index)>a&&(i=e.slice(a,i),s[o]?s[o]+=i:s[++o]=i),(r=r[0])===(n=n[0])?s[o]?s[o]+=n:s[++o]=n:(s[++o]=null,l.push({i:o,x:Gi(r,n)})),a=Xi.lastIndex;return a<e.length&&(i=e.slice(a),s[o]?s[o]+=i:s[++o]=i),s.length<2?l[0]?(e=l[0].x,function(t){return e(t)+""}):function(){return e}:(e=l.length,function(t){for(var r,n=0;n<e;++n)s[(r=l[n]).i]=r.x(t);return s.join("")})}t.geom.delaunay=function(e){return t.geom.voronoi().triangles(e)},t.geom.quadtree=function(t,e,r,n,i){var a,o=ei,s=ri;if(a=arguments.length)return o=Ui,s=Vi,3===a&&(i=r,n=e,r=e=0),l(t);function l(t){var l,c,u,h,f,p,d,g,v,m=ve(o),x=ve(s);if(null!=e)p=e,d=r,g=n,v=i;else if(g=v=-(p=d=1/0),c=[],u=[],f=t.length,a)for(h=0;h<f;++h)(l=t[h]).x<p&&(p=l.x),l.y<d&&(d=l.y),l.x>g&&(g=l.x),l.y>v&&(v=l.y),c.push(l.x),u.push(l.y);else for(h=0;h<f;++h){var b=+m(l=t[h],h),_=+x(l,h);b<p&&(p=b),_<d&&(d=_),b>g&&(g=b),_>v&&(v=_),c.push(b),u.push(_)}var w=g-p,k=v-d;function T(t,e,r,n,i,a,o,s){if(!isNaN(r)&&!isNaN(n))if(t.leaf){var l=t.x,c=t.y;if(null!=l)if(y(l-r)+y(c-n)<.01)A(t,e,r,n,i,a,o,s);else{var u=t.point;t.x=t.y=t.point=null,A(t,u,l,c,i,a,o,s),A(t,e,r,n,i,a,o,s)}else t.x=r,t.y=n,t.point=e}else A(t,e,r,n,i,a,o,s)}function A(t,e,r,n,i,a,o,s){var l=.5*(i+o),c=.5*(a+s),u=r>=l,h=n>=c,f=h<<1|u;t.leaf=!1,u?i=l:o=l,h?a=c:s=c,T(t=t.nodes[f]||(t.nodes[f]={leaf:!0,nodes:[],point:null,x:null,y:null,add:function(t){T(M,t,+m(t,++h),+x(t,h),p,d,g,v)}}),e,r,n,i,a,o,s)}w>k?v=d+w:g=p+k;var M={leaf:!0,nodes:[],point:null,x:null,y:null,add:function(t){T(M,t,+m(t,++h),+x(t,h),p,d,g,v)}};if(M.visit=function(t){!function t(e,r,n,i,a,o){if(!e(r,n,i,a,o)){var s=.5*(n+a),l=.5*(i+o),c=r.nodes;c[0]&&t(e,c[0],n,i,s,l),c[1]&&t(e,c[1],s,i,a,l),c[2]&&t(e,c[2],n,l,s,o),c[3]&&t(e,c[3],s,l,a,o)}}(t,M,p,d,g,v)},M.find=function(t){return function(t,e,r,n,i,a,o){var s,l=1/0;return function t(c,u,h,f,p){if(!(u>a||h>o||f<n||p<i)){if(d=c.point){var d,g=e-c.x,v=r-c.y,m=g*g+v*v;if(m<l){var y=Math.sqrt(l=m);n=e-y,i=r-y,a=e+y,o=r+y,s=d}}for(var x=c.nodes,b=.5*(u+f),_=.5*(h+p),w=(r>=_)<<1|e>=b,k=w+4;w<k;++w)if(c=x[3&w])switch(3&w){case 0:t(c,u,h,b,_);break;case 1:t(c,b,h,f,_);break;case 2:t(c,u,_,b,p);break;case 3:t(c,b,_,f,p)}}}(t,n,i,a,o),s}(M,t[0],t[1],p,d,g,v)},h=-1,null==e){for(;++h<f;)T(M,t[h],c[h],u[h],p,d,g,v);--h}else t.forEach(M.add);return c=u=t=l=null,M}return l.x=function(t){return arguments.length?(o=t,l):o},l.y=function(t){return arguments.length?(s=t,l):s},l.extent=function(t){return arguments.length?(null==t?e=r=n=i=null:(e=+t[0][0],r=+t[0][1],n=+t[1][0],i=+t[1][1]),l):null==e?null:[[e,r],[n,i]]},l.size=function(t){return arguments.length?(null==t?e=r=n=i=null:(e=r=0,n=+t[0],i=+t[1]),l):null==e?null:[n-e,i-r]},l},t.interpolateRgb=qi,t.interpolateObject=Hi,t.interpolateNumber=Gi,t.interpolateString=Yi;var Wi=/[-+]?(?:\d+\.?\d*|\.?\d+)(?:[eE][-+]?\d+)?/g,Xi=new RegExp(Wi.source,"g");function Zi(e,r){for(var n,i=t.interpolators.length;--i>=0&&!(n=t.interpolators[i](e,r)););return n}function Ji(t,e){var r,n=[],i=[],a=t.length,o=e.length,s=Math.min(t.length,e.length);for(r=0;r<s;++r)n.push(Zi(t[r],e[r]));for(;r<a;++r)i[r]=t[r];for(;r<o;++r)i[r]=e[r];return function(t){for(r=0;r<s;++r)i[r]=n[r](t);return i}}t.interpolate=Zi,t.interpolators=[function(t,e){var r=typeof e;return("string"===r?ge.has(e.toLowerCase())||/^(#|rgb\(|hsl\()/i.test(e)?qi:Yi:e instanceof Ut?qi:Array.isArray(e)?Ji:"object"===r&&isNaN(e)?Hi:Gi)(t,e)}],t.interpolateArray=Ji;var Ki=function(){return P},$i=t.map({linear:Ki,poly:function(t){return function(e){return Math.pow(e,t)}},quad:function(){return ra},cubic:function(){return na},sin:function(){return aa},exp:function(){return oa},circle:function(){return sa},elastic:function(t,e){var r;arguments.length<2&&(e=.45);arguments.length?r=e/Mt*Math.asin(1/t):(t=1,r=e/4);return function(n){return 1+t*Math.pow(2,-10*n)*Math.sin((n-r)*Mt/e)}},back:function(t){t||(t=1.70158);return function(e){return e*e*((t+1)*e-t)}},bounce:function(){return la}}),Qi=t.map({in:P,out:ta,"in-out":ea,"out-in":function(t){return ea(ta(t))}});function ta(t){return function(e){return 1-t(1-e)}}function ea(t){return function(e){return.5*(e<.5?t(2*e):2-t(2-2*e))}}function ra(t){return t*t}function na(t){return t*t*t}function ia(t){if(t<=0)return 0;if(t>=1)return 1;var e=t*t,r=e*t;return 4*(t<.5?r:3*(t-e)+r-.75)}function aa(t){return 1-Math.cos(t*Et)}function oa(t){return Math.pow(2,10*(t-1))}function sa(t){return 1-Math.sqrt(1-t*t)}function la(t){return t<1/2.75?7.5625*t*t:t<2/2.75?7.5625*(t-=1.5/2.75)*t+.75:t<2.5/2.75?7.5625*(t-=2.25/2.75)*t+.9375:7.5625*(t-=2.625/2.75)*t+.984375}function ca(t,e){return e-=t,function(r){return Math.round(t+e*r)}}function ua(t){var e,r,n,i=[t.a,t.b],a=[t.c,t.d],o=fa(i),s=ha(i,a),l=fa(((e=a)[0]+=(n=-s)*(r=i)[0],e[1]+=n*r[1],e))||0;i[0]*a[1]<a[0]*i[1]&&(i[0]*=-1,i[1]*=-1,o*=-1,s*=-1),this.rotate=(o?Math.atan2(i[1],i[0]):Math.atan2(-a[0],a[1]))*Lt,this.translate=[t.e,t.f],this.scale=[o,l],this.skew=l?Math.atan2(s,l)*Lt:0}function ha(t,e){return t[0]*e[0]+t[1]*e[1]}function fa(t){var e=Math.sqrt(ha(t,t));return e&&(t[0]/=e,t[1]/=e),e}t.ease=function(t){var e,n=t.indexOf("-"),i=n>=0?t.slice(0,n):t,a=n>=0?t.slice(n+1):"in";return i=$i.get(i)||Ki,a=Qi.get(a)||P,e=a(i.apply(null,r.call(arguments,1))),function(t){return t<=0?0:t>=1?1:e(t)}},t.interpolateHcl=function(e,r){e=t.hcl(e),r=t.hcl(r);var n=e.h,i=e.c,a=e.l,o=r.h-n,s=r.c-i,l=r.l-a;isNaN(s)&&(s=0,i=isNaN(i)?r.c:i);isNaN(o)?(o=0,n=isNaN(n)?r.h:n):o>180?o-=360:o<-180&&(o+=360);return function(t){return Wt(n+o*t,i+s*t,a+l*t)+""}},t.interpolateHsl=function(e,r){e=t.hsl(e),r=t.hsl(r);var n=e.h,i=e.s,a=e.l,o=r.h-n,s=r.s-i,l=r.l-a;isNaN(s)&&(s=0,i=isNaN(i)?r.s:i);isNaN(o)?(o=0,n=isNaN(n)?r.h:n):o>180?o-=360:o<-180&&(o+=360);return function(t){return Ht(n+o*t,i+s*t,a+l*t)+""}},t.interpolateLab=function(e,r){e=t.lab(e),r=t.lab(r);var n=e.l,i=e.a,a=e.b,o=r.l-n,s=r.a-i,l=r.b-a;return function(t){return te(n+o*t,i+s*t,a+l*t)+""}},t.interpolateRound=ca,t.transform=function(e){var r=i.createElementNS(t.ns.prefix.svg,"g");return(t.transform=function(t){if(null!=t){r.setAttribute("transform",t);var e=r.transform.baseVal.consolidate()}return new ua(e?e.matrix:pa)})(e)},ua.prototype.toString=function(){return"translate("+this.translate+")rotate("+this.rotate+")skewX("+this.skew+")scale("+this.scale+")"};var pa={a:1,b:0,c:0,d:1,e:0,f:0};function da(t){return t.length?t.pop()+",":""}function ga(e,r){var n=[],i=[];return e=t.transform(e),r=t.transform(r),function(t,e,r,n){if(t[0]!==e[0]||t[1]!==e[1]){var i=r.push("translate(",null,",",null,")");n.push({i:i-4,x:Gi(t[0],e[0])},{i:i-2,x:Gi(t[1],e[1])})}else(e[0]||e[1])&&r.push("translate("+e+")")}(e.translate,r.translate,n,i),function(t,e,r,n){t!==e?(t-e>180?e+=360:e-t>180&&(t+=360),n.push({i:r.push(da(r)+"rotate(",null,")")-2,x:Gi(t,e)})):e&&r.push(da(r)+"rotate("+e+")")}(e.rotate,r.rotate,n,i),function(t,e,r,n){t!==e?n.push({i:r.push(da(r)+"skewX(",null,")")-2,x:Gi(t,e)}):e&&r.push(da(r)+"skewX("+e+")")}(e.skew,r.skew,n,i),function(t,e,r,n){if(t[0]!==e[0]||t[1]!==e[1]){var i=r.push(da(r)+"scale(",null,",",null,")");n.push({i:i-4,x:Gi(t[0],e[0])},{i:i-2,x:Gi(t[1],e[1])})}else 1===e[0]&&1===e[1]||r.push(da(r)+"scale("+e+")")}(e.scale,r.scale,n,i),e=r=null,function(t){for(var e,r=-1,a=i.length;++r<a;)n[(e=i[r]).i]=e.x(t);return n.join("")}}function va(t,e){return e=(e-=t=+t)||1/e,function(r){return(r-t)/e}}function ma(t,e){return e=(e-=t=+t)||1/e,function(r){return Math.max(0,Math.min(1,(r-t)/e))}}function ya(t){for(var e=t.source,r=t.target,n=function(t,e){if(t===e)return t;var r=xa(t),n=xa(e),i=r.pop(),a=n.pop(),o=null;for(;i===a;)o=i,i=r.pop(),a=n.pop();return o}(e,r),i=[e];e!==n;)e=e.parent,i.push(e);for(var a=i.length;r!==n;)i.splice(a,0,r),r=r.parent;return i}function xa(t){for(var e=[],r=t.parent;null!=r;)e.push(t),t=r,r=r.parent;return e.push(t),e}function ba(t){t.fixed|=2}function _a(t){t.fixed&=-7}function wa(t){t.fixed|=4,t.px=t.x,t.py=t.y}function ka(t){t.fixed&=-5}t.interpolateTransform=ga,t.layout={},t.layout.bundle=function(){return function(t){for(var e=[],r=-1,n=t.length;++r<n;)e.push(ya(t[r]));return e}},t.layout.chord=function(){var e,r,n,i,a,o,s,l={},c=0;function u(){var l,u,f,p,d,g={},v=[],m=t.range(i),y=[];for(e=[],r=[],l=0,p=-1;++p<i;){for(u=0,d=-1;++d<i;)u+=n[p][d];v.push(u),y.push(t.range(i)),l+=u}for(a&&m.sort(function(t,e){return a(v[t],v[e])}),o&&y.forEach(function(t,e){t.sort(function(t,r){return o(n[e][t],n[e][r])})}),l=(Mt-c*i)/l,u=0,p=-1;++p<i;){for(f=u,d=-1;++d<i;){var x=m[p],b=y[x][d],_=n[x][b],w=u,k=u+=_*l;g[x+"-"+b]={index:x,subindex:b,startAngle:w,endAngle:k,value:_}}r[x]={index:x,startAngle:f,endAngle:u,value:v[x]},u+=c}for(p=-1;++p<i;)for(d=p-1;++d<i;){var T=g[p+"-"+d],A=g[d+"-"+p];(T.value||A.value)&&e.push(T.value<A.value?{source:A,target:T}:{source:T,target:A})}s&&h()}function h(){e.sort(function(t,e){return s((t.source.value+t.target.value)/2,(e.source.value+e.target.value)/2)})}return l.matrix=function(t){return arguments.length?(i=(n=t)&&n.length,e=r=null,l):n},l.padding=function(t){return arguments.length?(c=t,e=r=null,l):c},l.sortGroups=function(t){return arguments.length?(a=t,e=r=null,l):a},l.sortSubgroups=function(t){return arguments.length?(o=t,e=null,l):o},l.sortChords=function(t){return arguments.length?(s=t,e&&h(),l):s},l.chords=function(){return e||u(),e},l.groups=function(){return r||u(),r},l},t.layout.force=function(){var e,r,n,i,a,o,s={},l=t.dispatch("start","tick","end"),c=[1,1],u=.9,h=Ta,f=Aa,p=-30,d=Ma,g=.1,v=.64,m=[],y=[];function x(t){return function(e,r,n,i){if(e.point!==t){var a=e.cx-t.x,o=e.cy-t.y,s=i-r,l=a*a+o*o;if(s*s/v<l){if(l<d){var c=e.charge/l;t.px-=a*c,t.py-=o*c}return!0}if(e.point&&l&&l<d){c=e.pointCharge/l;t.px-=a*c,t.py-=o*c}}return!e.charge}}function b(e){e.px=t.event.x,e.py=t.event.y,s.resume()}return s.tick=function(){if((n*=.99)<.005)return e=null,l.end({type:"end",alpha:n=0}),!0;var r,s,h,f,d,v,b,_,w,k=m.length,T=y.length;for(s=0;s<T;++s)f=(h=y[s]).source,(v=(_=(d=h.target).x-f.x)*_+(w=d.y-f.y)*w)&&(_*=v=n*a[s]*((v=Math.sqrt(v))-i[s])/v,w*=v,d.x-=_*(b=f.weight+d.weight?f.weight/(f.weight+d.weight):.5),d.y-=w*b,f.x+=_*(b=1-b),f.y+=w*b);if((b=n*g)&&(_=c[0]/2,w=c[1]/2,s=-1,b))for(;++s<k;)(h=m[s]).x+=(_-h.x)*b,h.y+=(w-h.y)*b;if(p)for(!function t(e,r,n){var i=0,a=0;e.charge=0;if(!e.leaf)for(var o,s=e.nodes,l=s.length,c=-1;++c<l;)null!=(o=s[c])&&(t(o,r,n),e.charge+=o.charge,i+=o.charge*o.cx,a+=o.charge*o.cy);if(e.point){e.leaf||(e.point.x+=Math.random()-.5,e.point.y+=Math.random()-.5);var u=r*n[e.point.index];e.charge+=e.pointCharge=u,i+=u*e.point.x,a+=u*e.point.y}e.cx=i/e.charge;e.cy=a/e.charge}(r=t.geom.quadtree(m),n,o),s=-1;++s<k;)(h=m[s]).fixed||r.visit(x(h));for(s=-1;++s<k;)(h=m[s]).fixed?(h.x=h.px,h.y=h.py):(h.x-=(h.px-(h.px=h.x))*u,h.y-=(h.py-(h.py=h.y))*u);l.tick({type:"tick",alpha:n})},s.nodes=function(t){return arguments.length?(m=t,s):m},s.links=function(t){return arguments.length?(y=t,s):y},s.size=function(t){return arguments.length?(c=t,s):c},s.linkDistance=function(t){return arguments.length?(h="function"==typeof t?t:+t,s):h},s.distance=s.linkDistance,s.linkStrength=function(t){return arguments.length?(f="function"==typeof t?t:+t,s):f},s.friction=function(t){return arguments.length?(u=+t,s):u},s.charge=function(t){return arguments.length?(p="function"==typeof t?t:+t,s):p},s.chargeDistance=function(t){return arguments.length?(d=t*t,s):Math.sqrt(d)},s.gravity=function(t){return arguments.length?(g=+t,s):g},s.theta=function(t){return arguments.length?(v=t*t,s):Math.sqrt(v)},s.alpha=function(t){return arguments.length?(t=+t,n?t>0?n=t:(e.c=null,e.t=NaN,e=null,l.end({type:"end",alpha:n=0})):t>0&&(l.start({type:"start",alpha:n=t}),e=Te(s.tick)),s):n},s.start=function(){var t,e,r,n=m.length,l=y.length,u=c[0],d=c[1];for(t=0;t<n;++t)(r=m[t]).index=t,r.weight=0;for(t=0;t<l;++t)"number"==typeof(r=y[t]).source&&(r.source=m[r.source]),"number"==typeof r.target&&(r.target=m[r.target]),++r.source.weight,++r.target.weight;for(t=0;t<n;++t)r=m[t],isNaN(r.x)&&(r.x=g("x",u)),isNaN(r.y)&&(r.y=g("y",d)),isNaN(r.px)&&(r.px=r.x),isNaN(r.py)&&(r.py=r.y);if(i=[],"function"==typeof h)for(t=0;t<l;++t)i[t]=+h.call(this,y[t],t);else for(t=0;t<l;++t)i[t]=h;if(a=[],"function"==typeof f)for(t=0;t<l;++t)a[t]=+f.call(this,y[t],t);else for(t=0;t<l;++t)a[t]=f;if(o=[],"function"==typeof p)for(t=0;t<n;++t)o[t]=+p.call(this,m[t],t);else for(t=0;t<n;++t)o[t]=p;function g(r,i){if(!e){for(e=new Array(n),c=0;c<n;++c)e[c]=[];for(c=0;c<l;++c){var a=y[c];e[a.source.index].push(a.target),e[a.target.index].push(a.source)}}for(var o,s=e[t],c=-1,u=s.length;++c<u;)if(!isNaN(o=s[c][r]))retu
```
