## Supplemental Data 2 for "Rewiring phospholipid biosynthesis reveals robustness in membrane homeostasis and uncovers lipid regulatory players": typedarray.min.js

```
(function(global){"use strict";var undefined=void 0;var MAX_ARRAY_LENGTH=1e5;function Type(v){switch(typeof v){case"undefined":return"undefined";case"boolean":return"boolean";case"number":return"number";case"string":return"string";default:return v===null?"null":"object"}}function Class(v){return Object.prototype.toString.call(v).replace(/^\[object *|\]$/g,"")}function IsCallable(o){return typeof o==="function"}function ToObject(v){if(v===null||v===undefined)throw TypeError();return Object(v)}function ToInt32(v){return v>>0}function ToUint32(v){return v>>>0}var LN2=Math.LN2,abs=Math.abs,floor=Math.floor,log=Math.log,max=Math.max,min=Math.min,pow=Math.pow,round=Math.round;(function(){var orig=Object.defineProperty;var dom_only=!function(){try{return Object.defineProperty({},"x",{})}catch(_){return false}}();if(!orig||dom_only){Object.defineProperty=function(o,prop,desc){if(orig)try{return orig(o,prop,desc)}catch(_){}if(o!==Object(o))throw TypeError("Object.defineProperty called on non-object");if(Object.prototype.__defineGetter__&&"get"in desc)Object.prototype.__defineGetter__.call(o,prop,desc.get);if(Object.prototype.__defineSetter__&&"set"in desc)Object.prototype.__defineSetter__.call(o,prop,desc.set);if("value"in desc)o[prop]=desc.value;return o}}})();function makeArrayAccessors(obj){if(obj.length>MAX_ARRAY_LENGTH)throw RangeError("Array too large for polyfill");function makeArrayAccessor(index){Object.defineProperty(obj,index,{get:function(){return obj._getter(index)},set:function(v){obj._setter(index,v)},enumerable:true,configurable:false})}var i;for(i=0;i<obj.length;i+=1){makeArrayAccessor(i)}}function as_signed(value,bits){var s=32-bits;return value<<s>>s}function as_unsigned(value,bits){var s=32-bits;return value<<s>>>s}function packI8(n){return[n&255]}function unpackI8(bytes){return as_signed(bytes[0],8)}function packU8(n){return[n&255]}function unpackU8(bytes){return as_unsigned(bytes[0],8)}function packU8Clamped(n){n=round(Number(n));return[n<0?0:n>255?255:n&255]}function packI16(n){return[n>>8&255,n&255]}function unpackI16(bytes){return as_signed(bytes[0]<<8|bytes[1],16)}function packU16(n){return[n>>8&255,n&255]}function unpackU16(bytes){return as_unsigned(bytes[0]<<8|bytes[1],16)}function packI32(n){return[n>>24&255,n>>16&255,n>>8&255,n&255]}function unpackI32(bytes){return as_signed(bytes[0]<<24|bytes[1]<<16|bytes[2]<<8|bytes[3],32)}function packU32(n){return[n>>24&255,n>>16&255,n>>8&255,n&255]}function unpackU32(bytes){return as_unsigned(bytes[0]<<24|bytes[1]<<16|bytes[2]<<8|bytes[3],32)}function packIEEE754(v,ebits,fbits){var bias=(1<<ebits-1)-1,s,e,f,ln,i,bits,str,bytes;function roundToEven(n){var w=floor(n),f=n-w;if(f<.5)return w;if(f>.5)return w+1;return w%2?w+1:w}if(v!==v){e=(1<<ebits)-1;f=pow(2,fbits-1);s=0}else if(v===Infinity||v===-Infinity){e=(1<<ebits)-1;f=0;s=v<0?1:0}else if(v===0){e=0;f=0;s=1/v===-Infinity?1:0}else{s=v<0;v=abs(v);if(v>=pow(2,1-bias)){e=min(floor(log(v)/LN2),1023);f=roundToEven(v/pow(2,e)*pow(2,fbits));if(f/pow(2,fbits)>=2){e=e+1;f=1}if(e>bias){e=(1<<ebits)-1;f=0}else{e=e+bias;f=f-pow(2,fbits)}}else{e=0;f=roundToEven(v/pow(2,1-bias-fbits))}}bits=[];for(i=fbits;i;i-=1){bits.push(f%2?1:0);f=floor(f/2)}for(i=ebits;i;i-=1){bits.push(e%2?1:0);e=floor(e/2)}bits.push(s?1:0);bits.reverse();str=bits.join("");bytes=[];while(str.length){bytes.push(parseInt(str.substring(0,8),2));str=str.substring(8)}return bytes}function unpackIEEE754(bytes,ebits,fbits){var bits=[],i,j,b,str,bias,s,e,f;for(i=bytes.length;i;i-=1){b=bytes[i-1];for(j=8;j;j-=1){bits.push(b%2?1:0);b=b>>1}}bits.reverse();str=bits.join("");bias=(1<<ebits-1)-1;s=parseInt(str.substring(0,1),2)?-1:1;e=parseInt(str.substring(1,1+ebits),2);f=parseInt(str.substring(1+ebits),2);if(e===(1<<ebits)-1){return f!==0?NaN:s*Infinity}else if(e>0){return s*pow(2,e-bias)*(1+f/pow(2,fbits))}else if(f!==0){return s*pow(2,-(bias-1))*(f/pow(2,fbits))}else{return s<0?-0:0}}function unpackF64(b){return unpackIEEE754(b,11,52)}function packF64(v){return packIEEE754(v,11,52)}function unpackF32(b){return unpackIEEE754(b,8,23)}function packF32(v){return packIEEE754(v,8,23)}(function(){function ArrayBuffer(length){length=ToInt32(length);if(length<0)throw RangeError("ArrayBuffer size is not a small enough positive integer.");Object.defineProperty(this,"byteLength",{value:length});Object.defineProperty(this,"_bytes",{value:Array(length)});for(var i=0;i<length;i+=1)this._bytes[i]=0}global.ArrayBuffer=global.ArrayBuffer||ArrayBuffer;function $TypedArray$(){if(!arguments.length||typeof arguments[0]!=="object"){return function(length){length=ToInt32(length);if(length<0)throw RangeError("length is not a small enough positive integer.");Object.defineProperty(this,"length",{value:length});Object.defineProperty(this,"byteLength",{value:length*this.BYTES_PER_ELEMENT});Object.defineProperty(this,"buffer",{value:new ArrayBuffer(this.byteLength)});Object.defineProperty(this,"byteOffset",{value:0})}.apply(this,arguments)}if(arguments.length>=1&&Type(arguments[0])==="object"&&arguments[0]instanceof $TypedArray$){return function(typedArray){if(this.constructor!==typedArray.constructor)throw TypeError();var byteLength=typedArray.length*this.BYTES_PER_ELEMENT;Object.defineProperty(this,"buffer",{value:new ArrayBuffer(byteLength)});Object.defineProperty(this,"byteLength",{value:byteLength});Object.defineProperty(this,"byteOffset",{value:0});Object.defineProperty(this,"length",{value:typedArray.length});for(var i=0;i<this.length;i+=1)this._setter(i,typedArray._getter(i))}.apply(this,arguments)}if(arguments.length>=1&&Type(arguments[0])==="object"&&!(arguments[0]instanceof $TypedArray$)&&!(arguments[0]instanceof ArrayBuffer||Class(arguments[0])==="ArrayBuffer")){return function(array){var byteLength=array.length*this.BYTES_PER_ELEMENT;Object.defineProperty(this,"buffer",{value:new ArrayBuffer(byteLength)});Object.defineProperty(this,"byteLength",{value:byteLength});Object.defineProperty(this,"byteOffset",{value:0});Object.defineProperty(this,"length",{value:array.length});for(var i=0;i<this.length;i+=1){var s=array[i];this._setter(i,Number(s))}}.apply(this,arguments)}if(arguments.length>=1&&Type(arguments[0])==="object"&&(arguments[0]instanceof ArrayBuffer||Class(arguments[0])==="ArrayBuffer")){return function(buffer,byteOffset,length){byteOffset=ToUint32(byteOffset);if(byteOffset>buffer.byteLength)throw RangeError("byteOffset out of range");if(byteOffset%this.BYTES_PER_ELEMENT)throw RangeError("buffer length minus the byteOffset is not a multiple of the element size.");if(length===undefined){var byteLength=buffer.byteLength-byteOffset;if(byteLength%this.BYTES_PER_ELEMENT)throw RangeError("length of buffer minus byteOffset not a multiple of the element size");length=byteLength/this.BYTES_PER_ELEMENT}else{length=ToUint32(length);byteLength=length*this.BYTES_PER_ELEMENT}if(byteOffset+byteLength>buffer.byteLength)throw RangeError("byteOffset and length reference an area beyond the end of the buffer");Object.defineProperty(this,"buffer",{value:buffer});Object.defineProperty(this,"byteLength",{value:byteLength});Object.defineProperty(this,"byteOffset",{value:byteOffset});Object.defineProperty(this,"length",{value:length})}.apply(this,arguments)}throw TypeError()}Object.defineProperty($TypedArray$,"from",{value:function(iterable){return new this(iterable)}});Object.defineProperty($TypedArray$,"of",{value:function(){return new this(arguments)}});var $TypedArrayPrototype$={};$TypedArray$.prototype=$TypedArrayPrototype$;Object.defineProperty($TypedArray$.prototype,"_getter",{value:function(index){if(arguments.length<1)throw SyntaxError("Not enough arguments");index=ToUint32(index);if(index>=this.length)return undefined;var bytes=[],i,o;for(i=0,o=this.byteOffset+index*this.BYTES_PER_ELEMENT;i<this.BYTES_PER_ELEMENT;i+=1,o+=1){bytes.push(this.buffer._bytes[o])}return this._unpack(bytes)}});Object.defineProperty($TypedArray$.prototype,"get",{value:$TypedArray$.prototype._getter});Object.defineProperty($TypedArray$.prototype,"_setter",{value:function(index,value){if(arguments.length<2)throw SyntaxError("Not enough arguments");index=ToUint32(index);if(index>=this.length)return;var bytes=this._pack(value),i,o;for(i=0,o=this.byteOffset+index*this.BYTES_PER_ELEMENT;i<this.BYTES_PER_ELEMENT;i+=1,o+=1){this.buffer._bytes[o]=bytes[i]}}});Object.defineProperty($TypedArray$.prototype,"constructor",{value:$TypedArray$});Object.defineProperty($TypedArray$.prototype,"copyWithin",{value:function(target,start){var end=arguments[2];var o=ToObject(this);var lenVal=o.length;var len=ToUint32(lenVal);len=max(len,0);var relativeTarget=ToInt32(target);var to;if(relativeTarget<0)to=max(len+relativeTarget,0);else to=min(relativeTarget,len);var relativeStart=ToInt32(start);var from;if(relativeStart<0)from=max(len+relativeStart,0);else from=min(relativeStart,len);var relativeEnd;if(end===undefined)relativeEnd=len;else relativeEnd=ToInt32(end);var final;if(relativeEnd<0)final=max(len+relativeEnd,0);else final=min(relativeEnd,len);var count=min(final-from,len-to);var direction;if(from<to&&to<from+count){direction=-1;from=from+count-1;to=to+count-1}else{direction=1}while(count>0){o._setter(to,o._getter(from));from=from+direction;to=to+direction;count=count-1}return o}});Object.defineProperty($TypedArray$.prototype,"every",{value:function(callbackfn){if(this===undefined||this===null)throw TypeError();var t=Object(this);var len=ToUint32(t.length);if(!IsCallable(callbackfn))throw TypeError();var thisArg=arguments[1];for(var i=0;i<len;i++){if(!callbackfn.call(thisArg,t._getter(i),i,t))return false}return true}});Object.defineProperty($TypedArray$.prototype,"fill",{value:function(value){var start=arguments[1],end=arguments[2];var o=ToObject(this);var lenVal=o.length;var len=ToUint32(lenVal);len=max(len,0);var relativeStart=ToInt32(start);var k;if(relativeStart<0)k=max(len+relativeStart,0);else k=min(relativeStart,len);var relativeEnd;if(end===undefined)relativeEnd=len;else relativeEnd=ToInt32(end);var final;if(relativeEnd<0)final=max(len+relativeEnd,0);else final=min(relativeEnd,len);while(k<final){o._setter(k,value);k+=1}return o}});Object.defineProperty($TypedArray$.prototype,"filter",{value:function(callbackfn){if(this===undefined||this===null)throw TypeError();var t=Object(this);var len=ToUint32(t.length);if(!IsCallable(callbackfn))throw TypeError();var res=[];var thisp=arguments[1];for(var i=0;i<len;i++){var val=t._getter(i);if(callbackfn.call(thisp,val,i,t))res.push(val)}return new this.constructor(res)}});Object.defineProperty($TypedArray$.prototype,"find",{value:function(predicate){var o=ToObject(this);var lenValue=o.length;var len=ToUint32(lenValue);if(!IsCallable(predicate))throw TypeError();var t=arguments.length>1?arguments[1]:undefined;var k=0;while(k<len){var kValue=o._getter(k);var testResult=predicate.call(t,kValue,k,o);if(Boolean(testResult))return kValue;++k}return undefined}});Object.defineProperty($TypedArray$.prototype,"findIndex",{value:function(predicate){var o=ToObject(this);var lenValue=o.length;var len=ToUint32(lenValue);if(!IsCallable(predicate))throw TypeError();var t=arguments.length>1?arguments[1]:undefined;var k=0;while(k<len){var kValue=o._getter(k);var testResult=predicate.call(t,kValue,k,o);if(Boolean(testResult))return k;++k}return-1}});Object.defineProperty($TypedArray$.prototype,"forEach",{value:function(callbackfn){if(this===undefined||this===null)throw TypeError();var t=Object(this);var len=ToUint32(t.length);if(!IsCallable(callbackfn))throw TypeError();var thisp=arguments[1];for(var i=0;i<len;i++)callbackfn.call(thisp,t._getter(i),i,t)}});Object.defineProperty($TypedArray$.prototype,"indexOf",{value:function(searchElement){if(this===undefined||this===null)throw TypeError();var t=Object(this);var len=ToUint32(t.length);if(len===0)return-1;var n=0;if(arguments.length>0){n=Number(arguments[1]);if(n!==n){n=0}else if(n!==0&&n!==1/0&&n!==-(1/0)){n=(n>0||-1)*floor(abs(n))}}if(n>=len)return-1;var k=n>=0?n:max(len-abs(n),0);for(;k<len;k++){if(t._getter(k)===searchElement){return k}}return-1}});Object.defineProperty($TypedArray$.prototype,"join",{value:function(separator){if(this===undefined||this===null)throw TypeError();var t=Object(this);var len=ToUint32(t.length);var tmp=Array(len);for(var i=0;i<len;++i)tmp[i]=t._getter(i);return tmp.join(separator===undefined?",":separator)}});Object.defineProperty($TypedArray$.prototype,"lastIndexOf",{value:function(searchElement){if(this===undefined||this===null)throw TypeError();var t=Object(this);var len=ToUint32(t.length);if(len===0)return-1;var n=len;if(arguments.length>1){n=Number(arguments[1]);if(n!==n){n=0}else if(n!==0&&n!==1/0&&n!==-(1/0)){n=(n>0||-1)*floor(abs(n))}}var k=n>=0?min(n,len-1):len-abs(n);for(;k>=0;k--){if(t._getter(k)===searchElement)return k}return-1}});Object.defineProperty($TypedArray$.prototype,"map",{value:function(callbackfn){if(this===undefined||this===null)throw TypeError();var t=Object(this);var len=ToUint32(t.length);if(!IsCallable(callbackfn))throw TypeError();var res=[];res.length=len;var thisp=arguments[1];for(var i=0;i<len;i++)res[i]=callbackfn.call(thisp,t._getter(i),i,t);return new this.constructor(res)}});Object.defineProperty($TypedArray$.prototype,"reduce",{value:function(callbackfn){if(this===undefined||this===null)throw TypeError();var t=Object(this);var len=ToUint32(t.length);if(!IsCallable(callbackfn))throw TypeError();if(len===0&&arguments.length===1)throw TypeError();var k=0;var accumulator;if(arguments.length>=2){accumulator=arguments[1]}else{accumulator=t._getter(k++)}while(k<len){accumulator=callbackfn.call(undefined,accumulator,t._getter(k),k,t);k++}return accumulator}});Object.defineProperty($TypedArray$.prototype,"reduceRight",{value:function(callbackfn){if(this===undefined||this===null)throw TypeError();var t=Object(this);var len=ToUint32(t.length);if(!IsCallable(callbackfn))throw TypeError();if(len===0&&arguments.length===1)throw TypeError();var k=len-1;var accumulator;if(arguments.length>=2){accumulator=arguments[1]}else{accumulator=t._getter(k--)}while(k>=0){accumulator=callbackfn.call(undefined,accumulator,t._getter(k),k,t);k--}return accumulator}});Object.defineProperty($TypedArray$.prototype,"reverse",{value:function(){if(this===undefined||this===null)throw TypeError();var t=Object(this);var len=ToUint32(t.length);var half=floor(len/2);for(var i=0,j=len-1;i<half;++i,--j){var tmp=t._getter(i);t._setter(i,t._getter(j));t._setter(j,tmp)}return t}});Object.defineProperty($TypedArray$.prototype,"set",{value:function(index,value){if(arguments.length<1)throw SyntaxError("Not enough arguments");var array,sequence,offset,len,i,s,d,byteOffset,byteLength,tmp;if(typeof arguments[0]==="object"&&arguments[0].constructor===this.constructor){array=arguments[0];offset=ToUint32(arguments[1]);if(offset+array.length>this.length){throw RangeError("Offset plus length of array is out of range")}byteOffset=this.byteOffset+offset*this.BYTES_PER_ELEMENT;byteLength=array.length*this.BYTES_PER_ELEMENT;if(array.buffer===this.buffer){tmp=[];for(i=0,s=array.byteOffset;i<byteLength;i+=1,s+=1){tmp[i]=array.buffer._bytes[s]}for(i=0,d=byteOffset;i<byteLength;i+=1,d+=1){this.buffer._bytes[d]=tmp[i]}}else{for(i=0,s=array.byteOffset,d=byteOffset;i<byteLength;i+=1,s+=1,d+=1){this.buffer._bytes[d]=array.buffer._bytes[s]}}}else if(typeof arguments[0]==="object"&&typeof arguments[0].length!=="undefined"){sequence=arguments[0];len=ToUint32(sequence.length);offset=ToUint32(arguments[1]);if(offset+len>this.length){throw RangeError("Offset plus length of array is out of range")}for(i=0;i<len;i+=1){s=sequence[i];this._setter(offset+i,Number(s))}}else{throw TypeError("Unexpected argument type(s)")}}});Object.defineProperty($TypedArray$.prototype,"slice",{value:function(start,end){var o=ToObject(this);var lenVal=o.length;var len=ToUint32(lenVal);var relativeStart=ToInt32(start);var k=relativeStart<0?max(len+relativeStart,0):min(relativeStart,len);var relativeEnd=end===undefined?len:ToInt32(end);var final=relativeEnd<0?max(len+relativeEnd,0):min(relativeEnd,len);var count=final-k;var c=o.constructor;var a=new c(count);var n=0;while(k<final){var kValue=o._getter(k);a._setter(n,kValue);++k;++n}return a}});Object.defineProperty($TypedArray$.prototype,"some",{value:function(callbackfn){if(this===undefined||this===null)throw TypeError();var t=Object(this);var len=ToUint32(t.length);if(!IsCallable(callbackfn))throw TypeError();var thisp=arguments[1];for(var i=0;i<len;i++){if(callbackfn.call(thisp,t._getter(i),i,t)){return true}}return false}});Object.defineProperty($TypedArray$.prototype,"sort",{value:function(comparefn){if(this===undefined||this===null)throw TypeError();var t=Object(this);var len=ToUint32(t.length);var tmp=Array(len);for(var i=0;i<len;++i)tmp[i]=t._getter(i);if(comparefn)tmp.sort(comparefn);else tmp.sort();for(i=0;i<len;++i)t._setter(i,tmp[i]);return t}});Object.defineProperty($TypedArray$.prototype,"subarray",{value:function(start,end){function clamp(v,min,max){return v<min?min:v>max?max:v}start=ToInt32(start);end=ToInt32(end);if(arguments.length<1){start=0}if(arguments.length<2){end=this.length}if(start<0){start=this.length+start}if(end<0){end=this.length+end}start=clamp(start,0,this.length);end=clamp(end,0,this.length);var len=end-start;if(len<0){len=0}return new this.constructor(this.buffer,this.byteOffset+start*this.BYTES_PER_ELEMENT,len)}});function makeTypedArray(elementSize,pack,unpack){var TypedArray=function(){Object.defineProperty(this,"constructor",{value:TypedArray});$TypedArray$.apply(this,arguments);makeArrayAccessors(this)};if("__proto__"in TypedArray){TypedArray.__proto__=$TypedArray$}else{TypedArray.from=$TypedArray$.from;TypedArray.of=$TypedArray$.of}TypedArray.BYTES_PER_ELEMENT=elementSize;var TypedArrayPrototype=function(){};TypedArrayPrototype.prototype=$TypedArrayPrototype$;TypedArray.prototype=new TypedArrayPrototype;Object.defineProperty(TypedArray.prototype,"BYTES_PER_ELEMENT",{value:elementSize});Object.defineProperty(TypedArray.prototype,"_pack",{value:pack});Object.defineProperty(TypedArray.prototype,"_unpack",{value:unpack});return TypedArray}var Int8Array=makeTypedArray(1,packI8,unpackI8);var Uint8Array=makeTypedArray(1,packU8,unpackU8);var Uint8ClampedArray=makeTypedArray(1,packU8Clamped,unpackU8);var Int16Array=makeTypedArray(2,packI16,unpackI16);var Uint16Array=makeTypedArray(2,packU16,unpackU16);var Int32Array=makeTypedArray(4,packI32,unpackI32);var Uint32Array=makeTypedArray(4,packU32,unpackU32);var Float32Array=makeTypedArray(4,packF32,unpackF32);var Float64Array=makeTypedArray(8,packF64,unpackF64);global.Int8Array=global.Int8Array||Int8Array;global.Uint8Array=global.Uint8Array||Uint8Array;global.Uint8ClampedArray=global.Uint8ClampedArray||Uint8ClampedArray;global.Int16Array=global.Int16Array||Int16Array;global.Uint16Array=global.Uint16Array||Uint16Array;global.Int32Array=global.Int32Array||Int32Array;global.Uint32Array=global.Uint32Array||Uint32Array;global.Float32Array=global.Float32Array||Float32Array;global.Float64Array=global.Float64Array||Float64Array})();(function(){function r(array,index){return IsCallable(array.get)?array.get(index):array[index]}var IS_BIG_ENDIAN=function(){var u16array=new Uint16Array([4660]),u8array=new Uint8Array(u16array.buffer);return r(u8array,0)===18}();function DataView(buffer,byteOffset,byteLength){if(!(buffer instanceof ArrayBuffer||Class(buffer)==="ArrayBuffer"))throw TypeError();byteOffset=ToUint32(byteOffset);if(byteOffset>buffer.byteLength)throw RangeError("byteOffset out of range");if(byteLength===undefined)byteLength=buffer.byteLength-byteOffset;else byteLength=ToUint32(byteLength);if(byteOffset+byteLength>buffer.byteLength)throw RangeError("byteOffset and length reference an area beyond the end of the buffer");Object.defineProperty(this,"buffer",{value:buffer});Object.defineProperty(this,"byteLength",{value:byteLength});Object.defineProperty(this,"byteOffset",{value:byteOffset})}function makeGetter(arrayType){return function GetViewValue(byteOffset,littleEndian){byteOffset=ToUint32(byteOffset);if(byteOffset+arrayType.BYTES_PER_ELEMENT>this.byteLength)throw RangeError("Array index out of range");byteOffset+=this.byteOffset;var uint8Array=new Uint8Array(this.buffer,byteOffset,arrayType.BYTES_PER_ELEMENT),bytes=[];for(var i=0;i<arrayType.BYTES_PER_ELEMENT;i+=1)bytes.push(r(uint8Array,i));if(Boolean(littleEndian)===Boolean(IS_BIG_ENDIAN))bytes.reverse();return r(new arrayType(new Uint8Array(bytes).buffer),0)}}Object.defineProperty(DataView.prototype,"getUint8",{value:makeGetter(Uint8Array)});Object.defineProperty(DataView.prototype,"getInt8",{value:makeGetter(Int8Array)});Object.defineProperty(DataView.prototype,"getUint16",{value:makeGetter(Uint16Array)});Object.defineProperty(DataView.prototype,"getInt16",{value:makeGetter(Int16Array)});Object.defineProperty(DataView.prototype,"getUint32",{value:makeGetter(Uint32Array)});Object.defineProperty(DataView.prototype,"getInt32",{value:makeGetter(Int32Array)});Object.defineProperty(DataView.prototype,"getFloat32",{value:makeGetter(Float32Array)});Object.defineProperty(DataView.prototype,"getFloat64",{value:makeGetter(Float64Array)});function makeSetter(arrayType){return function SetViewValue(byteOffset,value,littleEndian){byteOffset=ToUint32(byteOffset);if(byteOffset+arrayType.BYTES_PER_ELEMENT>this.byteLength)throw RangeError("Array index out of range");var typeArray=new arrayType([value]),byteArray=new Uint8Array(typeArray.buffer),bytes=[],i,byteView;for(i=0;i<arrayType.BYTES_PER_ELEMENT;i+=1)bytes.push(r(byteArray,i));if(Boolean(littleEndian)===Boolean(IS_BIG_ENDIAN))bytes.reverse();byteView=new Uint8Array(this.buffer,byteOffset,arrayType.BYTES_PER_ELEMENT);byteView.set(bytes)}}Object.defineProperty(DataView.prototype,"setUint8",{value:makeSetter(Uint8Array)});Object.defineProperty(DataView.prototype,"setInt8",{value:makeSetter(Int8Array)});Object.defineProperty(DataView.prototype,"setUint16",{value:makeSetter(Uint16Array)});Object.defineProperty(DataView.prototype,"setInt16",{value:makeSetter(Int16Array)});Object.defineProperty(DataView.prototype,"setUint32",{value:makeSetter(Uint32Array)});Object.defineProperty(DataView.prototype,"setInt32",{value:makeSetter(Int32Array)});Object.defineProperty(DataView.prototype,"setFloat32",{value:makeSetter(Float32Array)});Object.defineProperty(DataView.prototype,"setFloat64",{value:makeSetter(Float64Array)});global.DataView=global.DataView||DataView})()})(this);
```
